# Identification of novel mutational signatures in Asian oral squamous cell carcinomas associated with bacterial infections

**DOI:** 10.1101/368753

**Authors:** Arnoud Boot, Alvin W.T. Ng, Fui Teen Chong, Szu-Chi Ho, Willie Yu, Daniel S.W. Tan, N. Gopalakrishna Iyer, Steven G. Rozen

**Author notes:** Corresponding author: SGR.

## Abstract

Mutational signatures can reveal the history of mutagenic processes that cells were exposed to prior to and during tumourigenesis. We expect that as-yet-undiscovered mutational processes will shed further light on mutagenesis leading to carcinogenesis. With this in mind, we analyzed the mutational spectra of 36 Asian oral squamous cell carcinomas. The mutational spectra of two samples from patients who presented with oral bacterial infections, showed novel mutational signatures. One of these novel signatures, SBS_A^n^T, is characterized by a preponderance of thymine mutations, strong transcriptional strand bias, and striking enrichment for adenines in the 4 base pairs 5’ of mutation sites. Examination of publicly available sequencing data revealed SBS_A^n^T in 25 tumours from several mucosal tissue types, all of which harbour human symbionts or are adjacent to tissues that harbour symbionts. Data in a preprint released while this manuscript was in revision strongly suggest that the bacterial compound colibactin causes SBS_A^n^T.

## Introduction

Mutagenesis is one of the major causes of cancer. A thorough mapping of mutational signatures promises to illuminate the mechanisms of carcinogenesis and help identify carcinogenic mutagenic compounds and processes. In recent years, the field of mutational-signature analysis has made huge strides in identifying distinct mutational processes. Currently, 65 distinct single base substitution (SBS) signatures have been described (Alexandrov et al. 2019). Most of these stem from defects in DNA repair and replication, endogenous mutagenic processes, or exposure to mutagenic compounds such as benzo[a]pyrene or aristolochic acid. However, the aetiology of 20 mutational signatures remains unknown (Alexandrov et al. 2019).

Although the mutational signatures of most common mutational processes are known, we expect that there are additional mutational processes that contribute to small numbers of tumours. An example of such a rare signature is SBS42, due to occupational exposure to haloalkanes (Mimaki et al. 2016; Alexandrov et al. 2019). This signature was not discovered in the original COSMIC signatures (Forbes et al. 2017), but was only discovered in cholangiocarcinomas from patients who worked at a printing company. SBS42 was extremely rare in other cancer types (Alexandrov et al. 2019). This example suggests that there are more rare mutational processes that are due to rare occupational exposures, dietary exposures, or genetic variants affecting DNA repair or replication mechanisms. Rare mutational processes will be challenging to find, but they will point to cancers that could be prevented if the responsible mutagens can be identified and exposure to them avoided. We might expect populations that have not be intensively studied to harbour such rare mutational signatures.

Head and neck squamous cell carcinoma (HNSCC) is the 6^th^ most common cancer worldwide, with more than 680,000 new cases every year (Ferlay et al. 2015). With 300,000 new cases per year, oral squamous cell carcinoma (OSCC) is the largest subtype (Ferlay et al. 2015). In OSCCs, 9 different mutational signatures have been detected, but >92% of mutations are due to mutational signatures associated with: aging (clock-like signatures SBS1 and SBS5), APOBEC cytidine deaminases (SBS2 and SBS13), and chewing tobacco (SBS29) (Alexandrov et al. 2019). With this in mind, we analyzed whole-exome sequencing data of 36 Asian OSCCs to search for possible rare mutational processes.

## Results

### Bacterial infection associated OSCCs show novel mutational signatures

We analyzed whole-exome sequencing data from 36 OSCCs treated in Singapore, including 18 previously published OSCCs (Vettore et al. 2015). Clinical information on these tumours is included in Supplemental Table S1. These tumours had significantly fewer somatic single base substitutions (SBSs) than the OSCCs and HNSCCs analyzed by the TCGA consortium (median 1.02 versus 1.66 and 2.44 mutations per megabase, *p* = 4.11×10^−5^ and 4.85×10^−10^ respectively, Wilcoxon rank sum tests) (Ellrott et al. 2018; Alexandrov et al. 2019). No difference in tumour mutation burden was observed between smokers and non-smokers. Strikingly, the two tumours from patients that presented with strong bacterial infection (62074759 and TC1) showed higher mutation burden, although not statistically significant (average mutation burden of 2.6 and 1.14 mutations per megabase respectively, p=0.078, Wilcoxon rank sum test). Experience has shown that mutational signature assignment to tumours with extremely low numbers of mutations is unreliable. Therefore we excluded 6 tumours that had < 10 SBSs from further analysis. The mutational spectra of the remaining 30 tumours are shown in Supplemental Fig S1.

We computationally reconstructed the mutational spectra of the 30 tumours using the mutational signatures previously observed in HNSCCs and OSCCs (Supplemental Fig S2A) (Alexandrov et al. 2019). The spectra of 62074759 and TC1 were poorly reconstructed (Fig. 1, Supplemental Fig S2B). Strikingly, examination of the pathology reports revealed that both 62074759 and TC1 had presented with strong oral bacterial infections, while none of the other 34 had mentions of bacterial infection. (*p*=0.0016, Fisher’s exact test, Supplemental Table S1). Both ofthese poorly reconstructed spectra showed unique distinctive mutation patterns. Clustering of the mutational spectra of the OSCC cohort together with the TCGA HNSCCs showed 62074759 and TC1 clustering apart, supporting these mutational spectra being distinct (Supplemental Fig S3). This led us to hypothesize that each was caused predominantly by a single, novel, mutational process, which in the case of TC1 appeared to be combined with APOBEC mutagenesis (Alexandrov et al. 2019). Both spectra showed T>A and T>C peaks with strong transcriptional strand bias, but were clearly distinct.

**Fig. 1:**
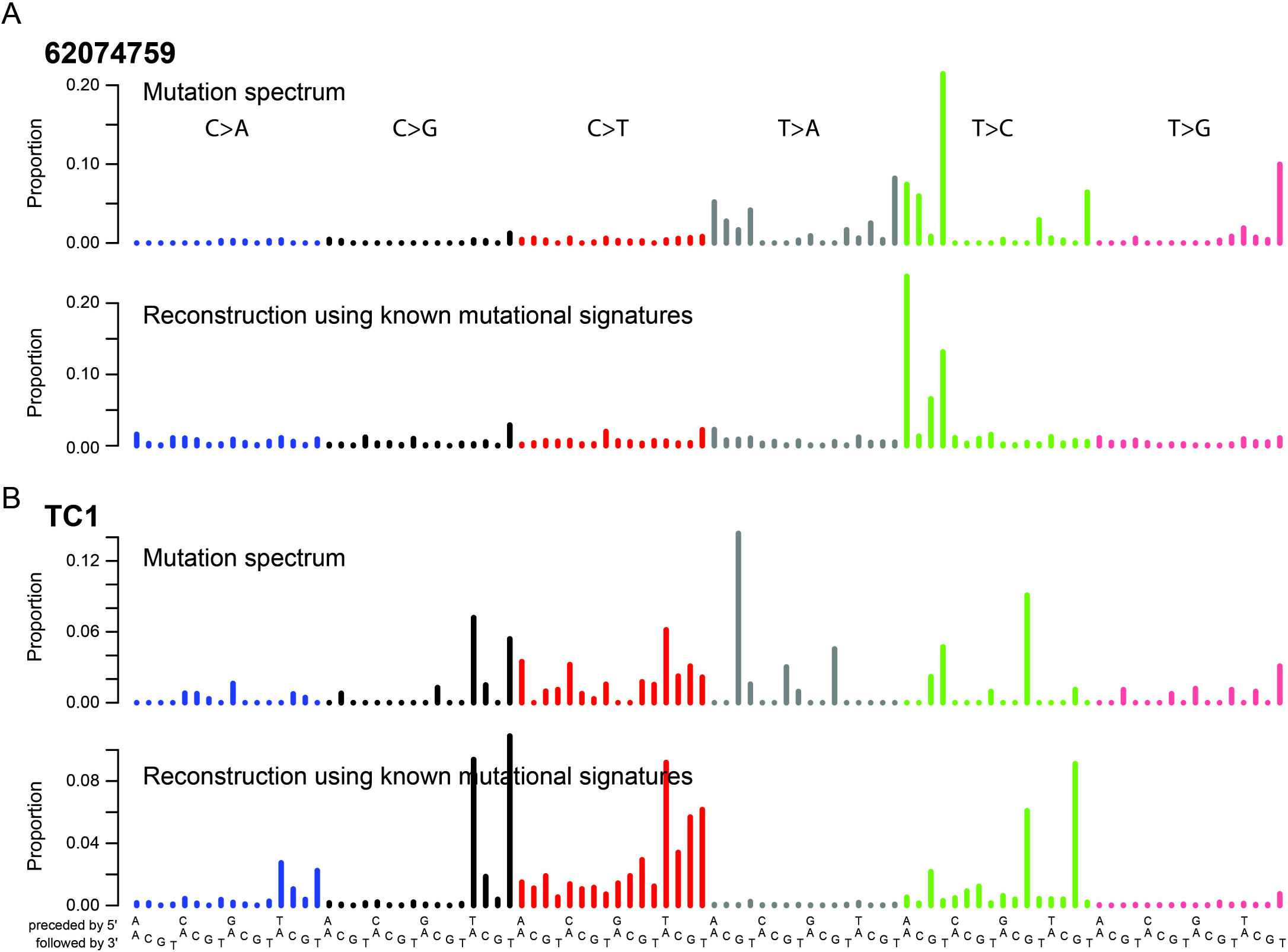
Two OSCC mutation spectra were poorly reconstructed using known mutational signatures. Mutational signature plots comparing the observed exome mutational spectra of 62074759 **(A)** and TC1 **(B)** to the corresponding reconstructed spectra.

### The SBS mutational spectrum in 62074759

During routine visual inspection of the read alignments supporting the somatic variants in 62074759, we noticed that 51 out of the 84 T>C mutations were directly preceded by at least 3 adenines (3 adenines directly 5’ of the T>C mutation). In addition, most of the T<u>T</u>T>T<u>N</u>T mutations were located within TTTT homopolymers. Because of the high risk of sequencing errors in and near homopolymers, we performed Sanger sequencing to validate 96 somatic SBSs detected in 62074759, all of which were confirmed.

We next sequenced the whole-genome of 62074759, identifying 34,905 somatic SBSs and 4,037 small insertions and deletions (indels). The whole-genome SBS mutation spectrum confirmed the spectrum observed in the exome (Fig. 2A, Supplemental Fig S1). The spectrum was dominated by A<u>T</u>>A<u>A</u> and A<u>T</u>>A<u>C</u> mutations with a main peak at A<u>T</u>T>A<u>C</u>T, and by T<u>T</u>T>T<u>A</u>T, T<u>T</u>T>T<u>C</u>T and T<u>T</u>T>T<u>G</u>T mutations. Similar to the exome data, the genome data showed a striking enrichment for adenines 5’ of T>C mutations. Among **all** SBSs, 79.5% had an adenine 3 bp 5’ of the mutation sites and 65.3% had an adenine 4 bp 5’ of the mutation (Fig. 2B). Thymine mutations predominantly occurred in AAWW<u>T</u>W motifs, with 93.5% and 75.2% having adenines 3 bp and 4 bp 5’ of the mutation, respectively. A<u>T</u>T>A<u>C</u>T SBSs mainly occurred in AAWA<u>T</u>T motifs, with 98.2% having an adenine 3 bp 5’ of the mutation. More broadly, we also observed strong enrichment for AAAA immediately 5’ of thymine SBSs (Fig. 2C). No enrichment of adenines 5’ of mutated cytosines was observed (Supplemental Fig S4).

**Fig. 2:**
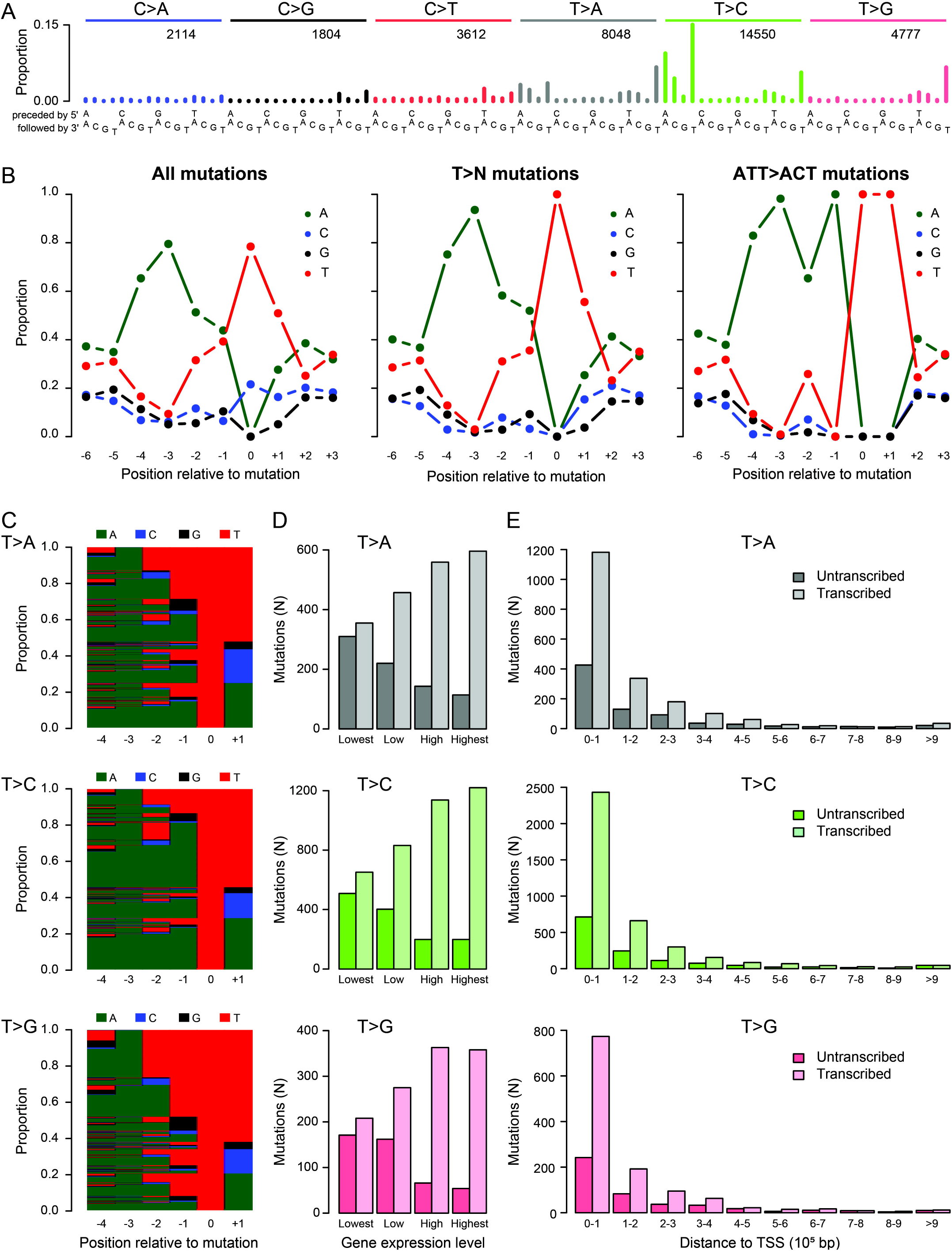
In-depth characterization of SBS_A^n^T in the whole-genome data from tumour 62074759. **(A)** SBS spectrum. **(B)** SBS sequence context preferences, revealing strong preference for adenines 3bp 3’ of mutated thymines. Thymine mutations predominantly occurred in AAWW<u>T</u>W motifs. **(C)** Per-mutation view of sequence context preferences of mutations from thymines. Each row represents one mutation, with bases indicated by colour as in panel B. **(D)** Transcriptional strand bias as a function of gene expression level and distance to the transcription start site **(E)**.

In 62074759, the mutational spectra of SBSs in trinucleotide context were essentially identical at a wide range of variant allele frequencies (VAFs) (Supplemental Fig S5). The presence of this signature in mutations with high VAFs as well as lower VAFs suggests that the underlying mutational process continued for a considerable period of time, which included both tumour initiation and tumour expansion.

Mutational processes associated with large adducts are known to generate more mutations on the non-transcribed strands of genes than on the transcribed strands, due to transcription-coupled nucleotide excision repair (TC-NER) of the adducts on transcribed strands (Tomkova et al. 2018). Therefore, to investigate whether this novel signature might have been caused by large adducts, we examined its transcriptional strand bias. We observed very strong enrichment of mutations when thymine is on the transcribed strand (and adenine is on the non-transcribed strand), which is indicative of adduct formation on adenines. Consistent with the activity of TC-NER, the bias of T>A, T>C and T>G mutations correlated strongly with transcriptional activity (Fig. 2D, *p*= 9.50×10^−41^, 6.33×10^−91^ and 5.69×10^−33^ respectively, Chi-squared tests). Furthermore, TC-NER proficiency decreases with increasing distance to the transcription start site (Huang et al. 2017; Boot et al. 2018). Consistent with this, T>A, T>C and T>G mutations all showed decreased transcriptional strand bias towards the 3’ end of transcripts (Fig. 2E, *p* = 4.32×10^−32^, 1.26×10^−78^ and 6.64×10^−24^ respectively, logistic regression with transcription strand as independent variable and distance to TSS as dependent variable). None of the cytosine mutation classes showed transcriptional strand bias. This, plus the absence of enrichment for adenines 5’ of cytosine mutations, suggests that the cytosine mutations in this sample were not caused by the same mutational process as the thymine mutations. In light of the striking preference for adenines 5’ of mutations from thymines in 62074759, we call this signature SBS_A^n^T.

### Insertions, deletions and dinucleotide substitutions associated with SBS_A^n^T

The vast majority of indels were deletions (98.6%), mainly of single thymines (Fig. 3A). The indel spectrum did not resemble any of the previously published indel signatures (Alexandrov et al. 2019). Like the SBSs, deletions of thymines in thymine mono-and dinucleotides showed strong enrichment for three preceding adenines (Fig. 3B+C). Thymine deletions in thymine tri-to octonucleotides, had very strong enrichment for single adenines immediately 5’ of the thymine repeat, but enrichment for adenines further 5’ decreased rapidly for longer repeats (Supplemental Fig S6). For thymine deletions outside of thymine repeats, we observed strand bias consistent with adenine adducts (*p* = 0.01, binomial test, Supplemental Fig S7). Contrastingly, thymine deletions in thymine tetranucleotides showed transcriptional strand bias in the opposite direction (*p* = 0.01, binomial test). Thymine deletions in thymine homopolymers of other lengths lacked transcriptional strand bias. We call this indel signature ID_A^n^T.

**Fig. 3:**
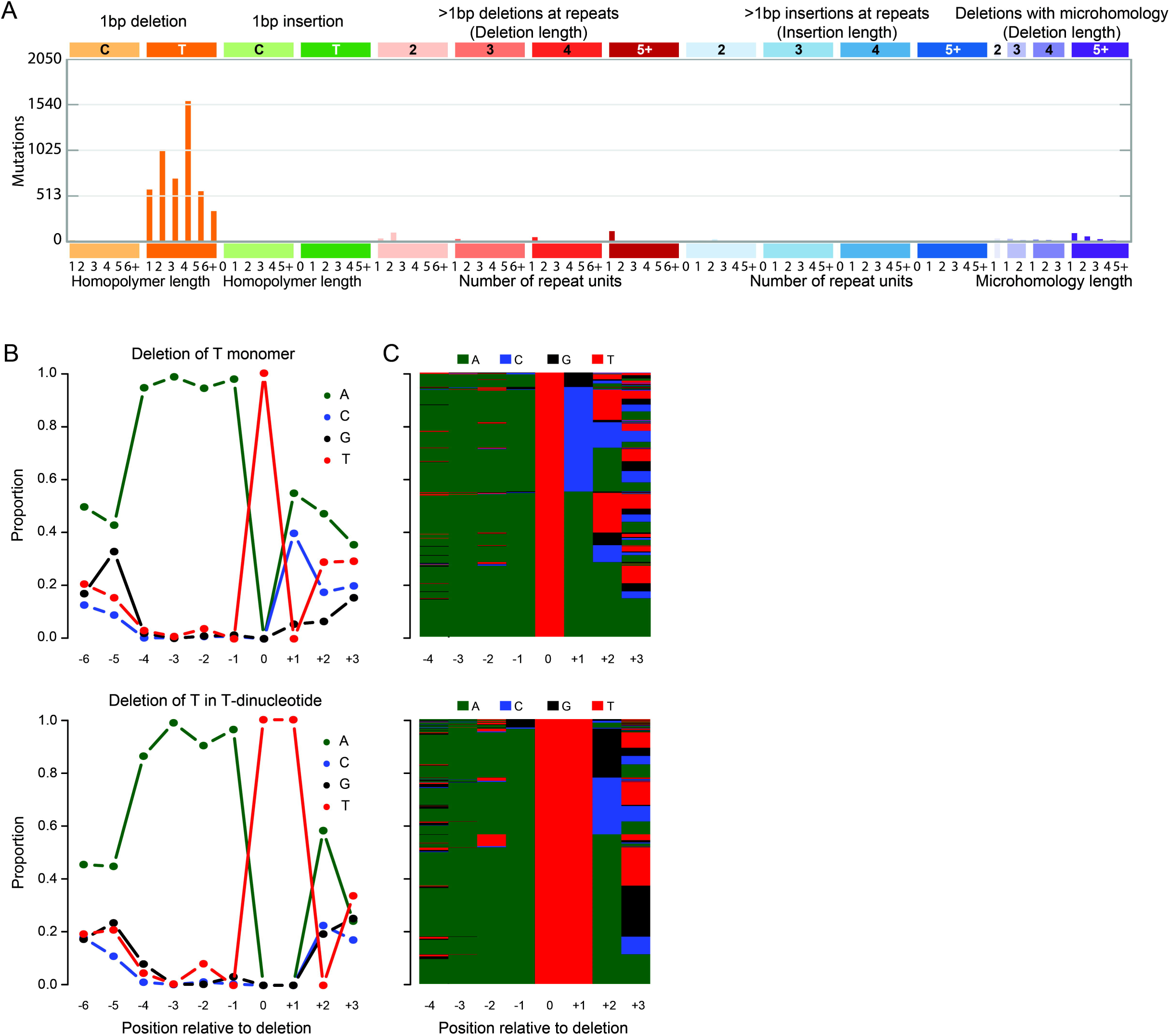
In-depth characterisation of the ID_A^n^T indel signature in the whole genome data from tumour 62074759. **(A)** Spectrum of indels in the classification proposed by the PCAWG consortium (Alexandrov et al. 2019). The indel spectrum is dominated by deletions of single thymines (orange). The numbers at the bottom indicate the lengths of the repeats in which the deletions occurred; "1" denotes deletions of thymines flanked by non-thymines, "2" denotes deletions of thymines in dinucleotides (TT>T), and so on. **(B)** Sequence contexts of deletions of single thymines not in a T-repeat (top) and in thymine dinucleotides (bottom). Supplemental Fig S6 provides analogous plots for thymine deletions in longer homopolymers. **(C)** Per-mutation view of sequence contexts of thymine deletions not in a T-repeat (top) and in a thymine dinucleotide (bottom); bases are indicated by colours as in panel B.

We detected 171 double base substitutions (DBSs) in tumour 62074759, most of which were CC>NN and TC>NN substitutions (Supplemental Fig S8). As the predominant mutational process in 62074759 caused almost exclusively thymine mutations, it is unlikely that these DBSs were caused by the same mutational process. The DBS spectrum best resembles the DBS spectrum observed in cisplatin exposed cell lines (cosine similarity of 0.859) (Boot et al. 2018). In concordance with cisplatin mutagenesis inducing these DBSs, prior to the development of the tumour that we sequenced, which was a recurrence, the initial tumour had been treated with several chemotherapeutic drugs, including cisplatin.

### SBS_A^n^T in publicly available sequencing data

To investigate whether SBS_A^n^T was also present in other tumours, we investigated 4,645 tumour genomes and 19,184 tumour exomes compiled for the PCAWG Mutational Signatures Working Group (Alexandrov et al. 2019). We searched for tumours that both show strong enrichment of adenines 3 and 4 bp 5’ of mutated thymines, as well as clear presence of the SBS_A^n^T mutational signature in the trinucleotide mutation spectrum. We found statistically significant enrichment for adenines 3 and 4 bp 5’ of thymine mutations in 39 whole-exome and 16 whole-genome samples (Supplemental Data S1). These included tumours of the bladder, colon, rectum, and prostate. Visual inspection of the mutation spectra showed POLE-associated mutagenesis (SBS10a) in 25 tumours, suggesting POLE mutagenesis sometimes shows sequence context specificity similar to SBS_A^n^T (Supplemental Fig S9).

To further increase confidence in the selection of tumours showing SBS_A^n^T we used the mSigAct signature presence test to assess whether SBS_A^n^T (interpreted as a SBS signature in trinucleotide context) was needed to explain candidate observed spectra (Supplemental Data S2) (Ng et al. 2017). Using the signature assignments previously reported for these tumours (Alexandrov et al. 2019), we compared reconstruction of the mutational spectra with and without SBS_A^n^T. We identified 25 tumours which were reconstructed significantly better when including SBS_A^n^T (Fig. 4). In these 25 tumours we identified 53 somatic SBSs that were likely caused by SBS A^n^T mutagenesis and that affected known oncogenes or tumour suppressor genes (Supplemental Table S2). Affected genes included *TP53*, *PTEN*, *KMT2A*, *KMT2C* and *EZH2*. Among the 25 tumours with likely SBS_A^n^T mutations, indel information was only available for the 6 PCAWG whole-genomes (Campbell et al. 2017). Five of these had thymine deletions with the expected sequence contexts (Supplemental Fig S10+S11).

**Fig. 4:**
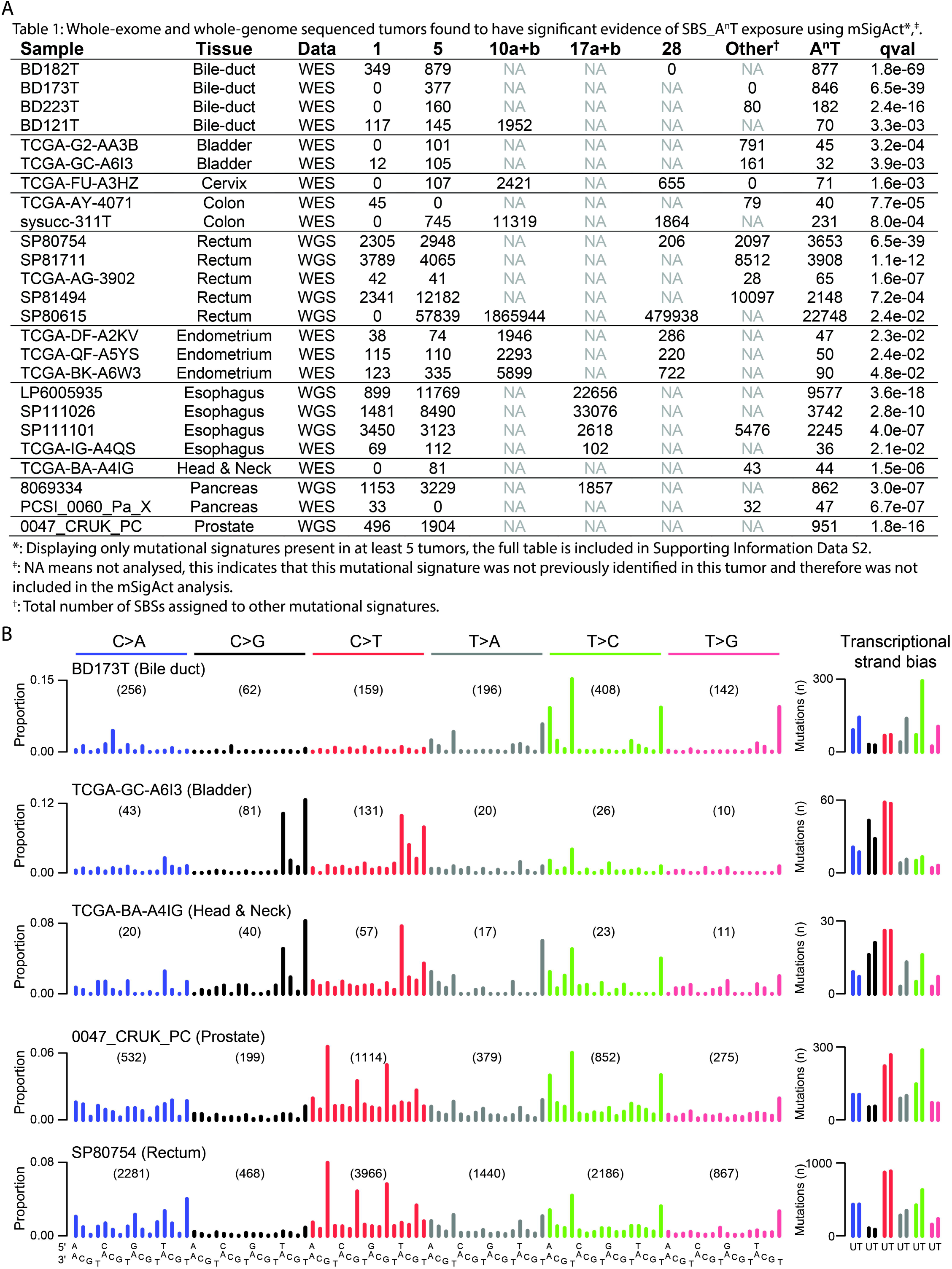
Discovery of SBS_A^n^T in publicly available mutation data. **(A)** Tumours found to be positive for SBS_A^n^T mutagenesis based on sequence context specificity of thymine mutations as well as mSigAct analysis. **(B)** Example spectra of tumours positive for SBS_A^n^T. The right panel shows the transcriptional strand bias (U = Untranscribed strand, T = transcribed strand); for the whole-genome samples, only SBSs in transcribed regions were included. The bile duct, bladder and HNSCC tumours are whole-exome data, the prostate and rectal tumours are whole-genome data.

### Exploring the aetiology of SBS_A^n^T

DNA repair defects can dramatically affect mutational signatures (Volkova et al. 2019). Therefore we first checked for defects in DNA repair genes that could have transformed the appearance of a known mutational process to the mutational signature we observed. We observed MSH6 p.V878A and ATR p.L1483X substitutions (Supplemental Table S2). However, MSH6 p.V878A is predicted to be benign, and ATR p.L1483X was only present at 7.4% variant allele frequency, and therefore could not have accounted for the vast majority of SBS_A^n^T mutations that had higher variant allele frequencies. We therefore concluded that these variants did not play a role in shaping SBS_A^n^T mutagenesis. Moreover, none of the other 25 SBS_A^n^T positive tumours showed mutations in these genes, nor did we observe any other recurrently affected DNA repair genes in these tumours (Supplemental Table S2). We next sought to identify the aetiology of SBS_A^n^T. The enrichment of mutations of T>A on the transcribed strand is indicative of a large molecule that adducts on adenines. Additionally, it is also expected to be an exceptionally large adduct, large enough to reach to 4 basepairs 5’ of the mutated site. Through literature study we identified a class of minor-groove binding compounds called Duocarmycins, which are produced by several species of *Streptomyces*, a common class of bacteria which are known human symbionts (Hurley and Rokem 1983; Ichimura et al. 1991; Seipke et al. 2012). The molecular structure of duocarmycin SA (duoSA), a naturally occurring duocarmycin, is shown in Fig. 5A. Fig. 5B shows duoSA intercalated in the minor-groove of the DNA helix (source: PDB ID: 1DSM) (Smith et al. 2000; Rose et al. 2018). Duocarmycins bind specifically to adenines in A/T-rich regions, which matches SBS_A^n^T’s sequence context (Reynolds et al. 1985; Baraldi et al. 1999; Woynarowski 2002).

**Fig. 5:**
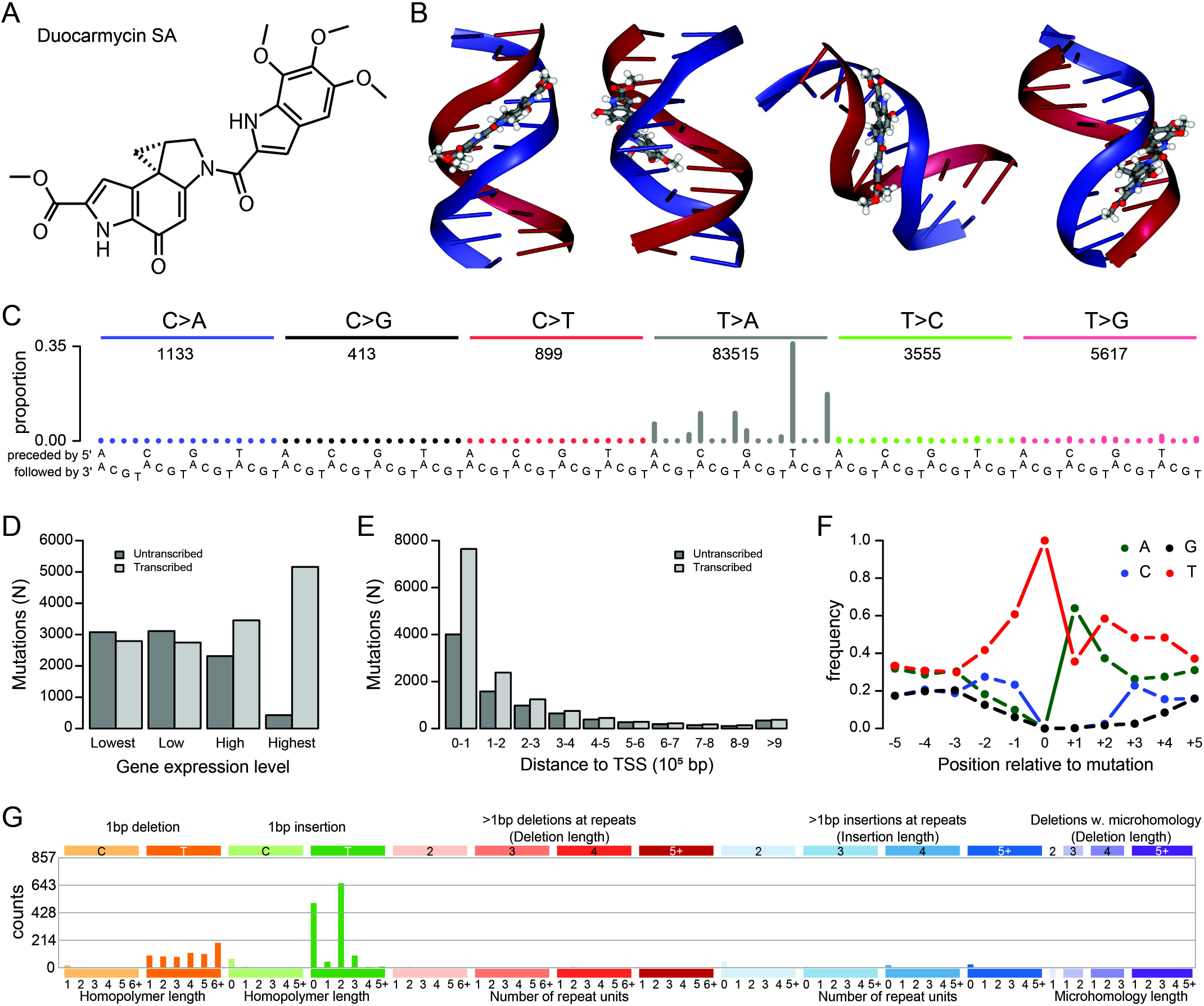
Mutational signature of duocarmycin SA. **(A)** Duocarmycin SA, one of the naturally occurring duocarmycins. **(B)** Several views of the conformation of Duocarmcin SA intercalated with DNA (source: PDB ID: <u>1DSM</u>) (Smith et al. 2000; Rose et al. 2018). Duocarmycins slot into the minor groove of the DNA helix. **(C)** SBS mutation spectrum of one of the duoSA treated HepG2 clones. **(D+E)** Transcriptional strand bias of T>A mutations induced by duoSA as a function of gene expression (D) and distance to transcription start site (E). **(F)** Extended sequence context specificity of T>A mutations induced by duoSA. **(G)** Indel spectrum of one of the duoSA treated HepG2 clones.

To investigate whether duocarmycins could be causing SBS_A^n^T, we sequenced four duoSA exposed HepG2 clones. The mutational spectra of duoSA exposed HepG2 clones are shown in Fig. 5C and Supplemental Fig S12. The mutational signature of duoSA is characterized by strong peaks of T>A mutations with always either an adenine directly 3’ or a thymine directly 5’ of the mutation site. DuoSA mutagenesis showed strong transcriptional strand bias, and extended sequence context preference of thymines 3’ of mutated thymines (Fig. 5D-F). Indels caused by duoSA treatment mainly comprised insertions and deletions of single thymines (Fig. 5G, Supplemental Fig S13). Insertions were mainly found either not next to thymines, or in 2bp thymine repeats (TT>TTT). Deletions occurred in any length of thymine repeats. Additionally we also observed TA>AT, CT>AA and TT>AA DBSs in all clones (Supplemental Fig S14). From these results we concluded that SBS_A^n^T is not caused by duoSA.

### Characterization of the mutational signature in TC1

We also sequenced the whole-genome of TC1, identifying 5,402 SBSs and 67 indels. Besides APOBEC-associated mutations, we observed prominent <u>T</u>G><u>A</u>G peaks and a strong G<u>T</u>G>G<u>C</u>G peak, all with strong transcriptional strand bias (Supplemental Fig S15). No extended sequence context preference was observed (Supplemental Fig S15E). As only the signature of SBS mutations in trinucleotide context was distinctive we screened for cosine similarity between the thymine (T>N) mutations, and T>A mutations specifically for all 23,829 tumours. We found no tumours in which presence of the TC1 mutational signature was visible in the mutation spectrum (Supplemental Fig S16).

### Identification of bacteria causing SBS_A^n^T and TC1 mutagenesis

To identify the bacterial species associated with SBS_A^n^T and TC1 mutagenesis, we extracted all reads from the WGS data that did not align to the human genome, and mapped them to bacterial reference genomes. Less than 0.1% of reads from both normal samples as well as tumour 62074759 were non-human, opposed to 1.5% from tumour TC1. Of the non-human reads, only a small proportion aligned to any of the bacterial genomes (Supplemental Figure S17). Focussing on tumour TC1, we identified several genera of bacteria including *Lachnoanaerobaculum*, *Prevotella*, *Anaerococcus* and *Streptococcus* (Supplemental Figure S17). All these bacterial genera are common oral symbionts (Downes et al. 2008; Labutti et al. 2009; Hedberg et al. 2012; Abranches et al. 2018). Because of the rareness of the mutational signatures discovered in this study, it is unlikely that such common oral bacteria would be causal. To explore whether other microorganisms (such as fungi) could be present, we also performed a nucleotide-BLAST on some of the non-human reads from all samples, but no high-confidence alignments were found.

## Discussion

We analyzed the mutational signatures of 36 Asian OSCCs, hypothesizing that there were still rare mutational processes to be discovered. Interestingly, we identified two novel mutational signatures. Strikingly, these two OSCCs were also the only tumours from our cohort of OSCCs with pathology reports that mentioned high levels of bacterial infection. The rarity of these signatures was illustrated by the fact that we only found 25 additional tumours with SBS_A^n^T and no additional tumours with the TC1 signature after examining a total of 23,829 tumours. In tumours from tissue types where we discovered SBS_A^n^T, only 0.4% showed SBS_A^n^T. Importantly, all tumours in which SBS_A^n^T was detected were from mucosal tissues that harbour bacterial symbionts or that are in direct contact with tissues that harbour symbionts.

Interestingly, since initial publication of this manuscript on bioRxiv, SBS_A^n^T has also been reported in normal colonic crypts from healthy individuals (Lee-Six 2019). SBS_A^n^T mutagenesis was found to be predominantly active early in life, and different patterns of SBS_A^n^T activity distribution over the colon were observed. These results fit with the hypothesis that bacterial compounds could be causing this signature. ‘Patchy’ exposure patterns are unlikely if there had been dietary or occupational exposure to chemicals, and occupational exposure is also improbable since SBS_A^n^T was found to be mostly active early in life. We postulate that early in life, while the microbiome is still being established, bacterial infections might have occurred in these patients. Later in life, microbiome homeostasis may have been established, preventing SBS_A^n^T mutagenesis later in life. For patient 62074759 we propose that the unusual initial treatment of the OSCC before surgery, which included 3 kinds of chemotherapy and radiotherapy, could have opened a window for bacterial infection after the oral microbiome had been disrupted by the treatments. The tumour sample we sequenced, was a recurrence 9 month post treatment. We can exclude the possibility of the treatments causing SBS_AnT, as the mutational signatures of 5-fluorouracil, cisplatin and radiotherapy have already been published, and gemcitabine, a cytosine analogue, would be unlikely to cause thymine mutations (Sherborne et al. 2015; Boot et al. 2018; Christensen 2019).

SBS_A^n^T shows strong transcriptional strand bias, which is commonly observed for mutational processes associated with bulky adducts (Huang et al. 2017; Ng et al. 2017; Boot et al. 2018). The depletion of adenine mutations on the transcribed strand (which corresponds to depletion of thymine mutations on the untranscribed strand) suggests that the mutational process causing SBS_A^n^T involves formation of a bulky adduct on adenine. Fig. 6A and B show a proposed model for adduct formation leading to SBS_A^n^T. The model assumes 2 independent adducts are formed, either directly adjacent to thymine homopolymers (Fig. 6A) or inside adenine homopolymers (Fig. 6B). We propose that adducts inside adenine homopolymers lead to T>A, T>C and T>G mutations in a T<u>T</u>T context as well as deletions of single thymines in thymine homopolymers. Conversely, adducts on adenines directly adjacent to thymine homopolymers would lead to T>A and T>C mutations in the AAAA<u>T</u> context as well as deletions of single thymines not in a homopolymers. The sequences for the adducts in the model are the reverse complement of each other, and we cannot exclude the possibility of an interstrand crosslink. However, if this were the case, we would expect to also observe multiple pairs of SBSs separated by 2 unaffected bases, which we did not.

**Fig. 6:**
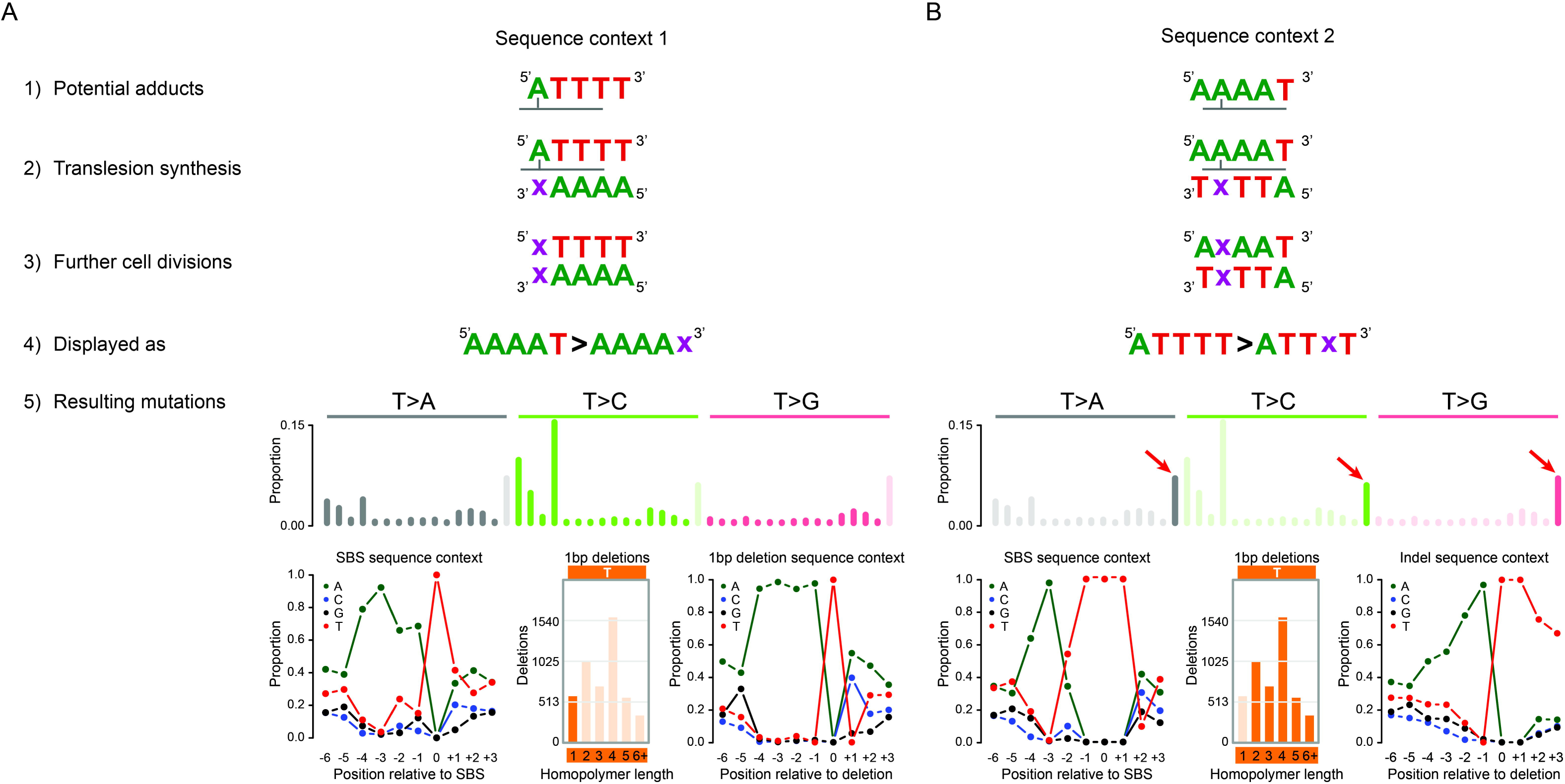
Proposed model for adduct formation leading to the mutation patterns of SBS_A^n^T and ID_A^n^T. **(A)** Potential adduct 1. **1**) Adducts are formed on adenines directly 5’ of thymine homopolymers. **2**) During translesion synthesis, an incorrect nucleotide (**x**) is incorporated opposite the adducted adenine. **3**) During further cell divisions, the mutation is maintained. **4**) Following the conventions of the mutational signature field, we display mutations as occurring from the pyrimidine of the Watson-Crick base pair. **5**) Potential adduct 1 would lead to SBSs directly adjacent to TTT trinucleotides and deletions of single thymines not in a thymine homopolymers. Both SBSs and deletions resulting from potential adduct 1 would be enriched for adenines up to 4 bp 5’ of the mutation site. **(B)** Potential adduct 2. **1**) Adducts are formed in adenine homopolymers with a thymine directly 3’. **2**) During translesion synthesis, an incorrect nucleotide (**x**) is incorporated opposite the adducted adenine. **3**) During further cell divisions, the mutation is maintained. **4**) Following the conventions of the mutational signature field, we display mutations as occurring from the pyrimidine of the Watson-Crick base pair. **5**) Potential adduct 2 would lead to SBSs inside TTT trinucleotides and deletions of single thymines inside thymine homopolymers. SBSs resulting from potential adduct 2 would be strongly enriched for adenines 3bp 5’ of the mutation site. Deletions resulting from potential adduct 2 would be strongly enriched for adenines up to 4 bp 5’ of the mutation site. The latter is due to the possible different locations of the adduct inside the homopolymers. We believe that for longer homopolymers (>3 thymines) the adduct will nearly always be situated opposite the 3^rd^ thymine, making the −3 position (relative to the adduct) the −1 position relative to the thymine homopolymers. This explains the nearly 100% presence of adenines directly 5’ of the thymine homopolymers.

Based on literature research for compounds that could induce mutagenesis with the characteristics of SBS_A^n^T, we experimentally established the mutational signature of duoSA. DuoSA is a naturally occurring minor-groove binding DNA alkylating agent produced by a subset of *Streptomyces* species. As reported, duoSA was highly mutagenic, causing T>A transversions in A/T rich regions (Woynarowski 2002). However, the mutational spectrum was clearly distinct from that of SBS_A^n^T. Additionally, in contrast to SBS_A^n^T, duoSA mutagenesis showed sequence context enrichment 3’ of the mutated site. Indels induced by duoSA were dominated by insertions of thymines, whilst SBS_A^n^T showed exclusively thymine deletions. We concluded that SBS_A^n^T was not caused by duoSA. Interestingly, after initial submission of our manuscript, a study of a large set of metastatic solid tumors was published, which detected the mutational signature of duoSA in two patients (Priestley et al. 2019). These patients had been treated with SYD985, a duocarmycin based antibody-drug conjugate.

In order to identify the bacteria associated with SBS_A^n^T and TC1 mutagenesis, we examined the whole-genome sequencing data for reads that map to bacterial genomes. The TC1 tumour data had a very high number of non-human sequencing reads. However, alignment to a set of 209 bacterial genomes failed to identify the bacteria associated with TC1 mutagenesis. Possibly a different genus of bacteria is present in this patient, for which the reference genome sequence is yet to be elucidated. In the 62074759 tumour data we also observed a low number of reads aligning to the same genera of bacteria also observed in tumour TC1. The absence of large numbers of non-human reads in tumour 62074759 is likely due to sampling, if the DNA we sequenced was from the centre of the tumour mass opposed to the edge, less contamination would be expected. Ideally, we would have tested the saliva of patients 62074759 and TC1 to identify the bacteria that cause SBS_A^n^T and TC1 mutagenesis, but no saliva samples were stored for these patients.

Bacteria have long been known to be associated with cancer. However, for most associations such as the association between *Salmonella* and gallbladder and colon cancer, and the association between *Chlamydia* and cervical carcinoma, only epidemiological evidence exists (van Elsland and Neefjes 2018). The only bacterium for which experimental evidence exists that it causes cancer is *Heliobacter pylori*, which has been shown to cause gastric cancer in gerbils (Watanabe et al. 1998). *H. pylori*, as well as most other cancer-associated bacteria are thought to stimulate carcinogenesis through the inflammation associated with the infection (van Elsland and Neefjes 2018). However, some bacteria have been reported to produce toxins able to induce double-strand DNA breaks (van Elsland and Neefjes 2018). For OSCC the association with bacterial infection is well known, but no mutagenic compounds have been reported to be produced by these bacteria (Karpinski 2019). An alternative mechanism through which (oral) bacteria could induce mutagenesis is by metabolizing ethanol. There have been several reports of conversion of ethanol into acetaldehyde by oral bacteria (Yokoi et al. 2015; Tagaino et al. 2019). Acetaldehyde is a known mutagen, forming DNA adducts mainly on guanines (Brooks and Zakhari 2014; Mizumoto et al. 2017). Due to acetaldehyde’s propensity to form adducts on guanines, it is unlikely to be the causal agent of SBS_A^n^T which primarily involves adenines and thymines.

In summary, we identified 2 novel mutational signatures in Asian OSCCs that had presented with strong oral bacterial infections. In the other 34 Asian OSCCs, of which none had presented with strong bacterial infections, no novel mutational signatures were discovered. Discovery of one of the novel mutational signatures; SBS_A^n^T, in 25 tumours from publicly available sequencing data confirmed the rarity of this mutational process. Importantly, these 25 tumours were all from tissues either harbouring or in direct contact with tissues that are known to harbour bacterial symbionts. This strongly supports our hypothesis that this mutagenic process is associated with bacterial infection. While our manuscript was in revision, a preprint was released that describes the sequence context specificity of double strand breaks induced by the bacterial toxin colibactin (Dziubańska-Kusibab et al. 2019). Colibactin-adduct-induced double-strand breaks were strongly enriched in AT-rich regions, with the AAWWTT motif to be most enriched at colibactin induced double-strand breaks, which fits exactly with the sequence context specificity we observed for the thymine mutations in 62074759 as shown in Figure 2B panel 2, positions −4 to +1 relative to the mutation site. Therefore, we conclude that SBS_A^n^T is most likely caused by colibactin. Furthermore, the transcriptional strand bias shown by SBS_A^n^T suggests that TC-NER can remove colibactin adducts during transcriptional elongation.

## Materials and Methods

### Samples

De-identified fresh frozen tissue samples and matching whole-blood were collected from OSCC patients operated on between 2012 and 2016 at the National Cancer Centre Singapore. In accordance with the Helsinki Declaration of 1975, written consent for research use of clinical material and clinico-pathologic data was obtained at the time of surgery. This study was approved by the SingHealth Centralized Institutional Review Board (CIRB 2007/438/B).

### Whole-exome and whole-genome sequencing

Whole-exome sequencing was performed at Novogene. (Beijing, China) on a HiSeq X Ten instrument with 150bp paired-end reads. Whole-genome sequencing was performed at BGI (Hong Kong) on the BGIseq500 platform, generating 100bp paired-end reads.

### Alignment and variant calling

Sequencing reads were trimmed by Trimmomatic (Bolger et al. 2014). Alignment and variant calling and filtering was performed as described previously (Boot et al. 2018). Annotation of somatic variants was performed using annovar (Wang et al. 2010). Sequencing reads that did not align to the human genome were subsequently aligned to 209 bacterial reference genomes from Ensembl (ftp://ftp.ensemblgenomes.org/pub/bacteria/release-35/fasta/bacteria_183_collection/). For driver gene analysis, only variants inside Tier 1 genes of the cancer gene census were considered (Sondka et al. 2018).

### Validation of SBSs by Sanger sequencing

We performed Sanger sequencing to validate 96 variants detected in the whole-exome sequencing of sample 62074759. We selected variants with >15% allele frequency to avoid variants below the detection limit of Sanger sequencing and excluded variants immediately adjacent to a homopolymer of ≥9 bp. PCR product purification and Sanger sequencing was performed at GENEWIZ^©^ (Suzhou, China).

### Signature assignment

We assigned mutational signatures to the mutational spectra of the 30 OSCCs with ≥10 mutations using SigProfiler and the SigProfiler reference mutational signatures (Alexandrov et al. 2019). As OSCC is a subset of HNSCC, all mutational signatures that were identified in HNSCCs and OSCCs in the International Cancer Genome Consortium’s Pan Cancer Analysis Working Group (PCAWG) analysis were included for reconstruction (Alexandrov et al. 2019). As the PCAWG mutational signatures are based on the trinucleotide abundance of the human genome, when analysing whole-exome sequencing data we adjusted to the mutational signature for exome trinucleotide frequency.

### Gene expression data

Single cell gene expression data for OSCC was downloaded from NCBI GSE103322 (Puram et al. 2017). We took the median gene expression for all tumour cells as the representative expression level of OSCCs.

### Identification of additional tumours with the signature in 62074759

Previously compiled whole-exome (N=19,184) and whole-genome (N=4,645) sequencing data was screened for presence of the signature in 62074759 (Alexandrov et al. 2019). This included 2,780 whole-genomes from the Pan Cancer of Whole Genomes consortium (Campbell et al. 2017), and 9,493 whole-exomes from the TCGA consortium (Ellrott et al. 2018). We examined tumours with ≥50 (exomes) or ≥500 (genomes) thymine mutations, to identify enrichment for mutations with the 5’ sequence context characteristic of the signature in 62074759 (Supplemental Data S1).

We then used the mSigAct signature presence test to test for the signature in 62074759 amongst the candidate tumours identified in the previous step (Supplemental Data S2) (Ng et al. 2017; Boot et al. 2018). This test provides a *p* value for the null hypothesis that a signature is not needed to explain an observed spectrum compared to the alternative hypothesis that the signature is needed.

### *In vitro* Duocarmycin SA exposure

Exposure of HepG2 cells to Duocarmycin SA was performed as described previously (Boot et al. 2018). In short, HepG2 cells were exposed to 100pM and 250pM Duocarmycin SA for 2 months followed by single cell cloning. For each concentration, 2 clones were whole-genome sequenced. Duocarmycin SA (CAS: 130288-24-3) was obtained from BOC Sciences (New York, USA).

### Data availability

FASTQ files for all patient sequencing data are at the European Genome-phenome Archive under accession EGAS00001003131. FASTQ files for the duocarmycin SA treated HepG2 clones are at the European Nucleotide archive under accession ERP116345.

## Abbreviations

bp: base pair
indels: insertions and deletions
SBS: single base substitutions
DBS: double base substitutions
HNSCC: head and neck squamous cell carcinoma
OSCC: oral squamous cell carcinoma
PCAWG: Pan-Cancer Analysis of Whole Genomes
TC-NER: transcription coupled nucleotide excision repair
VAF: variant allele frequency

## Acknowledgements

The results here are partly based on data generated by the TCGA Research Network (http://cancergenome.nih.gov/) and data assembled by the International Cancer Genome Consortium Pan Cancer Analysis Working Groups. This study was funded by NMRC/CIRG/1422/2015 to SGR.

## Authors’ contributions

AB and SGR designed the study, drafted the manuscript and prepared figures. AB, AWTN and WY performed bioinformatics analyses. SH performed cell line experiments. FTC, DSWT and NGI contributed materials. All authors read and approved the manuscript.

