## Supplemental Figs for "Identification of novel mutational signatures in Asian oral squamous cell carcinomas associated with bacterial infections"

| | |
|---|---|
| Supplemental_Fig_S1 | 2 |
| Supplemental_Fig_S2 | 6 |
| Supplemental_Fig_S3 | 7 |
| Supplemental_Fig_S4 | 8 |
| Supplemental_Fig_S5 | 9 |
| Supplemental_Fig_S6 | 10 |
| Supplemental_Fig_S7 | 11 |
| Supplemental_Fig_S8 | 12 |
| Supplemental_Fig_S9 | 13 |
| Supplemental_Fig_S10 | 20 |
| Supplemental_Fig_S11 | 21 |
| Supplemental_Fig_S12 | 27 |
| Supplemental_Fig_S13 | 28 |
| Supplemental_Fig_S14 | 29 |
| Supplemental_Fig_S15 | 30 |
| Supplemental_Fig_S16 | 31 |
| Supplemental_Fig_S17 | 38 |

Supplemental_Fig_S1: **SNS mutational spectra of the exome sequencing data from 30 OSCCs analyzed in this study.**

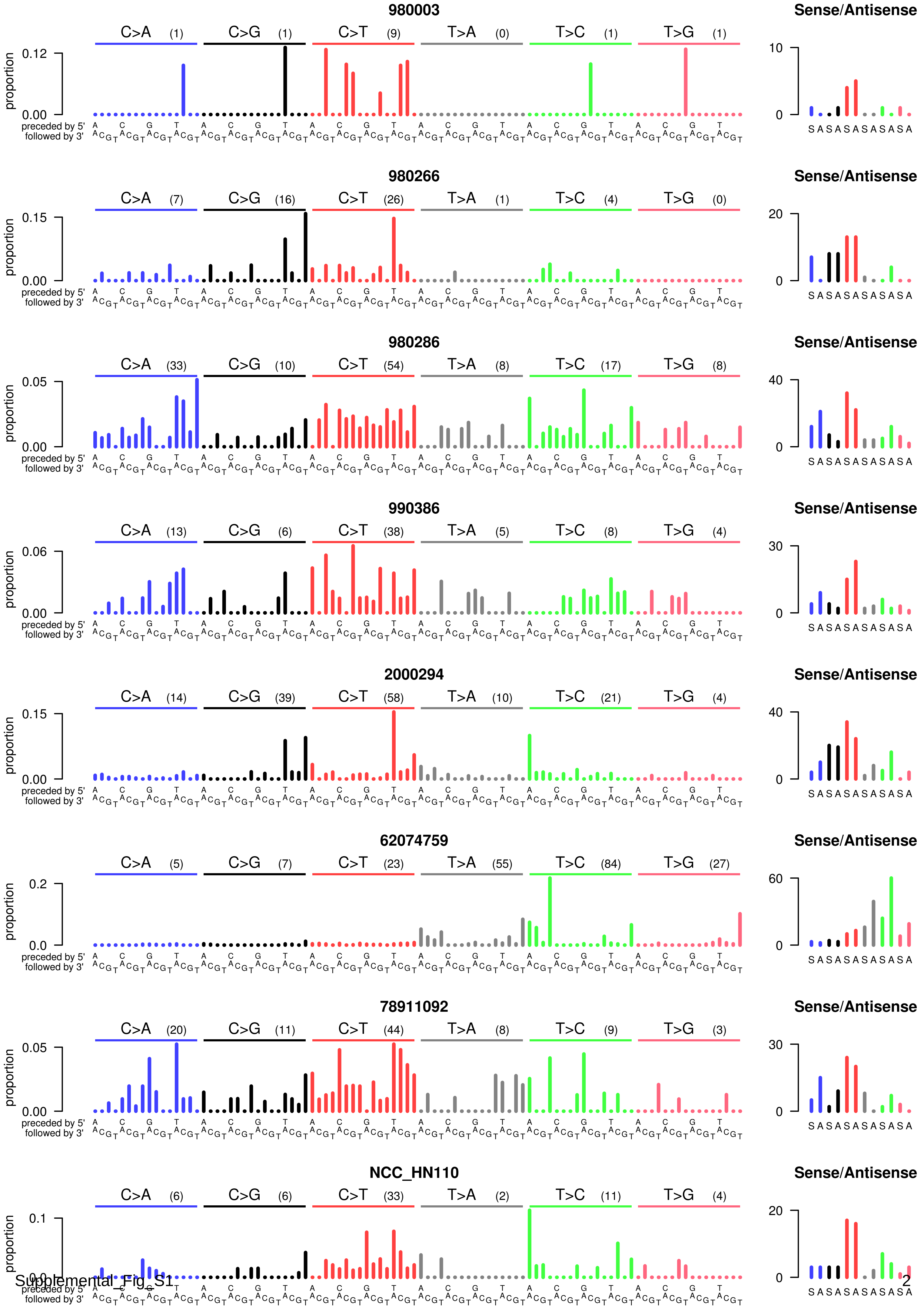

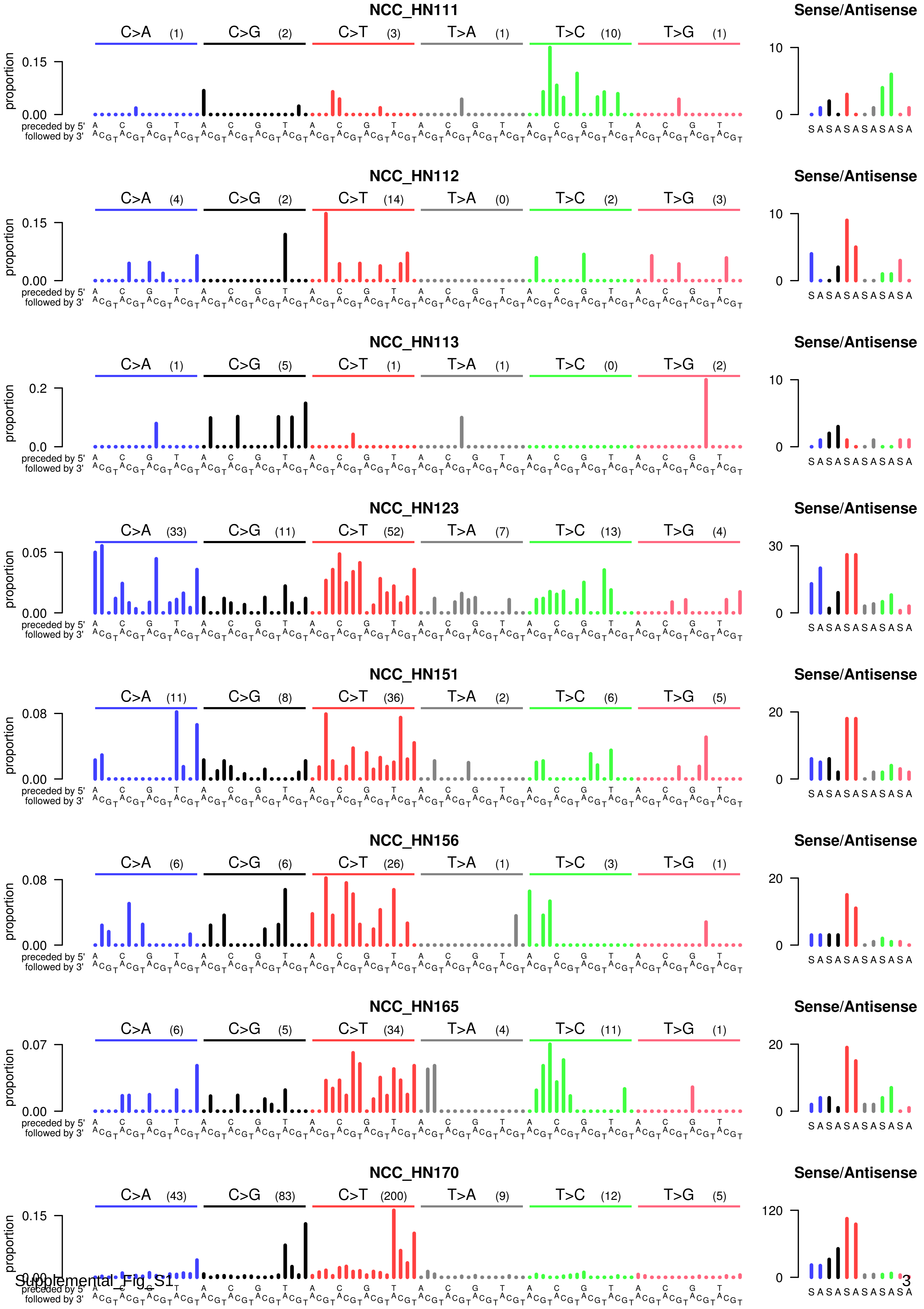

NCC_HN111
C>A (1)
C>G (2)
C>T (3)
T>A (1)
T>C (10)
T>G (1)
proportion
0.15
0.00
preceded by 5'
followed by 3'
Sense/Antisense
10
0
NCC_HN112
C>A (4)
C>G (2)
C>T (14)
T>A (0)
T>C (2)
T>G (3)
0.15
0.00
10
0
NCC_HN113
C>A (1)
C>G (5)
C>T (1)
T>A (1)
T>C (0)
T>G (2)
0.2
0.0
10
0
NCC_HN123
C>A (33)
C>G (11)
C>T (52)
T>A (7)
T>C (13)
T>G (4)
0.05
0.00
30
0
NCC_HN151
C>A (11)
C>G (8)
C>T (36)
T>A (2)
T>C (6)
T>G (5)
0.08
0.00
20
0
NCC_HN156
C>A (6)
C>G (6)
C>T (26)
T>A (1)
T>C (3)
T>G (1)
0.08
0.00
20
0
NCC_HN165
C>A (6)
C>G (5)
C>T (34)
T>A (4)
T>C (11)
T>G (1)
0.07
0.00
20
0
NCC_HN170
C>A (43)
C>G (83)
C>T (200)
T>A (9)
T>C (12)
T>G (5)
0.15
0.00
120
0
S A S A S A S A S A S A

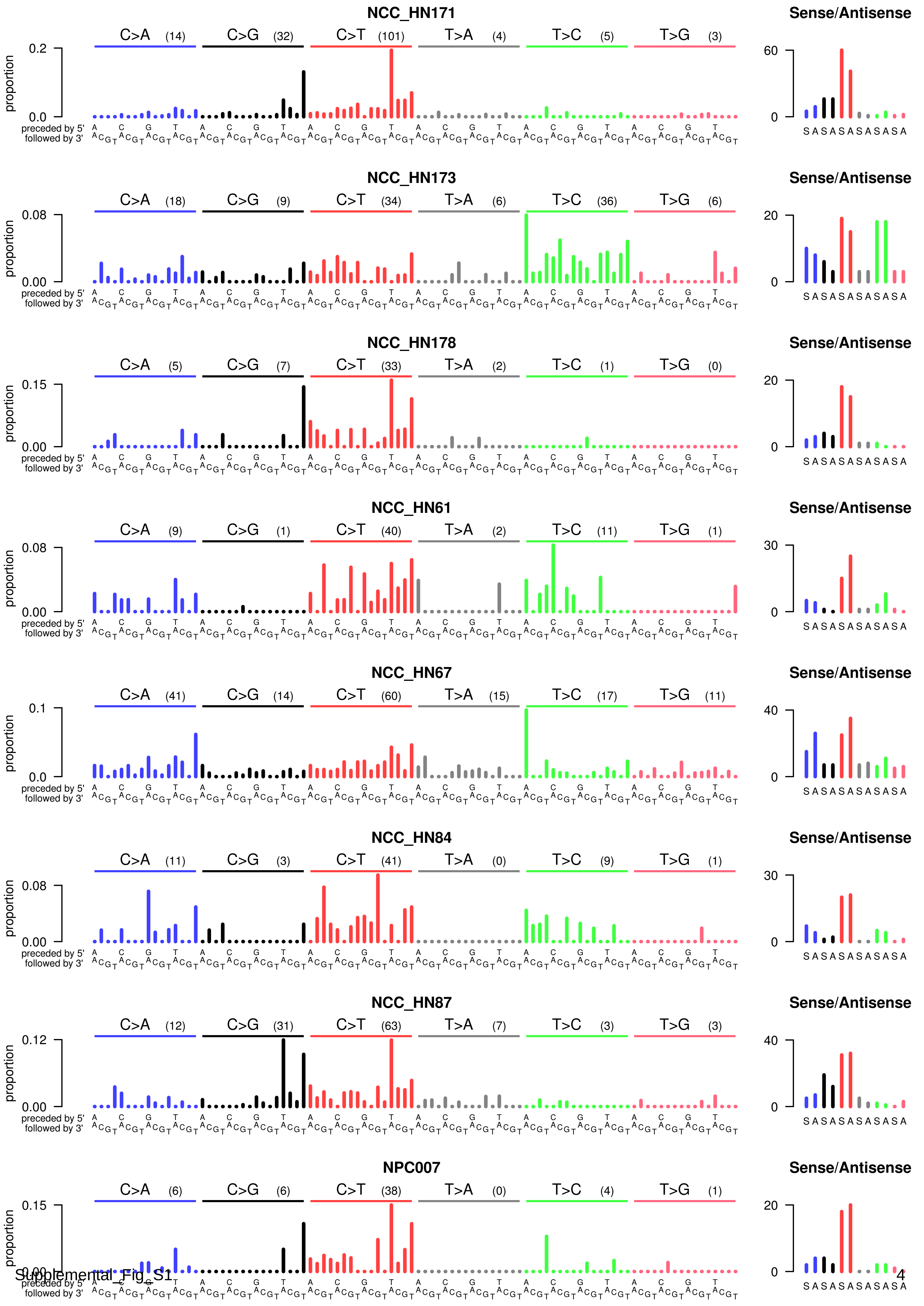
NCC_HN171
C>A (14) C>G (32) C>T (101) T>A (4) T>C (5) T>G (3)
Sense/Antisense
NCC_HN173
C>A (18) C>G (9) C>T (34) T>A (6) T>C (36) T>G (6)
Sense/Antisense
NCC_HN178
C>A (5) C>G (7) C>T (33) T>A (2) T>C (1) T>G (0)
Sense/Antisense
NCC_HN61
C>A (9) C>G (1) C>T (40) T>A (2) T>C (11) T>G (1)
Sense/Antisense
NCC_HN67
C>A (41) C>G (14) C>T (60) T>A (15) T>C (17) T>G (11)
Sense/Antisense
NCC_HN84
C>A (11) C>G (3) C>T (41) T>A (0) T>C (9) T>G (1)
Sense/Antisense
NCC_HN87
C>A (12) C>G (31) C>T (63) T>A (7) T>C (3) T>G (3)
Sense/Antisense
NPC007
C>A (6) C>G (6) C>T (38) T>A (0) T>C (4) T>G (1)
Sense/Antisense
proportion
preceded by 5'
followed by 3'
S A S A S A S A S A S A

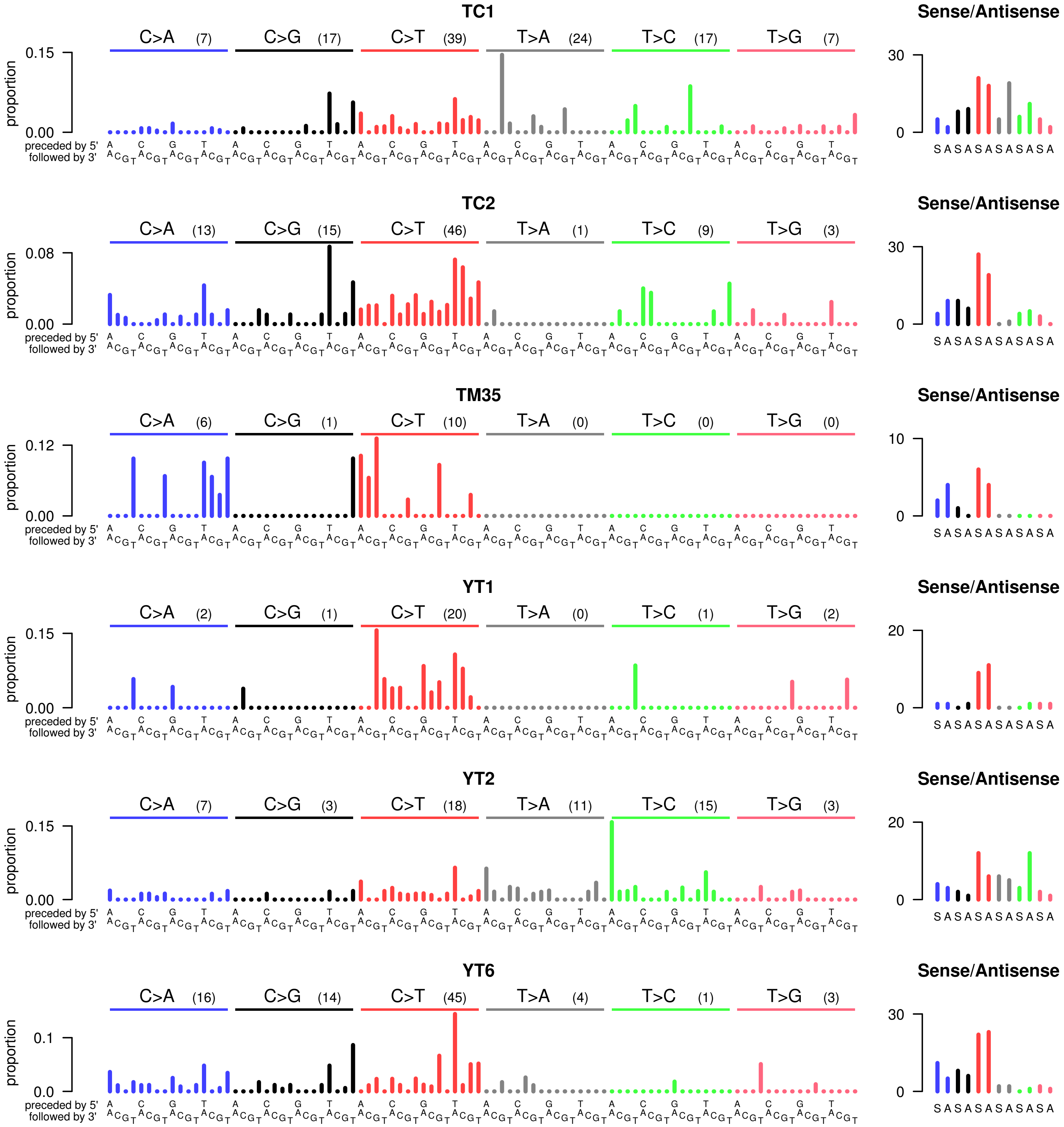
TC1
C>A (7) C>G (17) C>T (39) T>A (24) T>C (17) T>G (7)
TC2
C>A (13) C>G (15) C>T (46) T>A (1) T>C (9) T>G (3)
TM35
C>A (6) C>G (1) C>T (10) T>A (0) T>C (0) T>G (0)
YT1
C>A (2) C>G (1) C>T (20) T>A (0) T>C (1) T>G (2)
YT2
C>A (7) C>G (3) C>T (18) T>A (11) T>C (15) T>G (3)
YT6
C>A (16) C>G (14) C>T (45) T>A (4) T>C (1) T>G (3)
Sense/Antisense
proportion
preceded by 5'
followed by 3'
S A S A S A S A S A S A

Supplemental_Fig_S2: **Mutational signature analysis of 30 OSCCs. A**: Reconstruction of the patient mutation spectra using the mutational signatures established by the PCAWG consortium 1. **B**: Evaluation of reconstruction by cosine similarity and Pearson's r.

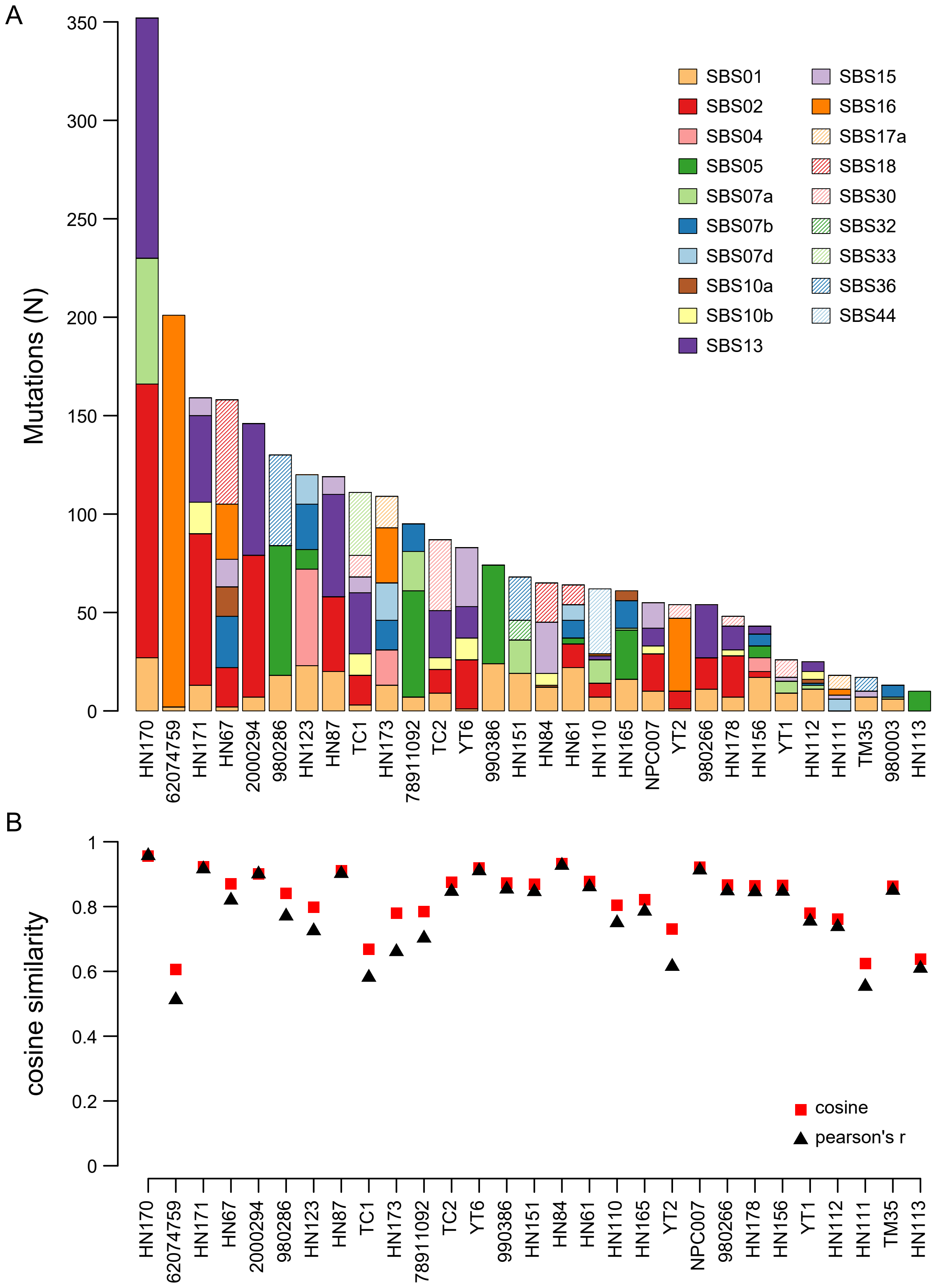

Supplemental_Fig_S3: **T-SNE clustering of singaporean OSCCs and TCGA HNSCCs**.

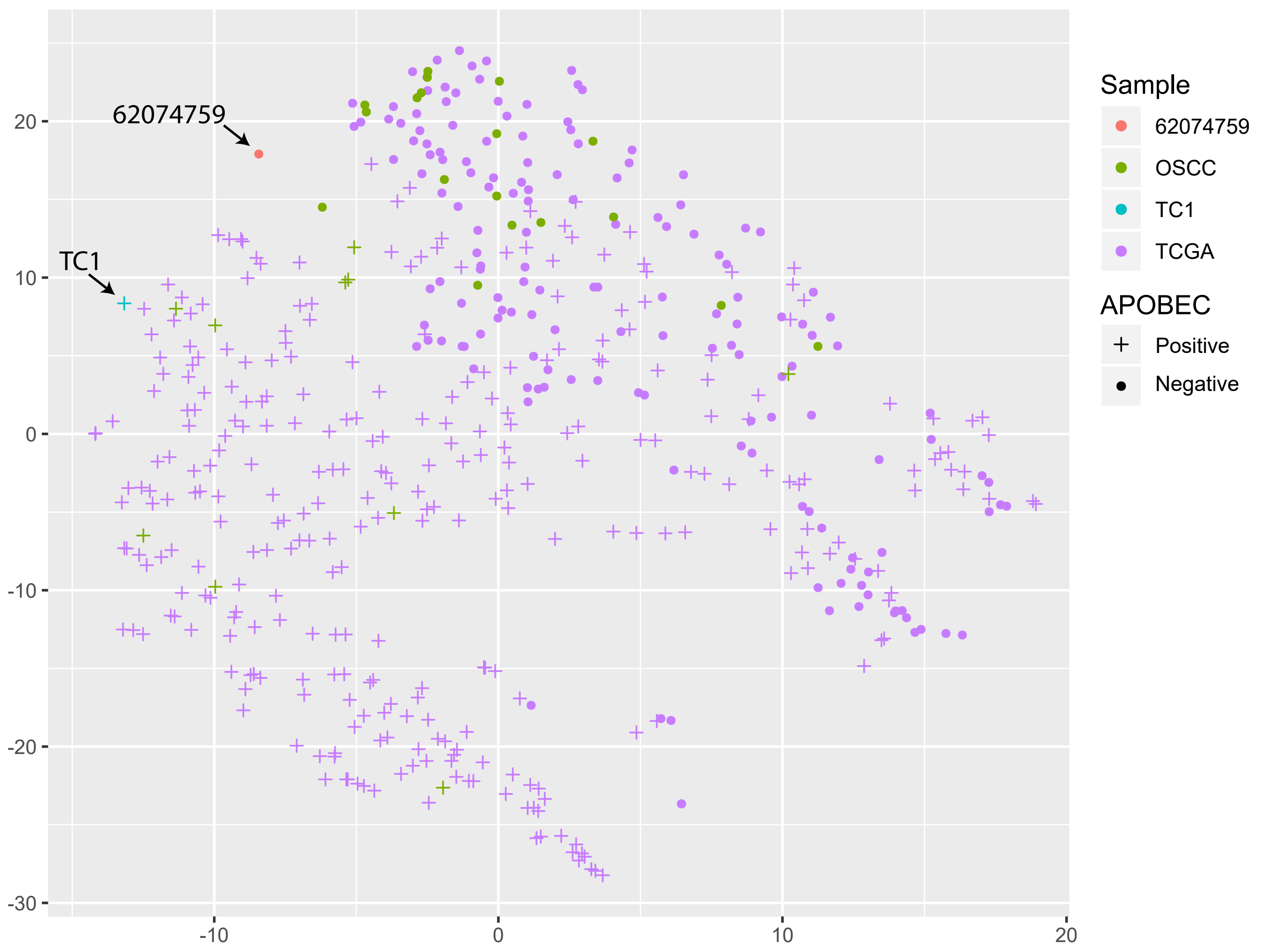

Supplemental_Fig_S4: **SNS sequence context of all (top), all T>N (bottom left) and all C>N (bottom right) mutations in sample 62074759.**

**Sequence context of all mutations in 62074759**

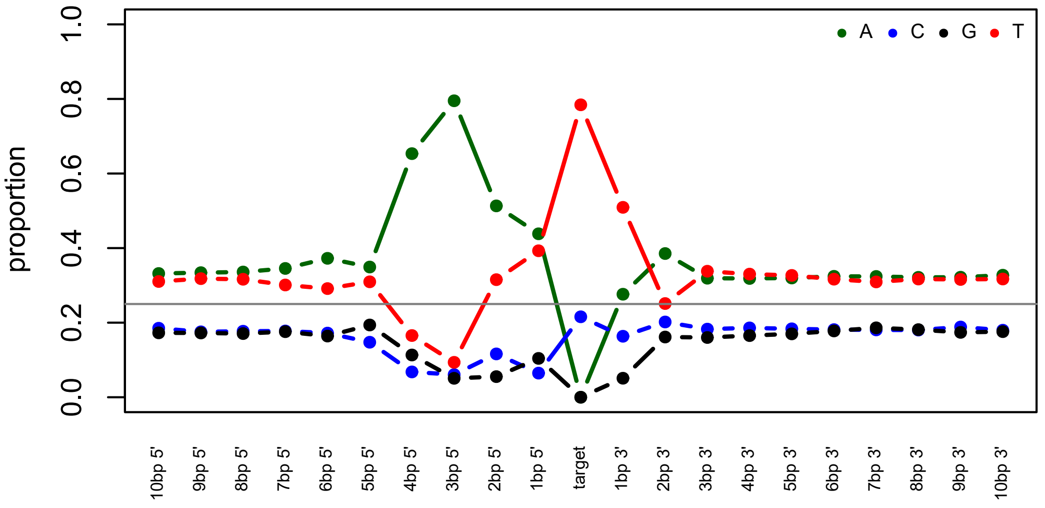


**Sequence context of T>N mutations in 62074759**

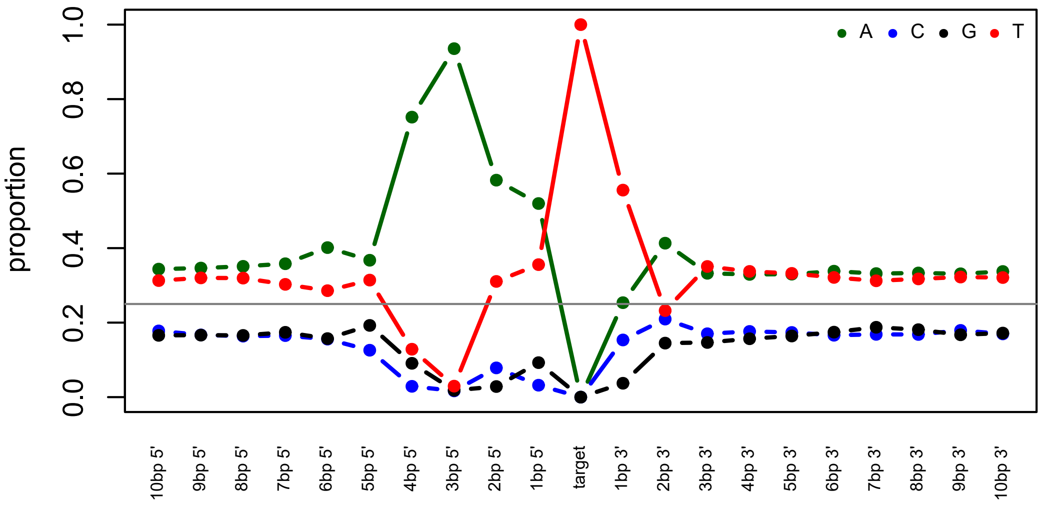


**Sequence context of C>N mutations in 62074759**

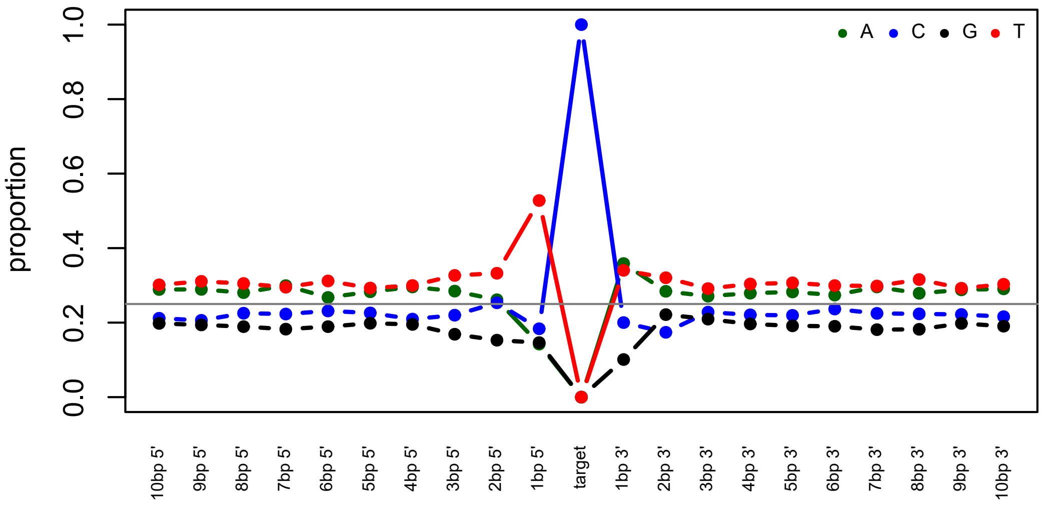

Supplemental_Fig_S5: **Variant allele frequency analysis of mutations in sample 62074759 suggestive of ongoing mutagenesis.**
(**A**) Variant allele frequency of mutations in sample 62074759.
(**B**) The mutational spectrum of sample 62074759 is similar across different variant allele frequencies.
(**C**) Table of cosine similarities of the different spectra shown in B

A

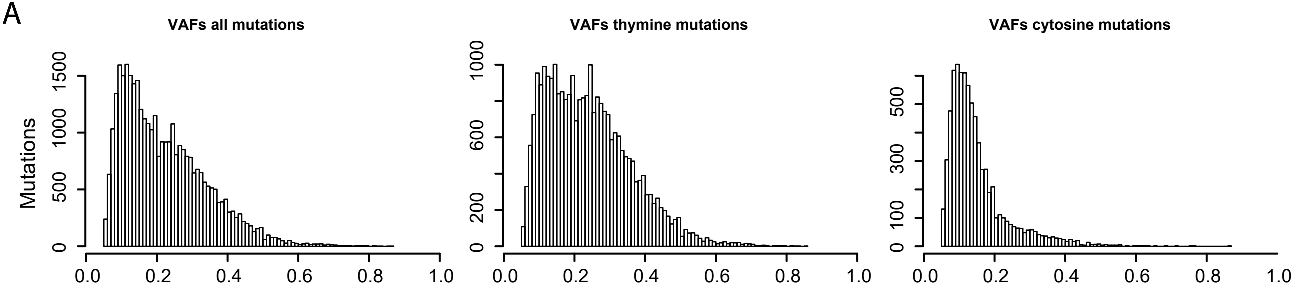


B

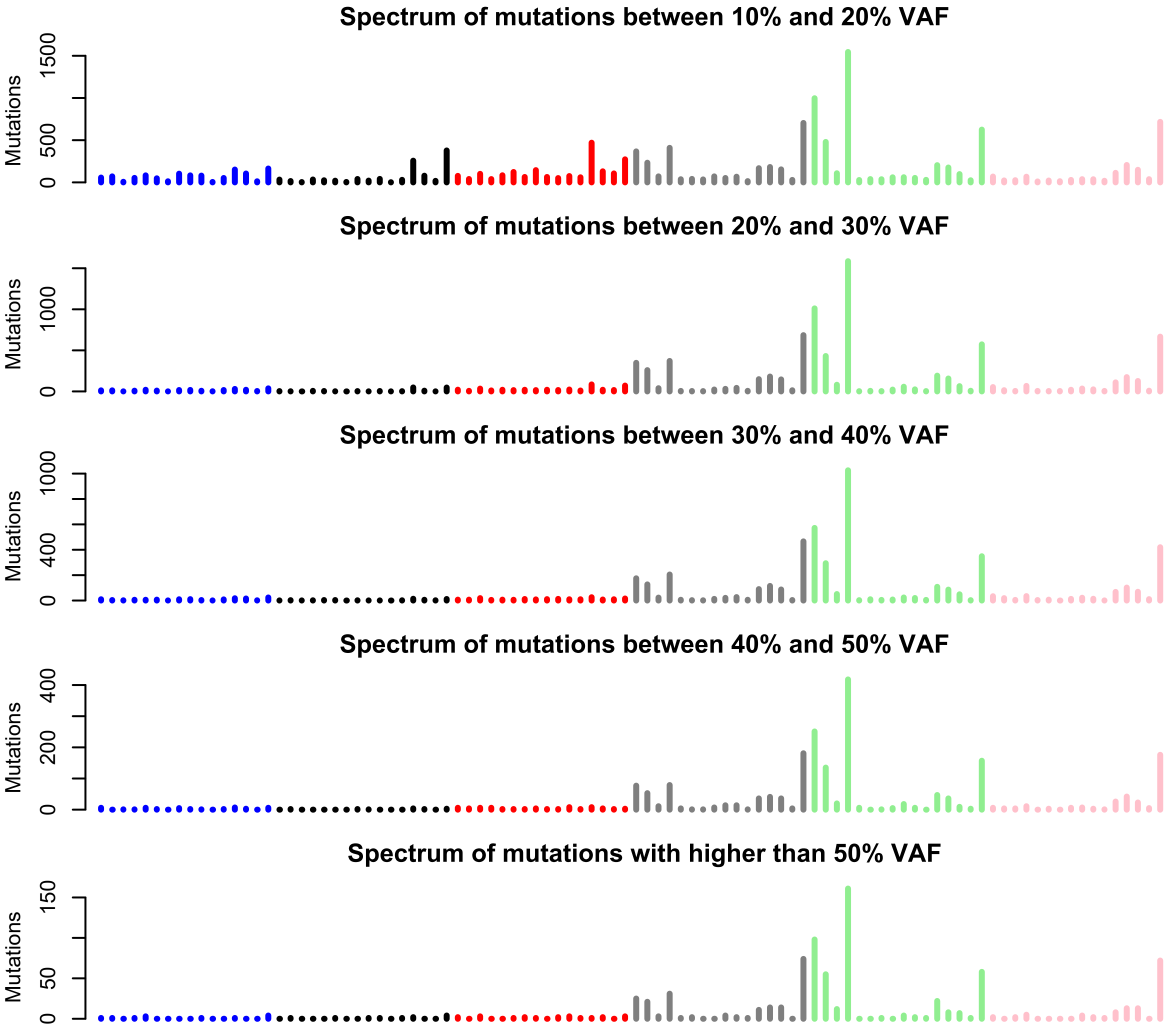


C

**Similarity of spectra of mutations with different VAFs (cosine similarity)**

| all | 10-20% | 20-30% | 30-40% | 40-50% | >50% | |
|---|---|---|---|---|---|---|
| | 0.988 | 0.990 | 0.985 | 0.983 | 0.981 | all |
| | | 0.958 | 0.948 | 0.945 | 0.943 | 10-20% |
| | | | 0.997 | 0.996 | 0.994 | 20-30% |
| | | | | 0.998 | 0.997 | 30-40% |
| | | | | | 0.997 | 40-50% |
| | | | | | | >50% |

Supplemental_Fig_S6: **Sequence context of deletions of single thymines in thymine-repeats in sample 62074759.**

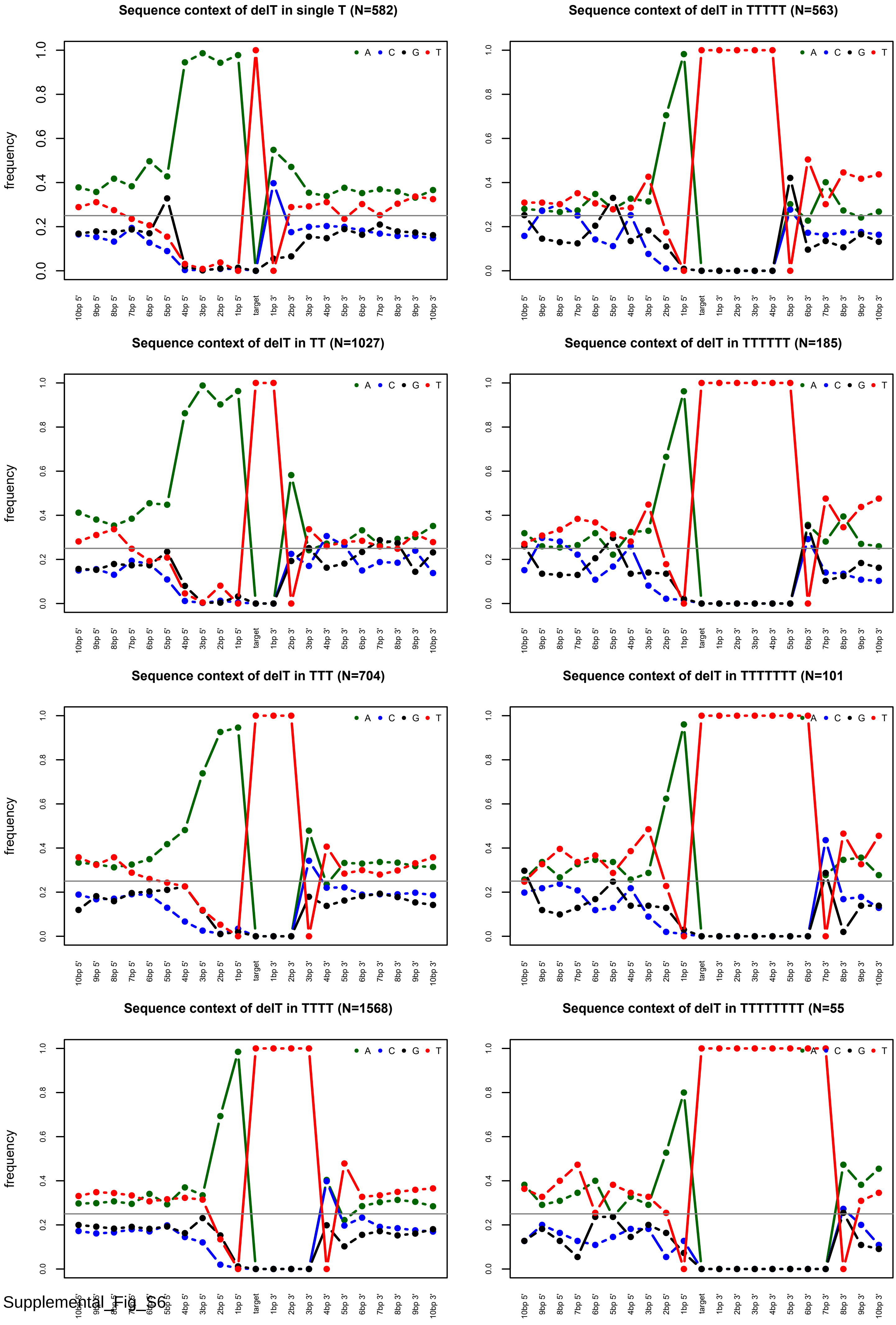

Supplemental_Fig_S7: **Transcriptional strand bias of deletions of a single thymine in thymine-repeats ranging from 1 to 8 thymines.**

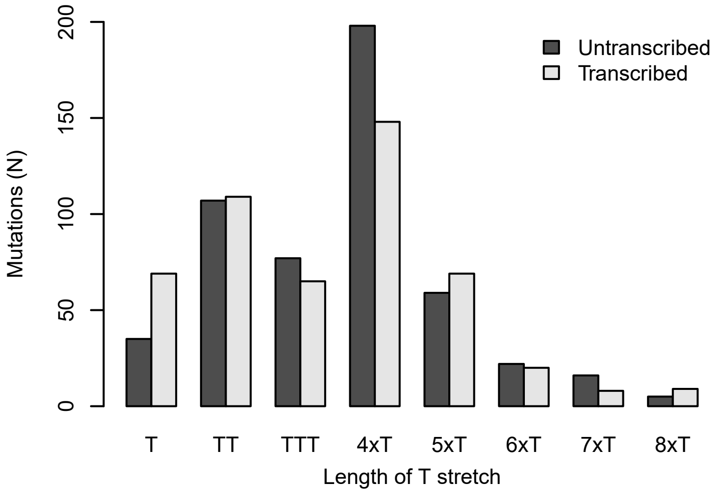


Binomial test results for transcriptional strand bias of deletions of thymines in sample 62074759

| | Transcribed | Untranscribed | pval |
|---|---|---|---|
| T | 68 | 35 | 0.001 |
| TT | 109 | 106 | 0.892 |
| TTT | 65 | 77 | 0.356 |
| TTTT | 148 | 198 | 0.008 |
| TTTTT | 69 | 59 | 0.426 |
| TTTTTT | 20 | 22 | 0.878 |
| TTTTTTT | 8 | 15 | 0.210 |
| TTTTTTTT | 9 | 5 | 0.424 |

Supplemental_Fig_S8: **Dinucleotide substitution spectrum of sample 62074759.**

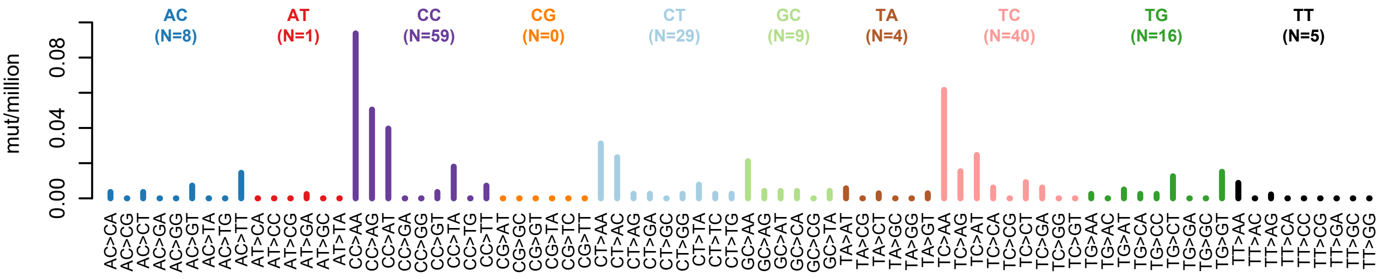

Supplemental_Fig_S9: **Mutational spectra of tumours from publicly available sequencing data showing likely exposure to SBS_AnT identified by examining the sequence context of thymine mutations in these samples.** Samples in which SBS_AnT was detected using mSigAct are marked with a *, samples with SBS10a presence are marked with a P

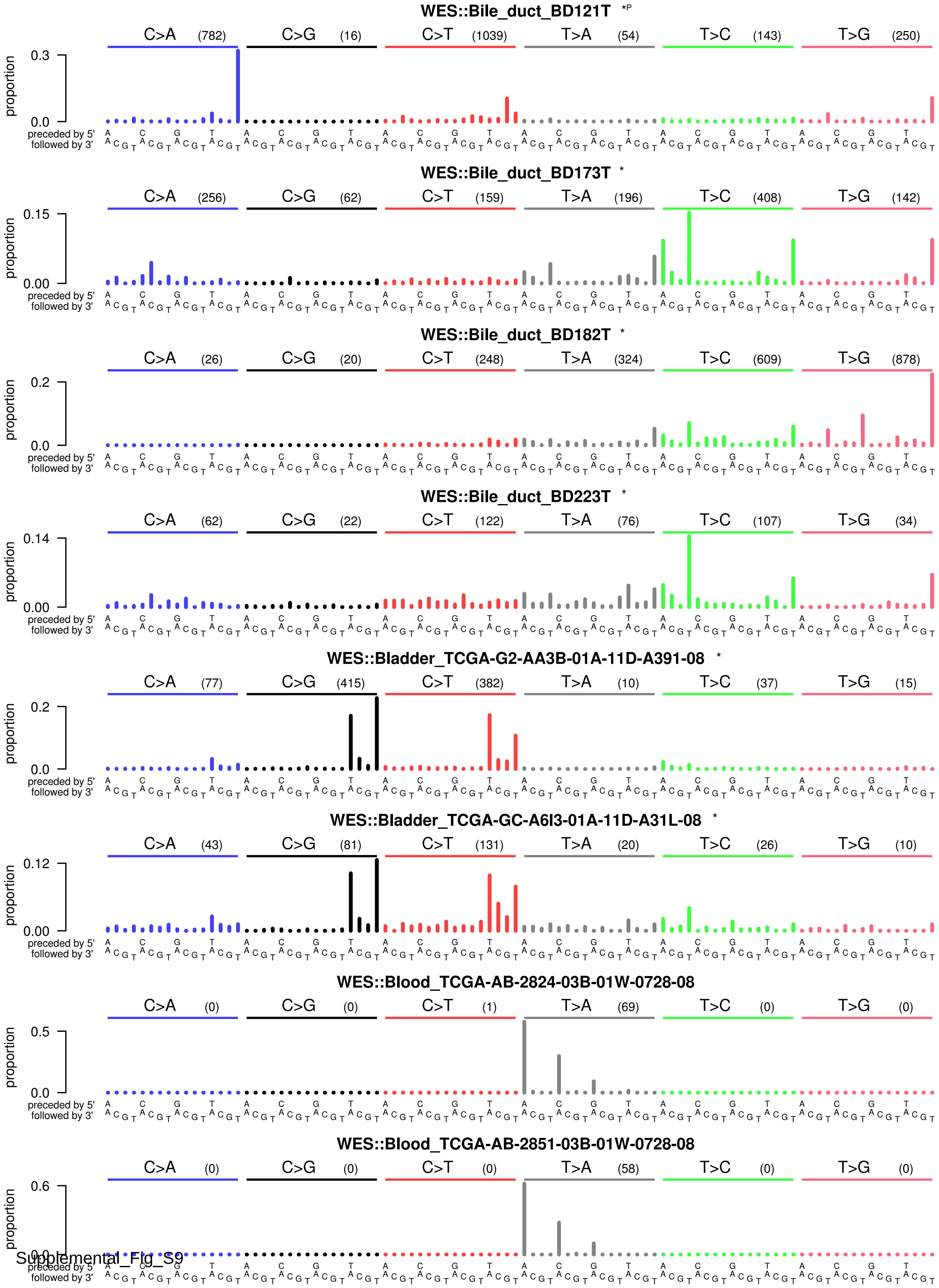

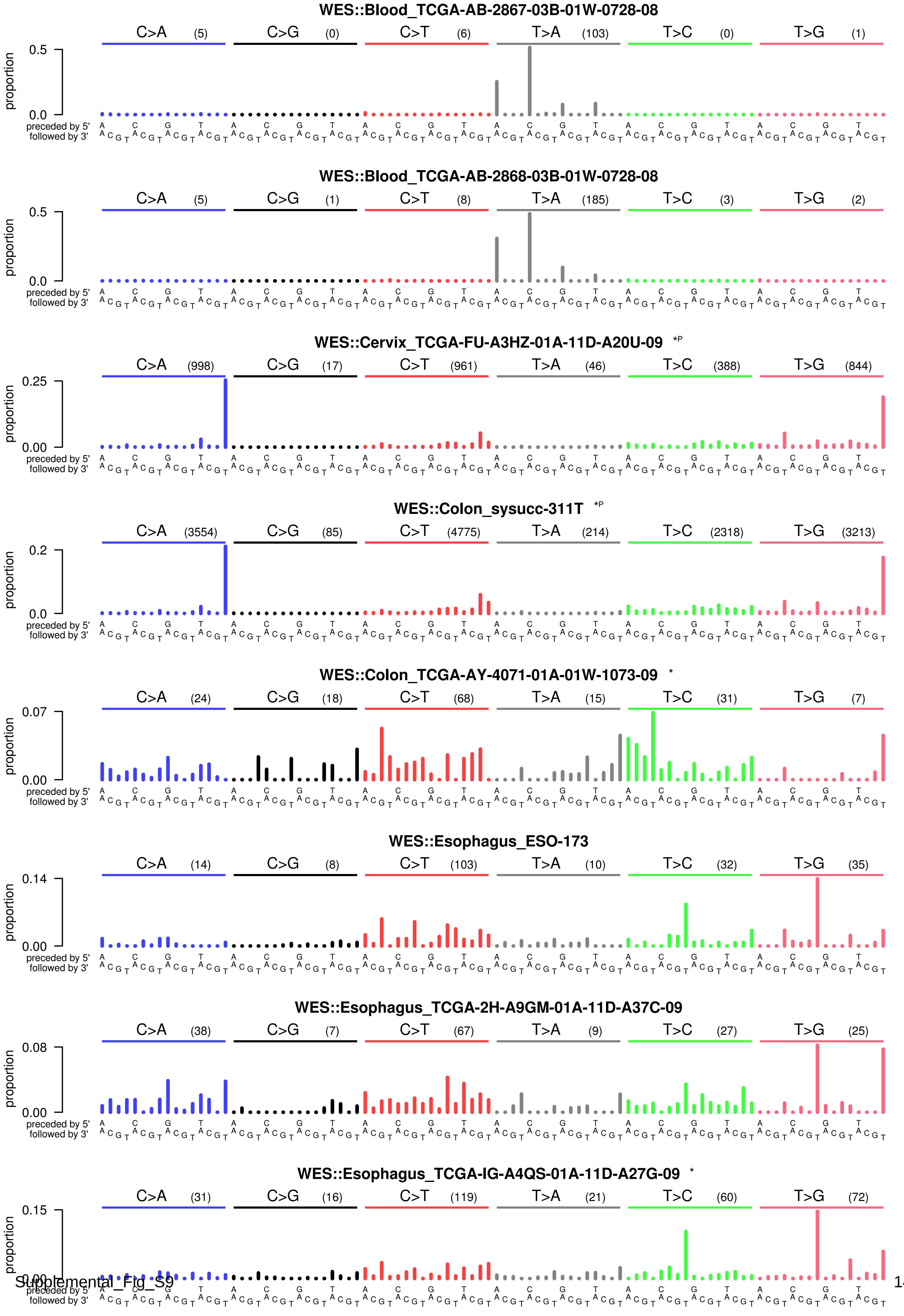
WES::Blood_TCGA-AB-2867-03B-01W-0728-08
C>A (5) C>G (0) C>T (6) T>A (103) T>C (0) T>G (1)
proportion
0.5
0.0
preceded by 5'
followed by 3'
WES::Blood_TCGA-AB-2868-03B-01W-0728-08
C>A (5) C>G (1) C>T (8) T>A (185) T>C (3) T>G (2)
proportion
0.5
0.0
preceded by 5'
followed by 3'
WES::Cervix_TCGA-FU-A3HZ-01A-11D-A20U-09 *P
C>A (998) C>G (17) C>T (961) T>A (46) T>C (388) T>G (844)
proportion
0.25
0.00
preceded by 5'
followed by 3'
WES::Colon_sysucc-311T *P
C>A (3554) C>G (85) C>T (4775) T>A (214) T>C (2318) T>G (3213)
proportion
0.2
0.0
preceded by 5'
followed by 3'
WES::Colon_TCGA-AY-4071-01A-01W-1073-09 *
C>A (24) C>G (18) C>T (68) T>A (15) T>C (31) T>G (7)
proportion
0.07
0.00
preceded by 5'
followed by 3'
WES::Esophagus_ESO-173
C>A (14) C>G (8) C>T (103) T>A (10) T>C (32) T>G (35)
proportion
0.14
0.00
preceded by 5'
followed by 3'
WES::Esophagus_TCGA-2H-A9GM-01A-11D-A37C-09
C>A (38) C>G (7) C>T (67) T>A (9) T>C (27) T>G (25)
proportion
0.08
0.00
preceded by 5'
followed by 3'
WES::Esophagus_TCGA-IG-A4QS-01A-11D-A27G-09 *
C>A (31) C>G (16) C>T (119) T>A (21) T>C (60) T>G (72)
proportion
0.15
0.00
preceded by 5'
followed by 3'

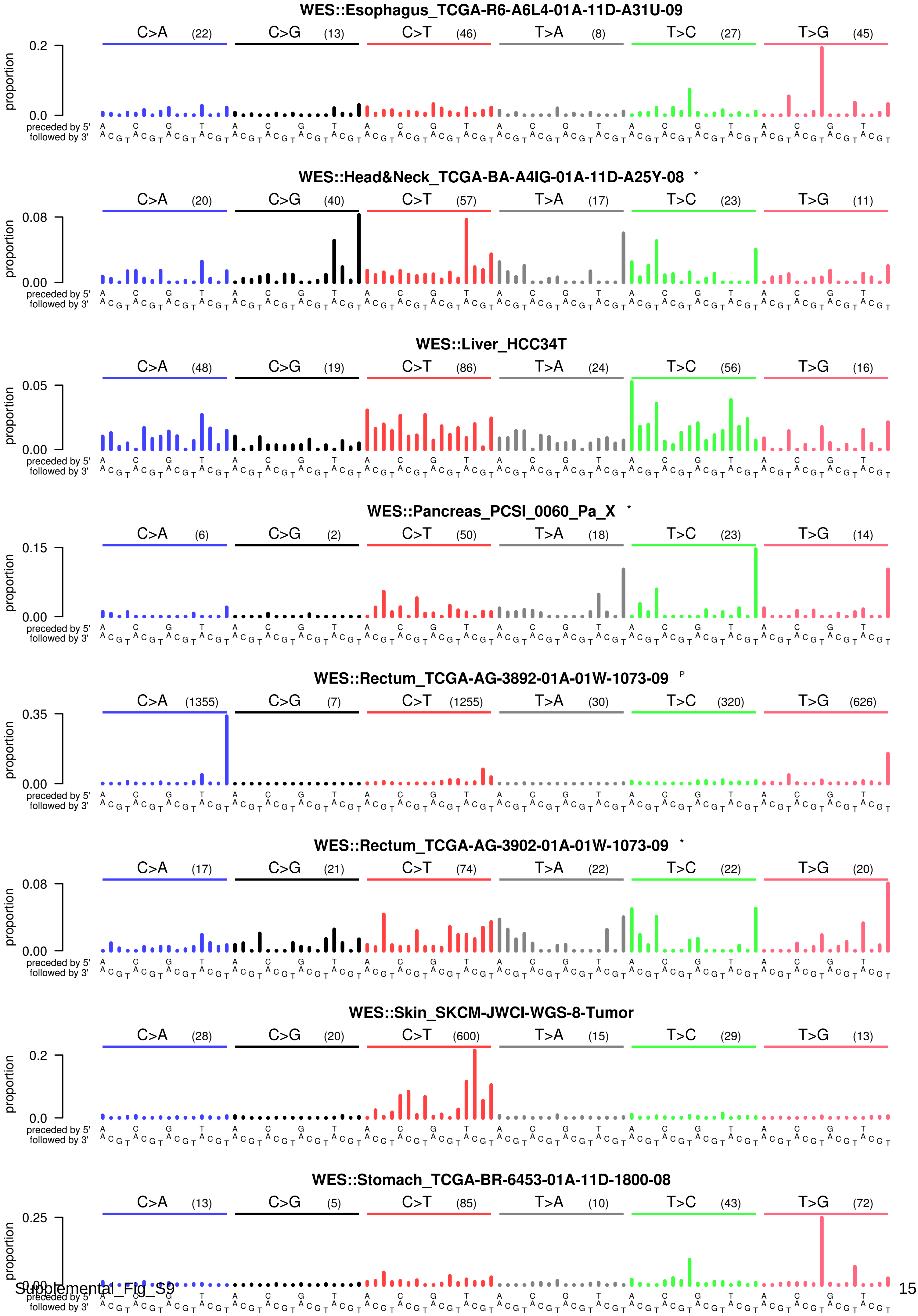

WES::Esophagus_TCGA-R6-A6L4-01A-11D-A31U-09
C>A (22) C>G (13) C>T (46) T>A (8) T>C (27) T>G (45)
proportion
0.2
0.0
preceded by 5'
followed by 3'
WES::Head&Neck_TCGA-BA-A4IG-01A-11D-A25Y-08 *
C>A (20) C>G (40) C>T (57) T>A (17) T>C (23) T>G (11)
0.08
0.00
WES::Liver_HCC34T
C>A (48) C>G (19) C>T (86) T>A (24) T>C (56) T>G (16)
0.05
0.00
WES::Pancreas_PCSI_0060_Pa_X *
C>A (6) C>G (2) C>T (50) T>A (18) T>C (23) T>G (14)
0.15
0.00
WES::Rectum_TCGA-AG-3892-01A-01W-1073-09 P
C>A (1355) C>G (7) C>T (1255) T>A (30) T>C (320) T>G (626)
0.35
0.00
WES::Rectum_TCGA-AG-3902-01A-01W-1073-09 *
C>A (17) C>G (21) C>T (74) T>A (22) T>C (22) T>G (20)
0.08
0.00
WES::Skin_SKCM-JWCI-WGS-8-Tumor
C>A (28) C>G (20) C>T (600) T>A (15) T>C (29) T>G (13)
0.2
0.0
WES::Stomach_TCGA-BR-6453-01A-11D-1800-08
C>A (13) C>G (5) C>T (85) T>A (10) T>C (43) T>G (72)
0.25
0.00

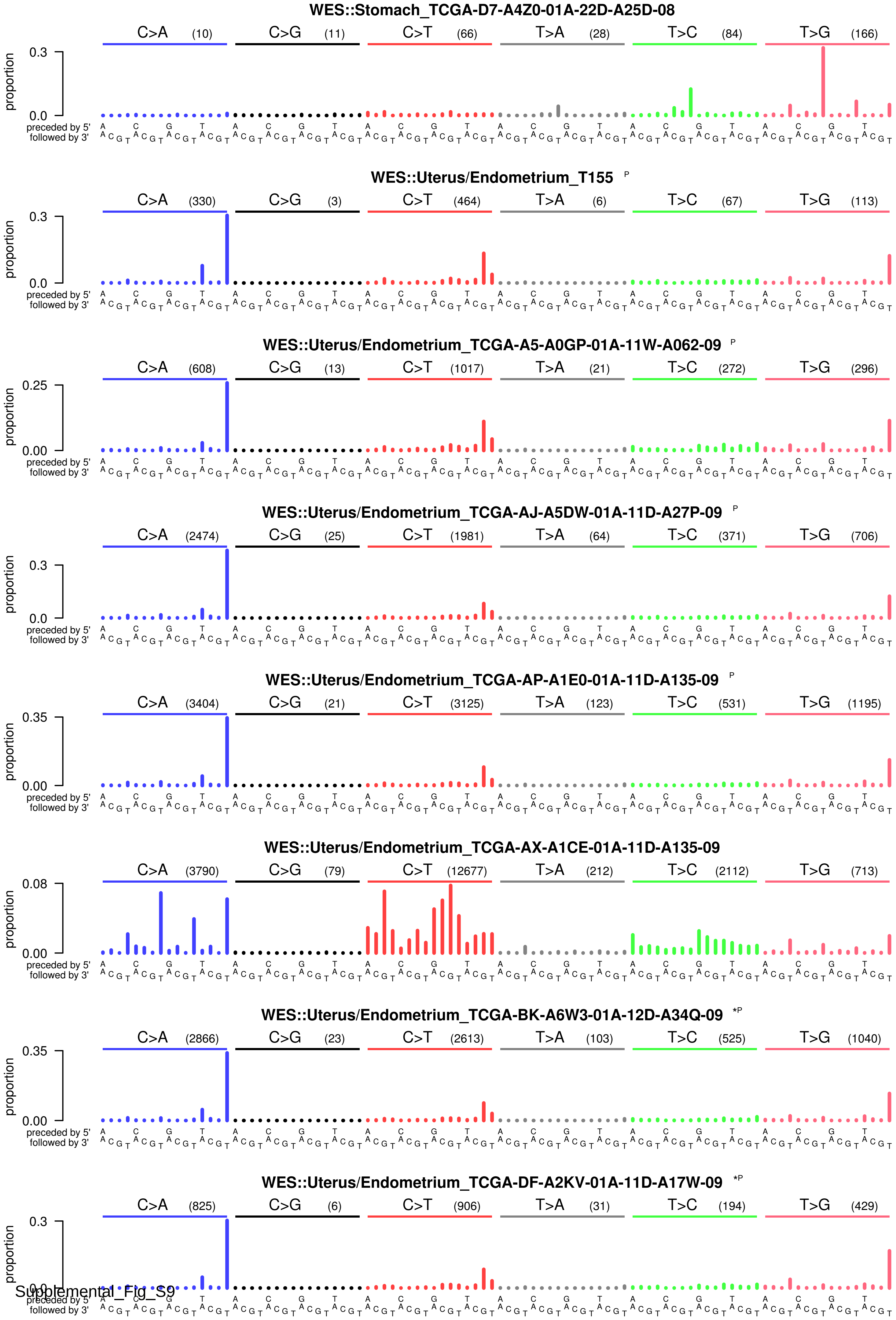
WES::Stomach_TCGA-D7-A4Z0-01A-22D-A25D-08
C>A (10) C>G (11) C>T (66) T>A (28) T>C (84) T>G (166)
WES::Uterus/Endometrium_T155 P
C>A (330) C>G (3) C>T (464) T>A (6) T>C (67) T>G (113)
WES::Uterus/Endometrium_TCGA-A5-A0GP-01A-11W-A062-09 P
C>A (608) C>G (13) C>T (1017) T>A (21) T>C (272) T>G (296)
WES::Uterus/Endometrium_TCGA-AJ-A5DW-01A-11D-A27P-09 P
C>A (2474) C>G (25) C>T (1981) T>A (64) T>C (371) T>G (706)
WES::Uterus/Endometrium_TCGA-AP-A1E0-01A-11D-A135-09 P
C>A (3404) C>G (21) C>T (3125) T>A (123) T>C (531) T>G (1195)
WES::Uterus/Endometrium_TCGA-AX-A1CE-01A-11D-A135-09
C>A (3790) C>G (79) C>T (12677) T>A (212) T>C (2112) T>G (713)
WES::Uterus/Endometrium_TCGA-BK-A6W3-01A-12D-A34Q-09 *P
C>A (2866) C>G (23) C>T (2613) T>A (103) T>C (525) T>G (1040)
WES::Uterus/Endometrium_TCGA-DF-A2KV-01A-11D-A17W-09 *P
C>A (825) C>G (6) C>T (906) T>A (31) T>C (194) T>G (429)
proportion
preceded by 5'
followed by 3'

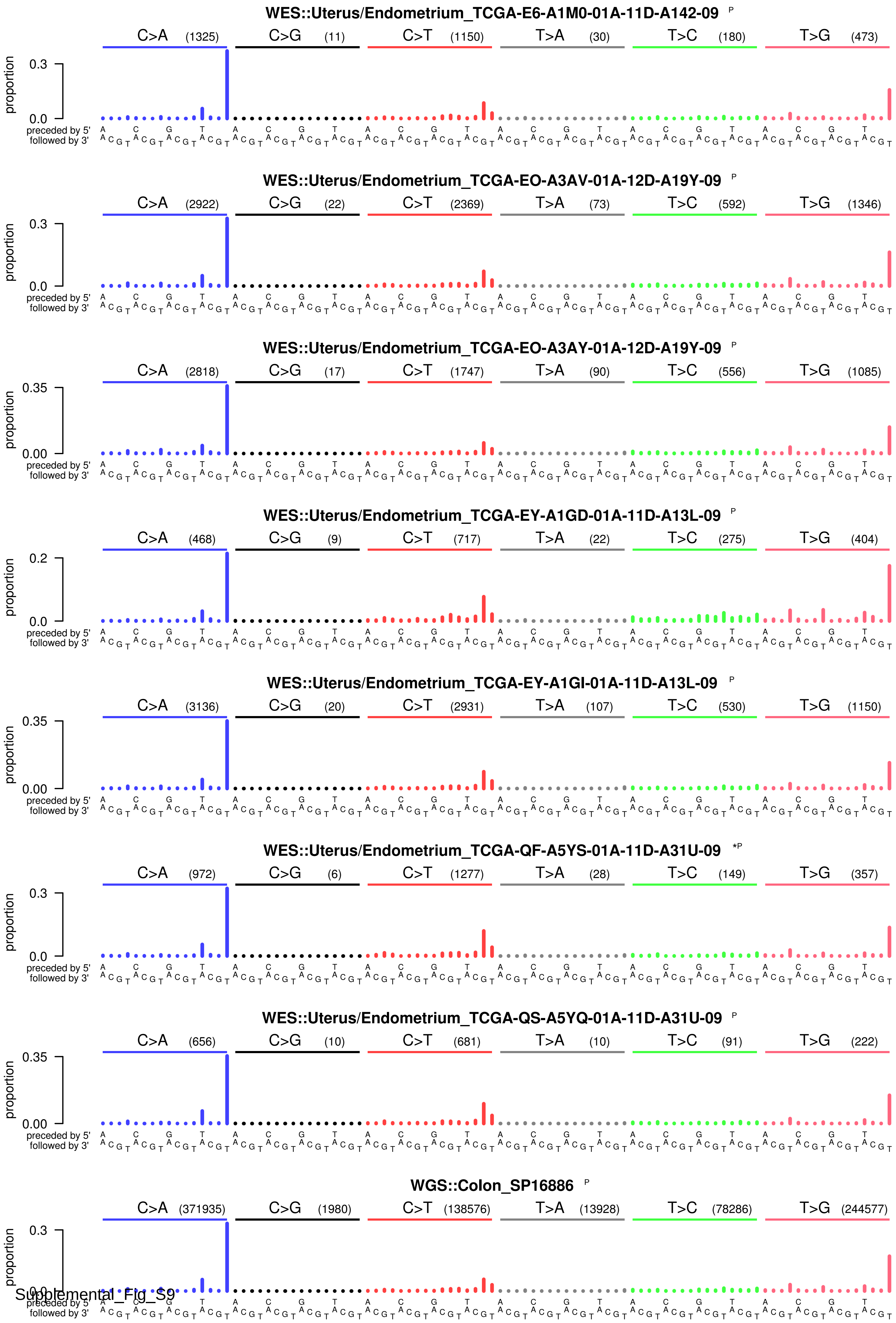
WES::Uterus/Endometrium_TCGA-E6-A1M0-01A-11D-A142-09 P
C>A (1325) C>G (11) C>T (1150) T>A (30) T>C (180) T>G (473)
WES::Uterus/Endometrium_TCGA-EO-A3AV-01A-12D-A19Y-09 P
C>A (2922) C>G (22) C>T (2369) T>A (73) T>C (592) T>G (1346)
WES::Uterus/Endometrium_TCGA-EO-A3AY-01A-12D-A19Y-09 P
C>A (2818) C>G (17) C>T (1747) T>A (90) T>C (556) T>G (1085)
WES::Uterus/Endometrium_TCGA-EY-A1GD-01A-11D-A13L-09 P
C>A (468) C>G (9) C>T (717) T>A (22) T>C (275) T>G (404)
WES::Uterus/Endometrium_TCGA-EY-A1GI-01A-11D-A13L-09 P
C>A (3136) C>G (20) C>T (2931) T>A (107) T>C (530) T>G (1150)
WES::Uterus/Endometrium_TCGA-QF-A5YS-01A-11D-A31U-09 *P
C>A (972) C>G (6) C>T (1277) T>A (28) T>C (149) T>G (357)
WES::Uterus/Endometrium_TCGA-QS-A5YQ-01A-11D-A31U-09 P
C>A (656) C>G (10) C>T (681) T>A (10) T>C (91) T>G (222)
WGS::Colon_SP16886 P
C>A (371935) C>G (1980) C>T (138576) T>A (13928) T>C (78286) T>G (244577)
proportion
preceded by 5'
followed by 3'

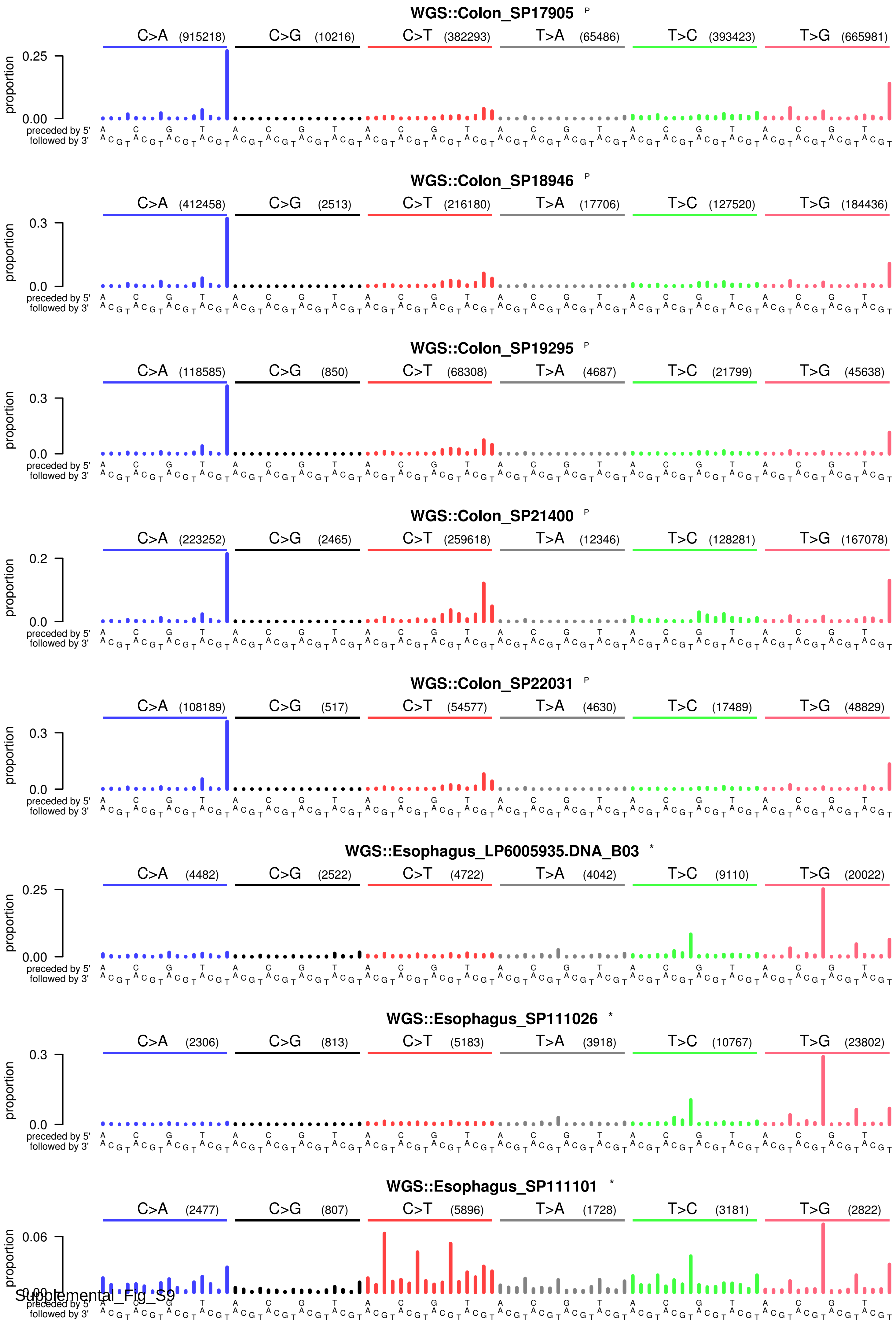
WGS::Colon_SP17905 P
C>A (915218) C>G (10216) C>T (382293) T>A (65486) T>C (393423) T>G (665981)
WGS::Colon_SP18946 P
C>A (412458) C>G (2513) C>T (216180) T>A (17706) T>C (127520) T>G (184436)
WGS::Colon_SP19295 P
C>A (118585) C>G (850) C>T (68308) T>A (4687) T>C (21799) T>G (45638)
WGS::Colon_SP21400 P
C>A (223252) C>G (2465) C>T (259618) T>A (12346) T>C (128281) T>G (167078)
WGS::Colon_SP22031 P
C>A (108189) C>G (517) C>T (54577) T>A (4630) T>C (17489) T>G (48829)
WGS::Esophagus_LP6005935.DNA_B03 *
C>A (4482) C>G (2522) C>T (4722) T>A (4042) T>C (9110) T>G (20022)
WGS::Esophagus_SP111026 *
C>A (2306) C>G (813) C>T (5183) T>A (3918) T>C (10767) T>G (23802)
WGS::Esophagus_SP111101 *
C>A (2477) C>G (807) C>T (5896) T>A (1728) T>C (3181) T>G (2822)
proportion
preceded by 5'
followed by 3'

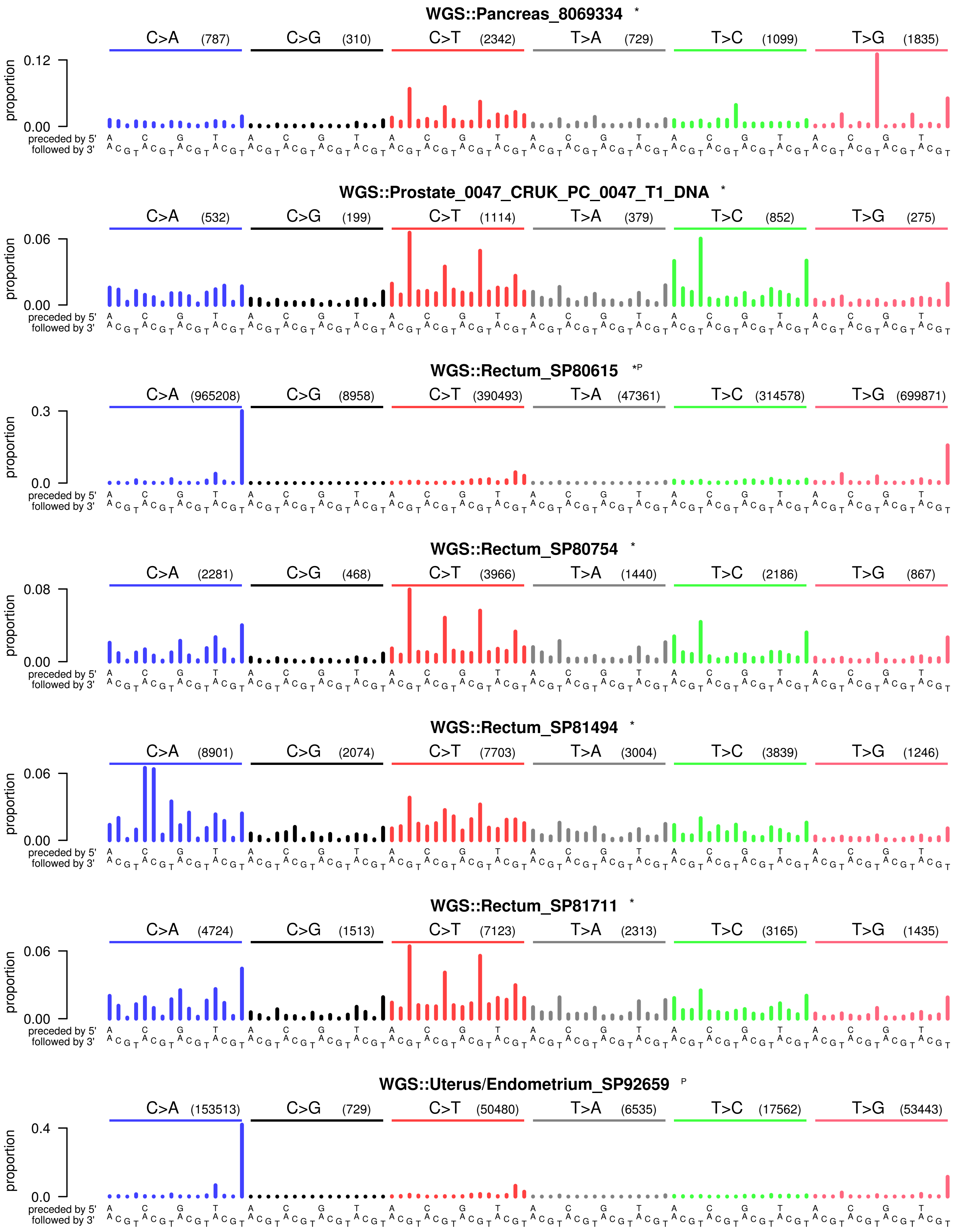
WGS::Pancreas_8069334 *
C>A (787)
C>G (310)
C>T (2342)
T>A (729)
T>C (1099)
T>G (1835)
proportion
0.12
0.00
preceded by 5'
followed by 3'
WGS::Prostate_0047_CRUK_PC_0047_T1_DNA *
C>A (532)
C>G (199)
C>T (1114)
T>A (379)
T>C (852)
T>G (275)
proportion
0.06
0.00
preceded by 5'
followed by 3'
WGS::Rectum_SP80615 *P
C>A (965208)
C>G (8958)
C>T (390493)
T>A (47361)
T>C (314578)
T>G (699871)
proportion
0.3
0.0
preceded by 5'
followed by 3'
WGS::Rectum_SP80754 *
C>A (2281)
C>G (468)
C>T (3966)
T>A (1440)
T>C (2186)
T>G (867)
proportion
0.08
0.00
preceded by 5'
followed by 3'
WGS::Rectum_SP81494 *
C>A (8901)
C>G (2074)
C>T (7703)
T>A (3004)
T>C (3839)
T>G (1246)
proportion
0.06
0.00
preceded by 5'
followed by 3'
WGS::Rectum_SP81711 *
C>A (4724)
C>G (1513)
C>T (7123)
T>A (2313)
T>C (3165)
T>G (1435)
proportion
0.06
0.00
preceded by 5'
followed by 3'
WGS::Uterus/Endometrium_SP92659 P
C>A (153513)
C>G (729)
C>T (50480)
T>A (6535)
T>C (17562)
T>G (53443)
proportion
0.4
0.0
preceded by 5'
followed by 3'

Supplemental_Fig_S10: **Indel spectra for SBS_$A^nT$ positive PCAWG tumours.** In addition to the $A^nT$ indel signature, these tumors also show clear evidence of signatures ID1 and ID2 (Alexandrov et al. 2019).

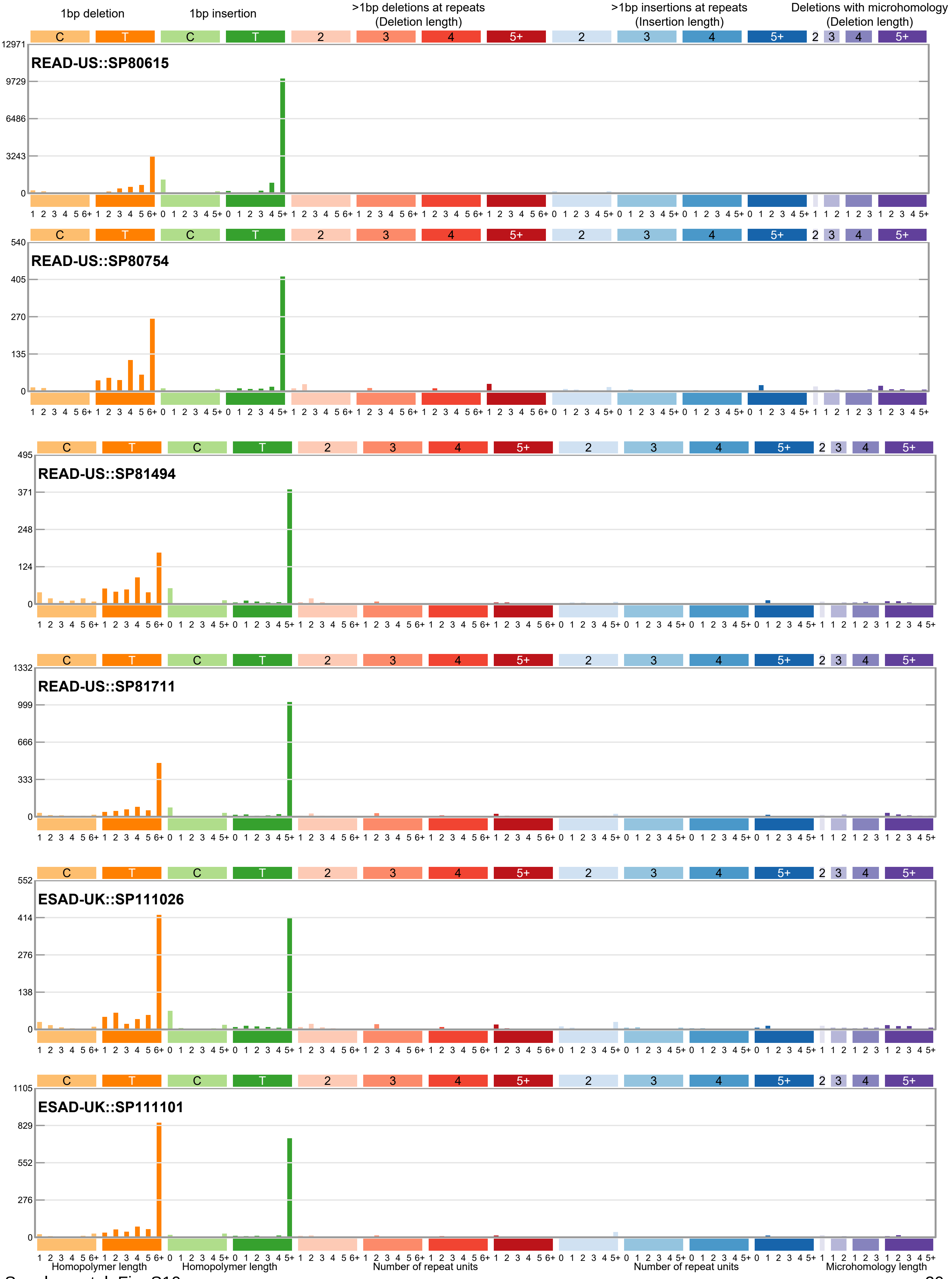

Supplemental_Fig_S11: **Sequence context of deletions of thymines by repeat length for SBS_A$^n$T positive PCAWG tumours.** All except for SP111026 show strong enrichment for adenines 5' of the repeat.

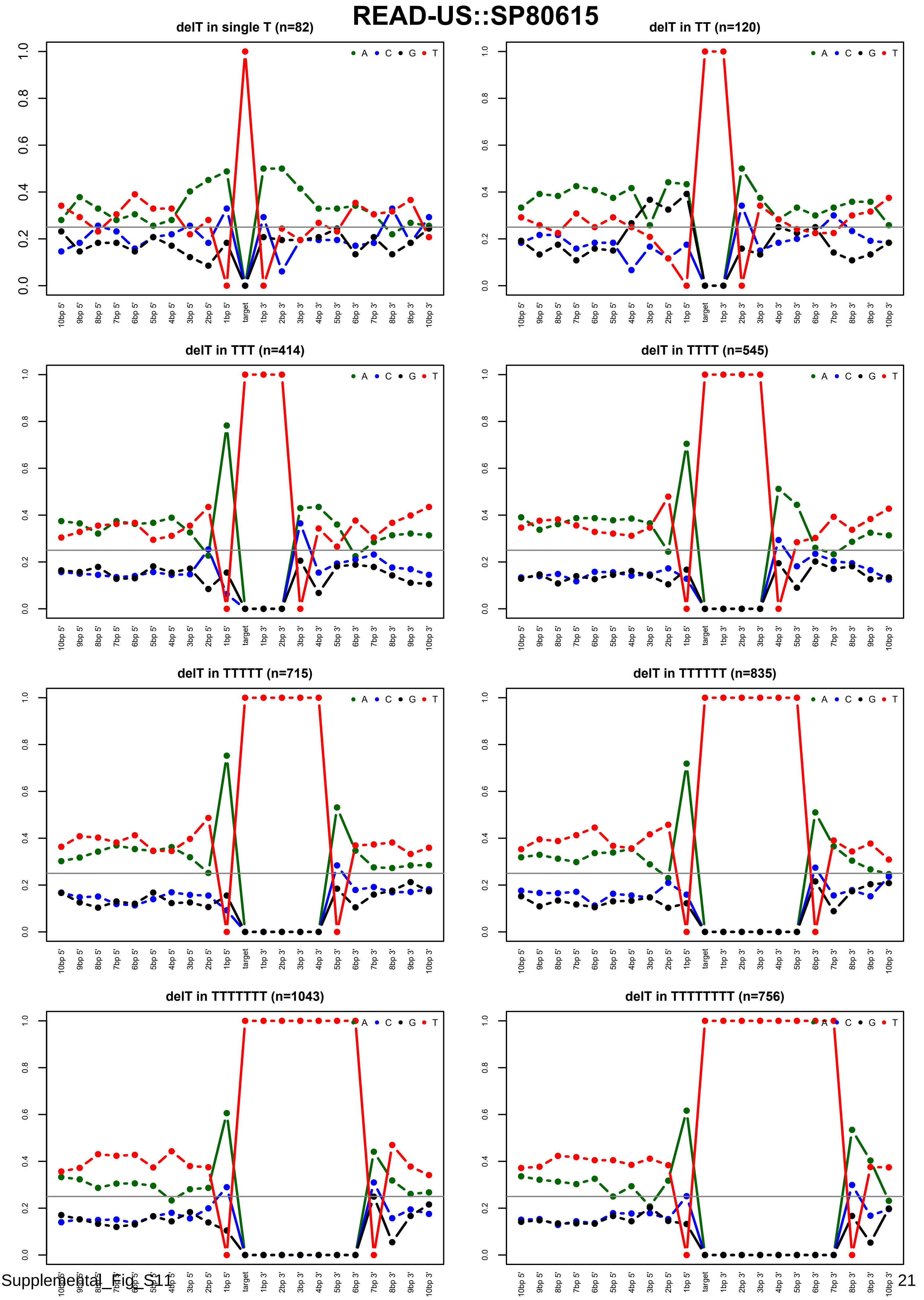

### READ-US::SP80754

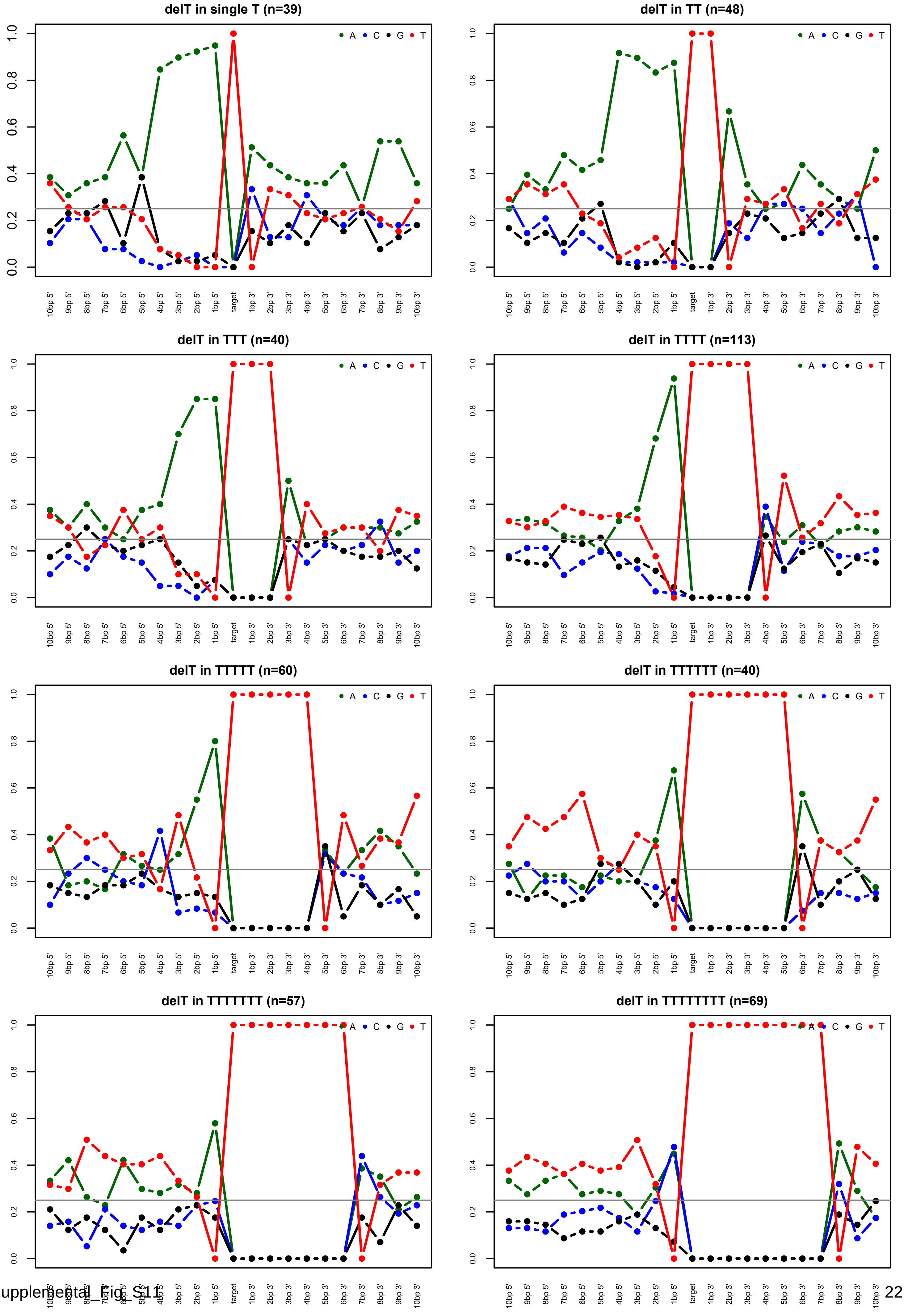

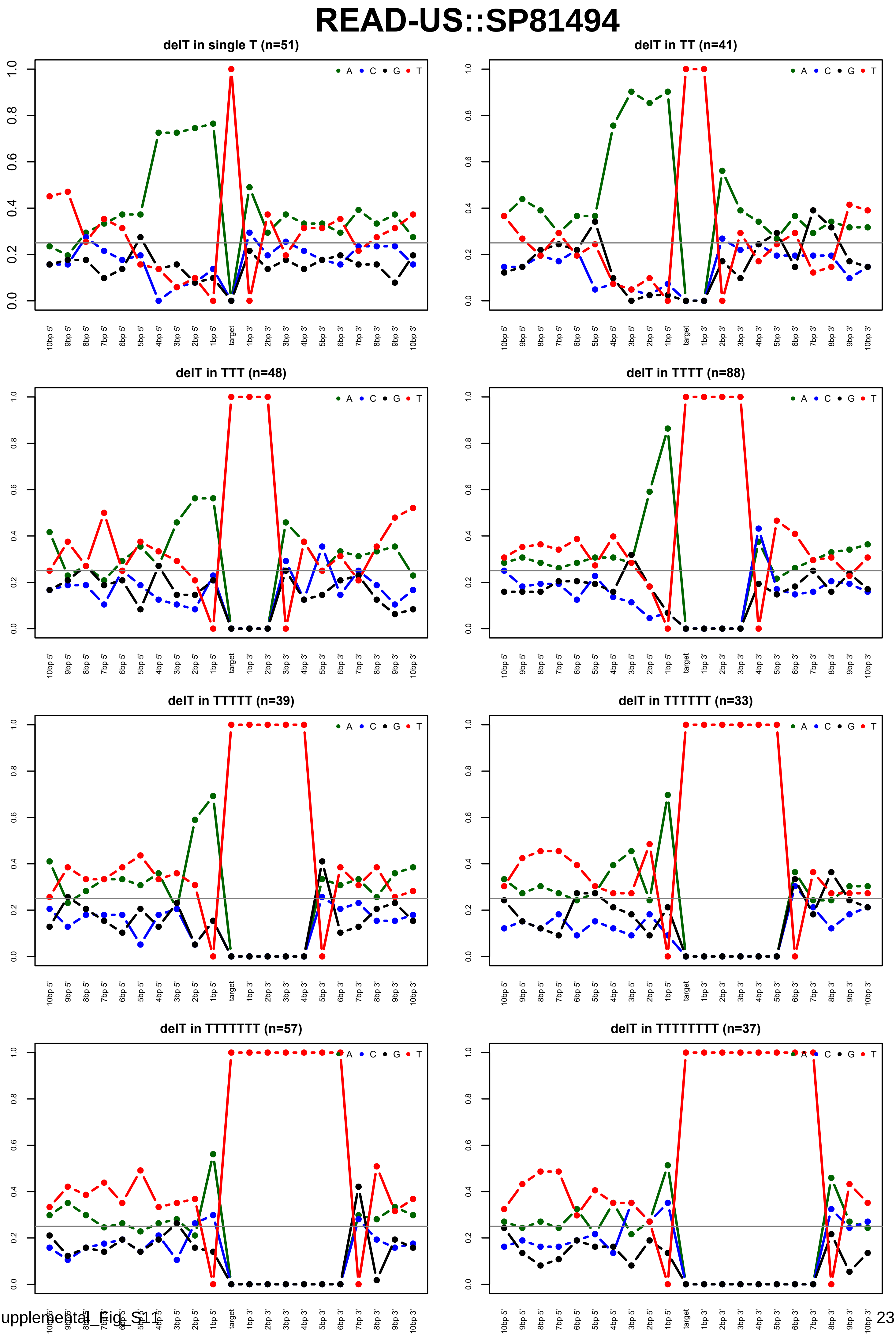
READ-US::SP81494
delT in single T (n=51)
delT in TT (n=41)
delT in TTT (n=48)
delT in TTTT (n=88)
delT in TTTTT (n=39)
delT in TTTTTT (n=33)
delT in TTTTTTT (n=57)
delT in TTTTTTTT (n=37)
A C G T
10bp 5' 9bp 5' 8bp 5' 7bp 5' 6bp 5' 5bp 5' 4bp 5' 3bp 5' 2bp 5' 1bp 5' target 1bp 3' 2bp 3' 3bp 3' 4bp 3' 5bp 3' 6bp 3' 7bp 3' 8bp 3' 9bp 3' 10bp 3'
0.0 0.2 0.4 0.6 0.8 1.0

### READ-US::SP81711

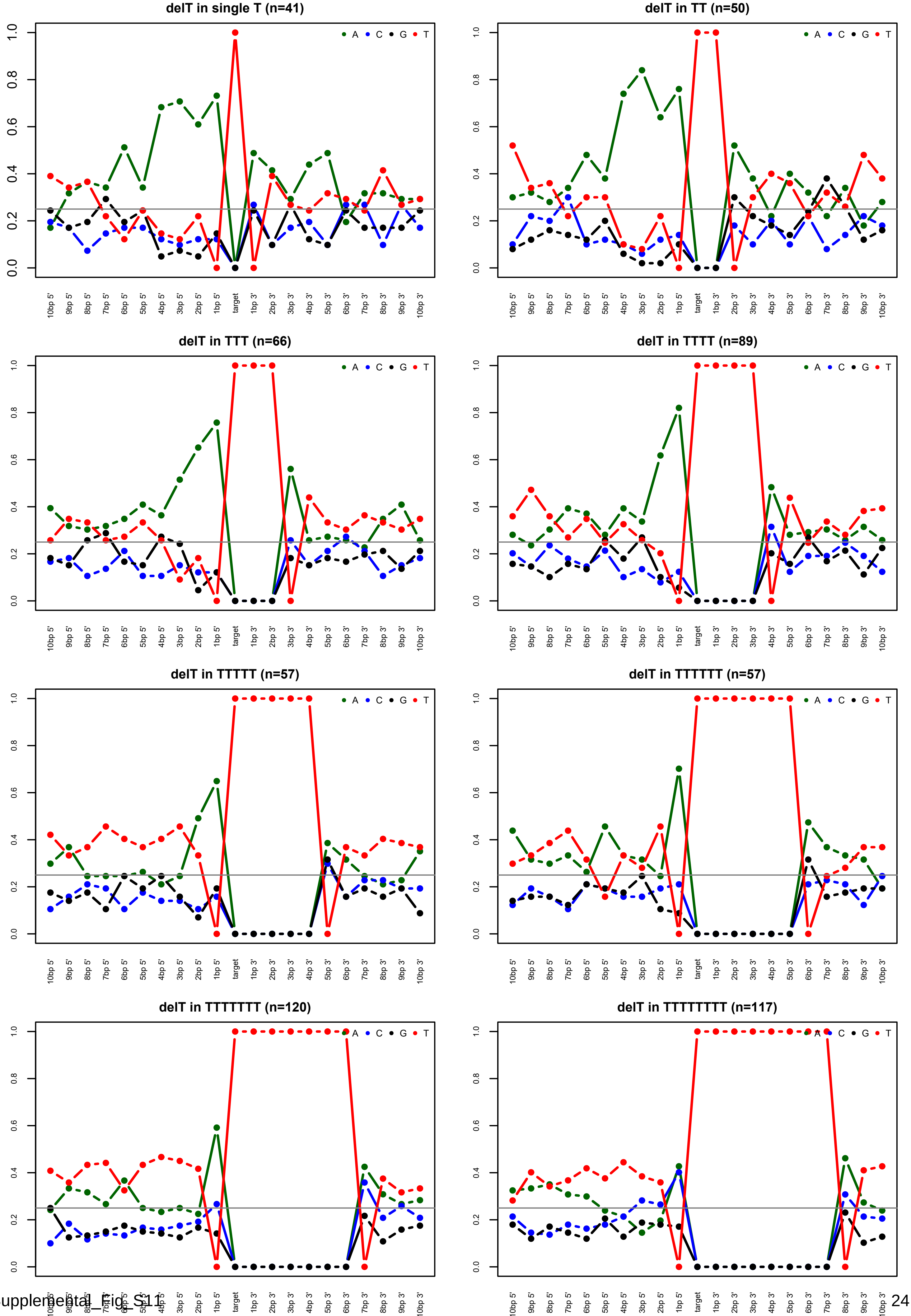

### ESAD-UK::SP111026

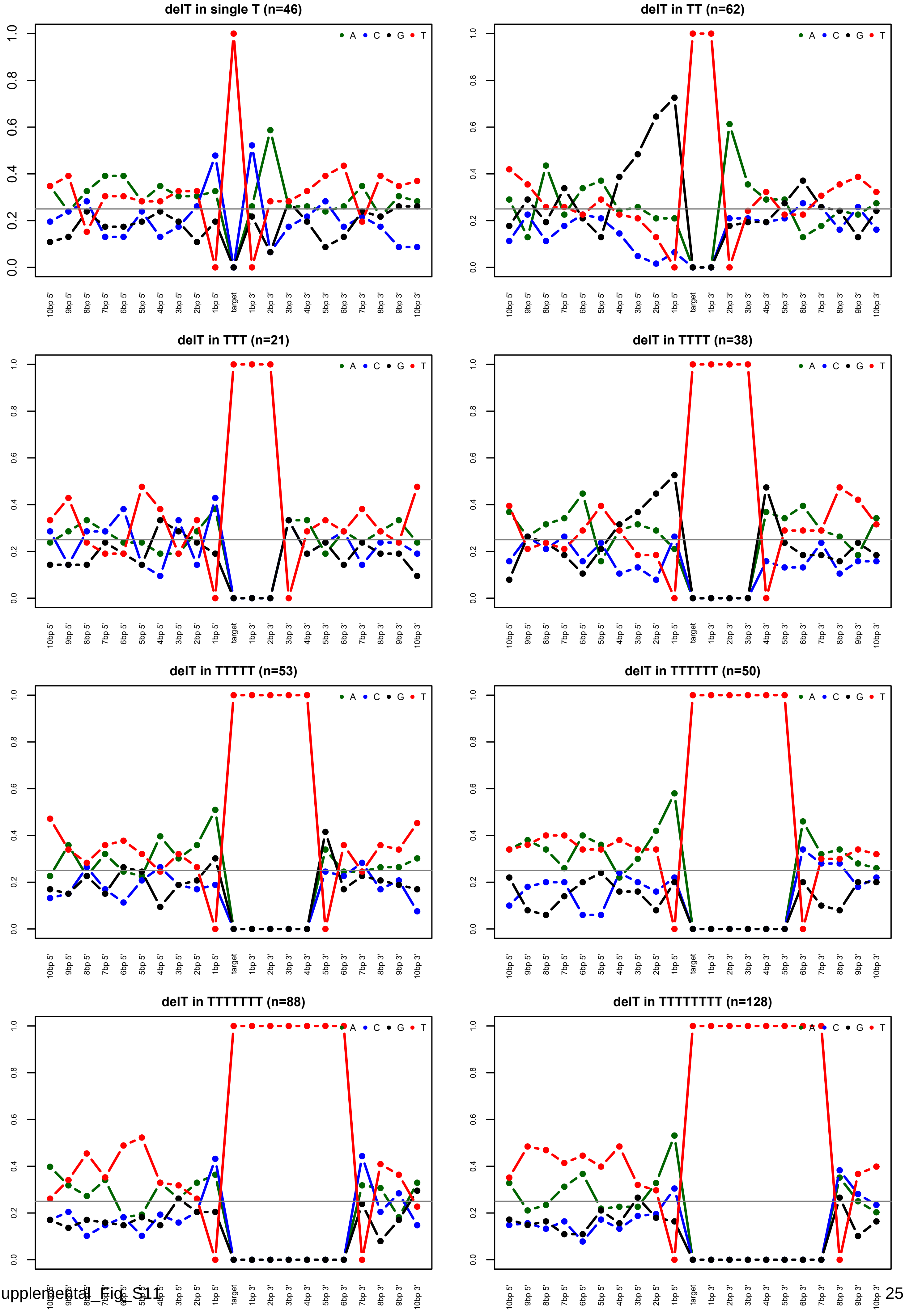

### ESAD-UK::SP111101

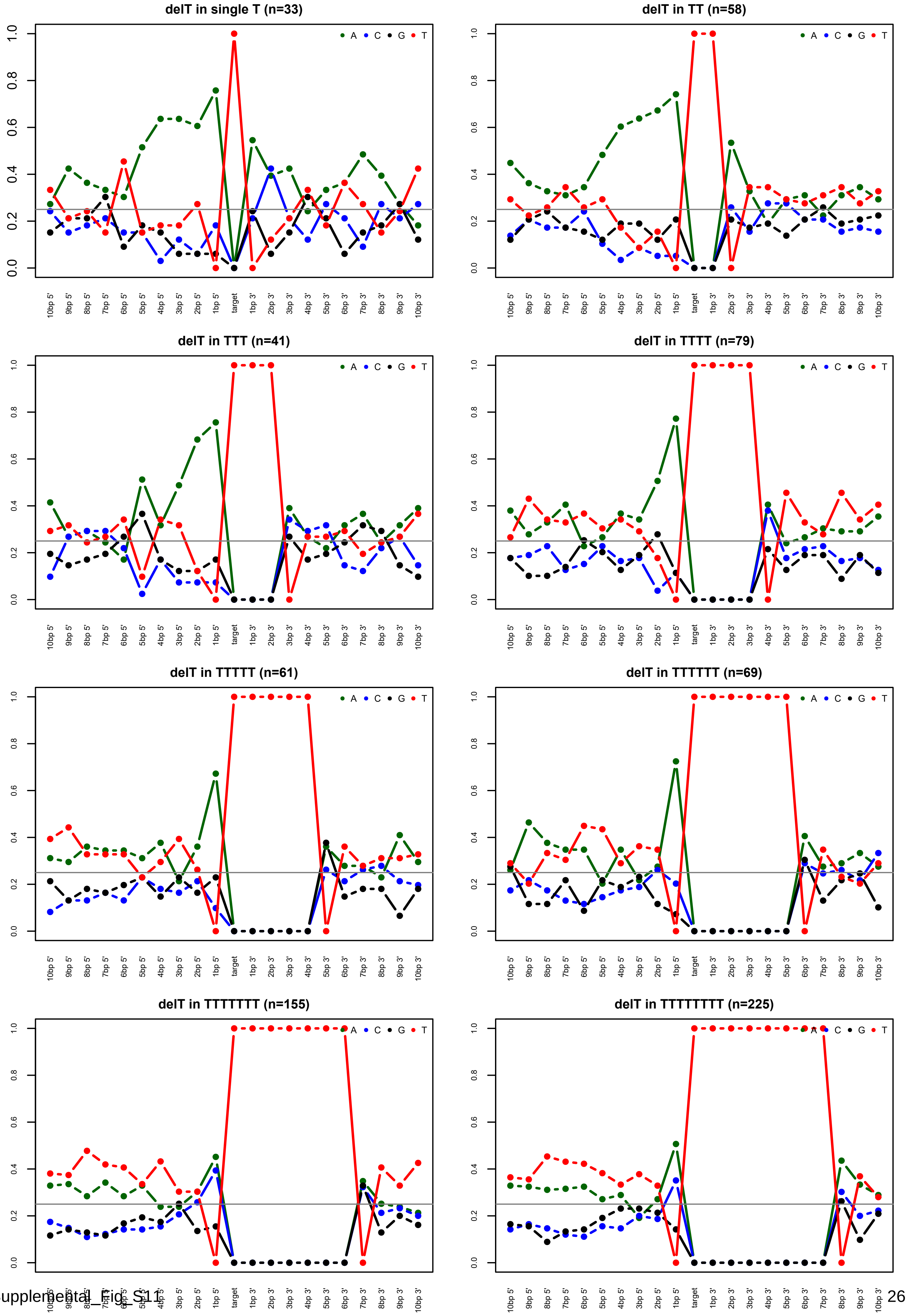

Supplemental_Fig_S12: **SNS mutational spectra of duocarmycin SA treated HepG2 clones.** Clones 1 and 2 were exposed to 100pM duocarmycin SA, clones 3 and 4 to 250pM.

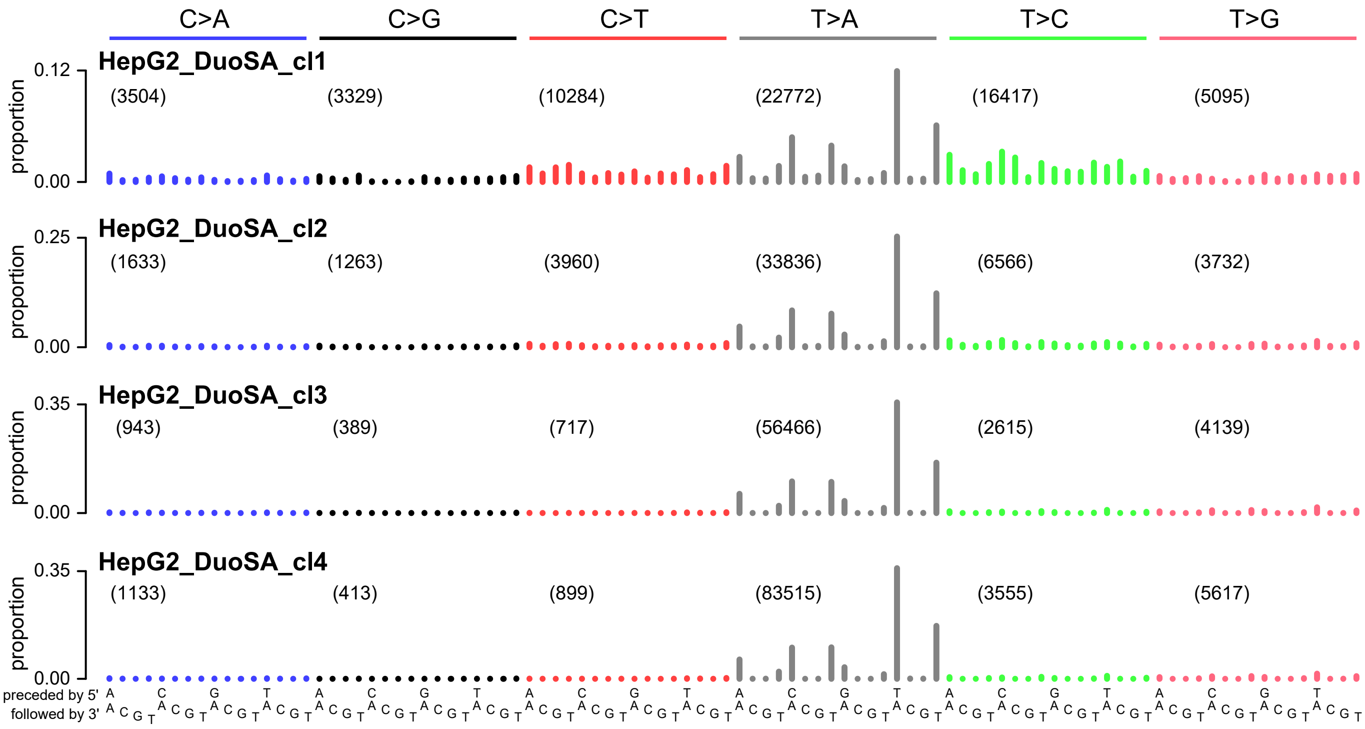

Supplemental_Fig_S13: **Indel mutational spectra of duocarmycin SA treated HepG2 clones.** Clones 1 and 2 were exposed to 100pM duocarmycin SA, clones 3 and 4 to 250pM.

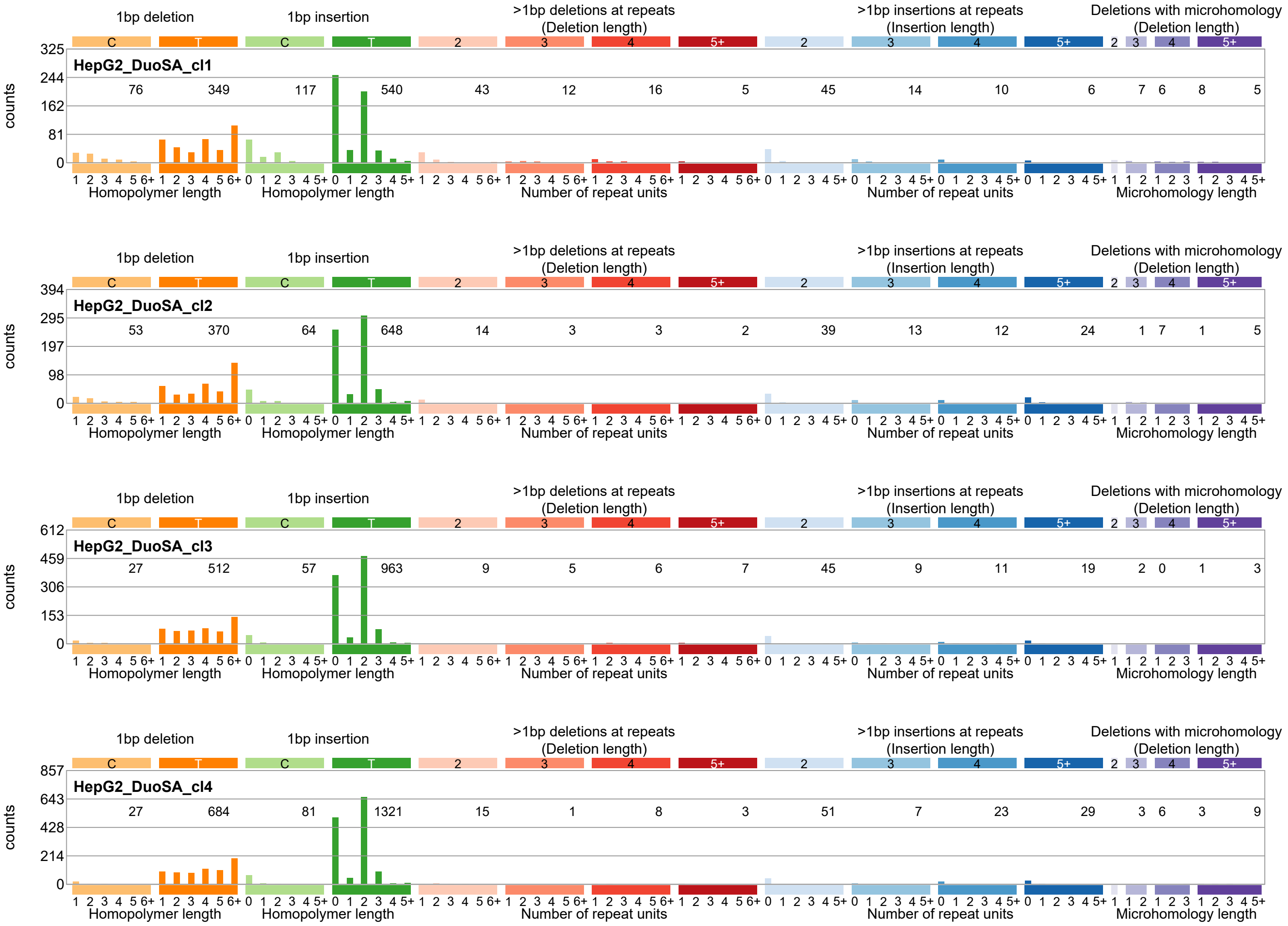

Supplemental_Fig_S14: **Double base substitution spectrum of Duocarmycin SA treated HepG2 clones.** Clones 1 and 2 were treated with 100pM duocarmycin SA, clones 3 and 4 were treated with 250pM duocarmycin SA.

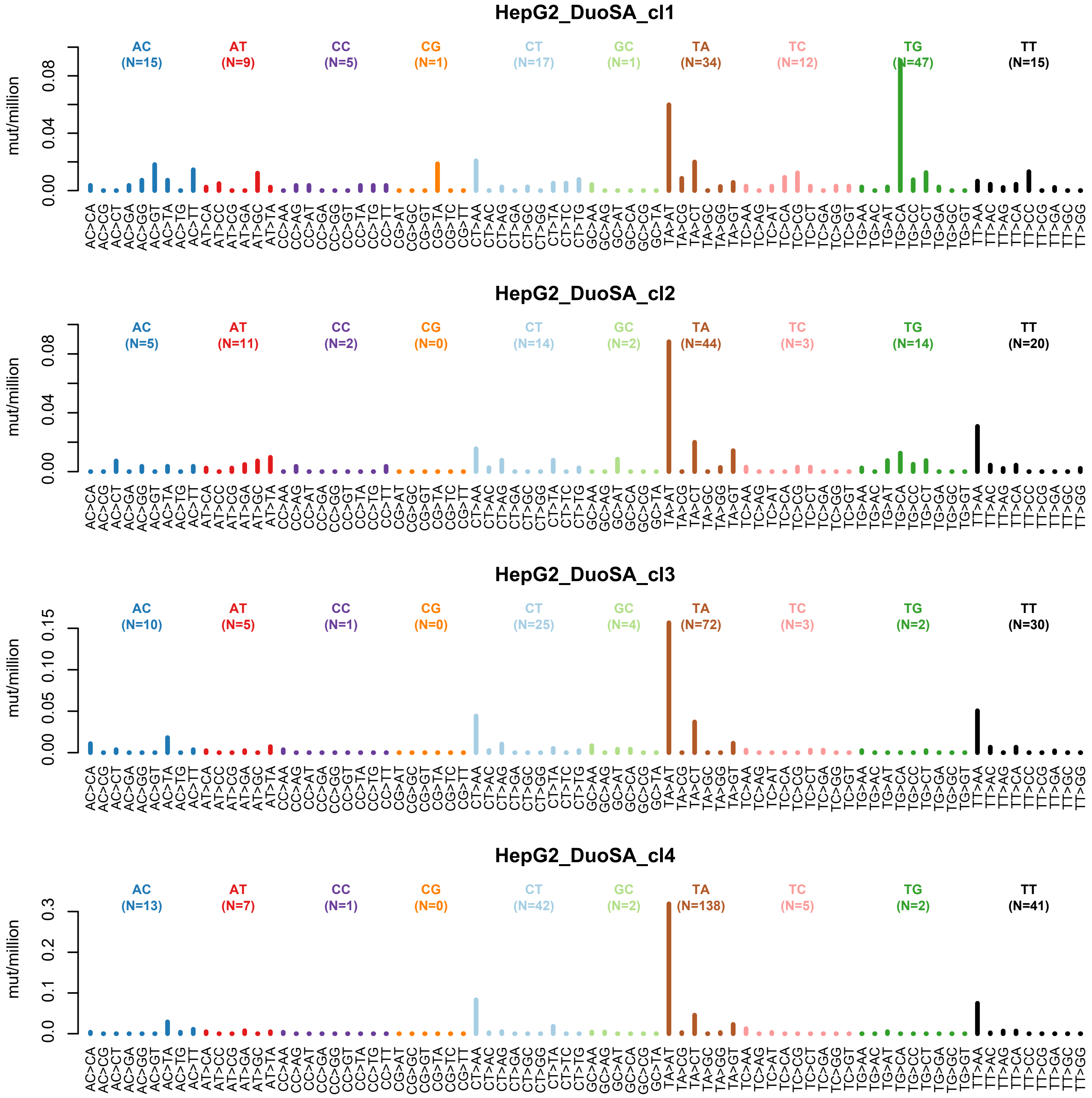

Supplemental_Fig_S15: **Characterization of the TC1 mutational spectrum.** (**A**) Whole-genome sequencing mutation spectrum of TC1. (**B**) Visualization of transcriptional strand bias in TC1 by plotting mutations on the transcribed (light) and untranscribed strand (dark) side by side. (**C**) Transcriptional strand bias as a function of gene expression for C>N (left) and T>N mutations (right). (**D**) Transcriptional strand bias as a function of distance to the transcription start site for C>N (left) and T>N mutations (right). (**E**) Extended sequence context specificity of TC1 mutations.

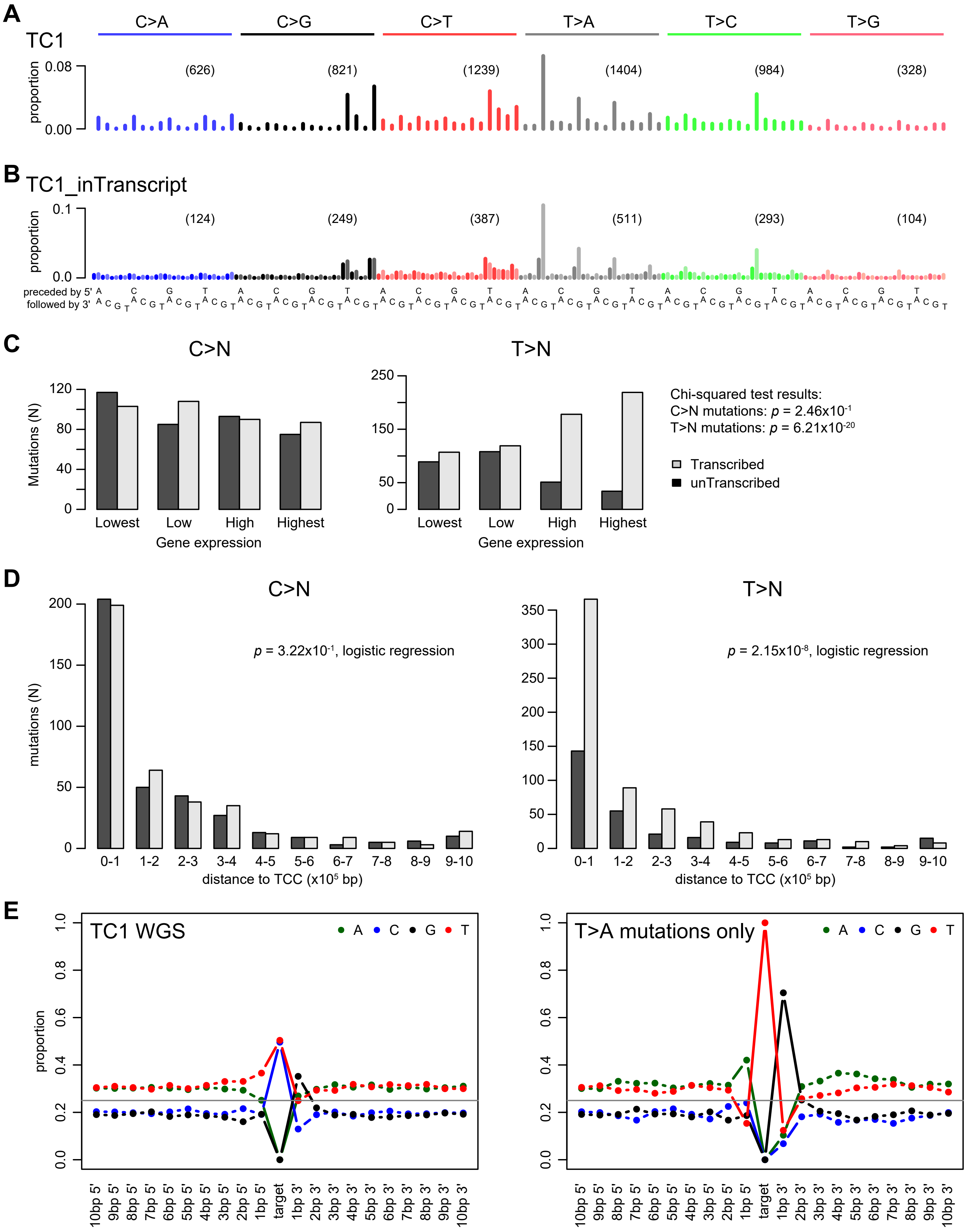

Supplemental_Fig_S16: **Identifying samples whose mutational spectrum resembles that of TC1.**

Background: Sample TC1 is not very highly mutated, and as a consequence we cannot extract a pure 'TC1 mutational signature' from the whole-genome sequencing data which could be used for mSigAct or sigProfiler analysis for presence of this signature in the publicly available sequencing data. Therefore we screened the publicly available sequencing data for tumors with presence of the same mutational processes by looking for tumors in which either the thymine mutations, or the T>A mutations specifically, bore high resemblance to that of TC1.
Below tables list the top 20 samples with the highest cosine similarity of T>N (left) or T>A (right) mutations compared to the TC1 spectrum from the 23,829 tumors investigated.
The mutation spectra of these tumors are found attached below.
Although we observe some similarity to the TC1 mutation spectrum in the samples below, the mutation counts are always very low. Therefore we cannot make solid conclusions on the presence of the same mutational process.

| Sample | cosine T>N | cosine T>A |
|---|---|---|
| Liver-HCC::TCGA-RC-A6M6-01A-11D-A32G-10 | 0.8133 | 0.8973 |
| Lung-AdenoCa::TCGA-44-8120-01A-11D-2238-08 | 0.8127 | 0.7769 |
| CNS-GBM::SK00102_P | 0.7937 | 0.9068 |
| Uterus-AdenoCa::TCGA-AX-A3G4-01A-11D-A20S-09 | 0.7932 | 0.9136 |
| Lung-SCC::TCGA-68-A59J-01A-21D-A26M-08 | 0.7882 | 0.7959 |
| Lung-AdenoCa::LUAD-NYU1051S | 0.7881 | 0.8168 |
| Lung-AdenoCa::TCGA-55-7570-01A-11D-2036-08 | 0.7817 | 0.8526 |
| Lung-AdenoCa::TCGA-49-4506-01A-01D-1265-08 | 0.7798 | 0.8500 |
| Lung-SCC::TCGA-98-A53H-01A-12D-A25L-08 | 0.7752 | 0.8631 |
| Lung-AdenoCa::TCGA-73-4658-01A-01D-1753-08 | 0.7723 | 0.8579 |
| Lung-SCC::TCGA-43-8118-01A-11D-2395-08 | 0.7700 | 0.7591 |
| Lung-SCC::TCGA-56-8307-01A-11D-2293-08 | 0.7699 | 0.7707 |
| Lung-AdenoCa::TCGA-49-AARQ-01A-11D-A410-08 | 0.7690 | 0.7660 |
| Lung-SCC::TCGA-39-5040-01A-21D-2122-08 | 0.7689 | 0.8090 |
| Lung-Small::S00050 | 0.7640 | 0.8592 |
| Lung-Small::585270 | 0.7630 | 0.8379 |
| Lung-AdenoCa::TCGA-MP-A4TK-01A-11D-A24P-08 | 0.7614 | 0.7600 |
| Lung-AdenoCa::TCGA-05-4249-01A-01D-1105-08 | 0.7604 | 0.8219 |
| Lung-AdenoCa::TCGA-NJ-A4YI-01A-11D-A25L-08 | 0.7594 | 0.8000 |
| Lung-SCC::TCGA-77-8144-01A-11D-2244-08 | 0.7579 | 0.8624 |

| Sample | cosine T>N | cosine T>A |
|---|---|---|
| Lung-SCC::TCGA-33-4547-01A-01D-1267-08 | 0.7455 | 0.9605 |
| Adrenal-neoplasm::TCGA-OR-A5J6-01A-31D-A29I-10 | 0.5917 | 0.9546 |
| Uterus-AdenoCa::TCGA-A5-A2K2-01A-11D-A18P-09 | 0.5914 | 0.9290 |
| Panc-AdenoCa::8014777 | 0.7065 | 0.9137 |
| Uterus-AdenoCa::TCGA-AX-A3G4-01A-11D-A20S-09 | 0.7932 | 0.9136 |
| Panc-AdenoCa::TCGA-IB-A5SS-01A-11D-A32N-08 | 0.6687 | 0.9108 |
| Prost-AdenoCa::TCGA-CH-5765-01A-11D-1576-08 | 0.5500 | 0.9108 |
| Liver-HCC::TCGA-DD-AADD-01A-11D-A40R-10 | 0.6629 | 0.9092 |
| CNS-GBM::SK00102_P | 0.7937 | 0.9068 |
| Transitional-cell-carcinoma::TCGA-DK-AA77-01A-11D-A | 0.7243 | 0.9042 |
| Sarcoma::TCGA-3B-A9HO-01A-11D-A387-09 | 0.5967 | 0.9004 |
| Kidney-Papillary::TCGA-G7-6796-01A-11D-1961-08 | 0.6828 | 0.8973 |
| Liver-HCC::TCGA-RC-A6M6-01A-11D-A32G-10 | 0.8133 | 0.8973 |
| Lymph-BNHL::TCGA-RQ-A68N-01A-11D-A31X-10 | 0.5947 | 0.8962 |
| Bladder-TCC::B106-Tumor | 0.4696 | 0.8946 |
| Head-SCC::HN_00378 | 0.5959 | 0.8916 |
| Sarcoma::TCGA-DX-A8BZ-01A-11D-A37C-09 | 0.6456 | 0.8910 |
| Kidney-RCC::TCGA-B0-5100-01A-01D-1421-08 | 0.6424 | 0.8889 |
| AML::CN-AML-CR-42-Dx | 0.5972 | 0.8865 |
| Transitional-cell-carcinoma::TCGA-E5-A4U1-01A-11D-A | 0.6997 | 0.8862 |

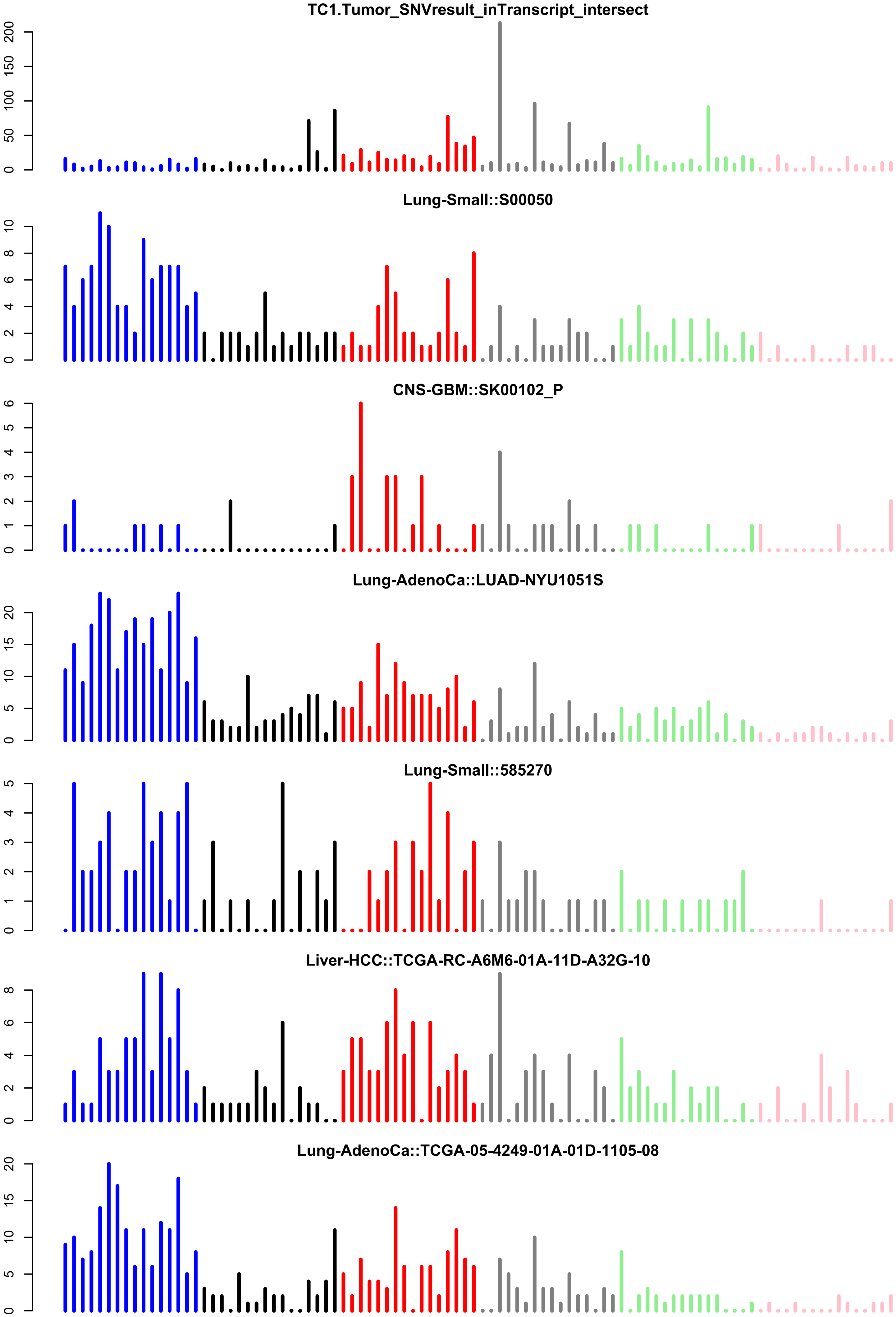
TC1.Tumor_SNVresult_inTranscript_intersect
Lung-Small::S00050
CNS-GBM::SK00102_P
Lung-AdenoCa::LUAD-NYU1051S
Lung-Small::585270
Liver-HCC::TCGA-RC-A6M6-01A-11D-A32G-10
Lung-AdenoCa::TCGA-05-4249-01A-01D-1105-08

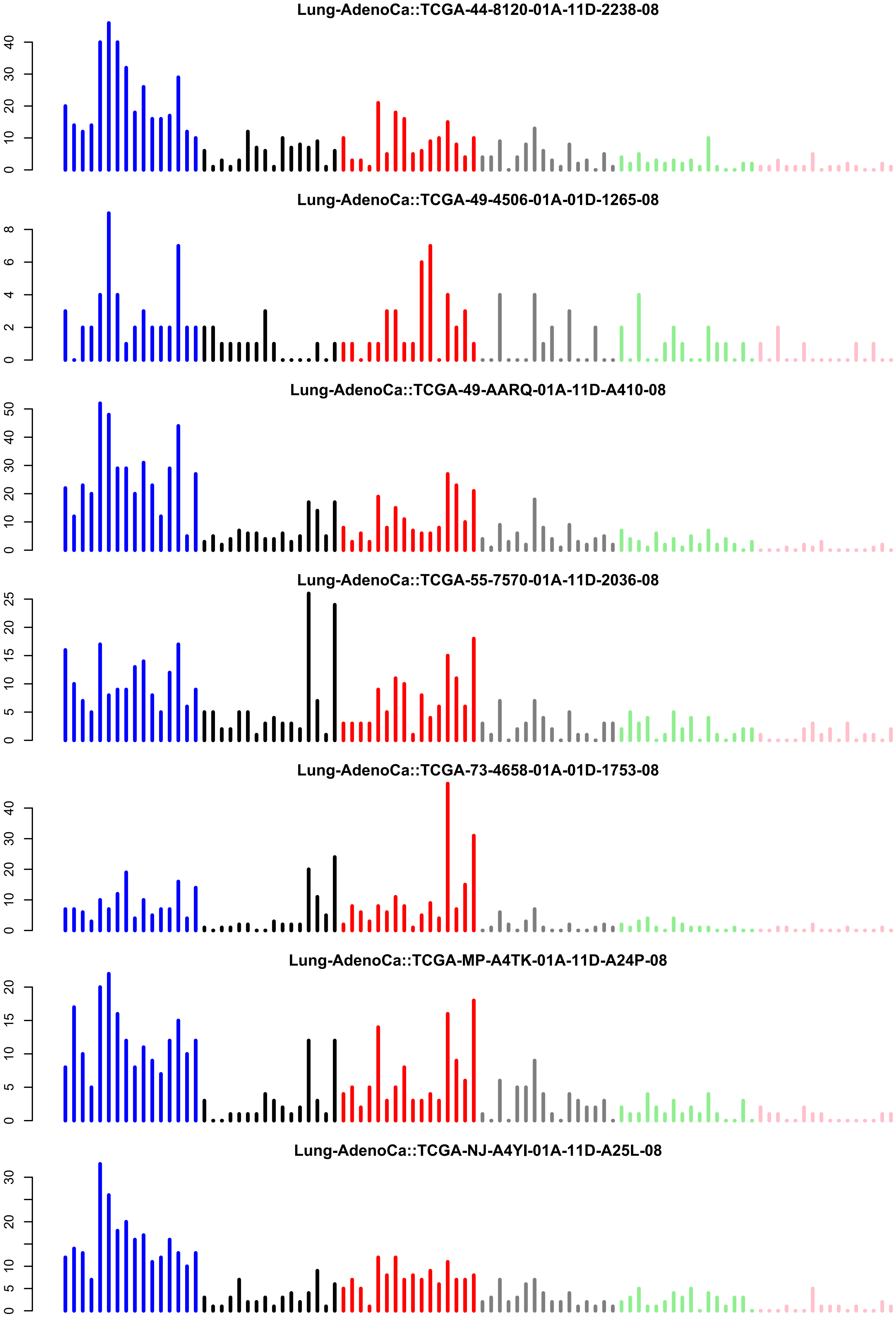
Lung-AdenoCa::TCGA-44-8120-01A-11D-2238-08
Lung-AdenoCa::TCGA-49-4506-01A-01D-1265-08
Lung-AdenoCa::TCGA-49-AARQ-01A-11D-A410-08
Lung-AdenoCa::TCGA-55-7570-01A-11D-2036-08
Lung-AdenoCa::TCGA-73-4658-01A-01D-1753-08
Lung-AdenoCa::TCGA-MP-A4TK-01A-11D-A24P-08
Lung-AdenoCa::TCGA-NJ-A4YI-01A-11D-A25L-08

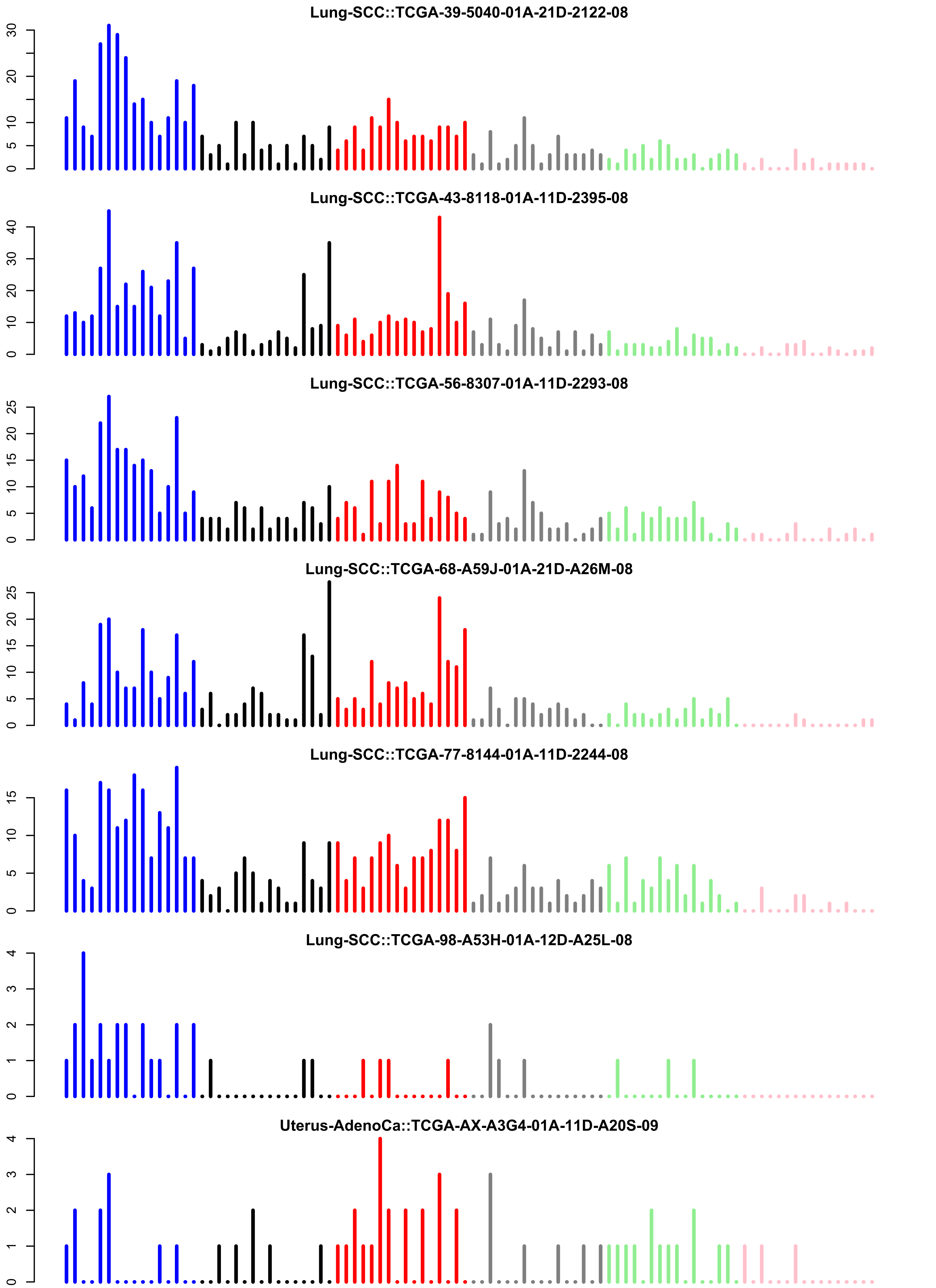
Lung-SCC::TCGA-39-5040-01A-21D-2122-08
Lung-SCC::TCGA-43-8118-01A-11D-2395-08
Lung-SCC::TCGA-56-8307-01A-11D-2293-08
Lung-SCC::TCGA-68-A59J-01A-21D-A26M-08
Lung-SCC::TCGA-77-8144-01A-11D-2244-08
Lung-SCC::TCGA-98-A53H-01A-12D-A25L-08
Uterus-AdenoCa::TCGA-AX-A3G4-01A-11D-A20S-09

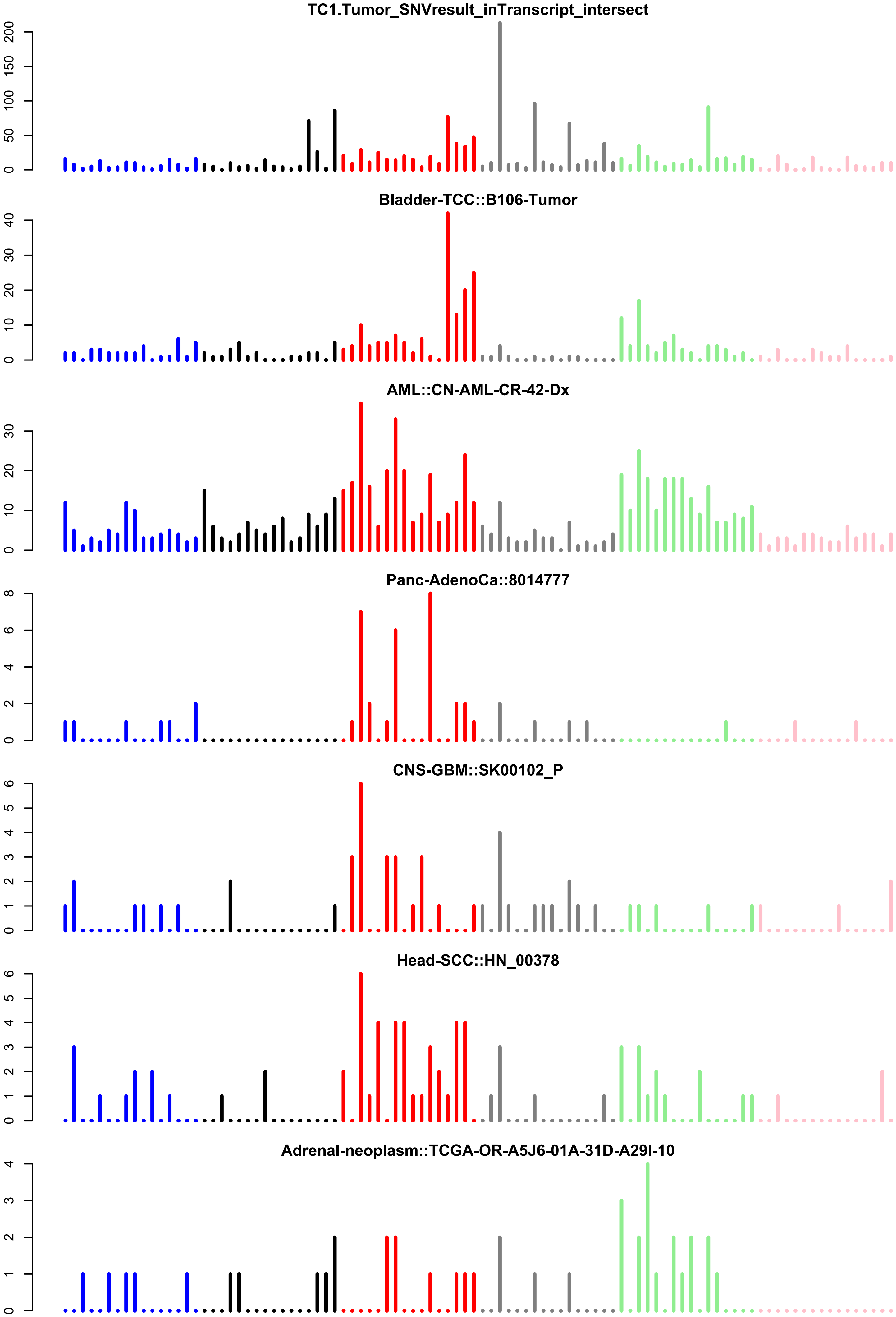
TC1.Tumor_SNVresult_inTranscript_intersect
0
50
100
150
200
Bladder-TCC::B106-Tumor
0
10
20
30
40
AML::CN-AML-CR-42-Dx
0
10
20
30
Panc-AdenoCa::8014777
0
2
4
6
8
CNS-GBM::SK00102_P
0
1
2
3
4
5
6
Head-SCC::HN_00378
0
1
2
3
4
5
6
Adrenal-neoplasm::TCGA-OR-A5J6-01A-31D-A29I-10
0
1
2
3
4

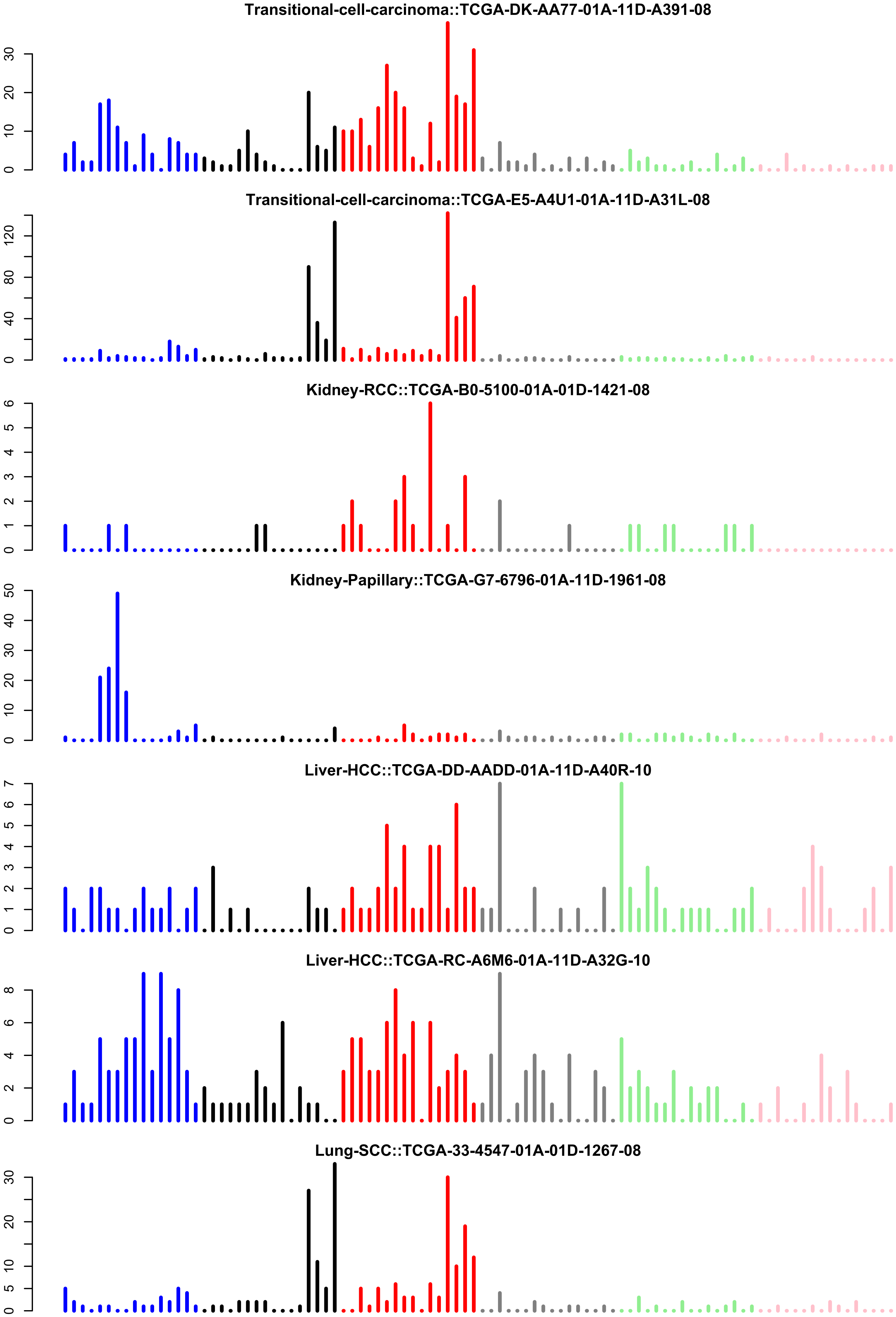
Transitional-cell-carcinoma::TCGA-DK-AA77-01A-11D-A391-08
0
10
20
30
Transitional-cell-carcinoma::TCGA-E5-A4U1-01A-11D-A31L-08
0
40
80
120
Kidney-RCC::TCGA-B0-5100-01A-01D-1421-08
0
1
2
3
4
5
6
Kidney-Papillary::TCGA-G7-6796-01A-11D-1961-08
0
10
20
30
40
50
Liver-HCC::TCGA-DD-AADD-01A-11D-A40R-10
0
1
2
3
4
5
6
7
Liver-HCC::TCGA-RC-A6M6-01A-11D-A32G-10
0
2
4
6
8
Lung-SCC::TCGA-33-4547-01A-01D-1267-08
0
5
10
20
30

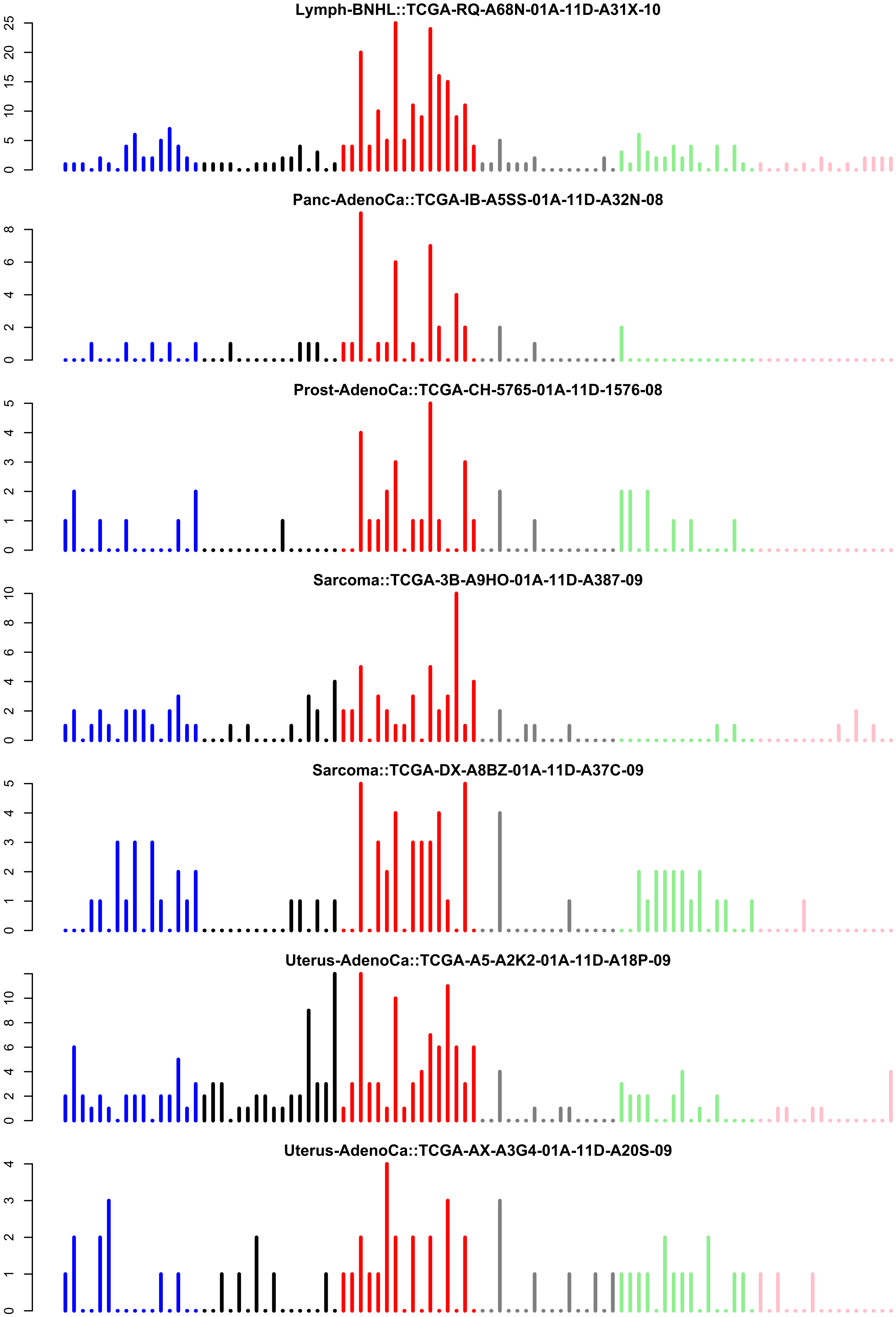
Lymph-BNHL::TCGA-RQ-A68N-01A-11D-A31X-10
Panc-AdenoCa::TCGA-IB-A5SS-01A-11D-A32N-08
Prost-AdenoCa::TCGA-CH-5765-01A-11D-1576-08
Sarcoma::TCGA-3B-A9HO-01A-11D-A387-09
Sarcoma::TCGA-DX-A8BZ-01A-11D-A37C-09
Uterus-AdenoCa::TCGA-A5-A2K2-01A-11D-A18P-09
Uterus-AdenoCa::TCGA-AX-A3G4-01A-11D-A20S-09

Supplemental_Fig_S17: **Alignment of non-human reads from WGS data of patients 62074759 and TC1 to bacterial genomes**

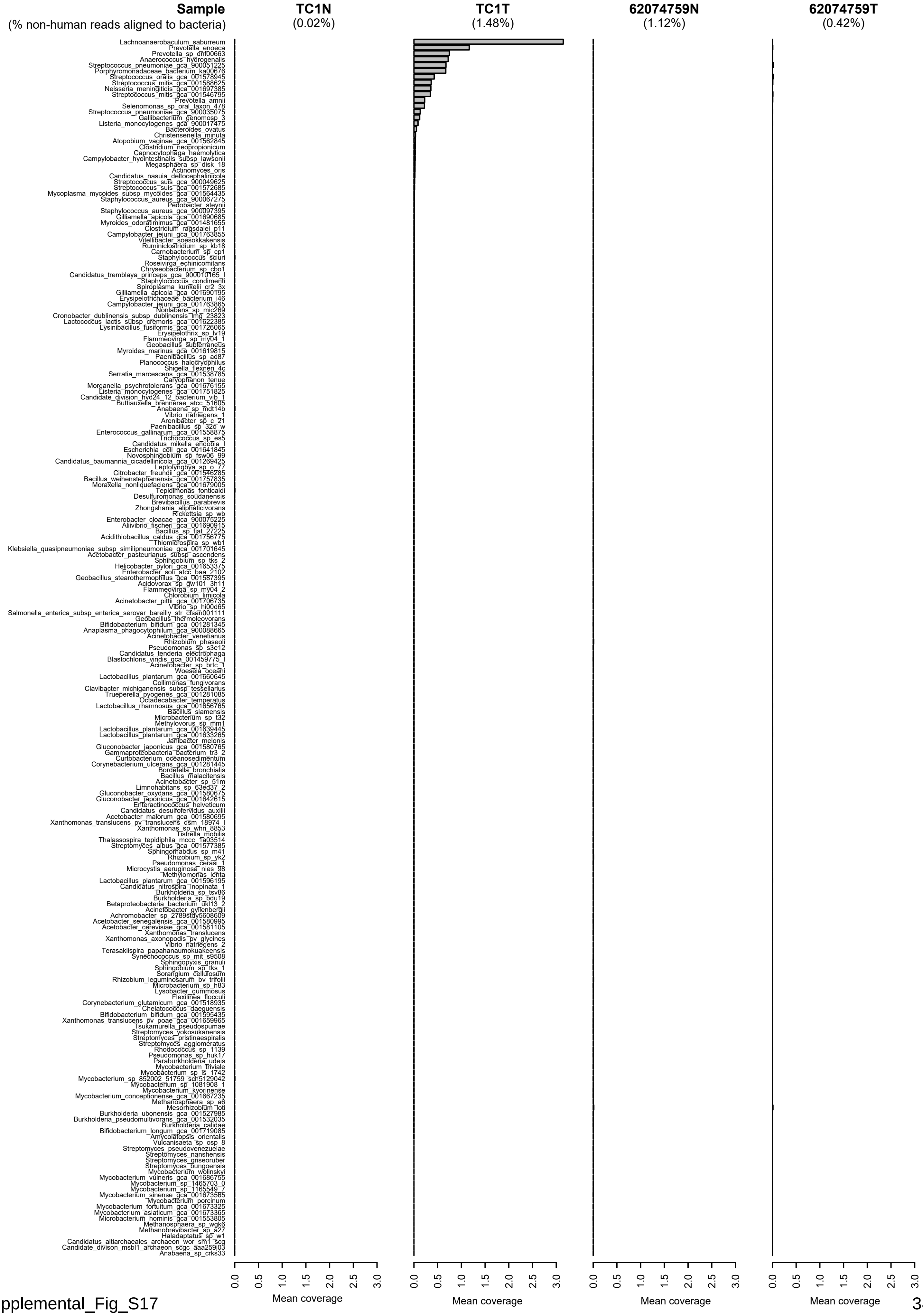
