## Supplemental Data S1 for "Identification of novel mutational signatures in Asian oral squamous cell carcinomas associated with bacterial infections": README.txt

Description of the information in "SBS86 samplesSelected.txt"

For each sample in the PCAWG mutational signature analysis, this table contains counts of mutations with specific characteristics that are associated with SBS86.

T2Ncount:		Sum of the number of T>A, T>C and T>G single nucleotide substitutions
A3bpUpstr:		Total number of T>A, T>C and T>G mutations where there is an adenine 3bp 5' of the mutated T
AAAT2AAAN:		Total number of T>A, T>C and T>G mutations where there are 3 consecutive adenines directly 5' of the mutated T
ATT2ACTcount:		Number of ATT>ACT mutations (this is the main peak of mutational signature SBS86)
A3bpUpstrATT2ACT:	Number of ATT>ACT mutations where there is an adenine 3bp 5' of the mutated T (i.e. ANATT>ANACT)
AAATT2AAACT:		Number of AAATT>AAACT mutations (this is equal to AAAT2AAAN, but then only for ATT>ACT mutations)
