## Supplemental Data S1 for "Identification of novel mutational signatures in Asian oral squamous cell carcinomas associated with bacterial infections": mSigAct.v0.10.R

```
#
# mSigAct.R
#
# v 0.9
#
# 2017 08 11
#
# Copyright 2017 by Alvin Wei Tian Ng, Steven G. Rozen
#
# The code is released under GPL-3
# https://www.gnu.org/licenses/gpl-3.0.en.html
#
# This file contains functions for sparse maximum likelihood assignment of
# mutational signature activities and for testing whether a given signature is
# needed to explain a spectrum

# Check R version and dependencies 
if (!(R.version$major >= "3")) stop('mSigAct only works with R 3.2 or newer')

if (! (R.version$minor >= '3.2')) stop('mSigAct only works with R 3.2 or newer') 

# Dependencies
library(nloptr)
library(sets)
library(parallel)

# Helper function, given signatures (sigs) and exposures (exp), return a
# *proportional* reconstruction; in general, it is *not necessarily* scaled to
# the actual spectrum counts.
prop.reconstruct <- function(sigs, exp) {
  stopifnot(length(exp) == ncol(sigs))
  as.matrix(sigs) %*% exp
}

# Define the objective function for use with nloptr (non-linear optimization)
# Negative binomial maximum likelihood objective function
#
# exp  is the matrix of exposures ("activities")
# sigs is the matrix of signatures
# spectrum is the spectrum to assess
# nbinom.size is the dispersion parameter for the negative
#             binomial distribution; smaller is more dispersed
#
# The result is
# -1 * the log(likelihood(spectrum | reconstruction))
# (nloptr minimizes the objective function.)
#
# The lower the objective function, the better
obj.fun.nbinom.maxlh <-function(exp, spectrum, sigs,
                                nbinom.size) {

  if (any(is.na(exp))) return(Inf)

  reconstruction <-  prop.reconstruct(sigs = sigs, exp = exp)

  ## catch errors with NA in the input or in reconstruction.
  if (any(is.na(reconstruction))) {
    save(reconstruction, spectrum, sigs, exp, nbinom.size,
         file='reconstruction.error.R')
  }
  stopifnot(!any(is.na(reconstruction)))

  loglh <- 0
  for (i in 1:length(spectrum)) { # Iterate over each channel in the
                                  # spectrum and sum the log
                                  # likelihoods.

    nbinom <- dnbinom(x=spectrum[i],
                      mu = reconstruction[i],
                      size=nbinom.size,
                      log=T)

    loglh <-loglh + nbinom
  }
  stopifnot(mode(loglh) == 'numeric' )
  -loglh
}

# Use nloptr (numerical non-linear optimization) to find an assignmeent of
# signature activites for one tumor. The nlpotr algorithm and the objective
# function are arguments.
nloptr.one.tumor <- function(spectrum, sigs,
                             algorithm='NLOPT_LN_COBYLA',
                             maxeval=1000, print_level=0,
                             xtol_rel=0.001, # 0.0001,
                             obj.fun,
                             ... # additional arguments for obj.fun
                             ) {
  if (class(sigs) == 'numeric') {
    # In this case we got a numeric vector, not a matrix.
    # We assume the caller intended it as a single
    # column matrix.
    sigs <- matrix(sigs, ncol=1)
  }

  # The if statement is an example of captuing state with
  # a call to save, which seems useful for debugging
  # in a call to mclapply (multi-core lapply).
  if (nrow(sigs)!= length(spectrum)) {
    save(spectrum, sigs, file='spectrum.sigs.debug.R')
  }
  stopifnot(nrow(sigs) == length(spectrum))

  # x0 is uniform distribution of mutations across signatures
  # (Each signature gets the same number of mutations)
  x0 <- rep(sum(spectrum) / ncol(sigs), ncol(sigs))

  res <- nloptr(x0=x0,
                eval_f=obj.fun,
                lb=rep(0, ncol(sigs)),
                opts=list(algorithm=algorithm,
                          xtol_rel=xtol_rel,
                          print_level=print_level,
                          maxeval=maxeval),
                spectrum=spectrum,
                sigs=sigs,
                ...)
  names(res$solution) <- colnames(sigs)
  res
}

# Returns a list with elements:
#
# loglh    - the log likelihood of the best solution (set of exposures) found
# exposure - the vector of exposures that generate loglh, in this case
# 'exposure' means the number of mutations ascribed to each signature
#
one.lh.and.exp <- function(spect, sigs, trace,
                           algorithm='NLOPT_LN_COBYLA',
                           obj.fun,
                           nbinom.size) {
  r <- nloptr.one.tumor(spect, sigs, maxeval=1e6,
                        xtol_rel = 1e-7,
                        algorithm=algorithm, obj.fun = obj.fun,
                        nbinom.size=nbinom.size)
  if (trace >  0) cat(r$objective, r$iterations, sum(r$solution), '\n')

  loglh <- r$objective

  if (loglh == Inf && trace > 0) cat("Got -Inf in one.lh.and.exp\n")

  # sum(recon) is likely to be close to, but not equal to the number
  # of mutations in the spectrum, so we scale exposures to get the
  # number of mutations in the spectrum
  exp <- r$solution * (sum(spect) / sum(r$solution)) # sum(recon))

  list(loglh=-loglh, exposure=exp)
}

## Assign activities and test whether signature activities can be removed

## Breadth-first first search down to fixed number of levels

# Helper function: is the set 'probe' a superset of any set in 'background'?
is.superset.of.any <- function(probe, background) {
  for (b in background) {
    if (set_is_proper_subset(b, probe)) return(T)
  }
  return(F)
}

sparse.assign.activity <- function(spect, sigs,
                                   max.level=5,
                                   p.thresh=0.05, trace=0,
                                   obj.fun,
                                   nbinom.size) {
  mode(spect) <-  'numeric'
  start <- one.lh.and.exp(spect, sigs, trace=0,
                          obj.fun=obj.fun,
                          nbinom.size=nbinom.size)
  lh.w.all <- start$loglh
  start.exp <- start$exposure
  non.0.exp.index <- which(start.exp > 0.5)
  if (trace > 0) {
    cat('Starting with',
        paste(names(start.exp)[non.0.exp.index], collapse=','),
        '\n')
    cat('max.level =', max.level, '\n')
  }
  if (length(non.0.exp.index) < 2) {
    if (trace > 0) cat('returning, only', length(non.0.exp.index),
                       'non-0 exposures\n')
    return(start.exp)
  }
  max.level <- min(max.level, length(non.0.exp.index) - 1)

  c.r.s <- set() # subsets of the signatures indices that cannot be removed

  max.p <- numeric(max.level)
  best.subset <- non.0.exp.index
  best.try <- start
  best.try.is.start <- T

  for (df in 1:max.level) {
    if (trace > 0) cat('df =', df, '\n')
    subsets <- set_combn(non.0.exp.index, df)
    for (subset in subsets) {
      # subset is of class set (package sets)
      if (is.superset.of.any(subset, c.r.s)) next
      subset.to.remove.v <- as.numeric(subset)
      subset.name <- paste(names(start.exp)[subset.to.remove.v], collapse=',')
      tmp.set <- setdiff(non.0.exp.index, subset.to.remove.v)
      try.sigs <- sigs[ , tmp.set]

      if (length(tmp.set) == 1) {
        try.sigs <- as.matrix(try.sigs)
        rownames(try.sigs) <- rownames(sigs)
        if (trace > 0) cat("New code\n")
      }

      # Get the max lh for try.sig
      try <- one.lh.and.exp(spect, try.sigs, trace=0,
                            obj.fun=obj.fun,
                            nbinom.size=nbinom.size)
      # try contains maxlh and exposure
      statistic <- 2 * (lh.w.all - try$loglh)
      chisq.p <- pchisq(statistic, df, lower.tail=F)
      if (trace > 0) {
        cat('Trying', subset.name, 'p =', chisq.p, '; ')
        if (trace > 1) cat('loglh =', try$loglh, '; statistic  =', statistic, ';')
      }
      if (chisq.p > p.thresh) {
        # This an acceptable set of exposures
        if (trace > 0) cat (' acceptable;')
        if (chisq.p > max.p[df]) {
          # This is the best so far
          max.p[df] <- chisq.p
          best.subset <- tmp.set
          best.try <- try
          best.try.is.start <- F
          if (trace > 0) cat('best\n')
        } else {
          if (trace > 0) cat('not best\n')
        }
      } else {
        c.r.s <- set_union(c.r.s, set(subset))
        if (trace > 0)  {
          cat('cannot remove\n')
        }
      }
    } # end for (subset in subsets)
    if (max.p[df] == 0) break
  } # end for df in 1:max.level

  # Need to return the exposures in the context of the orginal signatures matrix
  out.exp <- numeric(ncol(sigs)) #all zeros
  names(out.exp) <- colnames(sigs)
  if (best.try.is.start) {
    out.exp <- start.exp
  } else {
    out.exp[best.subset] <- best.try$exposure
  }
  if (trace > 0) {
    cat('max.p =', paste(max.p, collapse = ', '), '\n')
    cat('Ending with',
        paste(names(start.exp)[best.subset], collapse=','),
        '\n')
  }
  stopifnot(abs(sum(out.exp) - sum(spect)) < 1)
  out.exp
}

## Likelihood ratio test

is.present.p.m.likelihood <- function(spect,
                                 sigs,
                                 sig.to.test,
                                 trace=0,
                                 algorithm='NLOPT_LN_COBYLA',
                                 obj.fun,
                                 nbinom.size) {

  loglh.with <- one.lh.and.exp(spect, sigs, trace=trace,
                               algorithm=algorithm, obj.fun=obj.fun,
                               nbinom.size=nbinom.size)$loglh
  loglh.without <- one.lh.and.exp(spect, sigs[ ,-sig.to.test], trace=trace,
                                  algorithm=algorithm,
                                  obj.fun=obj.fun,
                                  nbinom.size=nbinom.size)$loglh
  statistic <- 2 * (loglh.with - loglh.without)
  chisq.p <- pchisq(statistic, 1, lower.tail=F)
  if (trace > 0) cat('statistic  =', statistic, '\nchisq p =', chisq.p, '\n')

  list(with=loglh.with,
       without=loglh.without,
       statistic=statistic,
       chisq.p=chisq.p)
}

signature.presence.test <-
    function(spect, sigs, target.sig.index,
                          trace=0,
                          obj.fun,
                          nbinom.size) {
  is.present.p.m.likelihood(spect,
                       sigs, target.sig.index,
                       trace=trace, obj.fun=obj.fun,
                       nbinom.size=nbinom.size)$chisq.p
}

## Cacluate exposures for entire set of tumors

compute.all.neg.log.lh <- function(spect, sigs, exp,
                                   obj.fun,
                                   nbinom.size) {
  out <- numeric(ncol(spect))
  names(out) <- colnames(spect)
  for (i in 1:ncol(spect)) {
    recon <- prop.reconstruct(sigs, exp[ , i])
    out[i] <- obj.fun(exp[ ,1], spect[ ,i], sigs,
                      nbinom.size=nbinom.size)
  }
  out
}

sanity.check.ex <- function(spectrum, sigs, exposure) {
  ex.sums <- margin.table(exposure, 2)
  all.reconstruct <- as.matrix(sigs)  %*% exposure
  rec.sums <- margin.table(all.reconstruct, 2)
  stopifnot(abs(ex.sums - rec.sums) < 1)
  spect.sums <- margin.table(spectrum, 2)
  stopifnot(abs(spect.sums - ex.sums) < 1)
}

plot.recon.and.loglh <- function(spect, sigs, ex, range,
                                 obj.fun,
                                 nbinom.size) { 
  plot.reconstruction(signatures      = sigs,
                      exposures.mat   = ex[ , range, drop=F],
                      input.genomes   = spect[ , range, drop=F],
                      normalize.recon = T)


  neg.ll <- compute.all.neg.log.lh(spect[ ,range,drop=F], sigs=sigs,
                                   exp = ex[ , range, drop=F],
                                   obj.fun=obj.fun,
                                   nbinom.size=nbinom.size)
  s.names <- names(neg.ll)
  l.range <- 1:length(s.names)
  plot(neg.ll, xaxt='n', ylab=('Neg ln likelihood'), xlab='', new=T)
  axis(side = 1, labels = s.names, at=l.range, las=2)
  abline(v=l.range, lty=3)
}

# plot.recon.by.range calls the objective function as one
# of its analyses.
plot.recon.by.range <- function(path, spect, sigs, ex, range,
                                obj.fun,
                                nbinom.size) {
  cairo_pdf(path,
            width=8.2677, height=11.6929, # for A4
            onefile=T)
  par(
    mfrow=c(3,1), # 3 graphs per page
    mar=c(1.5,1.1,4.6,1), # space between plot and figure, for axis labels etc
    oma=c(4,6,3,3) # outer margin between figure and device.
  )

  for (r in range) {
    plot.recon.and.loglh(spect, sigs, ex, r,
                         obj.fun=obj.fun,
                         nbinom.size=nbinom.size)
  }
  dev.off()
}


# process.one.group
#
# Test and plot one group of spectra
#
# Output is an R a list with the elements
# pval
# adj.p.val
# exposure
#
# Side effects are to generate 4 pdfs based on the input argument
# path.root. The names are:
# <path.root>.check.with.sig.pdf
# <path.root>.pos.with.sig.pdf ## FIX ME, probably removed
# <path.root>.exposures.pdf
# <path.root>.reconstruction.err.pdf

process.one.group <- function(spectra, sigs, target.sig.name,
                              path.root,
                              obj.fun,
                              nbinom.size,
                              trace=0,
                              col=NULL,
                              mc.cores=190) {

  target.sig.index <- which(colnames(sigs) == target.sig.name)
  # check if signatures sum to 1
  stopifnot(colSums(sigs) == rep(1, ncol(sigs)))
  # Need to match exactly one signature name
  stopifnot(length(target.sig.index) == 1)

  s.spectra <- sort.spectra.columns(spectra)
  s.spectra.to.list <- split(t(s.spectra), 1:ncol(s.spectra))

  out.pvals <-
      mclapply(
          X=s.spectra.to.list,
          FUN=signature.presence.test,
          sigs=sigs,
          target.sig.index=target.sig.index,
          trace=trace,
          obj.fun=obj.fun,
          nbinom.size=nbinom.size,
          mc.cores=mc.cores)

  out.pvals  <- unlist(out.pvals)
  names(out.pvals)  <- colnames(s.spectra)

  low.pval <- which(out.pvals < 0.05)
  if (length(low.pval) > 0) {
    # Have to wrap column-wise index of s.spectra in as.matrix in case
    # length(low.pval) == 1, in which case indexing returns a vector
    
    check.w.sig <- s.spectra[, low.pval, drop=F]
    # The column names are lost if length(low.pval) == 1
    colnames(check.w.sig) = colnames(s.spectra)[low.pval]
    spec.path <- paste(path.root, 'check.with.sig.pdf', sep='.')
    pdf.mut.sig.profile(path=spec.path, check.w.sig)

  }
  

  out.exp <-
      mclapply(
          X=s.spectra.to.list,
          FUN=sparse.assign.activity,
          sigs=sigs,
          obj.fun=obj.fun,
          nbinom.size=nbinom.size,
          mc.cores=mc.cores)

  out.exp  <-  do.call(cbind, out.exp)
  colnames(out.exp)  <-  colnames(s.spectra)
  sanity.check.ex(s.spectra, sigs, out.exp)

  # Plotting part
  hist.path <- paste(path.root, 'pval.histogram.pdf', sep='.')
  pdf(hist.path, useDingbats = F)
  hist(out.pvals, breaks=seq(from=0, to=1, by=0.01))
  dev.off()

  approx.num.per.row <- 30
  starts <- seq(from=1, to=ncol(s.spectra), by=approx.num.per.row)
  ranges <-
    lapply(starts,
           function(x) {
             x:(min(x+approx.num.per.row-1, ncol(s.spectra)))
             } )
  exp.path <- paste(path.root, 'exposures.pdf', sep='.')
  pdf.ex.by.range(exp.path, s.spectra, sigs, exp=out.exp,
                  range=ranges, col=col)
  recon.path <- paste(path.root, 'reconstruction.err.pdf', sep='.')

  plot.recon.by.range(recon.path, s.spectra,
                      sigs,
                      out.exp,
                      range = ranges,
                      obj.fun=obj.fun,
                      nbinom.size=nbinom.size)

  list(pval=out.pvals, exposure=out.exp)
}

# Self-contained test for mSigAct
# Takes a few minutes to run on a laptop

mSigAct.basic.test <- function() {
  liver.wes.sigs <-
    structure(
      c(0.00501651202811263, 0.00654743825297215, 0.00172198612846377,
        0.0027812414590599, 0.00456505896068523, 0.00546880487367255,
        0.00163313281919571, 0.00472311594276702, 0.00550473134775308,
        0.00523431509007715, 0.00117445523004569, 0.0031417255804599,
        0.00609165475854316, 0.00747396740197616, 0.00334532173007199,
        0.0073459935080331, 0.00081409404708735, 0.00184694609091571,
        0.000684772590591487, 0.00132237943387946, 0.000889348167196588,
        0.000522970017412516, 0.000926076106969921, 0.000907968031975508,
        0.00040260939331999, 0.00217567088867182, 4.73593032077635e-07,
        0.000627122436853997, 0.000932318288055095, 0.00138429522473908,
        0.000448244555915337, 0.00151722198714391, 0.0133407511750442,
        0.0102496239532803, 0.198362181314223, 0.00563207894511671, 0.0144626123888998,
        0.013780597232066, 0.174824403488048, 0.0112470908225736, 0.0166786834347028,
        0.0246910888370856, 0.197358877868819, 0.0124637064919174, 0.00720656395359164,
        0.0118386332541352, 0.111734723905107, 0.00662512020241512, 0.0010223223625619,
        0.00122210356639981, 0.00129413788895533, 0.0023701912387363,
        0.000387561477579557, 0.00125248107806446, 0.00105799892563039,
        0.00106791580207752, 0.000308232252560773, 0.000370962415771247,
        0.000775241687053448, 0.00091272981357242, 0.00141507820771524,
        0.00096374673775006, 0.000492758318963667, 0.00126302506972582,
        0.00353756909550728, 0.00323415662711626, 0.00466734940317019,
        0.00262402559901603, 0.00136879379537905, 0.00345675080783413,
        0.0049866809864481, 0.00328988254526685, 0.00502973866899031,
        0.00470056115266067, 0.00321508472775254, 0.00462070596505851,
        0.00204008599923715, 0.00234172065385724, 0.00376347665668317,
        0.00198513840129774, 0.000403597663136818, 0.000919532578622957,
        0.000638033906149146, 0.000895398084998278, 9.91989672157247e-05,
        0.00138103208303637, 0.00113734234259544, 0.00135173031562382,
        4.42903422784128e-05, 0.000135895620631068, 0.000767571099429845,
        0.000346842230809169, 0.000353141590496185, 0.000622740338405309,
        0.000918137355326322, 0.00127860432017847, 0.0231953617806992,
        0.0310888032579507, 0.0297329386873131, 0.0152422851349562, 0.0448579204216992,
        0.0642966842489053, 0.0226305057044867, 0.0399166334611914, 0.0353468454858301,
        0.0509945069459697, 0.0608147975482152, 0.0255568058953173, 0.0230714446047142,
        0.050646151397085, 0.0214960728567165, 0.0250430480980922, 0.00616424682939131,
        0.00543299474410789, 0.00503672731861588, 0.00338717447443471,
        0.0102170968422525, 0.0101576194986056, 0.016201384765712, 0.00868556376238888,
        0.00658052974470241, 0.011630326145572, 0.0166102090219795, 0.00594344323146914,
        0.00223723099197229, 0.00988447192694038, 0.00640748325536742,
        0.00338419568893138, 0.00762587236625729, 0.00754582603348318,
        0.00454930209423369, 0.00370080174058607, 0.0204341936845051,
        0.0150793526577237, 0.0195445276538749, 0.0185723225131932, 0.00817865839698729,
        0.0102244625455578, 0.00616185173396013, 0.00812270574967449,
        0.00244697139746969, 0.00658964795129359, 0.00227362309061425,
        0.00521166136095433, 0.00171554749700632, 0.0028260417008971,
        0.00647287954930368, 0.0011956802195276, 0.00345567214726311,
        0.0102702541314993, 0.0341887966265224, 0.00792269432149424,
        0.00433799937022567, 0.00490948604271471, 0.0120778221163859,
        0.00274047555291181, 0.00159866251473677, 0.00311784588463404,
        0.0052114557752384, 0.0020072142590673, 0.00300220811976106,
        0.0014492521543062, 0.00524303243493598, 0.00143481626343312,
        0.00239593268876909, 0.00240564511188272, 0.00999672415980188,
        0.0037202216813973, 0.00383504292150385, 0.00151061109006606,
        0.00381404908938503, 0.00146158696155297, 0.000994723342502879,
        0.00108446813378575, 0.0022879561940071, 0.000981304748877347,
        0, 0.00014492521543062, 0.000970931932395551, 7.97120146351735e-05,
        0, 0.00120282255594136, 0.00459849311350886, 0.000826715929199399,
        0, 0, 0.0016345924668793, 0.000121798913462747, 0, 0.000203337775084828,
        0.00190663016167259, 0.000490652374438673, 0.0112116004168735,
        0.0106453946767298, 0.00423564215876876, 0.00681900836822725,
        0.0111160022974026, 0.0061233762201889, 0.00850414475237292,
        0.0120159442874384, 0.0132004163114833, 0.0139558203323851, 0.00888644442581266,
        0.0120487300089916, 0.0121961612785602, 0.0133868799053278, 0.00952395037444146,
        0.0145799112672321, 0.00875751524435851, 0.00866285810947968,
        0.00441980031331175, 0.00866271507160414, 0.0086602536428113,
        0.0094382636744811, 0.00793672074453467, 0.0102742702225265,
        0.00617101046677666, 0.0081321691380021, 0.0034815493214597,
        0.00706605897377331, 0.00221325082674627, 0.00880795377045699,
        0.00321928341522986, 0.00531028418753381, 0.0163873415068199,
        0.0151540030441126, 0.0321535617580169, 0.0122042431554683, 0.0261607325666717,
        0.0216499601828251, 0.0390563679750653, 0.022289068608528, 0.0222426502412773,
        0.0271972347148896, 0.041736653438069, 0.0228113055833111, 0.00907864302257166,
        0.0229478529997239, 0.0183883754740834, 0.00942040341261076,
        0.00375713632510779, 0.00633085856681101, 0.00879555120943424,
        0.00526778707955995, 0.00274370905595059, 0.00678316962823705,
        0.0124029130343501, 0.00795946141104214, 0.00500367404387849,
        0.0044832702580177, 0.00730676786275787, 0.00366805029772905,
        0.00386155190736525, 0.0054704233912498, 0.00536121325271232,
        0.00539362789031376, 0.0149252020090427, 0.0117827427623739,
        0.0217434803004828, 0.0128489455250127, 0.00774645865171492,
        0.0136447950501854, 0.0221222265488299, 0.0154998269446264, 0.0120964529209212,
        0.0106737070145073, 0.0151231008921933, 0.0112657349078887, 0.00715612478481164,
        0.0113610283321237, 0.00945276569350174, 0.00915109890169289,
        0.00146107119698508, 0.00192221535238833, 0.00419614762892712,
        0.00179829249910159, 0.00111074934929267, 0.00380026796237097,
        0.00843670668880941, 0.00934411754845173, 0.00120828399986435,
        0.000365358416659753, 0.00637875499070535, 0.00237833279743381,
        0.00218248546063202, 0.00410689894605194, 0.00454403131799012,
        0.00704164750054199, 0.000762935772711461, 0.00198945594983114,
        0.000573703528543225, 0.000841643512363437, 0.00694049356123817,
        0.0178224752248122, 0.0165266089171643, 0.037259501372356, 0.00159332668342445,
        0.00523491056000767, 0.00397313639158346, 0.00608606928659376,
        0.000839345064546425, 0.00192793746193099, 0.00160564393192749,
        0.00277232555474576, 0.00058342147324994, 0.000852623978499059,
        0, 0.00079734648539694, 0, 0, 0, 0.000130506134404049, 0, 0.00270771235862466,
        0, 0.0011892319295643, 0, 0.000132961204271103, 0, 4.77987164611339e-05,
        0.0140021153579986, 0.0115814757079456, 0.10418456078345, 0.00660025701800801,
        0.00584100943272519, 0.00732126575625064, 0.163631589388627,
        0.00567701684657613, 0.043351763511507, 0.0697687217738953, 0.253334744206202,
        0.0366563253583348, 0.0036536196927315, 0.00445420034308195,
        0.0570733433985137, 0.00224653967367329, 0.000151441021594163,
        0.00168872397806437, 0, 0.00149176505900183, 3.25388380445231e-05,
        0.00169887853981198, 0.000564778904739306, 0.000535178324956747,
        0, 0.00186690186051041, 0.000384785775979266, 0.000903157482347644,
        5.01770407589817e-05, 0.000382928394237807, 0, 0.000220500729617936,
        0.00189301276992704, 0.00286571341732136, 0.00991945999974664,
        0.000647369742585698, 0.00201740795876043, 0.00261365929201843,
        0.0190612880349516, 0.00160553497487024, 0.00541665276097331,
        0.00393382892036123, 0.00737506070626927, 0.00180631496469529,
        0.00148022270238996, 0.00167531172479041, 0.00475753334239466,
        0.000913503022702878, 0, 0.000869948715972555, 0.000319982580636988,
        0.000816248805868924, 3.25388380445231e-05, 0.00261365929201843,
        0.00352986815462066, 0.00418412144966184, 0, 0.00106680106314881,
        0.000641309626632111, 0.00150526247057941, 0.000225796683415418,
        0.000909454936314792, 0.000493706290248502, 0.00226800750464163,
        0.00664448380322634, 0.00642104359845698, 0.0037505754166512,
        0.00391799498236882, 0.0178375094010451, 0.0159257709187271,
        0.00977759071157951, 0.00890821609831549, 0.0079143771419304,
        0.00971853412857413, 0.0054567857567032, 0.00363172427094147,
        0.00626566281705966, 0.0126549840923646, 0.00533263210226752,
        0.00450343376150557, 0.00267505192077944, 0.0020492692335501,
        0, 0.00212934509911349, 0.00251046428607301, 0.00270169328085549,
        0, 0.00238388881504217, 0.00153181493069621, 0.00190899777525563,
        0, 0.00121057475698049, 0.00227842284256715, 0.00345135929791763,
        0, 0.00137860217188946, 0.0104413316907842, 0.00737736924078036,
        0.00419181958331605, 0.00570664486562415, 0.0247082537629291,
        0.0133662720210745, 0.016063184740452, 0.0257209056359813, 0.0158287542838608,
        0.0159661632112289, 0.00836707149361157, 0.0154684552280841,
        0.0131009313447611, 0.0121436716037842, 0.00280664847487764,
        0.00827161303133676, 0.00281482719642406, 0.0063957216045446,
        0.00404312895158325, 0.0032471868237748, 0.00106361003536157,
        0.0108048538861134, 0.00420805723686217, 0.00505162443438572,
        0.00128054218104452, 0.00499987939784643, 0.00234289499553169,
        0.00256352376561434, 0.00274967502754125, 0.0121487887241491,
        0.00319307532780954, 0.00569339491427701, 0.0226156805781657,
        0.0311914422867791, 0.0428044304221966, 0.0135299450990617, 0.0195203724136946,
        0.0483705668157401, 0.0563336694612194, 0.0303097466063143, 0.0489167113159008,
        0.0351273578207672, 0.0505572077983153, 0.033491197583026, 0.0241199563819408,
        0.0314764071489317, 0.0276157866188933, 0.0224707395020933, 0.00058237804063946,
        0.000983957169929939, 0.00158209393757605, 0.0016235934118874,
        0.00081334885057061, 0.00615625395836692, 0.00610847018254186,
        0.00589356184011667, 0, 0.000512808143368865, 0.00135641289214992,
        0.00264621808063416, 0.00091655834251375, 0.00248497951175777,
        0.000949292665024459, 0.0019381769920943, 0.0138004655305096,
        0.0131636653737059, 0.00482135378723064, 0.00744536605065992,
        0.0208686766319906, 0.0141247178231626, 0.00763121768086928,
        0.0104690920842893, 0.0121728879389081, 0.0162045856048199, 0.0130090470183865,
        0.00841757495811504, 0.0115048312709967, 0.0184116092775063,
        0.0111518097532399, 0.0143332504799258, 0.00411442450599043,
        0.00325698937081384, 0, 0.00617626956475198, 0.00524998154263913,
        0.0070623589115813, 0.0097124588665609, 0.00997056388979938,
        0.00393082839693907, 0.00551645467398124, 0, 0.00387475672675137,
        0.00471509478319536, 0.00812650340524418, 0, 0.0154287855484552,
        0.0115718189230981, 0.0103137996742438, 0.0076703355705942, 0.00854524967178013,
        0.0275624030988555, 0.0138422234666993, 0.00554997649517766,
        0.0262973622593459, 0.0158501145037866, 0.0158598071876961, 0.00325226175459661,
        0.0154990269070055, 0.016219926054192, 0.0166339366576092, 0.0100366287779159,
        0.0176198556855139, 0.00390486570634956, 0.0071350396938804,
        0.00698468921662536, 0.00451577869677459, 0.00335602048816126,
        0.0159745177197918, 0.0175291410455944, 0.00892084240392306,
        0.00254402411128988, 0.00509392154552865, 0.00624693142526532,
        0.00271073418545209, 0.00407307848043987, 0.0079538178237248,
        0.00540064123715534, 0.00481316534592131, 0.0345652927339832,
        0.0167135861322404, 0.0303833980923203, 0.0267183572892497, 0.0090736850235471,
        0.0242113784190594, 0.0256195138358688, 0.0244393911690809, 0.0136529293972557,
        0.0126074558251834, 0.018128350018417, 0.0116643713434605, 0.0117879683080966,
        0.0167304443878349, 0.0141445365735021, 0.0104686346273788, 0.00250682736703923,
        0.00410509133072571, 0.00532582552767684, 0.00408570453517701,
        0.000124297055117084, 0.0102336754142416, 0.00903424961580637,
        0.0172841321576009, 0, 0.00407513723642292, 0.000979910811806324,
        0.00361431224726945, 0.0032584627843519, 0.00630820034295415,
        0.0042004987400097, 0.00980682439231467, 0.00147508378959959,
        0.00043626371379819, 3.24476568379931e-06, 0.00252699004168052,
        0.0012728501962151, 0.000559741657616993, 0.000969220589952107,
        3.6853215672797e-05, 0.00778419486399129, 0.00209506438514141,
        0.00435725473126722, 0.00389542028219519, 0.00161620697536921,
        0.00153447043333608, 0.00569052458149112, 0.00429835543324281,
        0.0013379515501418, 0.00207388342403985, 5.31149486669254e-05,
        0.00105581112685113, 0.00237324483852749, 0.000941174301020882,
        1.14622435731602e-05, 0.00111659343273038, 0.0015440267793825,
        0.000422251650657611, 1.48591205166154e-05, 0.00228470620381426,
        0.000347842955339254, 0.000781957395261438, 3.57246186920401e-05,
        0.00244608356926855, 0.00752871687353439, 0.00538489861164894,
        0.0189989592883657, 0.00455856756720577, 0.0262171926996803,
        0.00510443100792052, 0.0429316181273128, 0.011295019271077, 0.00246405386613427,
        0.000527057173493617, 0.0254956105410327, 0.00116180757015381,
        0.0142621917750927, 0.0200140094962501, 0.0426135442198519, 0.0170031740550405,
        0.000277588927001894, 0.00172474053577807, 0.000891754167861606,
        0.0010329286404854, 1.75596771138327e-05, 0.0119852395328507,
        0.00565790386535873, 0.0272729647570705, 6.80715846686e-06, 0.00307843276650053,
        0.00407168491936454, 0.00299304124046668, 0.000314545301154509,
        0.00221506727378633, 1.70175827943991e-05, 0.00219453039478919,
        0.00122143792080324, 0.00741658982164508, 0.00363547341054731,
        0.00158581682324754, 0.00433573822226979, 0.0464896701760888,
        0.0362367583415844, 0.0889197616358184, 9.08185332730424e-06,
        0.0115527500730291, 0.00535492998977843, 0.00628794337628364,
        0.000769059042177833, 0.00951011305742527, 0.00477666665857248,
        0.004073685899485, 2.40633066122837e-05, 0.00593891014197043,
        0.00285527814222454, 0.0156171288576823, 5.33982863926338e-05,
        0.0232281803051761, 0.0170354312654747, 0.228219671379932, 3.12202504056426e-05,
        0.0108598374686808, 0.0055029474707368, 0.0489948219212932, 6.00499981958012e-05,
        0.00825562073615158, 0.00364165284666774, 0.0328012246209437,
        0.00127871234044587, 0.00329903703682902, 0, 0.000774077111396133,
        0.00592265860572756, 0.00527376656105381, 0.00309691506798015,
        0.00548825764598788, 0.00163289312816462, 0.00309410339675284,
        0.00119650829529023, 0.000687978120692226, 0.00378954898798574,
        0.00345044276300746, 0.00141895118837052, 0.00326486738362238,
        0.000448585979859865, 0, 0, 0.000251287767305087, 0.000757653805510864,
        0.0011561874378598, 0.00131462420478063, 0.00217639933620949,
        0, 0, 0.00076949103566909, 0.000713542132538989, 0.000210904427321826,
        0.000791772813153455, 0.00109447730301379, 0.000233049619237001,
        0.00264690391487464, 0.00205244717616127, 0.00788075769415027,
        0, 0.0100644471300607, 0.00996180783390549, 0.0189291461986885,
        0.00515441539393097, 0.00240886303497761, 0.00552119272791768,
        0.0118296682232369, 0.00147635806104525, 0.00188626725551309,
        0.00250723863726625, 0.00414353625274554, 0.00107048986353727,
        0.0237250935105943, 0.0113099296315293, 0.0588115385218224, 0.00778467704970178,
        0.0535648856742036, 0.0664789684970726, 0.285156856611354, 0.0569840358097924,
        0.0451736355963403, 0.0119858184287638, 0.0611010577322677, 0.0086372479792596,
        0.0193960202683493, 0.0231862582486319, 0.0593852088124657, 0.0140703255587071,
        0.00805189717232644, 0.00107436775141007, 0.00425495704319472,
        0.00140259552102006, 0.00248248509671461, 0.00167407101290031,
        0.00367734243983605, 0.00217821202763508, 0.00354721215864901,
        0, 0.00179180772782759, 0.00127181694476048, 0.0022830415817287,
        0.000827109581089335, 0.00151490323133696, 0, 0.00078989177182593,
        0, 0.000941179916978616, 0, 0.000690595768796278, 0.00131775413794913,
        0.00560169805209201, 0.00150176416209596, 0, 0, 0.00228104562645118,
        0, 0.000785003398733137, 0, 0.00218745705001512, 0, 0.000273350881011472,
        0.00035014164329096, 0, 0, 0.000152091173998725, 0.000732119934407473,
        0.000400037892175032, 0.000633600694804806, 0.000312093259815045,
        0.000689168414207109, 0.00037979921900936, 0, 0.00137394731151464,
        0.000622010359989938, 0.000194918325917101, 0.000133206608533283,
        0, 0, 0, 0, 0.00018376676442616, 0.000281944865676523, 0, 0.000171631255336515,
        0, 0.00016673431722953, 0, 0, 0.000152969636393913, 0.000148075418125197,
        0, 0.000281886694483321, 0.0119184769882564, 0.0781618367472312,
        0.0142572731003575, 0.0197640733468222, 0.0559220324602966, 0.185172999554935,
        0.0369629174071061, 0.136774753704552, 0.0475239069838959, 0.197555356291589,
        0.0331623708679271, 0.101118055943648, 0.0096571547936287, 0.0388620797621748,
        0.0053981332799417, 0.015623351692409, 0, 0, 0.000145807681376713,
        0, 0.000259903584434659, 0, 0.000164200425130804, 0.000299248234464245,
        0, 0.000164110166149587, 0, 0, 0, 0, 0.000139608719653132, 0,
        0.000362312690749786, 0.000152334934592573, 0.000306332868677996,
        0.000118451351432588, 0, 0, 0.00030418067958214, 0, 0.000149821472213559,
        0.000482124974808281, 0, 0.000575361300165641, 0, 0.000148300016137133,
        0.000144425562258585, 0, 0, 0, 0, 0, 0, 0.00016692056416241,
        0, 0.000137811699825254, 0, 0, 0.000161233134503067, 0, 0, 0,
        0.000149242344564564, 0, 0.0163205434212236, 0.0182336644084891,
        0.0298303172739799, 0.0138338778695703, 0.0554472052305289, 0.0316917751041016,
        0.0517110882941659, 0.0166393829315822, 0.0460813899876904, 0.125977314697917,
        0.111745751949063, 0.0447844431094846, 0.0197133937639833, 0.0527803974391035,
        0.0237239009724563, 0.0183847313192857, 0.00683282000048767,
        0.00759948330485679, 0, 0.0021826936500059, 0.0117567733371852,
        0.0151203531540017, 0.00183860943922507, 0.00860582269328657,
        0.00416770430500731, 0.0107361060014385, 0.0135121506307147,
        0.00271375102706354, 0.00640634957173113, 0.00639590735224538,
        0.00295138654696075, 0.00370194126761636, 0.00359815382129189,
        0.00346759438304507, 0.00453108656885968, 0.00192347052382033,
        0.00526514256672385, 0, 0.00674238664368164, 0.00328968677767334,
        0.0199567386587386, 0.0182403952444163, 0.0303787839170938, 0.0161005910599866,
        0.0166550453652636, 0.0101746326932656, 0.00993859894329533,
        0.00335713865414839, 0.00037385284885917, 9.85710773916278e-06,
        0.00367478668507847, 1.77488120652309e-06, 0.00114284033490448,
        0.00223507865058502, 0.00619400293359609, 0.00180055296024357,
        0.00125417376293988, 0.00410788979753786, 0.000159375035172038,
        0, 0, 0.00485769709586381, 0.00638276412012252, 0.000108202536402443,
        0.0015143229974768, 0.000407944417847494, 0.00158062408469914,
        0.00254495838243838, 0.000763733123886472, 0.00369437214725243,
        0, 0.000986616113915783, 0.000715683495797367, 0.00567988662372263,
        0.000226074648274699, 0.00231411946903916, 0.000535806348108985,
        0.00142467116987079, 0.000274220703232252, 0.00014322083624978,
        0, 0.000951048939888829, 0.000491877926330202, 0.000384280731988633,
        0, 0.00181283808975711, 0.000102089451864033, 0.00156493333990863,
        0, 0.000908763153516299, 0.00238111473642821, 0.000450486277160574,
        0, 0, 0.000775189300463072, 2.77988648762916e-05),
      .Dim = c(96L,
               10L),
      .Dimnames =
        list(c("ACAA", "ACCA", "ACGA", "ACTA", "CCAA",
               "CCCA", "CCGA", "CCTA", "GCAA", "GCCA", "GCGA", "GCTA", "TCAA",
               "TCCA", "TCGA", "TCTA", "ACAG", "ACCG", "ACGG", "ACTG", "CCAG",
               "CCCG", "CCGG", "CCTG", "GCAG", "GCCG", "GCGG", "GCTG", "TCAG",
               "TCCG", "TCGG", "TCTG", "ACAT", "ACCT", "ACGT", "ACTT", "CCAT",
               "CCCT", "CCGT", "CCTT", "GCAT", "GCCT", "GCGT", "GCTT", "TCAT",
               "TCCT", "TCGT", "TCTT", "ATAA", "ATCA", "ATGA", "ATTA", "CTAA",
               "CTCA", "CTGA", "CTTA", "GTAA", "GTCA", "GTGA", "GTTA", "TTAA",
               "TTCA", "TTGA", "TTTA", "ATAC", "ATCC", "ATGC", "ATTC", "CTAC",
               "CTCC", "CTGC", "CTTC", "GTAC", "GTCC", "GTGC", "GTTC", "TTAC",
               "TTCC", "TTGC", "TTTC", "ATAG", "ATCG", "ATGG", "ATTG", "CTAG",
               "CTCG", "CTGG", "CTTG", "GTAG", "GTCG", "GTGG", "GTTG", "TTAG",
               "TTCG", "TTGG", "TTTG"),
             c("Signature.1", "Signature.4", "Signature.5",
               "Signature.6", "Signature.12", "Signature.16", "Signature.17",
               "Signature.AA", "Signature.23", "Signature.24")))


  short.taiwan.hcc2 <-
    structure(
      c(0L, 2L, 0L, 3L, 0L, 1L, 0L, 1L, 0L, 0L, 0L, 1L, 1L,
        1L, 0L, 1L, 0L, 0L, 0L, 0L, 1L, 0L, 0L, 0L, 0L, 1L, 0L, 0L, 0L,
        0L, 1L, 0L, 0L, 0L, 7L, 1L, 3L, 1L, 2L, 0L, 2L, 1L, 3L, 1L, 1L,
        1L, 0L, 1L, 0L, 1L, 1L, 0L, 0L, 3L, 3L, 0L, 0L, 0L, 1L, 0L, 0L,
        0L, 0L, 0L, 0L, 2L, 0L, 1L, 0L, 2L, 1L, 0L, 1L, 1L, 1L, 1L, 1L,
        0L, 0L, 0L, 0L, 0L, 0L, 0L, 0L, 0L, 0L, 1L, 0L, 0L, 1L, 0L, 0L,
        0L, 0L, 0L, 1L, 1L, 0L, 1L, 0L, 1L, 0L, 1L, 1L, 0L, 1L, 1L, 0L,
        1L, 0L, 1L, 0L, 2L, 0L, 2L, 1L, 0L, 0L, 0L, 2L, 3L, 0L, 0L, 0L,
        2L, 0L, 0L, 2L, 0L, 1L, 1L, 0L, 1L, 3L, 0L, 1L, 2L, 1L, 0L, 0L,
        2L, 1L, 0L, 0L, 0L, 1L, 0L, 0L, 0L, 0L, 2L, 2L, 1L, 0L, 0L, 0L,
        0L, 1L, 0L, 0L, 2L, 1L, 1L, 3L, 0L, 0L, 0L, 2L, 1L, 0L, 1L, 0L,
        0L, 0L, 0L, 0L, 0L, 0L, 0L, 0L, 1L, 0L, 1L, 0L, 0L, 0L, 0L, 0L,
        0L, 0L, 0L, 0L, 1L, 0L, 0L, 1L, 0L, 3L, 0L, 2L, 1L, 1L, 0L, 2L,
        0L, 0L, 2L, 0L, 0L, 0L, 0L, 0L, 0L, 0L, 0L, 0L, 0L, 0L, 0L, 1L,
        0L, 0L, 1L, 2L, 3L, 3L, 0L, 1L, 1L, 0L, 0L, 1L, 3L, 0L, 1L, 1L,
        2L, 1L, 0L, 0L, 0L, 2L, 1L, 2L, 0L, 2L, 1L, 0L, 0L, 2L, 1L, 0L,
        0L, 1L, 0L, 0L, 0L, 3L, 0L, 0L, 0L, 1L, 0L, 0L, 0L, 0L, 0L, 0L,
        1L, 2L, 0L, 0L, 0L, 0L, 0L, 0L, 1L, 1L, 1L, 0L, 0L, 0L, 0L, 1L,
        0L, 0L, 0L, 0L, 0L, 0L, 0L, 0L, 0L, 0L, 0L, 2L, 0L, 0L, 0L, 1L,
        1L, 2L, 0L, 0L, 0L, 0L, 0L, 1L, 0L, 0L, 1L, 0L, 0L, 0L, 1L, 2L,
        0L, 0L, 2L, 1L, 1L, 2L, 1L, 2L, 0L, 1L, 3L, 2L, 3L, 2L, 4L, 0L,
        0L, 0L, 1L, 1L, 1L, 1L, 0L, 0L, 1L, 0L, 0L, 0L, 0L, 1L, 0L, 0L,
        0L, 0L, 0L, 2L, 1L, 2L, 1L, 0L, 3L, 0L, 0L, 0L, 0L, 0L, 0L, 0L,
        1L, 0L, 0L, 0L, 0L, 0L, 0L, 0L, 0L, 0L, 1L, 0L, 0L, 0L, 0L, 0L,
        1L, 0L, 0L, 1L, 2L, 0L, 0L, 1L, 0L, 1L, 0L, 0L, 0L, 0L, 0L, 3L,
        1L, 0L, 1L, 0L, 0L, 0L, 1L, 0L, 0L, 0L, 0L, 0L, 1L, 0L, 0L, 0L,
        0L, 0L, 1L, 1L, 1L, 0L, 0L, 2L, 0L, 4L, 0L, 1L, 2L, 1L, 2L, 0L,
        0L, 1L, 0L, 0L, 0L, 2L, 1L, 2L, 0L, 2L, 0L, 1L, 1L, 0L, 0L, 1L,
        0L, 2L, 0L, 2L, 0L, 1L, 2L, 0L, 0L, 0L, 0L, 0L, 1L, 1L, 0L, 0L,
        1L, 1L, 1L, 0L, 0L, 1L, 0L, 0L, 1L, 0L, 0L, 0L, 0L, 0L, 0L, 0L,
        0L, 0L, 0L, 1L, 1L, 0L, 0L, 0L, 1L, 0L, 4L, 0L, 2L, 0L, 0L, 2L,
        0L, 0L, 1L, 1L, 0L, 0L, 0L, 0L, 1L, 1L, 0L, 0L, 1L, 1L, 0L, 0L,
        0L, 0L, 0L, 0L, 2L, 2L, 1L, 2L, 0L, 1L, 3L, 0L, 1L, 2L, 2L, 1L,
        2L, 1L, 1L, 0L, 0L, 0L, 0L, 0L, 0L, 0L, 0L, 0L, 0L, 0L, 0L, 1L,
        0L, 1L, 0L, 0L, 0L, 0L, 0L, 1L, 0L, 0L, 1L, 1L, 3L, 0L, 1L, 1L,
        1L, 0L, 0L, 0L, 1L, 0L, 0L, 0L, 0L, 0L, 0L, 0L, 0L, 0L, 1L, 0L,
        0L, 0L, 0L, 1L, 1L, 0L, 1L, 1L, 0L, 0L, 3L, 1L, 0L, 0L, 0L, 0L,
        3L, 0L, 2L, 0L, 0L, 0L, 0L, 0L, 0L, 0L, 0L, 1L, 2L, 0L, 0L, 0L,
        0L, 0L, 2L, 0L, 1L, 1L, 1L, 3L, 1L, 2L, 1L, 0L, 0L, 1L, 0L, 0L,
        0L, 2L, 2L, 1L, 0L, 1L, 1L, 0L, 1L, 2L, 0L, 1L, 0L, 0L, 0L, 0L,
        0L, 0L, 1L, 0L, 1L, 0L, 0L, 0L, 0L, 0L, 0L, 0L, 1L, 0L, 2L, 0L,
        0L, 1L, 0L, 0L, 0L, 0L, 0L, 0L, 0L, 0L, 0L, 0L, 1L, 0L, 0L, 0L,
        0L, 0L, 0L, 1L, 1L, 0L, 0L, 0L, 0L, 0L, 1L, 0L, 2L, 0L, 1L, 1L,
        1L, 0L, 0L, 0L, 1L, 0L, 0L, 0L, 0L, 0L, 0L, 0L, 0L, 0L, 0L, 0L,
        1L, 1L, 0L, 1L, 0L, 1L, 0L, 2L, 1L, 2L, 0L, 0L, 0L, 1L, 0L, 0L,
        0L, 0L, 3L, 1L, 1L, 2L, 0L, 0L, 0L, 0L, 0L, 1L, 0L, 0L, 0L, 0L,
        0L, 0L, 0L, 5L, 0L, 1L, 1L, 1L, 0L, 1L, 1L, 1L, 0L, 1L, 2L, 0L,
        0L, 0L, 1L, 0L, 0L, 0L, 0L, 1L, 0L, 0L, 0L, 0L, 0L, 0L, 0L, 0L,
        1L, 0L, 1L, 1L, 0L, 0L, 0L, 0L, 0L, 1L, 3L, 1L, 1L, 0L, 1L, 0L,
        1L, 0L, 0L, 0L, 0L, 0L, 1L, 0L, 0L, 0L, 2L, 0L, 0L, 0L, 0L, 0L,
        0L, 0L, 0L, 1L, 1L, 1L, 2L, 3L, 2L, 4L, 0L, 2L, 1L, 3L, 1L, 1L,
        0L, 0L, 1L, 0L, 0L, 0L, 1L, 0L, 1L, 1L, 0L, 0L, 0L, 0L, 0L, 0L,
        0L, 0L, 0L, 0L, 0L, 1L, 1L, 0L, 0L, 1L, 0L, 1L, 0L, 0L, 0L, 0L,
        0L, 0L, 0L, 0L, 0L, 0L, 0L, 0L, 1L, 0L, 0L, 0L, 0L, 0L, 1L, 0L,
        0L, 0L, 0L, 0L, 1L, 0L, 1L, 1L, 2L, 1L, 0L, 0L, 0L, 0L, 1L, 0L,
        1L, 0L, 0L, 0L, 0L, 0L, 0L, 0L, 1L, 0L, 1L, 0L, 0L, 0L, 0L, 1L,
        0L, 0L, 1L, 0L, 0L, 1L, 1L, 1L, 1L, 0L, 0L, 1L, 1L, 0L, 0L, 0L,
        0L, 0L, 0L, 0L, 1L, 0L, 0L, 2L, 0L, 3L, 1L, 1L, 1L, 1L, 1L, 1L,
        1L, 2L, 0L, 0L, 1L, 0L, 1L, 1L, 0L, 1L, 0L, 1L, 0L, 3L, 0L, 0L,
        0L, 0L, 1L, 0L, 0L, 0L, 0L, 0L, 0L, 1L, 0L, 0L, 0L, 0L, 0L, 0L,
        0L, 0L, 1L, 0L, 0L, 0L, 0L, 0L, 1L, 0L, 1L, 1L, 0L, 0L, 0L, 0L,
        1L, 0L, 0L, 0L, 0L, 0L, 0L, 0L, 1L, 0L, 0L, 0L, 0L, 0L, 0L, 0L,
        1L, 0L, 0L, 0L, 0L, 1L, 0L, 1L, 1L, 2L, 1L, 0L, 0L, 1L, 1L, 0L,
        1L, 0L, 0L, 0L, 0L, 0L, 0L, 1L, 2L, 4L, 1L, 0L, 0L, 0L, 0L, 0L,
        0L, 1L, 1L, 1L, 0L, 0L, 0L, 0L, 0L, 2L, 0L, 0L, 0L, 0L, 0L, 0L,
        0L, 0L, 0L, 0L, 0L, 0L, 0L, 1L, 0L, 0L, 0L, 0L, 0L, 0L, 0L, 0L,
        0L, 0L, 0L), .Dim = c(96L, 11L),
      .Dimnames =
        list(c("ACAA", "ACCA",
               "ACGA", "ACTA", "CCAA", "CCCA", "CCGA", "CCTA", "GCAA", "GCCA",
               "GCGA", "GCTA", "TCAA", "TCCA", "TCGA", "TCTA", "ACAG", "ACCG",
               "ACGG", "ACTG", "CCAG", "CCCG", "CCGG", "CCTG", "GCAG", "GCCG",
               "GCGG", "GCTG", "TCAG", "TCCG", "TCGG", "TCTG", "ACAT", "ACCT",
               "ACGT", "ACTT", "CCAT", "CCCT", "CCGT", "CCTT", "GCAT", "GCCT",
               "GCGT", "GCTT", "TCAT", "TCCT", "TCGT", "TCTT", "ATAA", "ATCA",
               "ATGA", "ATTA", "CTAA", "CTCA", "CTGA", "CTTA", "GTAA", "GTCA",
               "GTGA", "GTTA", "TTAA", "TTCA", "TTGA", "TTTA", "ATAC", "ATCC",
               "ATGC", "ATTC", "CTAC", "CTCC", "CTGC", "CTTC", "GTAC", "GTCC",
               "GTGC", "GTTC", "TTAC", "TTCC", "TTGC", "TTTC", "ATAG", "ATCG",
               "ATGG", "ATTG", "CTAG", "CTCG", "CTGG", "CTTG", "GTAG", "GTCG",
               "GTGG", "GTTG", "TTAG", "TTCG", "TTGG", "TTTG"),
             c("T68", "T41",
               "T74", "T16", "T46", "T82", "T50", "T15", "T24", "T80", "T95"
             )))

  short.analysis <-
    process.one.group(spectra=short.taiwan.hcc2,
                      sigs=liver.wes.sigs,
                      target.sig.name='Signature.AA',
                      path.root='mSigAct.basic.test.short.analysis',
                      obj.fun=obj.fun.nbinom.maxlh,
                      nbinom.size=5,
                      mc.cores=1)

  expected.short.pval <-
    structure(c(0.036297099364187, 0.76185665842143, 0.000540732195433939,
                0.999999809769497, 0.000672367936677276, 0.999997831038551, 0.120653696546247,
                0.996613255999323, 0.801907129656769, 1.68741386815852e-05, 1.46775631766299e-05
    ), .Names = c("T68", "T41", "T74", "T16", "T46", "T82", "T50",
                  "T15", "T24", "T80", "T95"))
  #cat(short.analysis$pval)
  stopifnot(all.equal(short.analysis$pval, expected.short.pval, tolerance = 0.005))

  expected.exp <-
    structure(c(10.5883215430212, 0, 43.0167421150595, 0, 0, 0, 0,
                6.39493634191927, 0, 0, 0, 0, 57, 0, 0, 0, 0, 0, 0, 0, 0, 0,
                0, 0, 0, 37.5493947700166, 0, 11.6064867320358, 7.84411849794756,
                0, 0, 0, 0, 0, 0, 37.8015117701765, 0, 0, 15.1984882298235, 0,
                0, 0, 41.1408335720938, 0, 0, 0, 0, 10.8591664279062, 0, 0, 0,
                0, 51, 0, 0, 0, 0, 0, 0, 0, 7.21731841509026, 14.5816564545762,
                0, 0, 0, 25.2010251303335, 0, 0, 0, 0, 0, 0, 45, 0, 0, 0, 0,
                0, 0, 0, 0, 0, 30.9066420035866, 13.0933579964134, 0, 0, 0, 0,
                0, 0, 0, 0, 0, 0, 0, 29.2344487210762, 0, 14.7655512789238, 0,
                0, 0, 0, 16.6275775292156, 0, 0, 0, 0, 12.3724224707844, 0, 0
    ), .Dim = 10:11,
    .Dimnames =
      list(c("Signature.1", "Signature.4",
             "Signature.5", "Signature.6", "Signature.12", "Signature.16",
             "Signature.17", "Signature.AA", "Signature.23", "Signature.24"
      ), c("T68", "T41", "T74", "T16", "T46", "T82", "T50", "T15",
           "T24", "T80", "T95")))

  stopifnot(all(all.equal(short.analysis$exposure, expected.exp, tolerance = 0.005)))

  # Create a spectrum that has only one signature, and a matrix of spectra that has only
  # on spectrum. These conditions often exercise errors.
  degenerate.spectrum <-
    matrix(c(round(liver.wes.sigs[ , 'Signature.AA'] * 1000, digits=0),
             round(liver.wes.sigs[ , 'Signature.AA'] * 500, digits=0)),
           ncol=2)

  colnames(degenerate.spectrum)  <-  c('Test1', 'Test2')
  rownames(degenerate.spectrum) <-  row.names(short.taiwan.hcc2)

  degenerate.analysis <-
    process.one.group(spectra=degenerate.spectrum,
                      sigs=liver.wes.sigs,
                      target.sig.name='Signature.AA',
                      path.root='mSigAct.basic.test.degenerate',
                      obj.fun=obj.fun.nbinom.maxlh,
                      nbinom.size=5,
                      mc.cores=1)

  degenerate.expected <-
    structure(list(pval = structure(c(2.59102885448936e-167, 3.85329163031964e-123
    ), .Names =
      c("Test1", "Test2")),

    exposure =
      structure(
        c(0.000000000000000000000000000167494431721009,
          0.0000000000000000000000000183002414718209,
          0.00000000000000000000000000000127078205831815,
          0.0000000000000000000000000000412941357417482,
          0.000000000000000000000000000013349661732206,
          0.000000000000000000000000107774231809069, 0, 998,
          0.0000000000000000000181192999334336,
          0.00000000000000000000000000016105377356955, 0, 0, 0, 0, 0, 0,
          0, 504, 0, 0), .Dim = c(10L, 2L),
        .Dimnames =
          list(c("Signature.1",
                 "Signature.4", "Signature.5", "Signature.6", "Signature.12",
                 "Signature.16", "Signature.17", "Signature.AA", "Signature.23",
                 "Signature.24"), c("Test1", "Test2")))),
    .Names = c("pval",
               "exposure"))

  stopifnot(all(all.equal(degenerate.analysis, degenerate.expected, tolerance =0.005)))

}
```
