## Supplemental Data S1 for "Identification of novel mutational signatures in Asian oral squamous cell carcinomas associated with bacterial infections": mSigTools.v0.13.R

```
#
# mSigTools
#
# v0.11
#
# An alpha version
#
# 2018 05 13
#
# Copyright 2017 by Alvin Wei Tian Ng, Steven G. Rozen
#
# The code is released under GPL-3
# https://www.gnu.org/licenses/gpl-3.0.en.html

# Dependencies
library(stringi)
library('lsa') # for cosine()

revc <- function(seq) {

  rq <- stri_reverse(seq)
  rq1 <- stri_trans_char(rq, 'ACGT', '1234')
  # We have to do this in two steps because
  # stri_trans_char computes something like the transitive
  # closure of the substitutions.
  stri_trans_char(rq1, '1234', 'TGCA')
}

xtract.col <- function(spec, sample.name.or.index) {
  tmp <- as.matrix(spec[ , sample.name.or.index])
  if (ncol(tmp) > 1) return(tmp)
  rownames(tmp) <- rownames(spec)
  colnames(tmp) <-
    ifelse(mode(sample.name.or.index) == 'numeric',
           colnames(spec)[sample.name.or.index],
           sample.name.or.index)
  tmp
}

######################################
# Funtions for reading catalogs, etc.
######################################

.canonical.96.row.order <-
  c("ACAA", "ACCA", "ACGA", "ACTA", "CCAA", "CCCA", "CCGA", "CCTA",
    "GCAA", "GCCA", "GCGA", "GCTA", "TCAA", "TCCA", "TCGA", "TCTA",
    "ACAG", "ACCG", "ACGG", "ACTG", "CCAG", "CCCG", "CCGG", "CCTG",
    "GCAG", "GCCG", "GCGG", "GCTG", "TCAG", "TCCG", "TCGG", "TCTG",
    "ACAT", "ACCT", "ACGT", "ACTT", "CCAT", "CCCT", "CCGT", "CCTT",
    "GCAT", "GCCT", "GCGT", "GCTT", "TCAT", "TCCT", "TCGT", "TCTT",
    "ATAA", "ATCA", "ATGA", "ATTA", "CTAA", "CTCA", "CTGA", "CTTA",
    "GTAA", "GTCA", "GTGA", "GTTA", "TTAA", "TTCA", "TTGA", "TTTA",
    "ATAC", "ATCC", "ATGC", "ATTC", "CTAC", "CTCC", "CTGC", "CTTC",
    "GTAC", "GTCC", "GTGC", "GTTC", "TTAC", "TTCC", "TTGC", "TTTC",
    "ATAG", "ATCG", "ATGG", "ATTG", "CTAG", "CTCG", "CTGG", "CTTG",
    "GTAG", "GTCG", "GTGG", "GTTG", "TTAG", "TTCG", "TTGG", "TTTG"
  )

.canonical.192.row.order <-
  
  c("AAAC","AACC","AAGC","AATC","CAAC","CACC","CAGC","CATC","GAAC",
    "GACC","GAGC","GATC","TAAC","TACC","TAGC","TATC","AAAG","AACG",
    "AAGG","AATG","CAAG","CACG","CAGG","CATG","GAAG","GACG","GAGG",
    "GATG","TAAG","TACG","TAGG","TATG","AAAT","AACT","AAGT","AATT",
    "CAAT","CACT","CAGT","CATT","GAAT","GACT","GAGT","GATT","TAAT",
    "TACT","TAGT","TATT","ACAA","ACCA","ACGA","ACTA","CCAA","CCCA",
    "CCGA","CCTA","GCAA","GCCA","GCGA","GCTA","TCAA","TCCA","TCGA",
    "TCTA","ACAG","ACCG","ACGG","ACTG","CCAG","CCCG","CCGG","CCTG",
    "GCAG","GCCG","GCGG","GCTG","TCAG","TCCG","TCGG","TCTG","ACAT",
    "ACCT","ACGT","ACTT","CCAT","CCCT","CCGT","CCTT","GCAT","GCCT",
    "GCGT","GCTT","TCAT","TCCT","TCGT","TCTT","AGAA","AGCA","AGGA",
    "AGTA","CGAA","CGCA","CGGA","CGTA","GGAA","GGCA","GGGA","GGTA",
    "TGAA","TGCA","TGGA","TGTA","AGAC","AGCC","AGGC","AGTC","CGAC",
    "CGCC","CGGC","CGTC","GGAC","GGCC","GGGC","GGTC","TGAC","TGCC",
    "TGGC","TGTC","AGAT","AGCT","AGGT","AGTT","CGAT","CGCT","CGGT",
    "CGTT","GGAT","GGCT","GGGT","GGTT","TGAT","TGCT","TGGT","TGTT",
    "ATAA","ATCA","ATGA","ATTA","CTAA","CTCA","CTGA","CTTA","GTAA",
    "GTCA","GTGA","GTTA","TTAA","TTCA","TTGA","TTTA","ATAC","ATCC",
    "ATGC","ATTC","CTAC","CTCC","CTGC","CTTC","GTAC","GTCC","GTGC",
    "GTTC","TTAC","TTCC","TTGC","TTTC","ATAG","ATCG","ATGG","ATTG",
    "CTAG","CTCG","CTGG","CTTG","GTAG","GTCG","GTGG","GTTG","TTAG",
    "TTCG","TTGG","TTTG"
    
  )

# Read 96-channel spectrum or signatures in Ludmil format

read.96.ludmil.format <- function(path) {
  cos <- read.table(path,
                    stringsAsFactors = F,
                    as.is = T,
                    header = T,
                    sep=',', 
                    check.names = F)
  stopifnot(nrow(cos)==96)
  ref.gt.var <- cos[ ,1]
  before.ref.after <- cos[ ,2]
  var <- substring(ref.gt.var, 3, 3)
  tmp <- paste(before.ref.after, var, sep='')
  rownames(cos) <- tmp
  out <- cos[ ,-(1:2), drop=F]
  out <- as.matrix(out)
  if (ncol(out) == 1) colnames(out) <- colnames(cos)[3]
  out[.canonical.96.row.order, drop=F,]
}

### Read 96-channel spectrum or signatures in Duke-NUS format
### This is a tab separated format with mutations information in
### the columns Before, Ref, After, Var

read.96.duke.nus.format <- function(path) {
  cos <- read.table(path,
                    stringsAsFactors = F,
                    as.is = T,
                    header = T,
                    sep='\t', 
                    check.names = F)
  stopifnot(nrow(cos)==96)
  # Not sure yet what will the form of the row labels
  # But Somatic.Mutation.Type has all the necessary information,
  # after which the first 3 columns are not needed
  tmp <- paste(cos$Before, cos$Ref, cos$After, cos$Var, sep='')
  rownames(cos) <- tmp
  out <- as.matrix(cos[ ,-(1:4), drop=F])
  out[.canonical.96.row.order, drop=F,]
}

### Read 192 channel spectra in Duke-NUS format
read.and.prep.192.duke.nus.catalog <- function(path) {
  df <- read.table(path,
                       stringsAsFactors = F,
                       as.is=T,
                       header=T, 
                   check.names = F)
  # Careful, df has 192 row
  stopifnot(nrow(df)==192)
  aa.must.complement <- df$Ref %in% c('A', 'G')

  xc <- df[aa.must.complement, ]
  rn1 <- revc(paste(xc$Before, xc$Ref, xc$After, sep = ''))
  rownames(xc) <- paste(rn1, revc(xc$Var), sep='')

  x1 <- df[!aa.must.complement, ]
  rownames(x1) <- paste(x1$Before, x1$Ref, x1$After, x1$Var, sep='')
  sort.string <- paste(x1$Ref, x1$Var, x1$Before, x1$After, sep = '')
  df.nocomp <- x1[order(sort.string), ]

  df.comp <- xc[rownames(df.nocomp), ]

  # For debugging
  # tmp.out <- cbind(df.comp[ ,1:4], df.nocomp[ , 1:4])

  # We no longer need the first 4 columns
  df.comp <- df.comp[ , -(1:4)]
  df.nocomp <- df.nocomp[ , -(1:4)]
  out <- df.comp + df.nocomp
  out <- as.matrix(out)
  out.order <- order(margin.table(out, 2), colnames(out), decreasing = T)
  out2 <- out[ , out.order]
  stopifnot(rownames(out2) == .canonical.96.row.order)
  list(channel.96=out2, channel.192=df[.canonical.192.row.order,])
}

# Add the four Duke-NUS format columns, Before, Ref, After, Var to the left of 
# matrix or data frame, i.e. splits the rownames and creates the four new
# columns, Before, Ref, After Var

duke.nus.rownames.to.cols <- function(mat.or.df) {
  newcols <- strsplit(rownames(mat.or.df), split='')
  newcols <- t(data.frame(newcols))
  colnames(newcols) <- c('Before', 'Ref', 'After', 'Var')
  rownames(newcols) <- rownames(mat.or.df)
  out <- cbind(newcols, data.frame(mat.or.df), stringsAsFactors=F)
  out
}

# works for spectrum or exposure matrices (columns are samples)
sort.spectra.columns <- function(spectrum) {
  if (ncol(spectrum) <= 1) return(spectrum) # ncol can be 0, or 1
  spect.total <- margin.table(spectrum, 2)
  spect.order <- order(spect.total, colnames(spectrum), method='radix', decreasing = c(T, F))
  spectrum[ , spect.order]
}
### new commands to read PCAWG SignatureAnalyzer and SignatureProfiler signatures and assigments
## function to read SA signatures and assignments 

read.sa.assignment <- function(sa.file){ ## reads in SignatureAnalyzer assignments from Synapse
  sa.assign <- read.table(sa.file, header = T)
  ## fix rownames so that it conforms to SBS** for each signature per row   
  row.names(sa.assign) <- gsub('_[PS]$', '', gsub('BI_SNV_', '', row.names(sa.assign)))
  
  ## remove signatures with 0 mutations attributed in all tumors  
  non.zero.sa <- rowSums(sa.assign)[rowSums(sa.assign) != 0]
  output <- as.matrix(sa.assign[names(non.zero.sa),])
  return(output) 
}

## function to read SA signatures
read.96.sa.signatures <- function(signature.file){
  sig.mat <- read.table(signature.file, header = T)
  stopifnot(nrow(sig.mat)==96)
  
  ## format signature names to SBS** format
  colnames(sig.mat) <- gsub('_[PS]$', '', 
                            gsub('BI.+SNV_', '', colnames(sig.mat)))
  
  ## format trinucleotide context features 
  var.base <- gsub('_at.+', '', 
                   gsub('\\w>', '', sig.mat[,'feature']))
  tri.context <- gsub('.+_', '', sig.mat[,'feature'])
  four.bases <- paste0(tri.context, var.base)
  
  ## out.sigs
  out.sigs <- sig.mat[, colnames(sig.mat) !='feature']
  rownames(out.sigs) <- four.bases
  sorted.sigs <- out.sigs[.canonical.96.row.order,]
  sorted.sigs
}

## function to read SP signatures and assignment 

read.192.sp.format <- function(path){
  cos <- read.table(path,
                    stringsAsFactors = F,
                    as.is = T,
                    header = T,
                    sep=',', 
                    check.names = F)
  
  stopifnot(nrow(cos)==192)
  ref.gt.var <- cos[ ,2]
  before.ref.after <- cos[ ,3]

  ## reverse complement 
  transcribed.strand.pos <- which(cos[,1] == 'T')
  
  before.ref.after[transcribed.strand.pos] <- revc(before.ref.after[transcribed.strand.pos])   
  var <- substring(ref.gt.var, 3, 3)
  var[transcribed.strand.pos] <- revc(var[transcribed.strand.pos])
  
  tmp <- paste(before.ref.after, var, sep='')
  rownames(cos) <- tmp
  out <- cos[ ,-(1:3), drop=F]
  out <- as.matrix(out)
  out[.canonical.192.row.order,, drop=F]
  ## need to set a standard format of output for 192
}

read.sp.assignment <- function(sp.file){ ## reads in SignatureProfiler assignments from Synapse
  sp.assign <- read.csv(sp.file, header = T)
  rownames(sp.assign) <- gsub('-', '.',
                              paste(sp.assign[,1],
                                    sp.assign[,2],
                                    sep = '..')
  )
  colnames(sp.assign) <- gsub('Signature.Subs.[0]{0,1}',
                              'SBS',
                              colnames(sp.assign))
  ## remove sample / tumor names and accuracy column
  sp.assign.cleaned <- t(sp.assign[,4:ncol(sp.assign)])
  
  non.zero.sa <- rowSums(sp.assign.cleaned)[rowSums(sp.assign.cleaned) != 0]
  output <- as.matrix(sp.assign.cleaned[names(non.zero.sa),])
}


######################################
# Funtions for plotting, etc.
######################################

pdf.mut.sig.profile <- function(path, spec.or.sig, show.counts=F) {
  spectrum.plot.pdf.setup(path)
  t.plot.spectra(duke.nus.rownames.to.cols(spec.or.sig),
                 show.counts=show.counts,
                 show.x.labels = T)
  dev.off()
}

### plot.ex.by.range
###
### Plot exposures broken up by ranges
plot.ex.by.range <- function(spectrum,
                             sigs,
                             exp,
                             ranges, # A list of vectors
                             col=NULL,
                             main=NULL,
                             xlab=NULL,
                             ylab=NULL,
                             mfrow=c(3,1) # 3 exposure graphs per page by default
) {

  par(
    mfrow=mfrow,
    # mar  =c(1.5,1,2,2), # space between plot and figure, for axis labels etc
    oma  =rep(3, 4) # outer margin between figure and device.
  )
  if (is.null(ylab)) ylab <- 'Number of mutations'
  first=T
  for (range in ranges) {
    if (first) {
      plot.exposures(exp[ ,range], signatures=sigs,
                     input.genomes = spectrum[ , range],
                     plot.proprtion = F,
                     col=col,
                     main=main,
                     ylab=ylab, xlab=xlab)
      first=F
    } else {
      plot.exposures(exp[ ,range], signatures=sigs, 
                     input.genomes = spectrum[ , range],
                     plot.proprtion = F, plot.legend=F,
                     main=main,
                     col=col,
                     ylab=ylab, xlab=xlab)
    }
  }
}


pdf.ex.by.range <- function(path,   # Out file path
                             spectrum,
                             sigs,
                             exp,
                             ranges, # A list of vectors
                             col=NULL,
                             main=NULL,
                             xlab=NULL,
                             ylab=NULL
) {
  pdf(path, width=8.2677, height=11.6929, # for A4
      onefile=T, useDingbats=F)
  plot.ex.by.range(spectrum, sigs, exp, ranges, col, main, xlab, ylab)
  dev.off()
}


### Utilities for managing different underlying trinucleotide frequencies

read.opportunity <- function(path) {
  op <- read.table(path, as.is=T, stringsAsFactors = F, header=T, sep='\t')

  # Fold the two strands to triplets centered on pyrimidines (C or T)
  purine <- grep('.[AG].', op$triplet, perl=T)
  op.pur <- op[purine, ]
  op.pyr <- op[-purine, ]
  new.names <- revc(op.pur$triplet)
  op.pur$triplet <- new.names
  op.pur <- op.pur[order(op.pur$triplet), ]
  op.pyr <- op.pyr[order(op.pyr$triplet), ]
  op.pur.oc <- op.pur$occurrences
  op.pyr.oc <- op.pyr$occurrences
  stopifnot(all(op.pur$triplet == op.pyr$triplet))
  tmp.occurrences <- c(op.pyr.oc + op.pur.oc)

  tmp <- data.frame(triplet=op.pur$triplet, occurrences=tmp.occurrences)
  rownames(tmp) <- tmp$triplet

  # Cacluate proportions of each triplet
  tmp$prop <- tmp$occurrences / sum(tmp$occurrences)
  tmp
}

.h19.96.WGS.op           <-
  read.opportunity('opportunity/triplet_counts-genome-hg19.tsv')
.h19.96.sureselect.v2.op <-
  read.opportunity("opportunity/triplet_counts-SureselectV2-hg19.tsv")
.h19.96.sureselect.v6.op <-
  read.opportunity("opportunity/triplet_counts-SureselectV6-hg19.tsv")
.mm10.96.WES.op <- read.opportunity('opportunity/triplet_counts-CCDS_exons-mm10.tsv')
.flat.96.op              <- .h19.96.WGS.op
.flat.96.op[ , 'prop']   <- 1/32
.flat.96.op[ , 'occurrences'] <-
  sum(.h19.96.WGS.op[ , 'occurrences']) / 32 # not sure if we need this

transform.96.sig.op1.op2 <- function(input.sig.mat, in.op, out.op) {
  out.sig.mat <- input.sig.mat

  stopifnot(rownames(out.op) == rownames(in.op))
  factor <- out.op$prop / in.op$prop
  names(factor) <- rownames(out.op)

  transform.one.triplet <- function(triplet) {
    rows <- grep(paste("^", triplet, sep=''), rownames(out.sig.mat))
    out.sig.mat[rows, ] <<- out.sig.mat[rows, ] * factor[triplet]
  }

  lapply(rownames(out.op), transform.one.triplet)
  out2 <- apply(out.sig.mat, MARGIN = 2, function (x) x / sum(x))
  # Each colmun in out2 sums to 1

  # lazy way to get new matrix in same shape as out2
  # out3's elements will be overwritten
  out3 <- out2

  for (i in 1:ncol(out2)) {
    out3[ ,i] <- out2[ ,i] * sum(input.sig.mat[ , i])
  }
  out3
}

### Get mutational signatures specified as proportions based on
### human WGS (whole genome sequence) count in file 
### signature.file.
###
### Return a list of three signature matrices: WGS, WES, and "flat".
###
### IMPORTANT: this replaces get.COSMIC.signatures and now requires the
### name of the signature file.
get.signatures <- function(exome.op, signature.file, debug=F) {

  # Read the mutational signatures in genome frequencies

  if (grepl('*csv', signature.file) ){
    cosmic <- read.96.ludmil.format(signature.file)
  } else{ 
    ## support signatureanalyzer signatures
    if (grepl('signatureana', tolower(signature.file))){
      cosmic <- read.96.sa.signatures(signature.file = signature.file)
    } else{
      cosmic <- read.96.duke.nus.format(signature.file)    
    } 
    
  }
  
  ## normalize signatures such that each colum sums to 1
  cosmic <- apply(cosmic, MARGIN = 2, function (x) x / sum(x))
  
  wgs.op <- .h19.96.WGS.op
  wes.op <- exome.op

  # For debugging / testing
  wes.and.wgs.op <-cbind(wes.op, wgs.op)

  wes.to.wgs.factor <- wes.op$prop / wgs.op$prop
  names(wes.to.wgs.factor) <- rownames(wes.op)

  transform.cosmic.to.wes <- function(input.sig.mat) {
    out.sig.mat <- input.sig.mat

    transform.one.triplet <- function(triplet) {
      rows <- grep(paste("^", triplet, sep=''), rownames(out.sig.mat))
      out.sig.mat[rows, ] <<- out.sig.mat[rows, ] * wes.to.wgs.factor[triplet]
    }

    lapply(rownames(wes.op), transform.one.triplet)
    out2 <- apply(out.sig.mat, MARGIN = 2, function (x) x / sum(x))
    out2
  }

  cosmic.wes <- transform.cosmic.to.wes(cosmic)

  genome.to.prop.adj.factors <- (1/32) / wgs.op$prop
  names(genome.to.prop.adj.factors) <- rownames(wgs.op)

  transform.genome.to.prop <- function(input.sig.mat, debug=F) {
    out.sig.mat <- input.sig.mat

    transform.one.triplet <- function(triplet) {
      rows <- grep(paste("^", triplet, sep=''), rownames(out.sig.mat))
      out.sig.mat[rows, ] <<- out.sig.mat[rows, ] * genome.to.prop.adj.factors[triplet]
    }

    if (debug) debug(transform.one.triplet)
    lapply(rownames(wgs.op), transform.one.triplet)
    out2 <- apply(out.sig.mat, MARGIN = 2, function (x) x / sum(x))
    out2
  }

  per.triplet.prop <- transform.genome.to.prop(cosmic, debug=F)

  if (debug) {
    pdf.mut.sig.profile('my.exon.spectra2.pdf', cosmic.wes)
    pdf.mut.sig.profile('genome.spectra2.pdf', cosmic)
    pdf.mut.sig.profile('per.triplet.spectra2.pdf', per.triplet.prop)
  }

  list(genome=as.matrix(cosmic), exome=cosmic.wes, flat=per.triplet.prop)

}

# Global variables used by these functions. (prefixed with '.')
# similar to original sanger colours, but better when printed.

# For 96 channels, 16 bars per major class, 6 major classes:
.bar.colours = c('C>A'='#4040FF',
                 'C>G'='#000000',
                 'C>T'='#FF4040',
                 'T>A'='#838383',
                 'T>C'='#40FF40',
                 'T>G'='#ff667f')

.bases = c('A', 'C', 'G', 'T') # order affects the order they are plotted in
# (reverse) complement bases
.pair = list(A="T", C="G", G="C", T="A", # for complementary base
             # for 1536 class plots (ie +-2 base context):
             AA='TT', AC='GT', AG='CT', AT='AT',
             CA='TG', CC='GG', CG='CG', CT='AG',
             GA='TC', GC='GC', GG='CC', GT='AC',
             TA='TA', TC='GA', TG='CA', TT='AA')

get_ylim = function(y) {
  if (y > 300) y = 200 * ceiling(y/200)   # round to nearest 200
  else if (y > 150) y = 100 * ceiling(y/100)   # round to nearest 100
  else if (y > 50) y = 30 * ceiling(y/30) # nearest 30
  else y = 10 * ceiling(y/10)             # nearest 10
  return(y)
}

t.plot.spectra = function(counts.data.frame,
                        show.counts=T,
                        show.class.names=F,
                        all.labels=F,
                        show.x.labels=F,
                        trace=F,
                        show.paren=F) {
  region <- 'genome'
  
  num.classes <- 96

  base.frequencies.triplets <- flat.64.opportunity()

  spectrum.names <- colnames(counts.data.frame) # first 4 columns are the bases/variant info

  context.before <- .bases
  context.after  <- .bases

  ref.bases = c('C', 'T') # pyrimidines - we don't care about strand
  var.bases = .bases

  first.col <- 5

  for (which.col in c(first.col:length(spectrum.names))) {
    spectrum.name = spectrum.names[which.col]
    display.name = spectrum.name
    if (trace) cat('\rplotting', display.name, '\n')

    # get counts per (unstranded) mutation class (6 classes)
    Ref.A = counts.data.frame$Ref=='A'
    Ref.C = counts.data.frame$Ref=='C'
    Ref.G = counts.data.frame$Ref=='G'
    Ref.T = counts.data.frame$Ref=='T'
    Var.A = counts.data.frame$Var=='A'
    Var.C = counts.data.frame$Var=='C'
    Var.G = counts.data.frame$Var=='G'
    Var.T = counts.data.frame$Var=='T'
    unstranded.counts = c(
      "C>A"=sum(counts.data.frame[(Ref.C & Var.A)|(Ref.G & Var.T), which.col]),
      "C>G"=sum(counts.data.frame[(Ref.C & Var.G)|(Ref.G & Var.C), which.col]),
      "C>T"=sum(counts.data.frame[(Ref.C & Var.T)|(Ref.G & Var.A), which.col]),
      "T>A"=sum(counts.data.frame[(Ref.A & Var.T)|(Ref.T & Var.A), which.col]),
      "T>C"=sum(counts.data.frame[(Ref.A & Var.G)|(Ref.T & Var.C), which.col]),
      "T>G"=sum(counts.data.frame[(Ref.A & Var.C)|(Ref.T & Var.G), which.col])
    )

     
    if (trace) print(unstranded.counts)
    percents = rep(0, num.classes)
    maj.class.counter = 0 # major class. Ie B₁>B₂
    maj.class.names = c() # only those (eg C>A) present in input data
    
    contexts.used = c() # debug
    for (ref in ref.bases) {
      for (variant in var.bases) {
        if (variant == ref) next
        maj.class.counter = maj.class.counter + 1 # 6 major classes
        maj.class.names[maj.class.counter] = sprintf('%s>%s', ref, variant)
        counter = 0 # before & after combination - 16 combos (for 96 classes)
        # 32 combos if we have 192 classes and want the main plot to show
        # strand counts (instead of in a separate plot)
        for (before in context.before)
          for (after in context.after) {
            counter = counter + 1
            count.sense =
              counts.data.frame[counts.data.frame$Ref==ref & counts.data.frame$Var==variant & counts.data.frame$Before==before & counts.data.frame$After==after, which.col]
            # reverse complement, for other strand
            count.antisense =
              counts.data.frame[counts.data.frame$Ref==.pair[ref] & counts.data.frame$Var==.pair[variant] & counts.data.frame$Before==.pair[after] & counts.data.frame$After==.pair[before], which.col]
            if (length(count.sense)==0) count.sense = 0
            if (length(count.antisense)==0) count.antisense = 0
            # correct for the opportunity
            weighting.sense =
              base.frequencies.triplets[sprintf("%s%s%s",before,ref,after), 1]
            weighting.antisense =
              base.frequencies.triplets[sprintf("%s%s%s",.pair[after],.pair[ref],.pair[before]), 1]
            class.offset = (maj.class.counter-1) * 16
            contexts.used[class.offset+counter] = 
              paste(before, ref, variant, after, sep='')
            # we combine the weightings for both strands instead of taking
            # individual triplets into account, so that it's easier to compare
            # whole genome data...
            percents[class.offset+counter] <-
              (count.sense + count.antisense)/(weighting.sense+weighting.antisense)
          }
      } # var base
    } # ref base

    if (sum(percents)==0) {
      plot(NA, xlim=c(0,1), ylim=c(0,1), main=display.name, xaxt='n', xlab='',
           yaxt='n', ylab='proportion', type='h', bty='n')
      text(.5, .5, 'no data')
      # we are showing another figure for each spectrum, so leave it blank
      if (region != 'genome') plot.new()
      next
    }

    percents = percents/sum(percents) # after applying the opportunity weighting

    values.per.class <- 16 # ie 6 major classes

    # 'cols.per.class' is used for the coloured bars at the top.
    # get colours only for classes present in input data
    # do this way to handle if we
    # aren't showing all 6 major classes. (eg 256 classes for B1>B2 +-2)
    cols.per.class = .bar.colours[maj.class.names]
    # get colours for each bar
    cols <- rep(cols.per.class, each=values.per.class)

    main.extra <- ifelse(show.counts,
                         paste(' (', sum(unstranded.counts), ')', sep=''),
                         '')

    plot(percents,
         main=paste(display.name, main.extra, sep=''),
         xaxt='n',
         xlim=c(0, length(percents)),
         xlab='',
         yaxt='n',
         ylab='proportion',
         type='h',
         lwd=3,
         bty='n',
         col=cols, xpd=NA)

    # min, max, # ticks calculated automatically:
    y.axis.values = par('yaxp')
    axis(side=2, at=c(0,y.axis.values[2]), las=1) # only show max value on axis
    # mutation class labels at the top of the figure:
    yb = par('usr')[4]
    # for A4 portrait: move left a bit if we are showing the mutation count

    if (show.class.names) {
      mtext(maj.class.names, side=3,
            at=1:6*values.per.class-(values.per.class/2),
            line=.3, cex=.8)
    }

    yt = yb*1.03

    # don't try to print counts if our input files has proportions instead.
    # just silently ignore.
    if (show.counts && any(unstranded.counts>=1)) {
      # if floats (eg reconstructed or Févotte output), round to ints
      # only keep major classes present in input file (eg for 256 class [+-2])
      unstranded.counts = round(unstranded.counts)[maj.class.names]
      # add the mutation count at the top of each figure, in smaller text
      num.format <- ifelse(show.paren, "(%d)", "%d")
      mtext(sprintf(num.format, unstranded.counts), side=3,
            at=1:6*values.per.class - 1, line=.3, cex=.6)
    }

    if (show.class.names) {
      # draw lines above each of the main 6 mutation types:
      n_classes = length(cols.per.class) # normally 6, maybe 1 (if 256)
      rect(xleft=0:(n_classes-1)*values.per.class+1, yb,
           xright=1:n_classes*values.per.class, yt,
           col=cols.per.class, border=NA, xpd=NA)
    }

    # draw the 5' and 3' base labels?
    if (is.na(all.labels)) {
      draw.labels=F
    } else {
      if (all.labels) {
        draw.labels = T
      } else if (all.labels==F) { # only draw for last figure on page
        num.rows = par('mfrow')[1] # (rows, cols)
        if (num.rows > 1) { # using layout/par('mfrow') plotting method
          # check if current spectrum is a multiple of num.rows
          draw.labels = (which.col-first.col) %% num.rows == num.rows-1
          # always draw for last spectrum
          if (which.col == length(spectrum.names)) draw.labels = T
        } else { # using grid() instead of mfrow or layout()?
          # TODO. ???
        }
      }
    }
    
    if (show.x.labels && draw.labels) {
      # context base labels:
      sz=.4
      y.step = .15
      mtext("preceded by 5'", side=1, at=-.5, line=0, cex=.5, adj=1)
      # 0,4,8,12 for 16 classes; 0,8,16,24 for 32 (bars for both strands)
      offsets = 0:3 * (values.per.class/4)
      # only used if > 1 major class. (& our first bar is at 1, not 0)
      start.of.class = (0:5)*values.per.class +1 # major classes
      mtext('A',side=1, at=start.of.class+offsets[1], line=-.1, adj=.5, cex=sz)
      mtext('C',side=1, at=start.of.class+offsets[2], line=-.1, adj=.5, cex=sz)
      mtext('G',side=1, at=start.of.class+offsets[3], line=-.1, adj=.5, cex=sz)
      mtext('T',side=1, at=start.of.class+offsets[4], line=-.1, adj=.5, cex=sz)
      
      # note: at -.7 because it doesn't line up with 5' !?
      mtext("followed by 3'", side=1, at=-.7, line=.65, cex=.5, adj=1)
      
      labels.5p = (0:23)*4 +1 # where we have each 5' label
      bars.per.3p = 1
      mtext('A', side=1, at=labels.5p+0*bars.per.3p, line=.4, adj=.5, cex=sz)
      mtext('C', side=1, at=labels.5p+1*bars.per.3p, line=.4+y.step, adj=.5, cex=sz)
      mtext('G', side=1, at=labels.5p+2*bars.per.3p, line=.4+2*y.step, adj=.5, cex=sz)
      mtext('T', side=1, at=labels.5p+3*bars.per.3p, line=.4+3*y.step, adj=.5, cex=sz)

    }
    
  } # end for each spectrum
  if (trace) cat('\n')
}

plot.exposures <- 
  function(s.weights, # This is actually the exposure "counts"
           # (or floats approximating the exposure counts)
           signames=NULL,
           scale.num=NULL,
           signatures=NULL,
           input.genomes=NULL,
           plot.proprtion=T,
           plot.legend=T,
           ylim=NULL,
           main=NULL,
           ylab=NULL,
           xlab=NULL,
           col=NULL
  ) {
    
    # note - might be reals > 1, not necessary colSum==1
    s.weights <- as.matrix(s.weights) # in case it is a data frame
    signature_proportions <- t(s.weights)
    num.sigs = dim(s.weights)[1]
    num.samples = dim(s.weights)[2]
    
    if (is.null(col)) {
      if (num.sigs <= 8) {
        col = # c('skyblue', 'black', 'grey', 'yellow', 'blue', 'brown', 'green4', 'red')
          c('red', 'black', 'grey', 'yellow', 'blue', 'brown', 'green4', 'skyblue')
        
      } else {
        # lots of signatures; use shaded lines to differentiate
        col = rainbow(num.sigs)
      }
    }
    if (num.sigs <= 12) {
      p.dense = -1 # will repeat as needed, -1 = solid
      p.angle = 0  # ditto
    } else {
      # lots of signatures; use shaded lines to differentiate
      p.dense = c(-1,35,35,50,50) # will repeat as needed, -1 = solid
      p.angle = c(0,0,135,45,135)  # ditto
    }
    # for legend, we need to put in reverse order. make sure this matches
    # order used in barplot
    num.repeats = ceiling(num.sigs/length(p.dense))
    p.dense.rev = rev(rep(p.dense,num.repeats)[1:num.sigs])
    p.angle.rev = rev(rep(p.angle,num.repeats)[1:num.sigs])
    
    # add names (if not already set as row.names in the original input frame)
    # for sorting. (needs a "Sample" column in the signature_proportions frame)
    # if (length(colnames(s.weights))==0) colnames(s.weights) = signature_proportions$Sample
    if (is.null(scale.num)){
      ylabel = '# mutations'
    } else { # show as rate instead of count
      if (scale.num < 3000 || scale.num > 3300) {
        warning('assuming "scale.num" should be divisor for human genome. using 3000')
        scale.num = 3000
      }
      s.weights = s.weights/scale.num
      ylabel ='# mutations/Mbase'
    }
    l.cex = if (num.sigs > 15) .5 else 1 # char expansion for legend (was 0.7)
    direction = 2 # 1=always horizontal, 2=perpendicular to axis
    
    # if we weights file and counts file have samples in different order
    if (!all(colnames(s.weights)==colnames(input.genomes))) {
      input.genomes = input.genomes[,colnames(s.weights)]
      warning('weights file and counts file are ordered differently; re-ordering counts.')
    }
    
    # ignore column names; we'll plot them separately to make them fit
    bp = barplot(s.weights,
                 ylim=ylim,
                 las=1,
                 col=col,
                 ylab=ylab,
                 yaxt='s',
                 xaxt='n',
                 xlab=xlab,
                 density=p.dense, angle=p.angle,
                 border=ifelse(num.samples>200,NA,1),
                 main=main, cex.main=1.2)
    
    # get max y values for plot region, put legend at top right
    dims = par('usr') # c(x.min, x.max, y.min, y.max)
    y.max = dims[4]
    
    if (plot.legend) {
      # less space between rows (y.intersp), and between box & label (x.intersp)
      # reverse the order, so sig 1 is at bottom (to match bargraph)
      legend.x <- ncol(s.weights) * .7
      legend.y <- y.max * 0.8
      legend(x=legend.x, y=legend.y,
             rev(row.names(s.weights)),
             density=p.dense.rev, angle=p.angle.rev,
             bg=NA, xpd=NA,
             fill=col[num.sigs:1],
             x.intersp=.4, y.intersp=.8,
             bty='n', cex=l.cex * 0.9)
      text(x=legend.x, y = legend.y, "Mutational signature", adj=-0.09)
    }
    
    # now add sample names, rotated to hopefully fit
    # don't even try to show all if there are too many
    if (num.samples <= 200) {
      if (length(bp)<50) size.adj = .75
      else if (length(bp)<80) size.adj = .65
      else if (length(bp)<100) size.adj = 0.4 # .5
      else if (length(bp)<120) size.adj = .4
      else if (length(bp)<150) size.adj = .3
      else size.adj = .3
      mtext(colnames(s.weights), side=1, at=bp, las=direction, cex=size.adj)
    }
    
    if (plot.proprtion) {
      # add proportion panel; eg col should sum() to 1. matrix divided by
      # a vector goes col-wise, not row-wise, so transpose twice :(
      bp = barplot(t(t(s.weights)/colSums(s.weights)), las=1, col=col, ylab='Proportion',
                   density=p.dense, angle=p.angle.rev,
                   main='', axisnames=F, border=NA)
    }
    
  }

plot.reconstruction <-
  function(signatures, # Mutation classes x signatures
           exposures.mat, # Signatures x samples
           input.genomes, # Mutation classes x samples
           normalize.recon, # Normalize to the number of mutations in the spectrum
           padding=0          # How many samples wide the plot should be. (we'll add empty
  ) {
    
    num.sigs = ncol(signatures)
    num.samples = ncol(input.genomes)
    recon = signatures %*% exposures.mat
    # original matrix with mutation class counts per sample
    mat = input.genomes # as.matrix(input.genomes)
    
    # Calculate reconstruction errors
    cos.sim = sapply(1:num.samples, function(i) cosine(mat[,i], recon[,i]) )
    pearson.cor = sapply(1:num.samples, function(i) cor(mat[,i], recon[,i], method='pearson') )
    
    # Euclidian distance
    Eu.dist = apply(mat - recon, 2, function(x) sqrt(sum((x)^2)) ) # raw dist
    
    # KL divergence:
    norm.mat = mat/colSums(mat) # normalized by total number of mutations
    norm.recon = recon/sum(recon)
    kl = vector('numeric', length=num.samples)
    for (i in 1:num.samples) { # each column of sample/recon. matrix
      zero = norm.mat[,i] == 0 | norm.recon[,i] == 0 # use only non-zero rows
      if (all(zero)) { # shouldn't happen...
        kl[i] = 0
      } else {
        kl[i] = sum( norm.mat[!zero,i] *
                       log(norm.mat[!zero,i]/norm.recon[!zero,i]) )
      }
    }
    
    if (normalize.recon) {
      Eu.dist <- Eu.dist / colSums(mat)
      # kl <- kl / colSums(mat)
    }
    
    if (padding && ncol(mat) < padding) {
      # add empty columns so that reconstruction error plots have the
      # samples lined up with the exposure/proportion plots. (Eg if plotting
      # 50 samples at a time, and last figure only has 40 samples.)
      NAs = rep(NA, padding-dims[2])
      cos.sim = c(cos.sim, NAs)
      pearson.cor = c(pearson.cor, NAs)
      Eu.dist = c(Eu.dist, NAs)
      kl = c(kl, NAs)
    }
    # don't need so much top/bottom margin space for these 2 plots
    # new.mar = par('mar'); new.mar[1] = 2; new.mar[3] = 2
    # old.pars = par(mar=new.mar)
    
    # plot the cosine and pearson on the same figure, since they are both 0<=x<=1
    
    y.low <- min(c(cos.sim, pearson.cor)) - 0.1
    
    if (is.na(y.low)) {y.low = 0}
    
    plot(cos.sim, type='p', pch=16, col='grey50', main='Reconstruction error',
         ylim=c(y.low, 1),
         xlab='',
         xaxt='n',
         ylab='', bty='n',
         las=2, new=T)
    points(pearson.cor, type='p', pch=1, col='black') # same scale as cos.sim
    axis(side=1, # below
         labels=colnames(input.genomes),
         at=1:length(cos.sim),
         las=2)
    abline(v=1:length(cos.sim), lty=3)
    legend(0.1 * length(cos.sim), y.low + (1 - y.low) * .25,
           legend=c('Cosine similarity',"Pearson corr."),
           col=c('grey50','black'), pch=c(16,1), xpd=NA, bty='n')
    
    
    if (normalize.recon) {
      Eu.label = 'Euclidean dist / number of mutations'
      # kl.label = 'KL divergence / number of mutations'
    } else {
      Eu.label = 'Euclidean dist'
      # kl.label = 'KL divergence'
    }
    kl.label = 'KL divergence'
    # now plot kl and Euclidean on another graph.
    kl.max = max(na.omit(kl))
    plot(kl, type='p', pch=16, col='blue', ylim=c(0,kl.max), xlab='', ylab='', axes=F,
         new=T)
    axis(2, ylim=c(0, kl.max), col.axis='blue', las=1)
    par(new=TRUE) # allow a second plot on the same figure
    eu.max = max(na.omit(Eu.dist))
    plot(Eu.dist, type='p', pch=16, col='red',  xlab="", ylab="", ylim=c(0,eu.max), axes=F)
    axis(side=1, # below
         labels=colnames(input.genomes),
         at=1:length(Eu.dist),
         las=2)
    abline(v=1:length(Eu.dist), lty=3)
    legend(0.1 * length(kl), 0.25 * eu.max,legend=c(Eu.label, kl.label),
           col=c('red', 'blue'), pch=16, xpd=NA, bty='n')
    ## add text to second y axis
    
    axis(4, ylim=c(0, eu.max), col.axis='red', las=1)
    
  }


flat.64.opportunity = function() {
  # don't weight - set all to 1
  base.frequencies.triplets = data.frame(occurrences=rep(1/64, 64))
  i = 1
  for (b1 in .bases) for (b2 in .bases) for (b3 in .bases) {
    rownames(base.frequencies.triplets)[i] = paste(b1,b2,b3,sep='')
    i = i + 1
  }
  return(base.frequencies.triplets)
}

spectrum.plot.pdf.setup <- function(path) {
  pdf(path, width=8.2677, height=11.6929, onefile=T, useDingbats = F) # for A4
  par(
    mfrow=c(8,1), # 8 signatures per page
    mar=c(1.5,1.1,4.6,1), # space between plot and figure, for axis labels etc
    oma=c(2,4.1,1,1) # outer margin between figure and device.
  )
}

### Example call
### 
### plot.one.exome('test.exome.pdf',
###                exome.100[ , 1, drop=F],
###                 exome.op=.h19.96.sureselect.v6.op)
plot.one.exome <- function(path, spec, exome.op, show.class.names=F) {
  save.name <- colnames(spec)

  spectrum.plot.pdf.setup(path)

  colnames(spec) <- paste(save.name, 'exome', sep='-')
  t.plot.spectra(duke.nus.rownames.to.cols(spec),
                 show.class.names=show.class.names,
                 show.counts=T)

  wgs.spec <- transform.96.sig.op1.op2(input.sig.mat = spec,
                                       in.op=exome.op,
                                       out.op = .h19.96.WGS.op)
  colnames(wgs.spec) <- paste(save.name, 'genome', sep='-')
  t.plot.spectra(duke.nus.rownames.to.cols(wgs.spec), show.counts=F)

  flat.spec <- transform.96.sig.op1.op2(input.sig.mat = spec,
                                        in.op=exome.op,
                                        out.op = .flat.96.op)
  colnames(flat.spec) <- paste(save.name, 'flat', sep='-')
  t.plot.spectra(duke.nus.rownames.to.cols(flat.spec), show.counts=F,
                 show.x.labels = T)

  dev.off()
}

### Example call
### 
### plot.one.genome('test.genome.pdf', 
###                  genome.100[ , 1, drop=F], 
###                  exome.op=.h19.96.sureselect.v6.op)
plot.one.genome <- function(path, spec, exome.op) {
  save.name <- colnames(spec)

  spectrum.plot.pdf.setup(path)
  colnames(spec) <- paste(save.name, 'genome', sep='-')
  t.plot.spectra(duke.nus.rownames.to.cols(spec), show.counts=T)

  flat.spec <- transform.96.sig.op1.op2(input.sig.mat = spec,
                                       out.op= .flat.96.op,
                                       in.op = .h19.96.WGS.op)
  colnames(flat.spec) <- paste(save.name, 'flat', sep='-')
  t.plot.spectra(duke.nus.rownames.to.cols(flat.spec),
                 show.counts=F,
                 show.x.labels = T)

  dev.off()

  }

# FIX ME, MOVE THIS TO mSigActTesR
test.sig.spec.transform <- function(in.wgs.spec) {

  flat.spec <-
    transform.96.sig.op1.op2(input.sig.mat = in.wgs.spec,
                             in.op = .h19.96.WGS.op, out.op = .flat.96.op)
  new.wgs.spec <-
    transform.96.sig.op1.op2(input.sig.mat = flat.spec,
                             in.op = .flat.96.op, out.op=.h19.96.WGS.op)

  stopifnot(abs(new.wgs.spec - in.wgs.spec) < 1e-12)

  wes.from.flat.spec <-
    transform.96.sig.op1.op2(input.sig.mat = flat.spec,
                             in.op = .flat.96.op,
                             out.op = .h19.96.sureselect.v2.op)

  flat.from.wes.spec <-
    transform.96.sig.op1.op2(input.sig.mat = wes.from.flat.spec,
                             out.op = .flat.96.op,
                             in.op = .h19.96.sureselect.v2.op)

  stopifnot(abs(flat.spec - flat.from.wes.spec) < 1e-12)

  wgs.from.wes.spec <-
    transform.96.sig.op1.op2(input.sig.mat = wes.from.flat.spec,
                             out.op = .h19.96.WGS.op,
                             in.op = .h19.96.sureselect.v2.op)

  stopifnot(abs(wgs.from.wes.spec - in.wgs.spec) < 1e-12)

  wes.from.wgs.spec <-
    transform.96.sig.op1.op2(input.sig.mat = in.wgs.spec,
                             in.op = .h19.96.WGS.op,
                             out.op = .h19.96.sureselect.v2.op)

  stopifnot(abs(wes.from.wgs.spec - wes.from.flat.spec) < 1e-12)
}

### code to generate a matrix of counts using a matrix of exposure and a matrix of signatures 
### can be used with generate.synthetic.exposures to generate counts for synthetic data
convert.exp.to.mat <- function(exposure.matrix, ## matrix of exposures
                               signature.matrix ## matrix of signatures to multiply exposures to get counts
){
  ## signatures are loaded before calling the function so that any signature
  ## matrix of any dimension can be used
  ## get.signatures can be used to load signatures 
  
  ## check if dimensions of matrix are the same 
  stopifnot(ncol(signature.matrix) == nrow(exposure.matrix))
  round(signature.matrix %*% exposure.matrix)
}


## add negative binomial noise on a matrix of counts, with columns being each sample 
add.neg.binom.noise <- function(counts.mat,       ## matrix of counts without noise added
                                proportion = 0.2, ## proportion of total mutations per sample to add as noise
                                dispersion = 10){ ## dispersion parameter for negative binom model
  

  new.counts.mat <- apply(counts.mat, 2, function(x){
    
    noise <- rnbinom(length(x), 
                     dispersion, 
                     mu = rep(1, length(x))/length(x) * proportion * sum(x)) ## equal probability of noise in each class 
    x+noise
  }) 
}
```
