## Supplemental Data S1 for "Identification of novel mutational signatures in Asian oral squamous cell carcinomas associated with bacterial infections": Sup.Data1.html

### Supplemental Data 1 for Boot et al., “Mutational signature analysis of Asian OSCCs reveals novel mutational signature with exceptional sequence context specificity”

These are the scripts used to search for the presence of the novel mutational signature (SBS\_A^n^T) in the data set of 19,184 whole-exomes and 4,645 whole-genomes that was compiled for the PCAWG on Mutational Signatures (Alexandrov et al., 2018).

#### Background

SBS\_A^n^T is characterized by extremely strong enrichment for adenines 4bp and 3bp 5’ of mutated thymines.

We will use this characteristic to search for this mutational signature in the whole-exome and whole-genome data.

To test for enrichment we perform binomial tests, comparing: a) the proportion of T>N mutations with an adenine 3 or 4bp 5’ against b) the proportion thymines in the human genome or exome that has an adenine 3 or 4bp 5’

The null hypothesis is that the proportion of T>N mutations with A at -3 (or -4) is equal to the proportion of thymines in the human genome/exome with A at -3 (or -4)

#### Methods / code

We retrieved the variant calls for 19,184 whole-exomes and 4,645 whole-genomes, and counted:

- T2Ncount: Sum of the number of T>A, T>C and T>G single nucleotide substitutions
- ANNTcount: Total number of T>A, T>C and T>G mutations where there is an adenine 3bp 5’ of the mutated T
- ANNNTcount: Total number of T>A, T>C and T>G mutations where there is an adenine 4bp 5’ of the mutated T

We load these data into data frame *df*, which is keyed by the name of the sample and also contains the field *dataType* indicating whether the data are whole-genome or whole-exome.

```
df<-read.csv("input/mutCounts_allExomes+allGenomes.txt",sep="\t",as.is=T)
```

```
## keep only samples with >= 50 (exome) or >= 500 (genome) thymine mutations
df<-df[c(which(grepl("WGS",df$dataType) & df$T2Ncount>499),
         which(grepl("WES",df$dataType) & df$T2Ncount>49)), ]

## load the tri- tetra- and pentanucleotide abundances. After load we have
## variable *opp* which contains abundance of tri, tetra, and penta nucleotides
## for the human genome and exome.
load("input/trinucContent_hs37d5.RData")
```

```
source('src/mSigTools.v0.13.R')
```

```
## Loading required package: SnowballC
```

```
rev.comp <- function(seq) { revc(toupper(seq)) }
```

The following function tests for statistically significant enrichment for the adenines at the -3 and -4 position relative to mutations from thymines.

```
enrichmentTest<-function(df){
  df$pval_ANNT2ANNN<-NA
  df$pval_ANNNT2ANNNN<-NA
  
  # Create columns for the reverse complements of tetra and penta nucleotides
  tmpOpp4<-opp$`4bp`
  for(ii in 1:nrow(tmpOpp4)){tmpOpp4$revComp[ii]<-rev.comp(tmpOpp4$NA.[ii])}
  tmpOpp5<-opp$`5bp`
  for(ii in 1:nrow(tmpOpp5)){tmpOpp5$revComp[ii]<-rev.comp(tmpOpp5$NA.[ii])} 
  
  for(i in 1:nrow(df)){
    dataType<-df$dataType[i]
    if(grepl("WGS",dataType)) dataType <- "genome"
    if(grepl("WES",dataType)) dataType <- "exome..agilent.V6."
    
    # test1: 
    #
    # - H0: proportion of T>N mutations with A at -3 is the same as the
    # proportion of all T's in the genome with A at -3. 
    # 
    # - HA: proportion of T>N mutations with A at -3 is > the above
    
    obs.mut<-df$T2Ncount[i]         ## total number of T>N mutations
    obs.mut.target<-df$ANNTcount[i] ## total number of T>N mutations with an A 3bp 5'
    
    abundance_T2N<-sum(tmpOpp4[substr(tmpOpp4$NA.,4,4) == "T" |
                                 substr(tmpOpp4$revComp,1,1) == "A",
                               dataType])
    
    abundance_ANNT2ANNN<-sum(tmpOpp4[(substr(tmpOpp4$NA.,4,4) == "T" &
                                        substr(tmpOpp4$NA.,1,1) == "A")
                                     |
                                       (substr(tmpOpp4$revComp,1,1) == "A" &
                                          substr(tmpOpp4$revComp,4,4) == "T"),
                                     dataType])
    
    stopifnot(mode(obs.mut.target) == "numeric")
    stopifnot(mode(obs.mut) == "numeric")
    testResult<-binom.test(obs.mut.target,
                           obs.mut,
                           abundance_ANNT2ANNN/abundance_T2N,
                           alternative = "greater")
    df$pval_ANNT2ANNN[i]<-testResult$p.value
    
    #  test2:
    #
    # - H0: proportion of T>N mutations with A at -4 is the same as the
    # proportion of all T's in the genome with A at -4.
    #
    # - HA: proportion of T>N mutations with A at -4 is > the above
    
    obs.mut<-df$T2Ncount[i]         ## total number of T>N mutations
    obs.mut.target<-df$ANNNTcount[i] ## total number of T>N mutations with an A 3bp 5'
    abundance_T2N<-sum(tmpOpp5[substr(tmpOpp5$NA.,5,5) == "T" |
                                 substr(tmpOpp5$revComp,1,1) == "A",
                               dataType])
    
    abundance_ANNNT2ANNNN<-sum(tmpOpp5[(substr(tmpOpp5$NA.,5,5) == "T" &
                                          substr(tmpOpp5$NA.,1,1) == "A" ) 
                                       |
                                         (substr(tmpOpp5$revComp,1,1) == "A" &
                                            substr(tmpOpp5$revComp,5,5) == "T")
                                       ,dataType])
    
    stopifnot(mode(obs.mut.target) == "numeric")
    stopifnot(mode(obs.mut) == "numeric")
    testResult<-binom.test(obs.mut.target,
                           obs.mut,
                           abundance_ANNNT2ANNNN/abundance_T2N,
                           alternative = "greater")
    df$pval_ANNNT2ANNNN[i]<-testResult$p.value
  }
  return(df)
}
```

Find samples with mutations from thymine that have statistically significant enrichment for adenines at the -3 and -4 positions and with proportions of adenines above determined cutoffs.

```
## add the counts for sample 62074759
df<-rbind(df,'62074759T'=list("WGS",34905,27375,22695,18549))
df<-enrichmentTest(df)

## only look at tumors that fit all criteria
result<-df[ p.adjust(df$pval_ANNT2ANNN,method="BH") < 0.05 &
              p.adjust(df$pval_ANNNT2ANNNN,method="BH") < 0.05 ,]

## plot scatter of enrichment for adenines 3 and 4bp 5' of mutated thymines
plot(df$ANNTcount/df$T2Ncount,df$ANNNTcount/df$T2Ncount,
     pch=16,cex=0.5,
     main="Identification of samples with enrichment of adenines\n(3 and 4bp 5' of mutated thymines)",
     xlab="Proportion thymines with adenine 3bp 5'",
     ylab="Proportion thymines with adenine 4bp 5'")
points(result$ANNTcount/result$T2Ncount,result$ANNNTcount/result$T2Ncount,
       pch=16,cex=0.5,col="magenta")
points(result$ANNTcount[nrow(result)]/result$T2Ncount[nrow(result)],
       result$ANNNTcount[nrow(result)]/result$T2Ncount[nrow(result)],
       pch=8,cex=0.8,col="red",lwd=1.5)
## expected ratio of A's 5' of thymines (abundance in the genome)
points(c(325868953,3890707)/c(1074123658,14758195),
       c(318941964,3724456)/c(1074123658,14758195),
       pch=c(3,4),cex=0.8,col="lightgreen",lwd=3)
legend("topleft",
       legend=c("not significant","significant","62074759",
                "expected_exomes","expected_genomes"),
       col=c("black","magenta","red","lightgreen","lightgreen"),
       pch=c(16,16,8,4,3),
       pt.lwd=c(1,1,1.5,2,2),
       pt.cex=1.1,cex=0.7,bty="n",ncol=1)
```

```
selected <- result[result$ANNTcount/result$T2Ncount > 0.4 | 
                   result$ANNNTcount/result$T2Ncount > 0.4,]

## format to print output table to html
selected$pval_ANNT2ANNN<-format(selected$pval_ANNT2ANNN,digits=3)
selected$pval_ANNNT2ANNNN<-format(selected$pval_ANNNT2ANNNN,digits=3)
library(knitr)
knitr::kable(selected,row.names = T,
             align = "c",
             caption = paste("Whole-exome and whole-genome sequenced samples",
                             "with likely AnT exposure based on enrichment of",
                             "adenines 5' of mutated thymines"))
```

Whole-exome and whole-genome sequenced samples with likely AnT exposure based on enrichment of adenines 5’ of mutated thymines

|  | dataType | totalMuts | T2Ncount | ANNTcount | ANNNTcount | pval\_ANNT2ANNN | pval\_ANNNT2ANNNN |
| --- | --- | --- | --- | --- | --- | --- | --- |
| Eso-AdenoCa::LP6005935-DNA\_B03\_\_\_ICGC:ESAD-UK | WGS\_Other | 45144 | 33324 | 13342 | 12543 | 6.67e-309 | 7.14e-228 |
| Panc-AdenoCa::8069334\_\_\_ICGC:PACA-AU | WGS\_Other | 7164 | 3682 | 1483 | 1331 | 1.19e-37 | 7.78e-19 |
| Prost-AdenoCa::0047\_CRUK\_PC\_0047\_T1\_DNA\_\_\_ICGC:PRAD-UK | WGS\_Other | 3369 | 1514 | 863 | 730 | 3.36e-102 | 2.09e-53 |
| COAD-US::SP22031\_\_\_PCAWG | WGS\_ICGC | 234336 | 71001 | 31769 | 33139 | 0.00e+00 | 0.00e+00 |
| COAD-US::SP16886\_\_\_PCAWG | WGS\_ICGC | 850298 | 337390 | 149095 | 152004 | 0.00e+00 | 0.00e+00 |
| COAD-US::SP19295\_\_\_PCAWG | WGS\_ICGC | 260008 | 72187 | 31520 | 32393 | 0.00e+00 | 0.00e+00 |
| COAD-US::SP17905\_\_\_PCAWG | WGS\_ICGC | 2439746 | 1129261 | 475376 | 468709 | 0.00e+00 | 0.00e+00 |
| COAD-US::SP21400\_\_\_PCAWG | WGS\_ICGC | 794330 | 308195 | 127033 | 121523 | 0.00e+00 | 0.00e+00 |
| COAD-US::SP18946\_\_\_PCAWG | WGS\_ICGC | 962132 | 330311 | 143781 | 139479 | 0.00e+00 | 0.00e+00 |
| ESAD-UK::SP111026\_\_\_PCAWG | WGS\_ICGC | 47007 | 38593 | 15452 | 14388 | 0.00e+00 | 1.02e-241 |
| ESAD-UK::SP111101\_\_\_PCAWG | WGS\_ICGC | 17017 | 7777 | 3192 | 2964 | 4.64e-89 | 6.24e-61 |
| READ-US::SP80615\_\_\_PCAWG | WGS\_ICGC | 2433765 | 1066128 | 458226 | 453231 | 0.00e+00 | 0.00e+00 |
| READ-US::SP81494\_\_\_PCAWG | WGS\_ICGC | 26908 | 8134 | 3434 | 3092 | 1.54e-113 | 2.87e-62 |
| READ-US::SP81711\_\_\_PCAWG | WGS\_ICGC | 20415 | 6971 | 3155 | 2827 | 5.01e-151 | 1.36e-87 |
| READ-US::SP80754\_\_\_PCAWG | WGS\_ICGC | 11274 | 4522 | 2738 | 2290 | 0.00e+00 | 4.21e-196 |
| UCEC-US::SP92659\_\_\_PCAWG | WGS\_ICGC | 282392 | 77606 | 36861 | 38199 | 0.00e+00 | 0.00e+00 |
| Biliary-AdenoCa::BD121T\_\_\_ICGC:BTCA-JP | WES\_Other | 2284 | 447 | 180 | 171 | 2.04e-10 | 9.16e-10 |
| Biliary-AdenoCa::BD173T\_\_\_ICGC:BTCA-JP | WES\_Other | 1231 | 749 | 678 | 513 | 3.89e-300 | 4.91e-136 |
| Biliary-AdenoCa::BD182T\_\_\_ICGC:BTCA-JP | WES\_Other | 2117 | 1814 | 974 | 771 | 5.30e-132 | 8.77e-58 |
| Biliary-AdenoCa::BD223T\_\_\_ICGC:BTCA-JP | WES\_Other | 425 | 218 | 148 | 113 | 4.29e-37 | 4.17e-17 |
| ColoRect-AdenoCa::sysucc-311T\_\_\_ICGC:COCA-CN | WES\_Other | 14191 | 5763 | 2307 | 2190 | 1.41e-109 | 3.78e-101 |
| Liver-HCC::HCC34T\_\_\_ICGC:LINC-JP | WES\_Other | 249 | 96 | 40 | 38 | 9.32e-04 | 1.40e-03 |
| Panc-AdenoCa::PCSI\_0060\_Pa\_X\_\_\_ICGC:PACA-CA | WES\_Other | 113 | 55 | 43 | 33 | 2.03e-15 | 5.09e-08 |
| Skin-Melanoma::SKCM-JWCI-WGS-8-Tumor\_\_\_doi:10.1016/j.cell.2012.06.024 | WES\_Other | 705 | 57 | 24 | 24 | 7.80e-03 | 3.96e-03 |
| Eso-AdenoCa::ESO-173\_\_\_PMID:23525077 | WES\_Other | 202 | 77 | 35 | 30 | 2.75e-04 | 5.52e-03 |
| Uterus-AdenoCa::T155\_\_\_PMID:23104009 | WES\_Other | 983 | 186 | 83 | 73 | 8.50e-08 | 1.84e-05 |
| ACUTE MYELOID LEUKEMIA::TCGA-AB-2824-03B-01W-0728-08\_\_\_TCGA-LAML | WES\_TCGA | 72 | 71 | 46 | 35 | 1.74e-11 | 1.10e-05 |
| ACUTE MYELOID LEUKEMIA::TCGA-AB-2851-03B-01W-0728-08\_\_\_TCGA-LAML | WES\_TCGA | 60 | 60 | 45 | 24 | 7.02e-15 | 8.47e-03 |
| ACUTE MYELOID LEUKEMIA::TCGA-AB-2867-03B-01W-0728-08\_\_\_TCGA-LAML | WES\_TCGA | 115 | 104 | 85 | 40 | 1.00e-31 | 2.01e-03 |
| ACUTE MYELOID LEUKEMIA::TCGA-AB-2868-03B-01W-0728-08\_\_\_TCGA-LAML | WES\_TCGA | 212 | 197 | 159 | 82 | 1.46e-56 | 3.68e-07 |
| BLADDER UROTHELIAL CARCINOMA::TCGA-G2-AA3B-01A-11D-A391-08\_\_\_TCGA-BLCA | WES\_TCGA | 946 | 66 | 38 | 26 | 1.04e-07 | 7.94e-03 |
| BLADDER UROTHELIAL CARCINOMA::TCGA-GC-A6I3-01A-11D-A31L-08\_\_\_TCGA-BLCA | WES\_TCGA | 346 | 64 | 38 | 27 | 3.33e-08 | 2.25e-03 |
| CERVICAL SQUAMOUS CELL CARCINOMA AND ENDOCERVICAL ADENOCARCINOMA::TCGA-FU-A3HZ-01A-11D-A20U-09\_\_\_TCGA-CESC | WES\_TCGA | 3256 | 1279 | 493 | 523 | 5.53e-21 | 1.46e-34 |
| COLON ADENOCARCINOMA::TCGA-AY-4071-01A-01W-1073-09\_\_\_TCGA-COAD | WES\_TCGA | 165 | 53 | 31 | 24 | 9.56e-07 | 1.21e-03 |
| ESOPHAGEAL CARCINOMA::TCGA-2H-A9GM-01A-11D-A37C-09\_\_\_TCGA-ESCA | WES\_TCGA | 173 | 61 | 31 | 24 | 4.60e-05 | 1.07e-02 |
| ESOPHAGEAL CARCINOMA::TCGA-IG-A4QS-01A-11D-A27G-09\_\_\_TCGA-ESCA | WES\_TCGA | 321 | 154 | 57 | 66 | 2.81e-03 | 1.42e-06 |
| ESOPHAGEAL CARCINOMA::TCGA-R6-A6L4-01A-11D-A31U-09\_\_\_TCGA-ESCA | WES\_TCGA | 163 | 81 | 34 | 37 | 1.86e-03 | 5.31e-05 |
| HEAD AND NECK SQUAMOUS CELL CARCINOMA::TCGA-BA-A4IG-01A-11D-A25Y-08\_\_\_TCGA-HNSC | WES\_TCGA | 170 | 52 | 35 | 25 | 9.57e-10 | 3.15e-04 |
| RECTUM ADENOCARCINOMA::TCGA-AG-3892-01A-01W-1073-09\_\_\_TCGA-READ | WES\_TCGA | 3593 | 976 | 406 | 397 | 1.85e-24 | 3.72e-26 |
| RECTUM ADENOCARCINOMA::TCGA-AG-3902-01A-01W-1073-09\_\_\_TCGA-READ | WES\_TCGA | 178 | 66 | 43 | 38 | 6.01e-11 | 2.46e-08 |
| STOMACH ADENOCARCINOMA::TCGA-BR-6453-01A-11D-1800-08\_\_\_TCGA-STAD | WES\_TCGA | 228 | 125 | 53 | 44 | 9.01e-05 | 8.40e-03 |
| STOMACH ADENOCARCINOMA::TCGA-D7-A4Z0-01A-22D-A25D-08\_\_\_TCGA-STAD | WES\_TCGA | 369 | 279 | 116 | 99 | 3.82e-08 | 9.16e-05 |
| UTERINE CORPUS ENDOMETRIAL CARCINOMA::TCGA-A5-A0GP-01A-11W-A062-09\_\_\_TCGA-UCEC | WES\_TCGA | 2229 | 589 | 241 | 214 | 2.81e-14 | 1.62e-09 |
| UTERINE CORPUS ENDOMETRIAL CARCINOMA::TCGA-AJ-A5DW-01A-11D-A27P-09\_\_\_TCGA-UCEC | WES\_TCGA | 5623 | 1141 | 454 | 460 | 1.67e-22 | 5.21e-29 |
| UTERINE CORPUS ENDOMETRIAL CARCINOMA::TCGA-AP-A1E0-01A-11D-A135-09\_\_\_TCGA-UCEC | WES\_TCGA | 8411 | 1853 | 726 | 749 | 1.75e-32 | 1.57e-46 |
| UTERINE CORPUS ENDOMETRIAL CARCINOMA::TCGA-AX-A1CE-01A-11D-A135-09\_\_\_TCGA-UCEC | WES\_TCGA | 19621 | 3038 | 1379 | 1097 | 4.74e-110 | 2.32e-40 |
| UTERINE CORPUS ENDOMETRIAL CARCINOMA::TCGA-BK-A6W3-01A-12D-A34Q-09\_\_\_TCGA-UCEC | WES\_TCGA | 7184 | 1672 | 672 | 649 | 6.42e-34 | 2.15e-34 |
| UTERINE CORPUS ENDOMETRIAL CARCINOMA::TCGA-DF-A2KV-01A-11D-A17W-09\_\_\_TCGA-UCEC | WES\_TCGA | 2391 | 654 | 254 | 273 | 4.93e-12 | 2.43e-20 |
| UTERINE CORPUS ENDOMETRIAL CARCINOMA::TCGA-E6-A1M0-01A-11D-A142-09\_\_\_TCGA-UCEC | WES\_TCGA | 4792 | 1011 | 418 | 401 | 1.61e-24 | 5.95e-24 |
| UTERINE CORPUS ENDOMETRIAL CARCINOMA::TCGA-EO-A3AV-01A-12D-A19Y-09\_\_\_TCGA-UCEC | WES\_TCGA | 7328 | 2014 | 821 | 811 | 1.19e-43 | 1.79e-49 |
| UTERINE CORPUS ENDOMETRIAL CARCINOMA::TCGA-EO-A3AY-01A-12D-A19Y-09\_\_\_TCGA-UCEC | WES\_TCGA | 6317 | 1731 | 728 | 663 | 2.49e-44 | 3.93e-33 |
| UTERINE CORPUS ENDOMETRIAL CARCINOMA::TCGA-EY-A1GD-01A-11D-A13L-09\_\_\_TCGA-UCEC | WES\_TCGA | 1895 | 701 | 282 | 256 | 2.54e-15 | 2.52e-11 |
| UTERINE CORPUS ENDOMETRIAL CARCINOMA::TCGA-EY-A1GI-01A-11D-A13L-09\_\_\_TCGA-UCEC | WES\_TCGA | 7878 | 1787 | 722 | 708 | 3.74e-37 | 1.09e-40 |
| UTERINE CORPUS ENDOMETRIAL CARCINOMA::TCGA-QF-A5YS-01A-11D-A31U-09\_\_\_TCGA-UCEC | WES\_TCGA | 2789 | 534 | 228 | 231 | 5.70e-16 | 1.06e-19 |
| UTERINE CORPUS ENDOMETRIAL CARCINOMA::TCGA-QS-A5YQ-01A-11D-A31U-09\_\_\_TCGA-UCEC | WES\_TCGA | 1670 | 323 | 128 | 139 | 2.11e-07 | 2.57e-12 |
| 62074759T | WGS | 34905 | 27375 | 22695 | 18549 | 0.00e+00 | 0.00e+00 |

```
write.table(selected,file="candidatesW_AnT.txt",sep="\t",quote=F,row.names = T)
```
