## Supplementary figures and images for "Identification of novel mutational signatures in Asian oral squamous cell carcinomas associated with bacterial infections"

### TCGA-AB-2824-03B-01W-0728-08.exposures.pdf

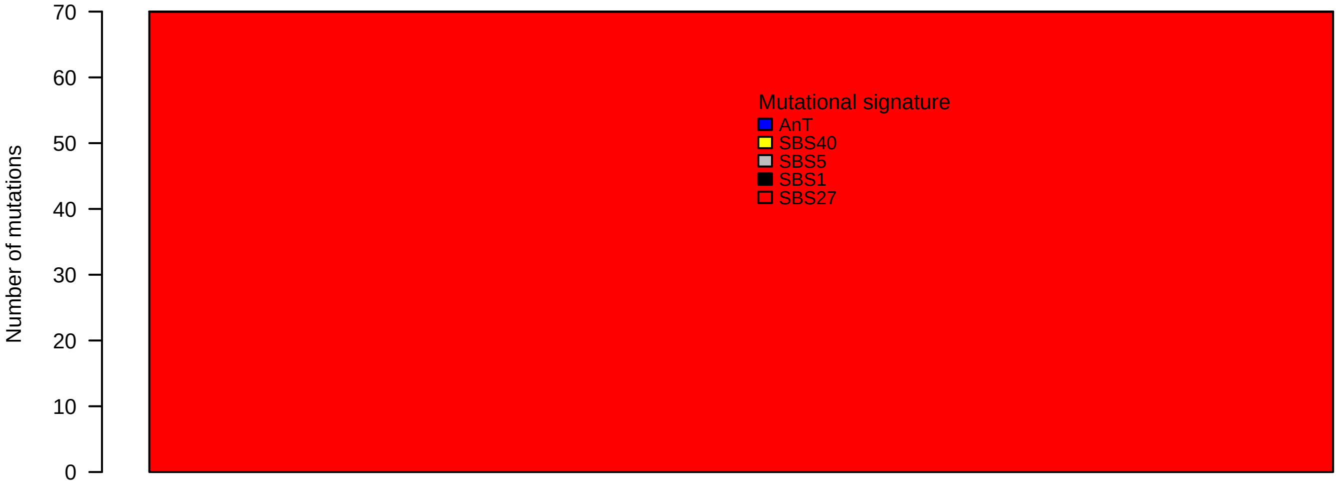

### TCGA-AB-2824-03B-01W-0728-08.pval.histogram.pdf

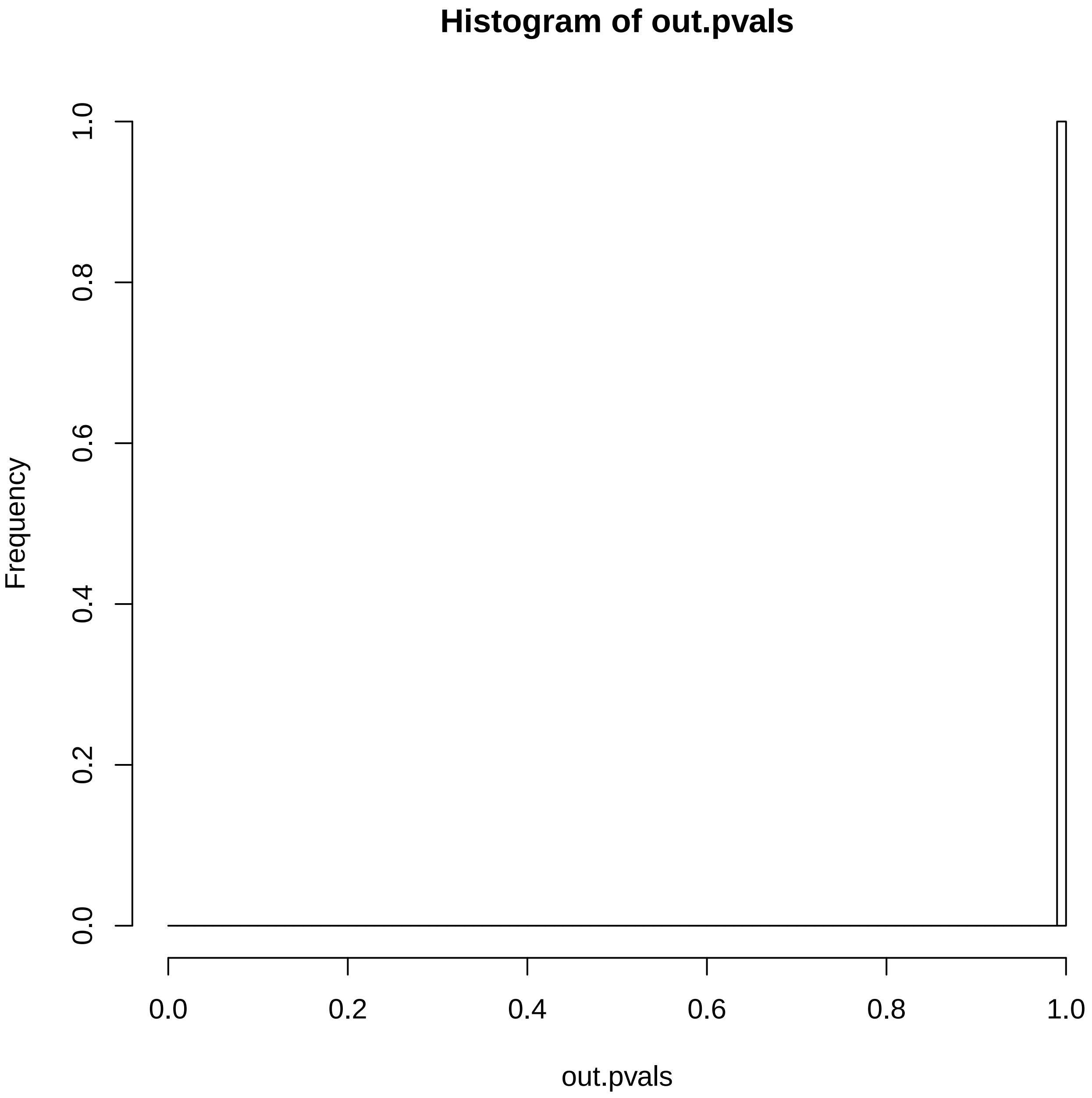

### TCGA-AB-2824-03B-01W-0728-08.reconstruction.err.pdf

##### Reconstruction error

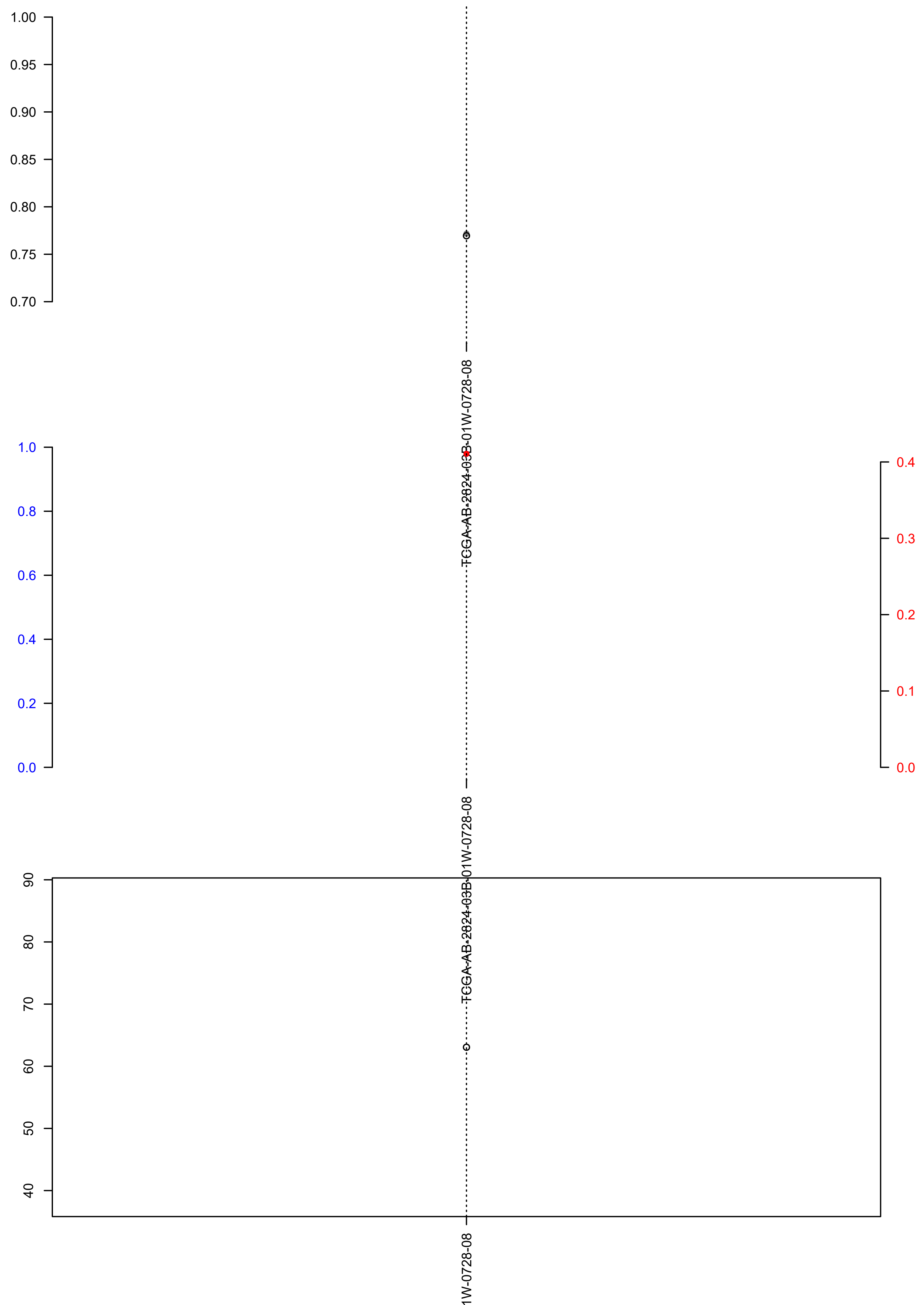

### TCGA-AB-2851-03B-01W-0728-08.exposures.pdf

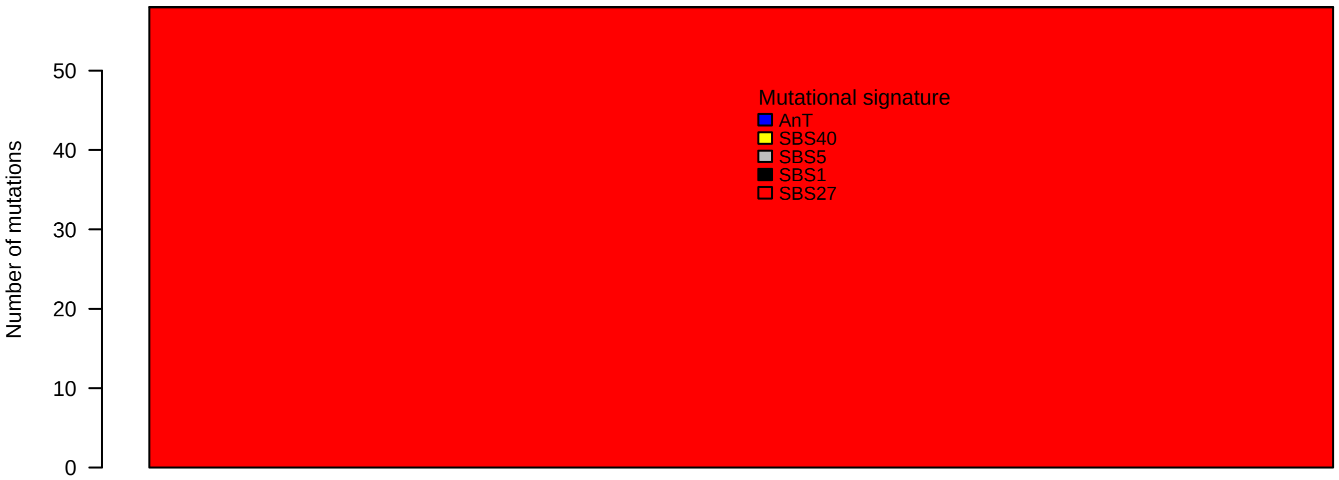

### TCGA-AB-2851-03B-01W-0728-08.pval.histogram.pdf

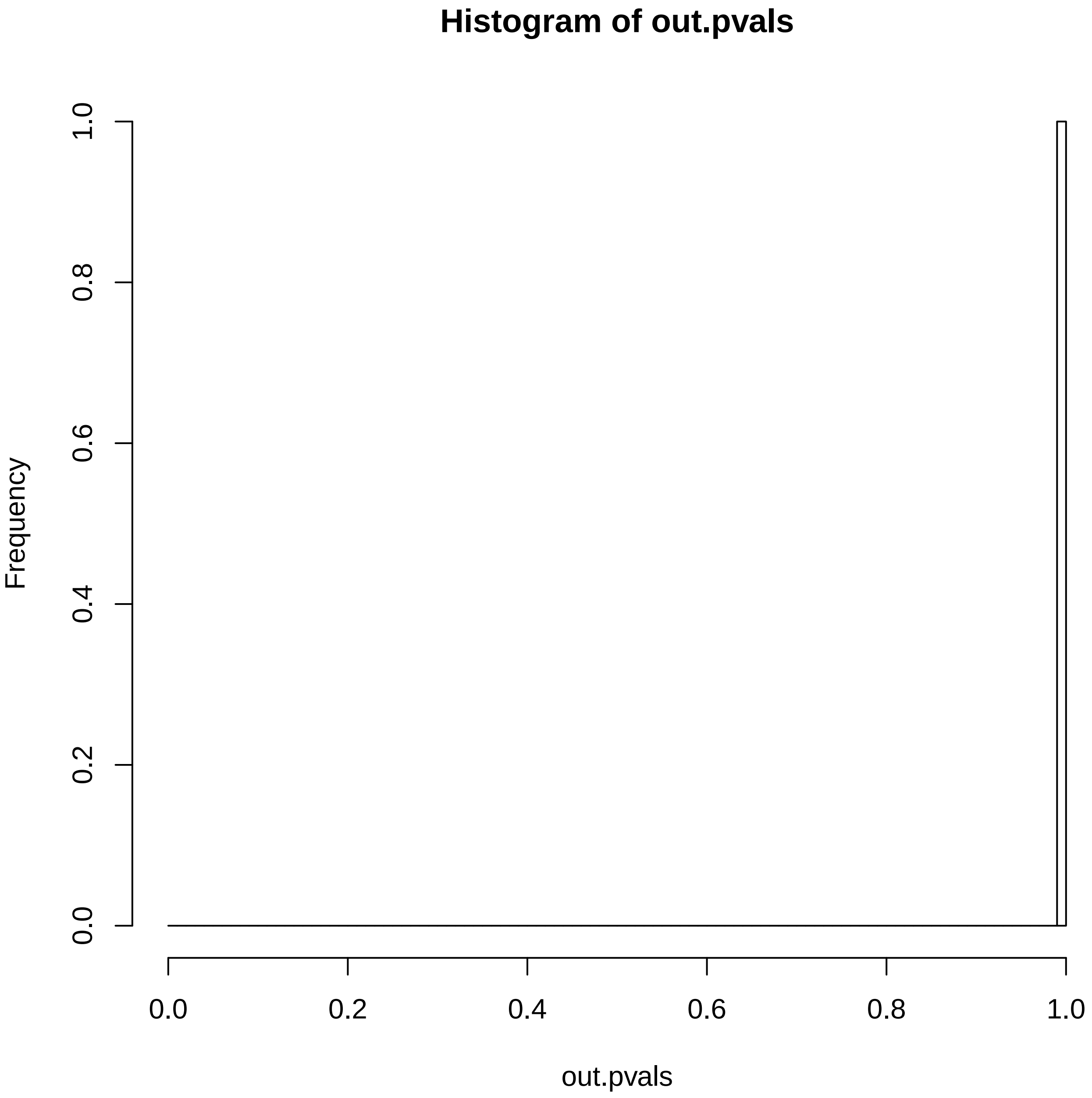

### TCGA-AB-2851-03B-01W-0728-08.reconstruction.err.pdf

**Reconstruction error**

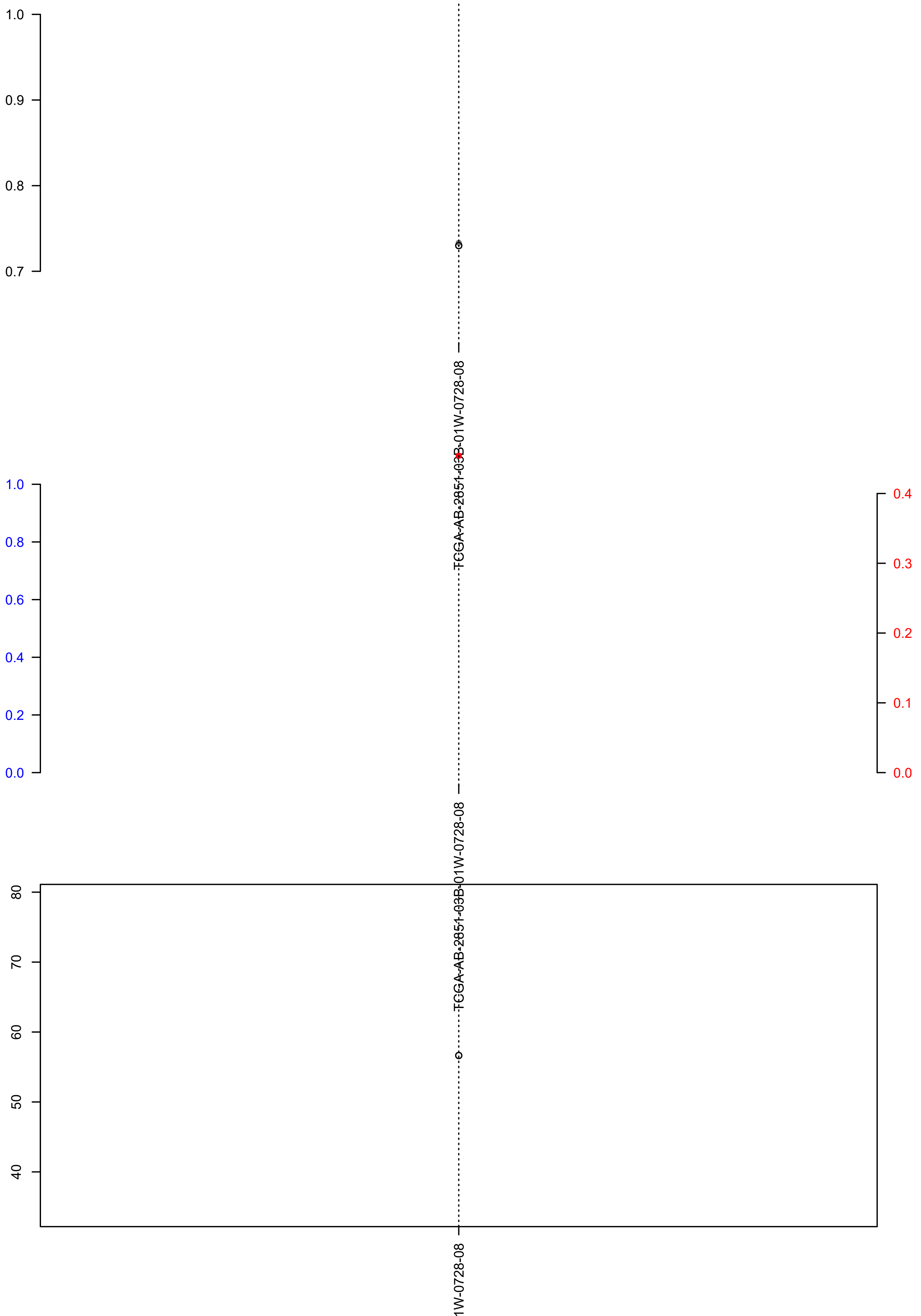

### TCGA-AB-2867-03B-01W-0728-08.exposures.pdf

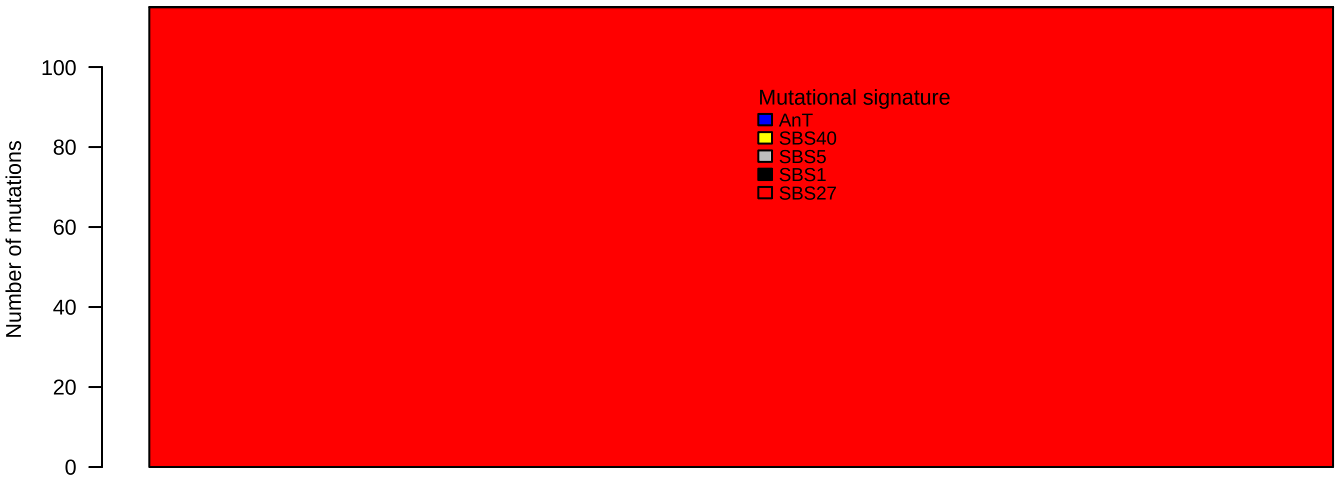

### TCGA-AB-2867-03B-01W-0728-08.reconstruction.err.pdf

##### Reconstruction error

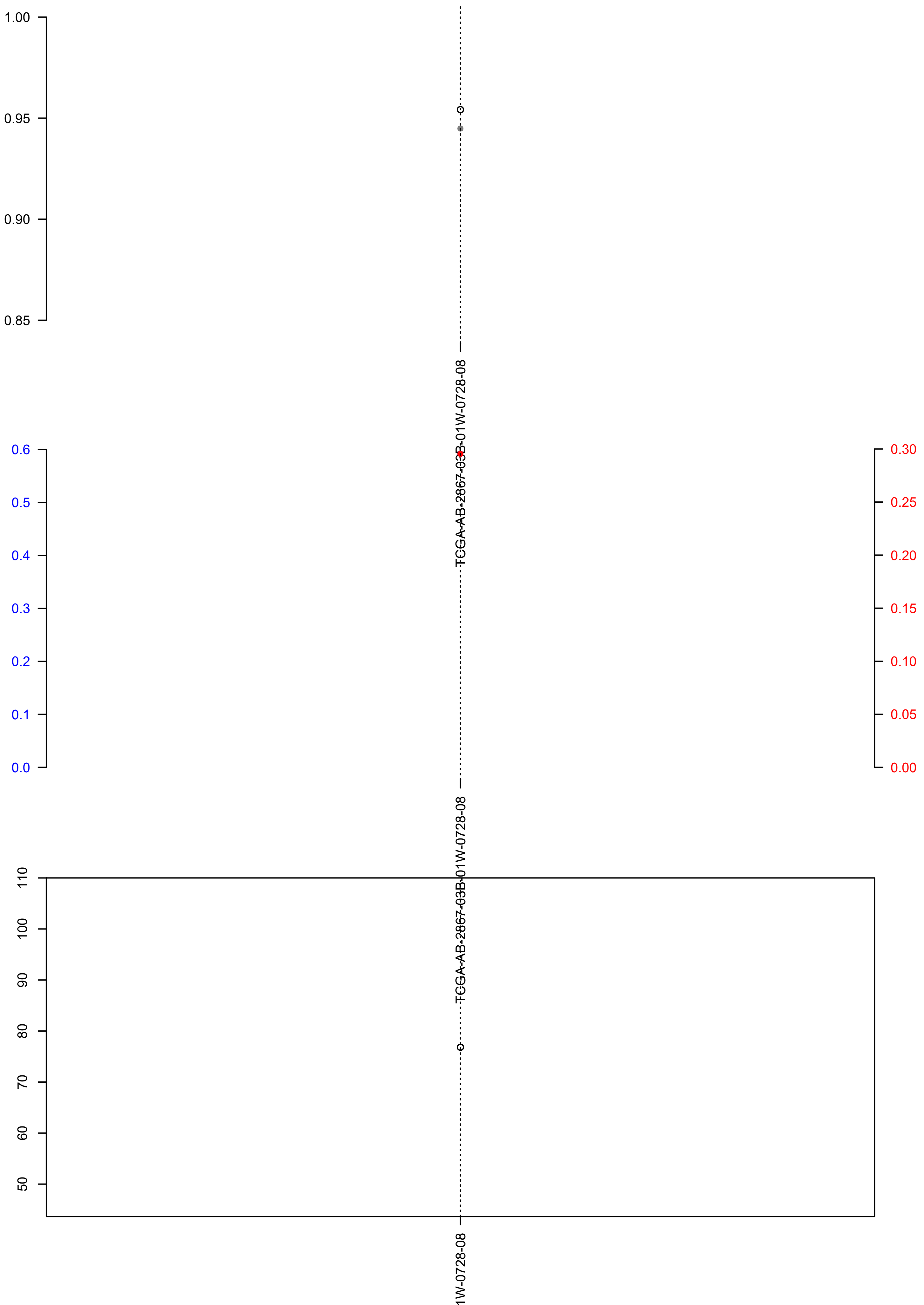

### TCGA-AB-2868-03B-01W-0728-08.check.with.sig.pdf

#### TCGA.AB.2868.03B.01W.0728.08

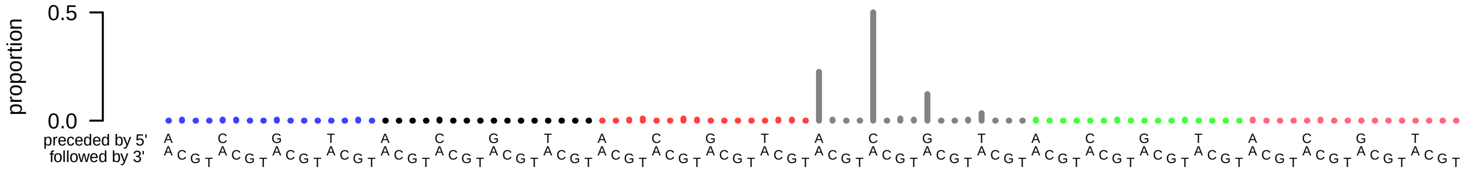

### TCGA-AB-2868-03B-01W-0728-08.exposures.pdf

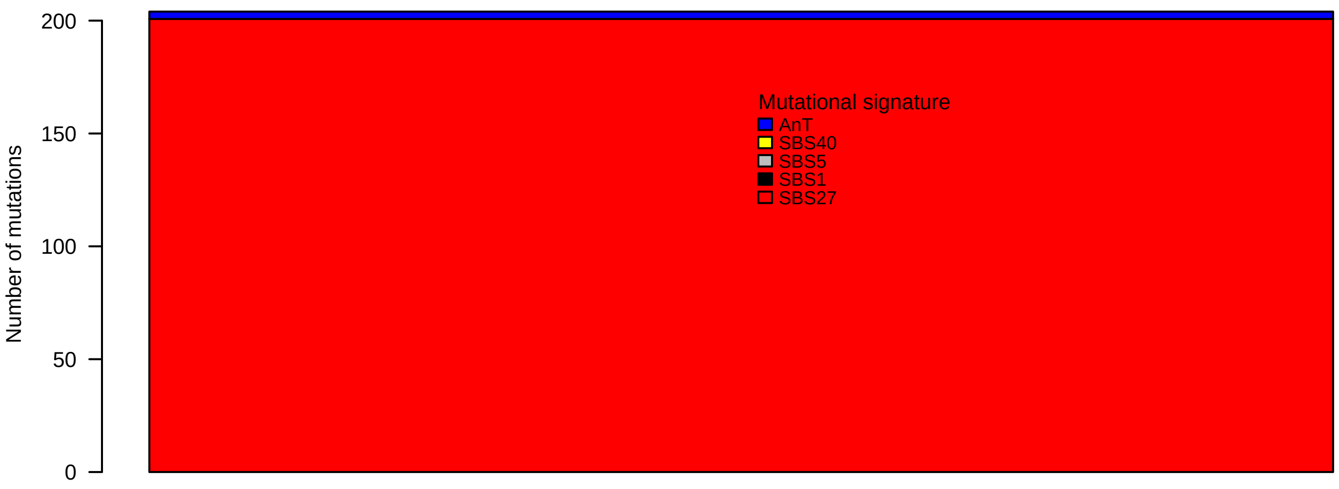

### TCGA-AB-2868-03B-01W-0728-08.pval.histogram.pdf

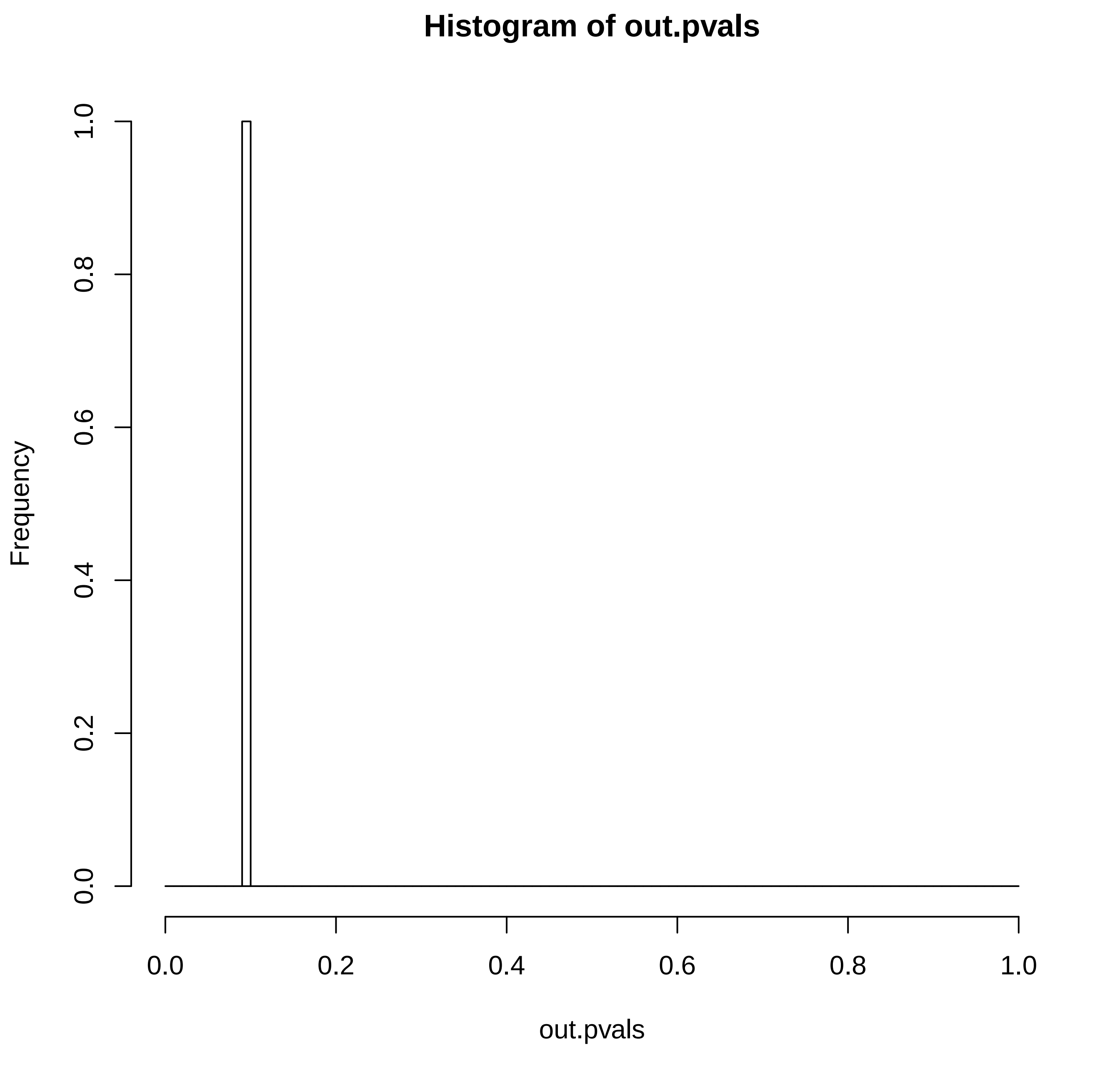

### TCGA-AB-2868-03B-01W-0728-08.reconstruction.err.pdf

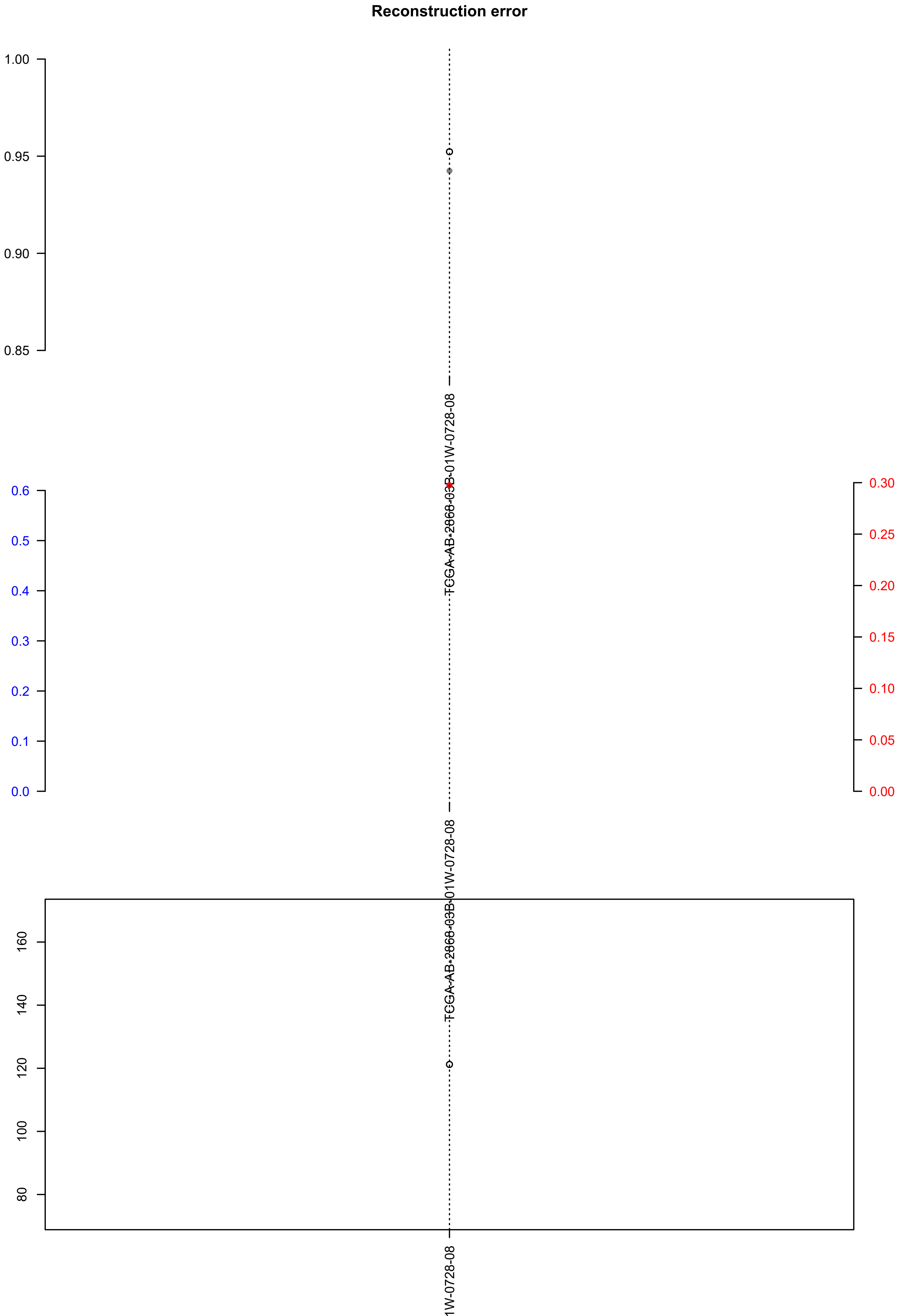

### TCGA-AG-3892-01A-01W-1073-09.exposures.pdf

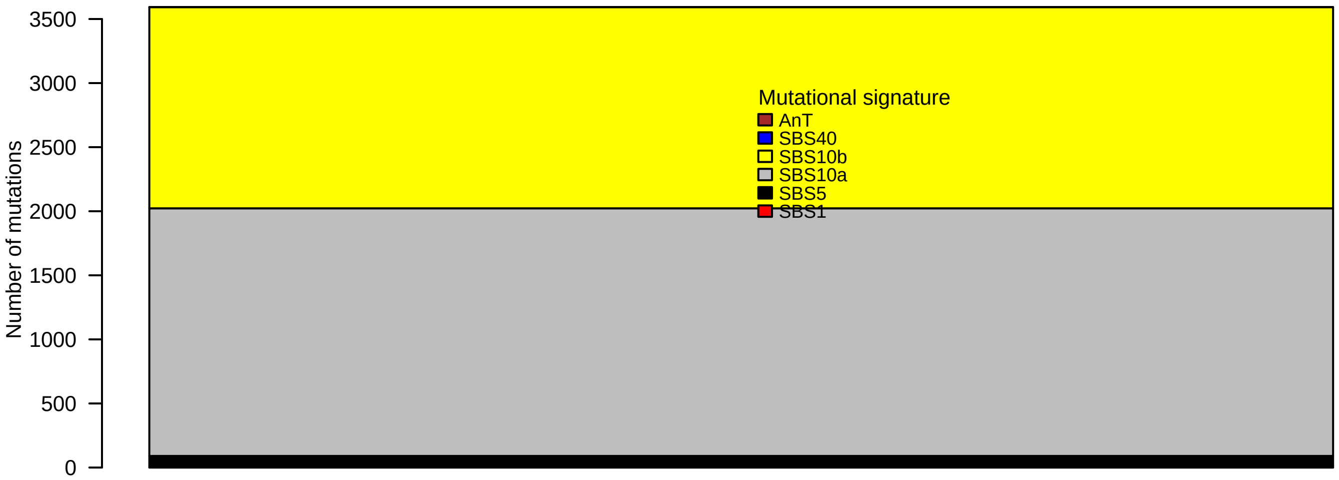

### TCGA-AG-3892-01A-01W-1073-09.pval.histogram.pdf

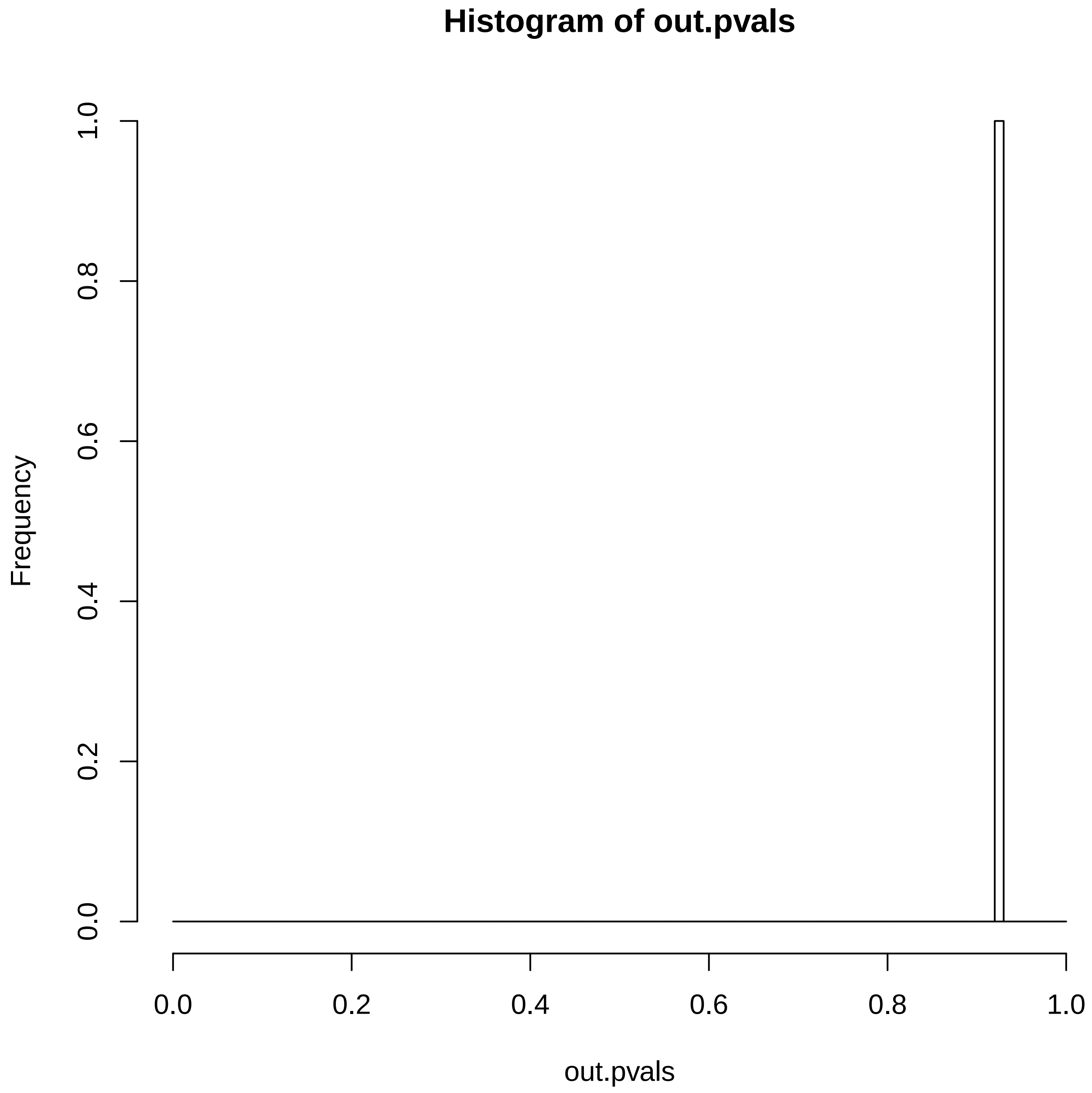

### TCGA-AG-3892-01A-01W-1073-09.reconstruction.err.pdf

**Reconstruction error**

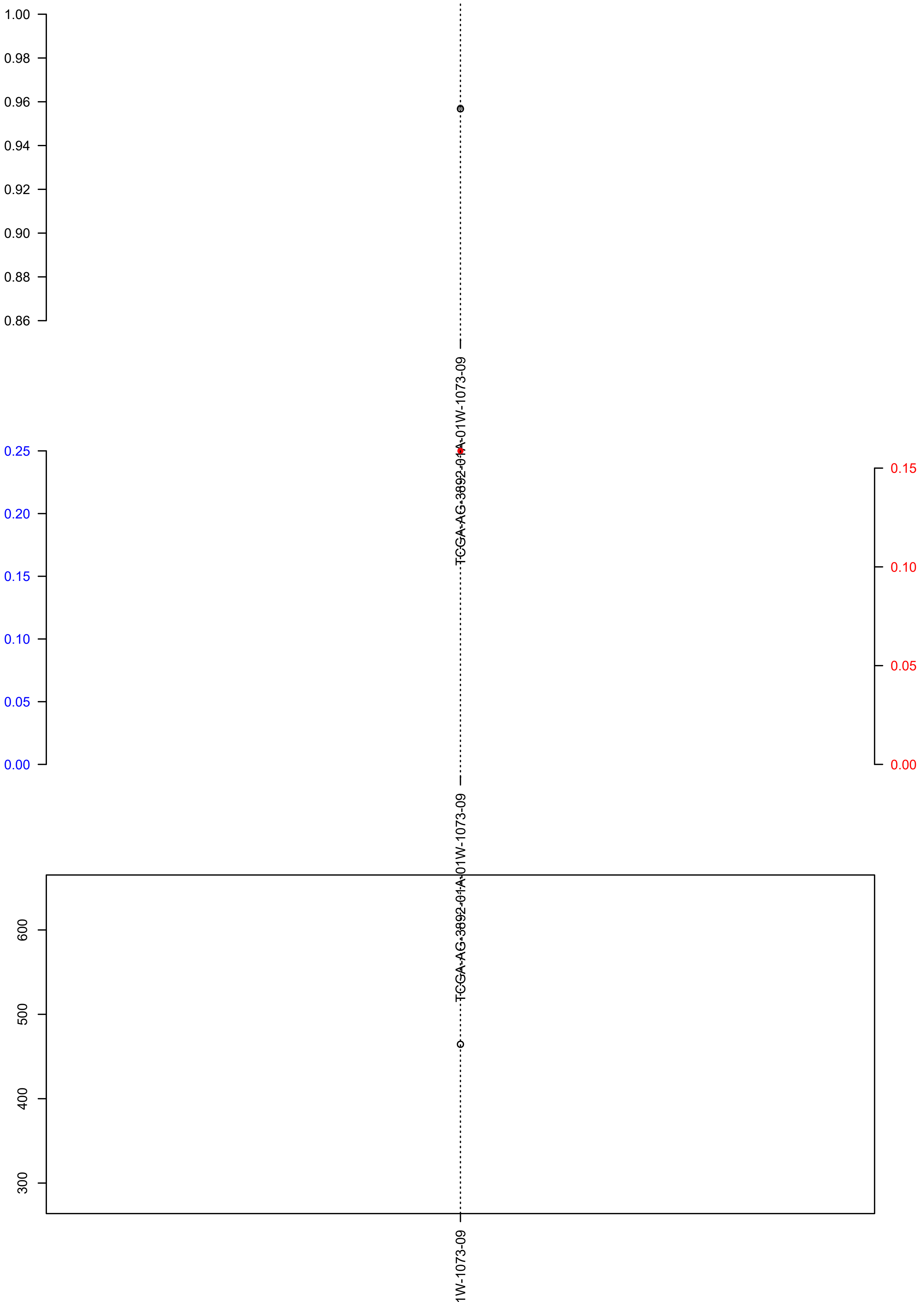

### TCGA-AG-3902-01A-01W-1073-09.check.with.sig.pdf

##### TCGA.AG.3902.01A.01W.1073.09

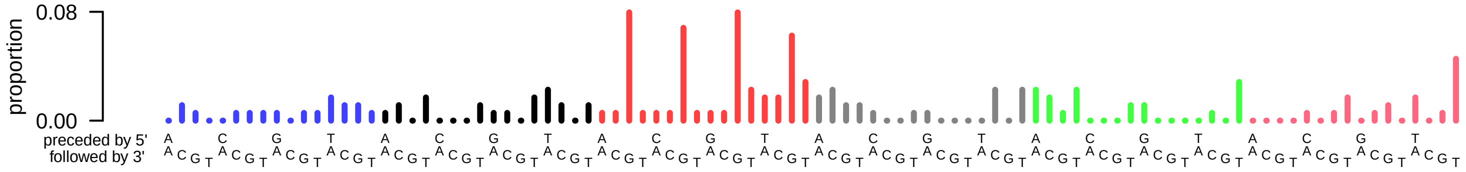

### TCGA-AG-3902-01A-01W-1073-09.exposures.pdf

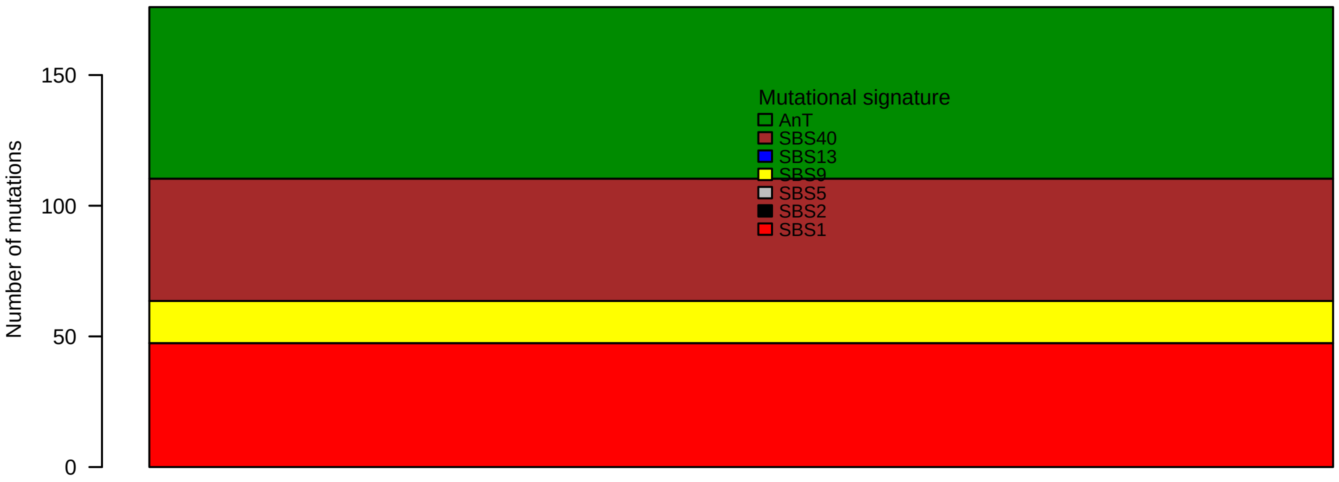

### TCGA-AG-3902-01A-01W-1073-09.pval.histogram.pdf

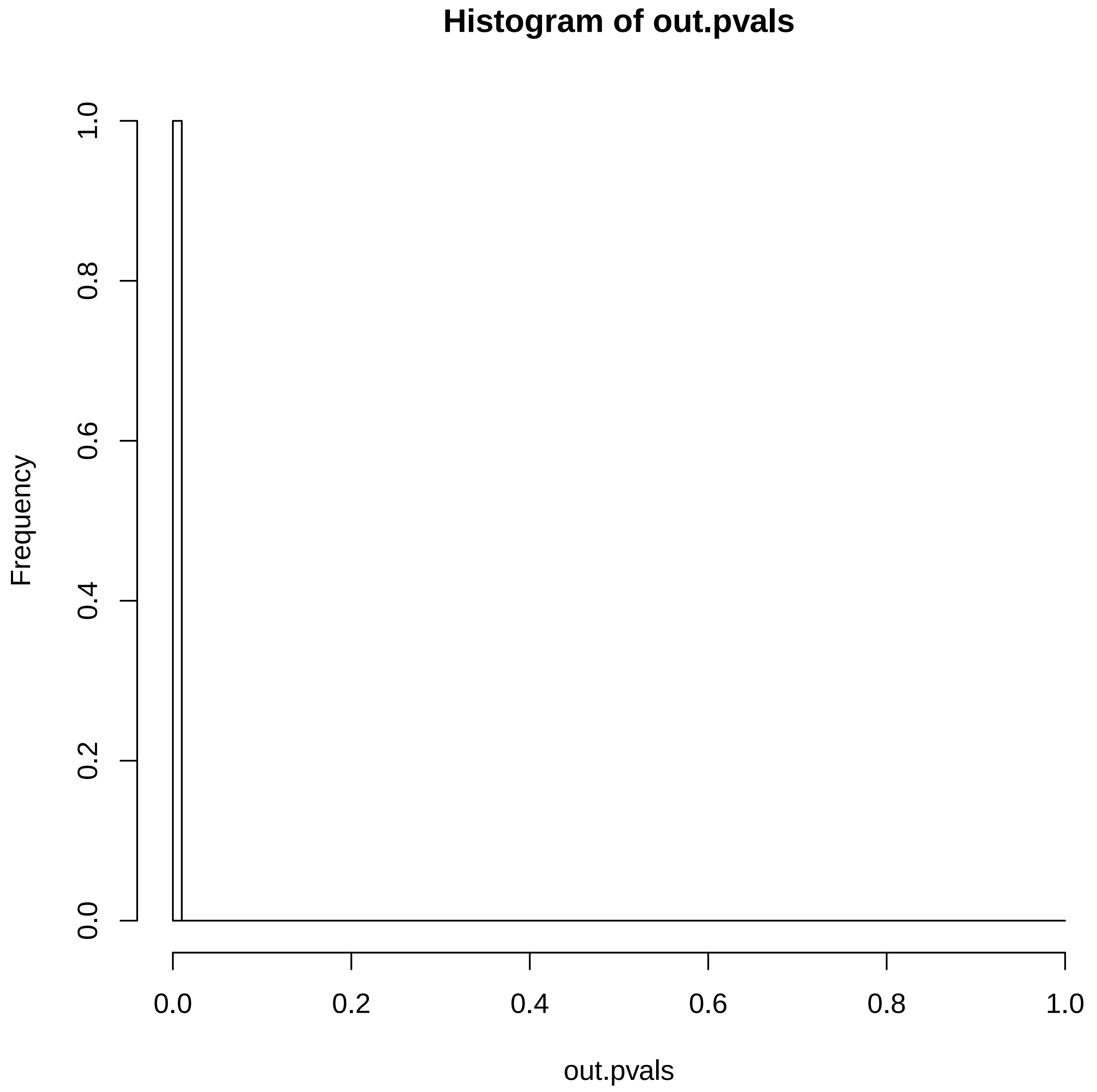

### TCGA-AG-3902-01A-01W-1073-09.reconstruction.err.pdf

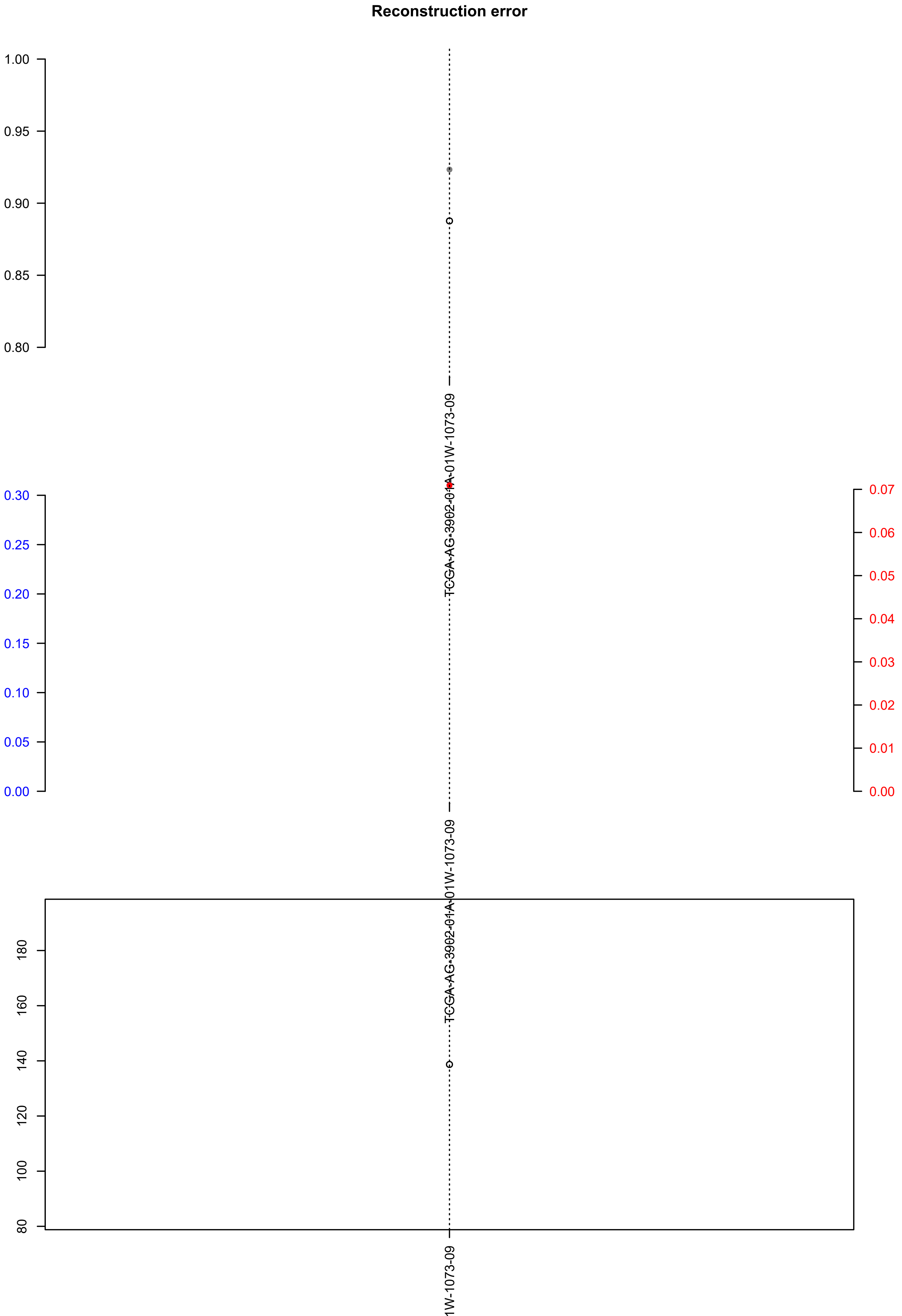

### TCGA-AJ-A5DW-01A-11D-A27P-09.exposures.pdf

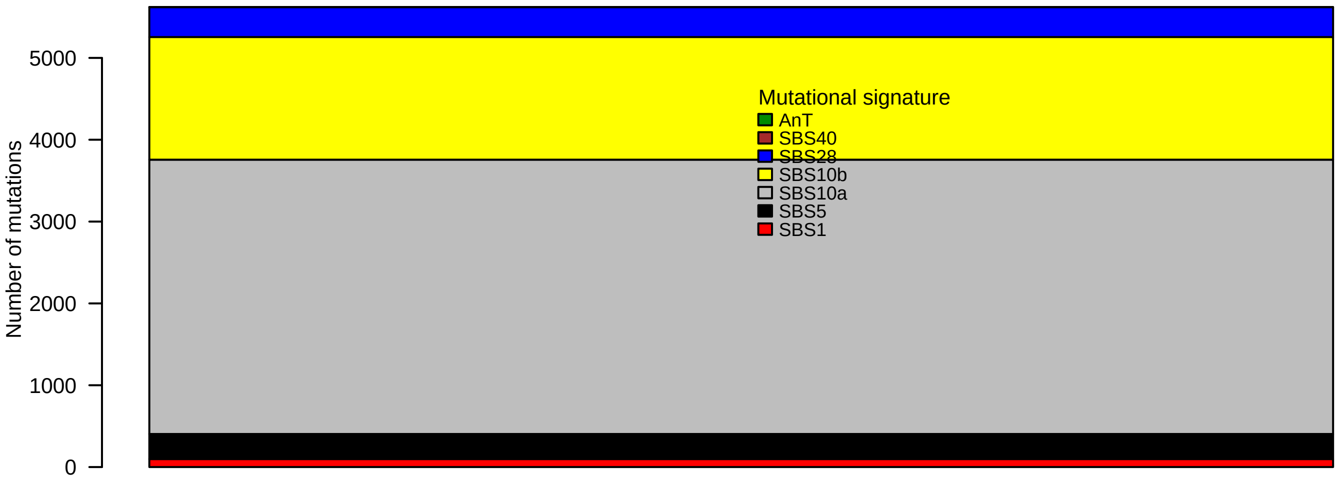

### TCGA-AJ-A5DW-01A-11D-A27P-09.pval.histogram.pdf

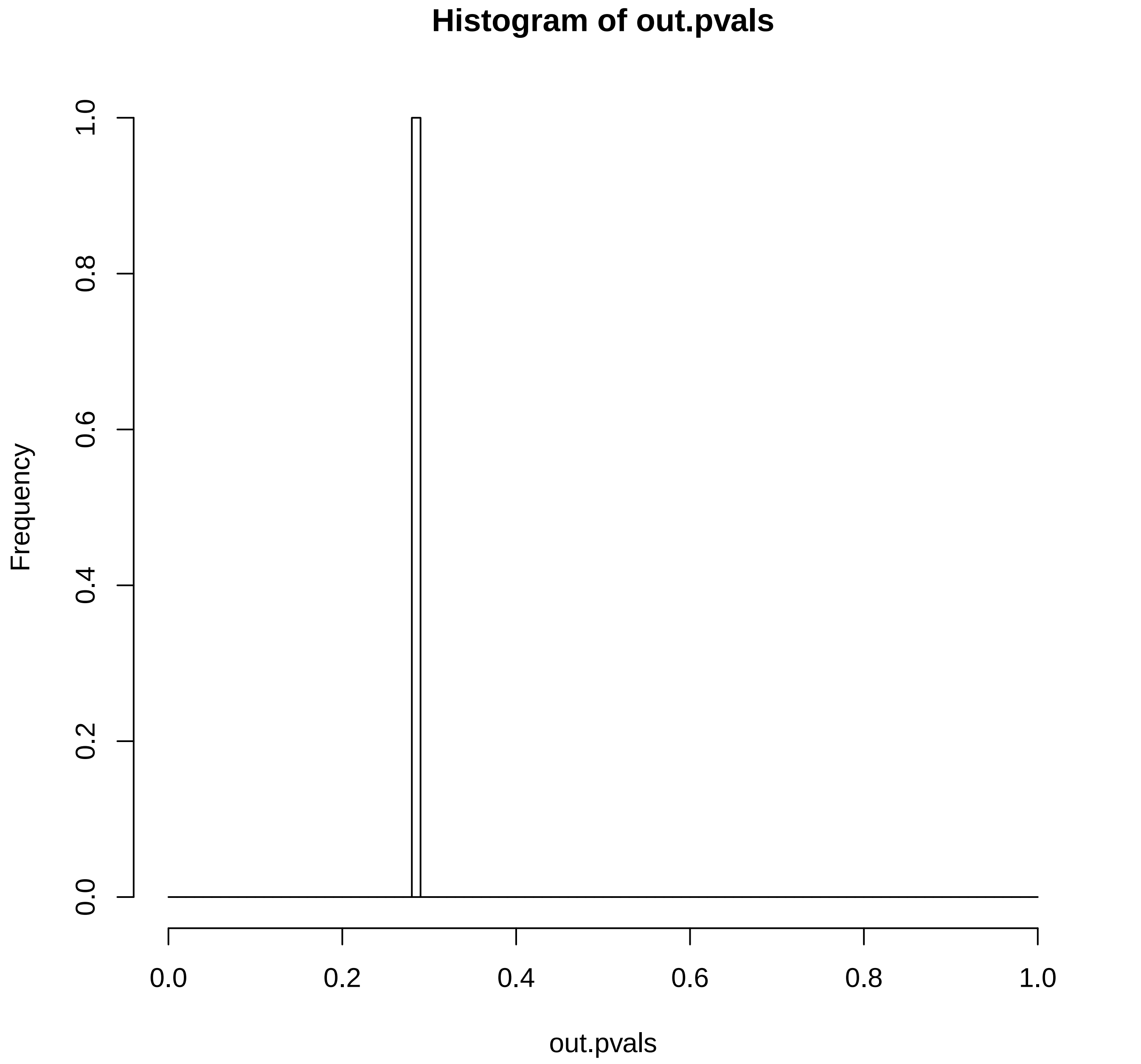

### TCGA-AJ-A5DW-01A-11D-A27P-09.reconstruction.err.pdf

##### Reconstruction error

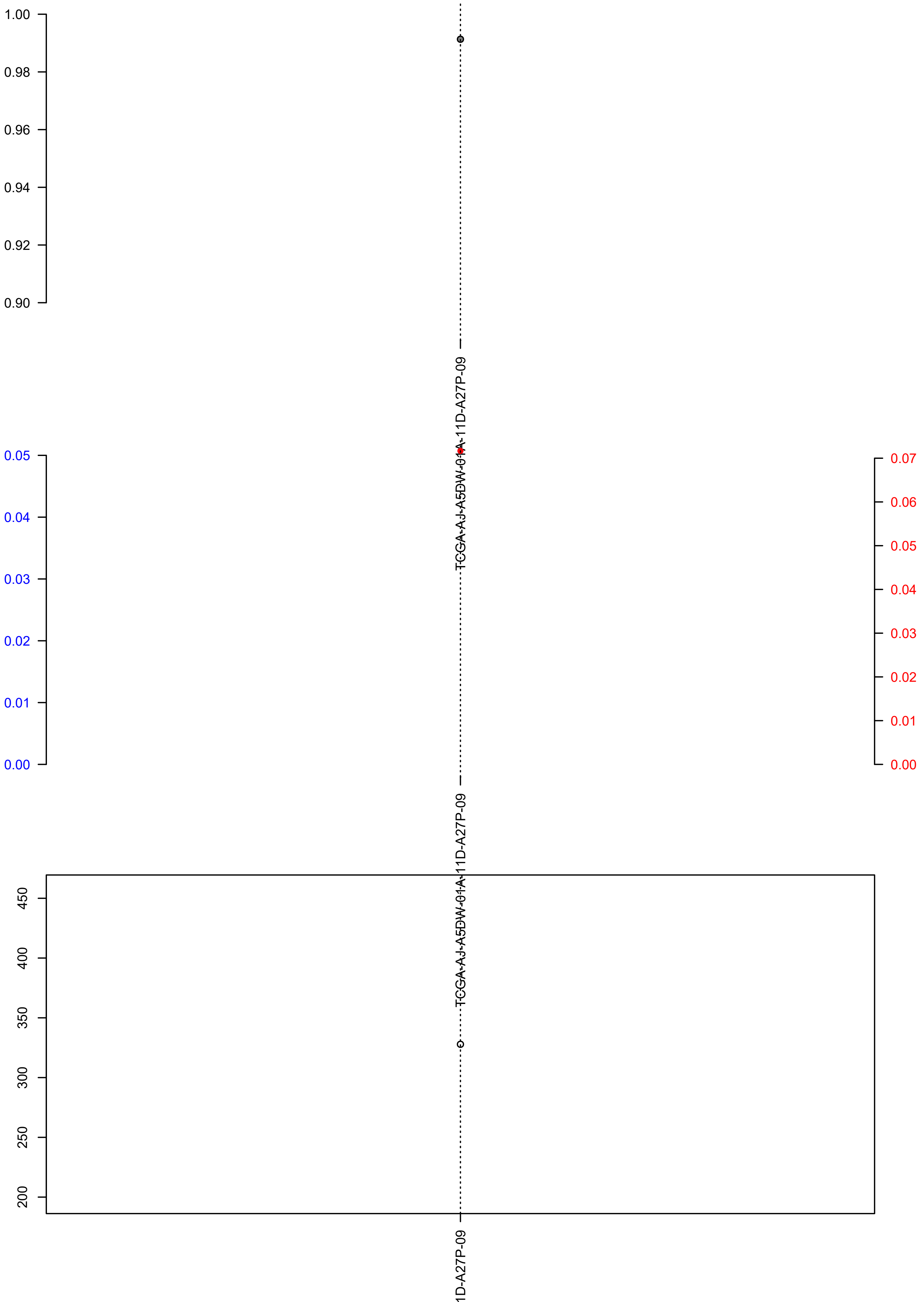

### TCGA-AP-A1E0-01A-11D-A135-09.exposures.pdf

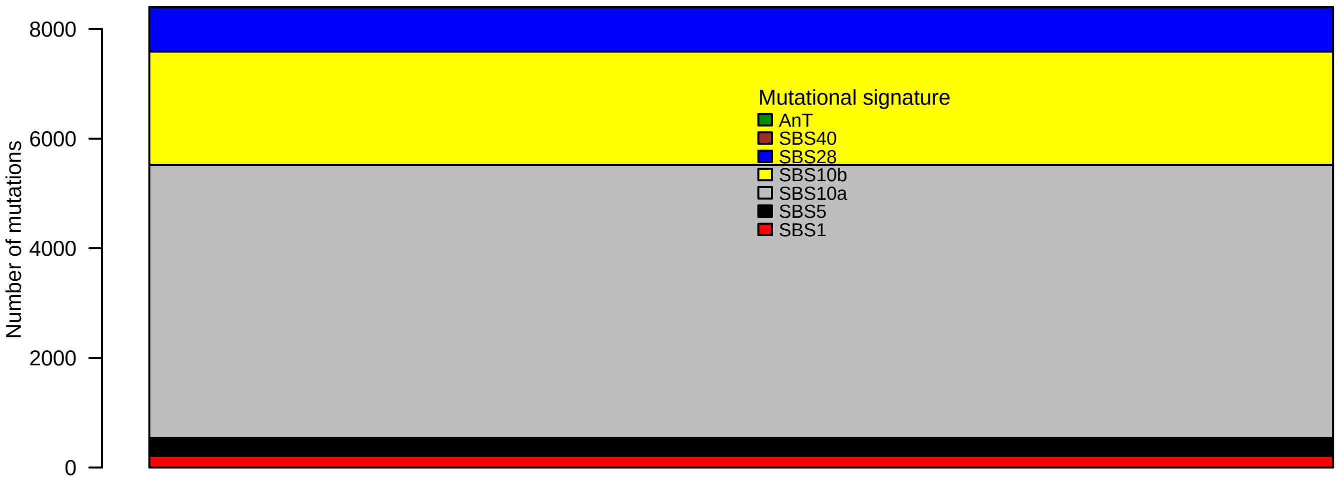

### TCGA-AP-A1E0-01A-11D-A135-09.pval.histogram.pdf

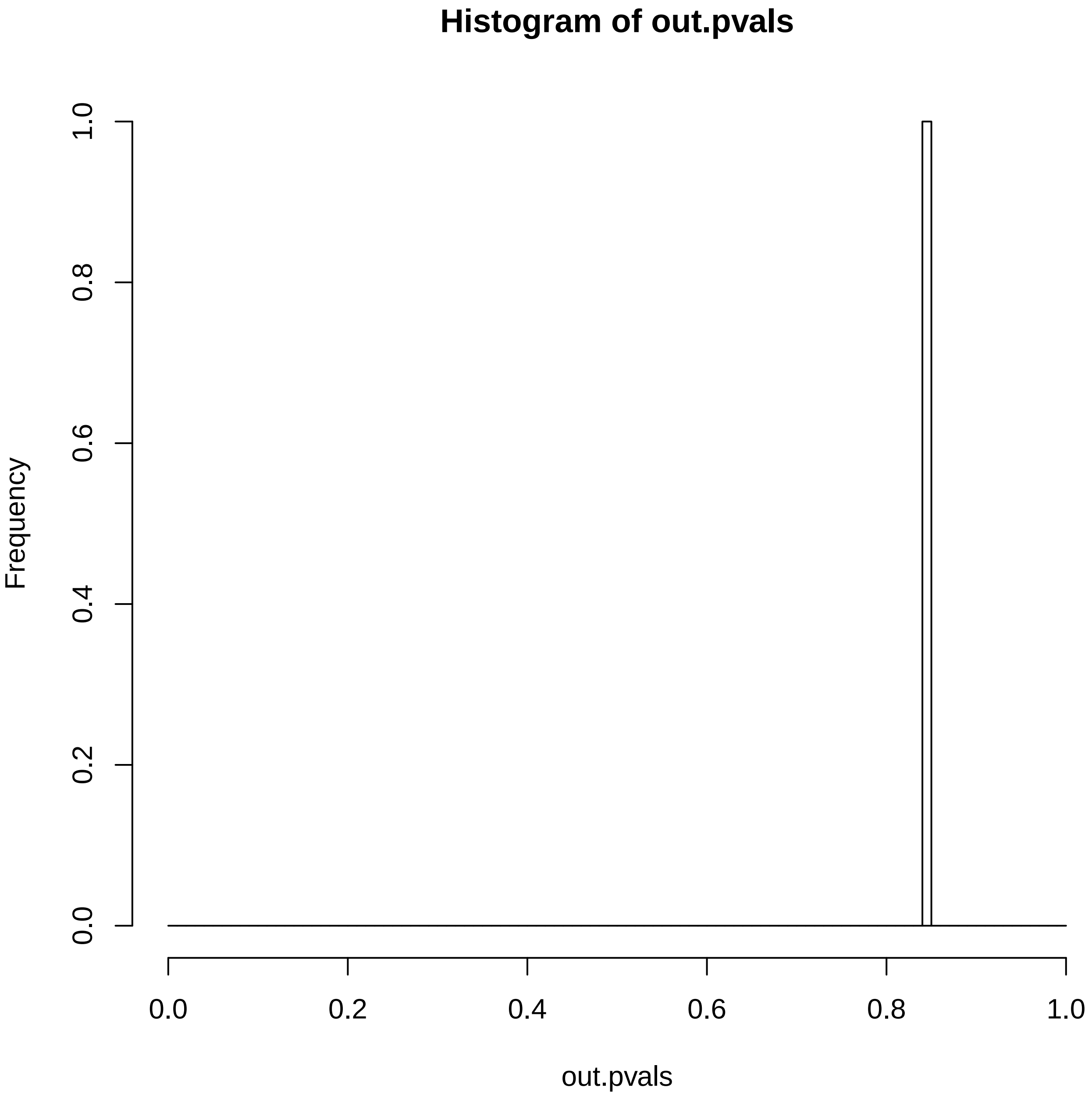

### TCGA-AP-A1E0-01A-11D-A135-09.reconstruction.err.pdf

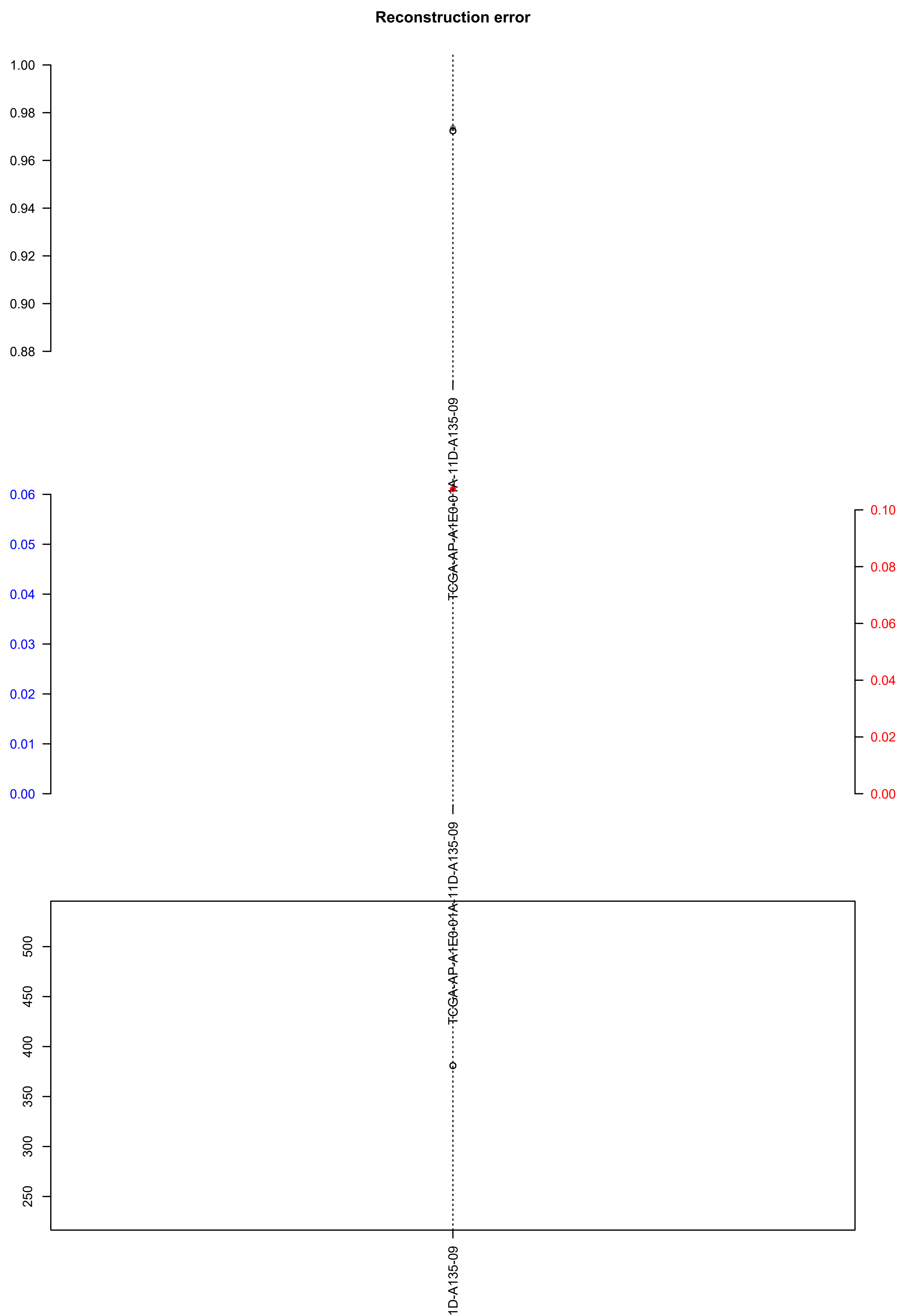

### TCGA-AX-A1CE-01A-11D-A135-09.check.with.sig.pdf

#### TCGA.AX.A1CE.01A.11D.A135.09

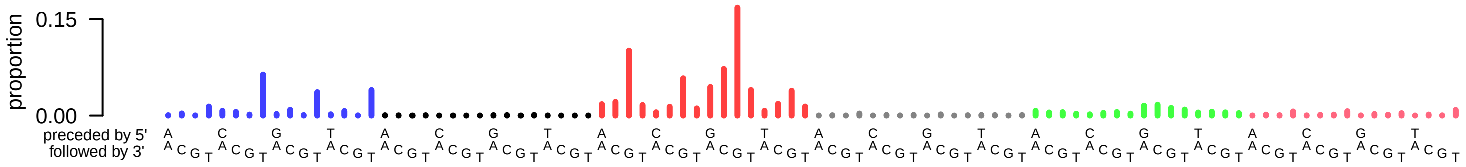

### TCGA-AX-A1CE-01A-11D-A135-09.exposures.pdf

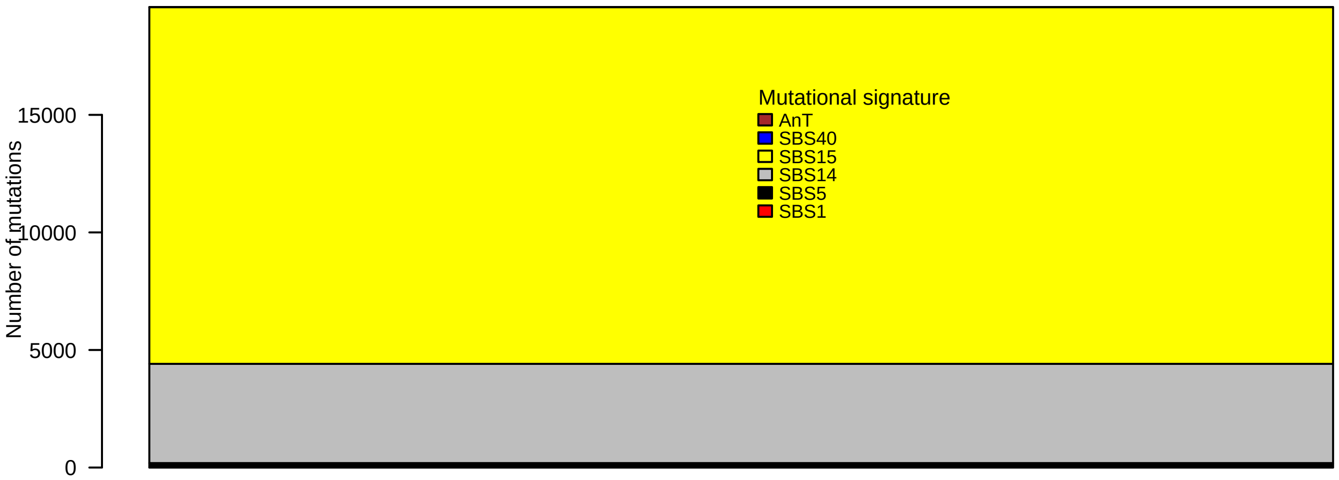

### TCGA-AX-A1CE-01A-11D-A135-09.pval.histogram.pdf

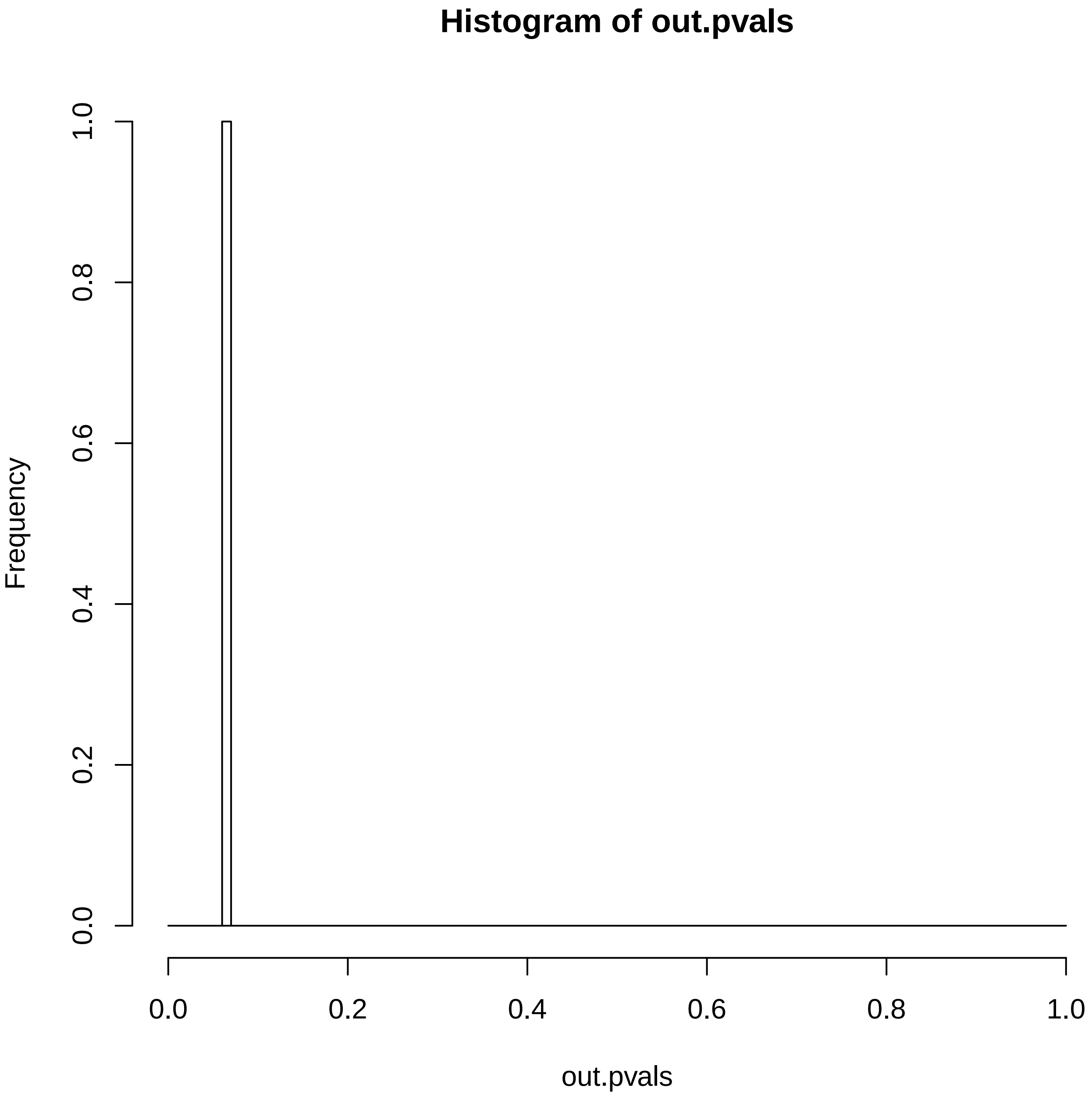

### TCGA-AX-A1CE-01A-11D-A135-09.reconstruction.err.pdf

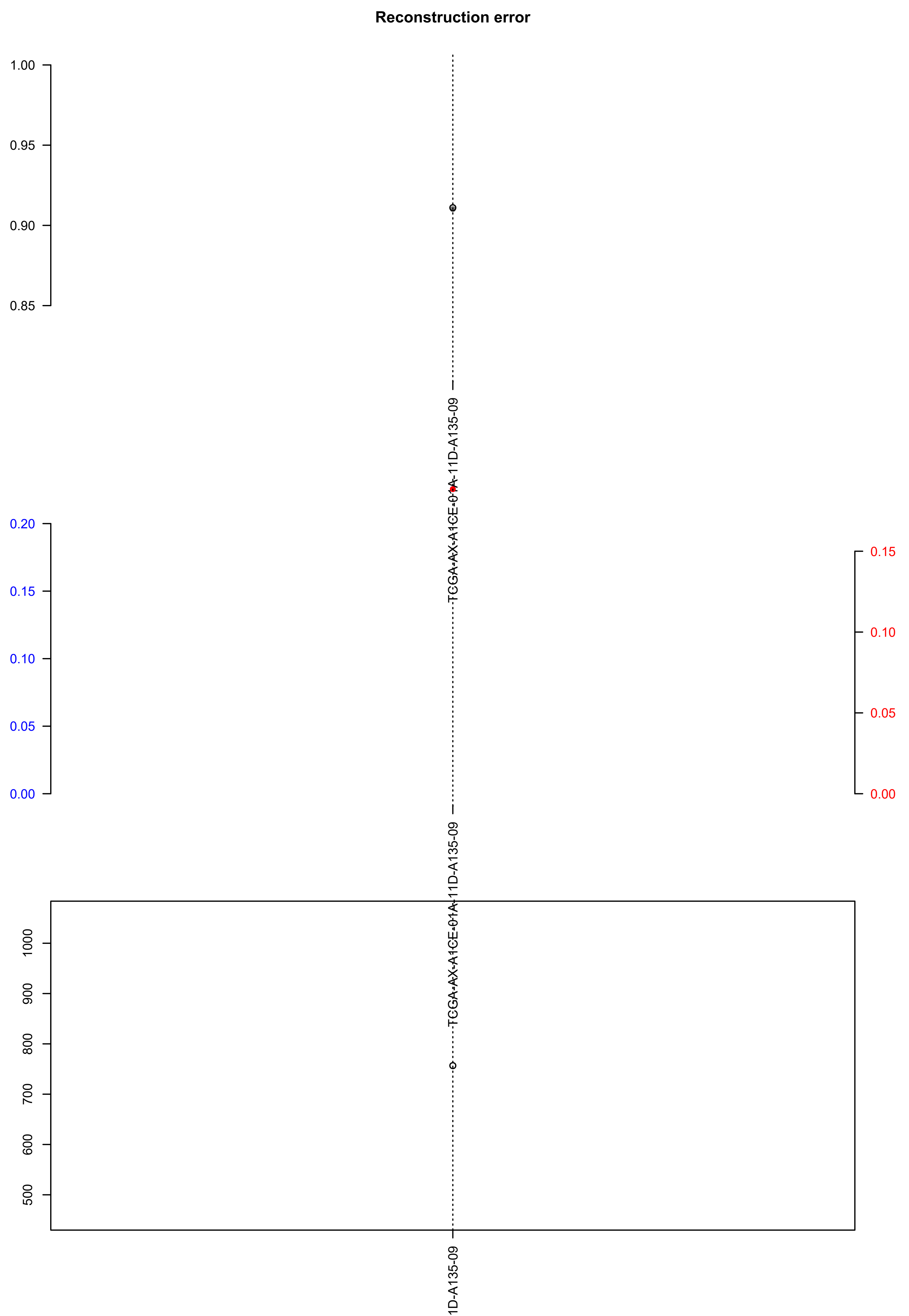

### TCGA-AY-4071-01A-01W-1073-09.check.with.sig.pdf

#### TCGA.AY.4071.01A.01W.1073.09

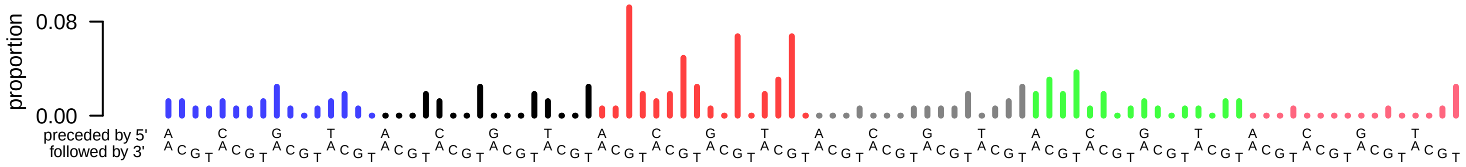

### TCGA-AY-4071-01A-01W-1073-09.exposures.pdf

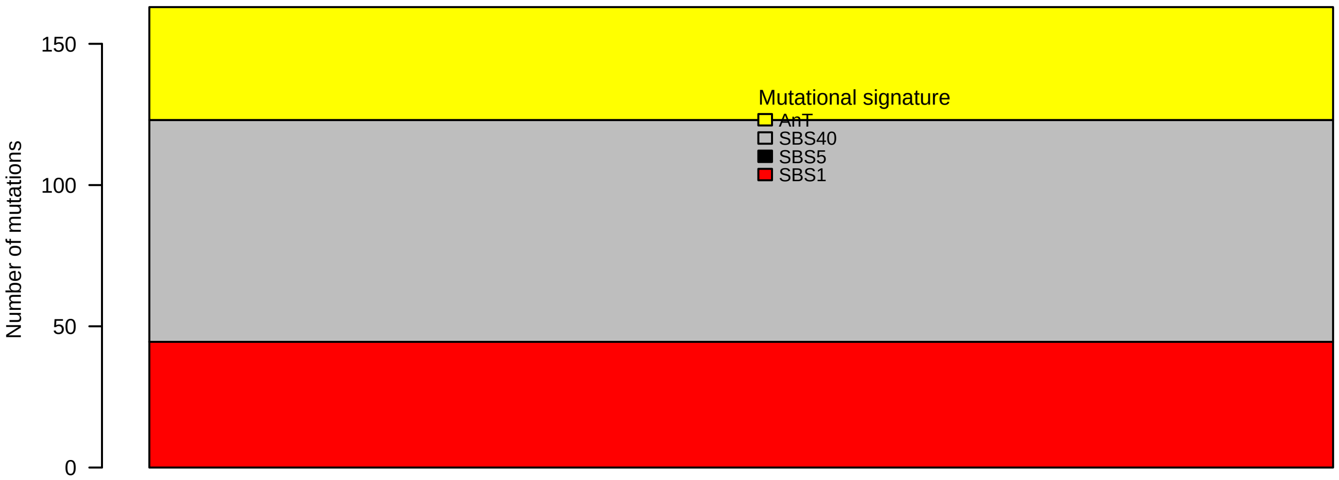

### TCGA-AY-4071-01A-01W-1073-09.pval.histogram.pdf

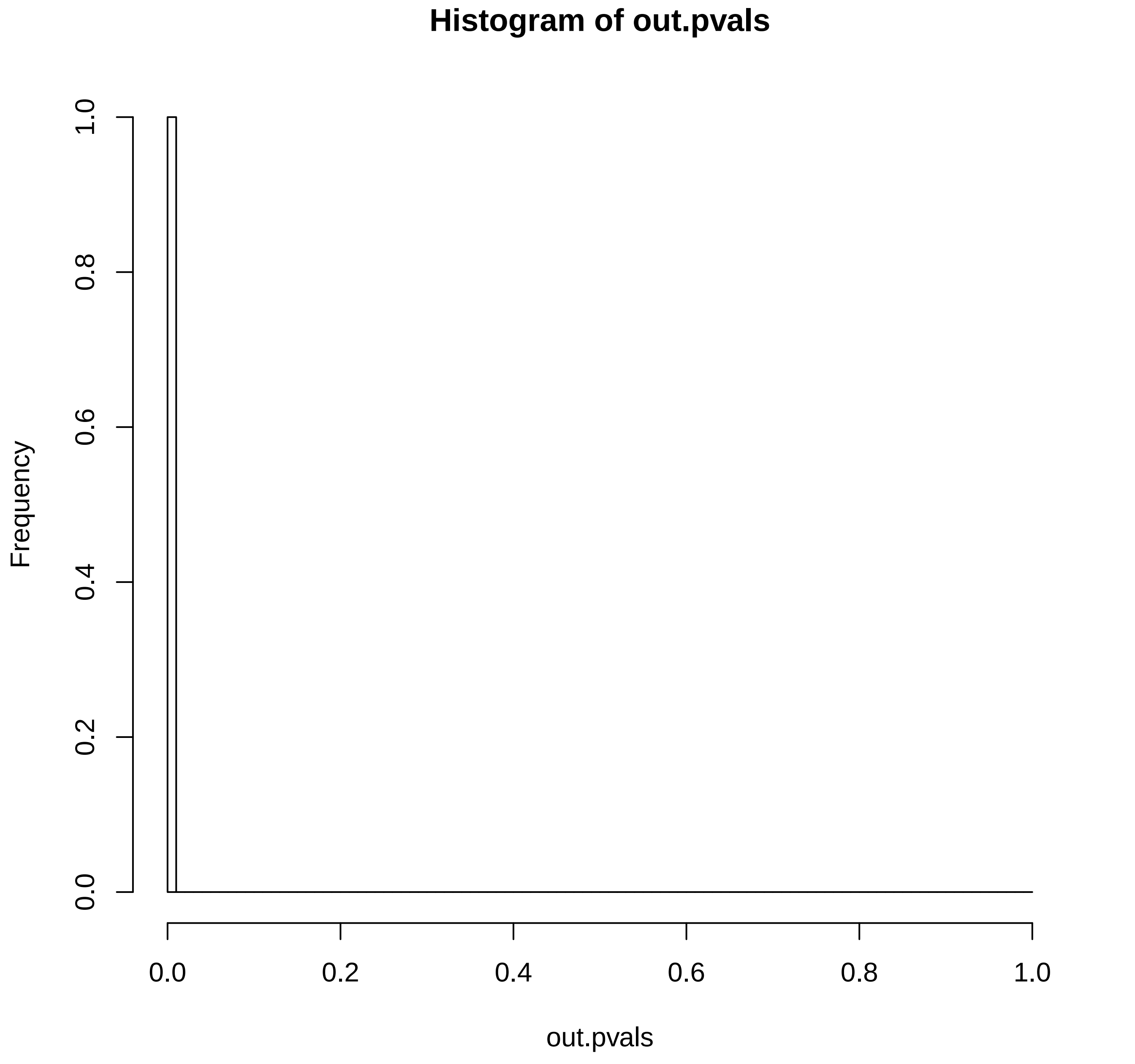

### TCGA-AY-4071-01A-01W-1073-09.reconstruction.err.pdf

Reconstruction error

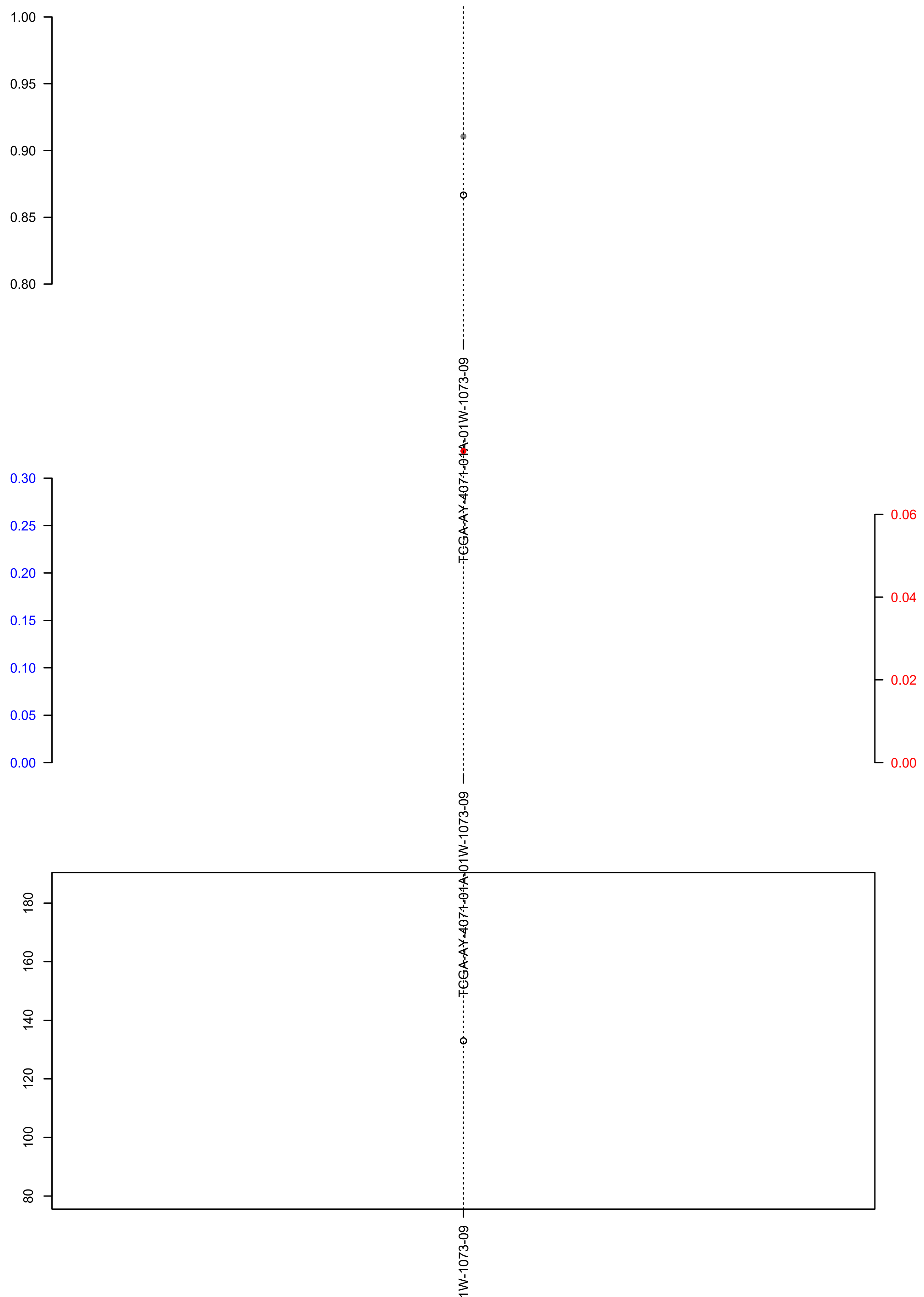

### TCGA-BA-A4IG-01A-11D-A25Y-08.check.with.sig.pdf

#### TCGA.BA.A4IG.01A.11D.A25Y.08

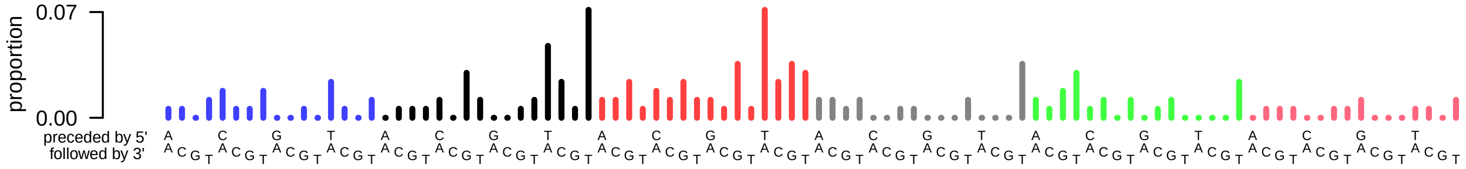

### TCGA-BA-A4IG-01A-11D-A25Y-08.exposures.pdf

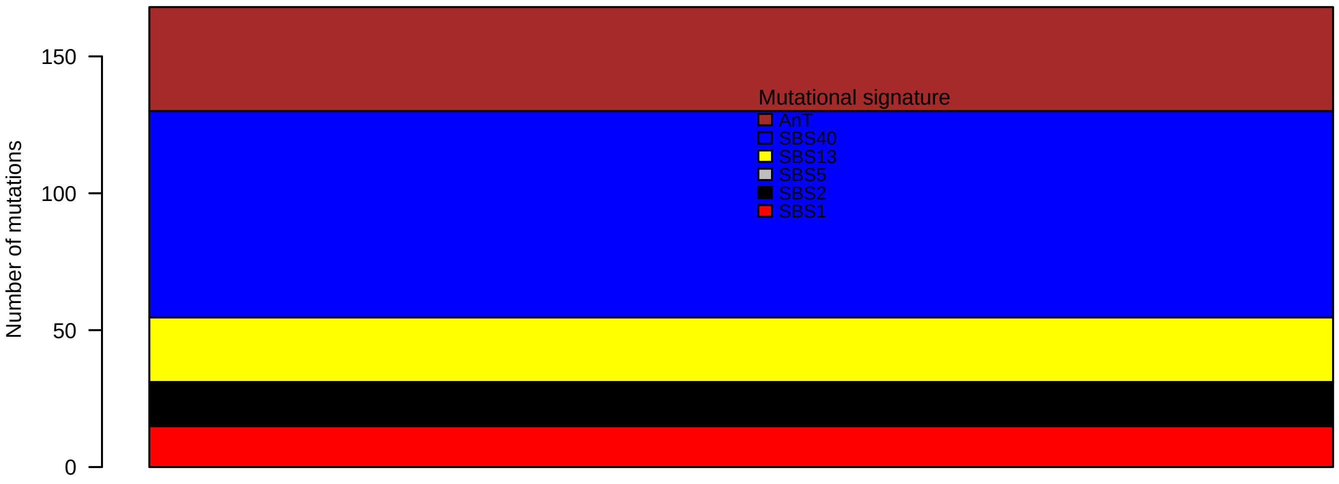

### TCGA-BA-A4IG-01A-11D-A25Y-08.pval.histogram.pdf

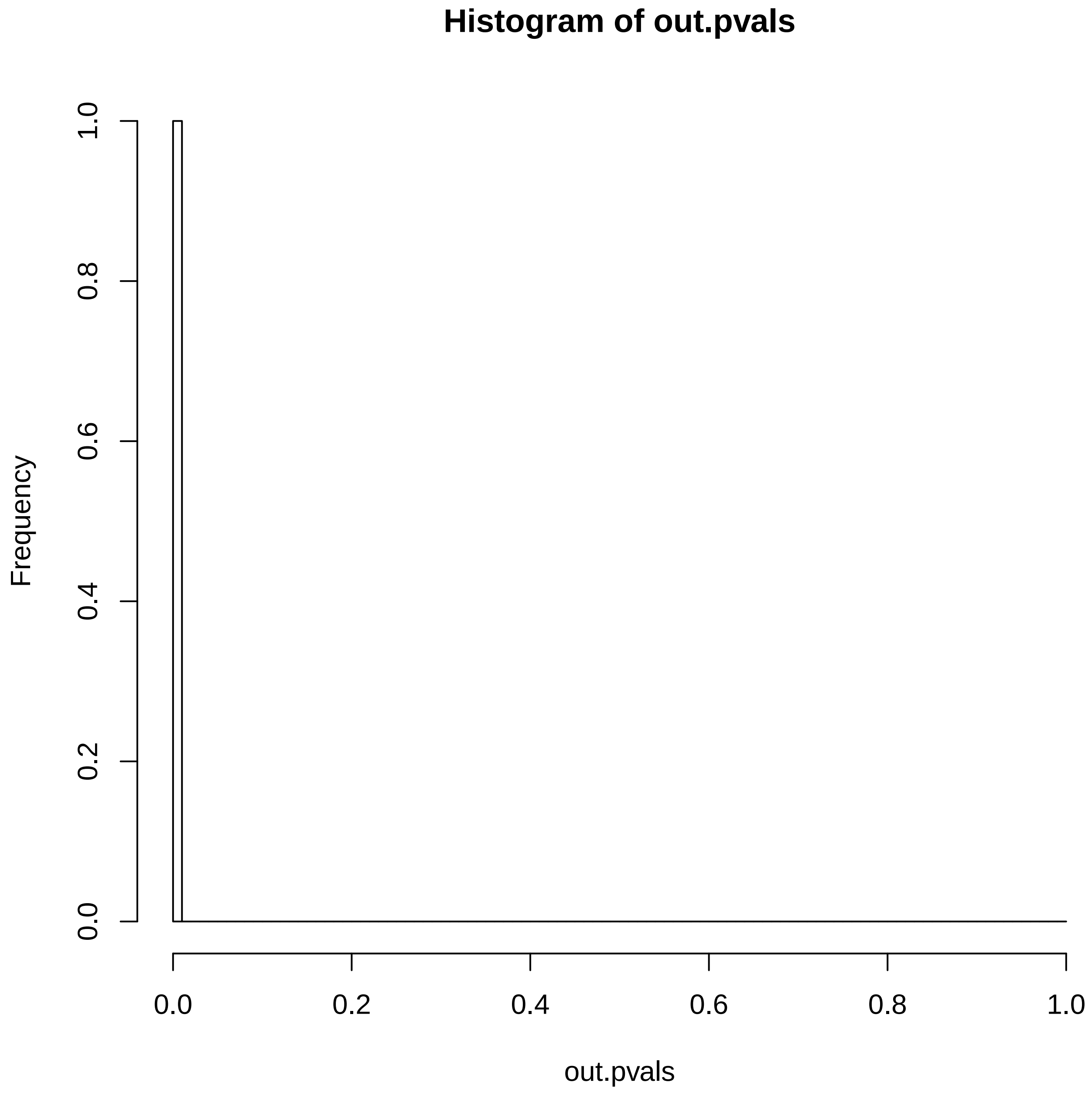

### TCGA-BA-A4IG-01A-11D-A25Y-08.reconstruction.err.pdf

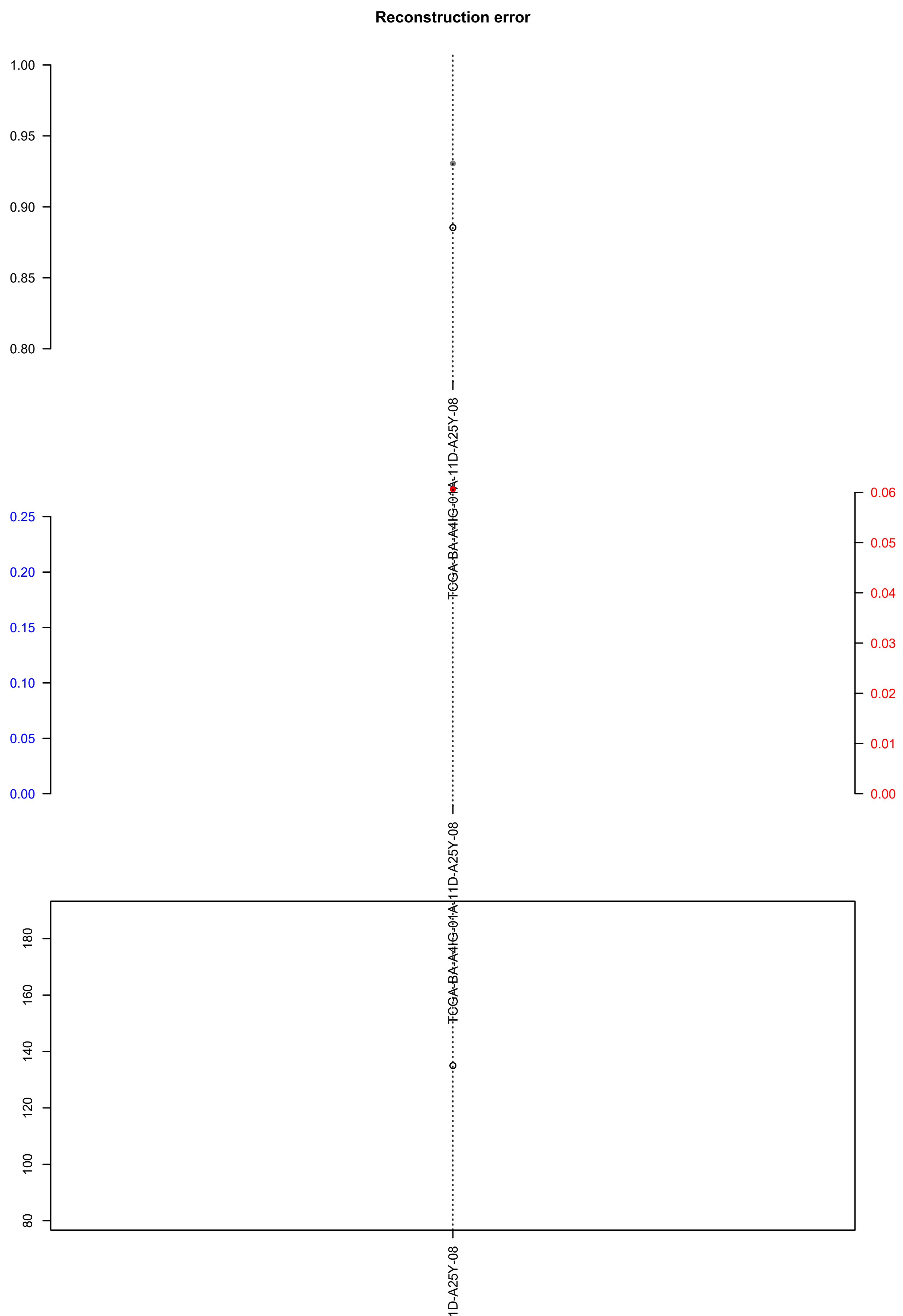

### TCGA-BK-A6W3-01A-12D-A34Q-09.check.with.sig.pdf

#### TCGA.BK.A6W3.01A.12D.A34Q.09

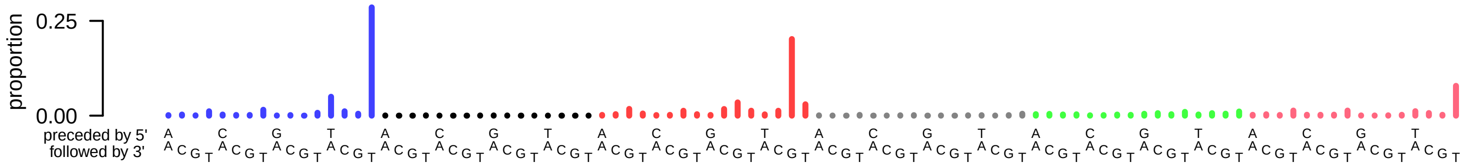

### TCGA-BK-A6W3-01A-12D-A34Q-09.exposures.pdf

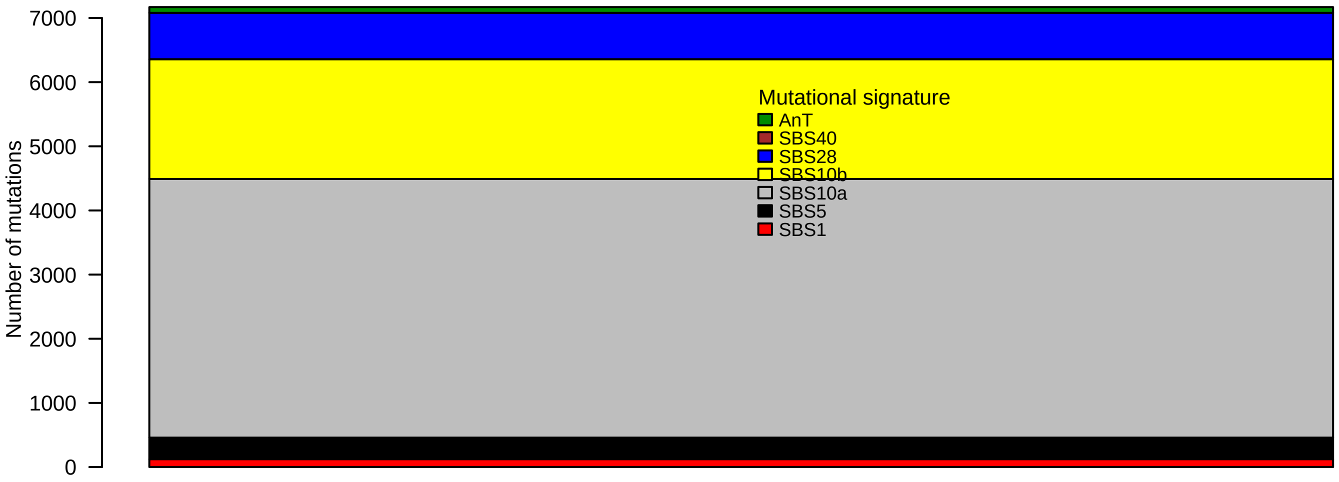

### TCGA-BK-A6W3-01A-12D-A34Q-09.pval.histogram.pdf

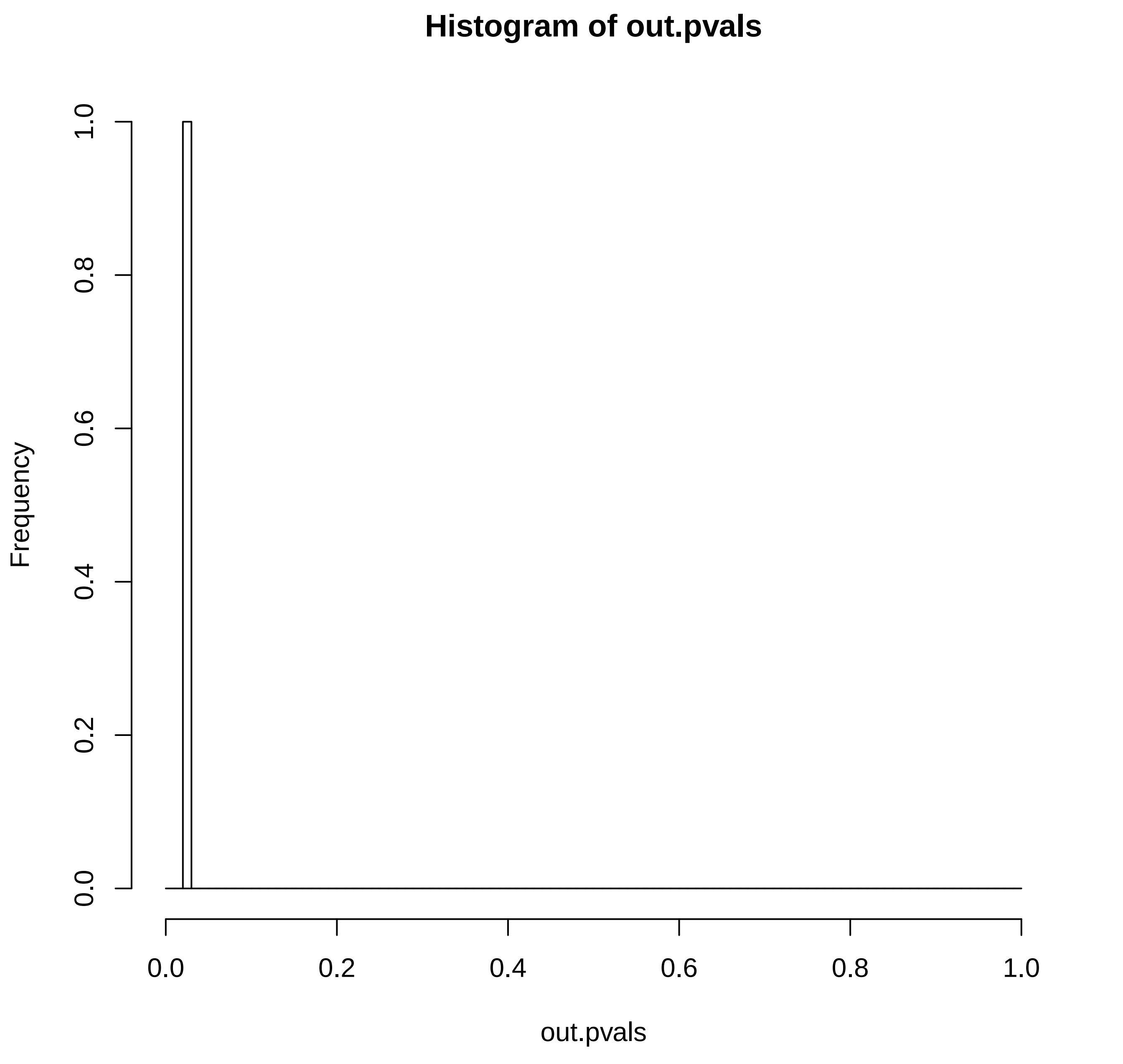

### TCGA-BK-A6W3-01A-12D-A34Q-09.reconstruction.err.pdf

**Reconstruction error**

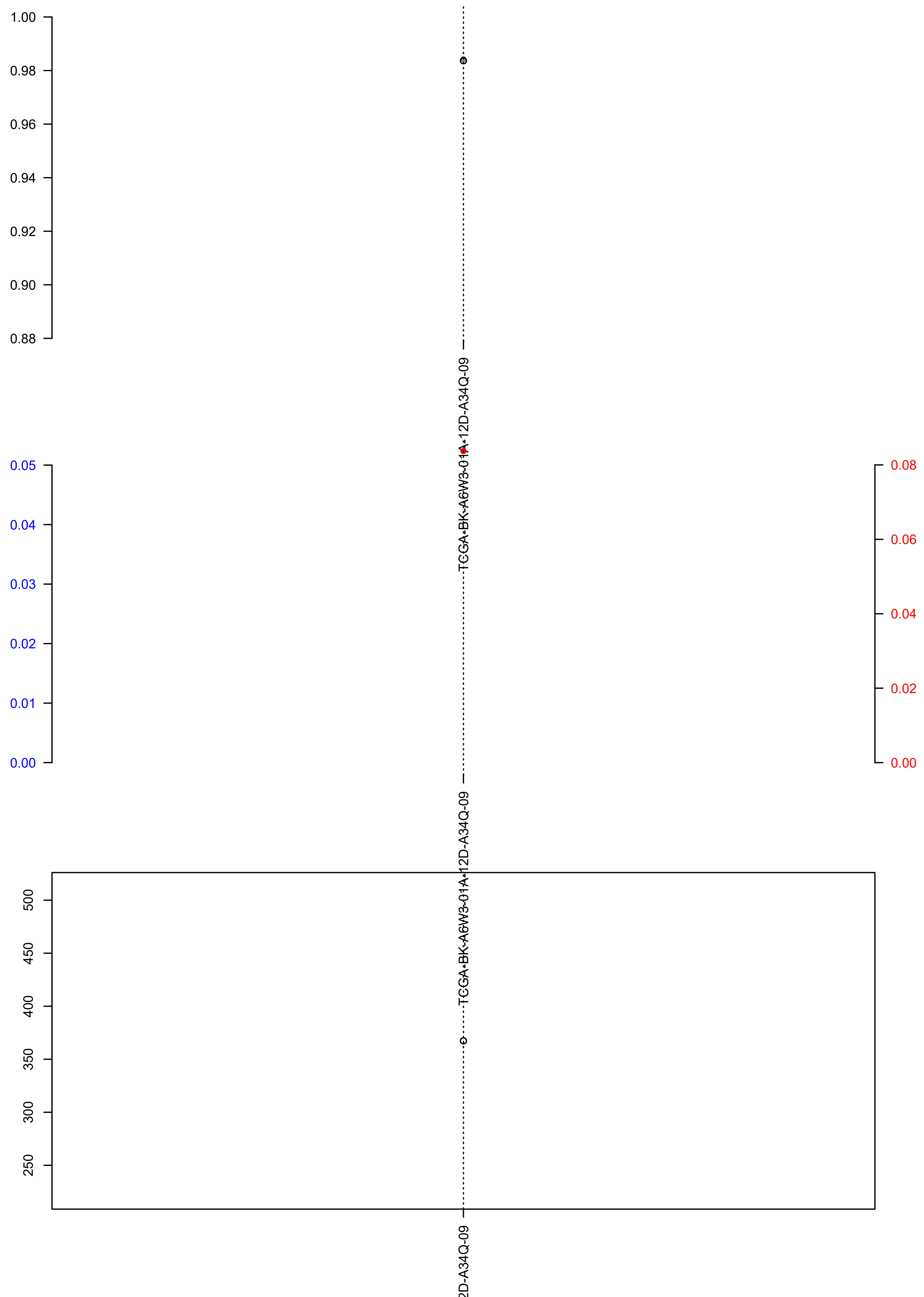

### TCGA-BR-6453-01A-11D-1800-08.exposures.pdf

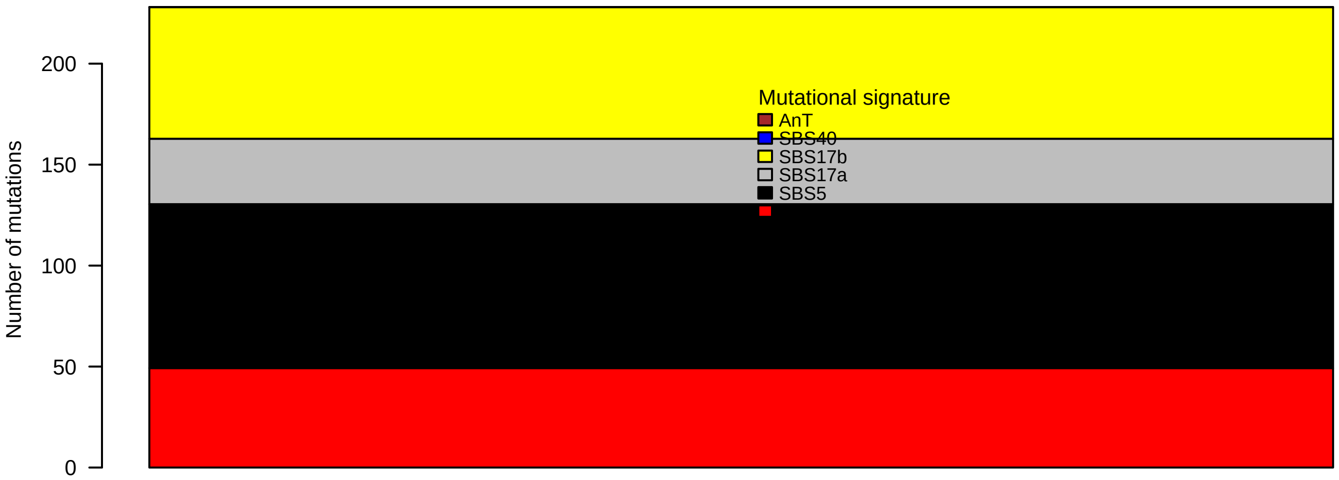

### TCGA-BR-6453-01A-11D-1800-08.pval.histogram.pdf

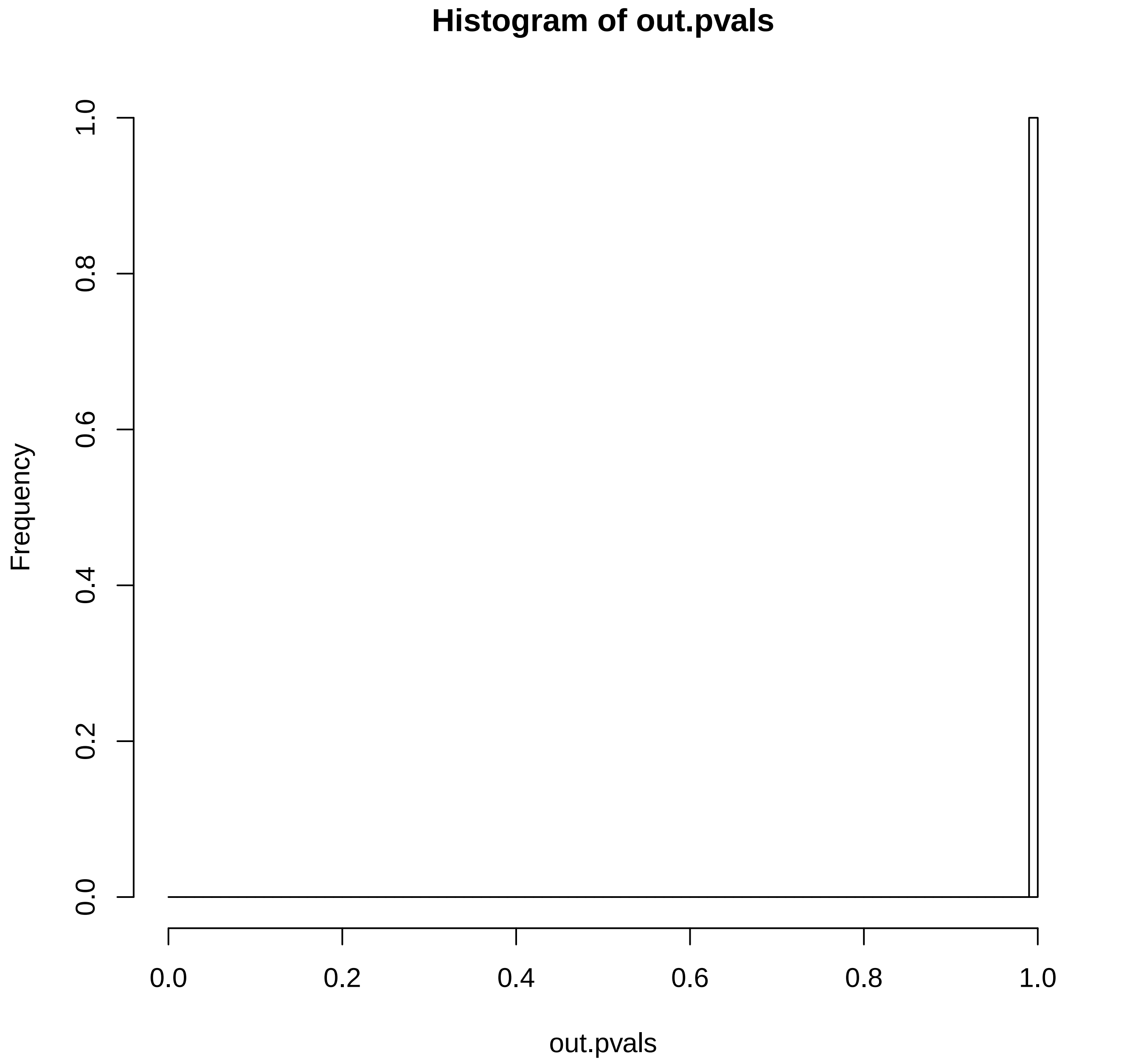

### TCGA-BR-6453-01A-11D-1800-08.reconstruction.err.pdf

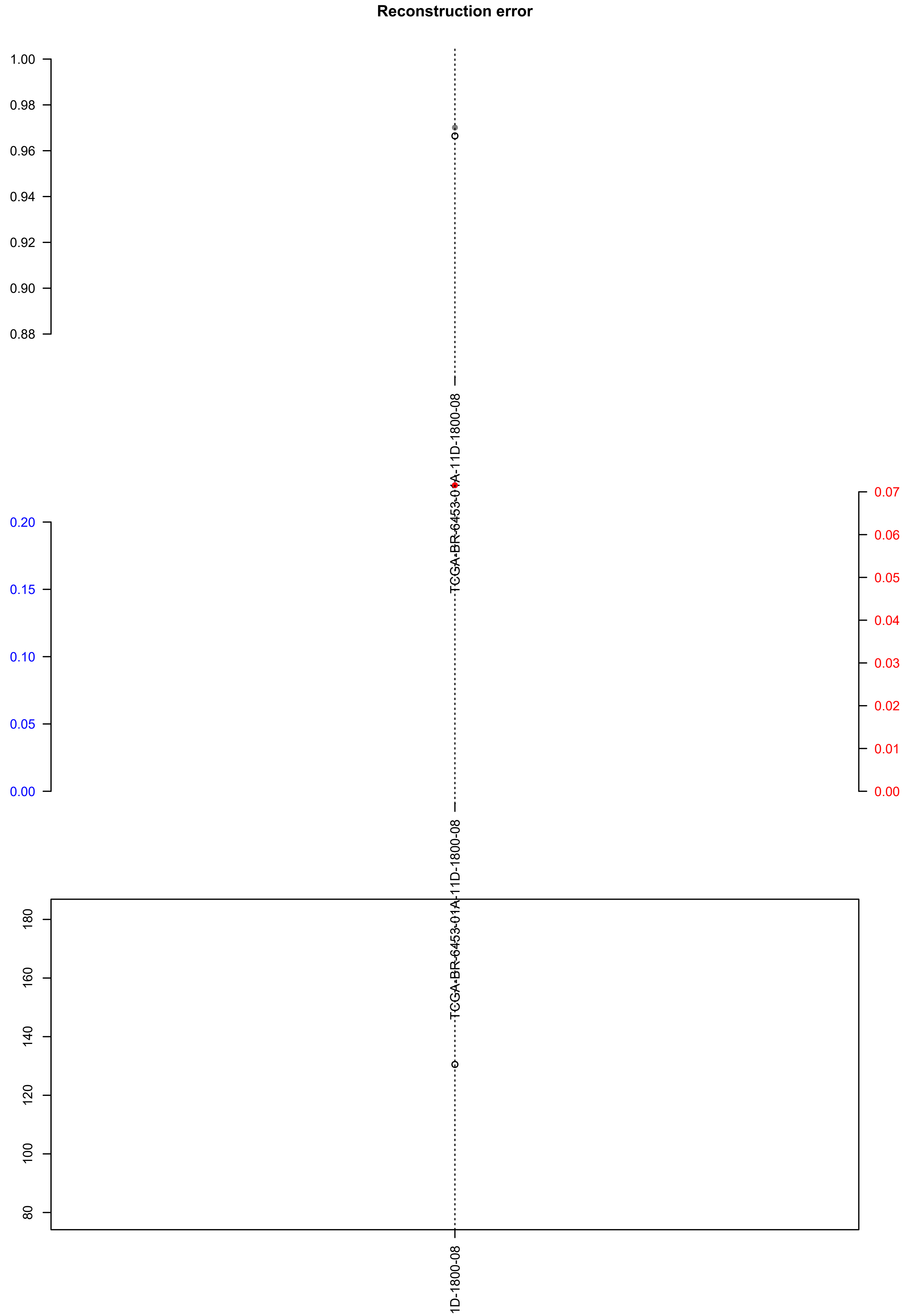

### TCGA-D7-A4Z0-01A-22D-A25D-08.exposures.pdf

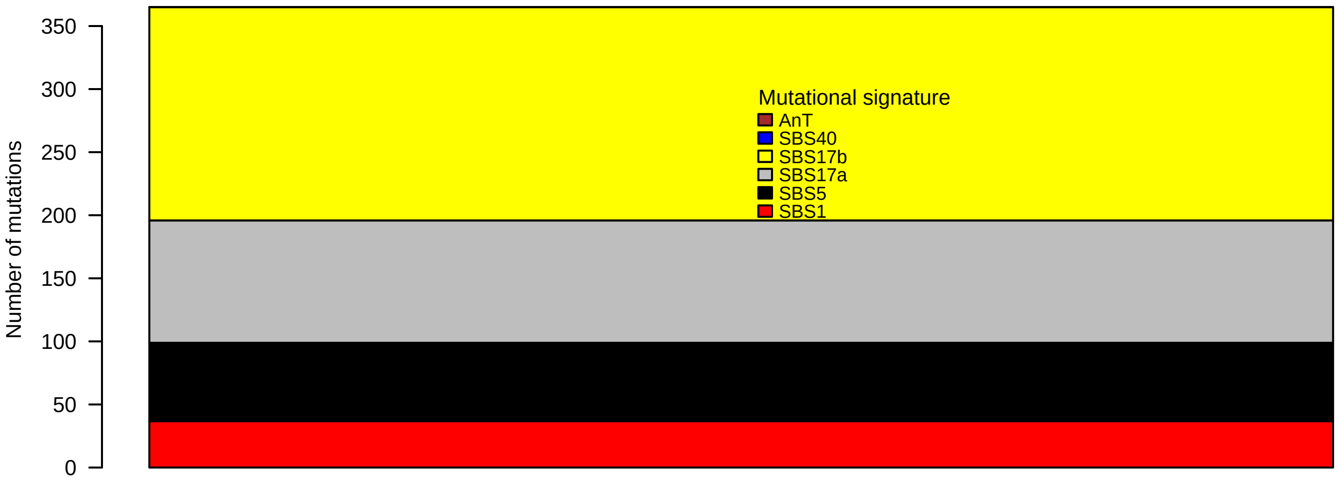

### TCGA-D7-A4Z0-01A-22D-A25D-08.pval.histogram.pdf

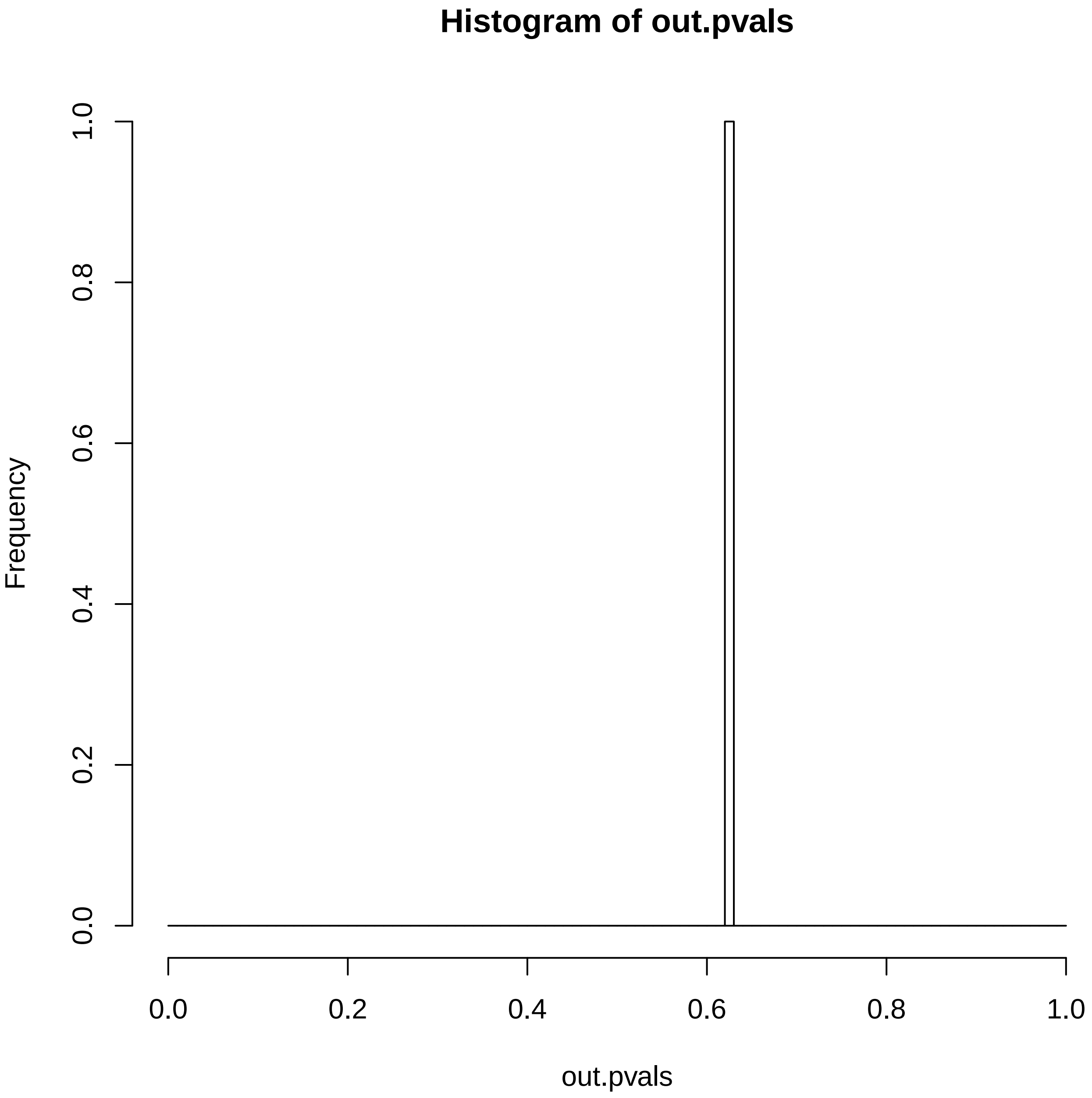

### TCGA-D7-A4Z0-01A-22D-A25D-08.reconstruction.err.pdf

##### Reconstruction error

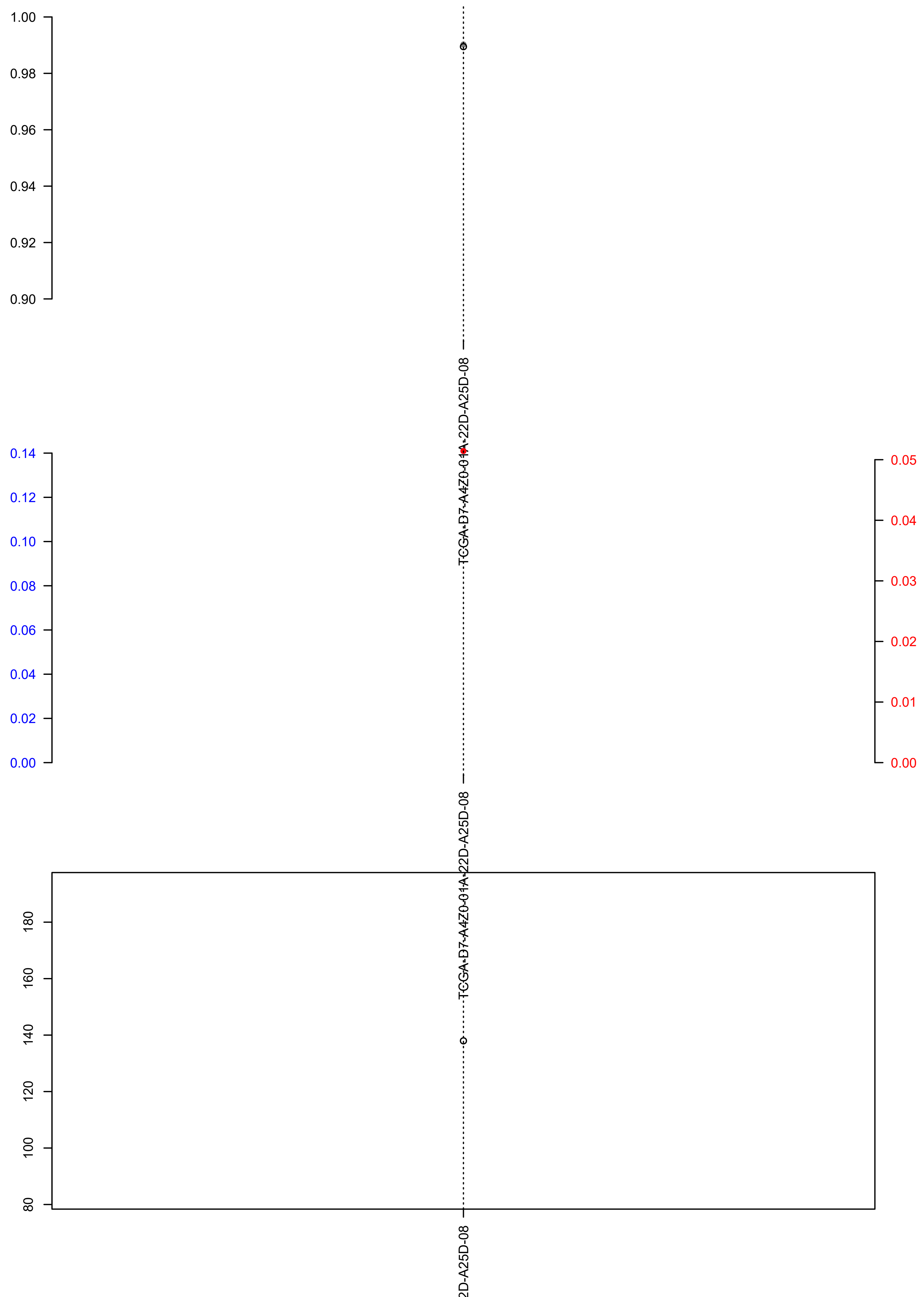

### TCGA-DF-A2KV-01A-11D-A17W-09.check.with.sig.pdf

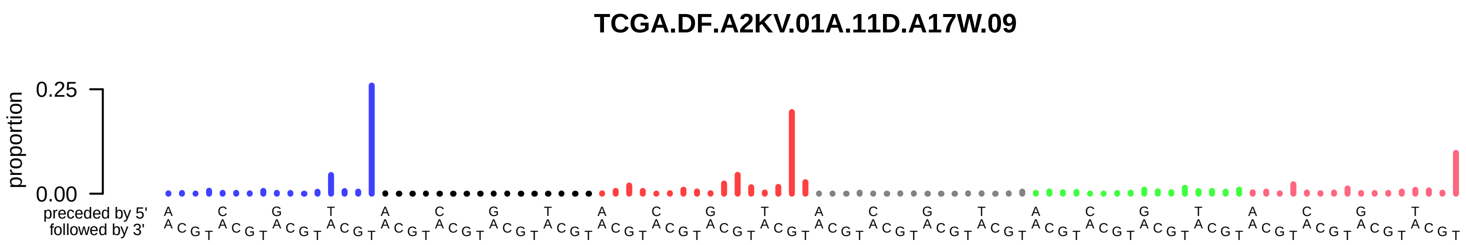

### TCGA-DF-A2KV-01A-11D-A17W-09.exposures.pdf

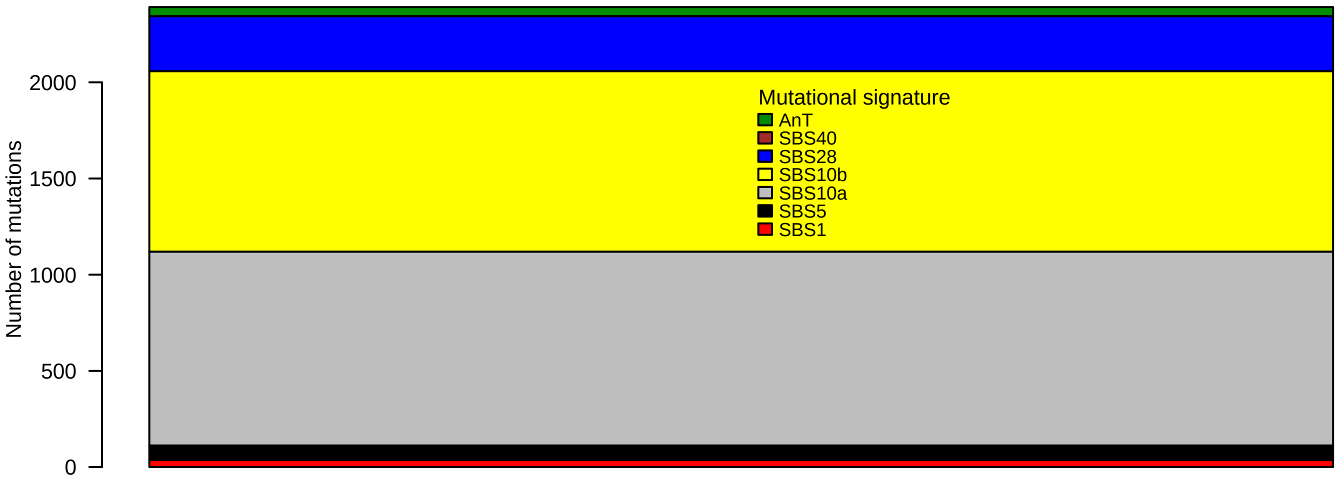

### TCGA-DF-A2KV-01A-11D-A17W-09.pval.histogram.pdf

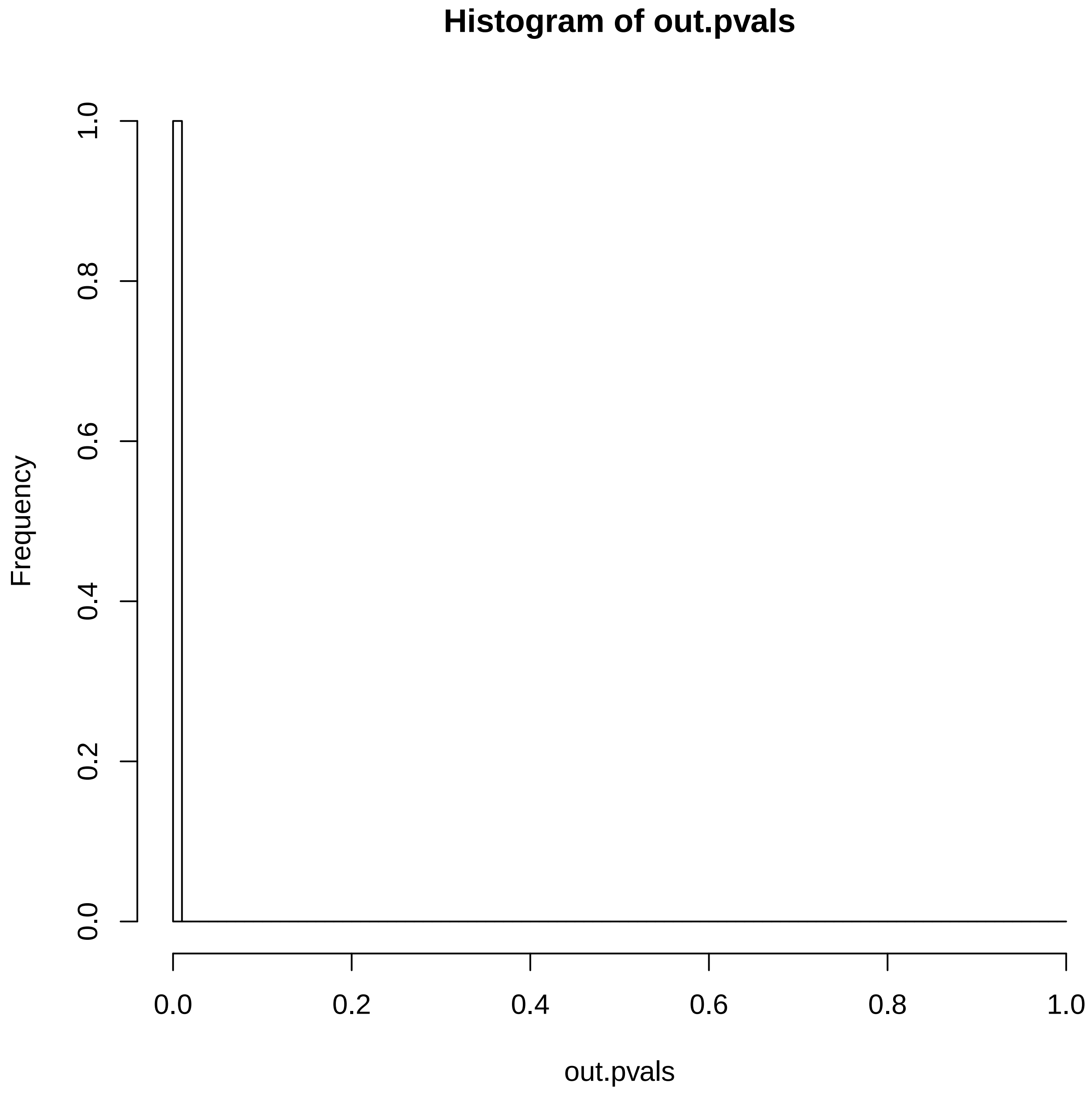

### TCGA-DF-A2KV-01A-11D-A17W-09.reconstruction.err.pdf

##### Reconstruction error

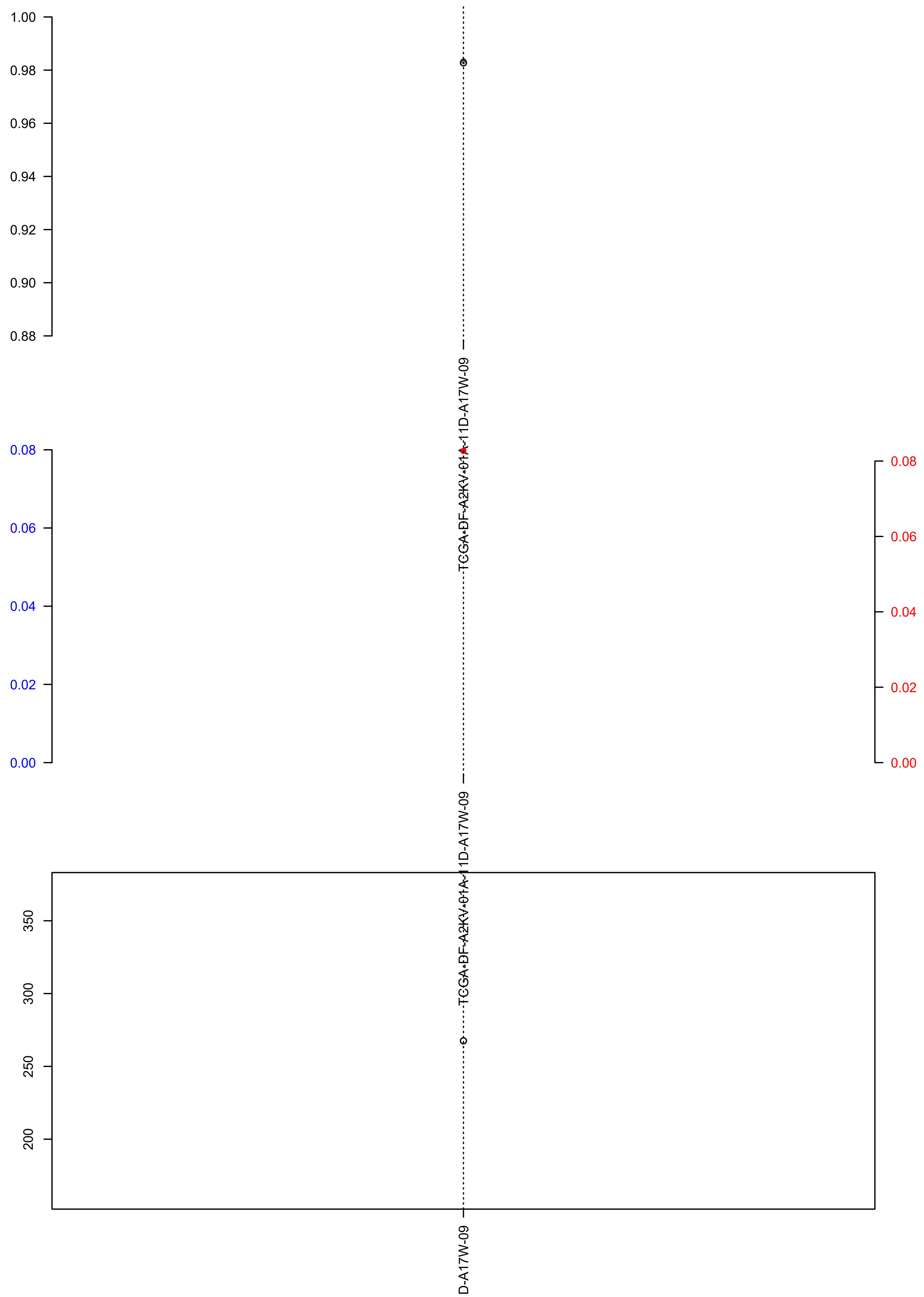

### TCGA-E6-A1M0-01A-11D-A142-09.exposures.pdf

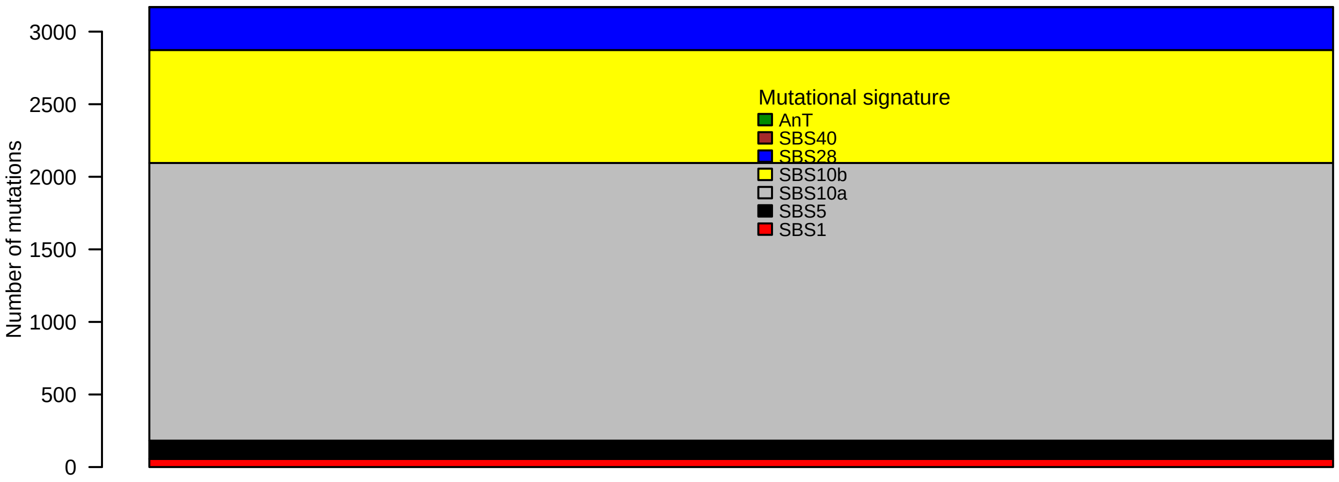

### TCGA-E6-A1M0-01A-11D-A142-09.pval.histogram.pdf

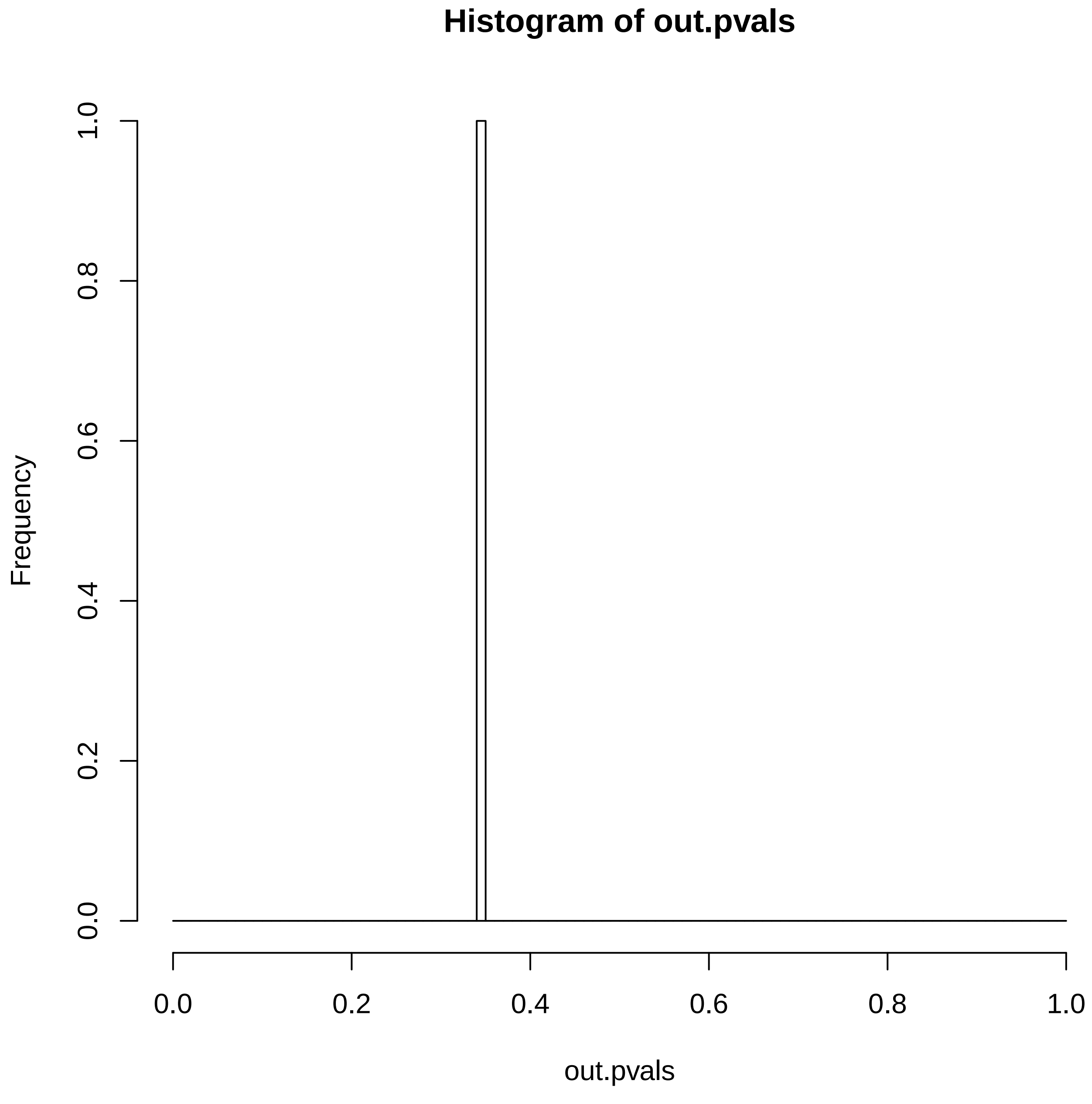

### TCGA-E6-A1M0-01A-11D-A142-09.reconstruction.err.pdf

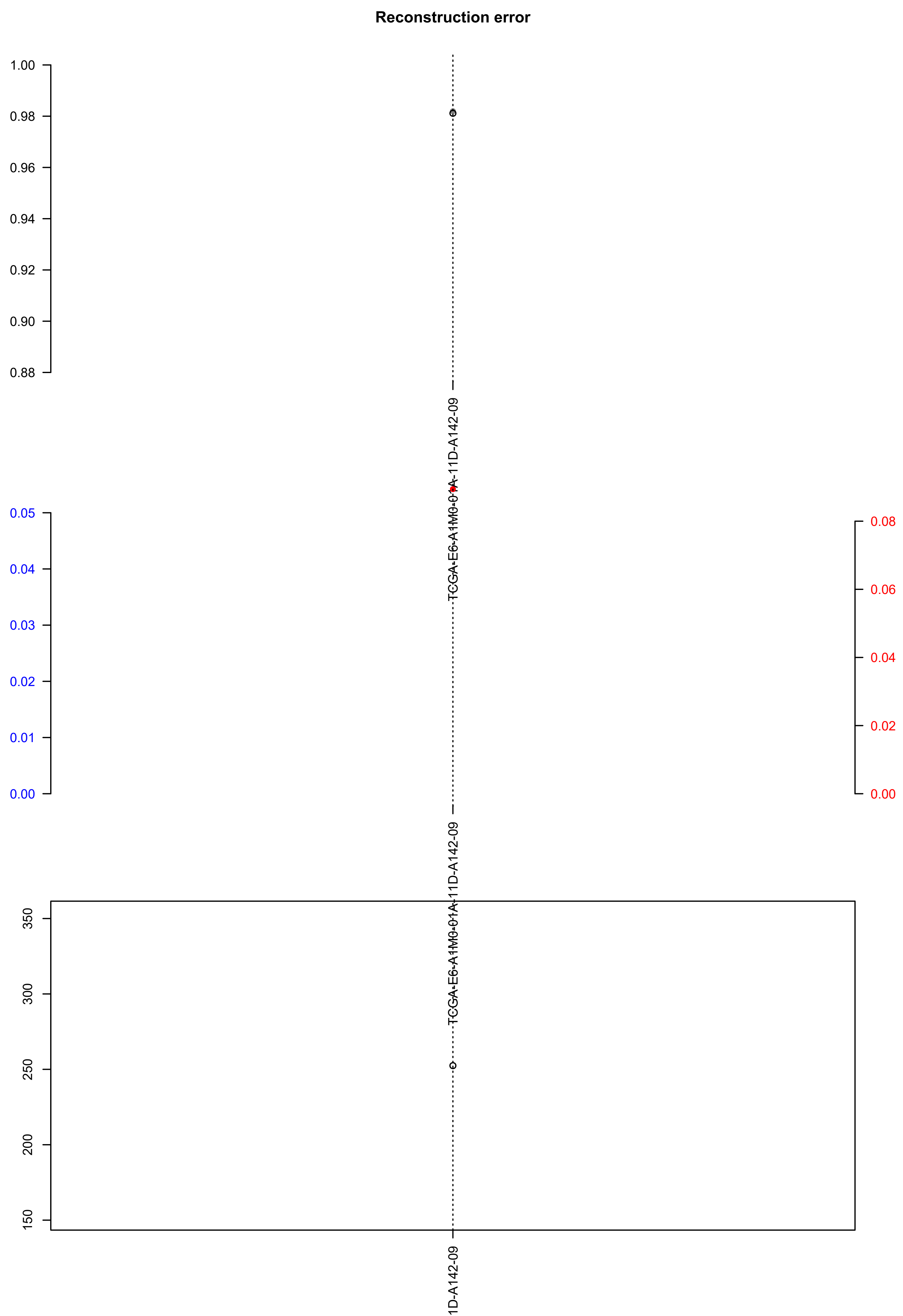

### TCGA-EO-A3AV-01A-12D-A19Y-09.check.with.sig.pdf

#### TCGA.EO.A3AV.01A.12D.A19Y.09

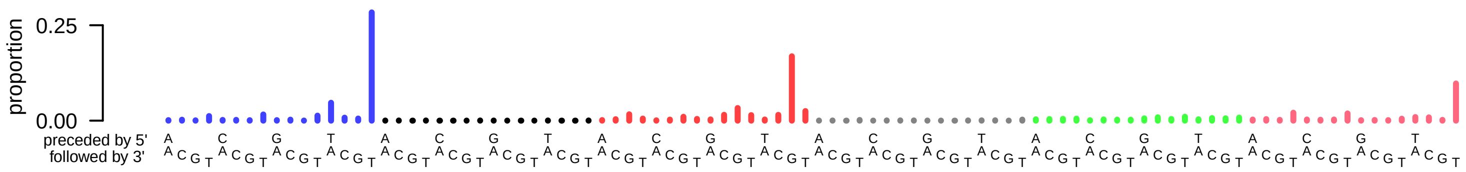

### TCGA-EO-A3AV-01A-12D-A19Y-09.exposures.pdf

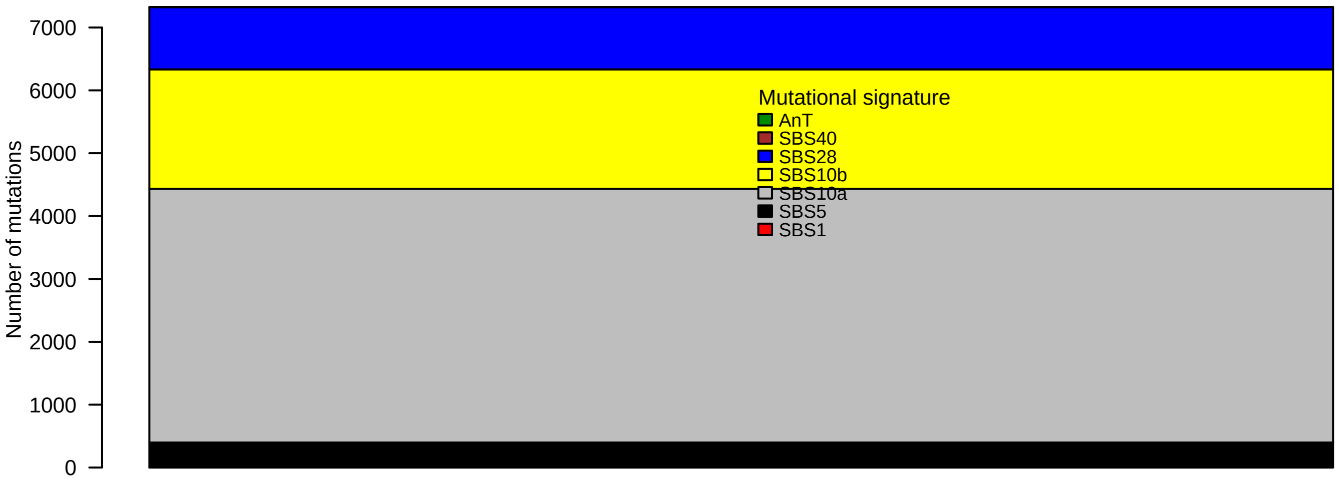

### TCGA-EO-A3AV-01A-12D-A19Y-09.pval.histogram.pdf

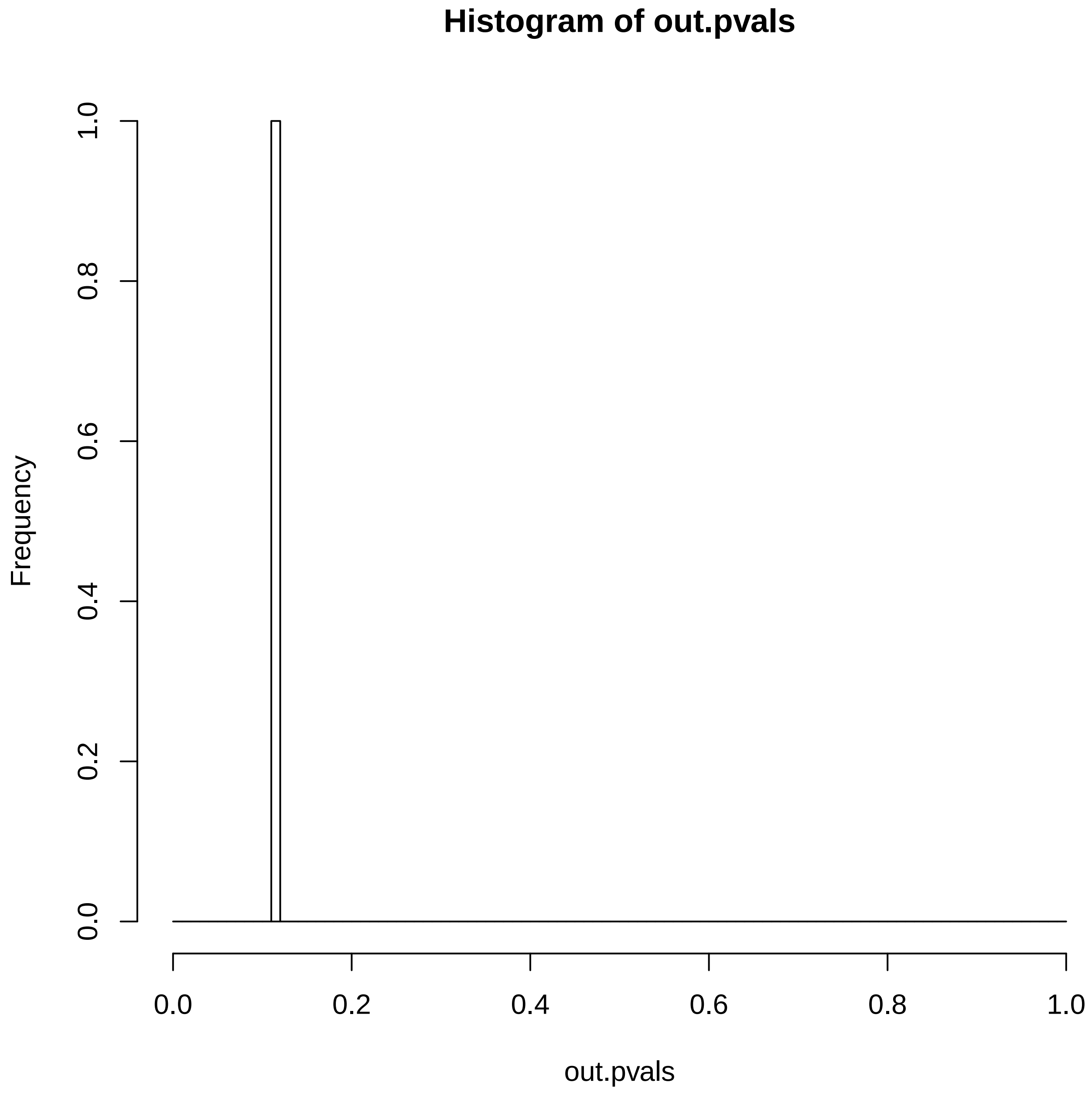

### TCGA-EO-A3AV-01A-12D-A19Y-09.reconstruction.err.pdf

##### Reconstruction error

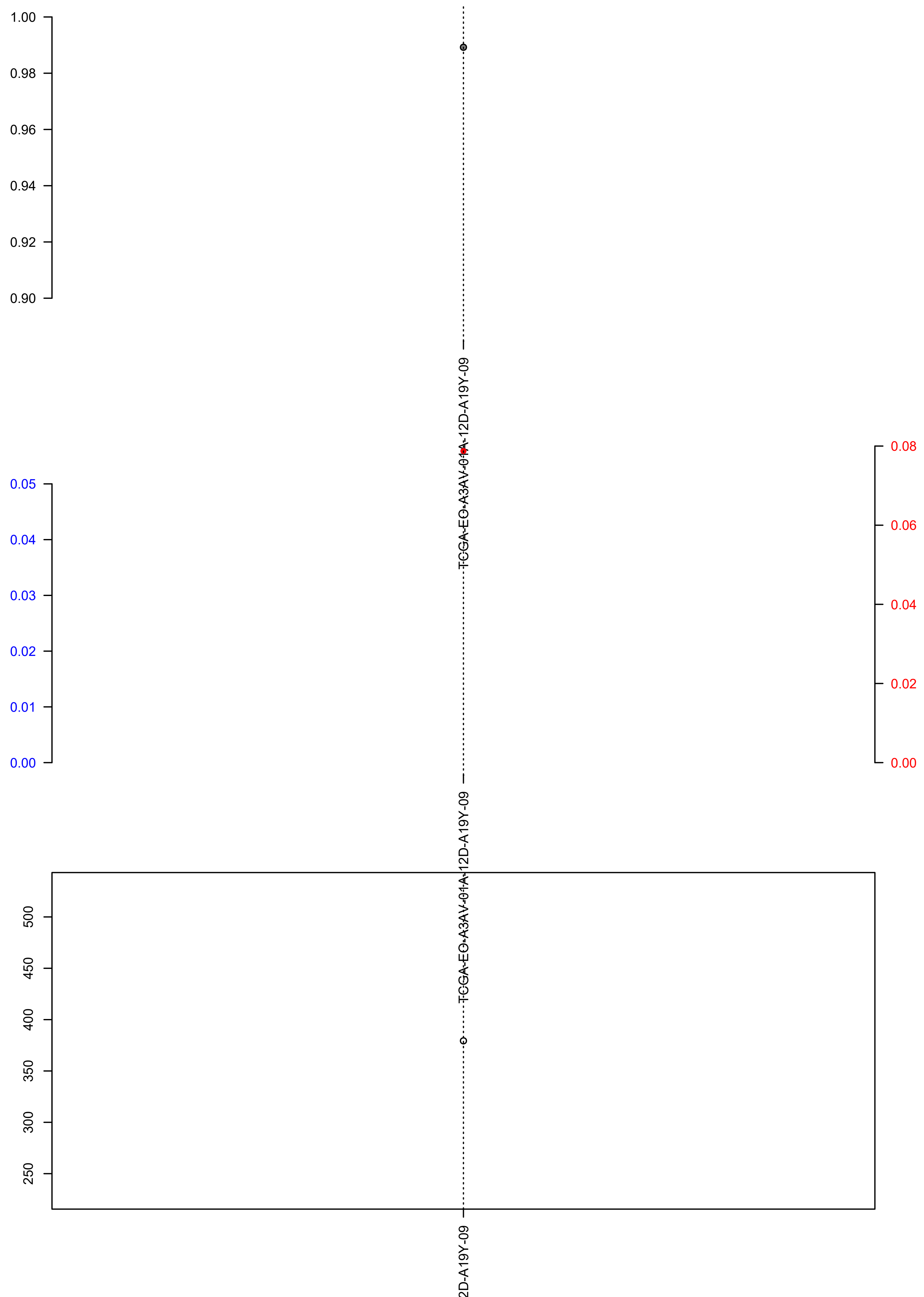

### TCGA-EO-A3AY-01A-12D-A19Y-09.check.with.sig.pdf

#### TCGA.EO.A3AY.01A.12D.A19Y.09

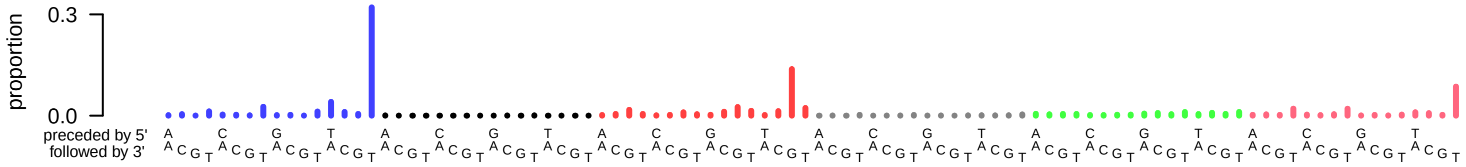

### TCGA-EO-A3AY-01A-12D-A19Y-09.exposures.pdf

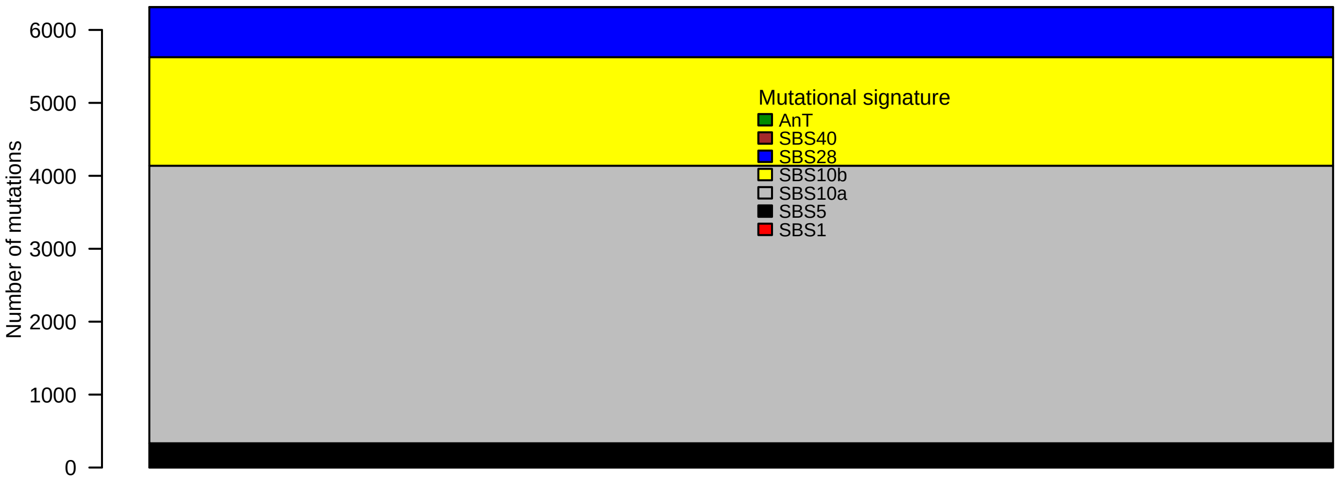

### TCGA-EO-A3AY-01A-12D-A19Y-09.pval.histogram.pdf

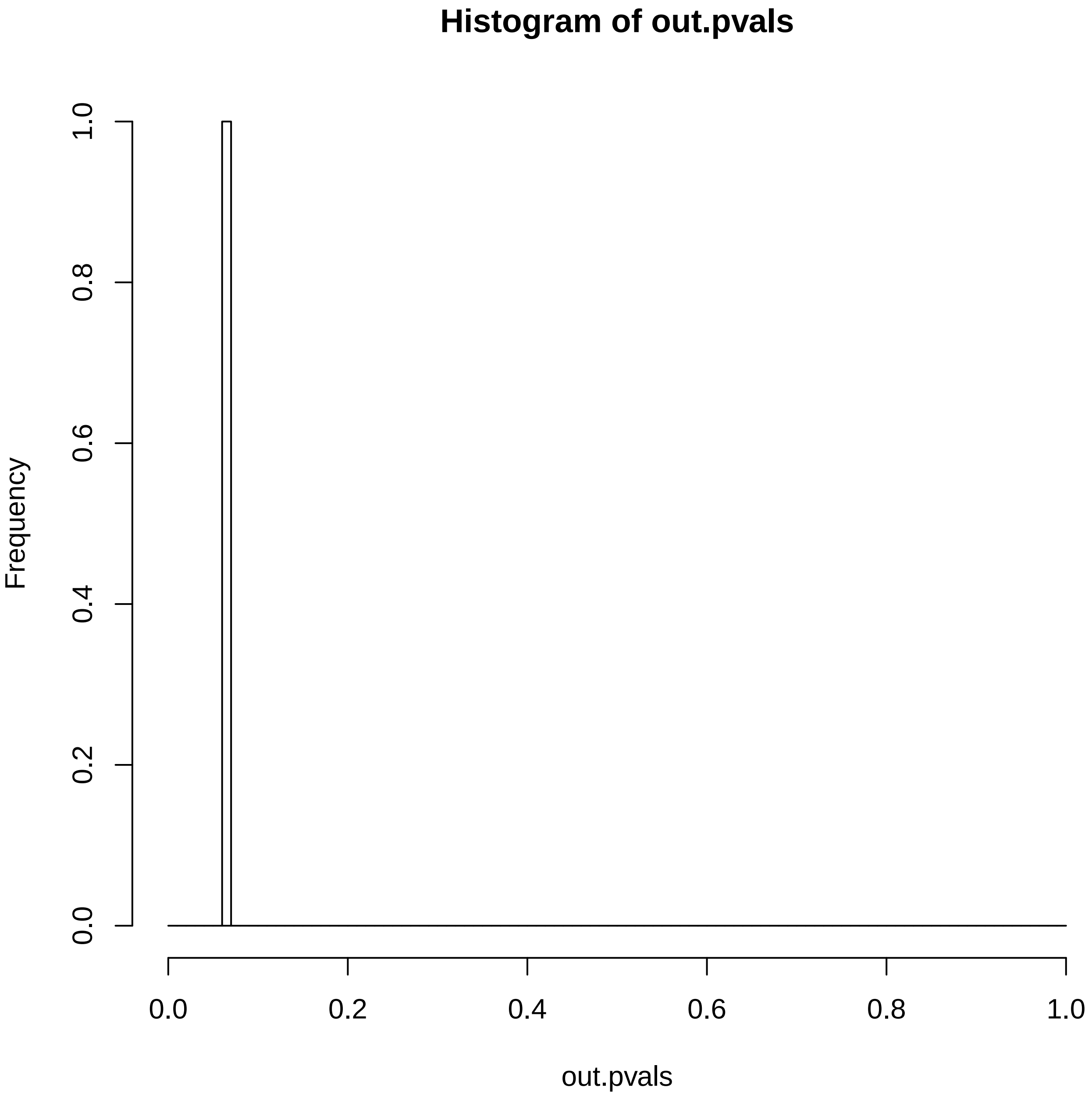

### TCGA-EO-A3AY-01A-12D-A19Y-09.reconstruction.err.pdf

##### Reconstruction error

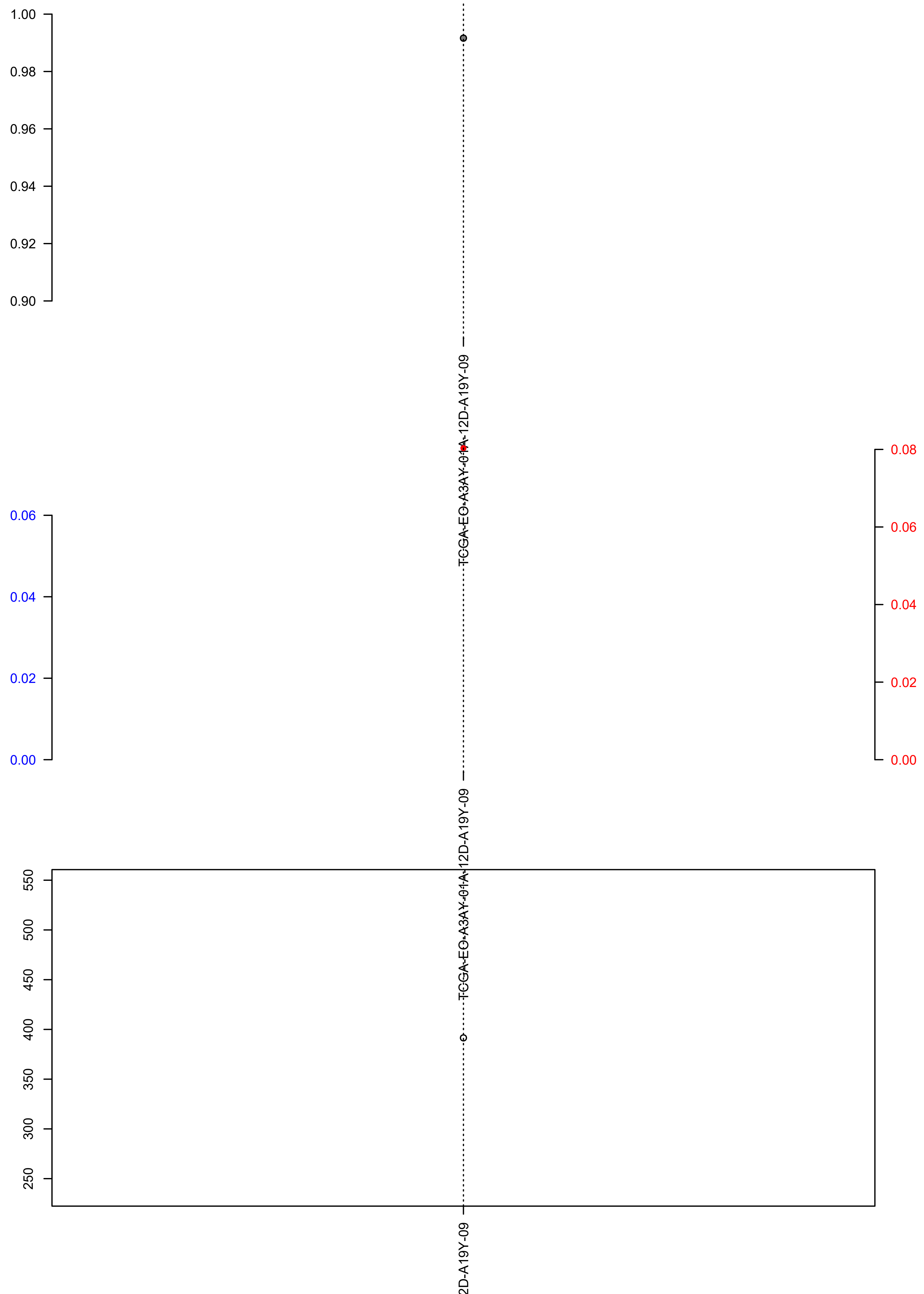

### TCGA-EY-A1GD-01A-11D-A13L-09.check.with.sig.pdf

##### TCGA.EY.A1GD.01A.11D.A13L.09

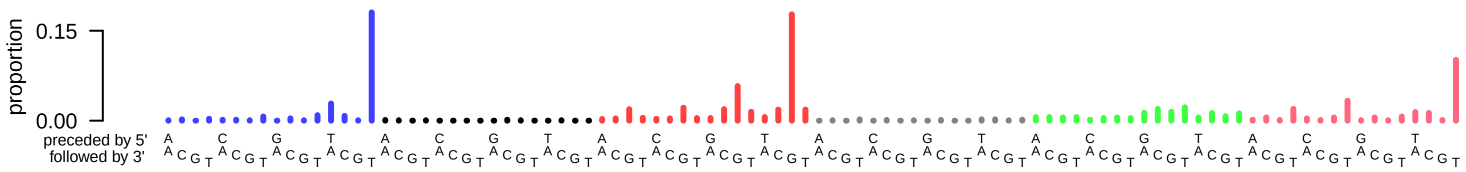

### TCGA-EY-A1GD-01A-11D-A13L-09.exposures.pdf

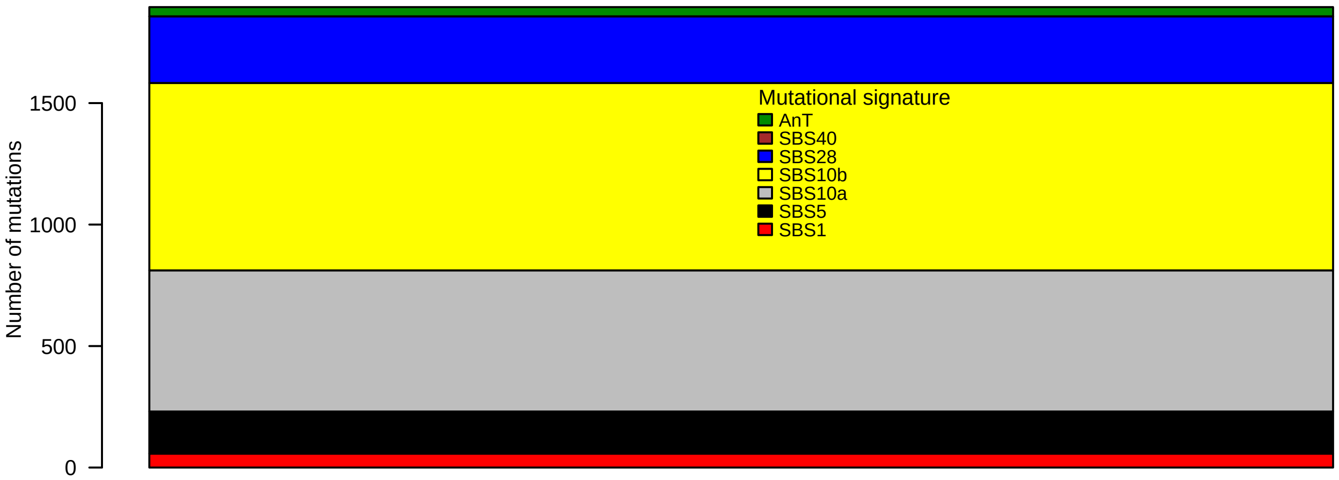

### TCGA-EY-A1GD-01A-11D-A13L-09.pval.histogram.pdf

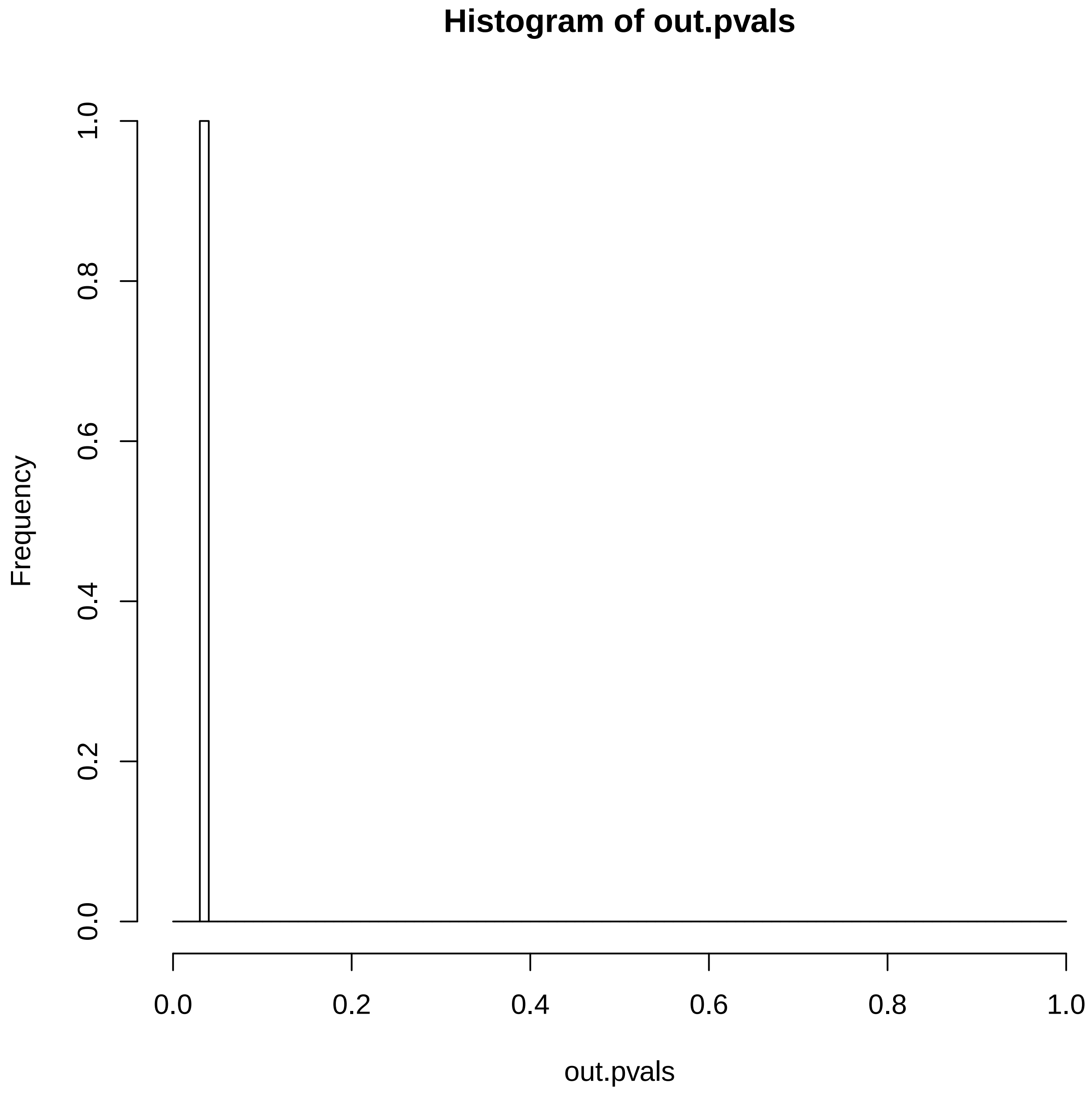

### TCGA-EY-A1GD-01A-11D-A13L-09.reconstruction.err.pdf

**Reconstruction error**

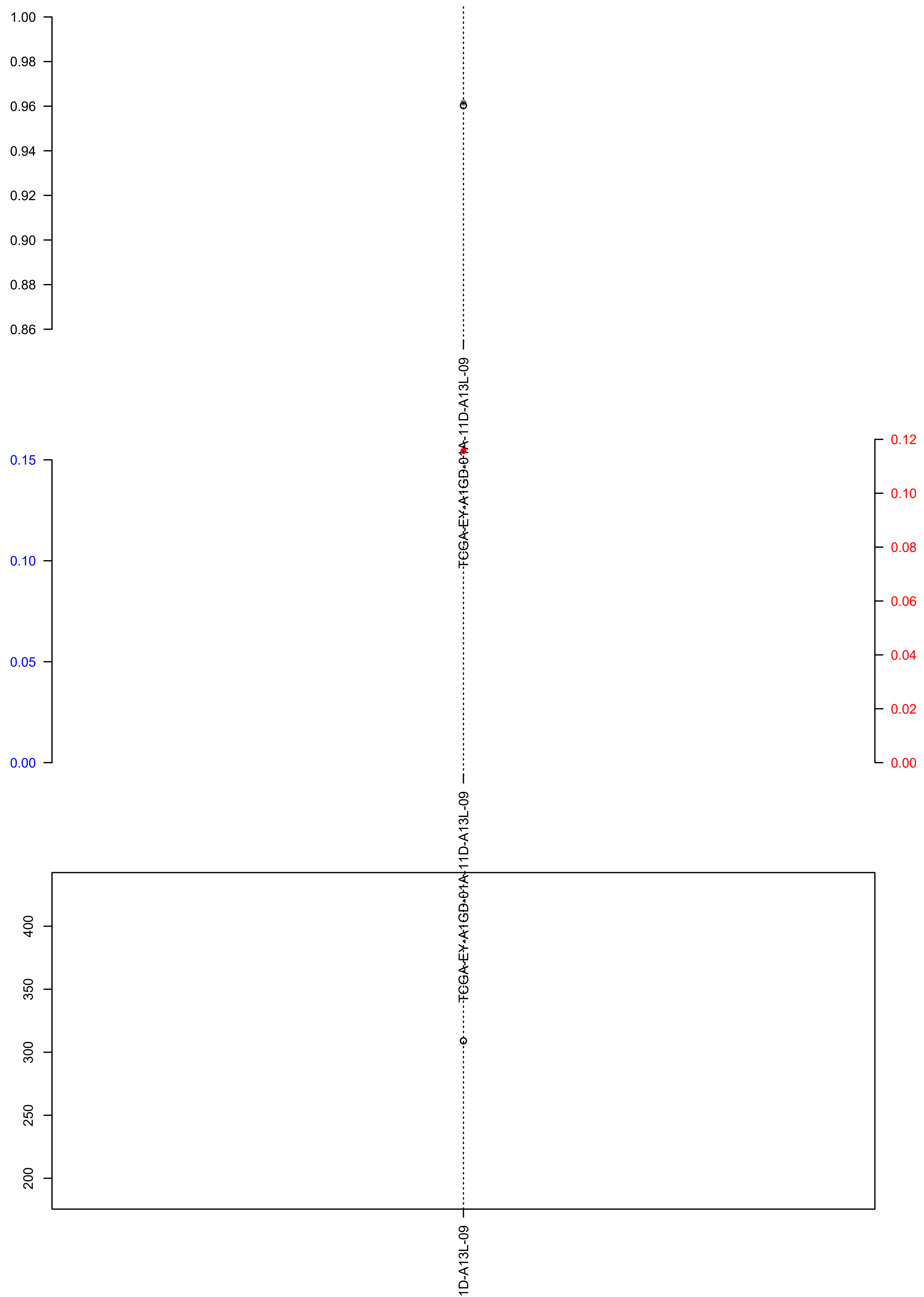

### TCGA-EY-A1GI-01A-11D-A13L-09.check.with.sig.pdf

#### TCGA.EY.A1GI.01A.11D.A13L.09

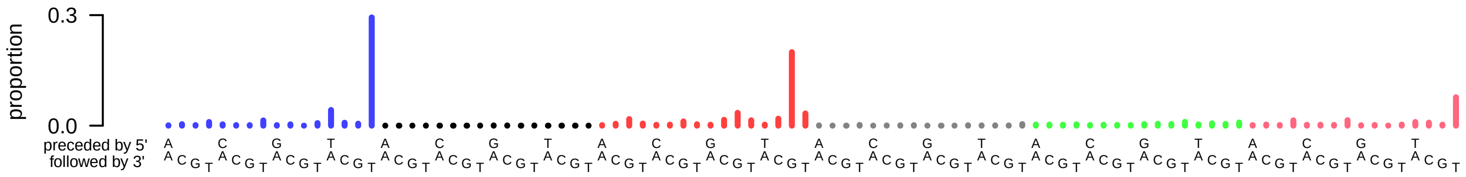

### TCGA-EY-A1GI-01A-11D-A13L-09.exposures.pdf

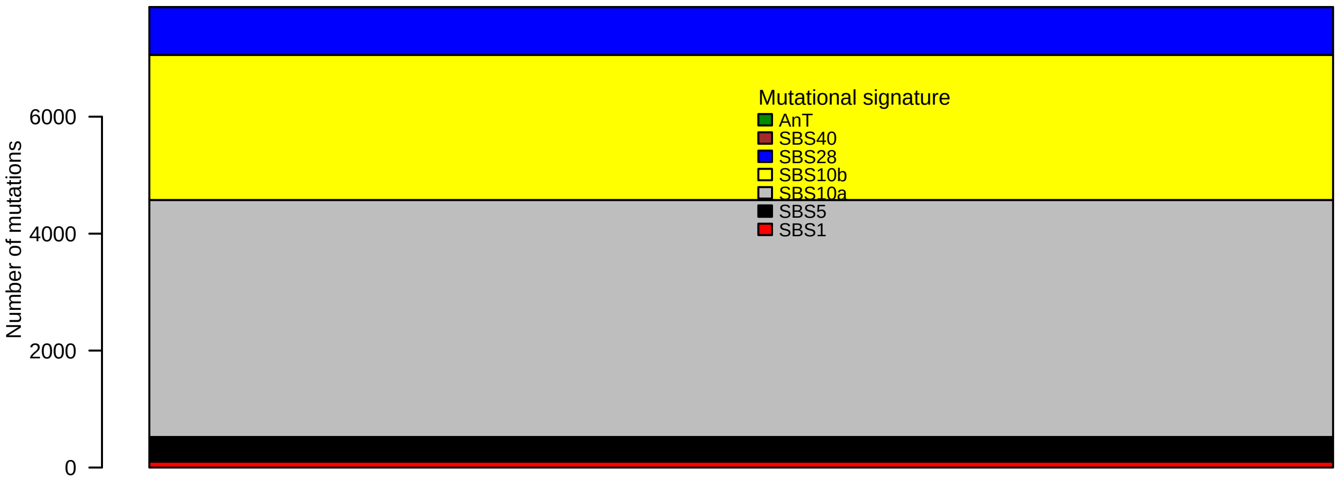

### TCGA-EY-A1GI-01A-11D-A13L-09.pval.histogram.pdf

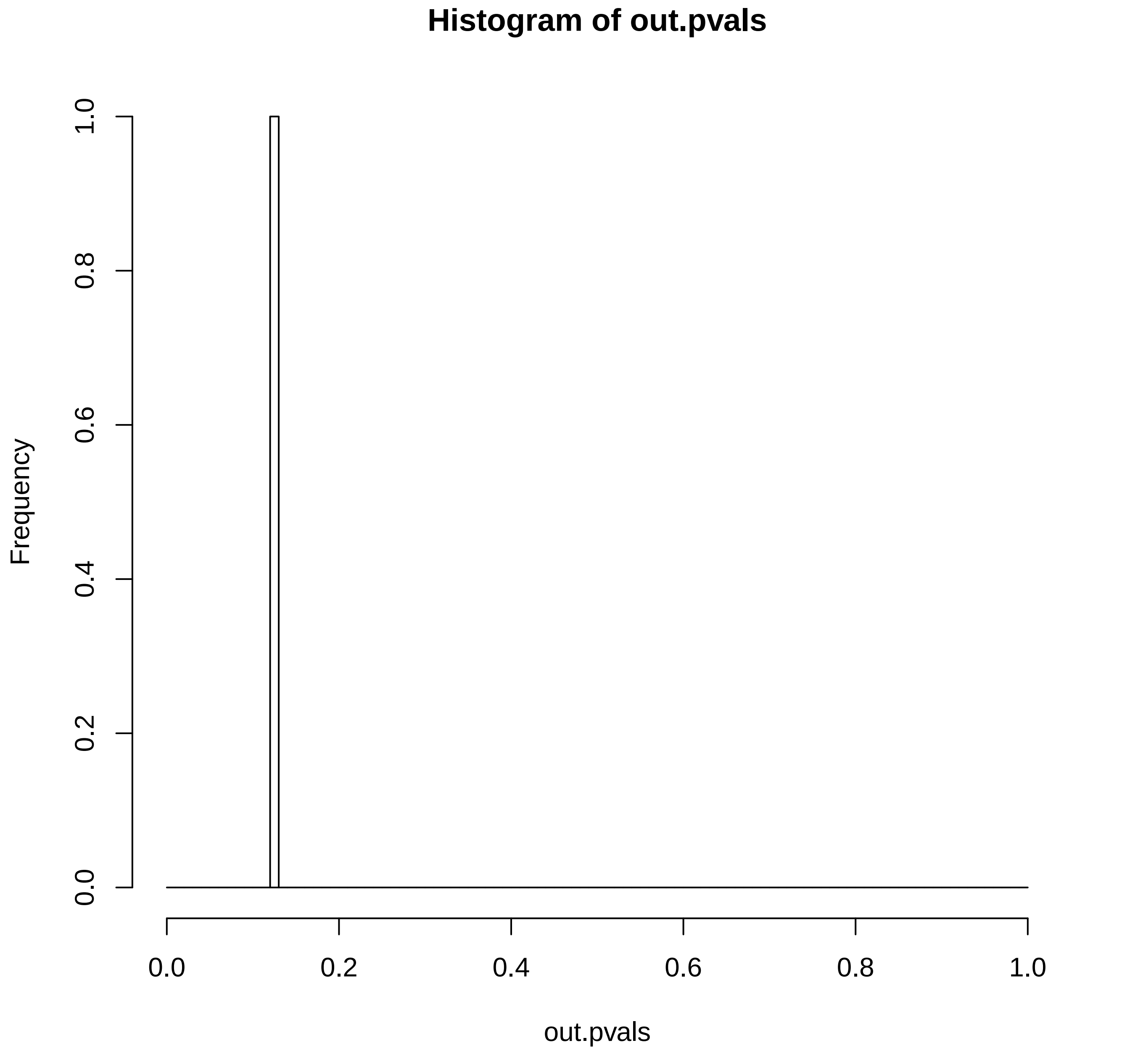

### TCGA-EY-A1GI-01A-11D-A13L-09.reconstruction.err.pdf

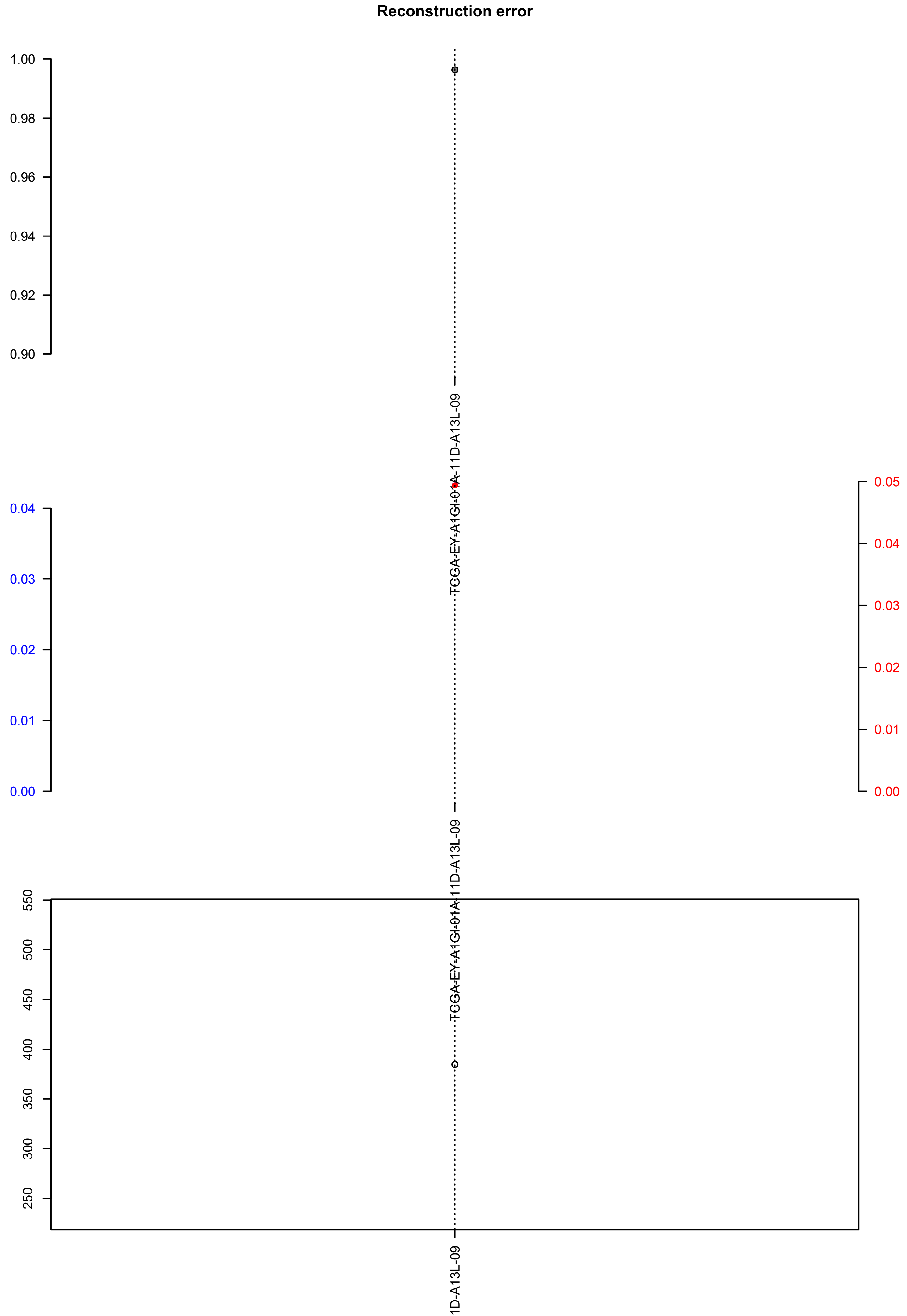

### TCGA-FU-A3HZ-01A-11D-A20U-09.check.with.sig.pdf

#### TCGA.FU.A3HZ.01A.11D.A20U.09

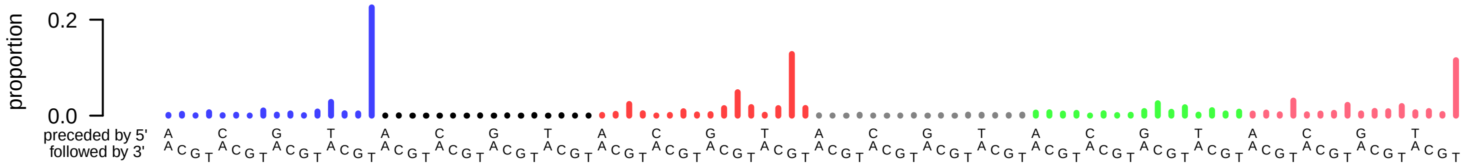

### TCGA-FU-A3HZ-01A-11D-A20U-09.exposures.pdf

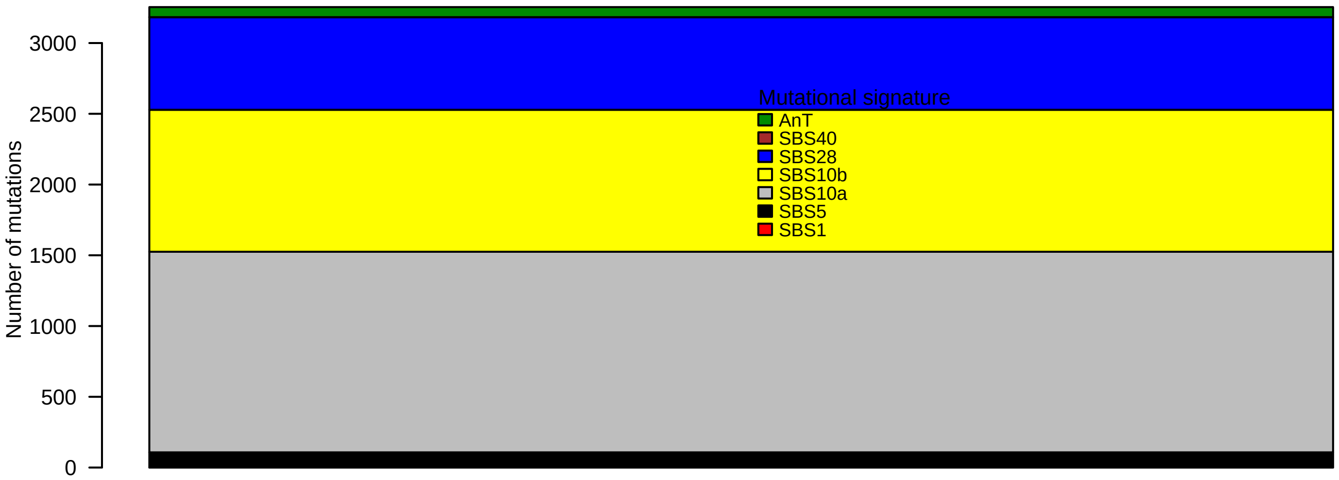

### TCGA-FU-A3HZ-01A-11D-A20U-09.pval.histogram.pdf

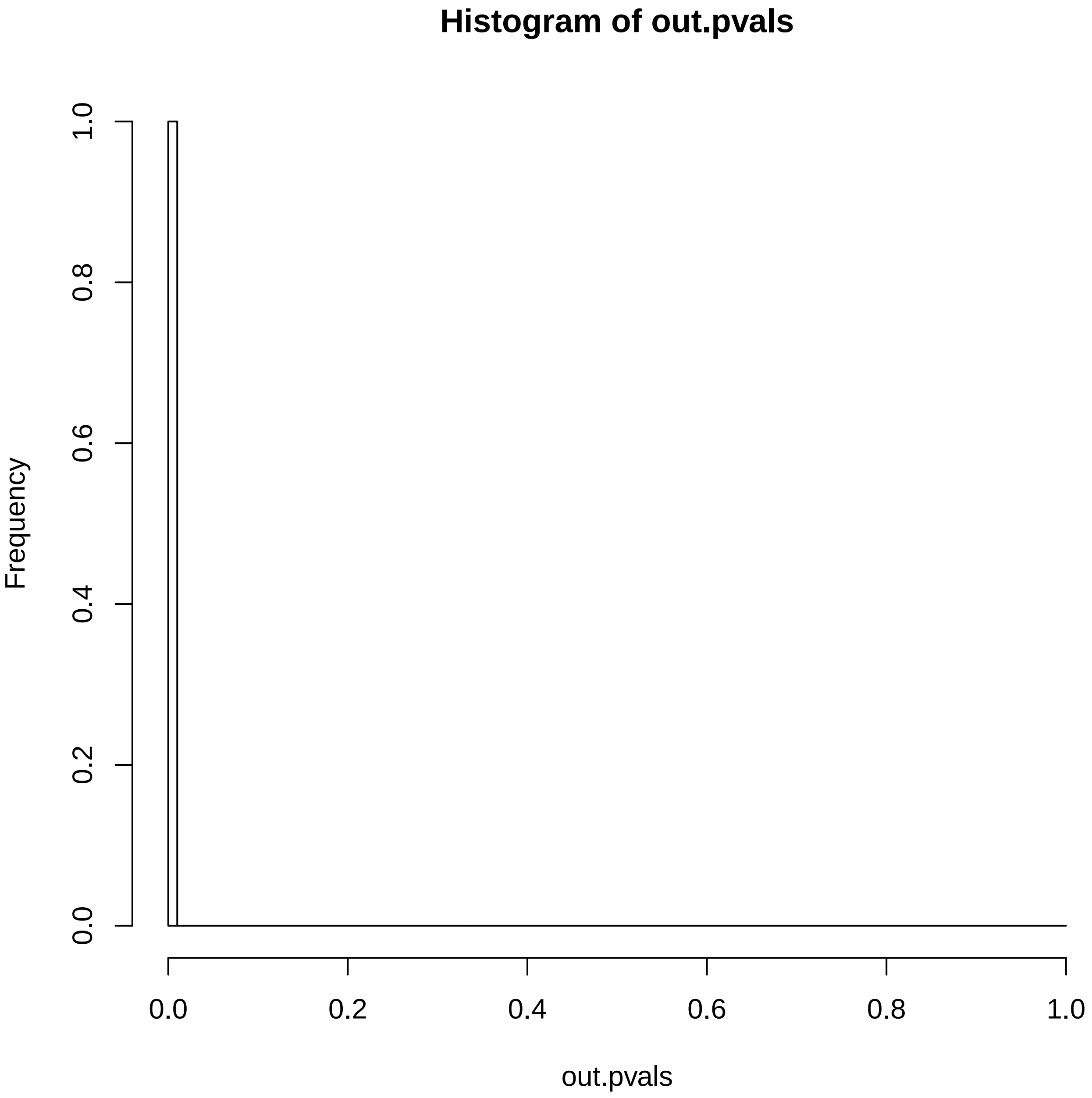

### TCGA-FU-A3HZ-01A-11D-A20U-09.reconstruction.err.pdf

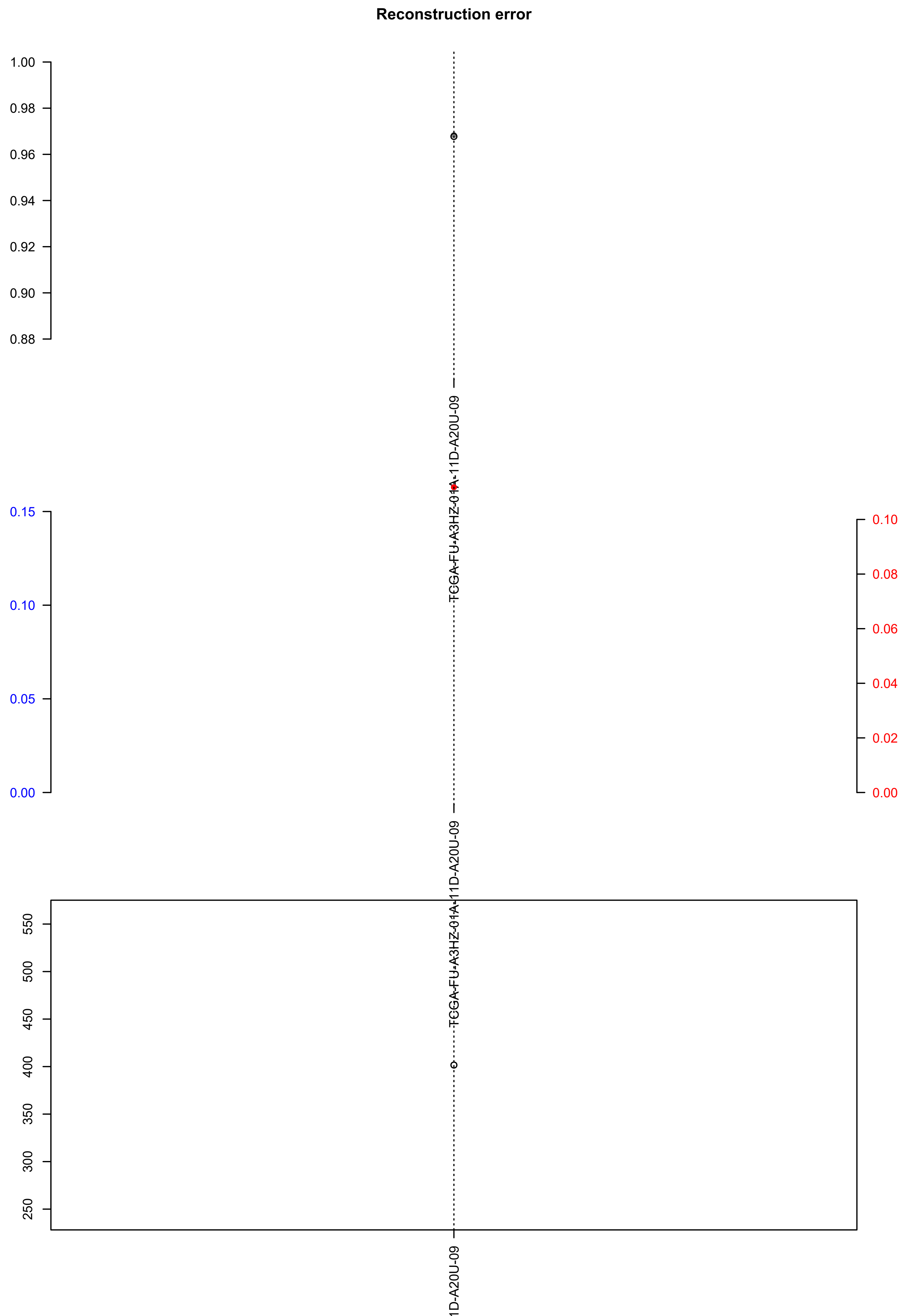

### TCGA-G2-AA3B-01A-11D-A391-08.check.with.sig.pdf

#### TCGA.G2.AA3B.01A.11D.A391.08

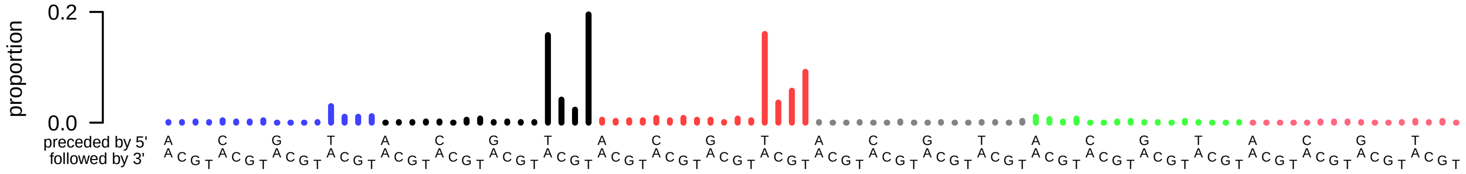

### TCGA-G2-AA3B-01A-11D-A391-08.exposures.pdf

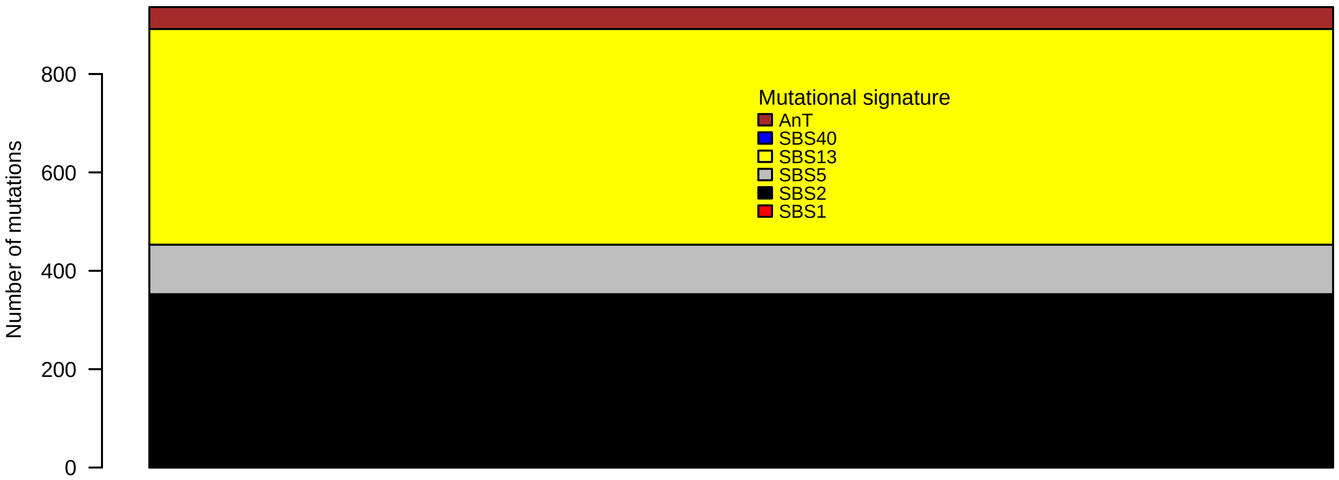

### TCGA-G2-AA3B-01A-11D-A391-08.pval.histogram.pdf

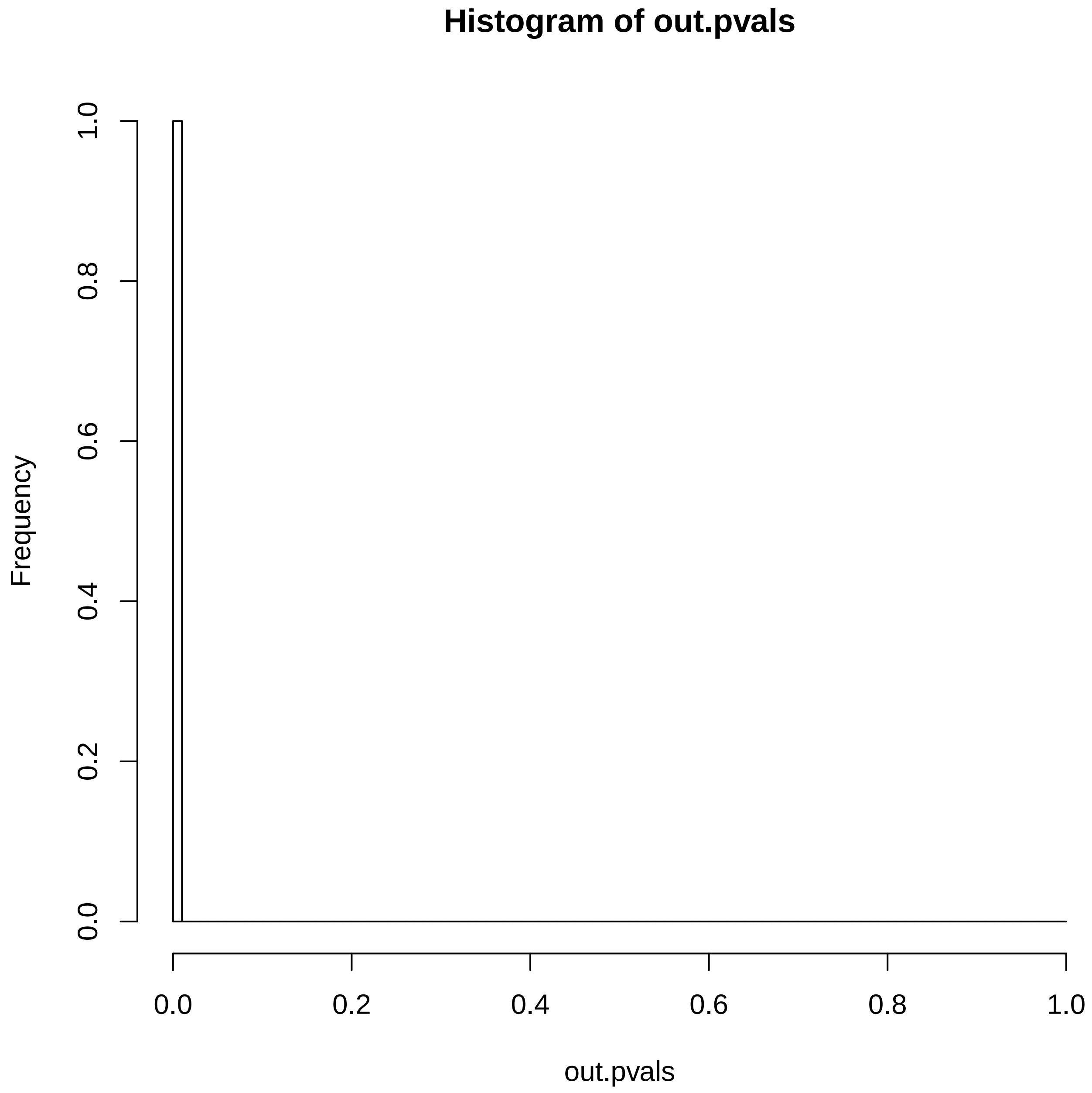

### TCGA-G2-AA3B-01A-11D-A391-08.reconstruction.err.pdf

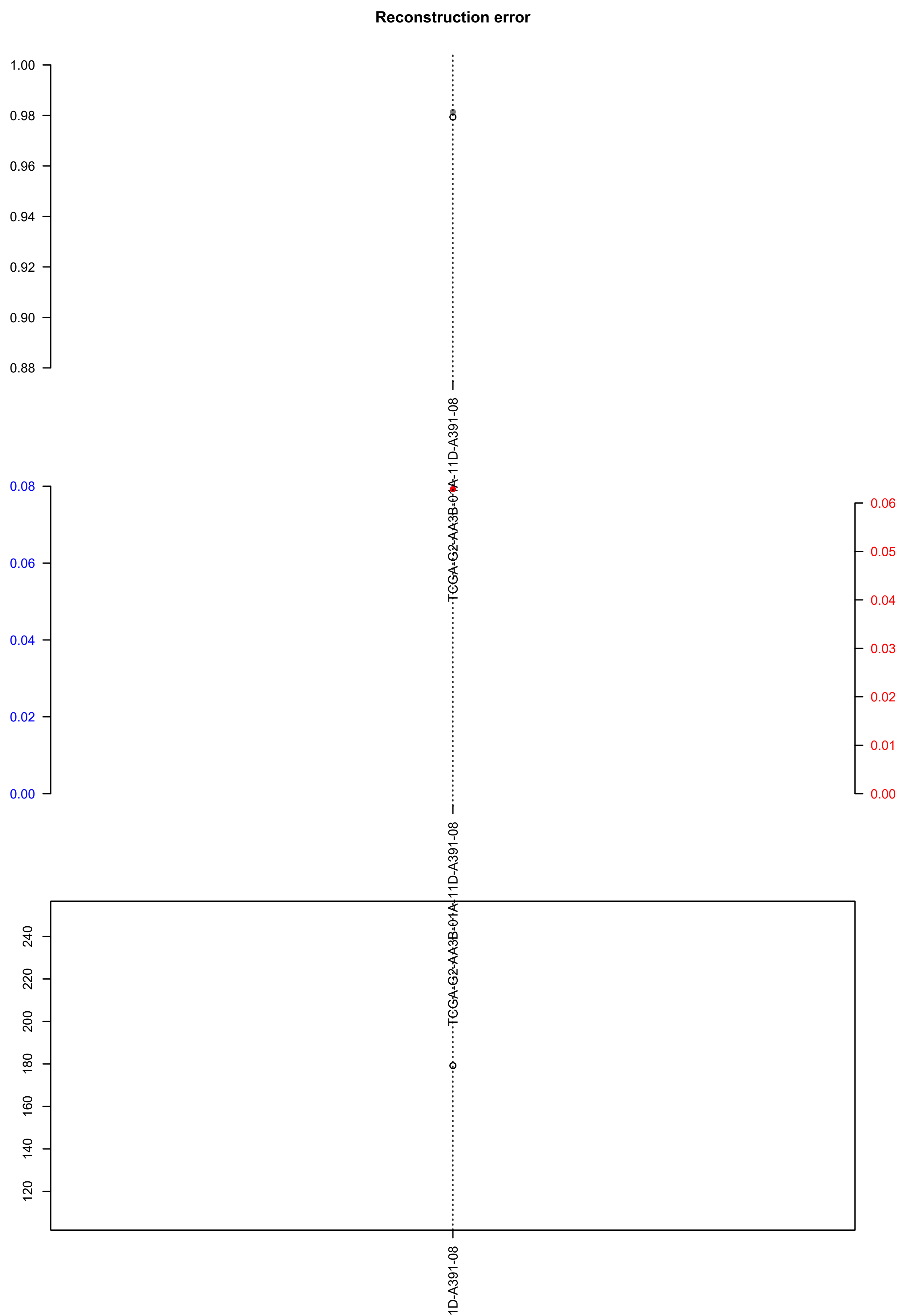

### TCGA-GC-A6I3-01A-11D-A31L-08.check.with.sig.pdf

##### TCGA.GC.A6I3.01A.11D.A31L.08

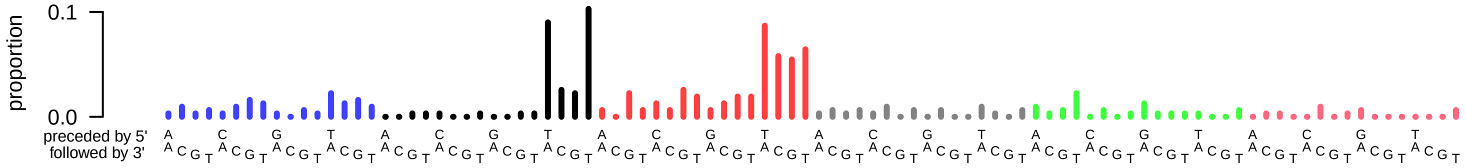

### TCGA-GC-A6I3-01A-11D-A31L-08.exposures.pdf

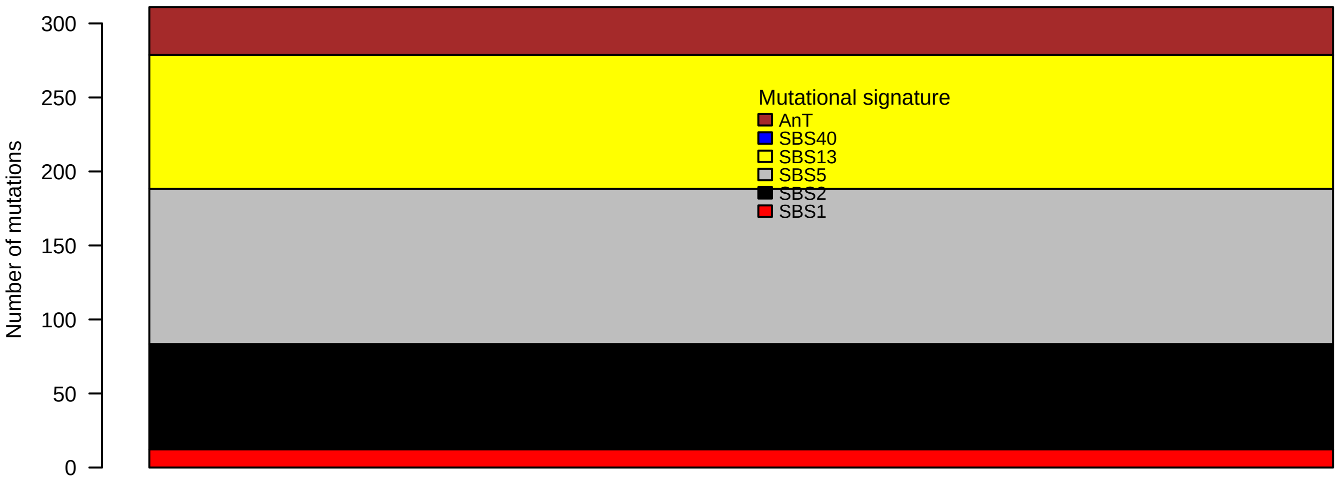

### TCGA-GC-A6I3-01A-11D-A31L-08.pval.histogram.pdf

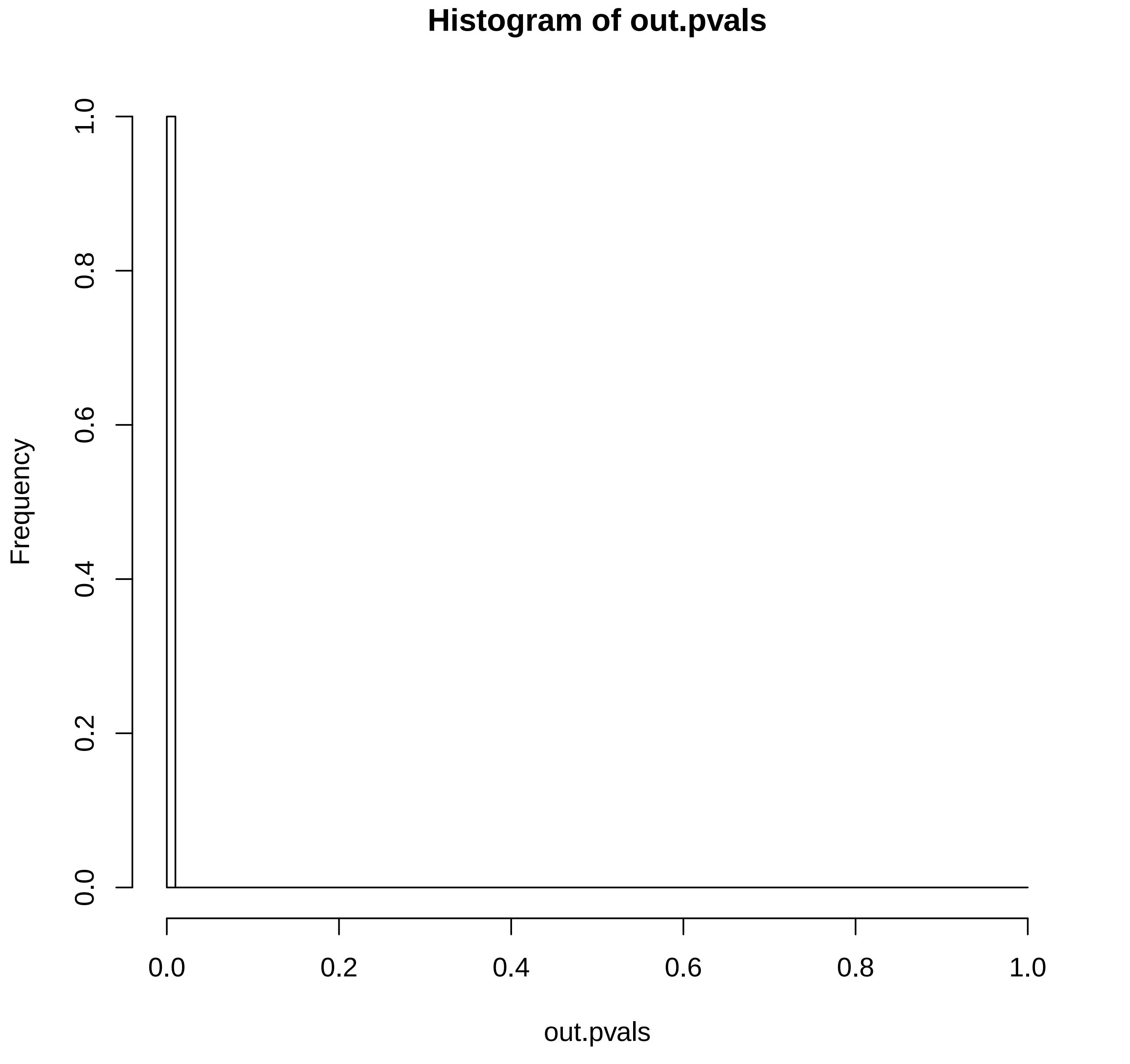

### TCGA-GC-A6I3-01A-11D-A31L-08.reconstruction.err.pdf

##### Reconstruction error

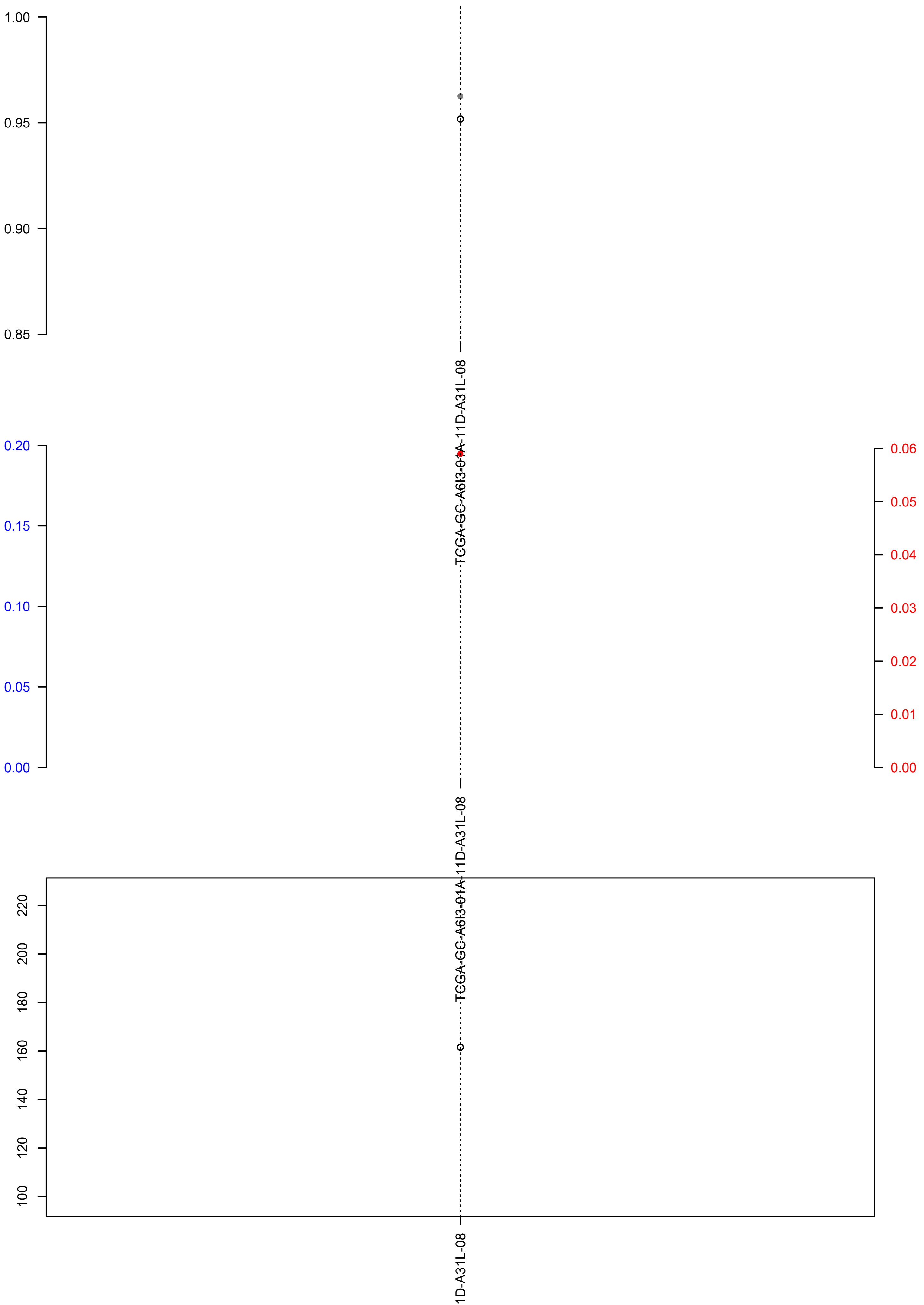

### TCGA-IG-A4QS-01A-11D-A27G-09.check.with.sig.pdf

##### TCGA.IG.A4QS.01A.11D.A27G.09

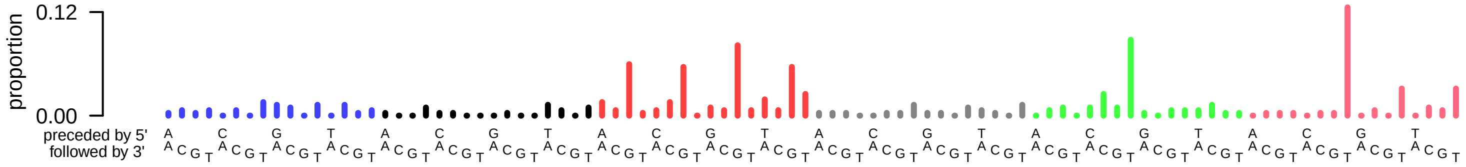

### TCGA-IG-A4QS-01A-11D-A27G-09.exposures.pdf

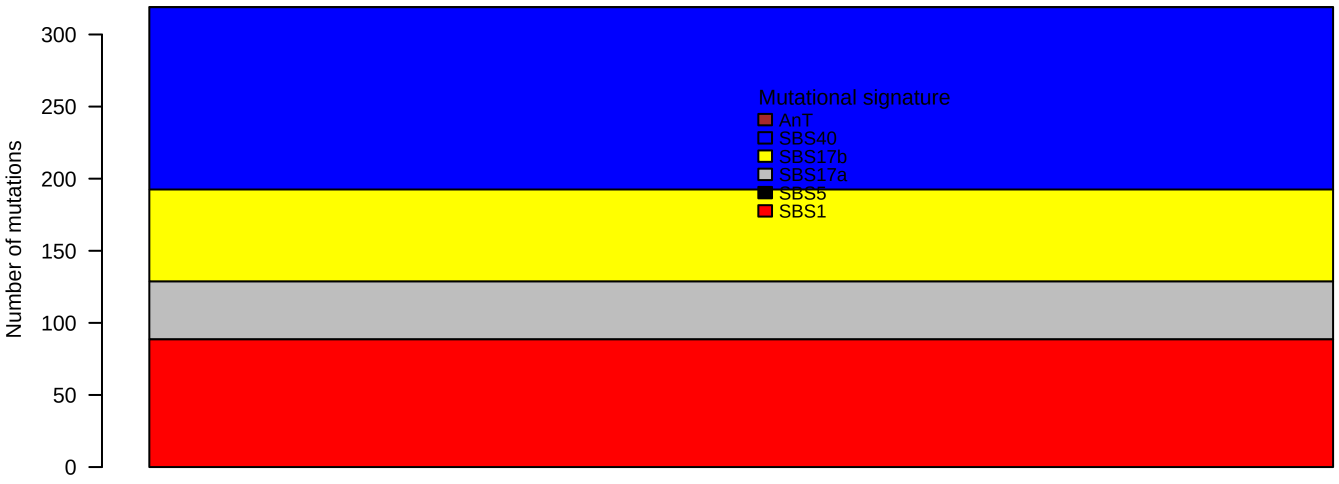

### TCGA-IG-A4QS-01A-11D-A27G-09.pval.histogram.pdf

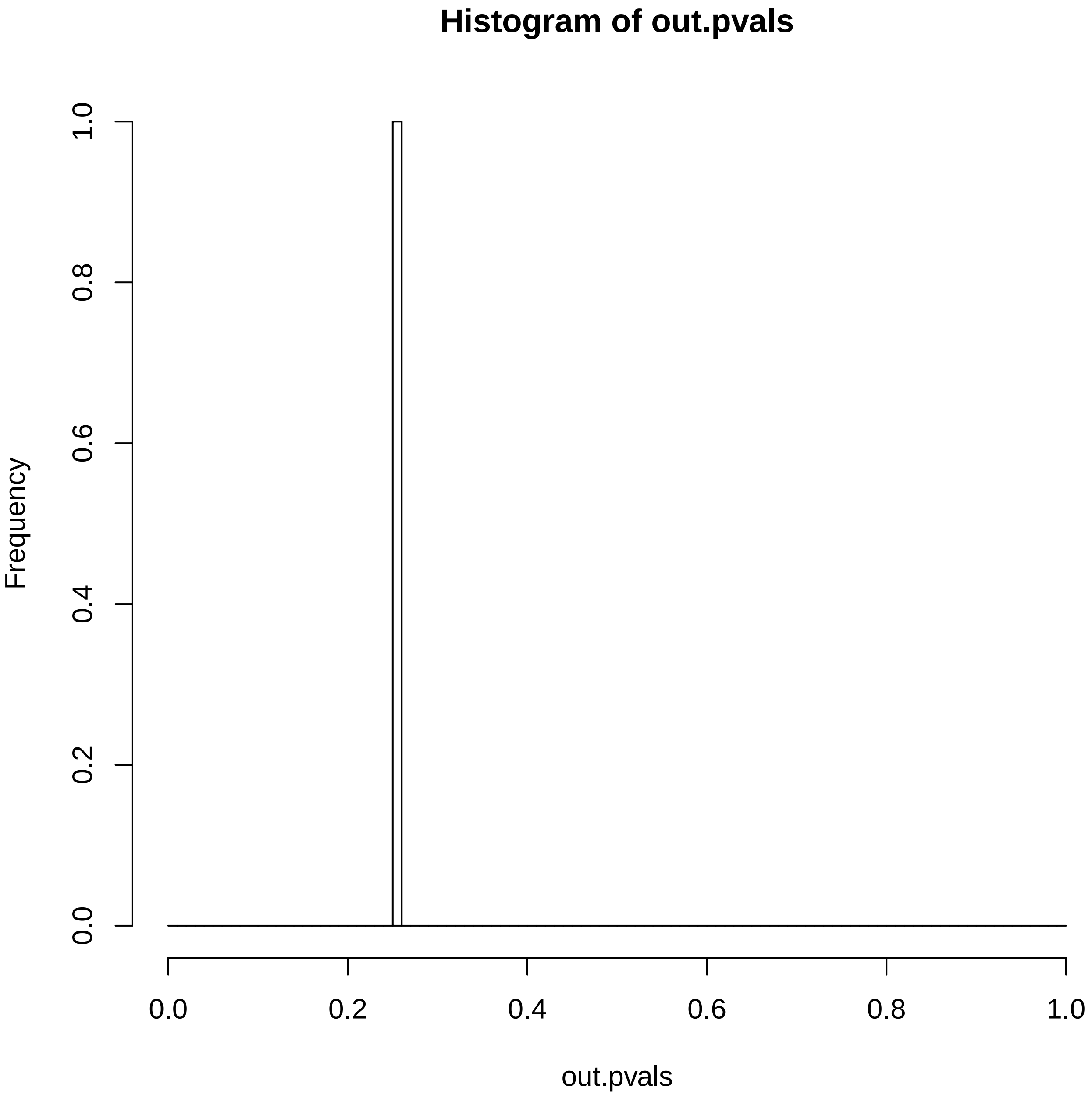

### TCGA-IG-A4QS-01A-11D-A27G-09.reconstruction.err.pdf

**Reconstruction error**

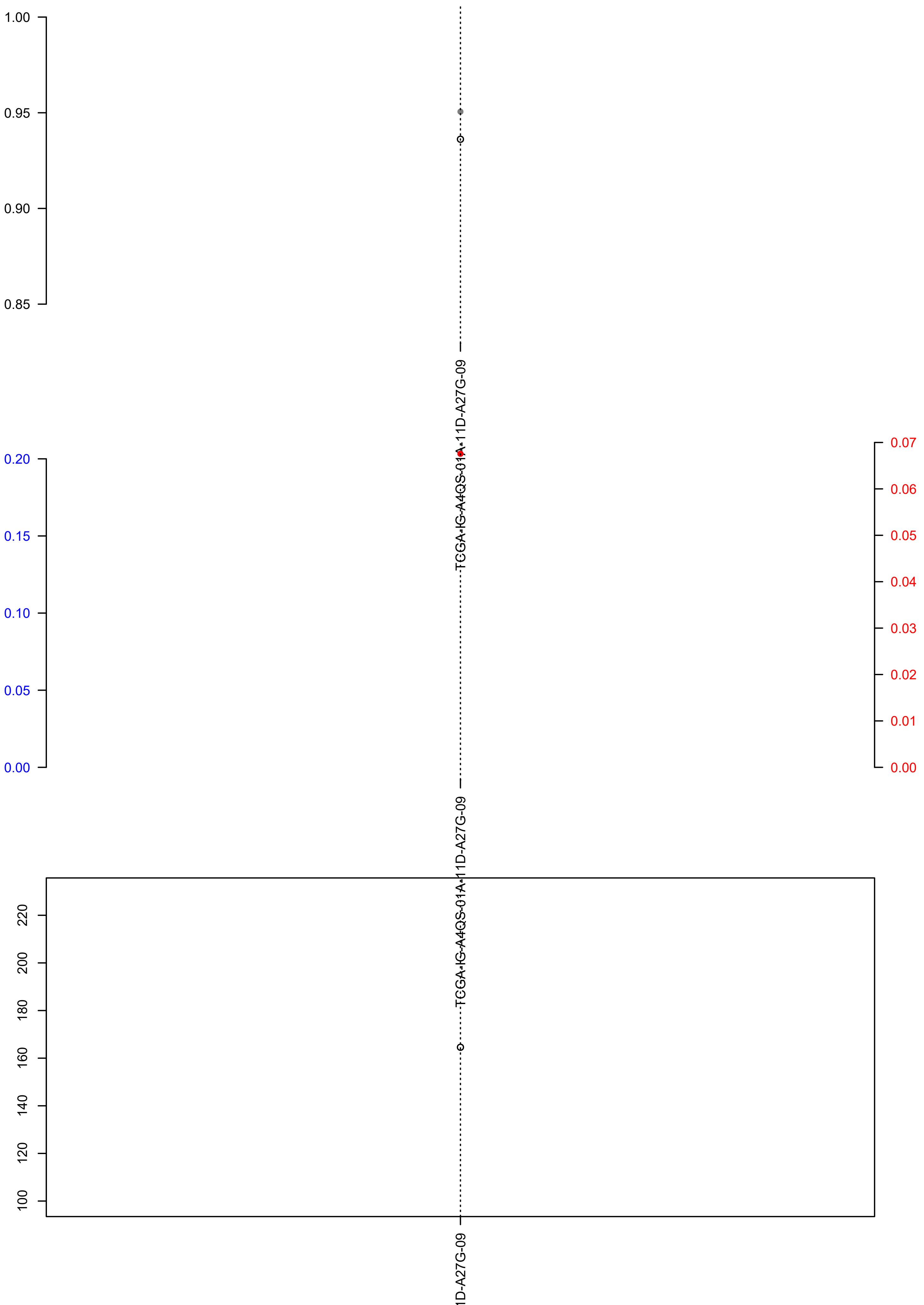

### TCGA-QF-A5YS-01A-11D-A31U-09.check.with.sig.pdf

#### TCGA.QF.A5YS.01A.11D.A31U.09

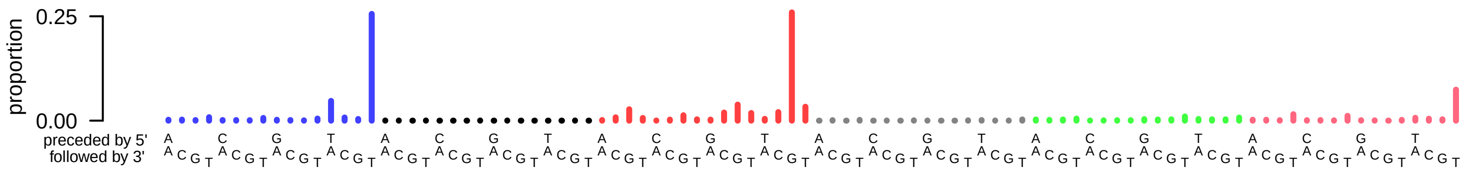

### TCGA-QF-A5YS-01A-11D-A31U-09.exposures.pdf

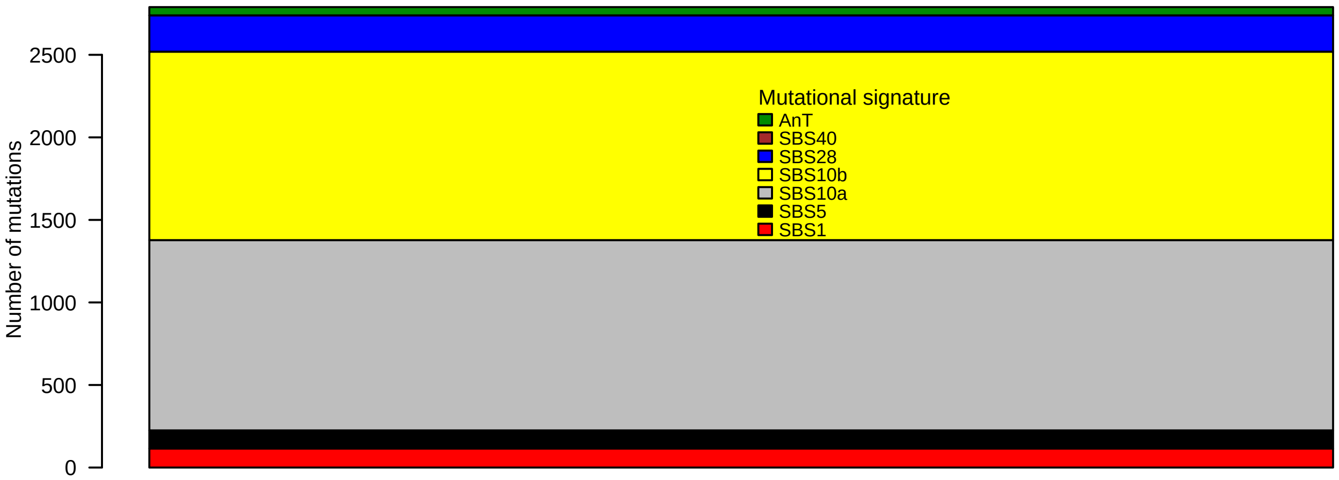

### TCGA-QF-A5YS-01A-11D-A31U-09.pval.histogram.pdf

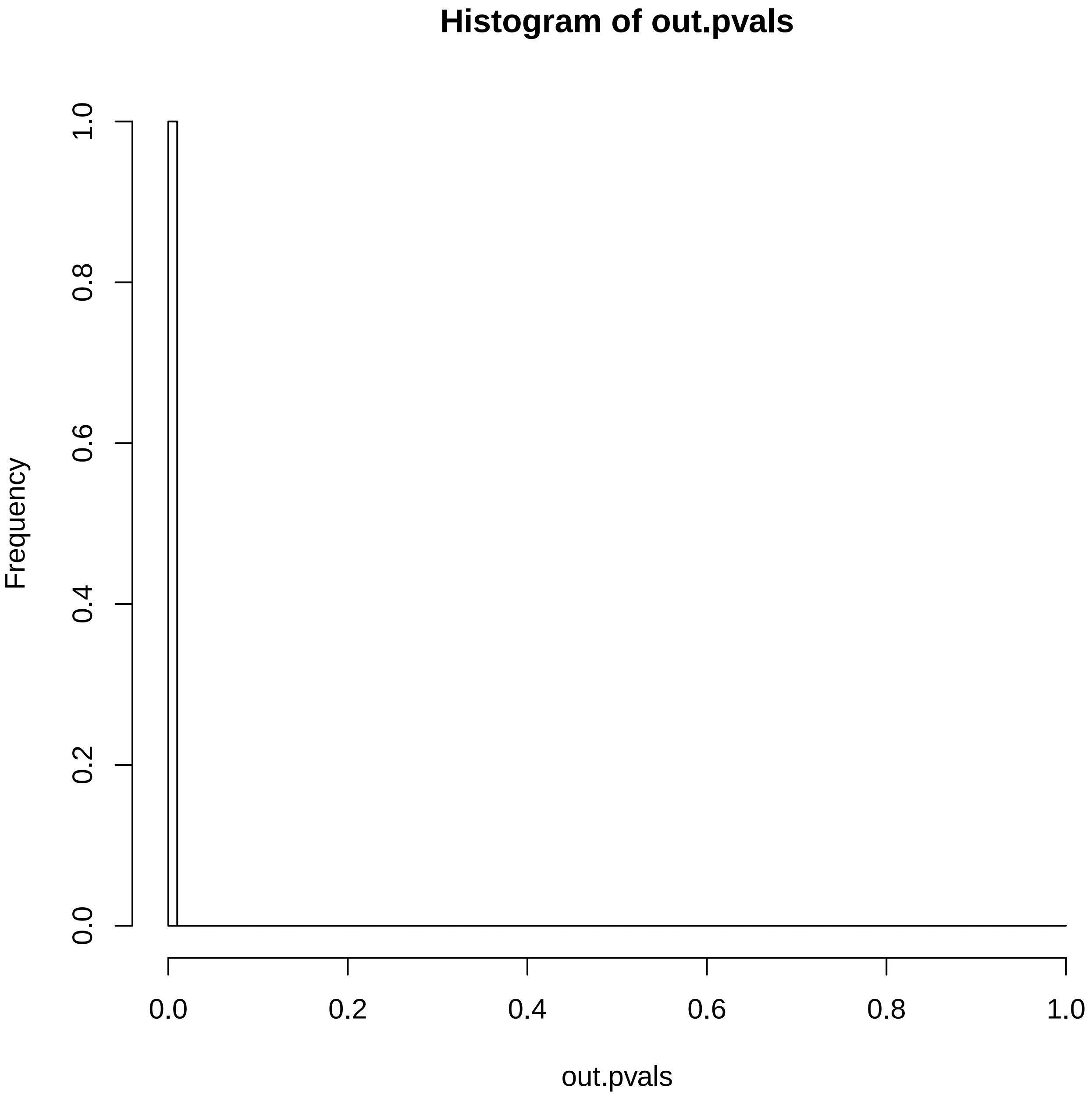

### TCGA-QF-A5YS-01A-11D-A31U-09.reconstruction.err.pdf

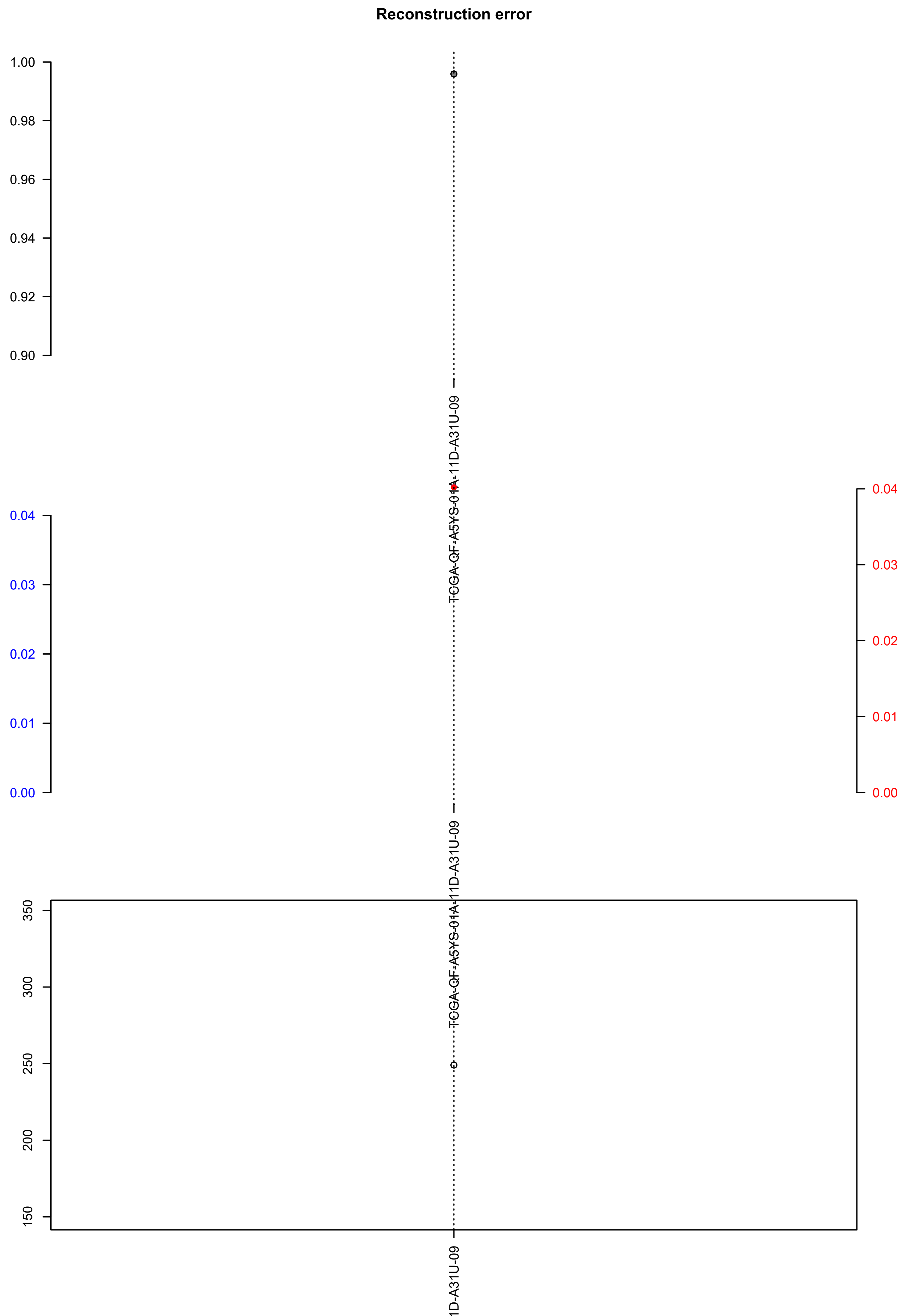

### TCGA-QS-A5YQ-01A-11D-A31U-09.exposures.pdf

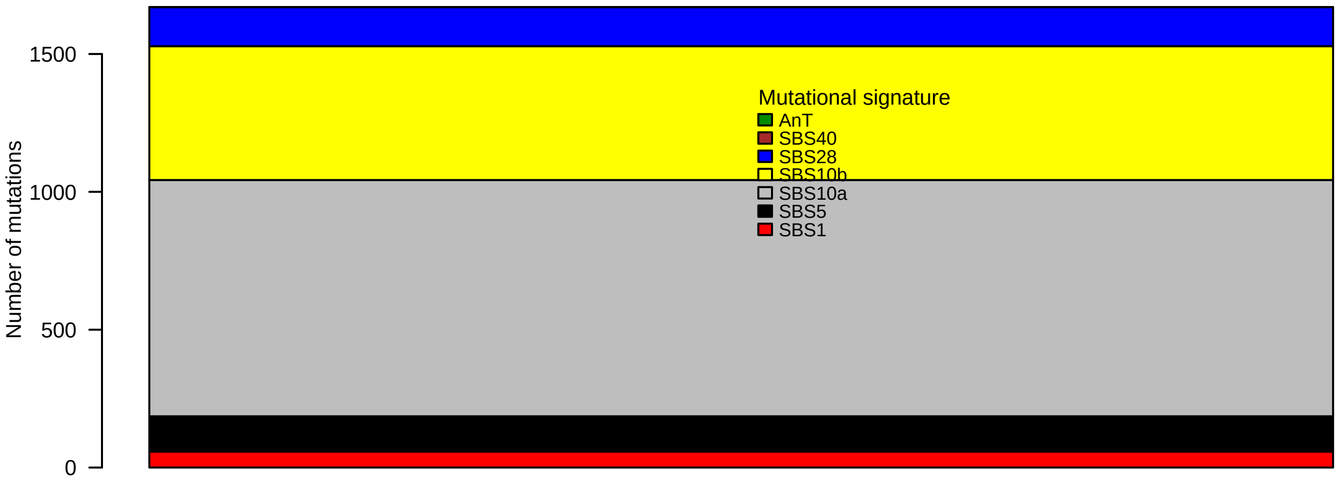

### TCGA-QS-A5YQ-01A-11D-A31U-09.pval.histogram.pdf

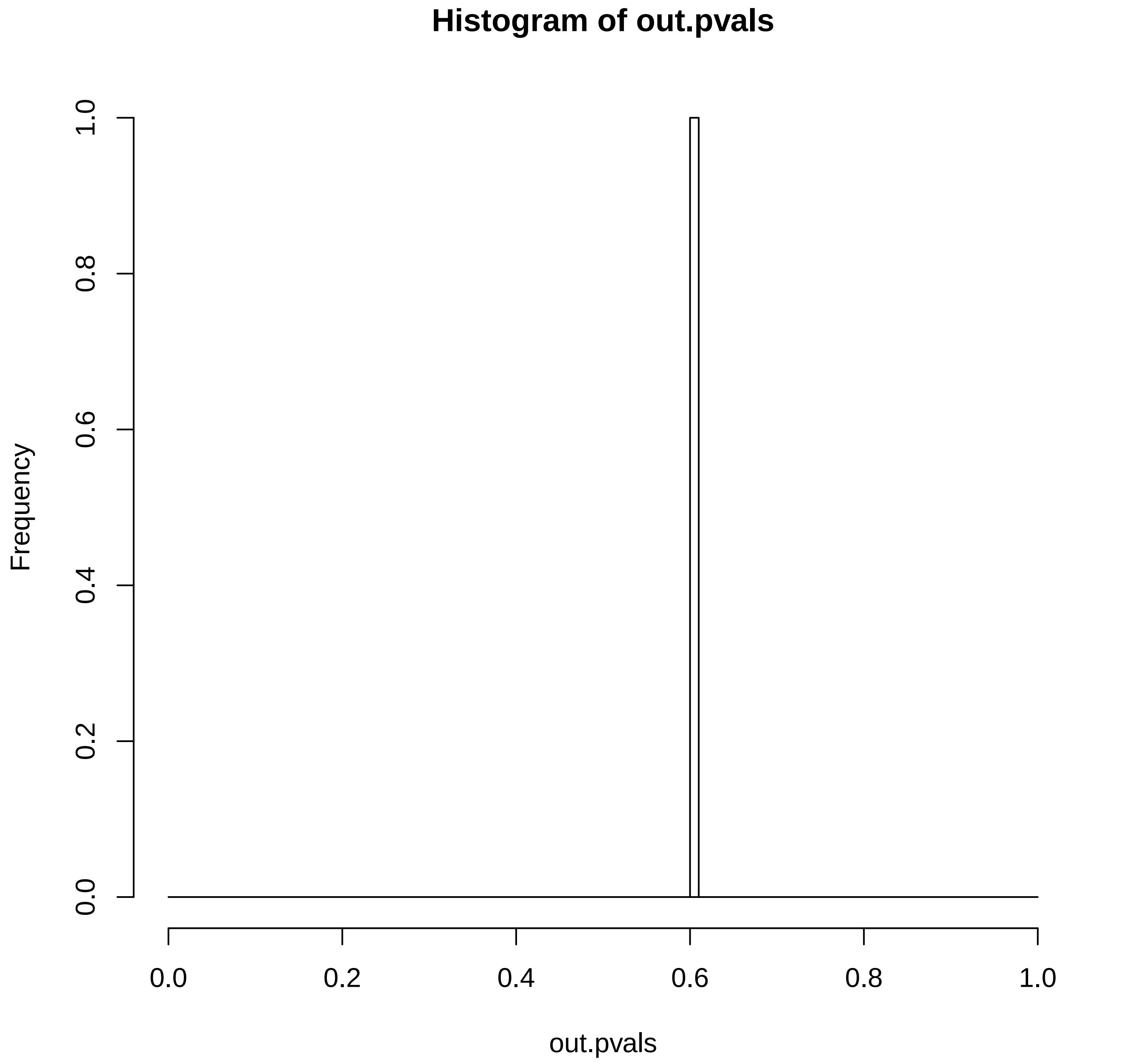

### TCGA-QS-A5YQ-01A-11D-A31U-09.reconstruction.err.pdf

**Reconstruction error**

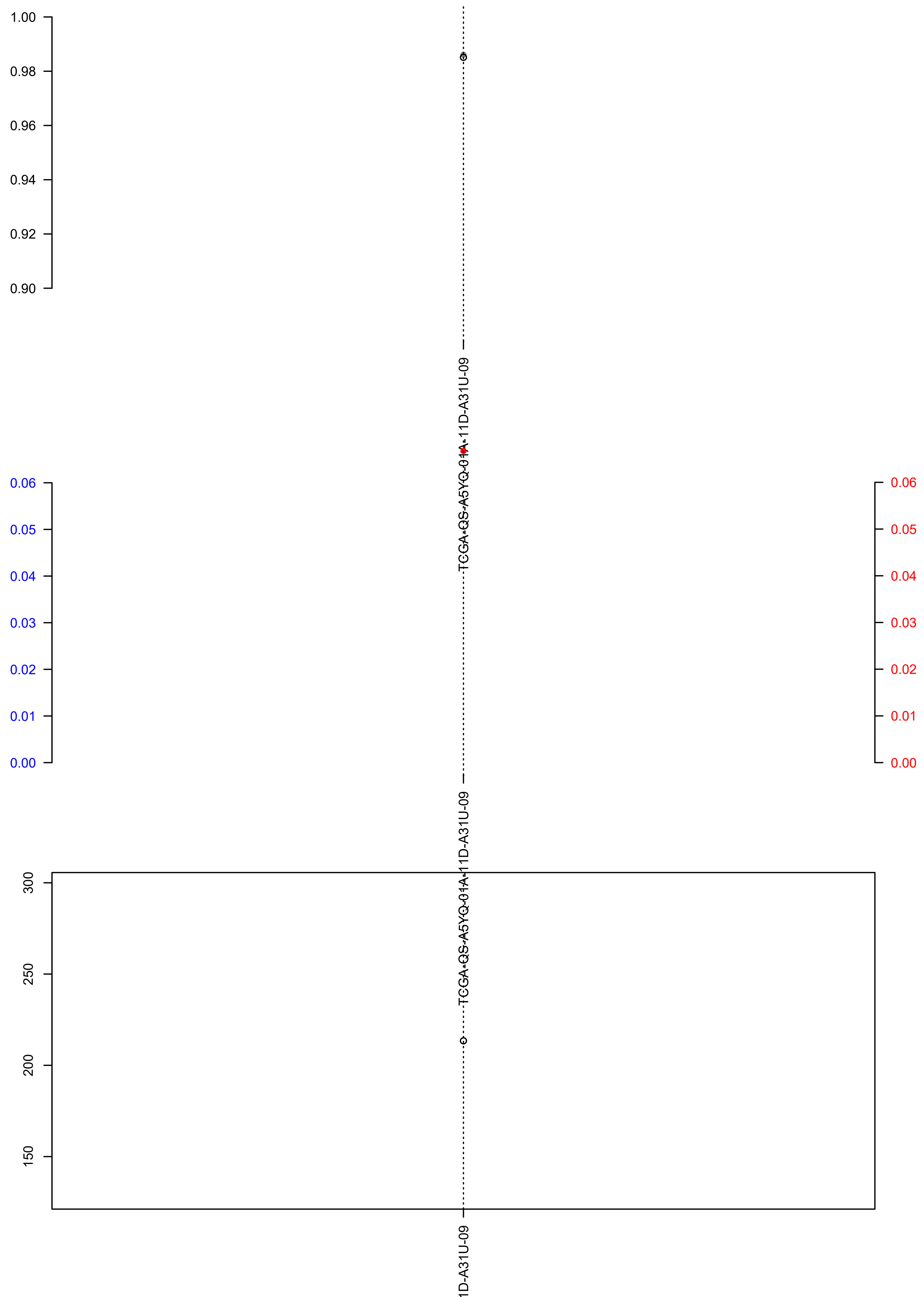

### TCGA-R6-A6L4-01A-11D-A31U-09.exposures.pdf

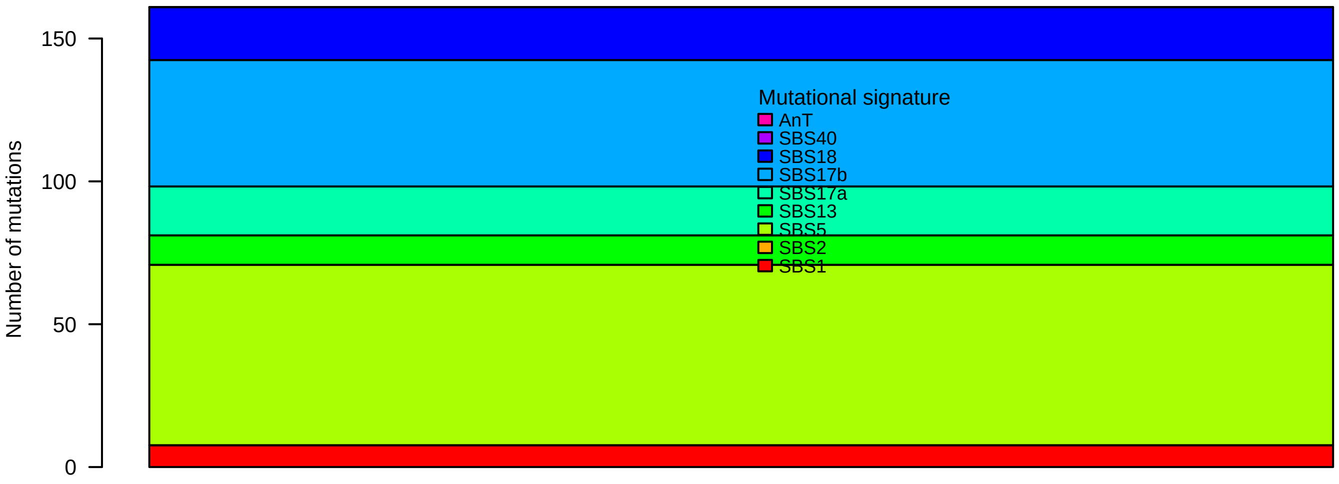
