## Supplementary figures and images for "Identification of novel mutational signatures in Asian oral squamous cell carcinomas associated with bacterial infections"

### 0047_CRUK_PC_0047_T1_DNA.check.with.sig.pdf

#### X0047_CRUK_PC_0047_T1_DNA

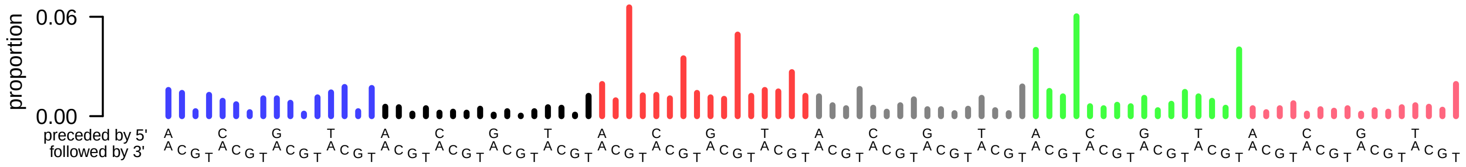

### 0047_CRUK_PC_0047_T1_DNA.exposures.pdf

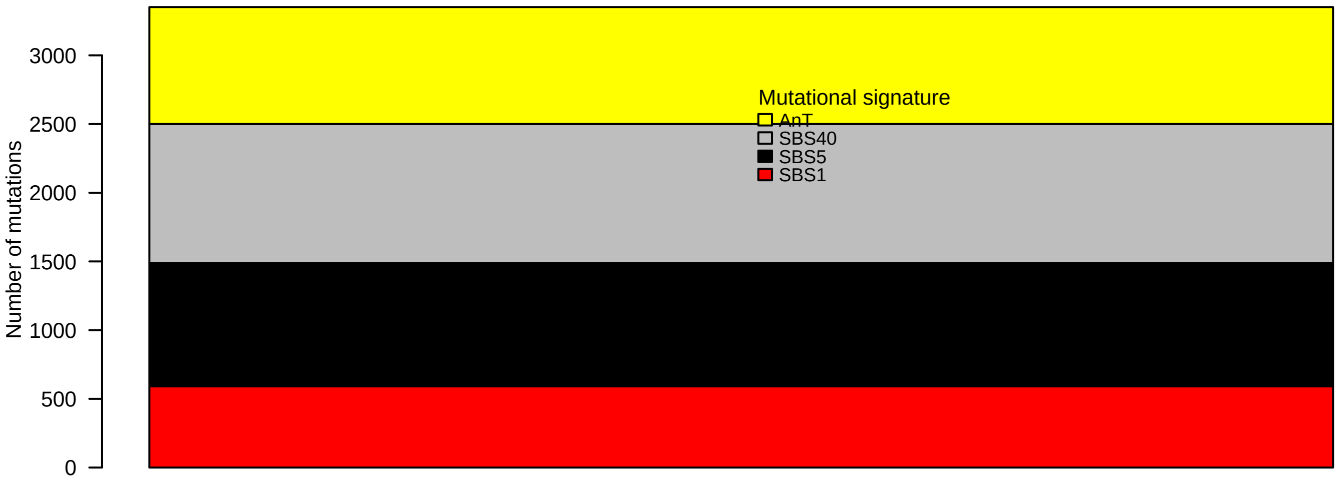

### 0047_CRUK_PC_0047_T1_DNA.pval.histogram.pdf

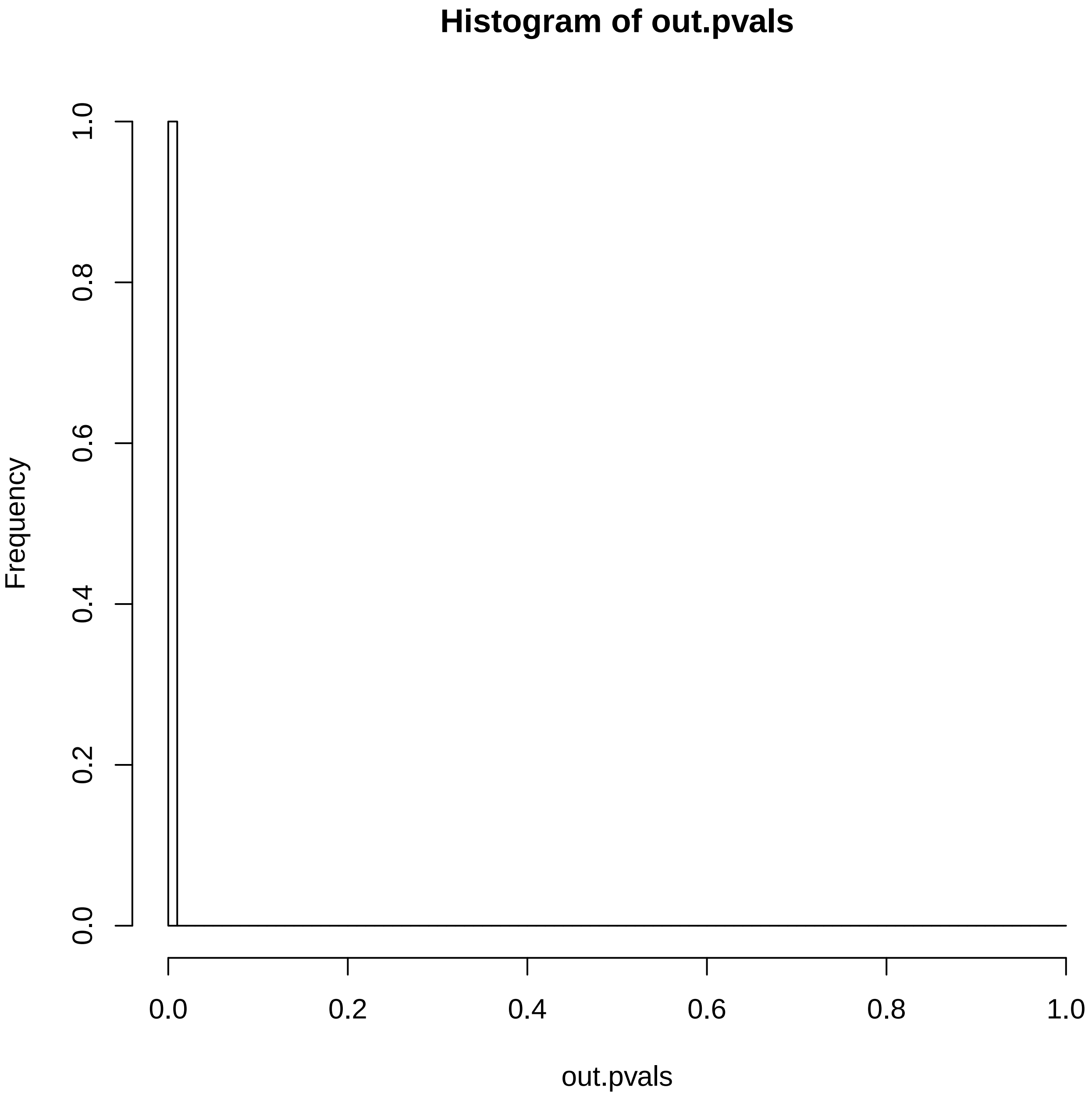

### 0047_CRUK_PC_0047_T1_DNA.reconstruction.err.pdf

#### Reconstruction error

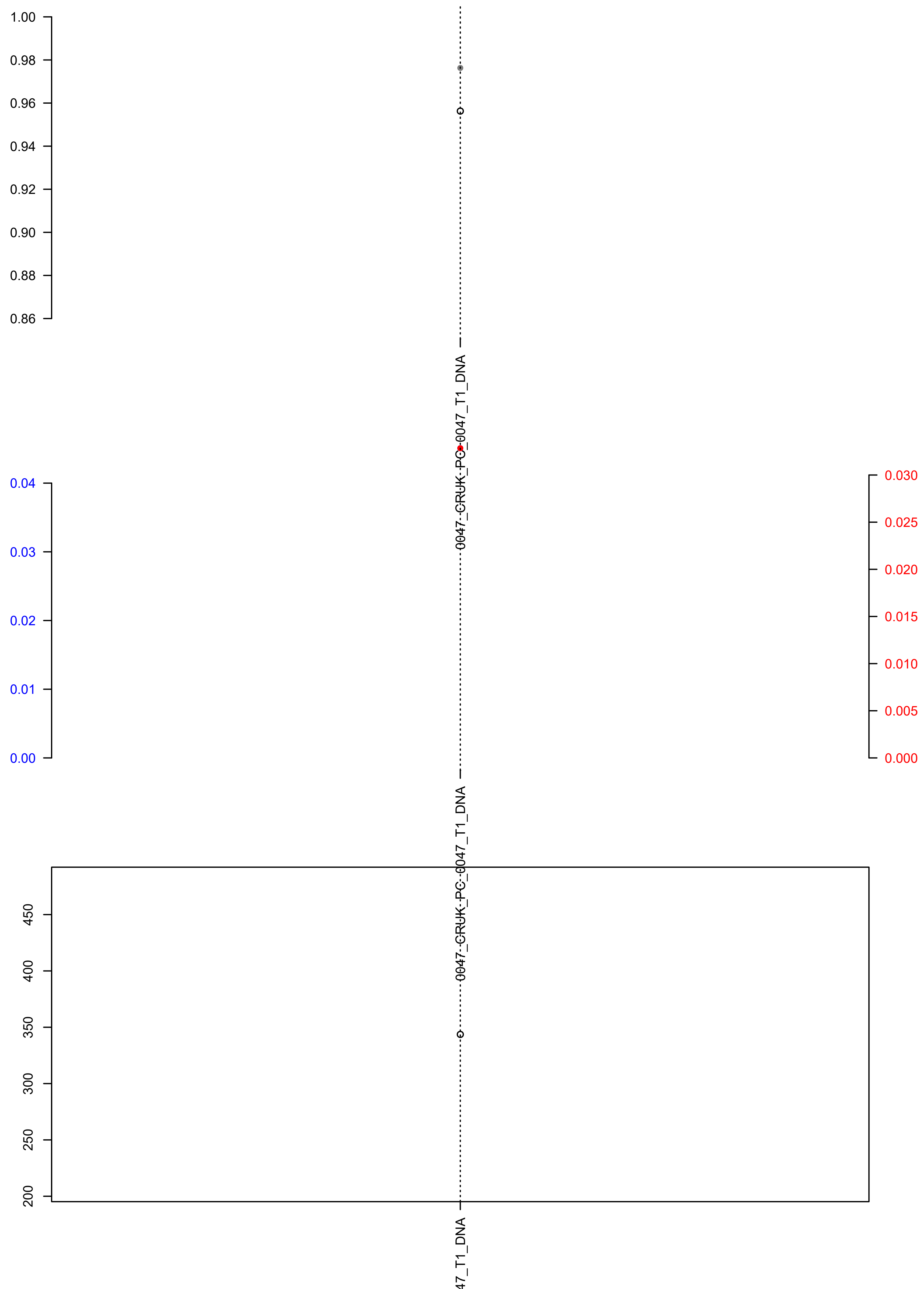

### 8069334.check.with.sig.pdf

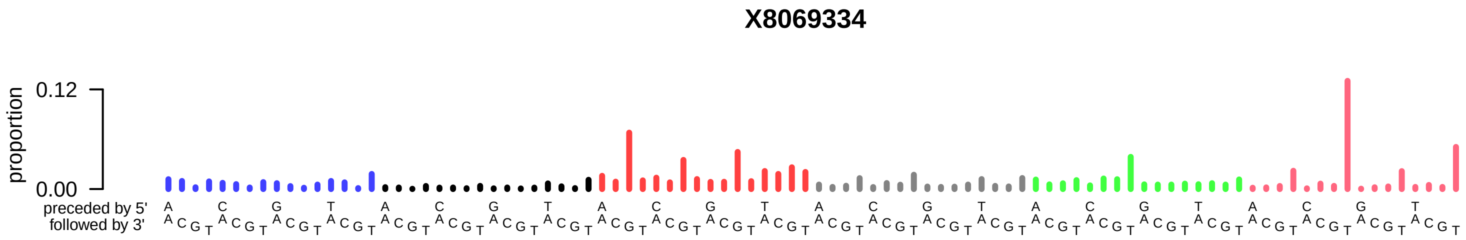

### 8069334.exposures.pdf

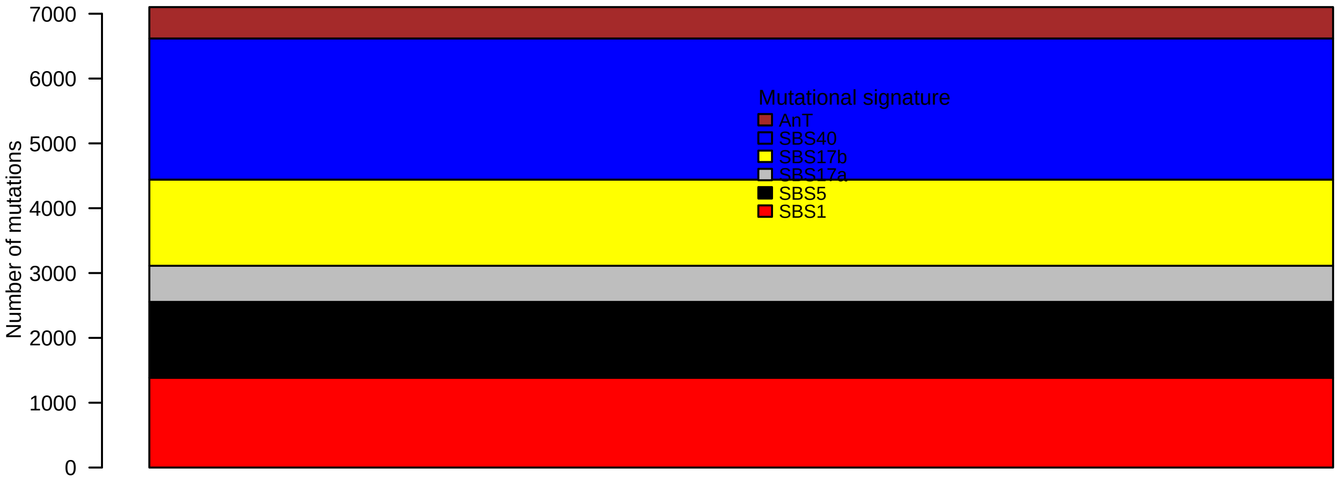

### 8069334.pval.histogram.pdf

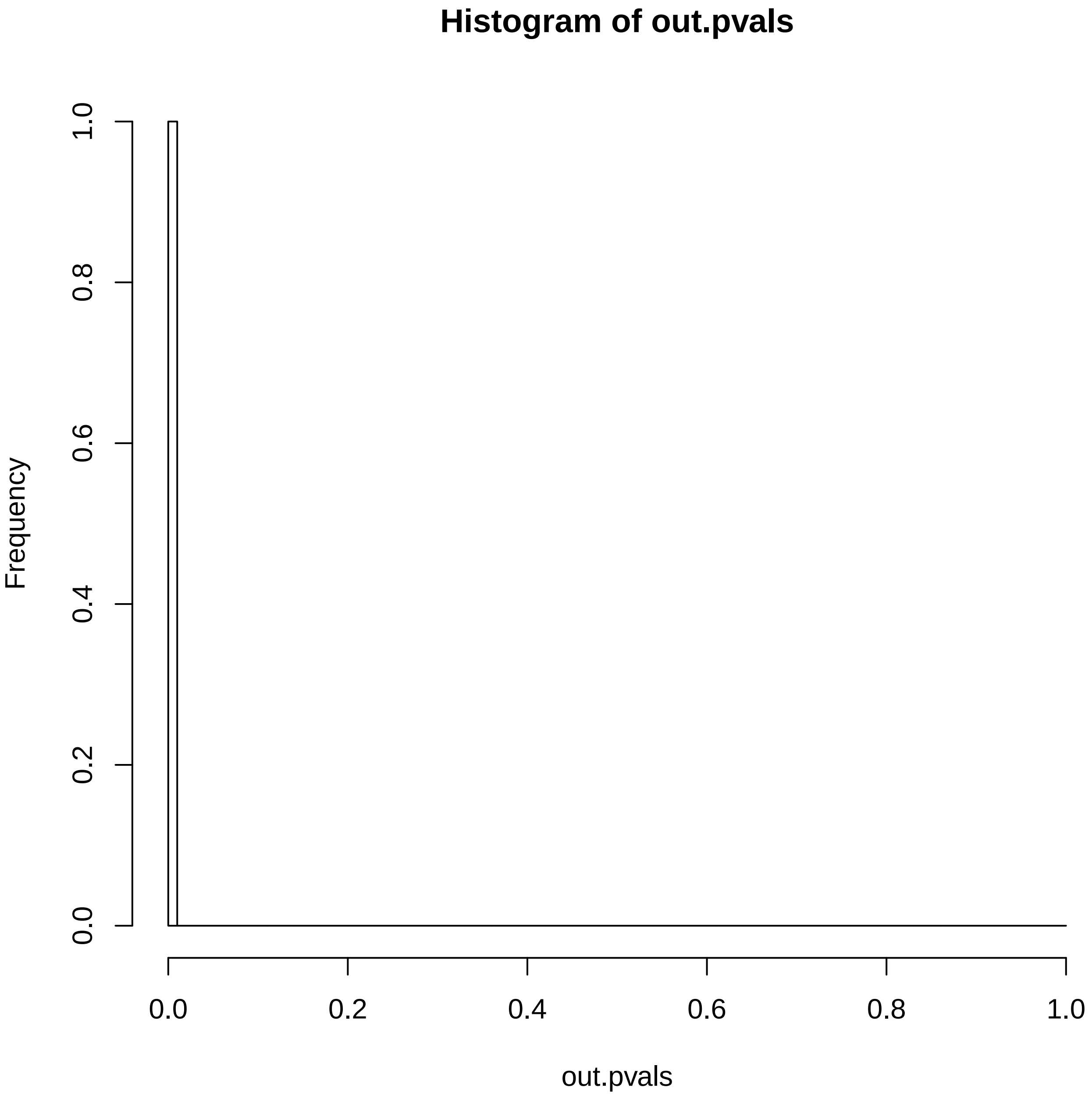

### 8069334.reconstruction.err.pdf

##### Reconstruction error

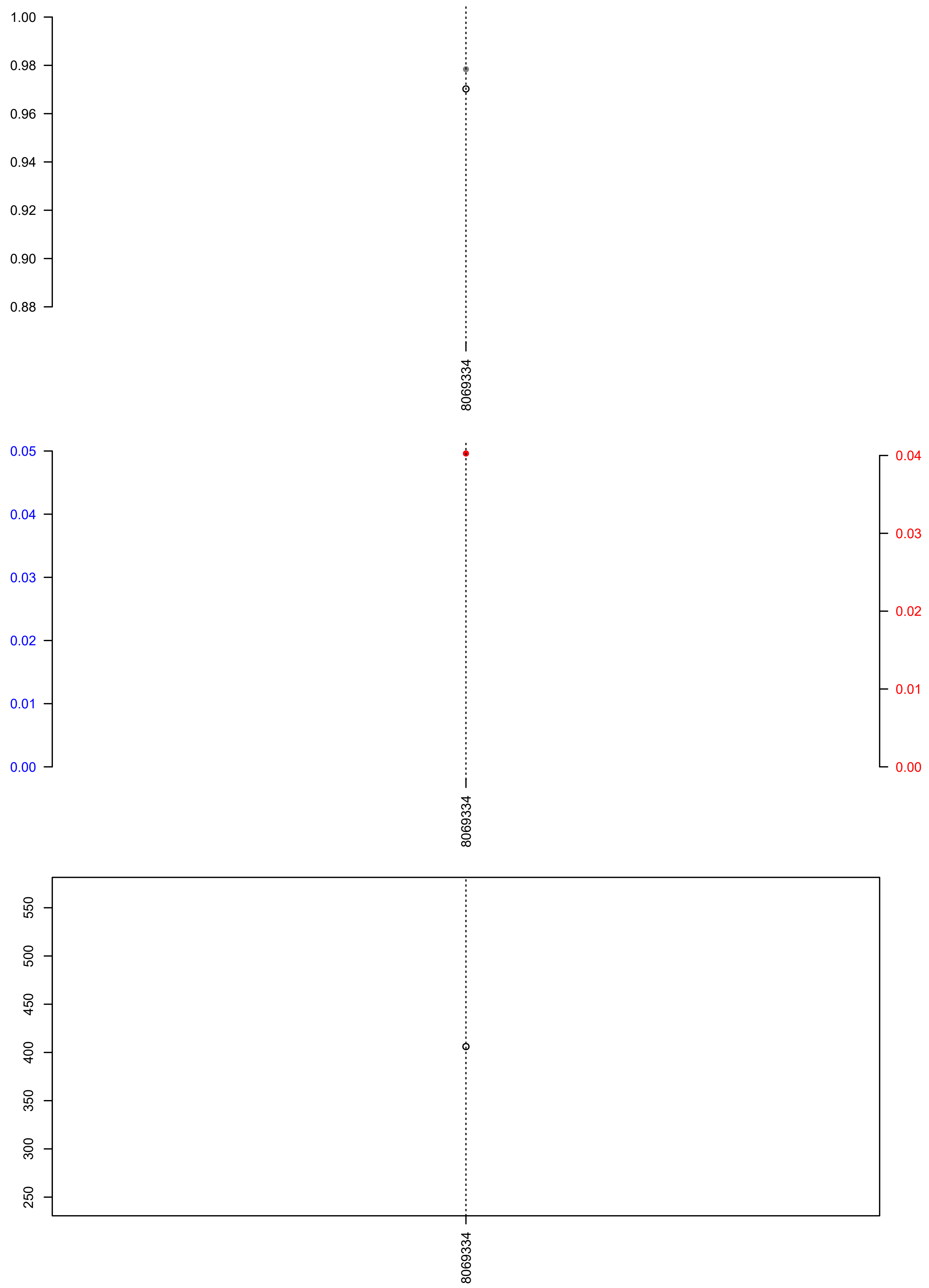

### BD121T.check.with.sig.pdf

## BD121T

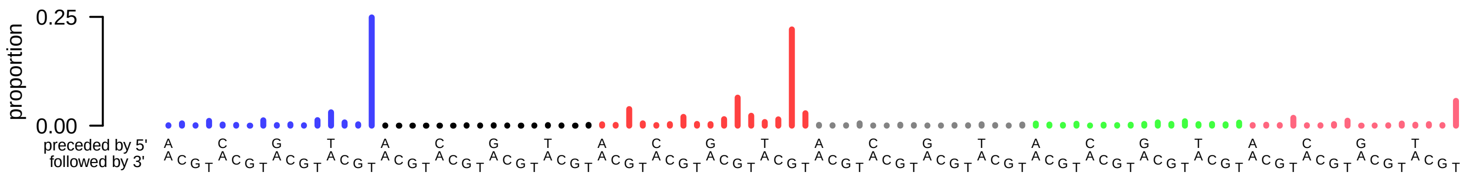

### BD121T.exposures.pdf

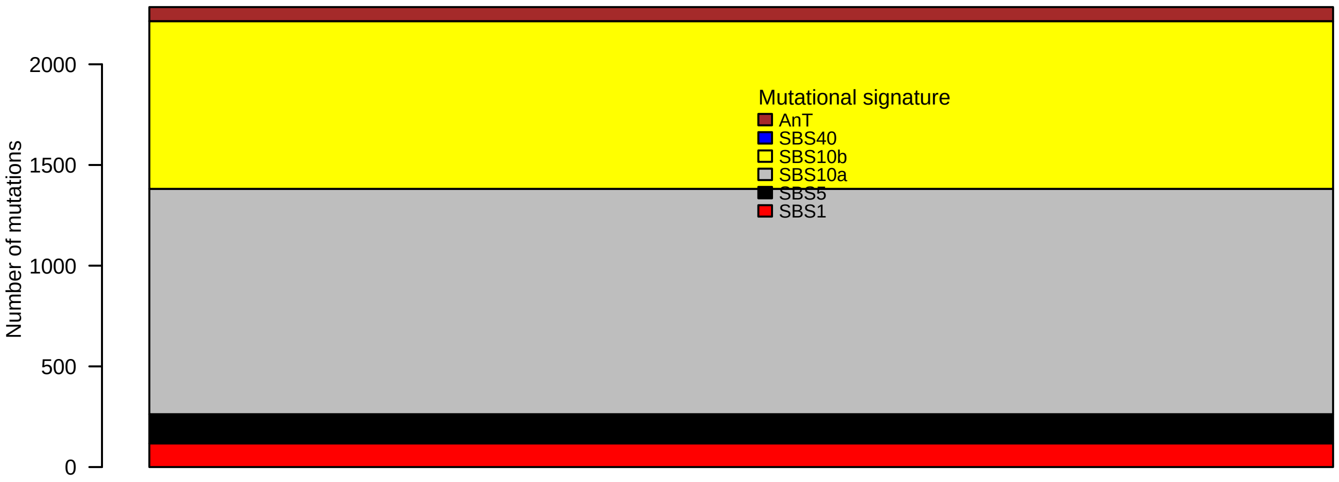

### BD121T.pval.histogram.pdf

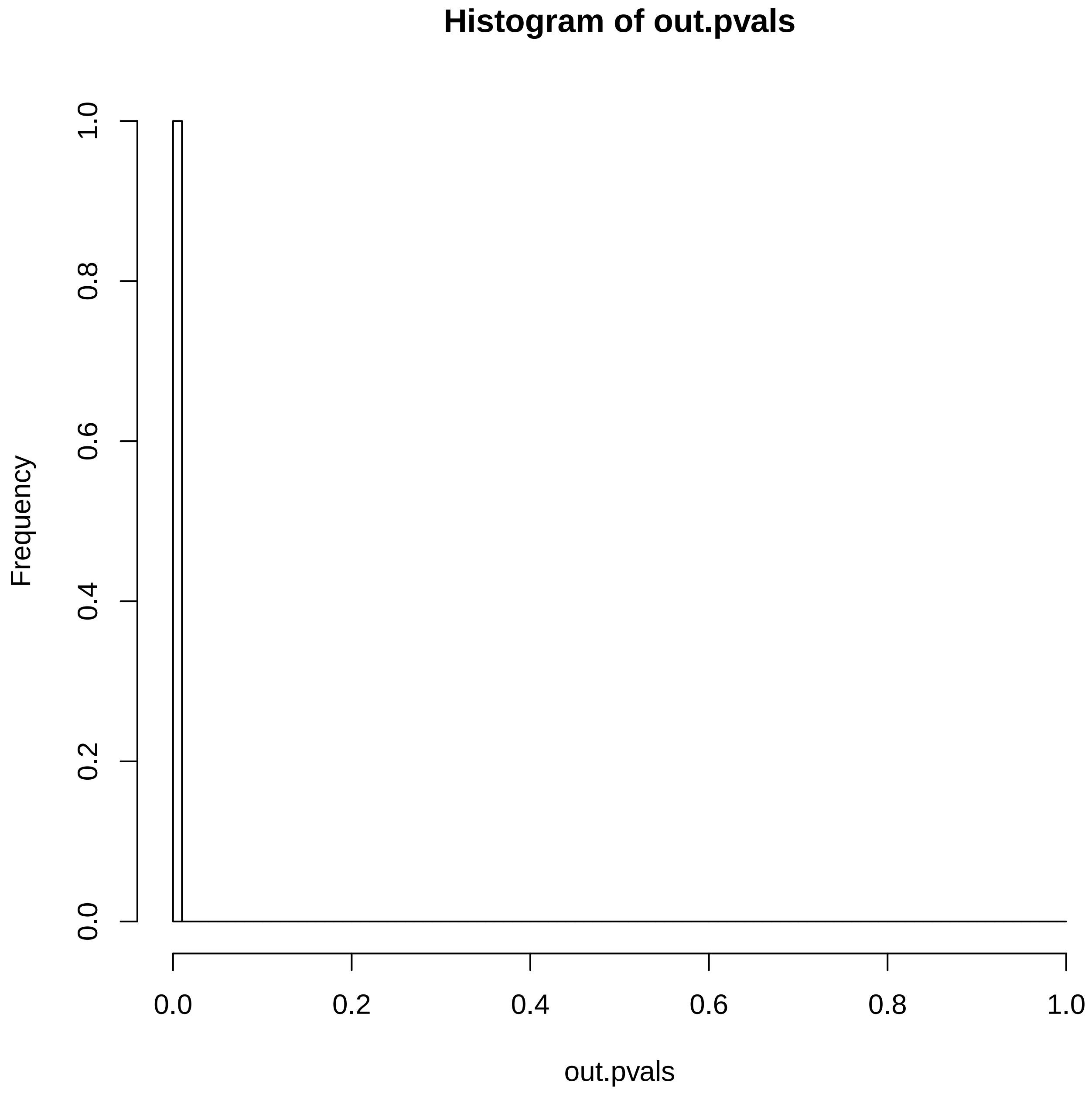

### BD121T.reconstruction.err.pdf

##### Reconstruction error

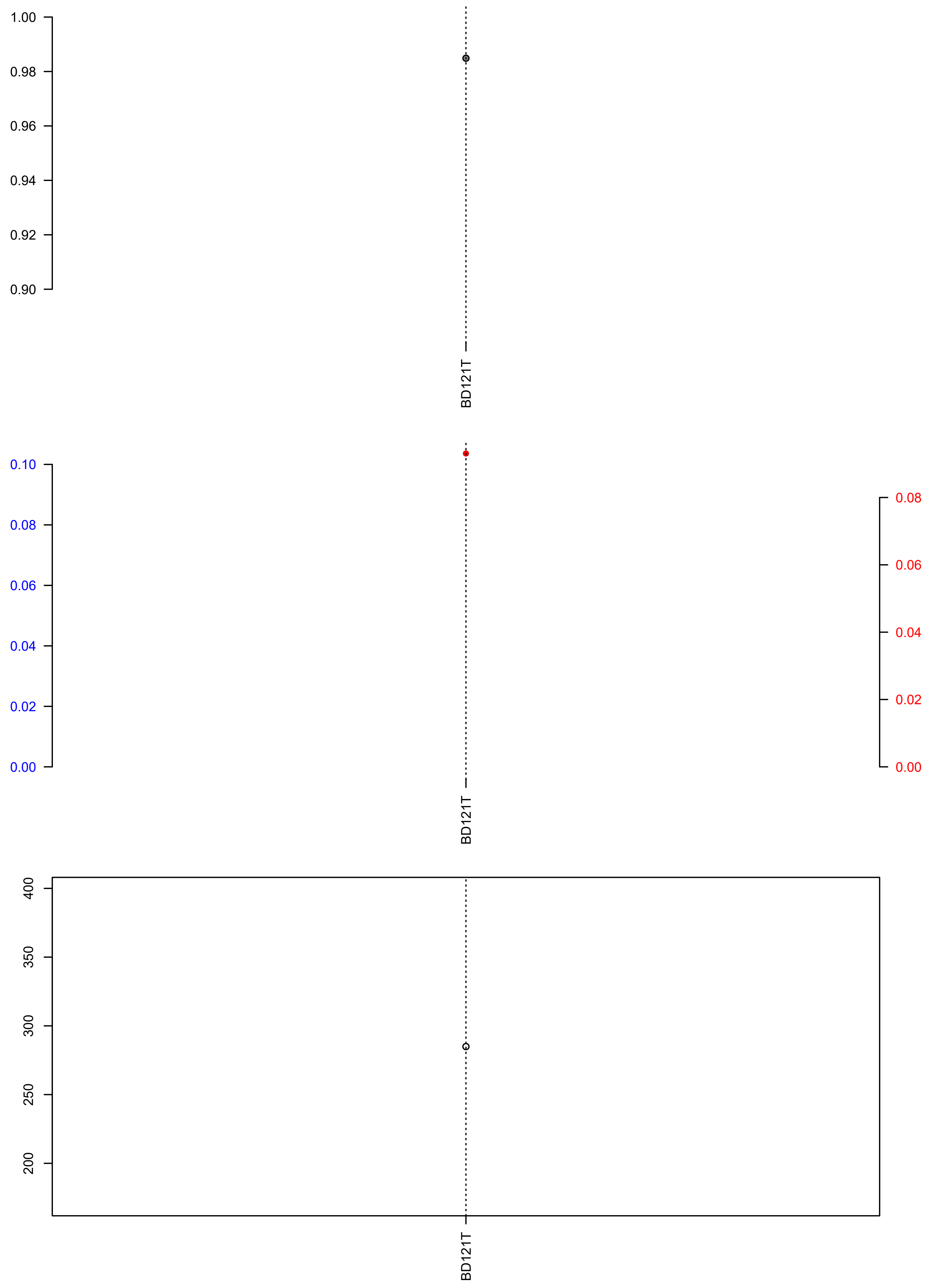

### BD173T.check.with.sig.pdf

# BD173T

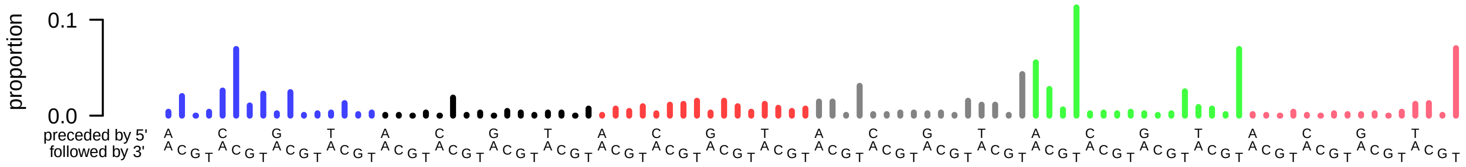

### BD173T.exposures.pdf

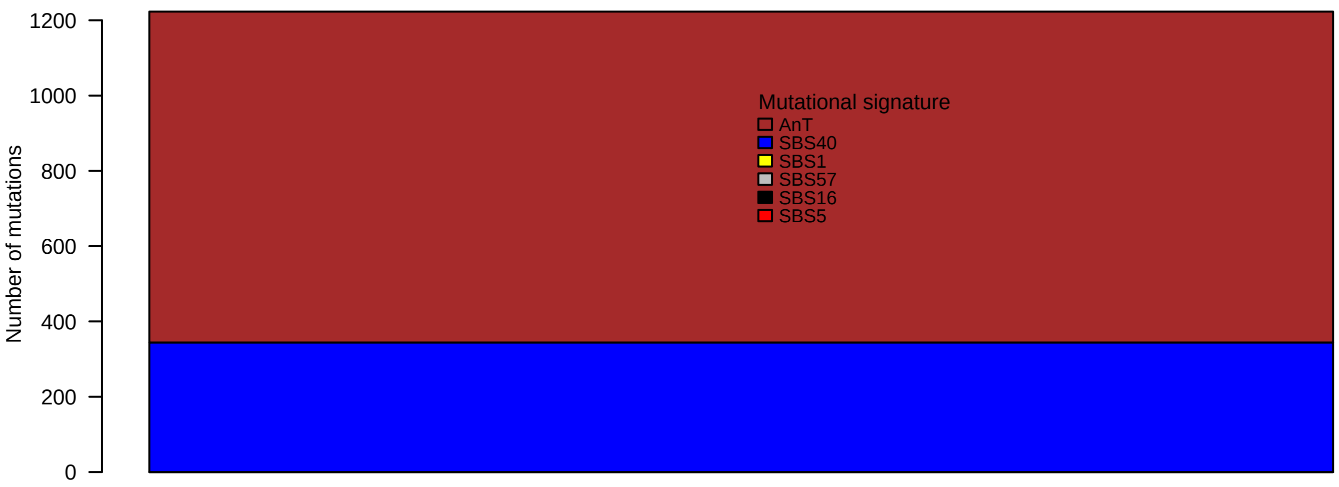

### BD173T.pval.histogram.pdf

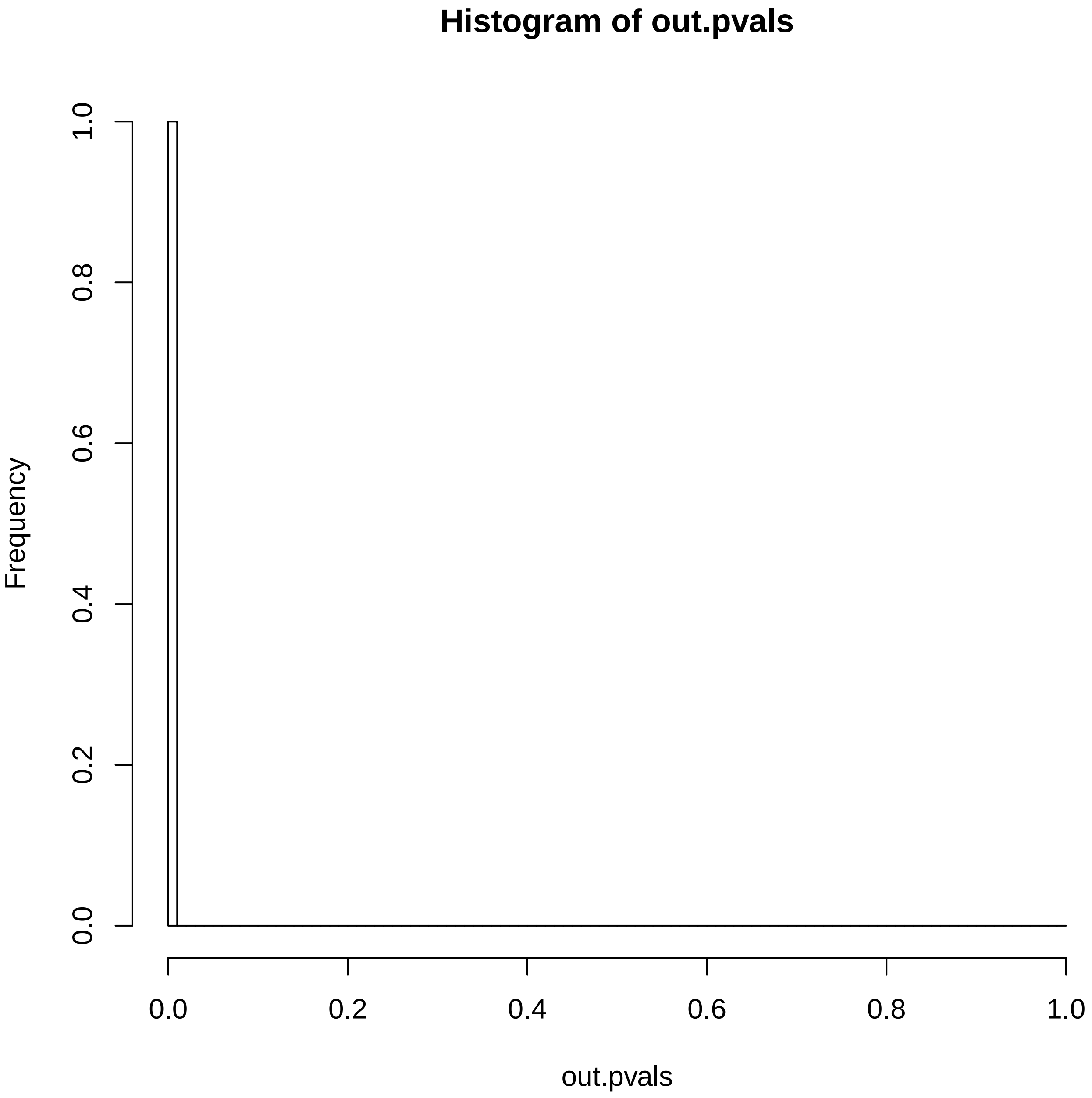

### BD173T.reconstruction.err.pdf

##### Reconstruction error

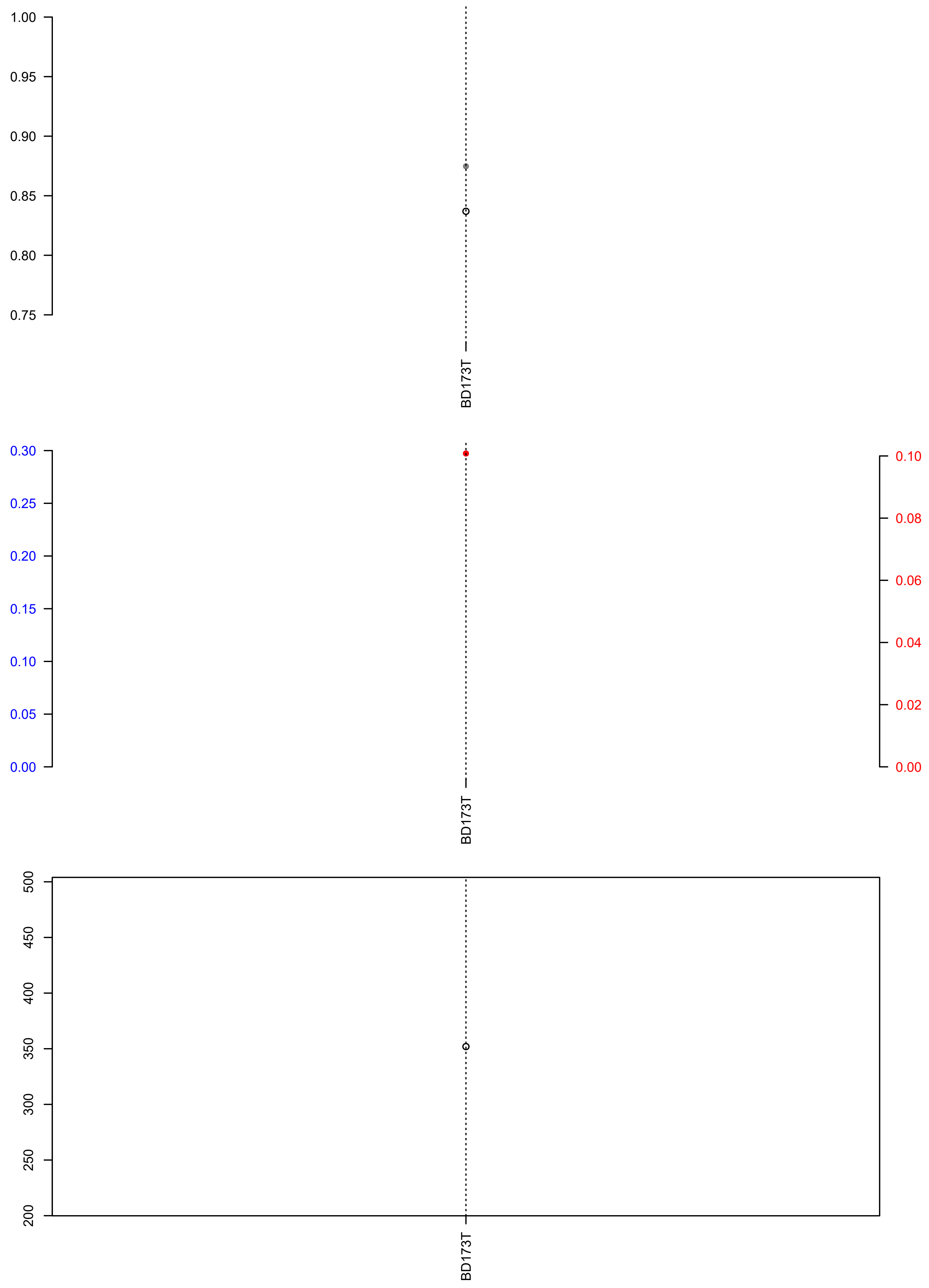

### BD182T.check.with.sig.pdf

# BD182T

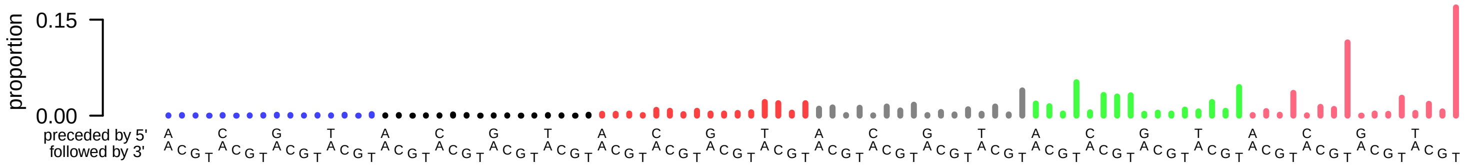

### BD182T.exposures.pdf

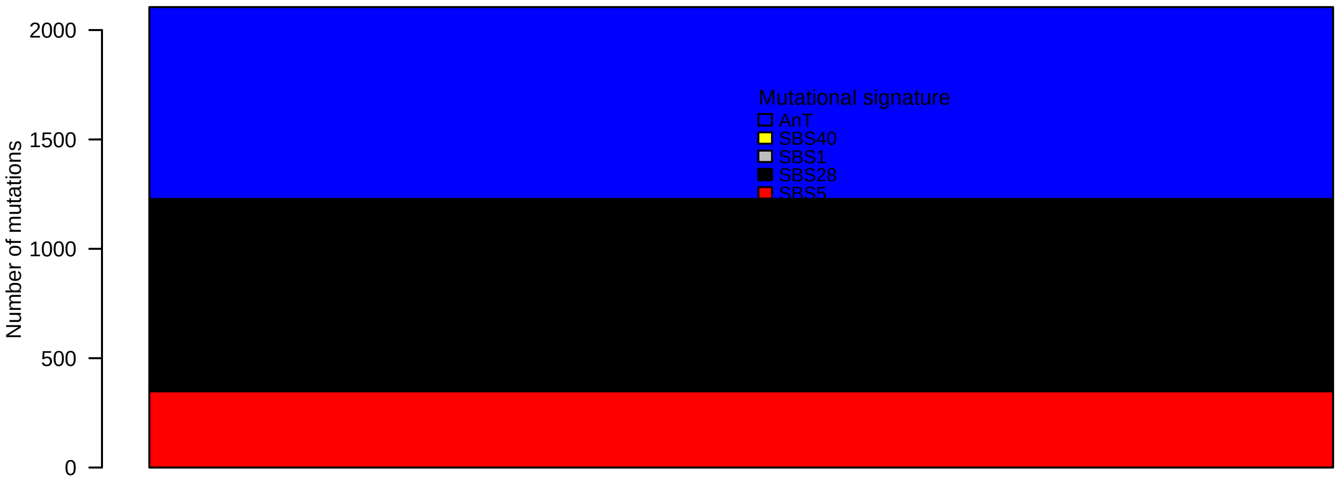

### BD182T.reconstruction.err.pdf

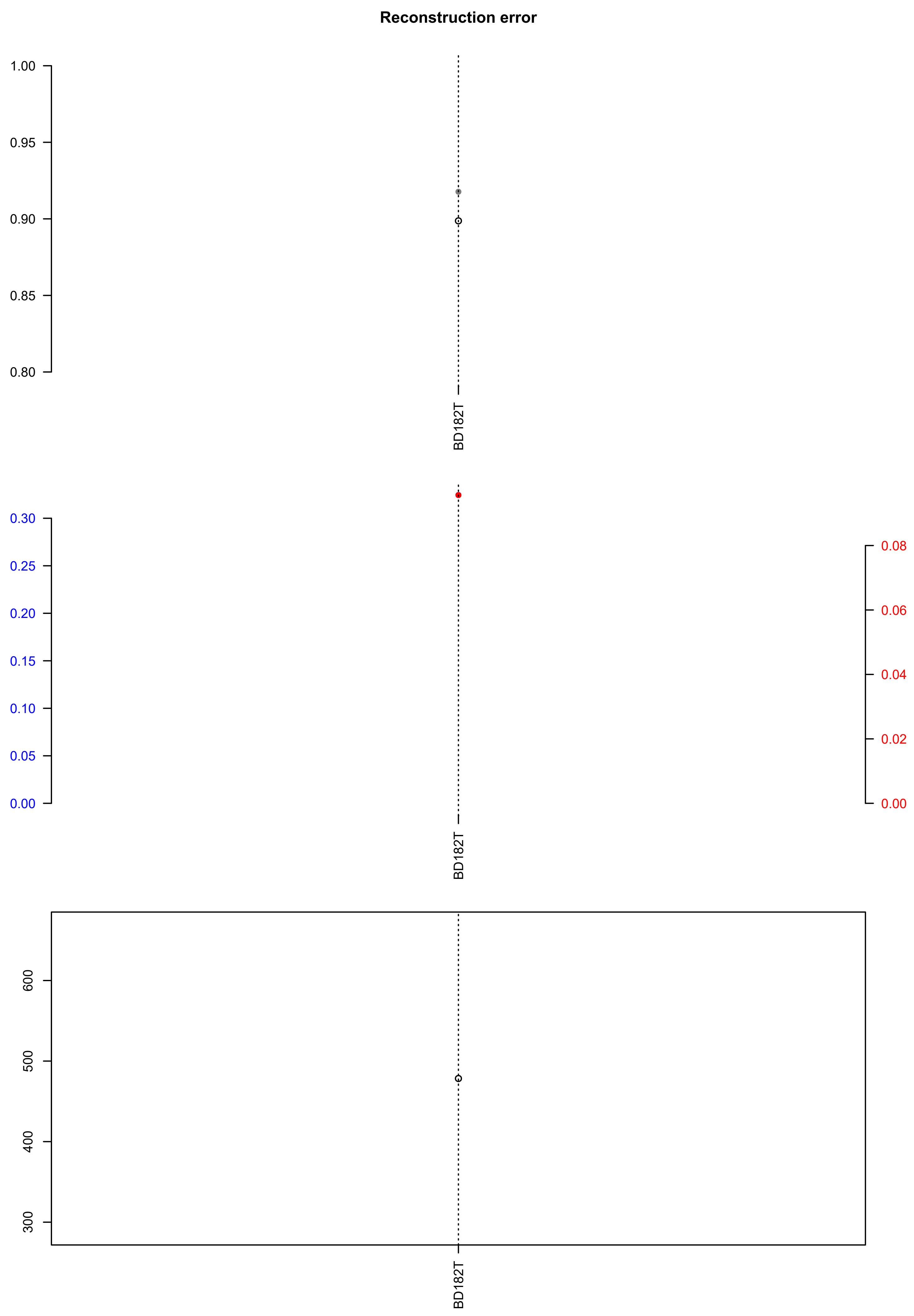

### BD223T.check.with.sig.pdf

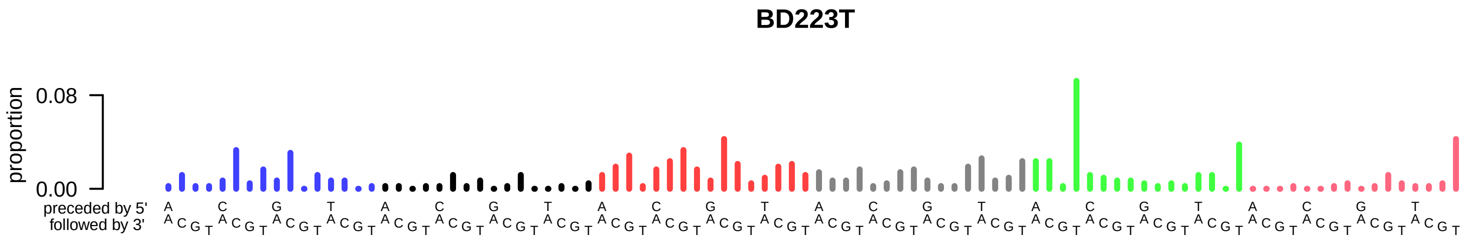

### BD223T.exposures.pdf

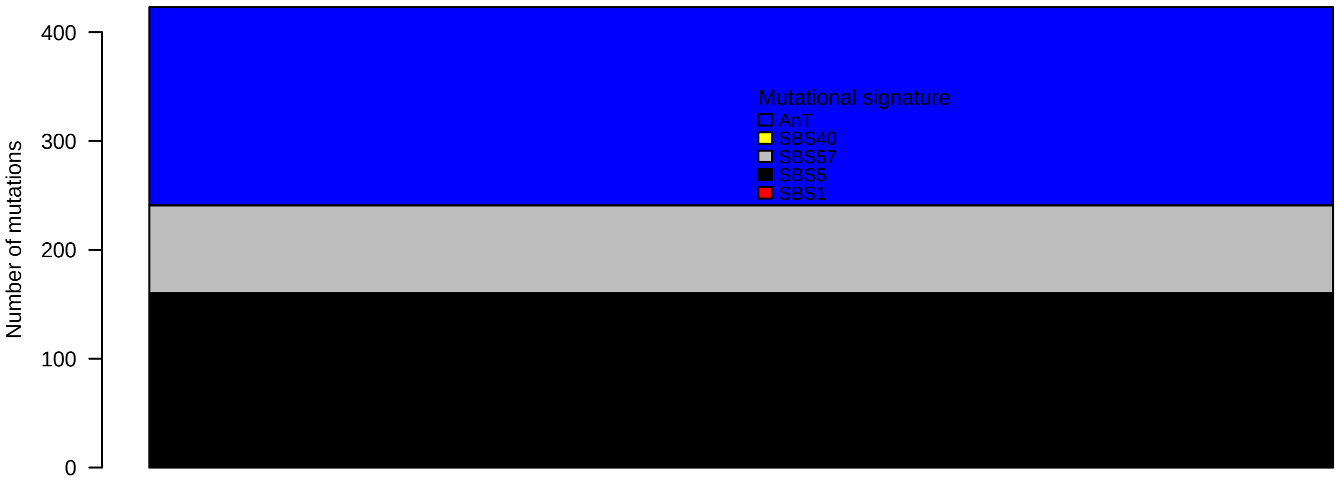

### BD223T.pval.histogram.pdf

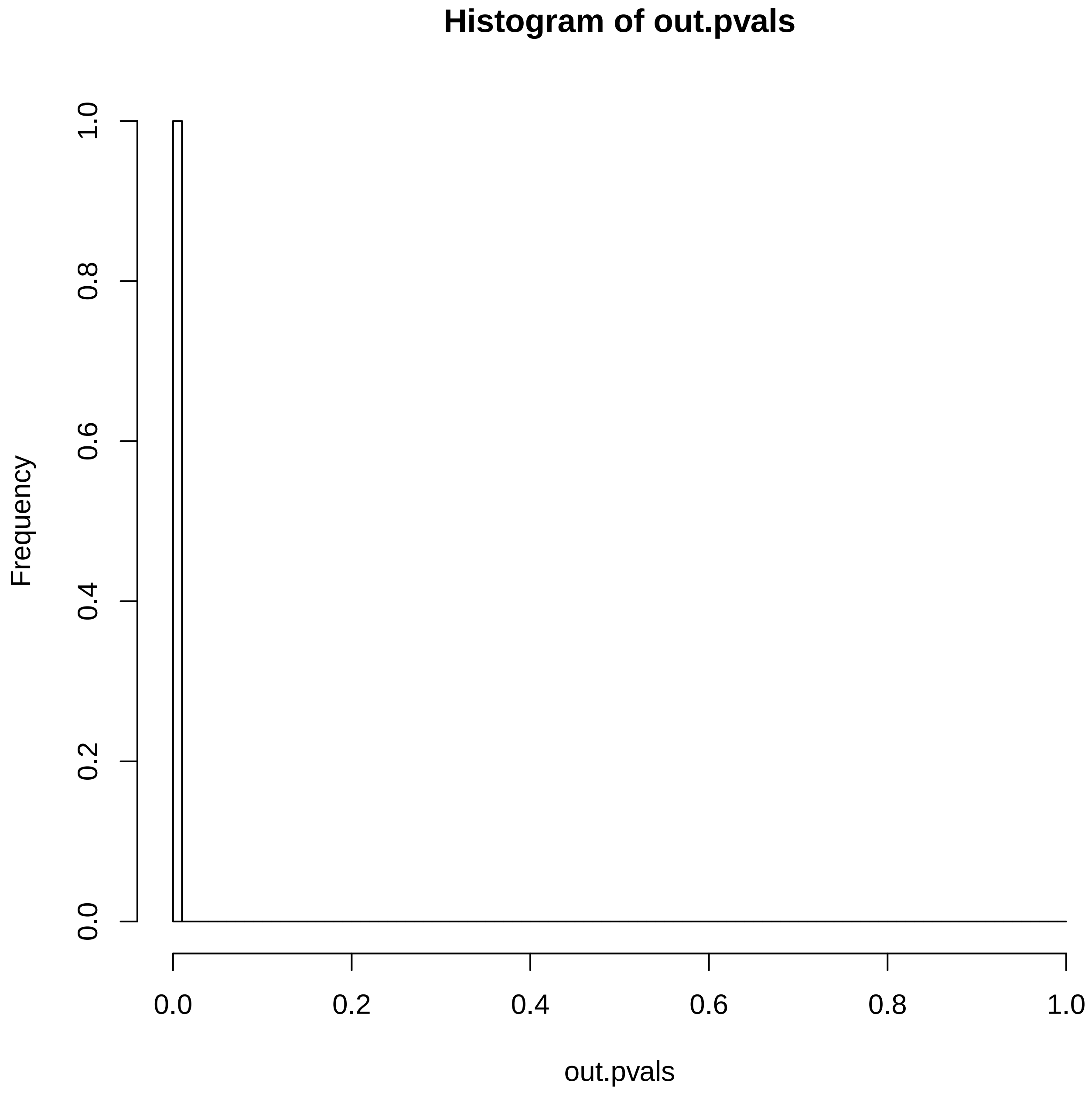

### BD223T.reconstruction.err.pdf

##### Reconstruction error

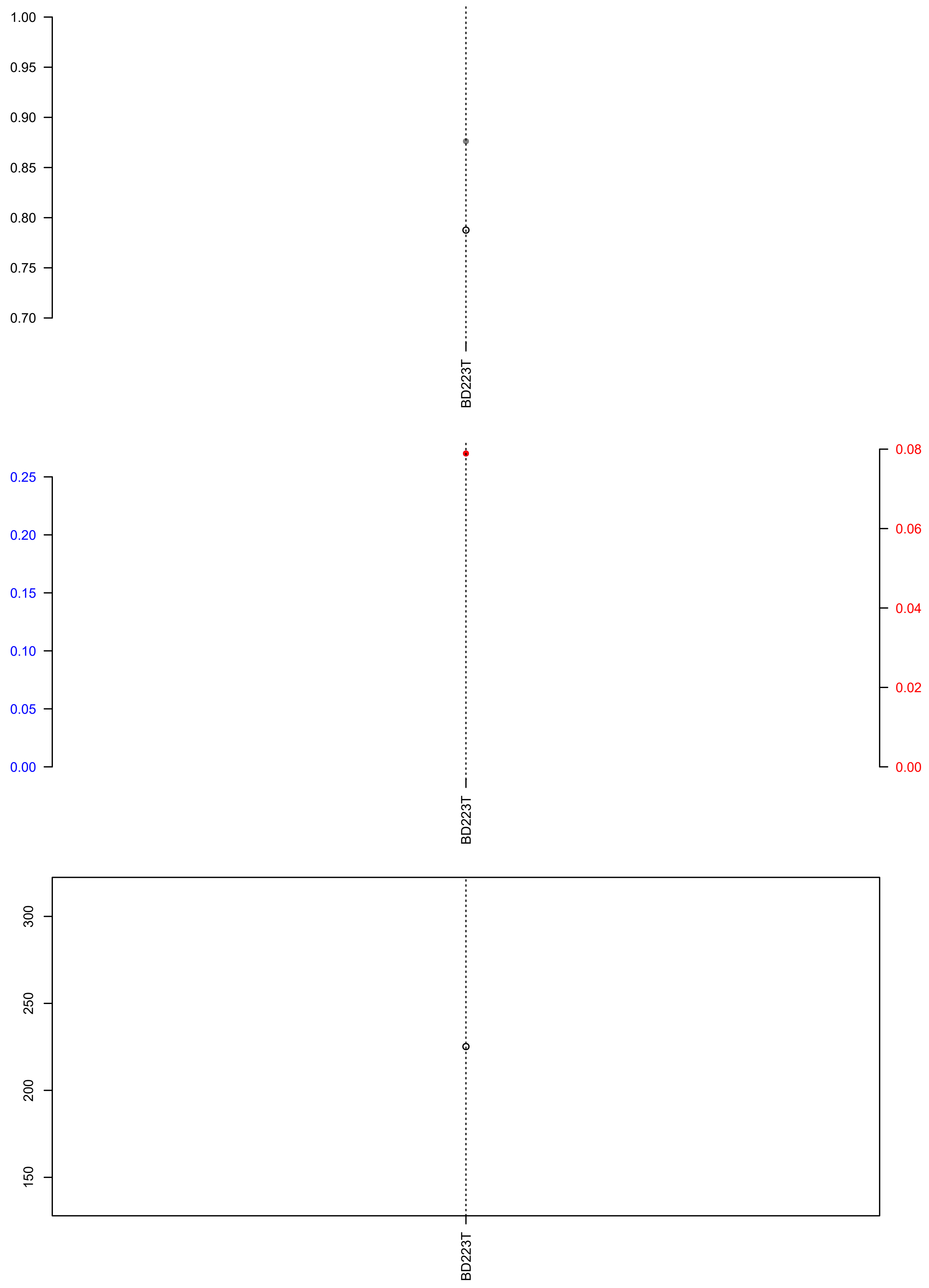

### ESO-173.exposures.pdf

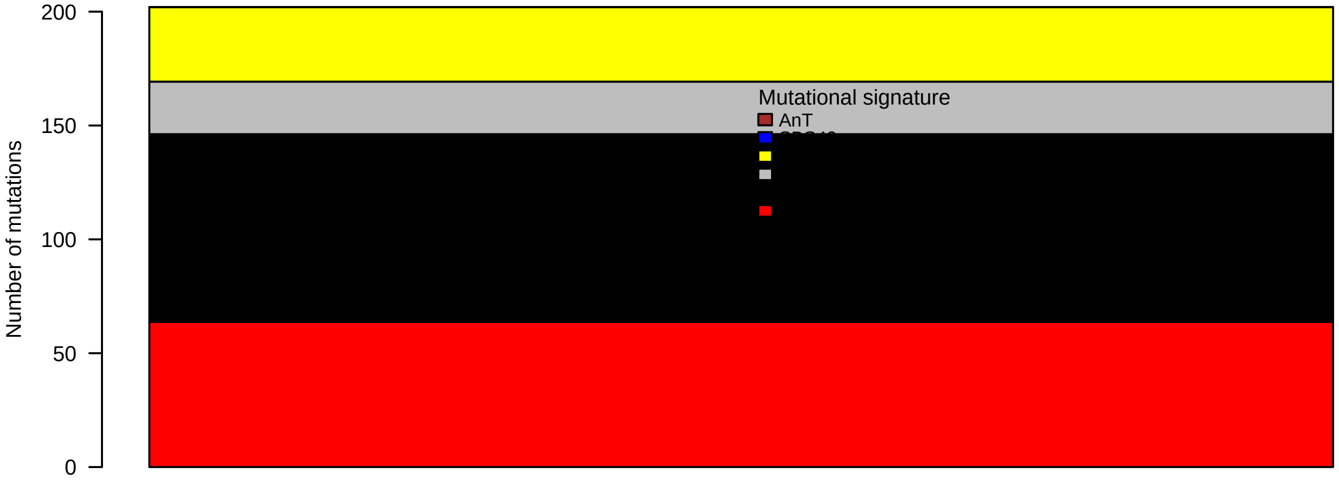

### ESO-173.pval.histogram.pdf

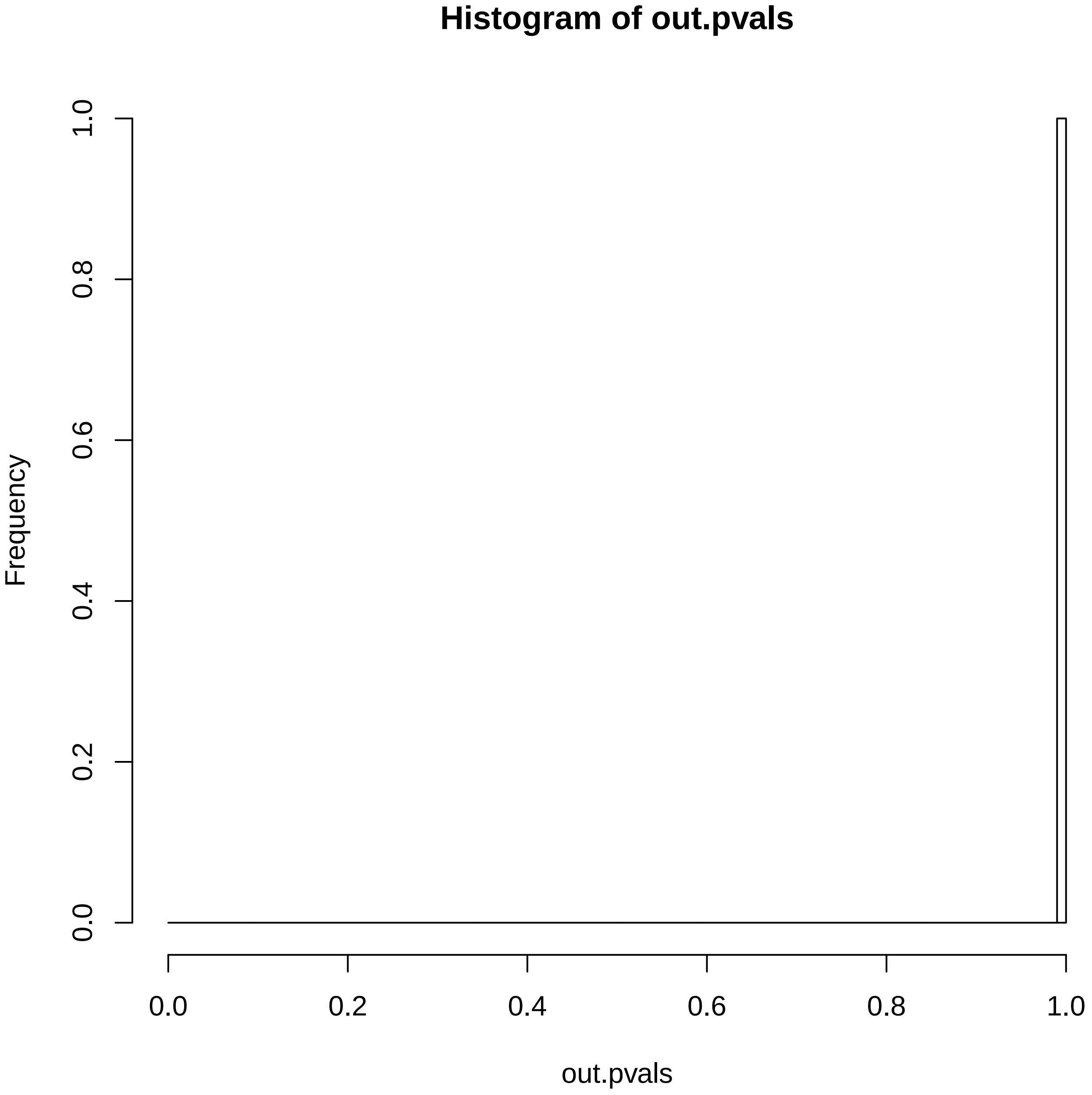

### ESO-173.reconstruction.err.pdf

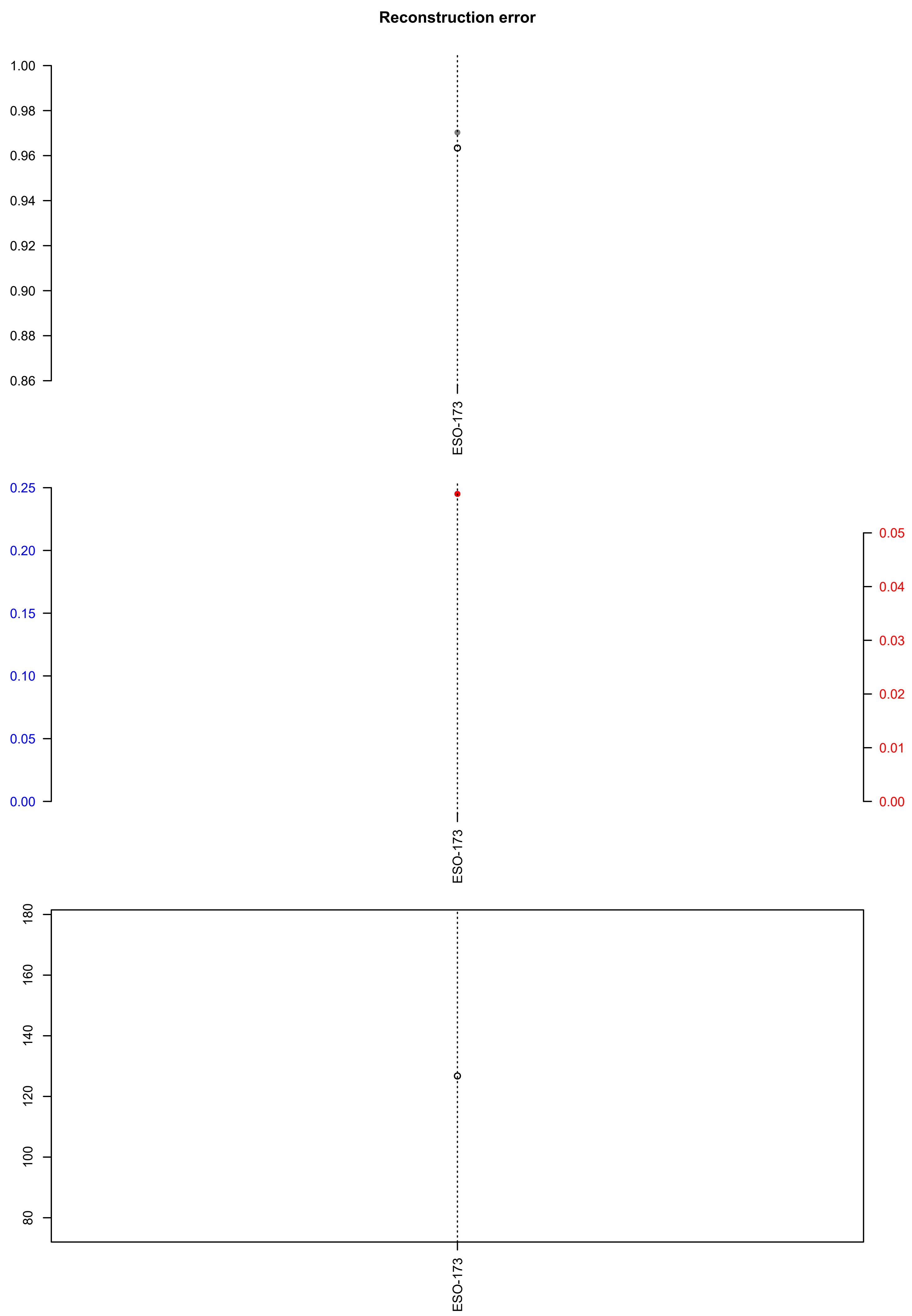

### HCC34T.exposures.pdf

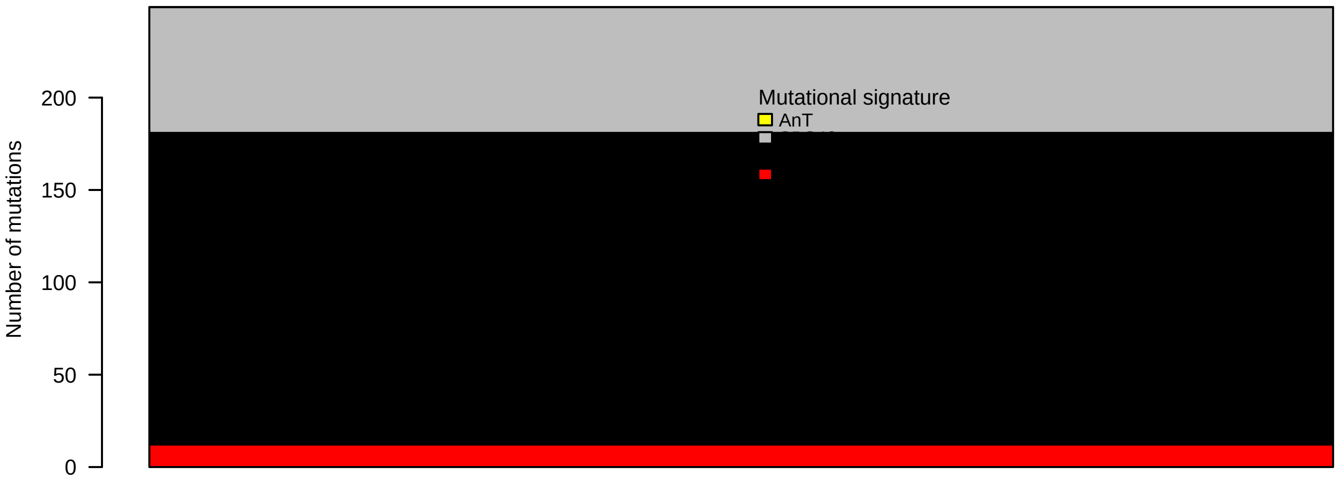

### HCC34T.pval.histogram.pdf

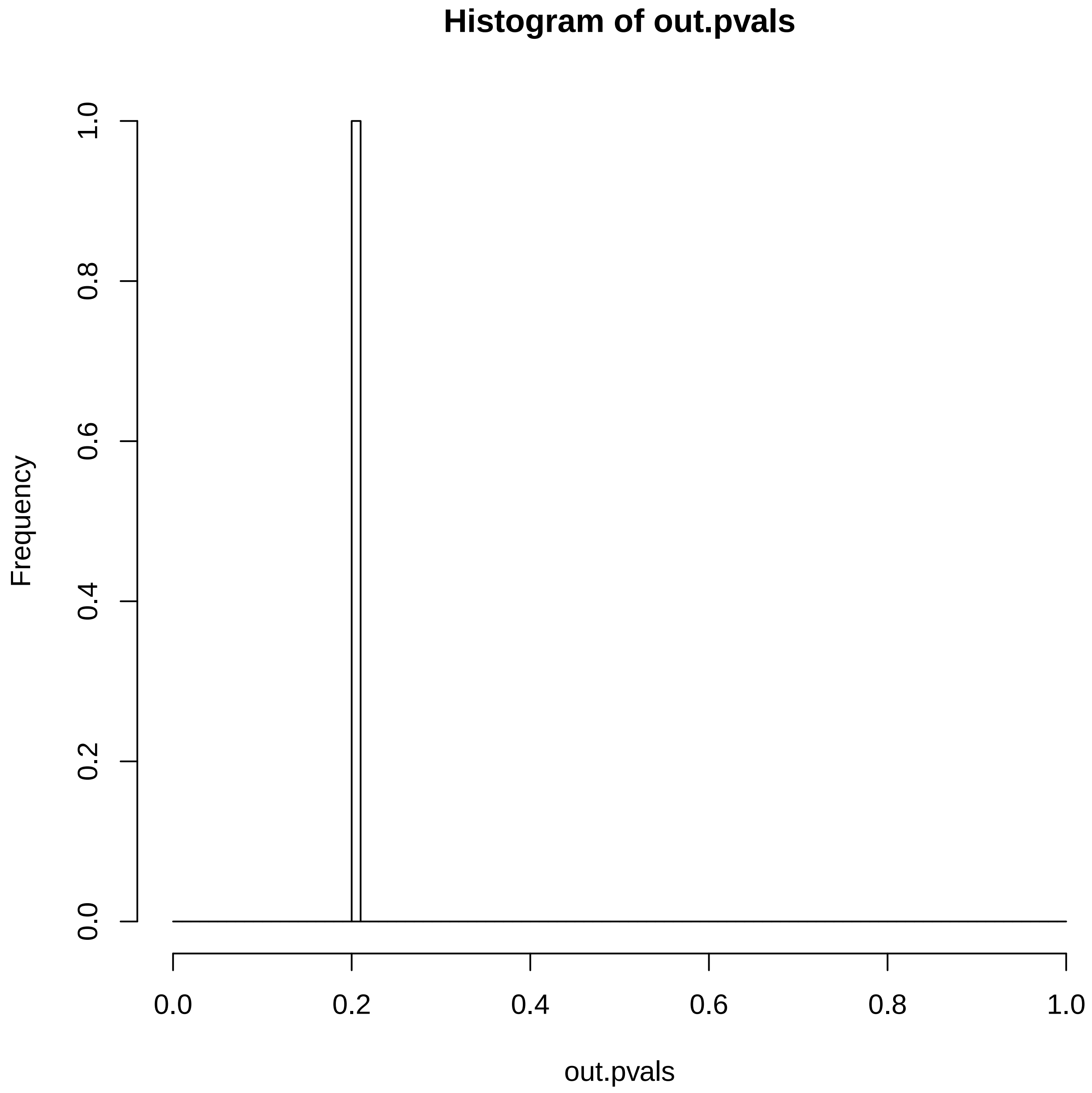

### HCC34T.reconstruction.err.pdf

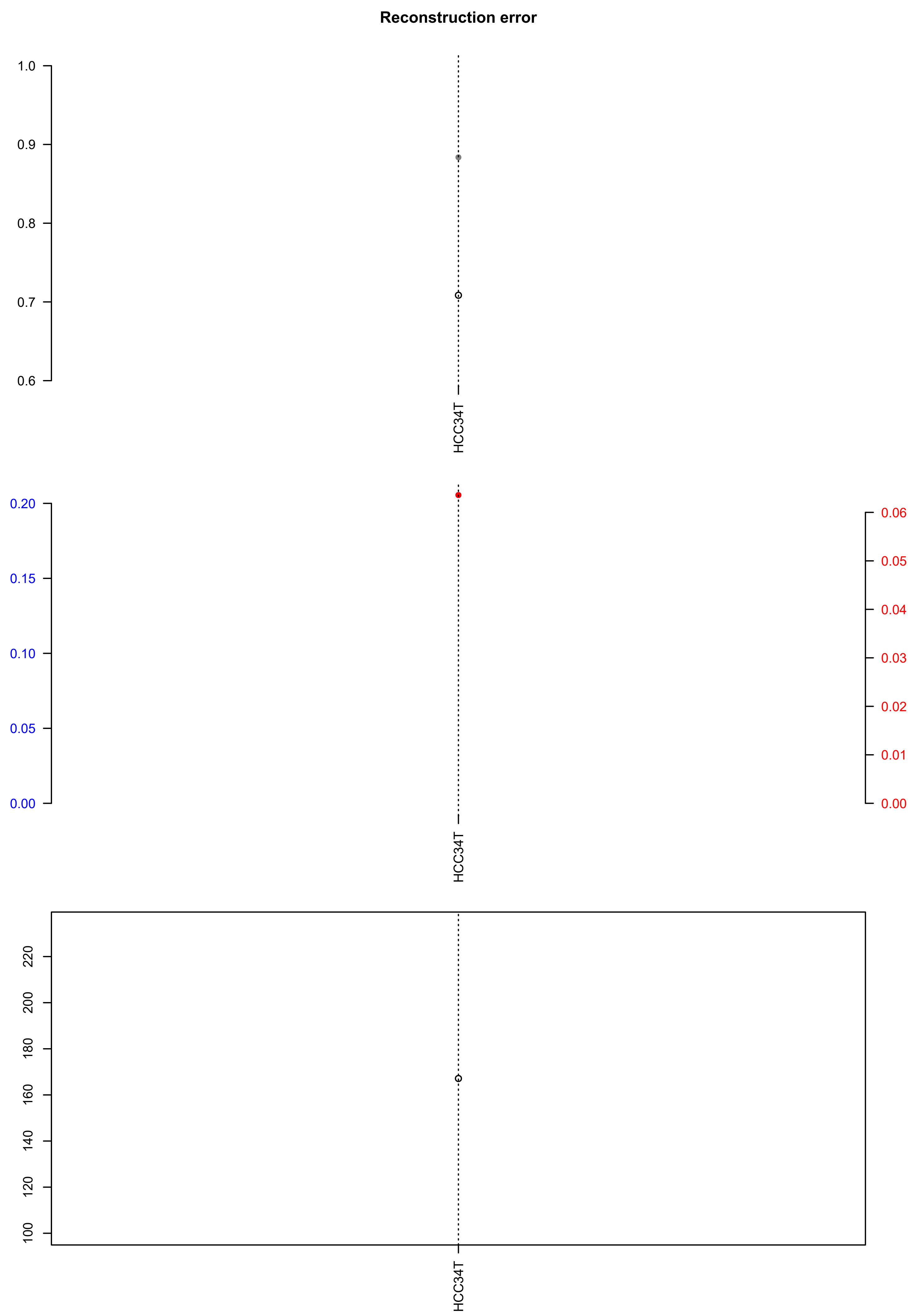

### LP6005935-DNA_B03.check.with.sig.pdf

##### LP6005935.DNA_B03

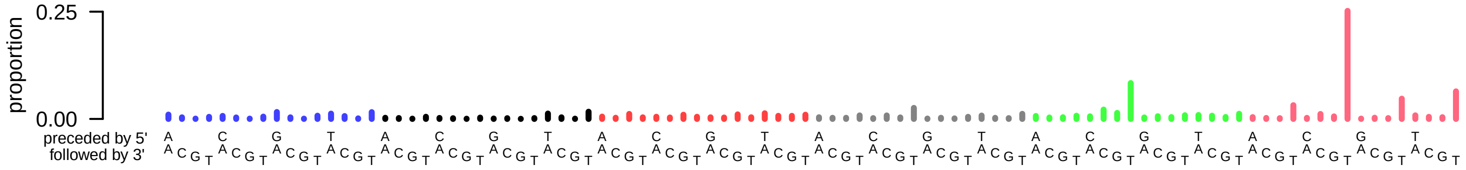

### LP6005935-DNA_B03.exposures.pdf

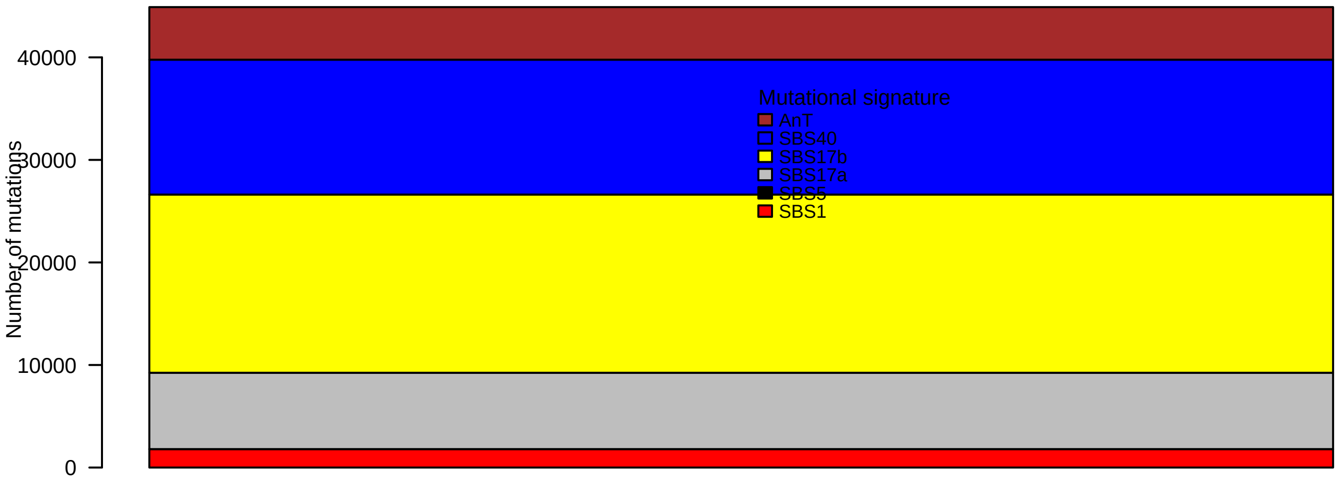

### LP6005935-DNA_B03.pval.histogram.pdf

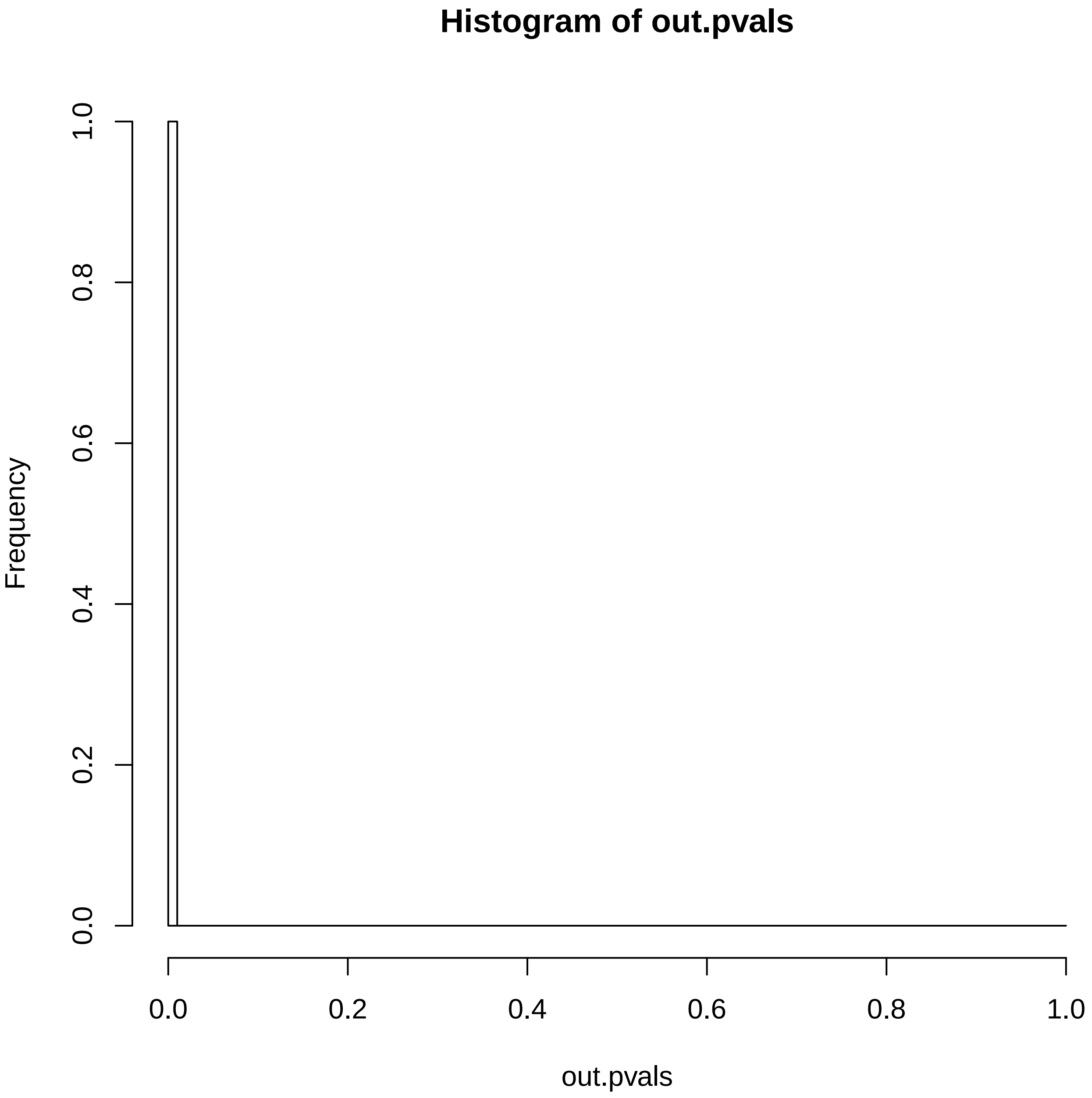

### LP6005935-DNA_B03.reconstruction.err.pdf

##### Reconstruction error

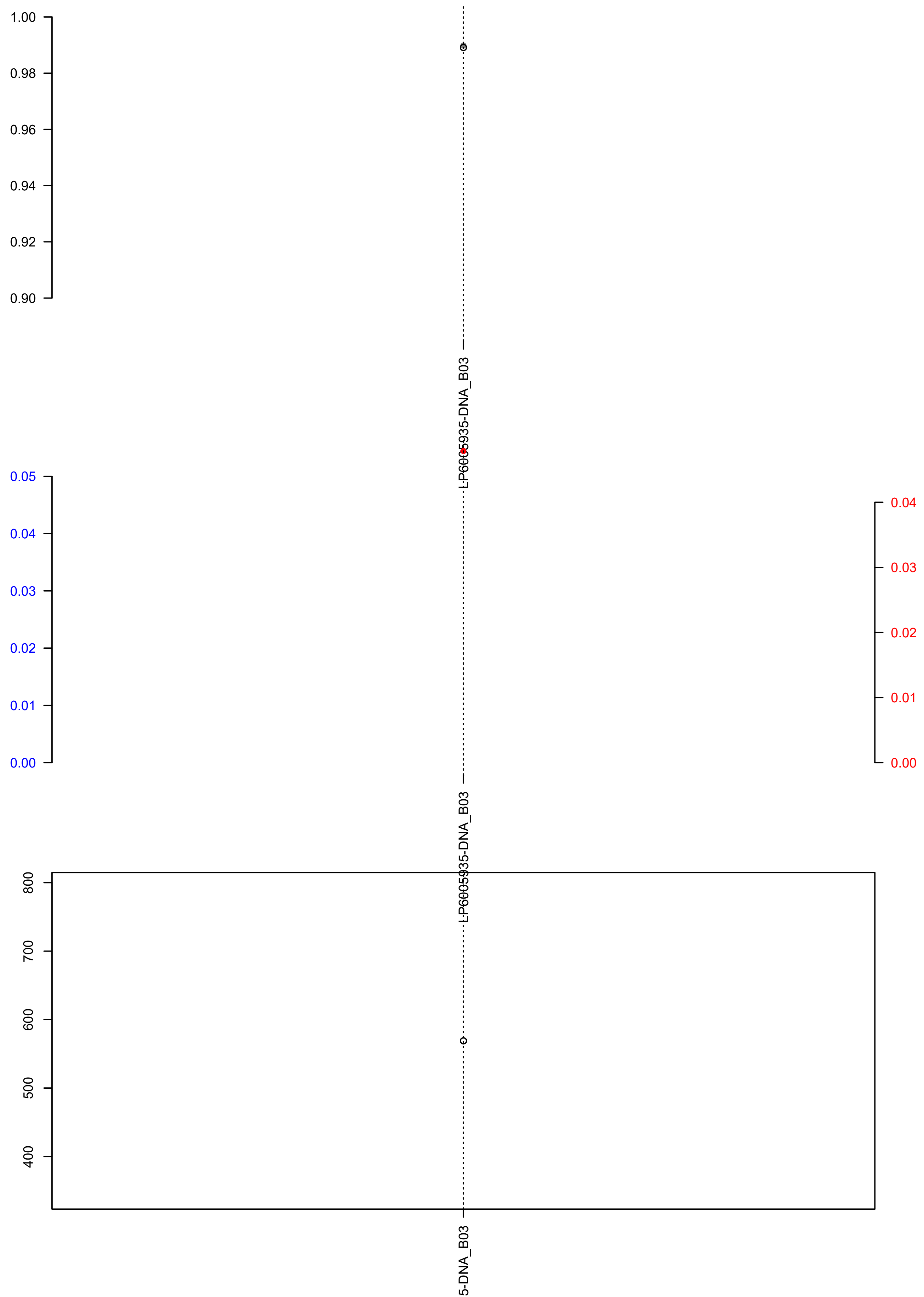

### PCSI_0060_Pa_X.check.with.sig.pdf

##### PCSI_0060_Pa_X

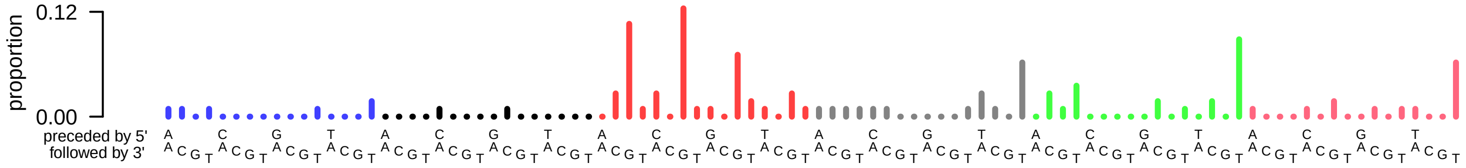

### PCSI_0060_Pa_X.exposures.pdf

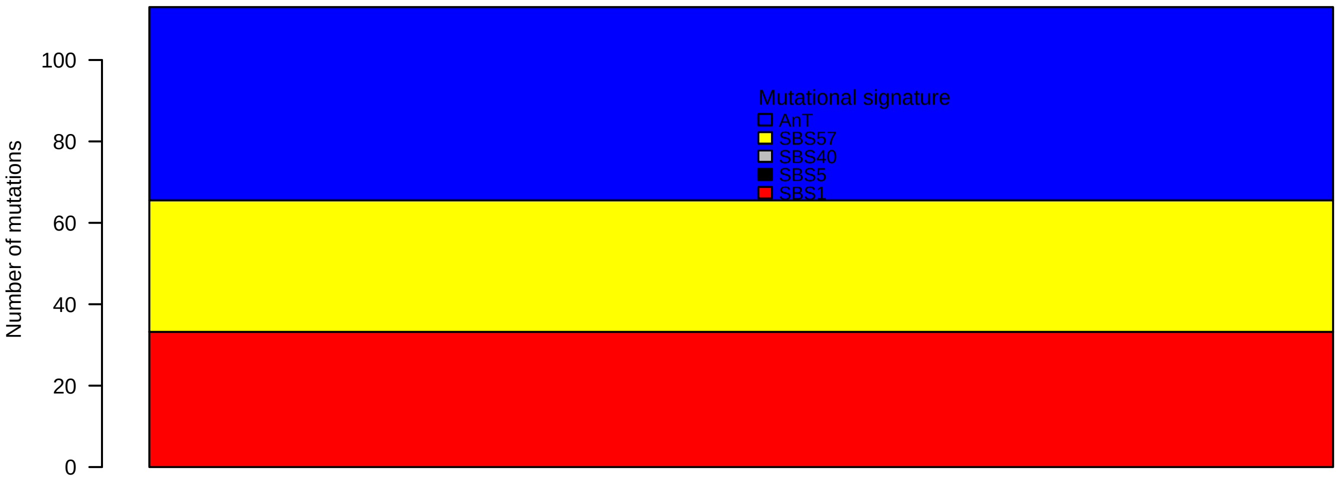

### PCSI_0060_Pa_X.pval.histogram.pdf

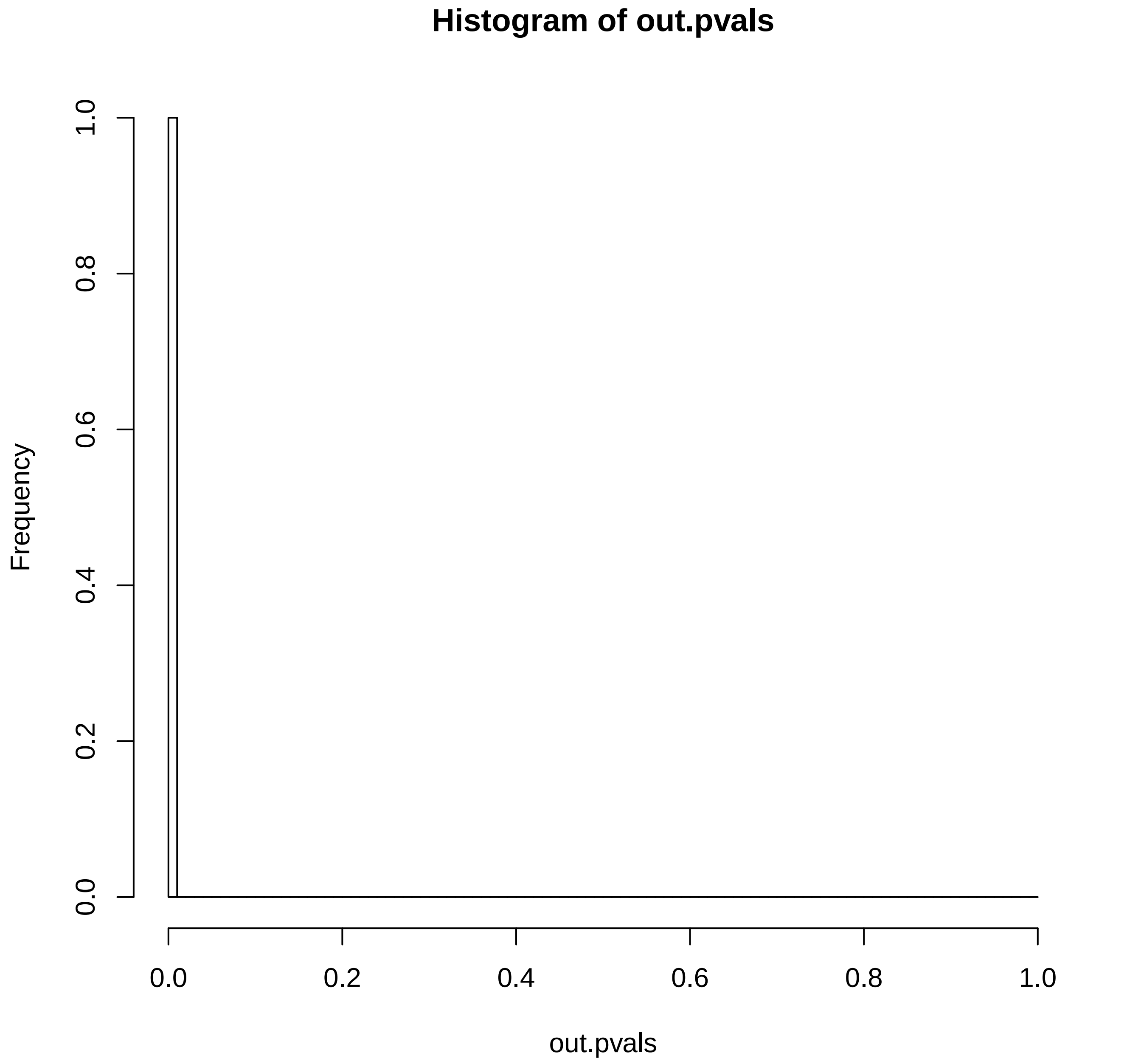

### PCSI_0060_Pa_X.reconstruction.err.pdf

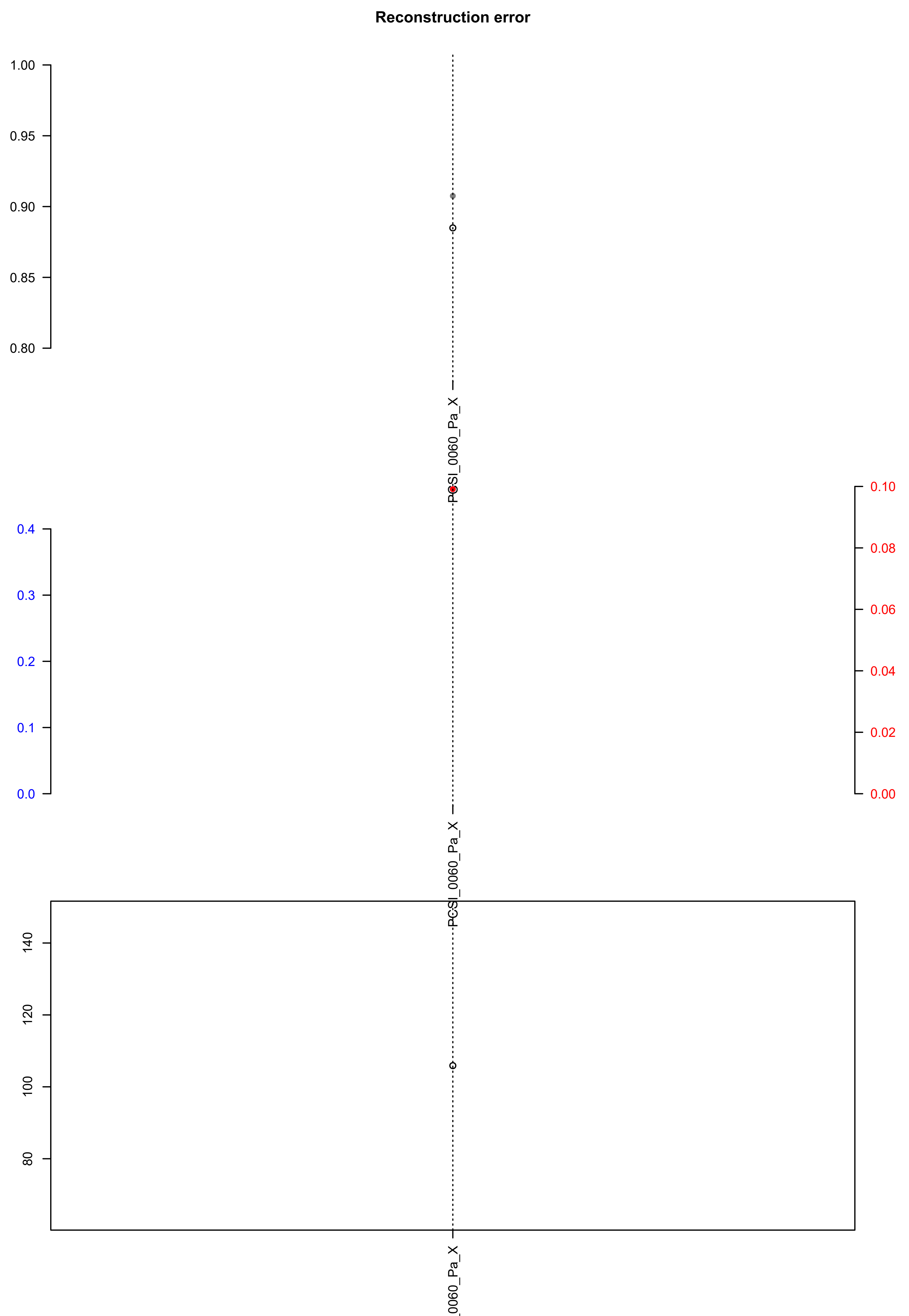

### SKCM-JWCI-WGS-8-Tumor.exposures.pdf

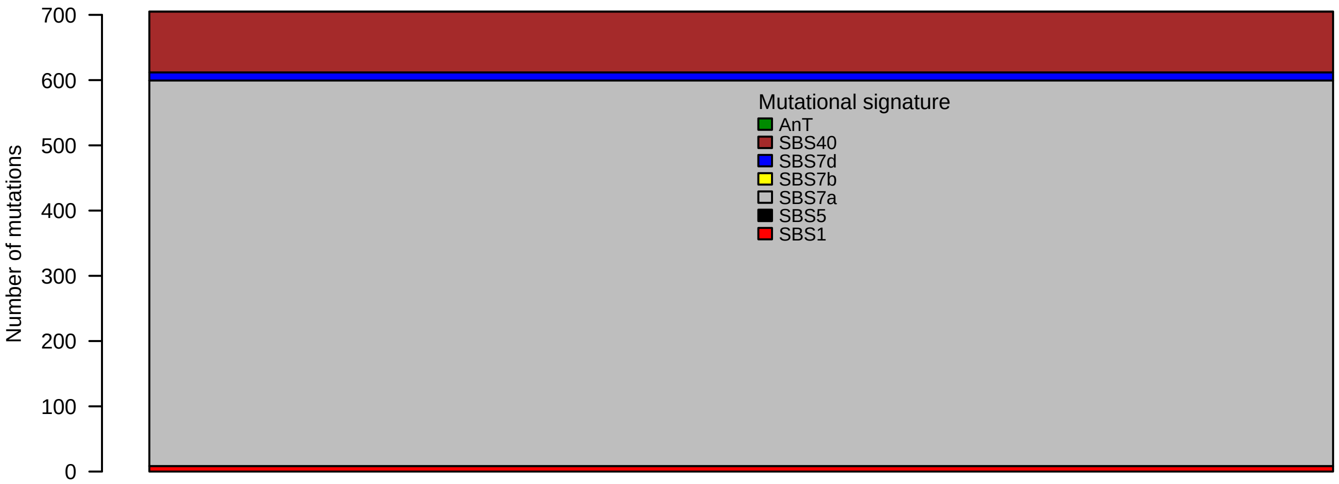

### SKCM-JWCI-WGS-8-Tumor.pval.histogram.pdf

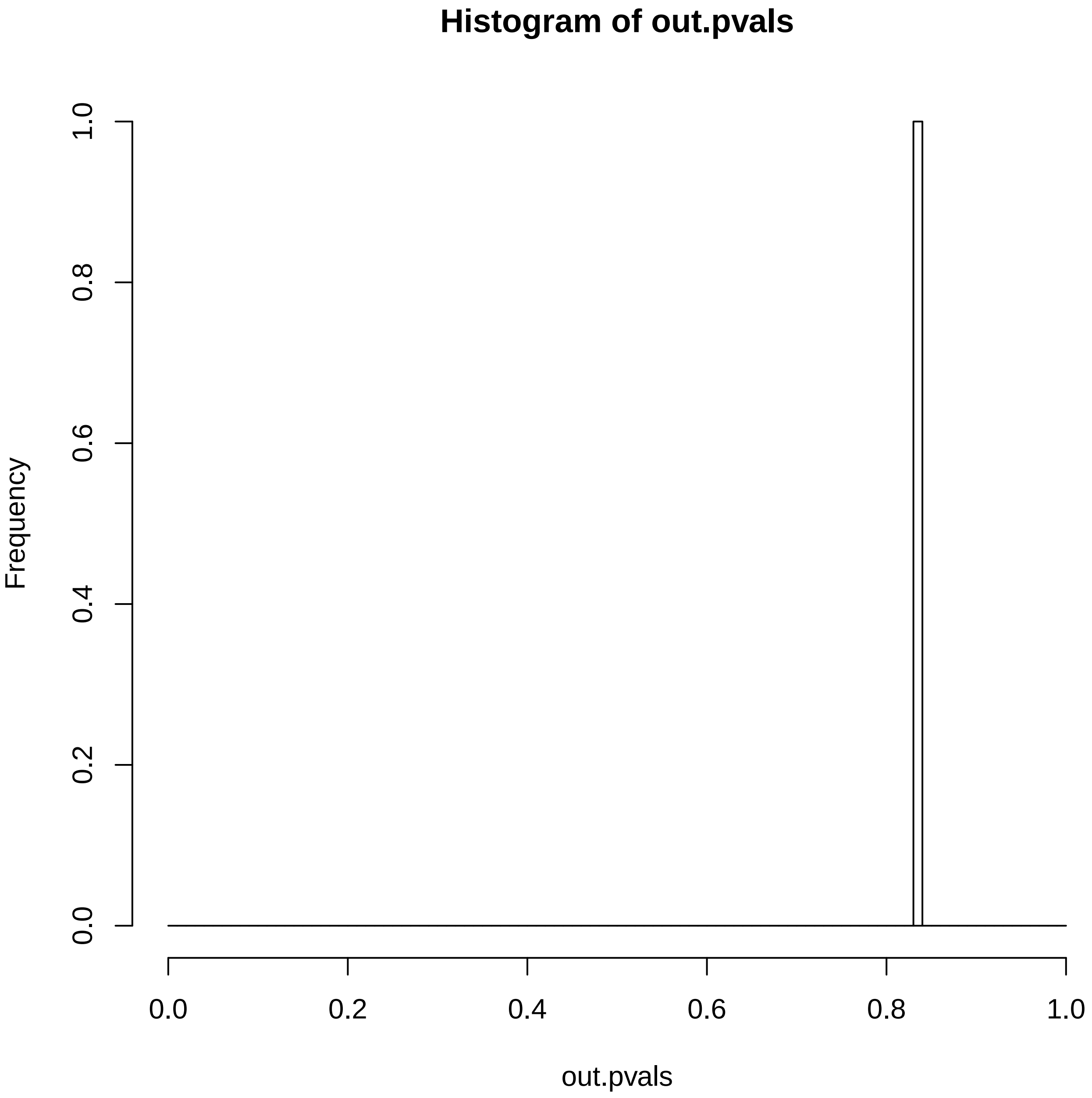

### SKCM-JWCI-WGS-8-Tumor.reconstruction.err.pdf

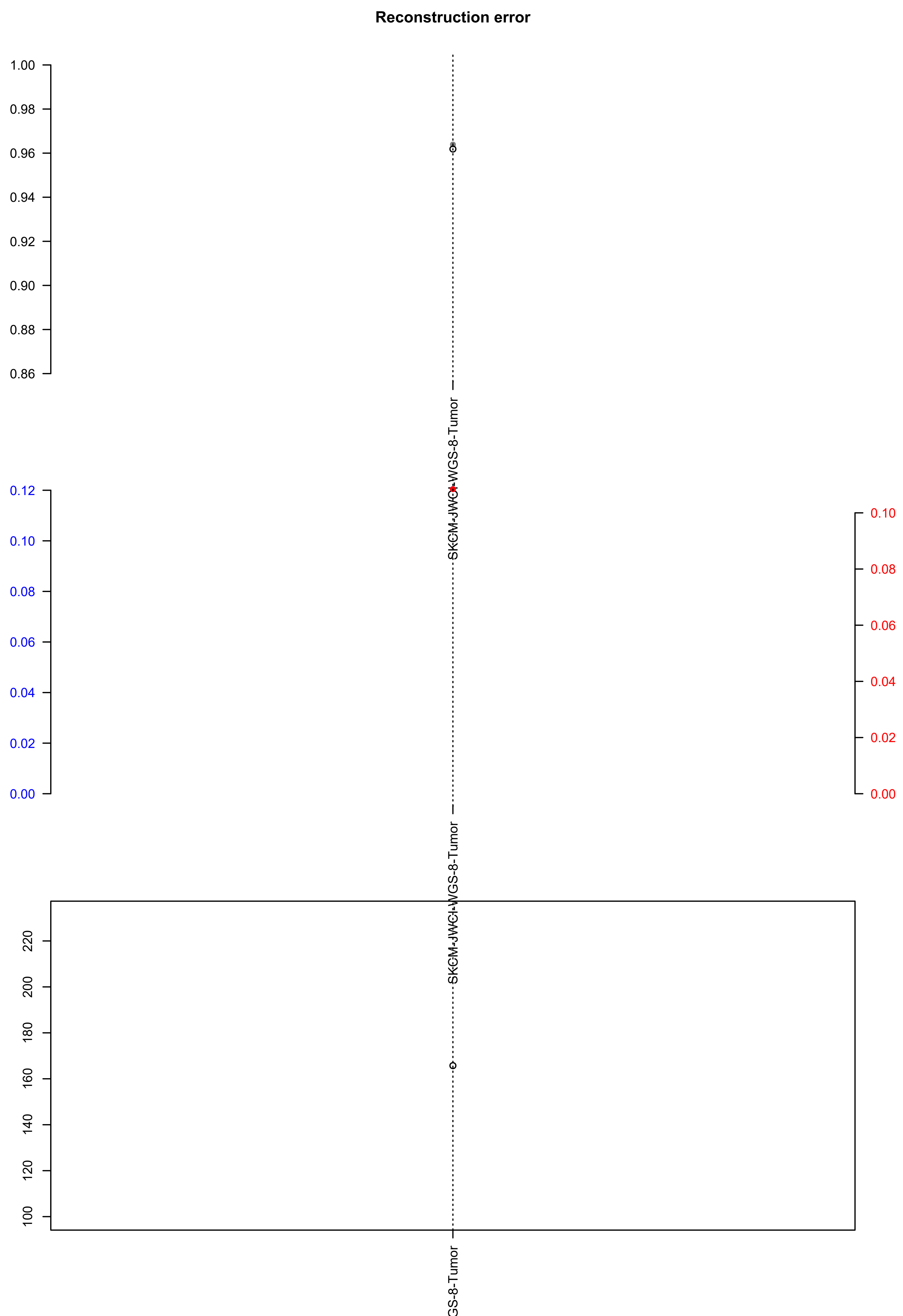

### SP16886.exposures.pdf

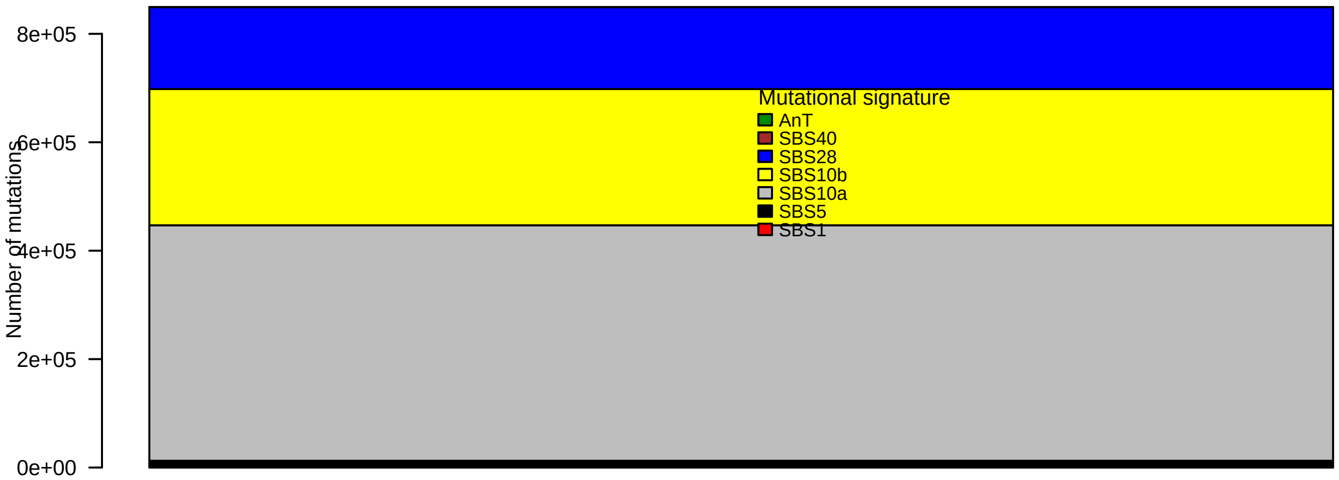

### SP16886.pval.histogram.pdf

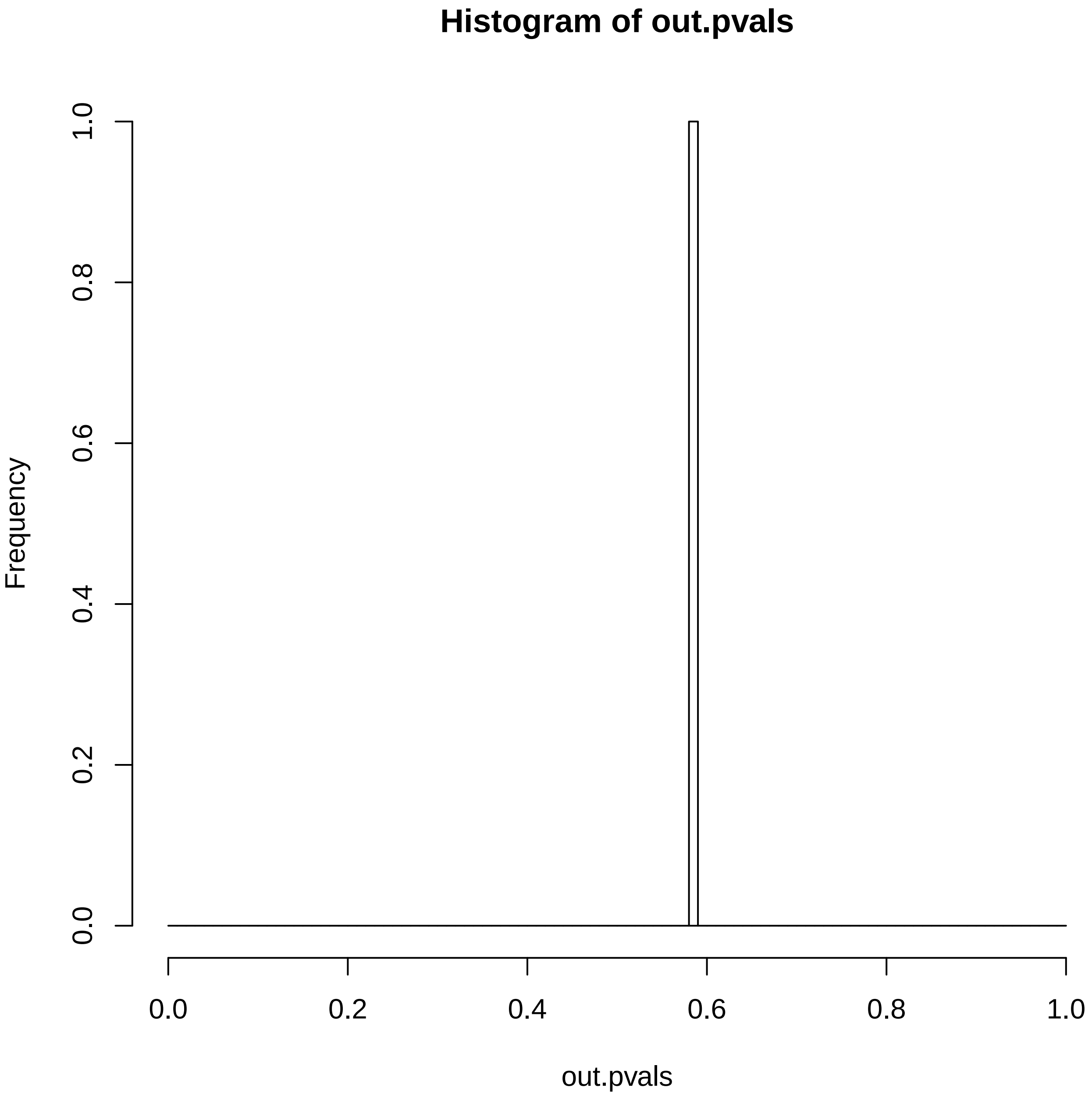

### SP16886.reconstruction.err.pdf

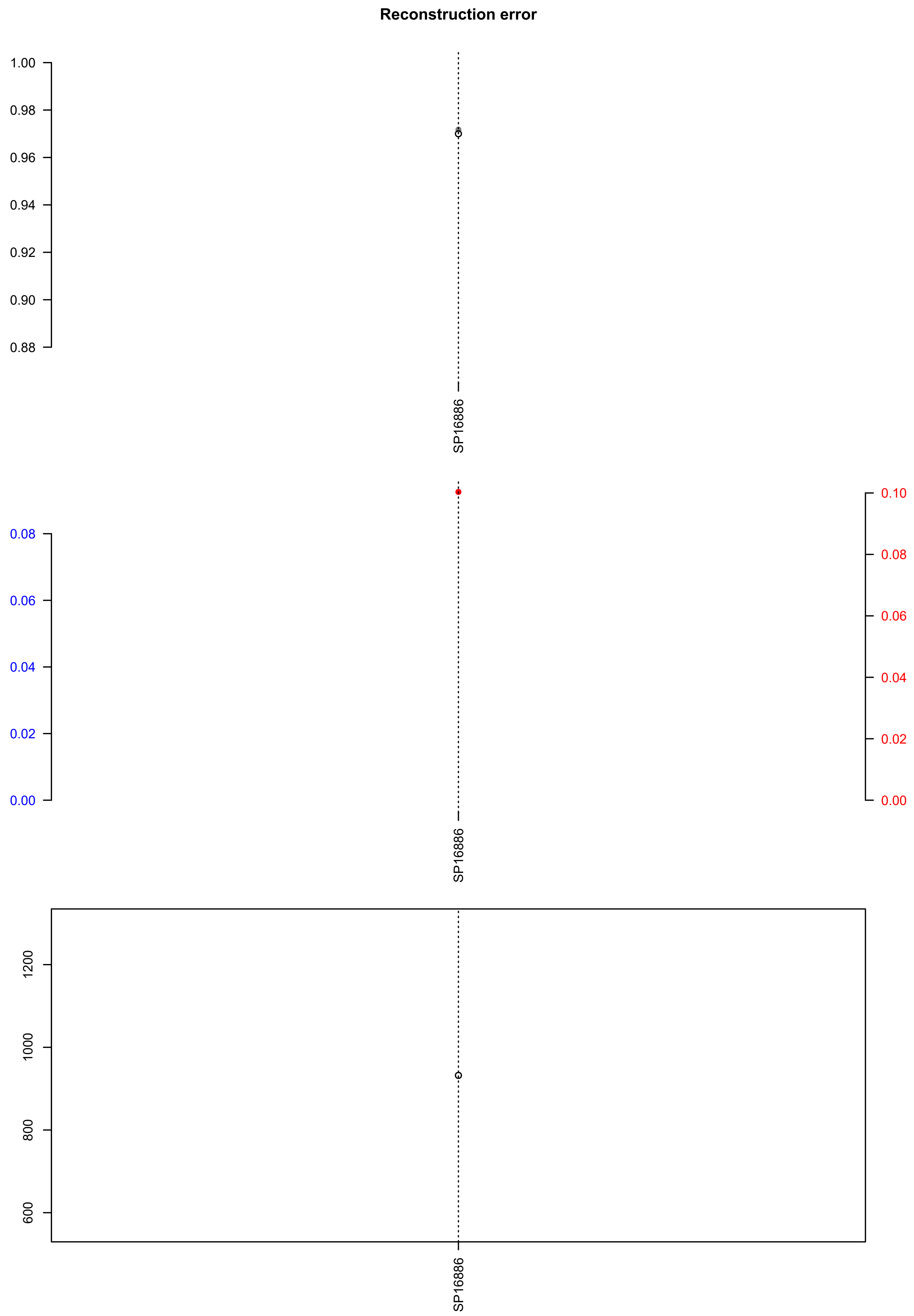

### SP17905.exposures.pdf

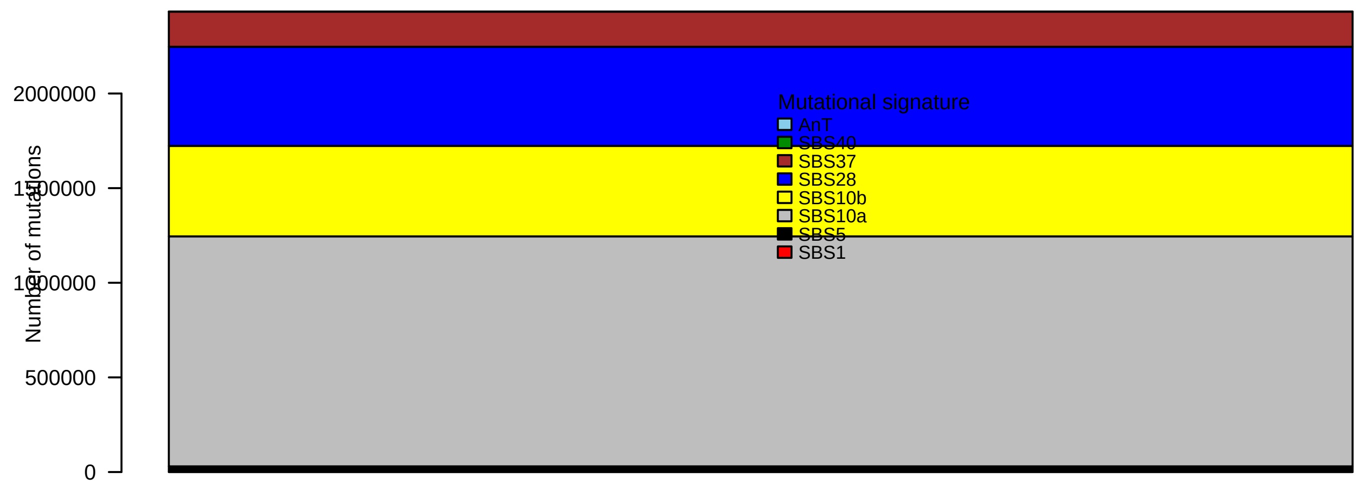

### SP17905.pval.histogram.pdf

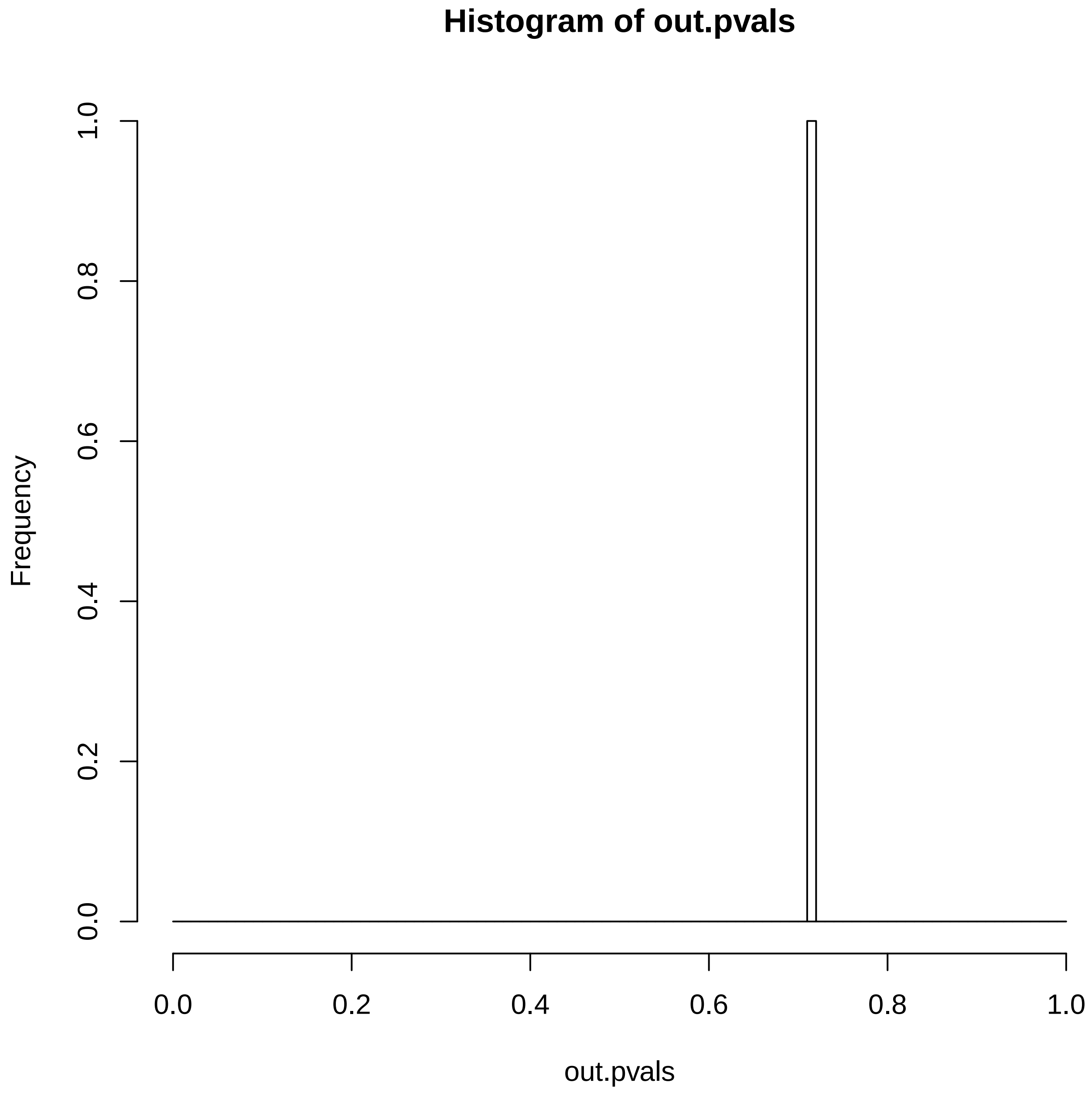

### SP17905.reconstruction.err.pdf

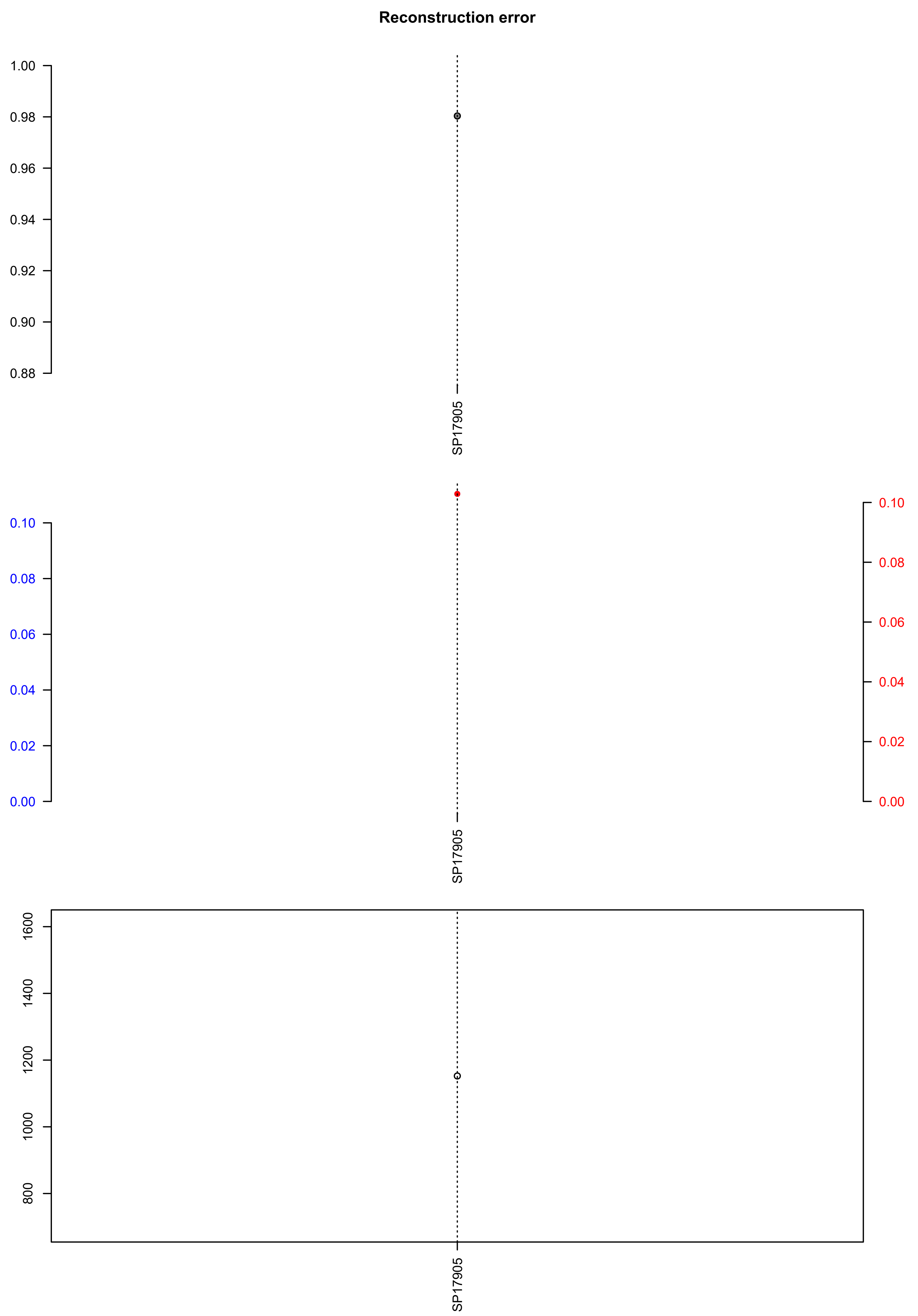

### SP18946.exposures.pdf

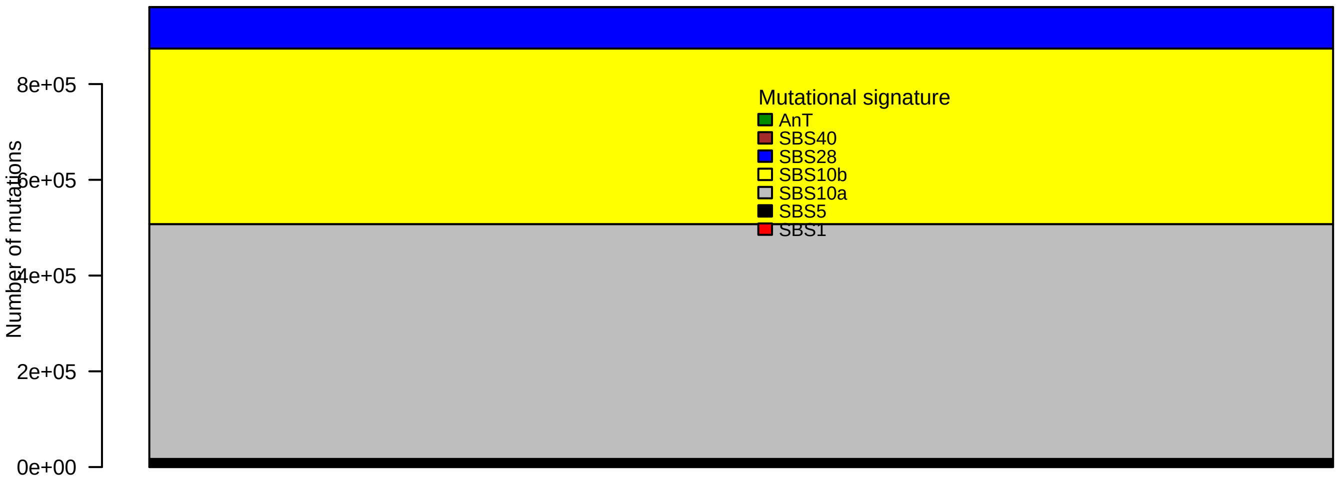

### SP18946.pval.histogram.pdf

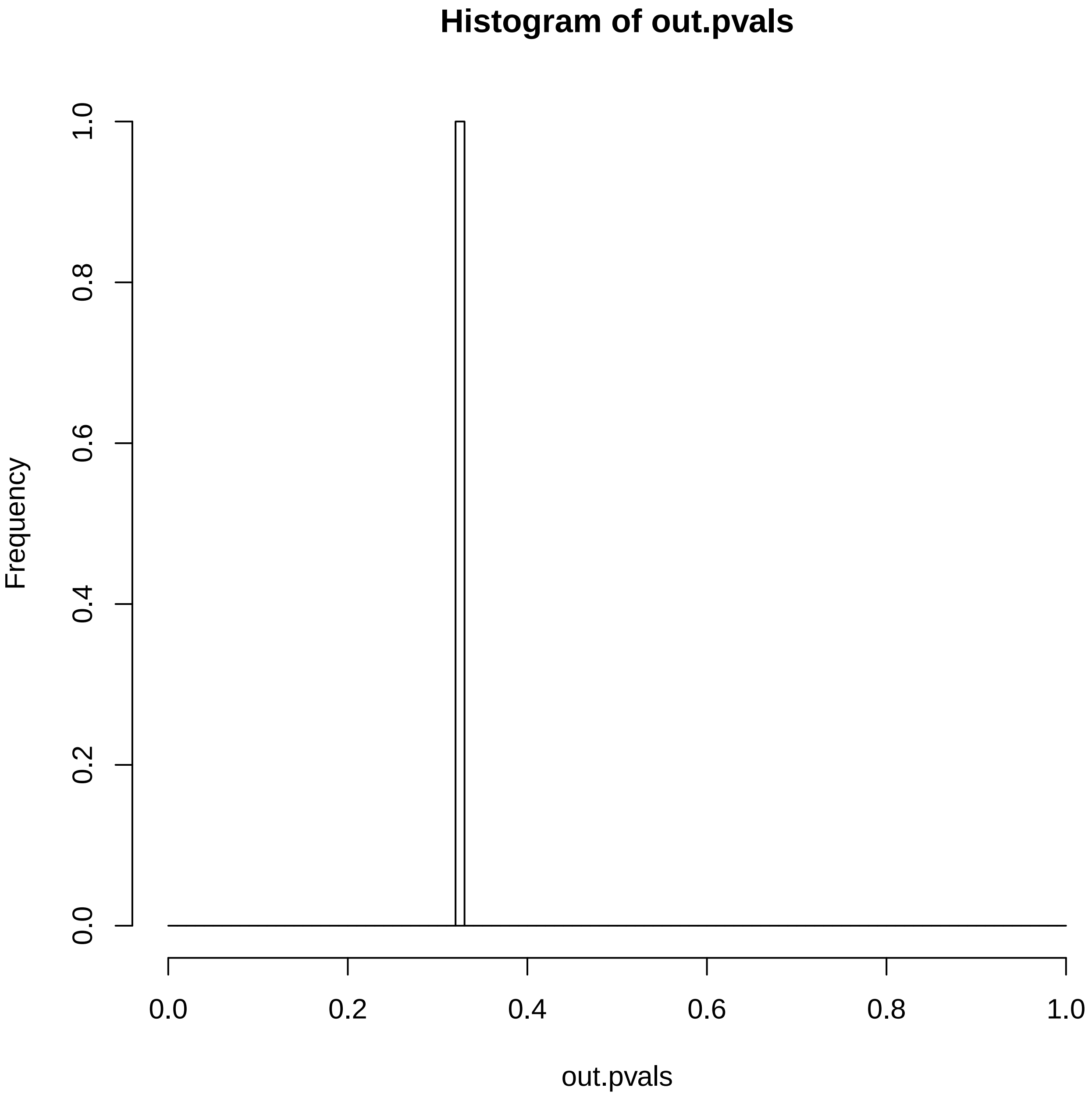

### SP18946.reconstruction.err.pdf

#### Reconstruction error

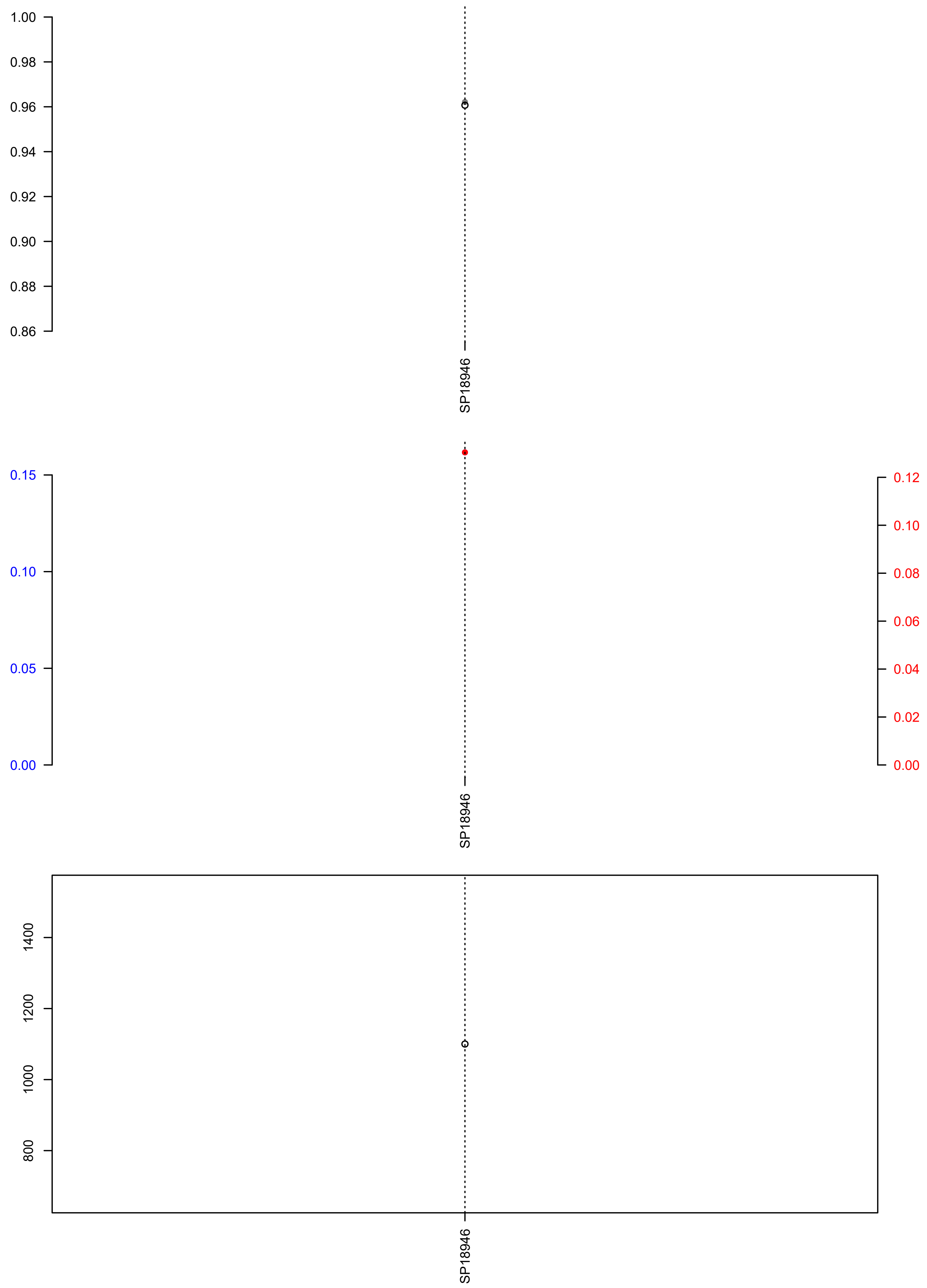

### SP19295.check.with.sig.pdf

# SP19295

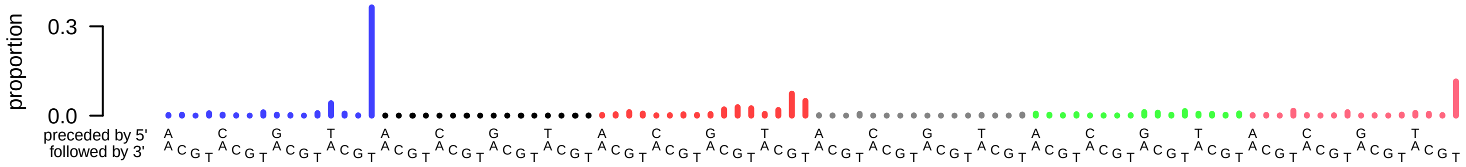

### SP19295.exposures.pdf

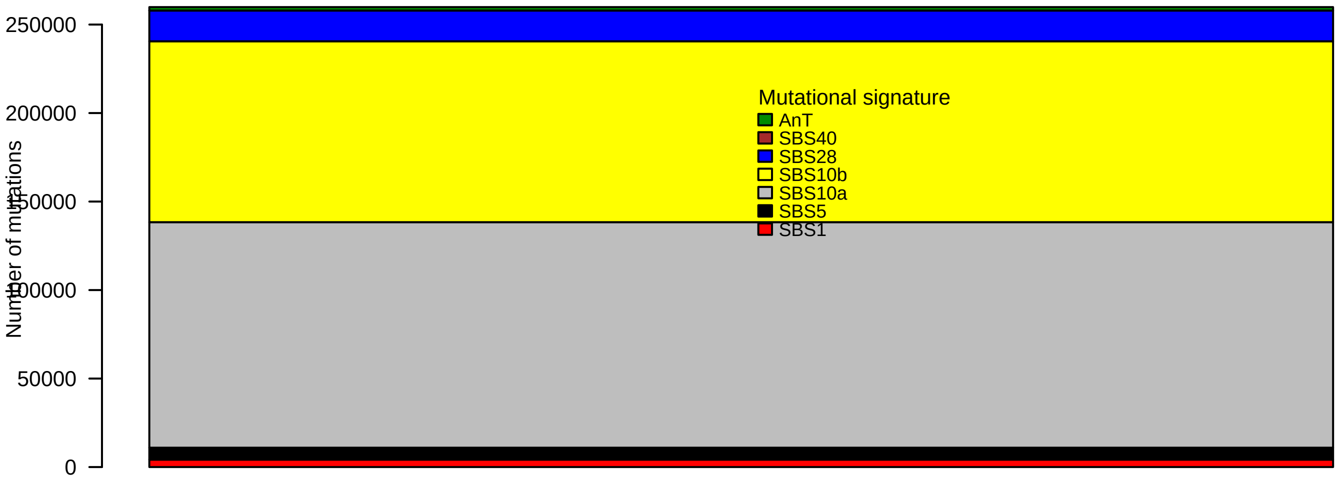

### SP19295.pval.histogram.pdf

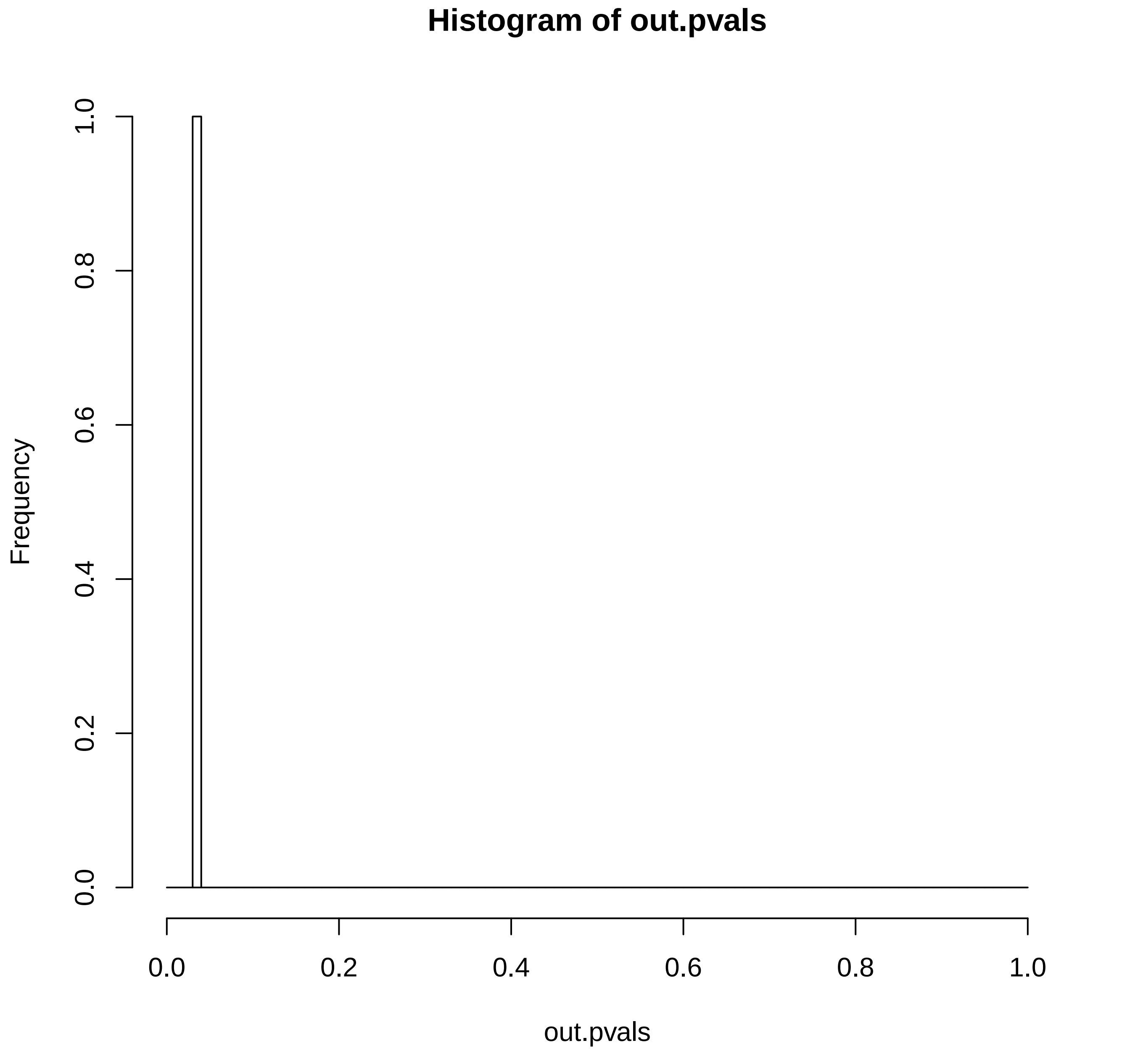

### SP19295.reconstruction.err.pdf

##### Reconstruction error

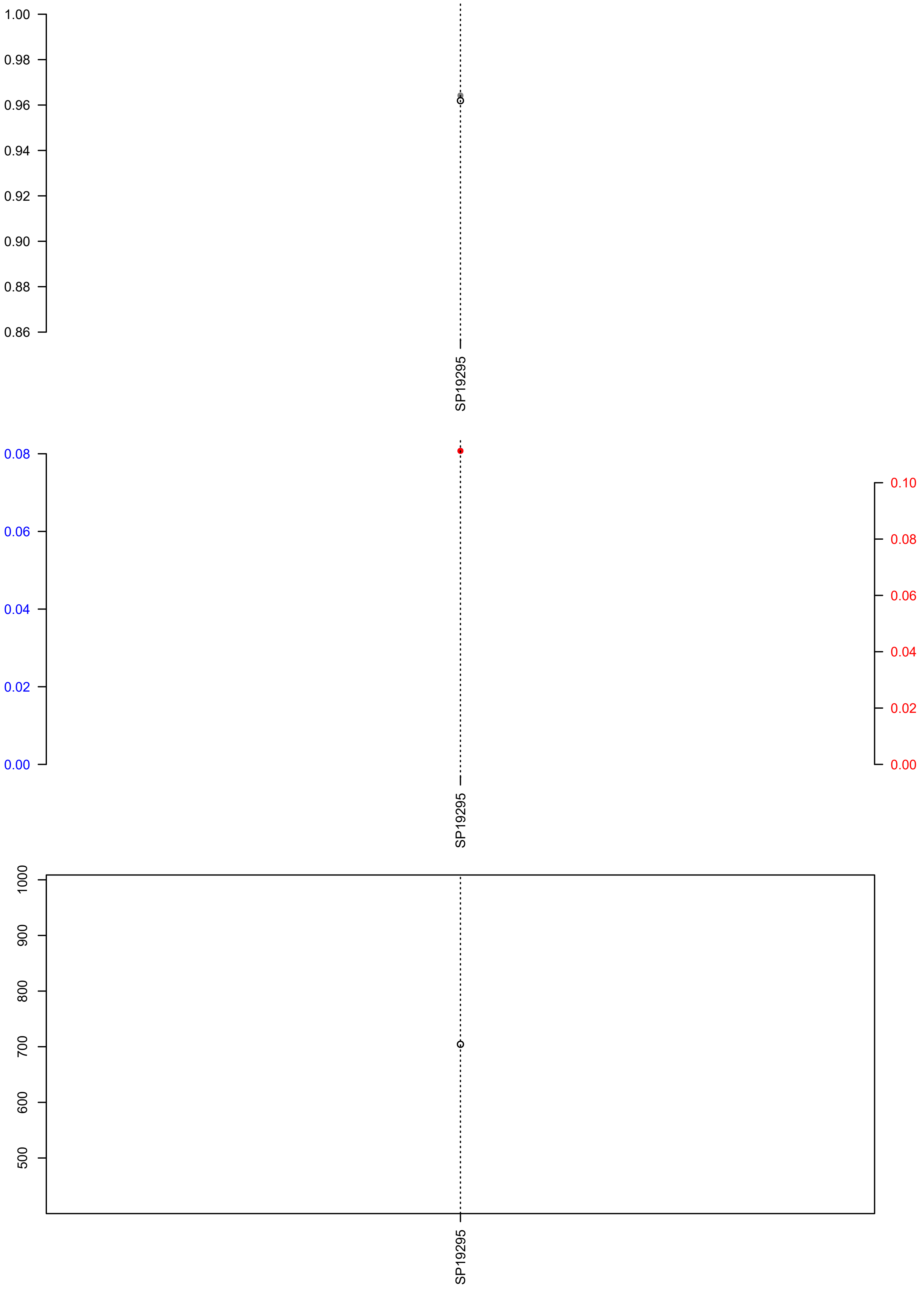

### SP21400.check.with.sig.pdf

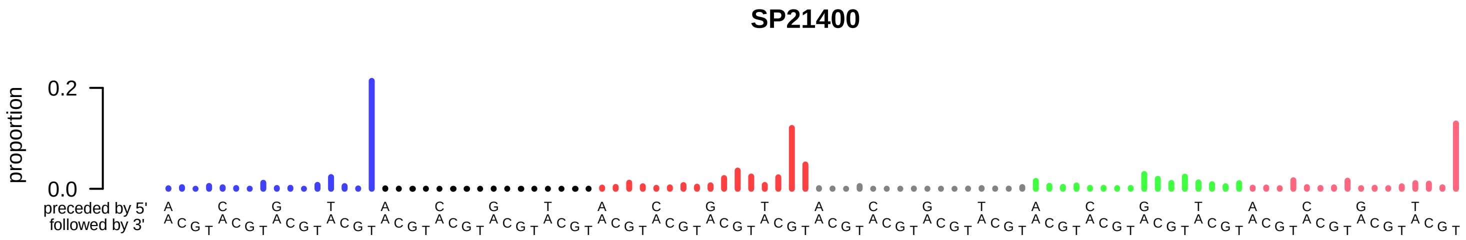

### SP21400.exposures.pdf

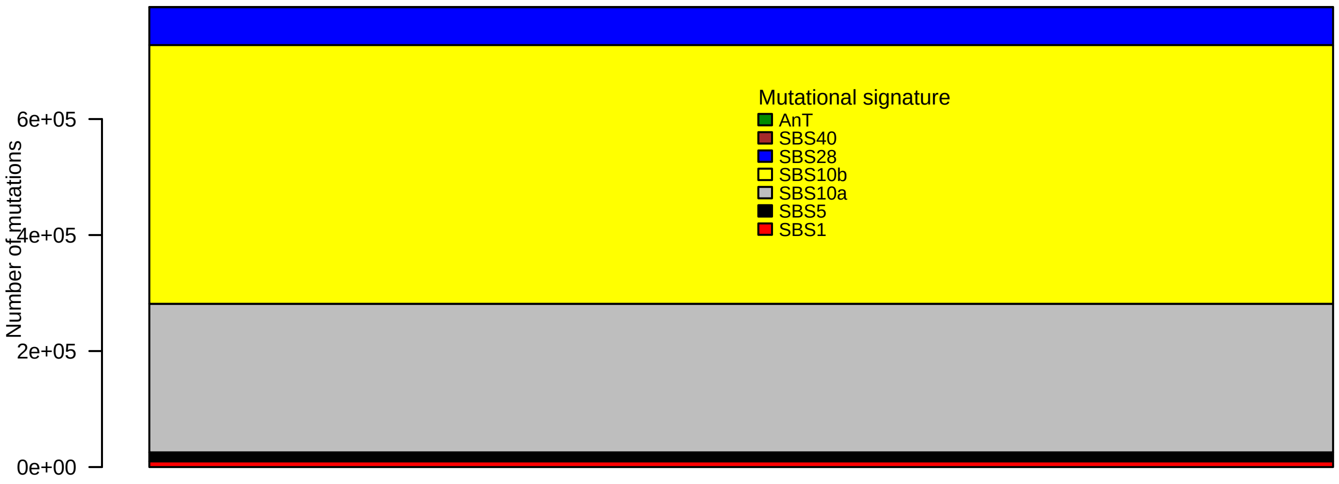

### SP21400.pval.histogram.pdf

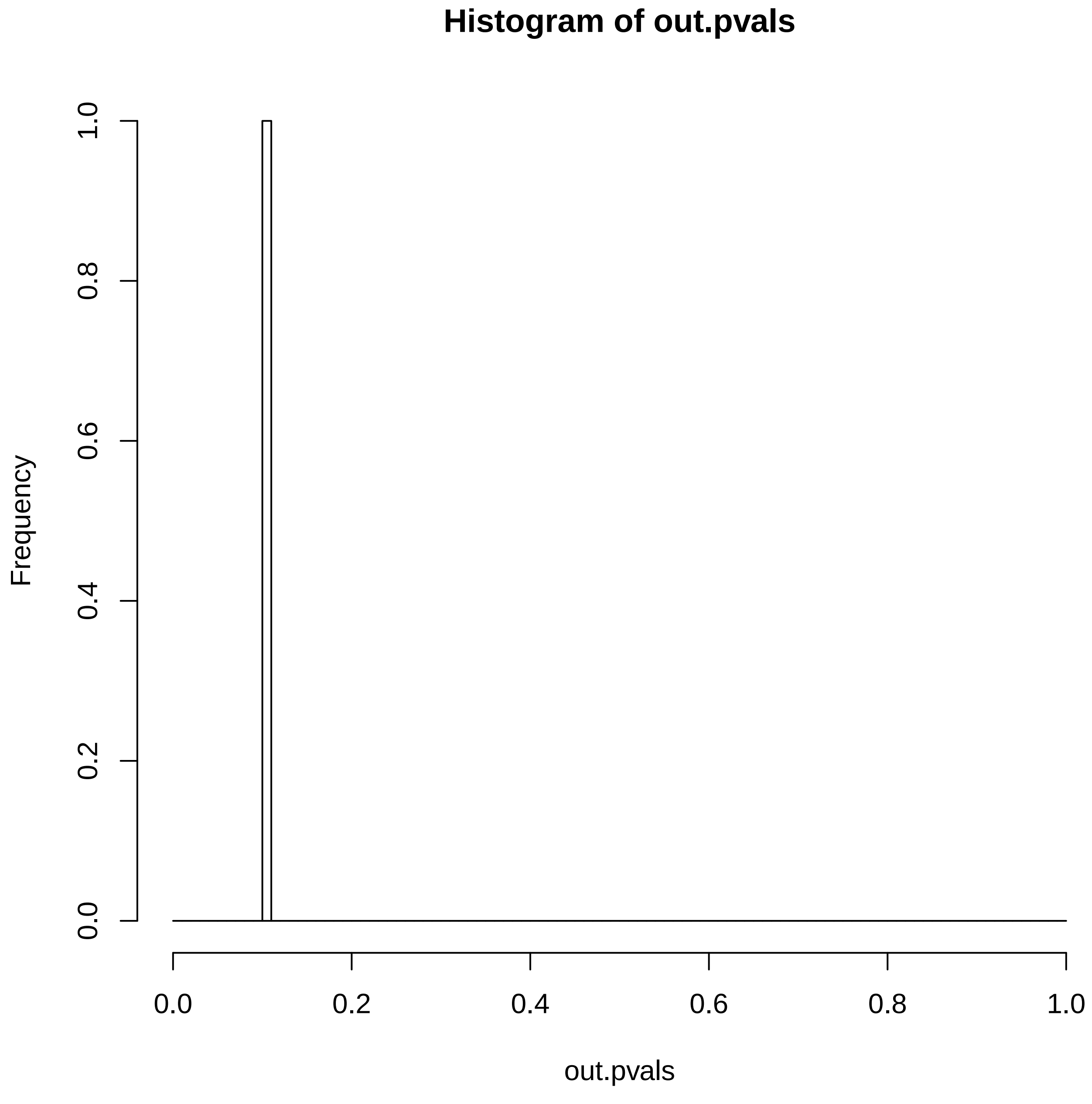

### SP21400.reconstruction.err.pdf

##### Reconstruction error

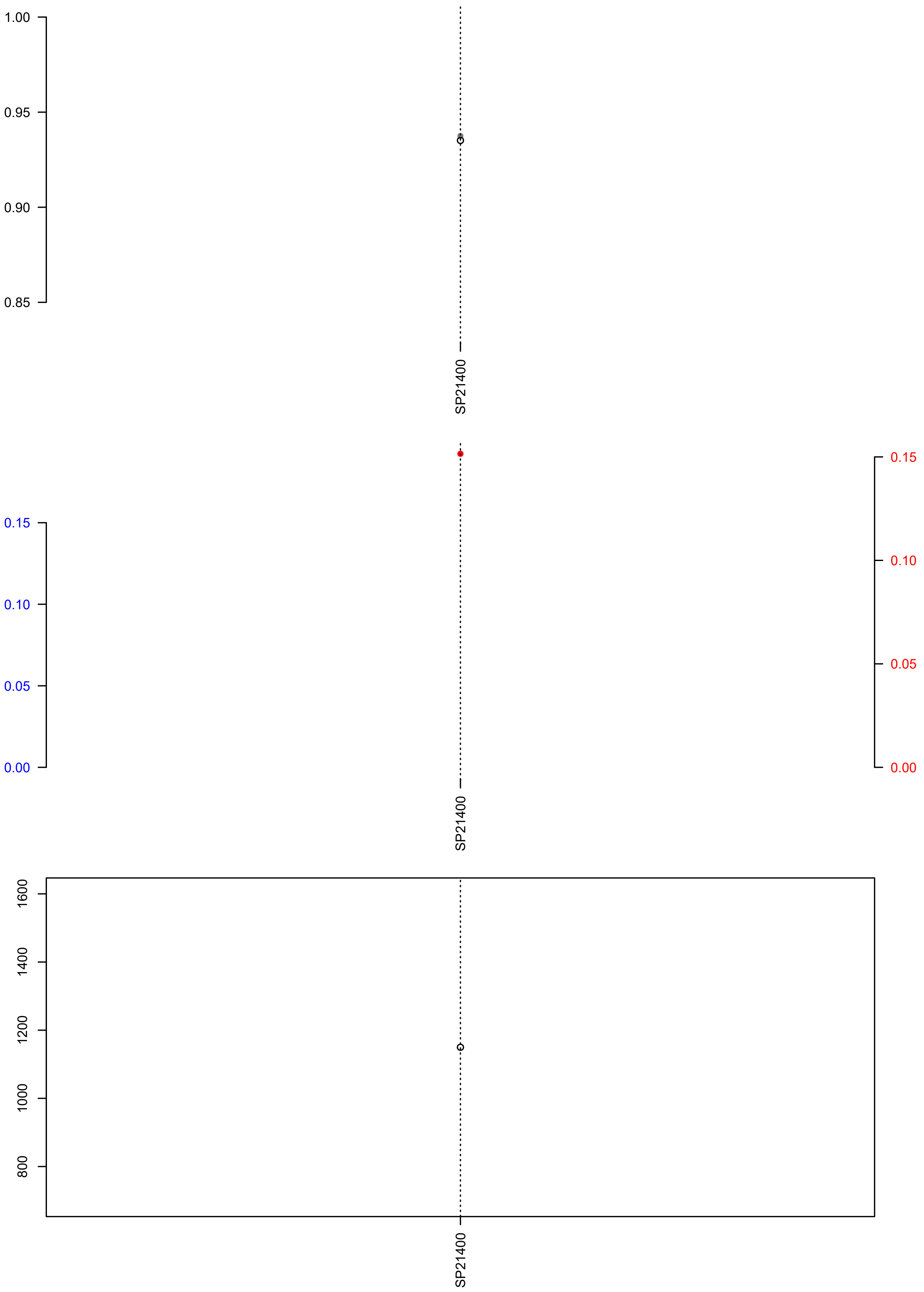

### SP22031.exposures.pdf

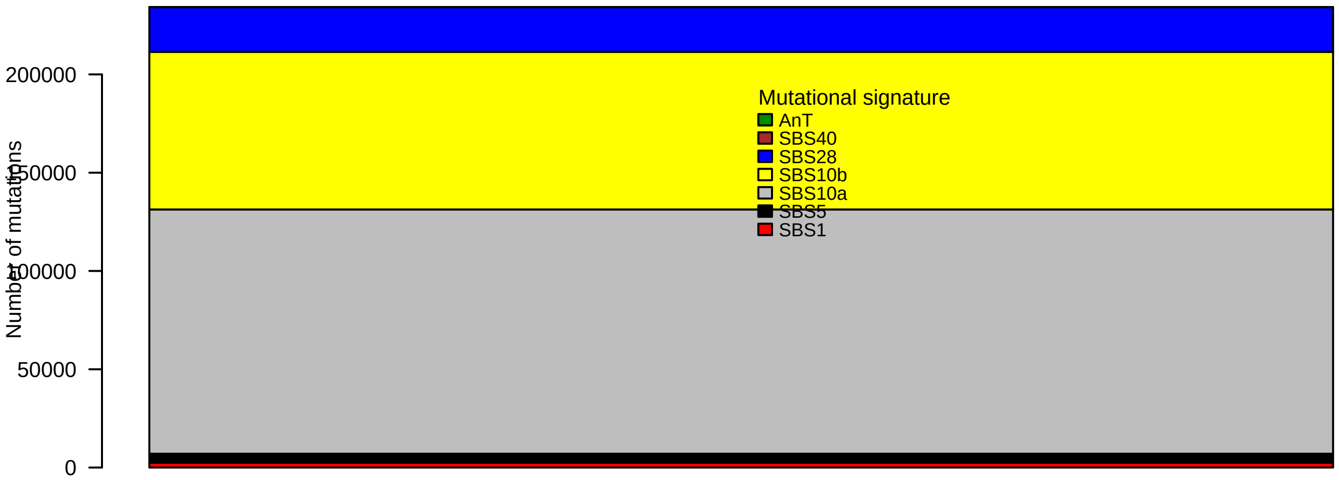

### SP22031.pval.histogram.pdf

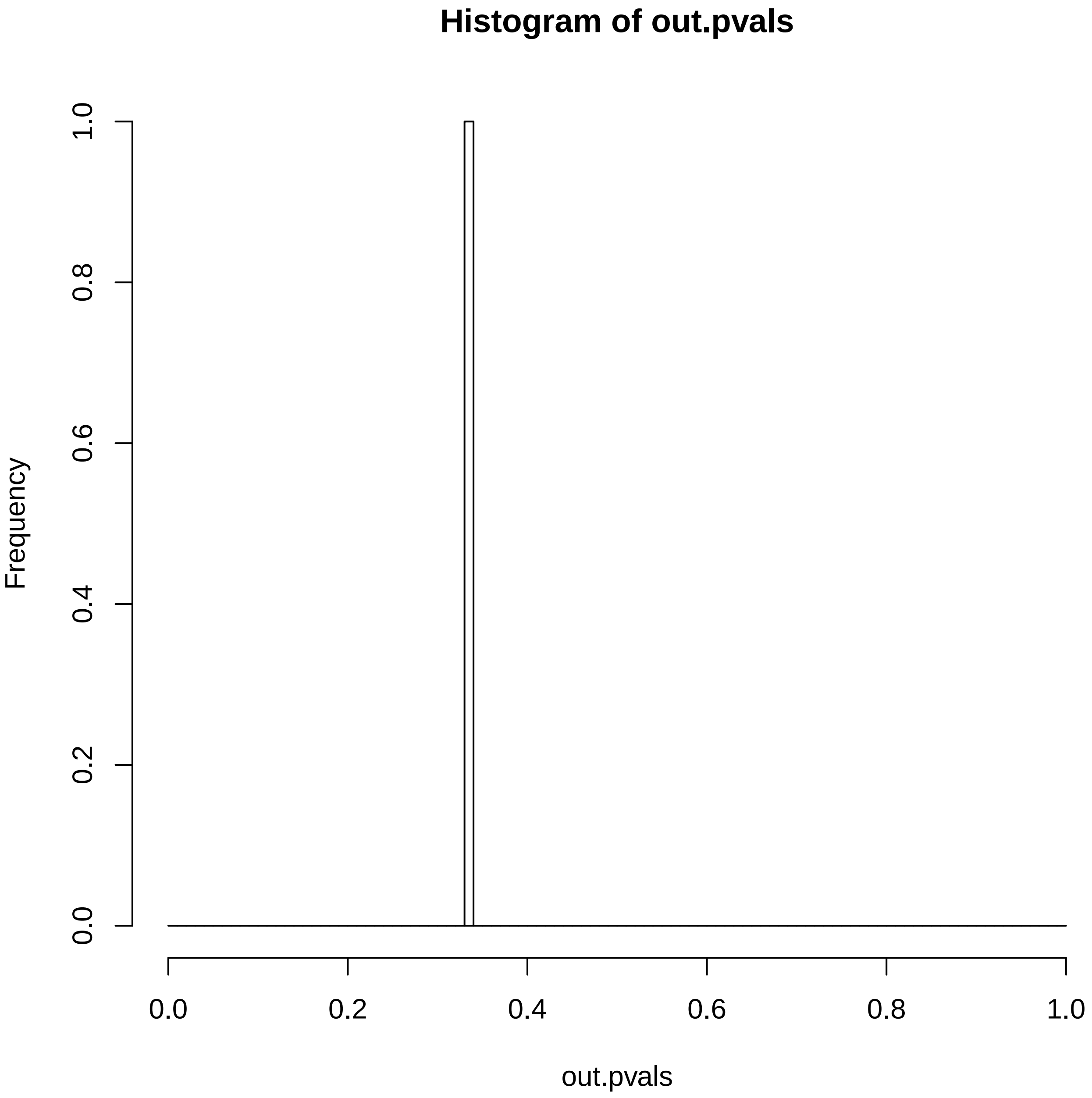

### SP22031.reconstruction.err.pdf

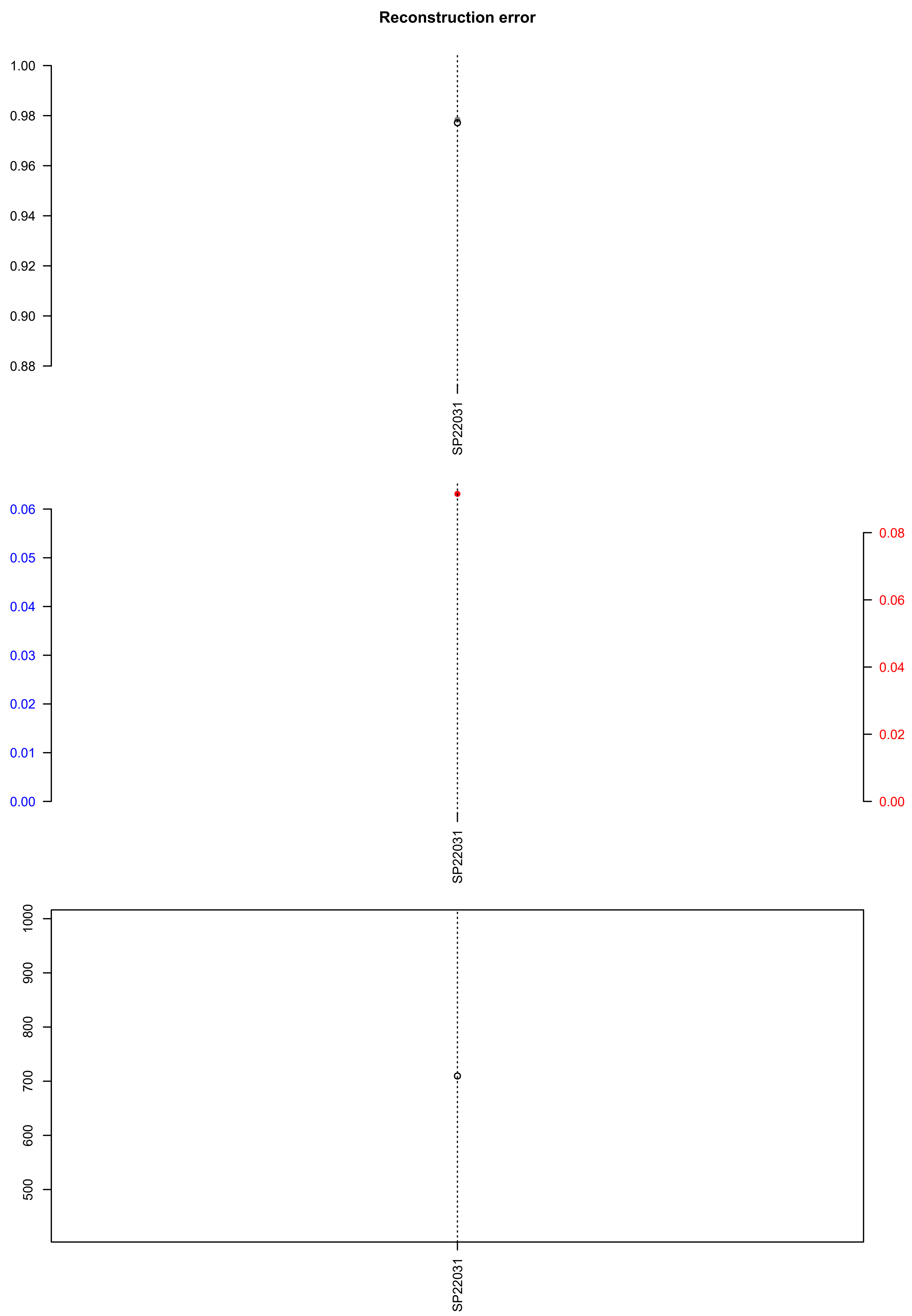

### SP80615.check.with.sig.pdf

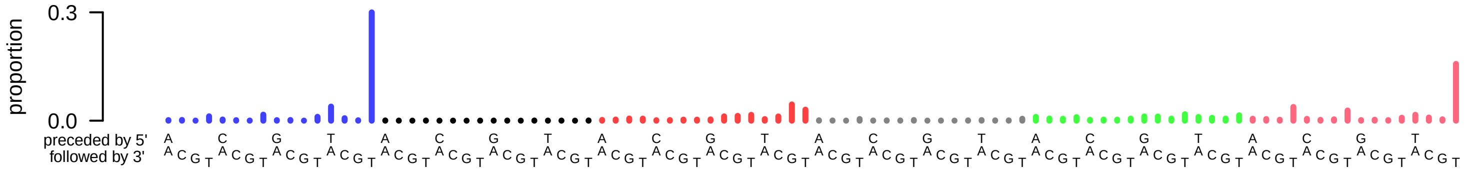

### SP80615.exposures.pdf

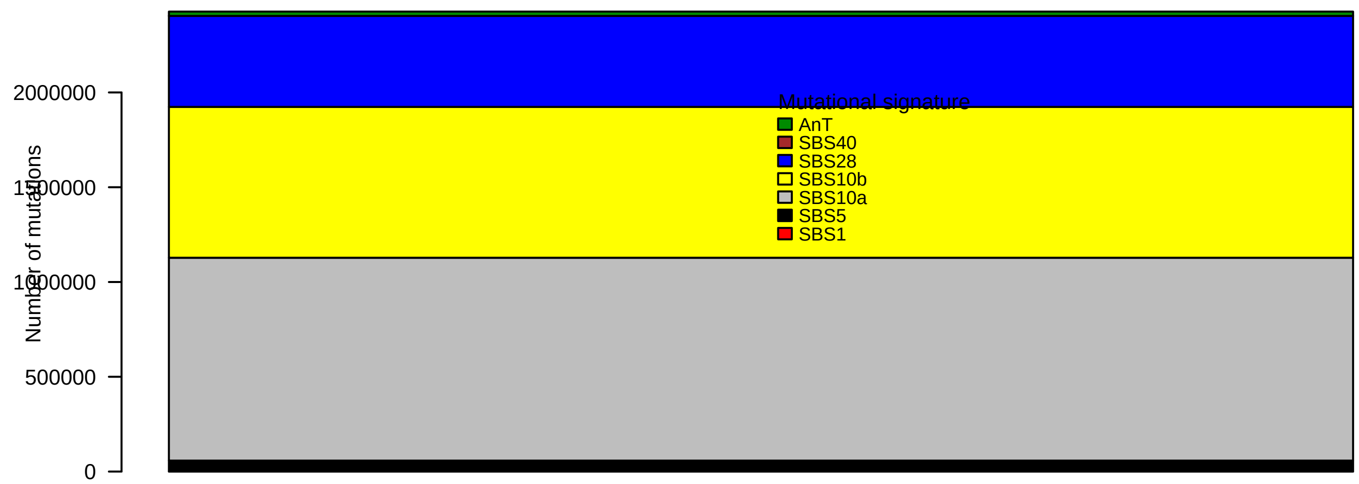

### SP80615.pval.histogram.pdf

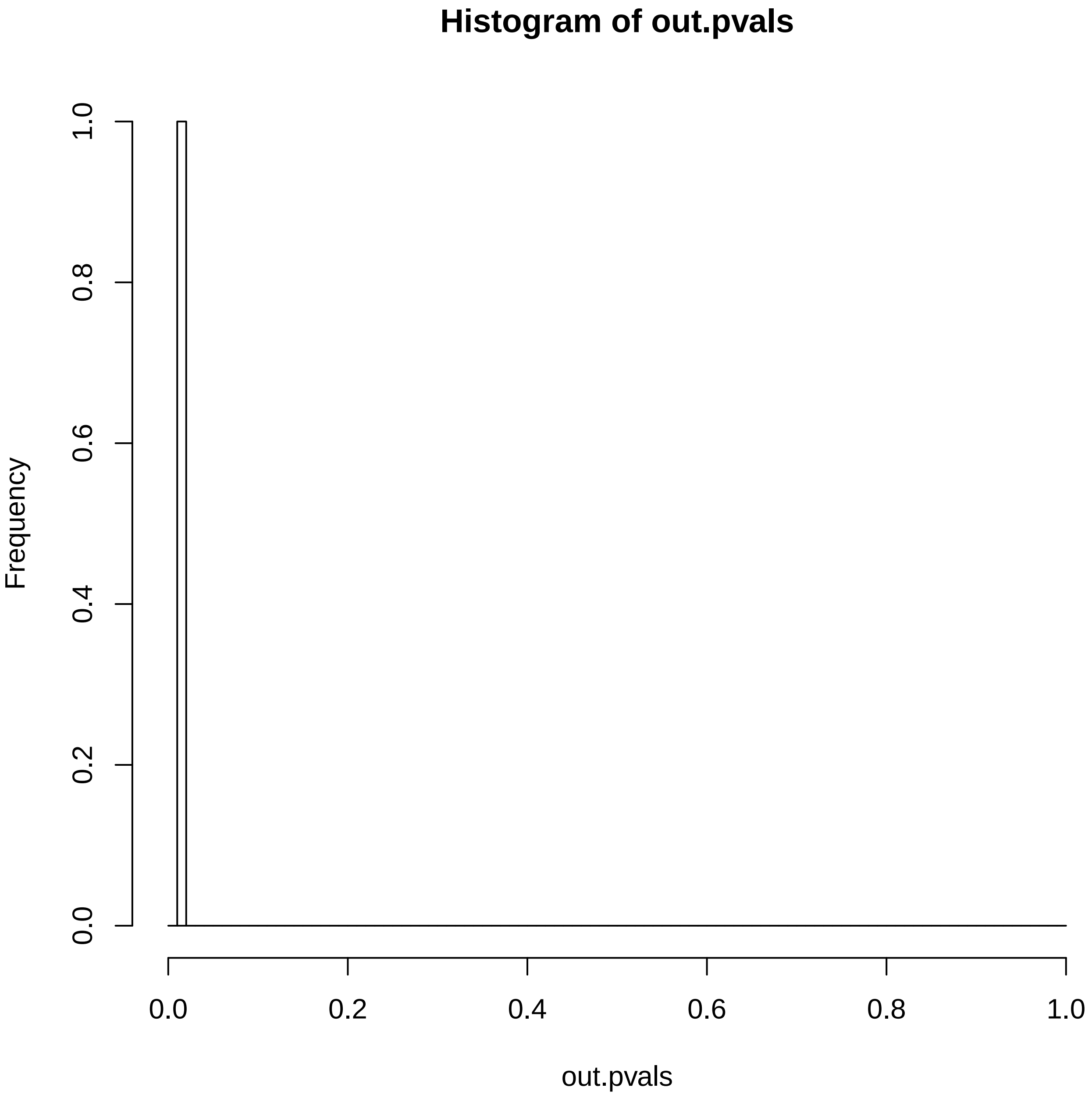

### SP80615.reconstruction.err.pdf

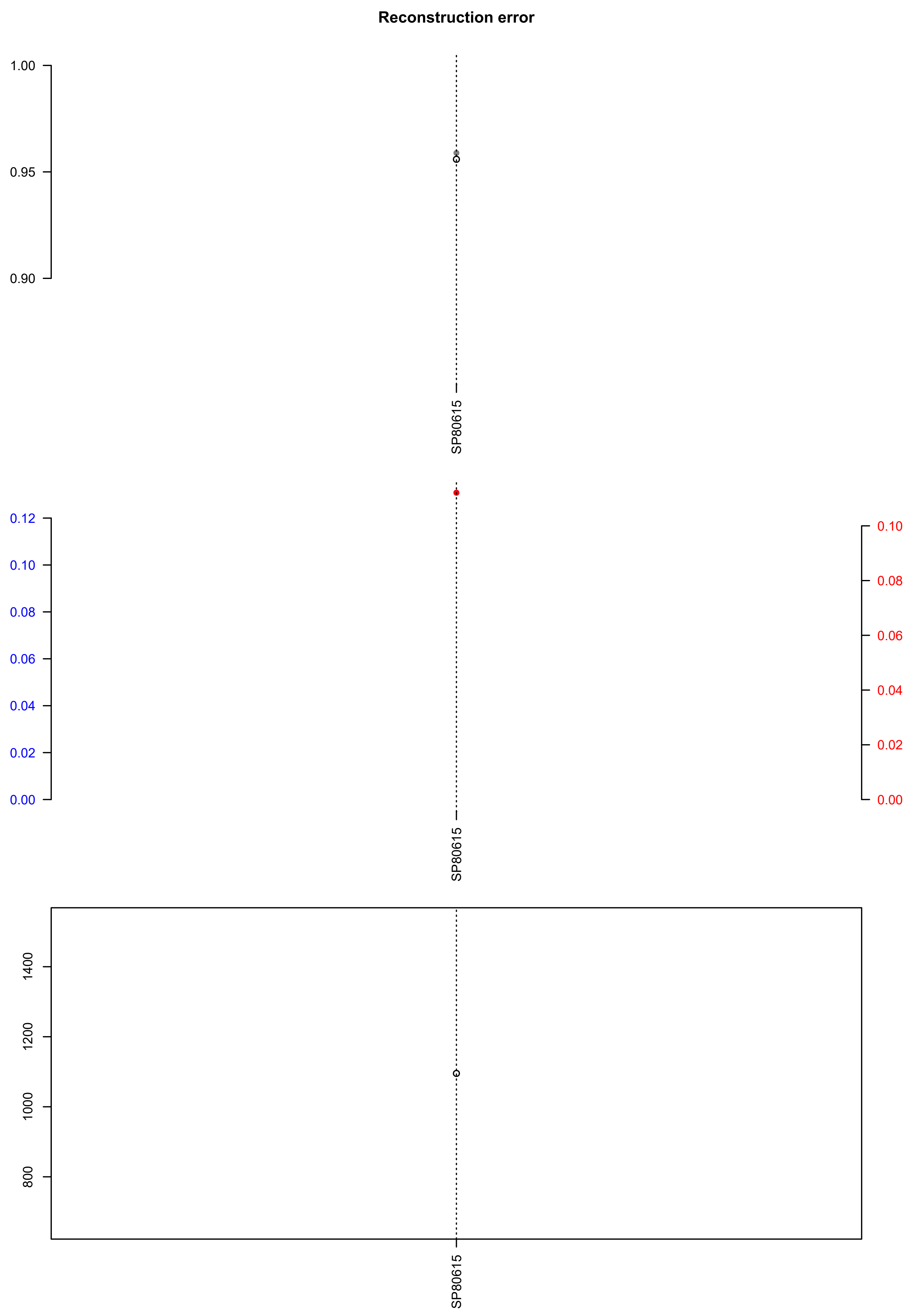

### SP80754.check.with.sig.pdf

# SP80754

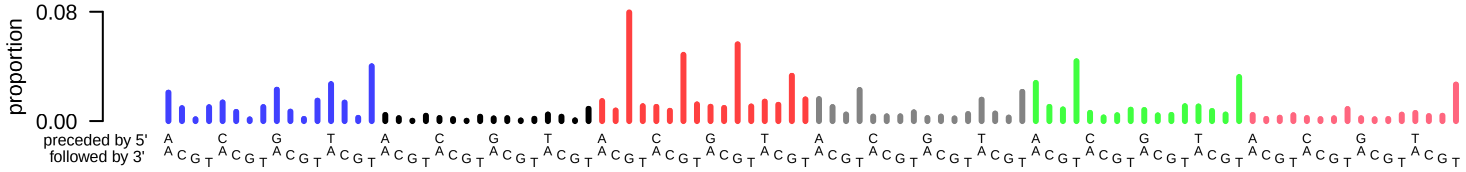

### SP80754.exposures.pdf

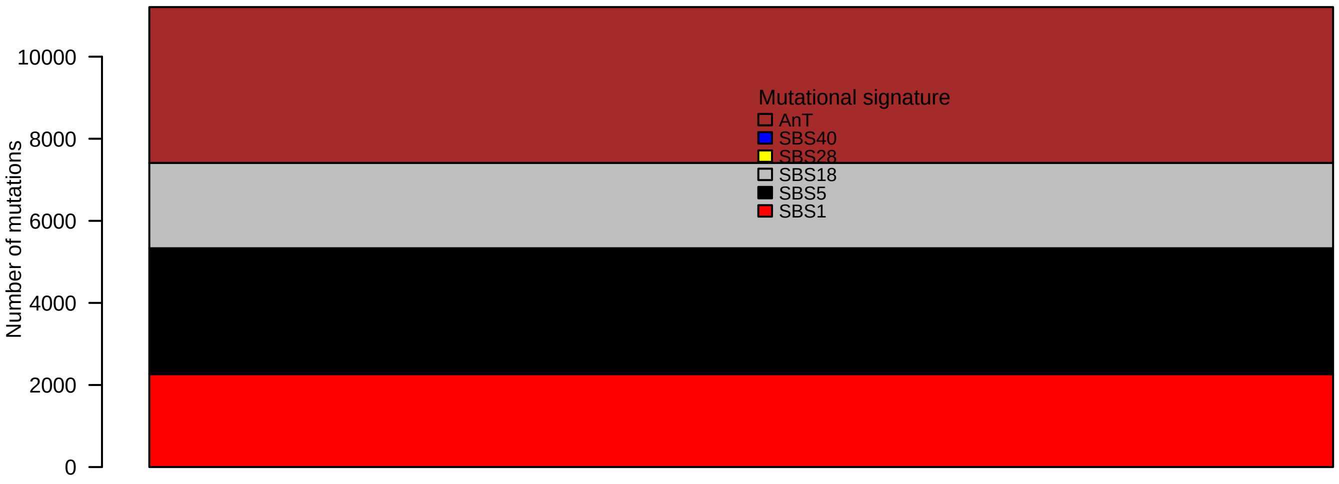

### SP80754.pval.histogram.pdf

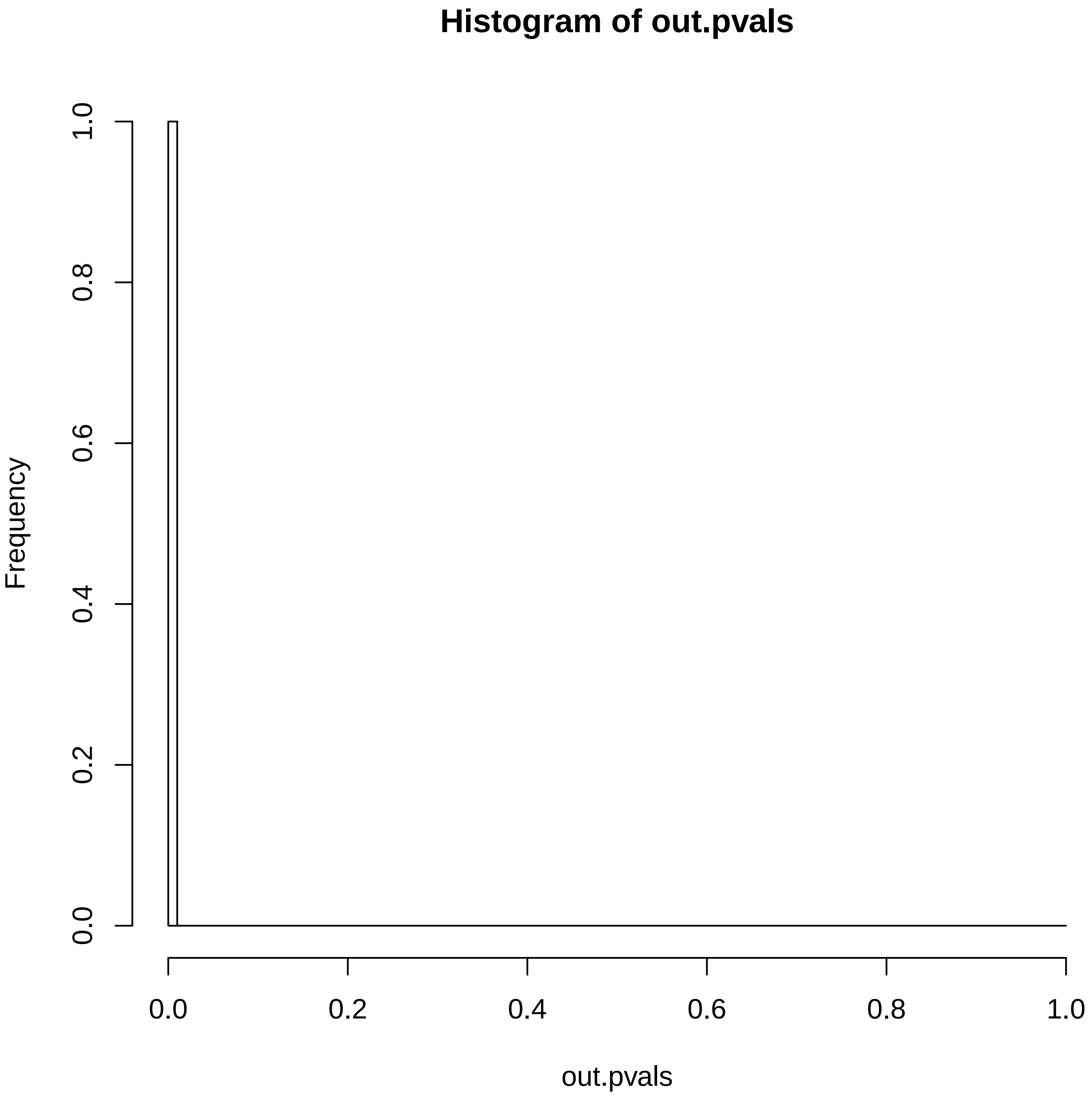

### SP80754.reconstruction.err.pdf

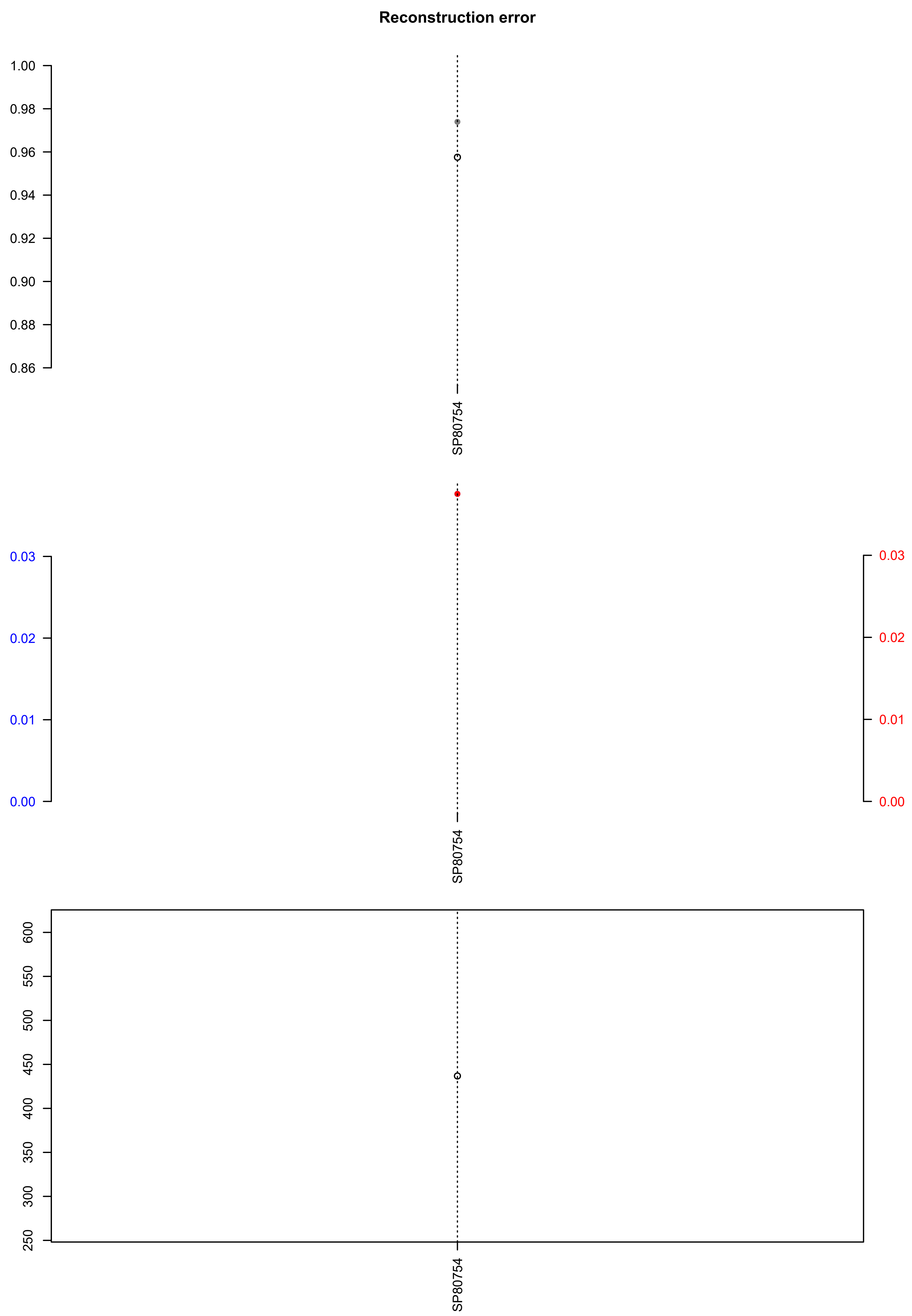

### SP81494.check.with.sig.pdf

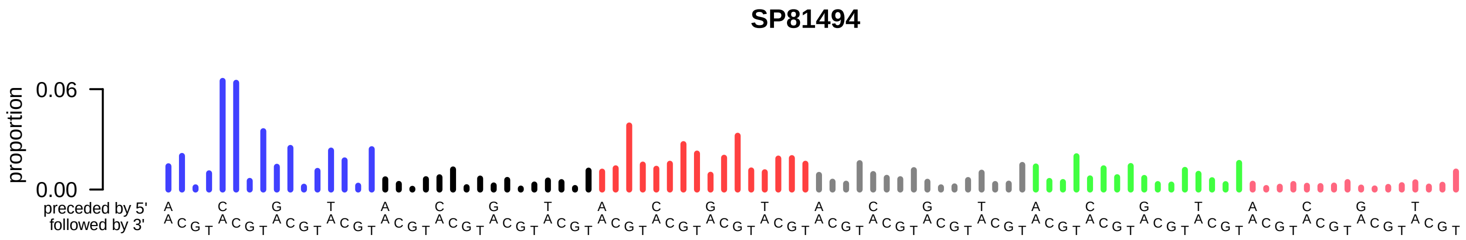

### SP81494.exposures.pdf

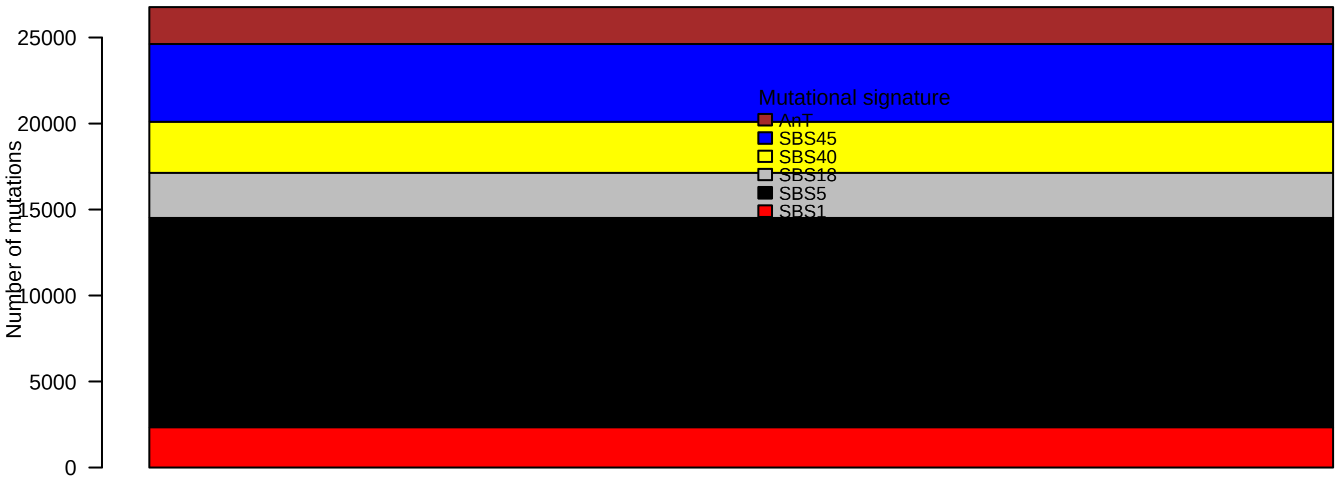

### SP81494.pval.histogram.pdf

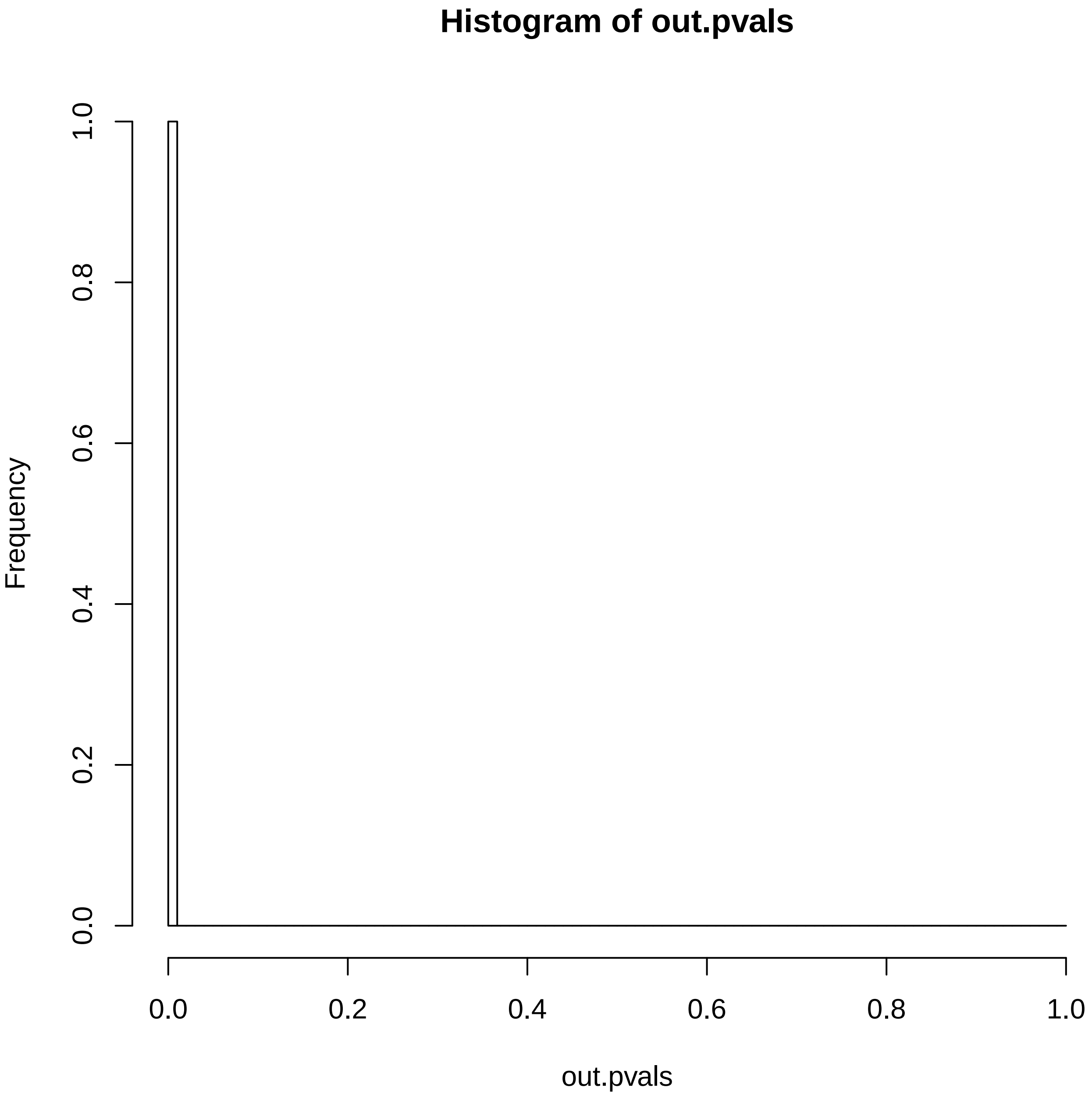

### SP81494.reconstruction.err.pdf

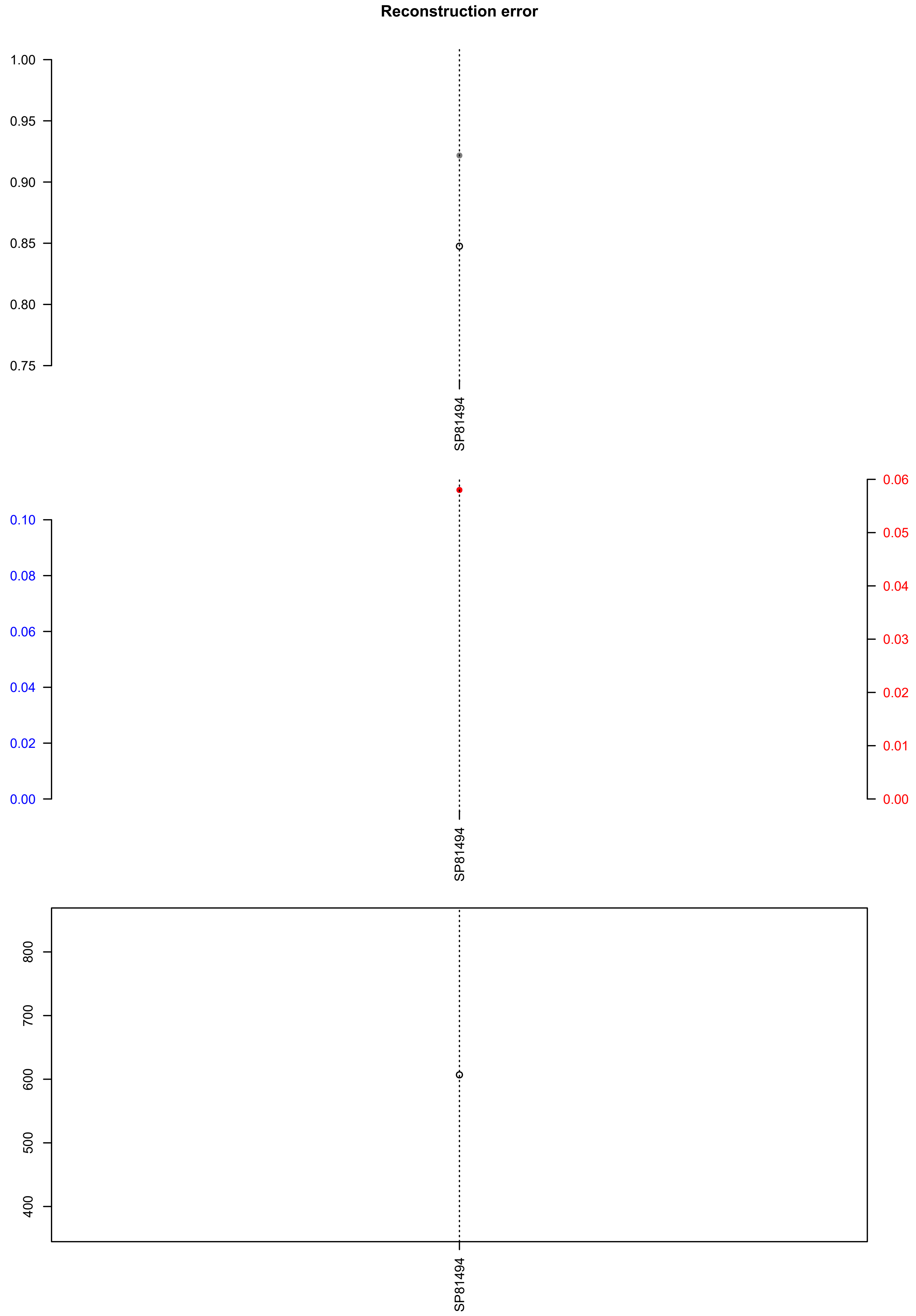

### SP81711.check.with.sig.pdf

# SP81711

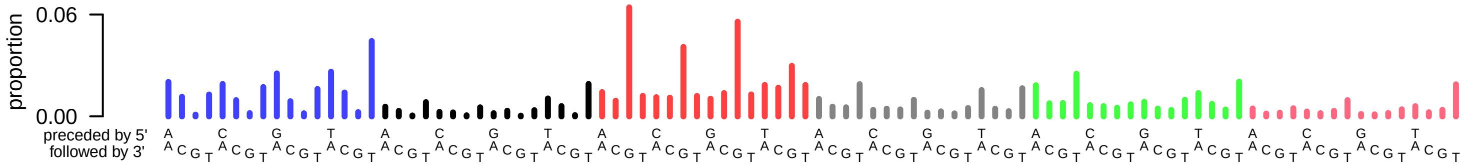

### SP81711.exposures.pdf

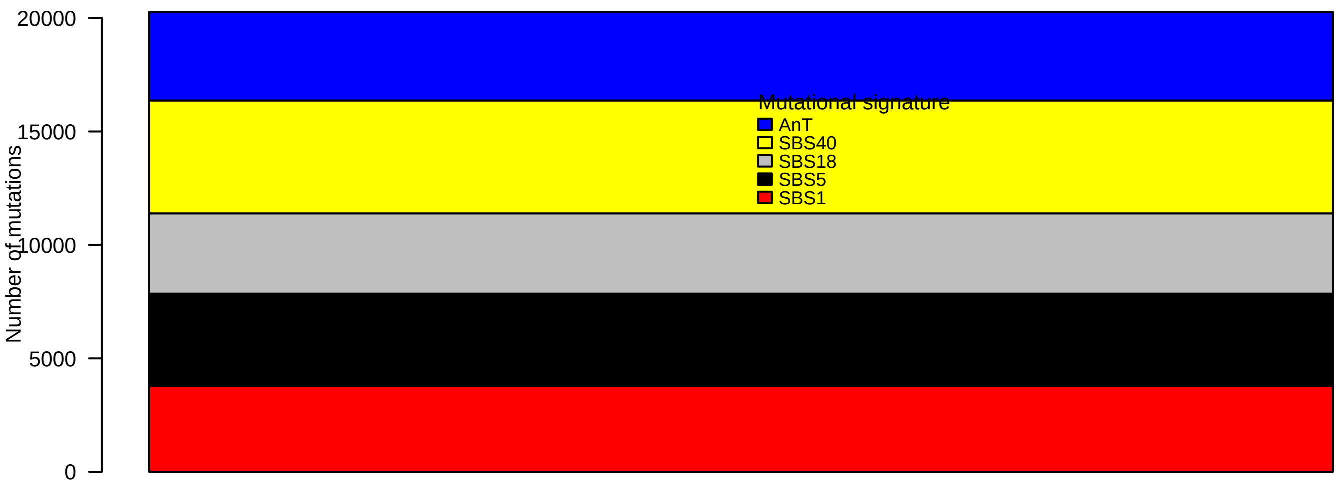

### SP81711.pval.histogram.pdf

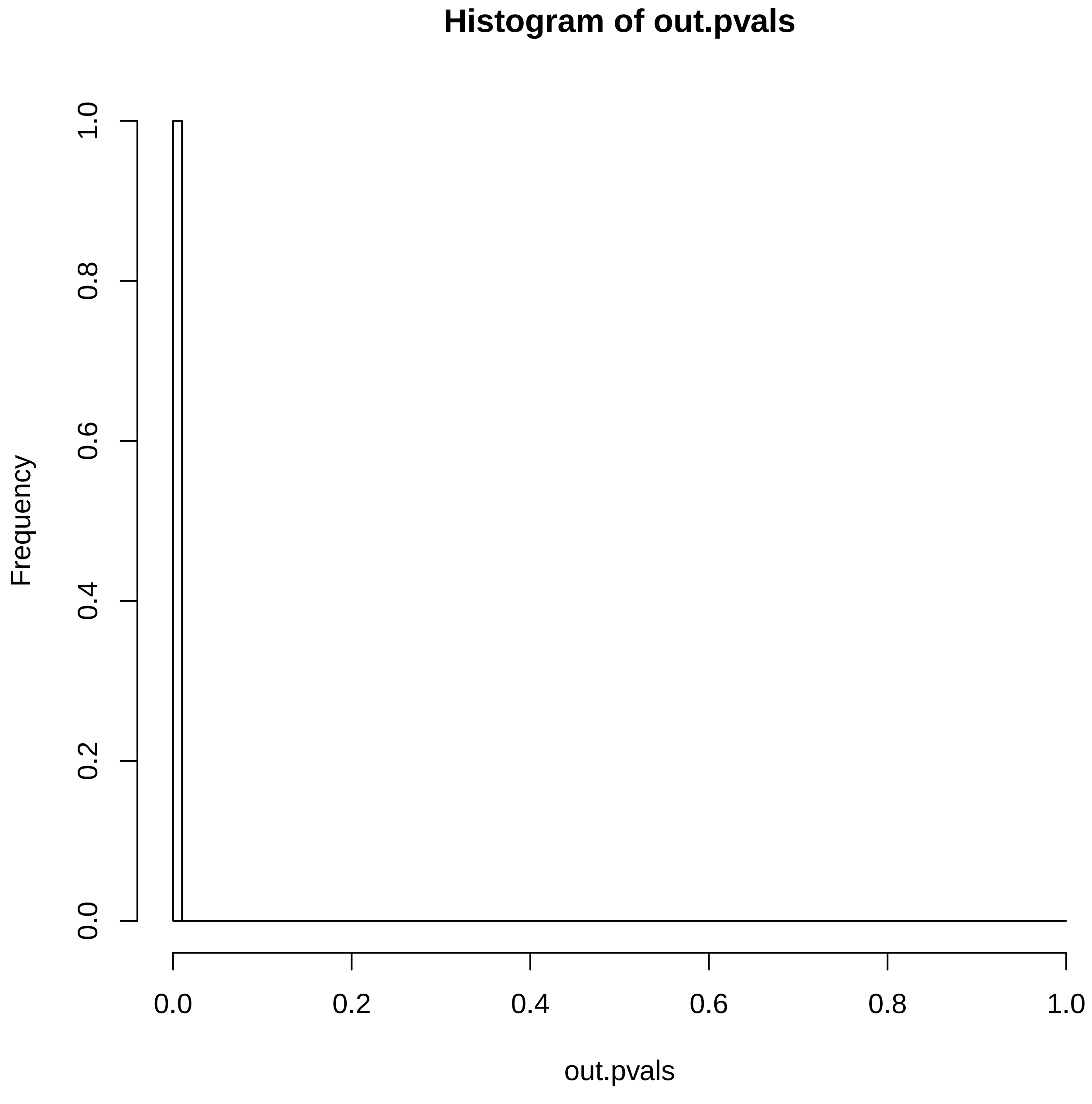

### SP81711.reconstruction.err.pdf

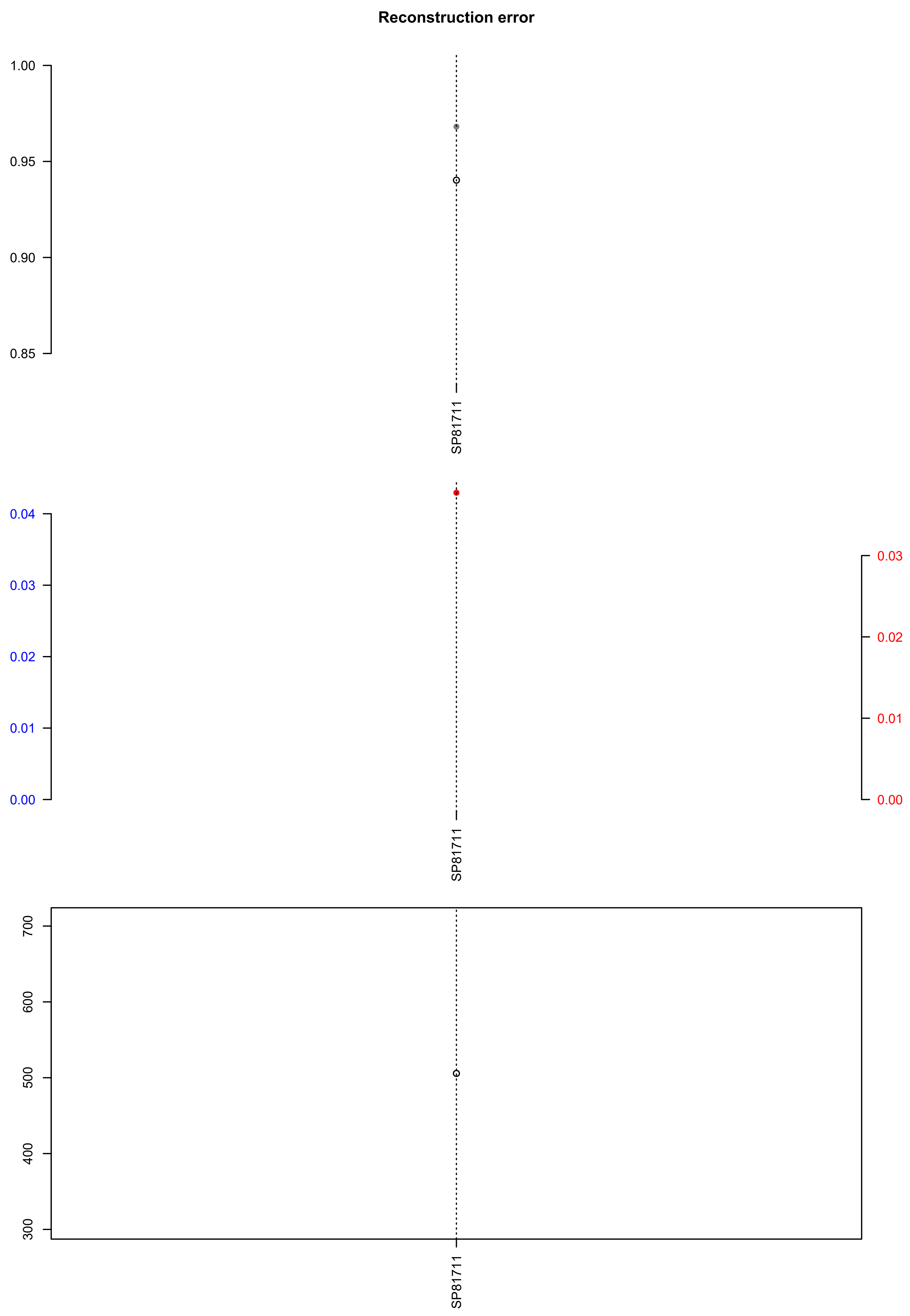

### SP92659.exposures.pdf

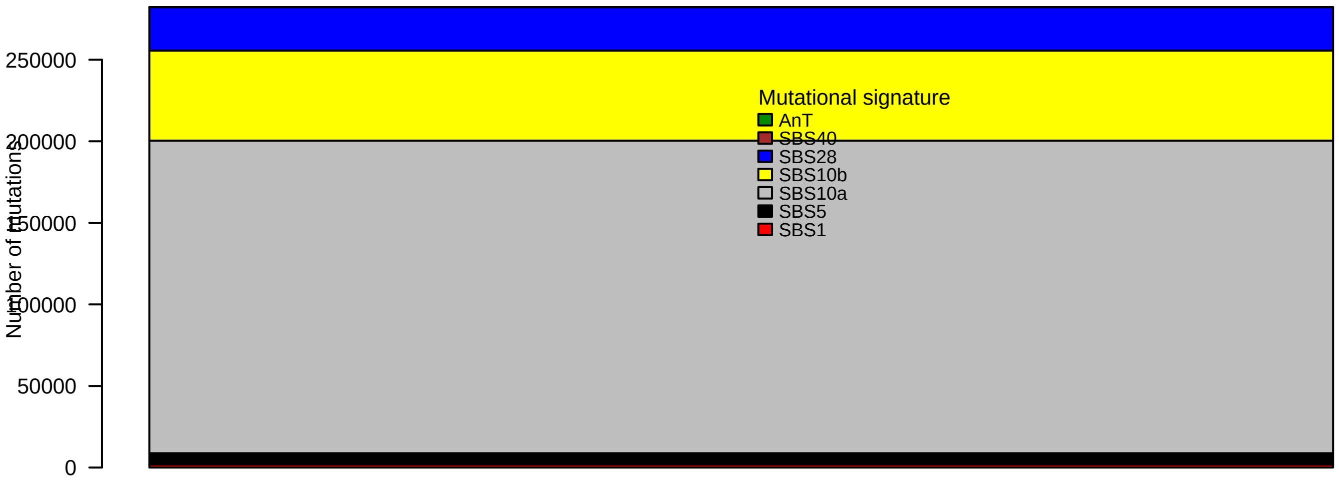

### SP92659.pval.histogram.pdf

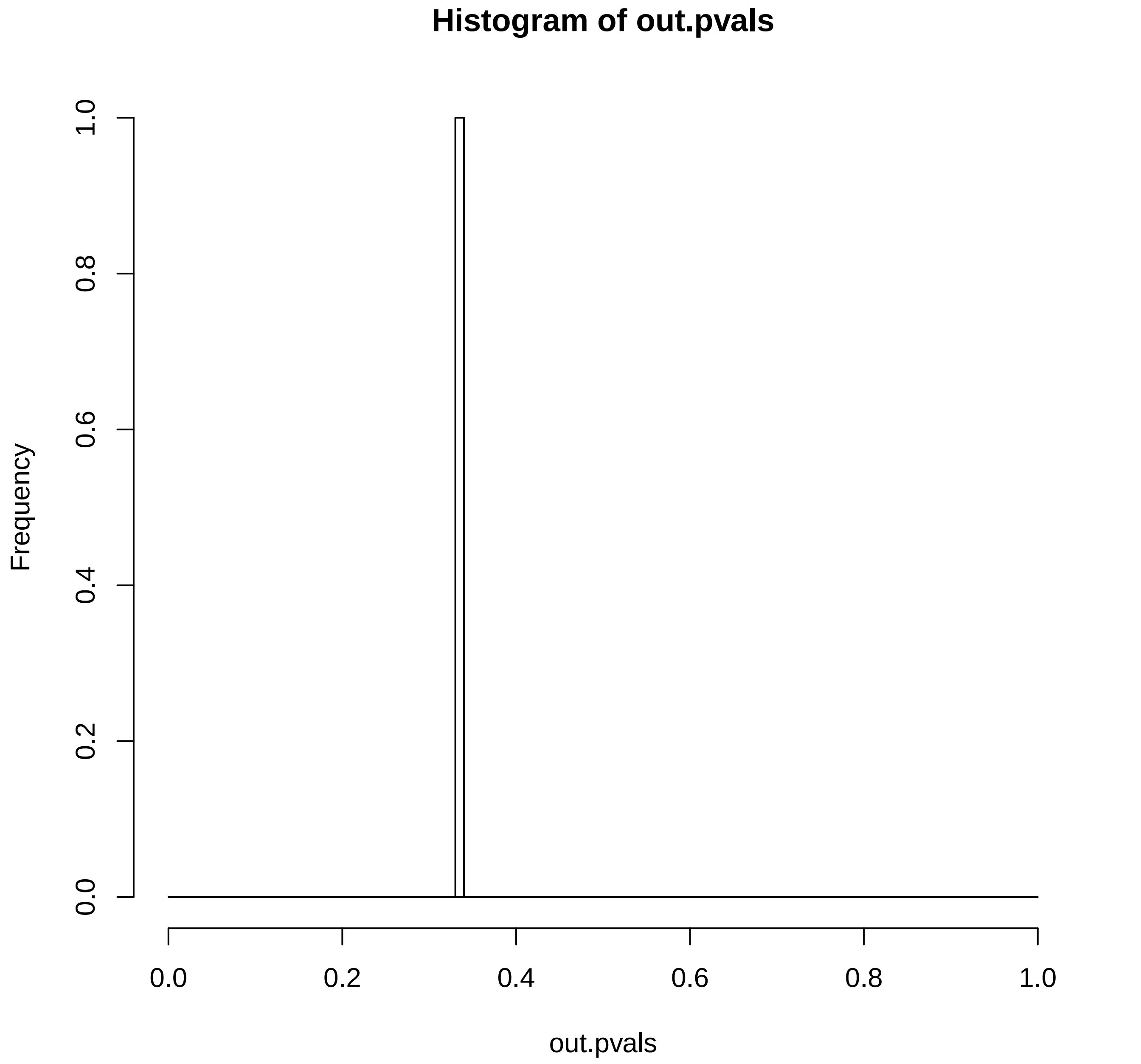

### SP92659.reconstruction.err.pdf

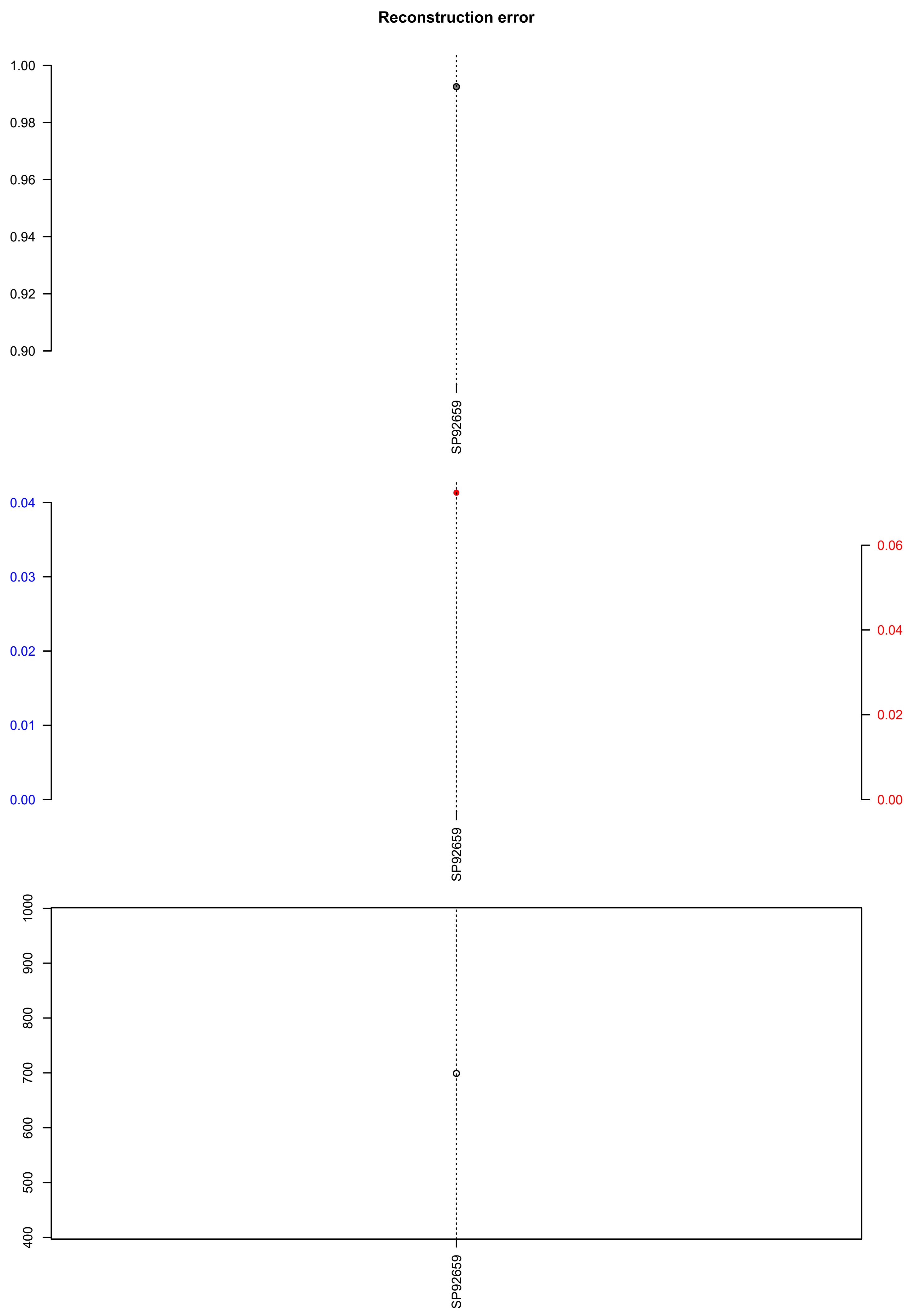

### SP111026.check.with.sig.pdf

# SP111026

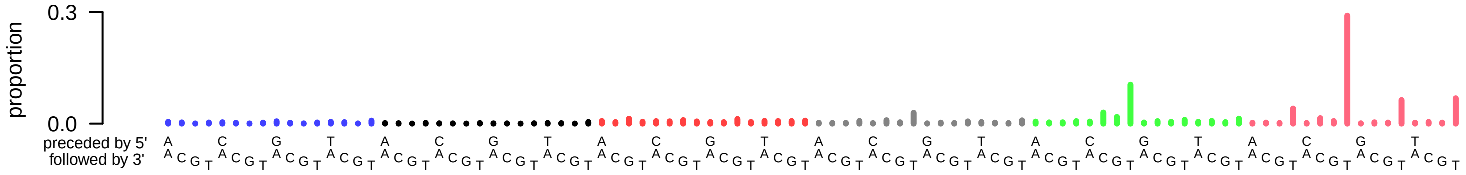

### SP111026.exposures.pdf

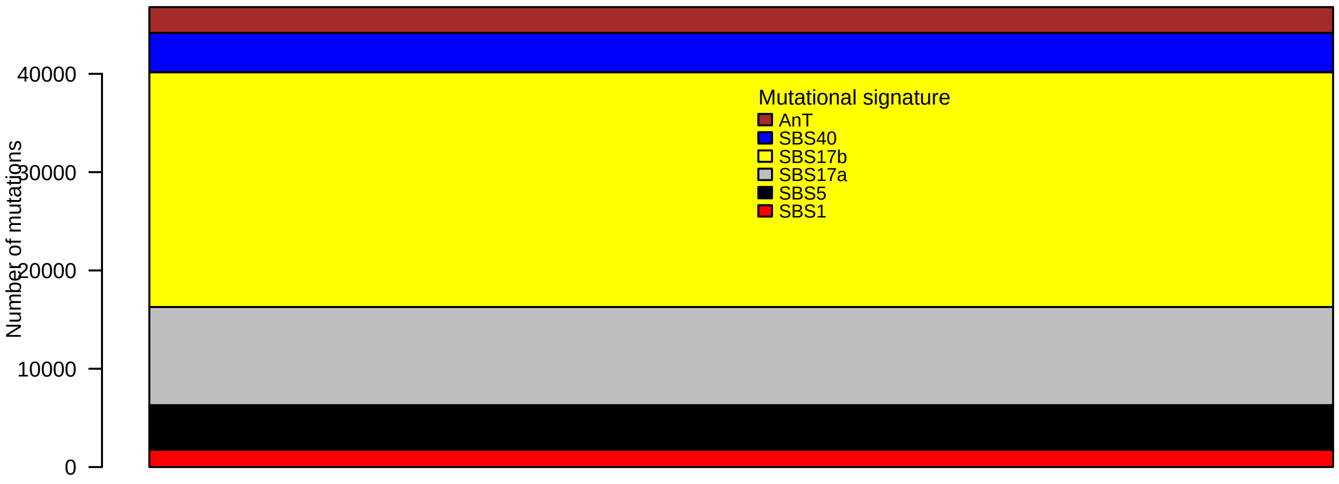

### SP111026.pval.histogram.pdf

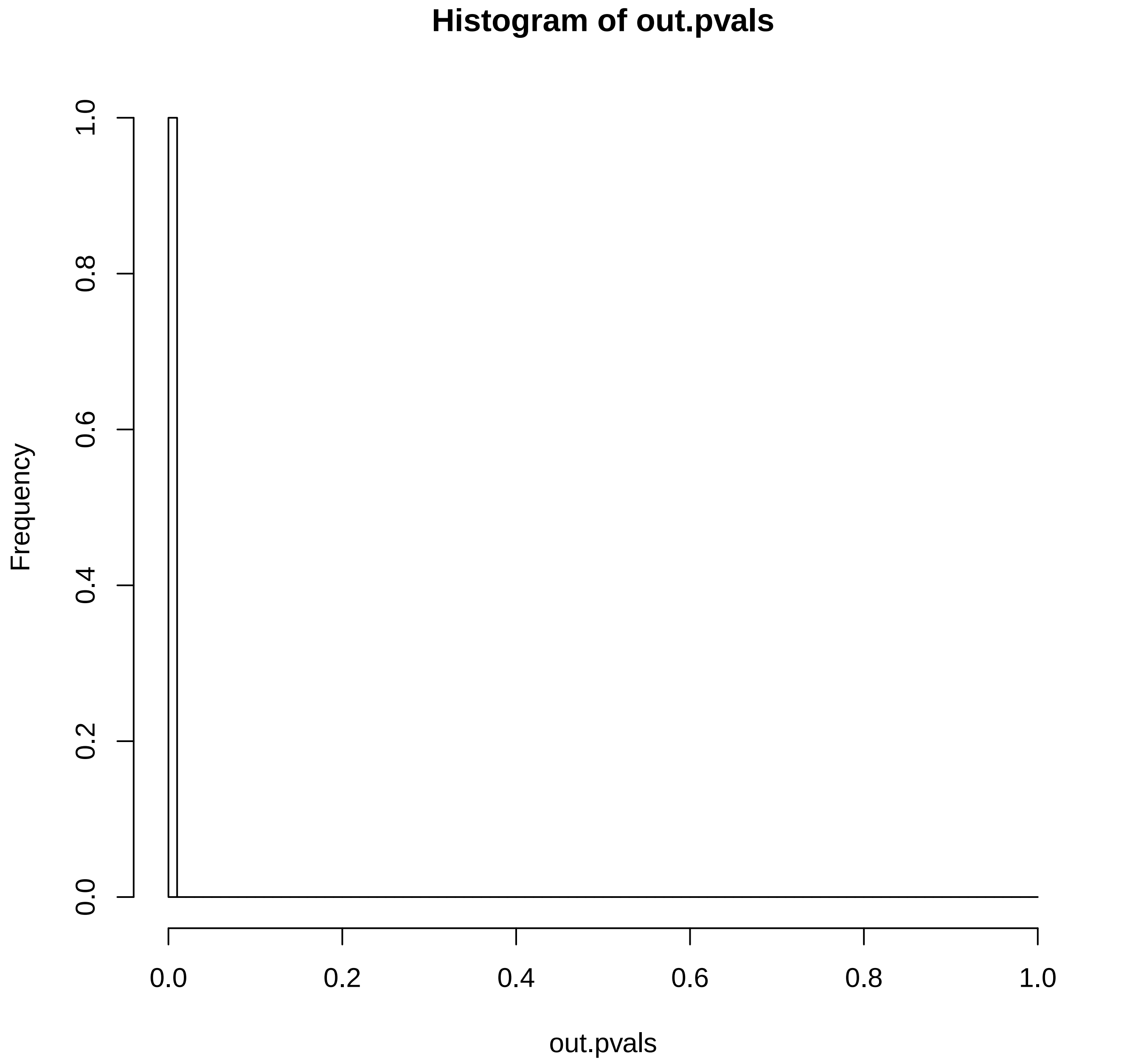

### SP111026.reconstruction.err.pdf

**Reconstruction error**

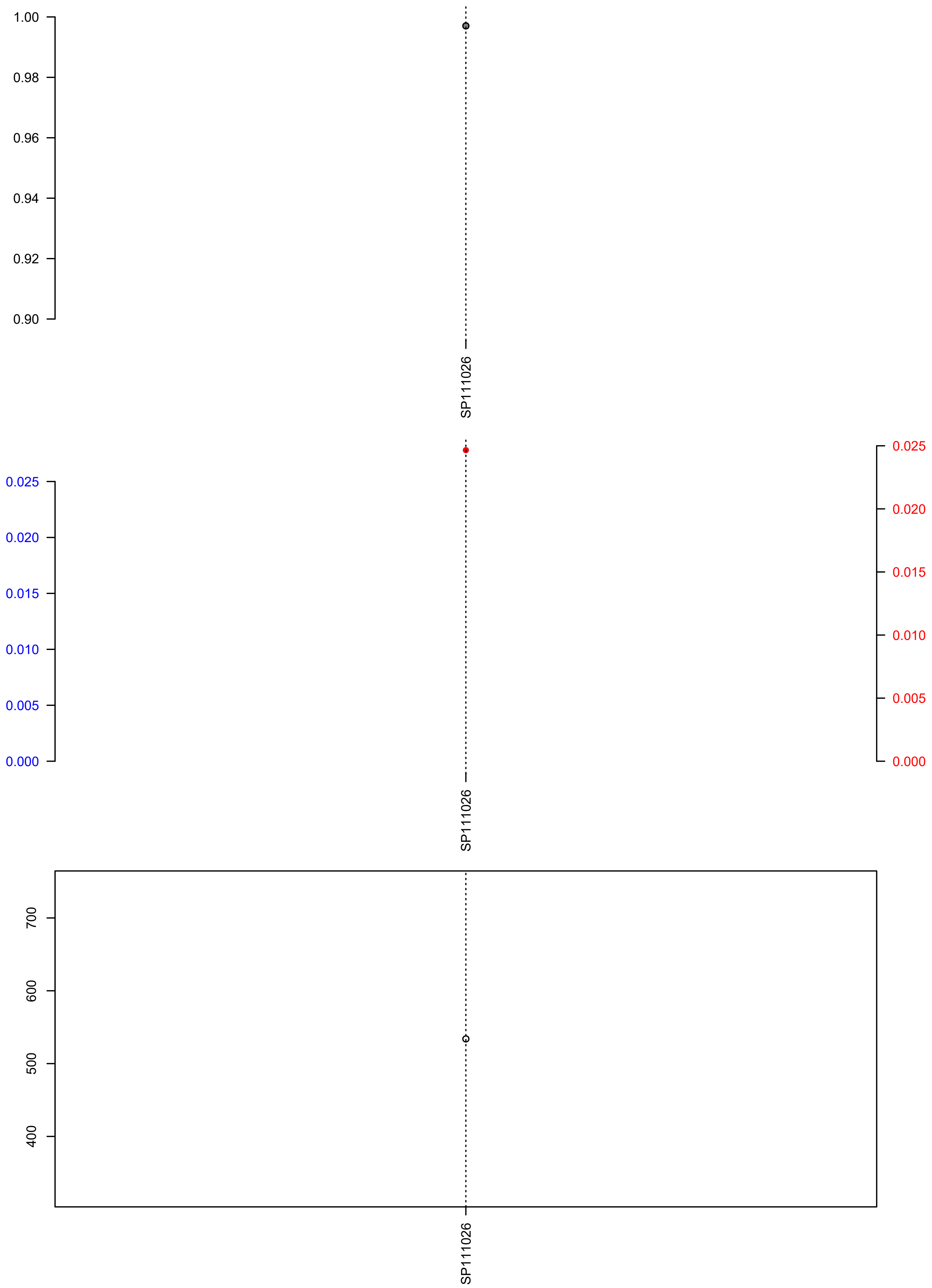

### SP111101.check.with.sig.pdf

# SP111101

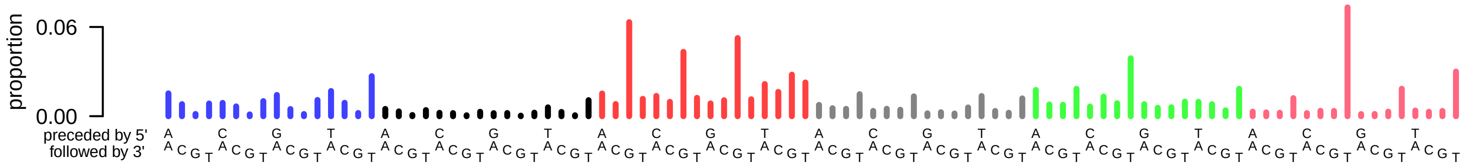

### SP111101.exposures.pdf

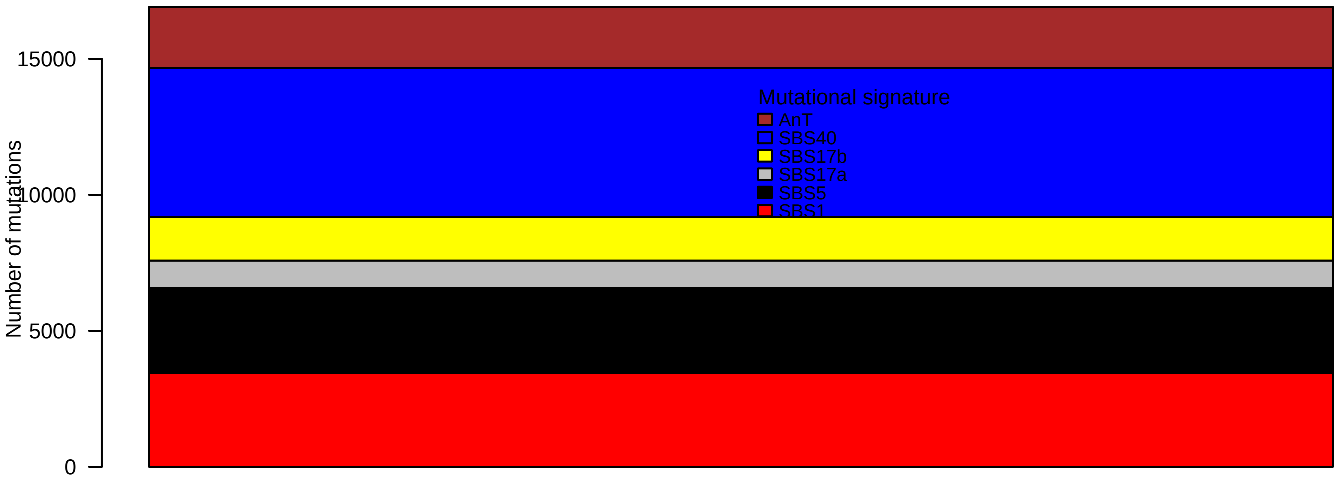

### SP111101.pval.histogram.pdf

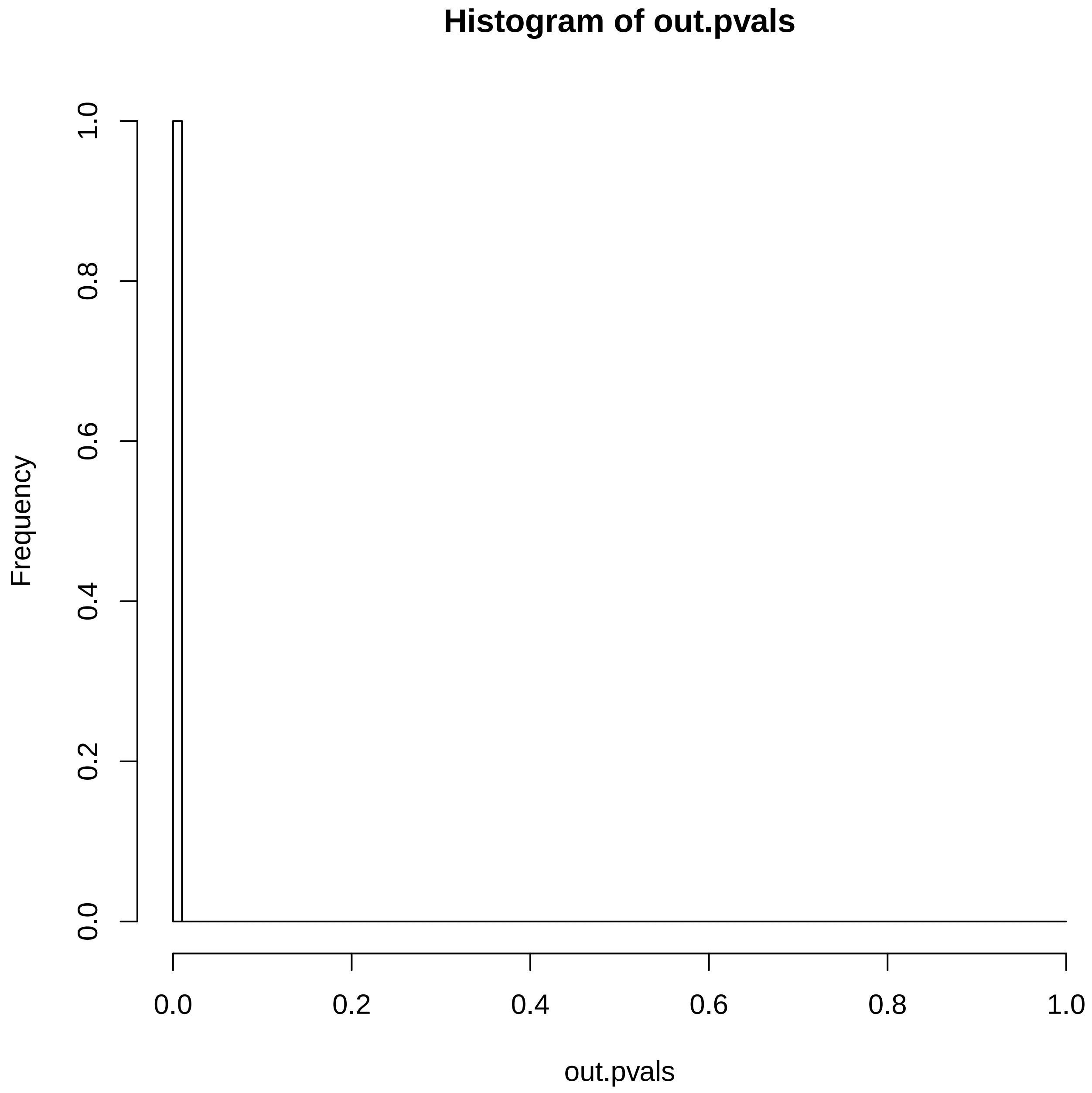

### SP111101.reconstruction.err.pdf

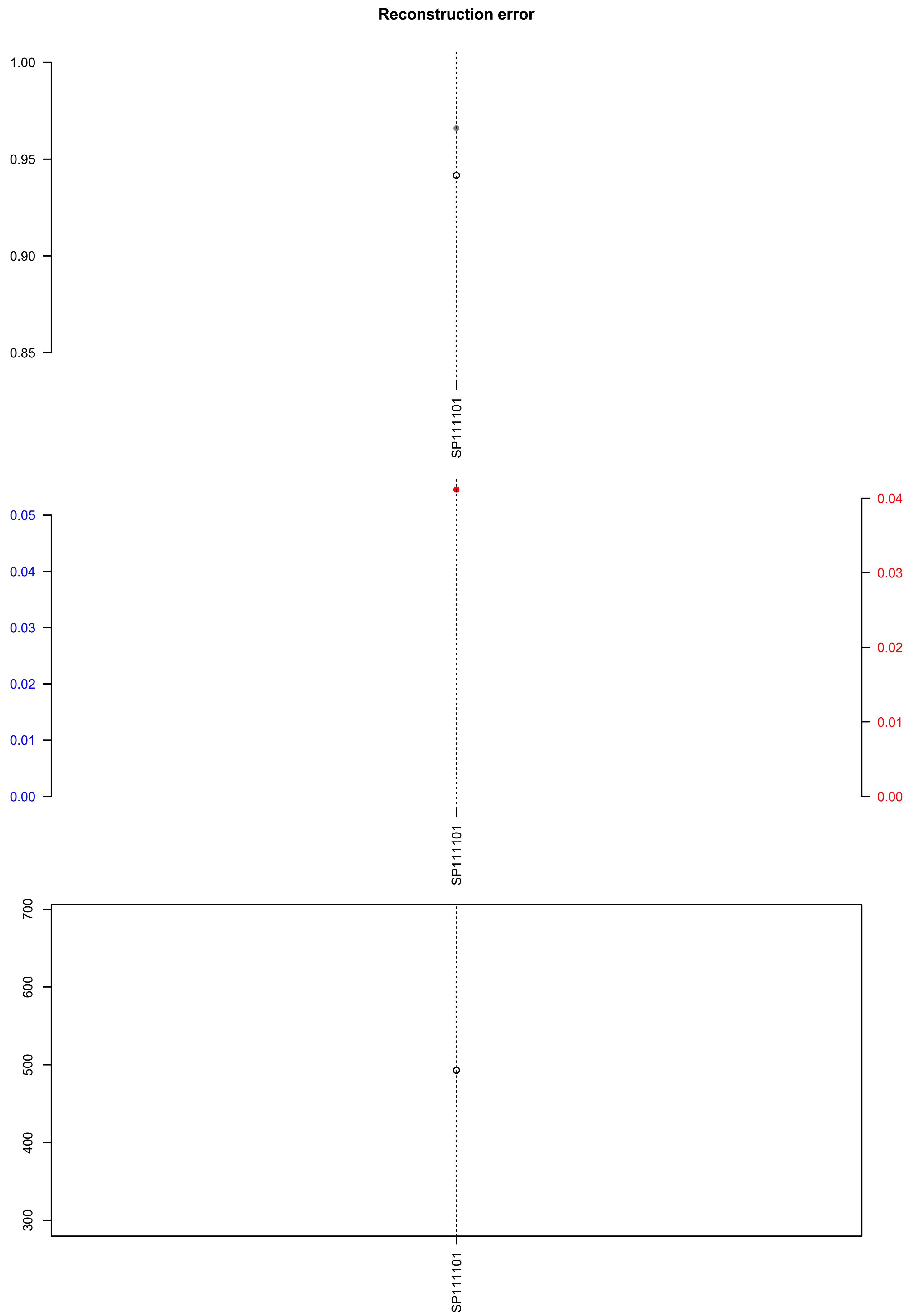

### sysucc-311T.check.with.sig.pdf

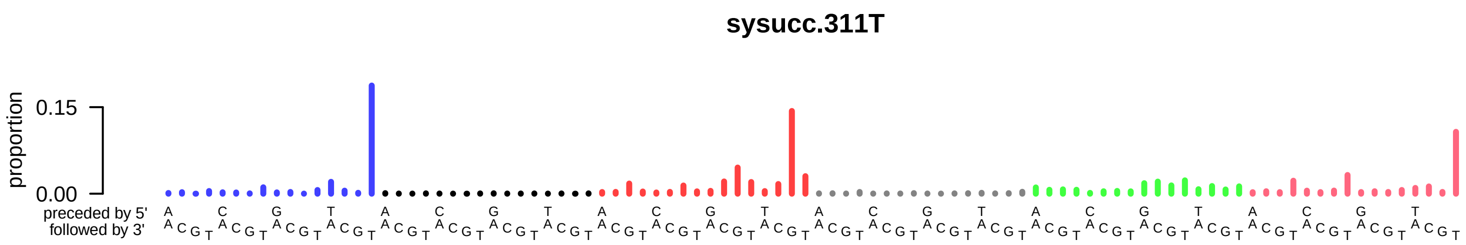

### sysucc-311T.exposures.pdf

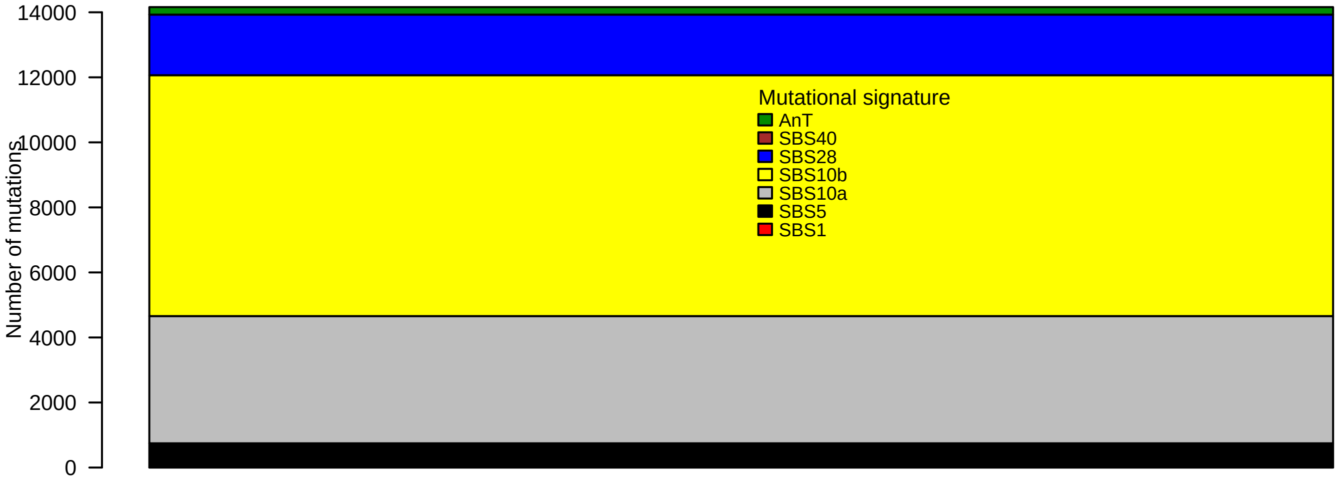

### sysucc-311T.pval.histogram.pdf

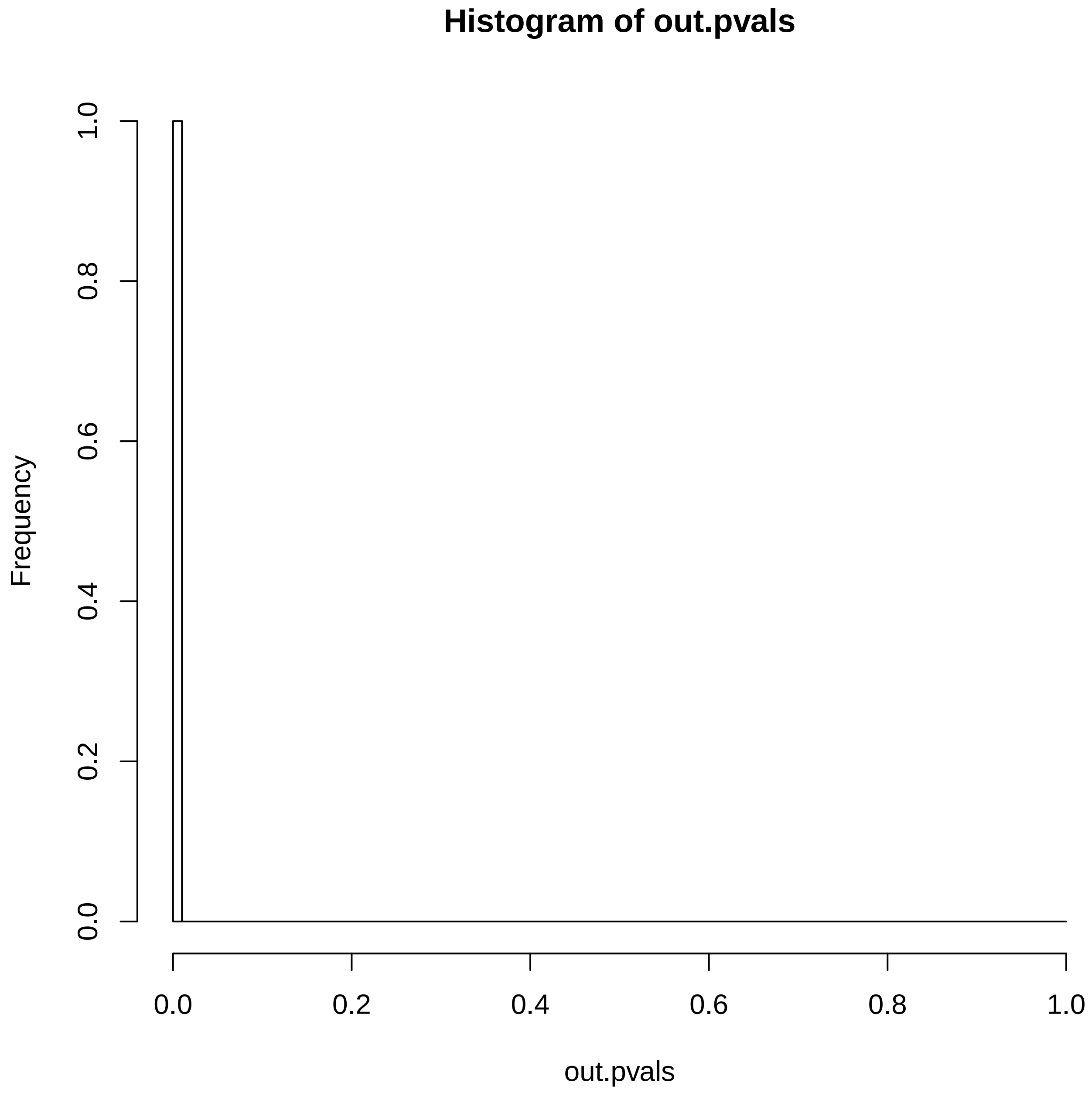

### sysucc-311T.reconstruction.err.pdf

#### Reconstruction error

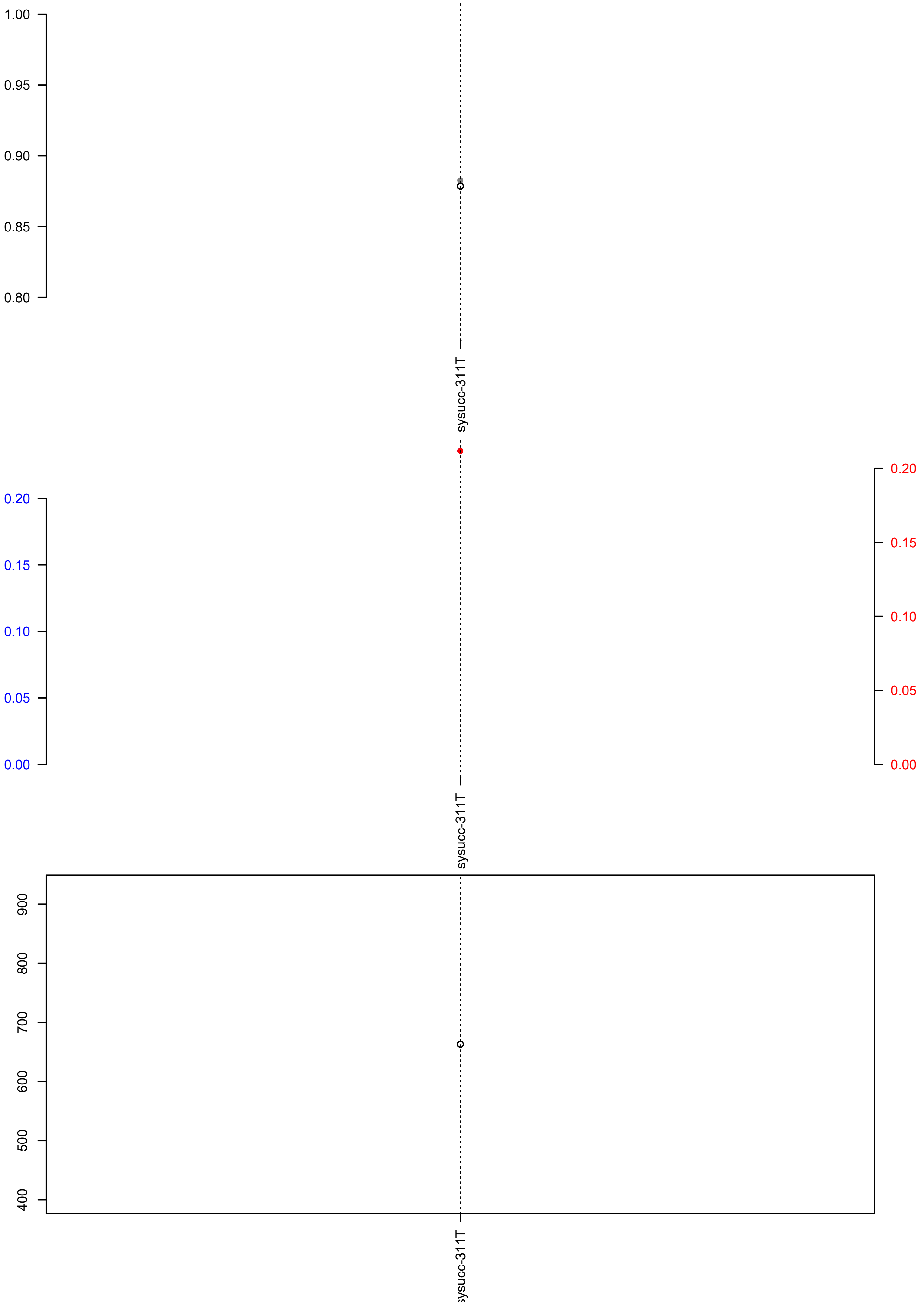

### T155.exposures.pdf

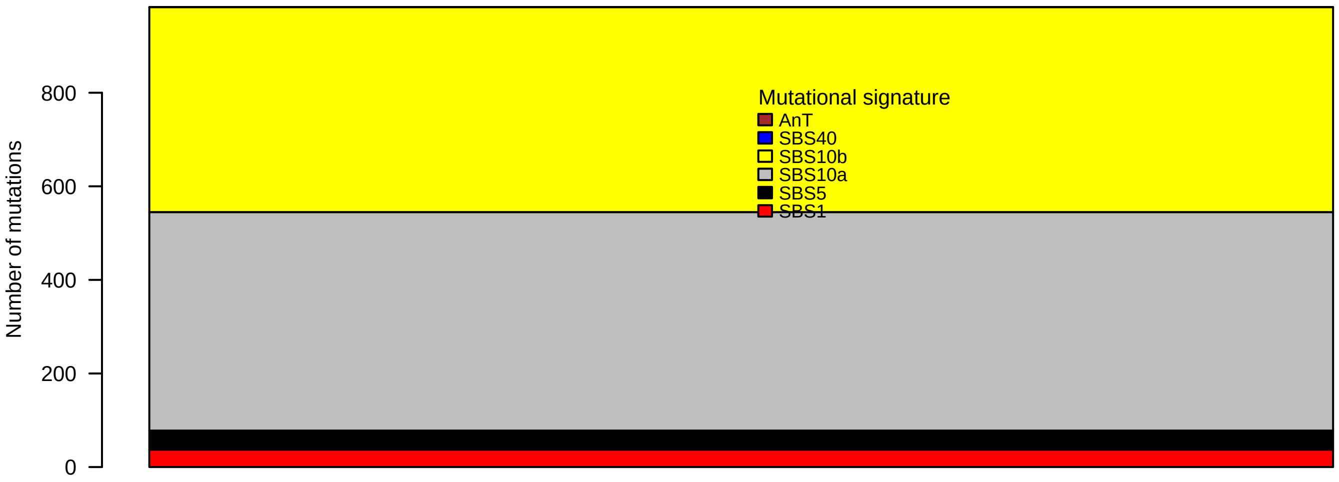

### T155.pval.histogram.pdf

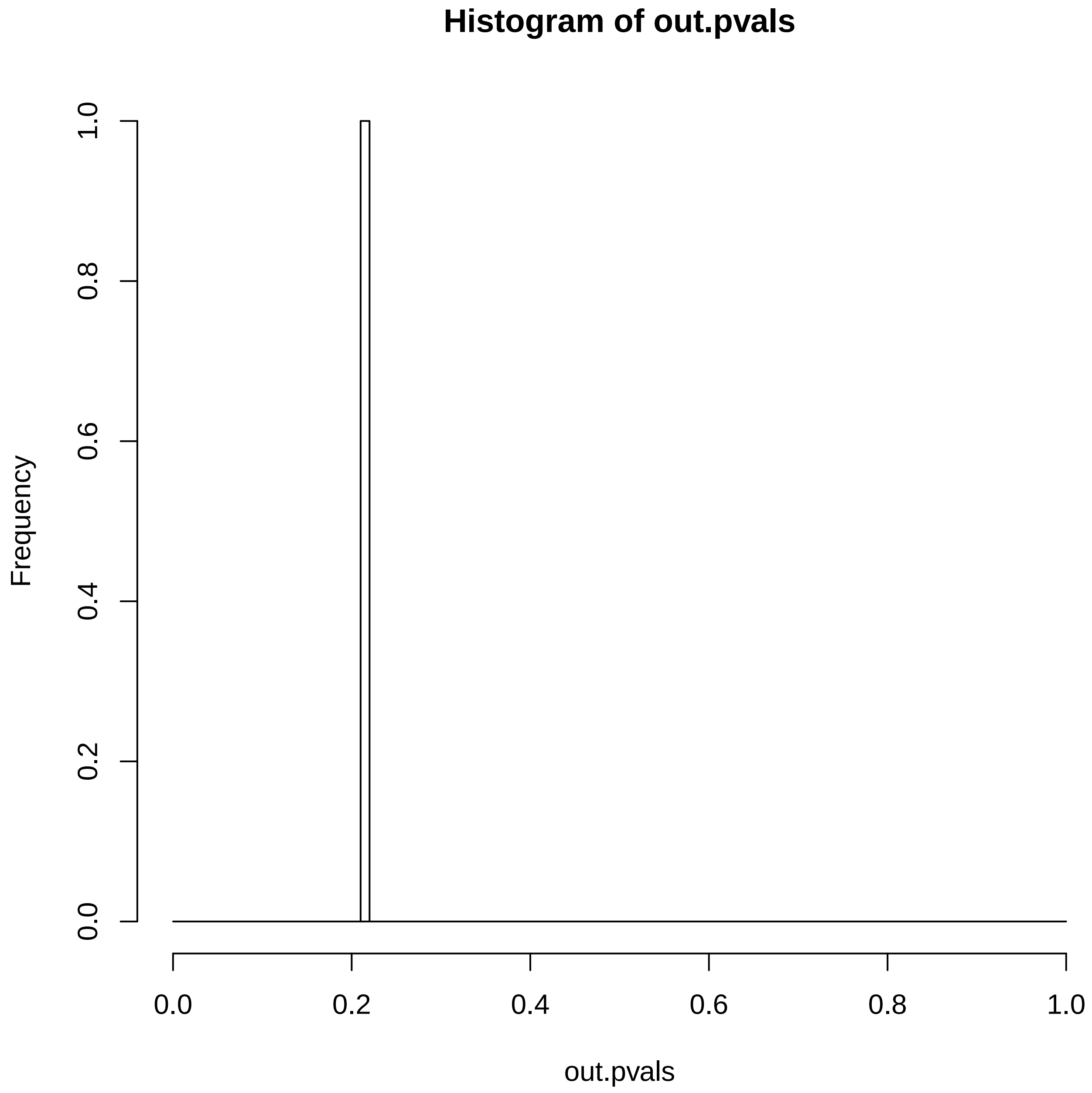

### T155.reconstruction.err.pdf

##### Reconstruction error

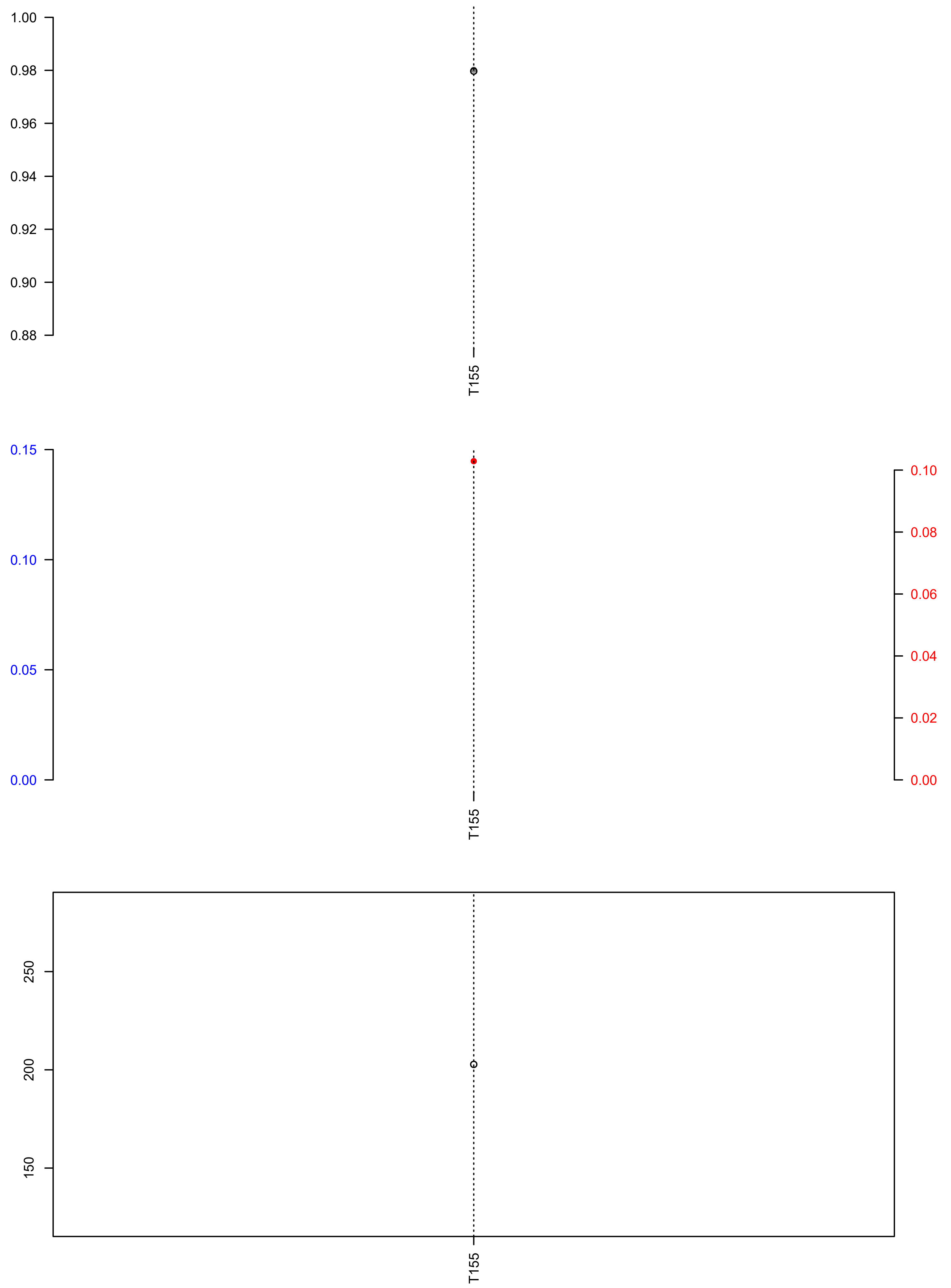

### TCGA-2H-A9GM-01A-11D-A37C-09.exposures.pdf

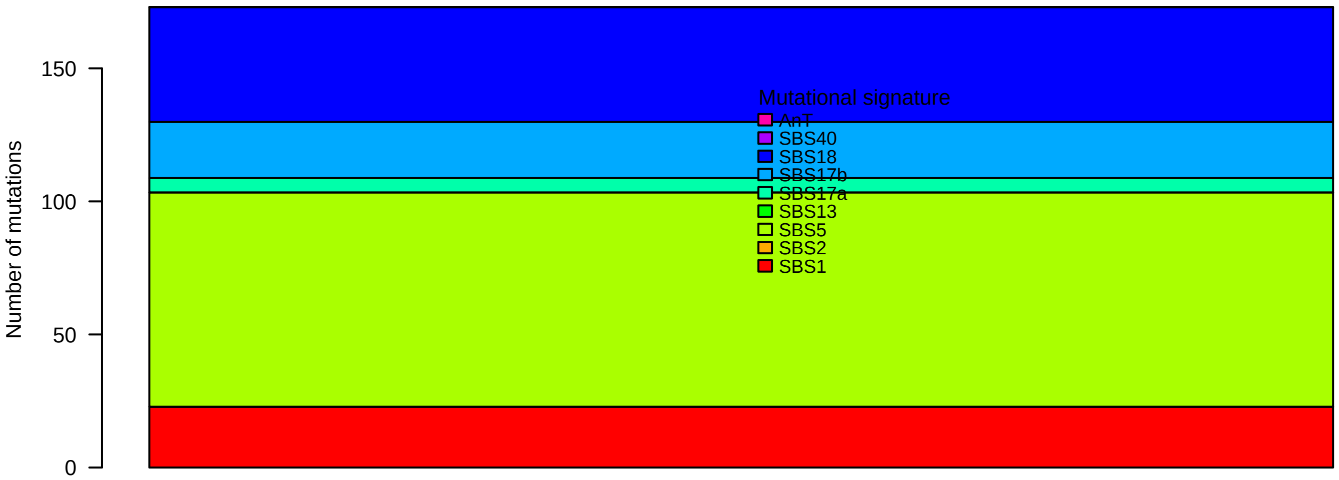

### TCGA-2H-A9GM-01A-11D-A37C-09.pval.histogram.pdf

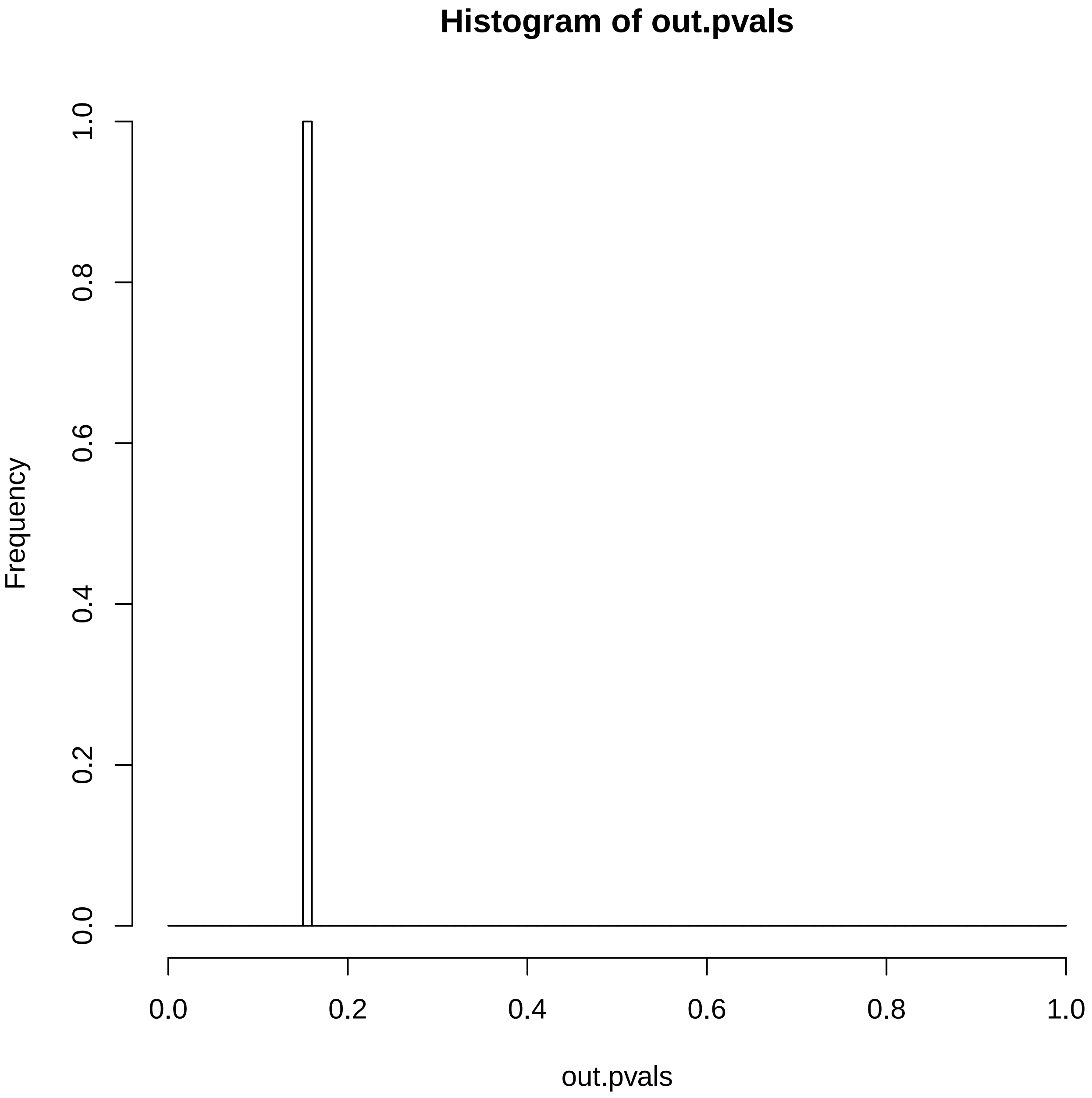

### TCGA-2H-A9GM-01A-11D-A37C-09.reconstruction.err.pdf

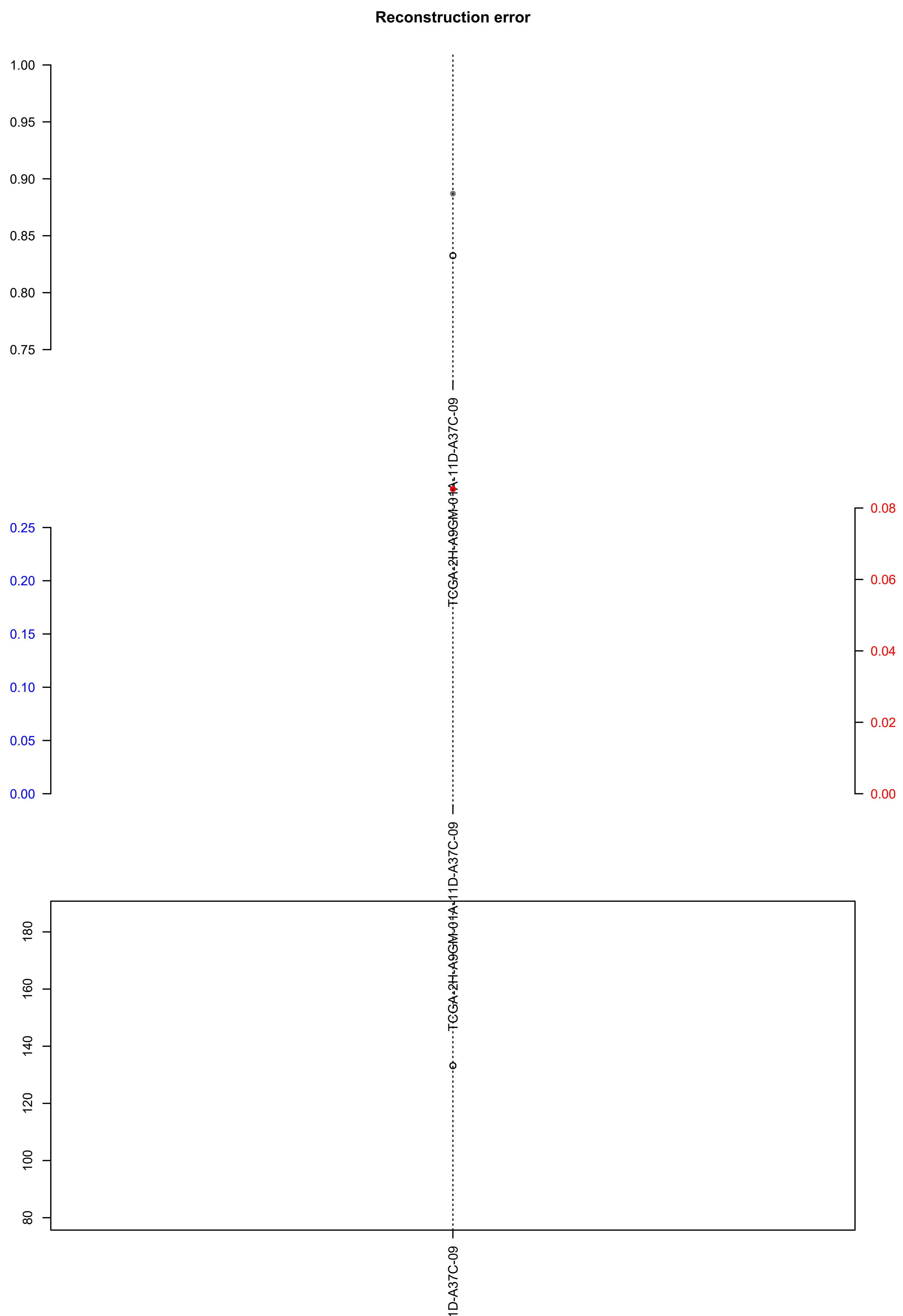

### TCGA-A5-A0GP-01A-11W-A062-09.exposures.pdf

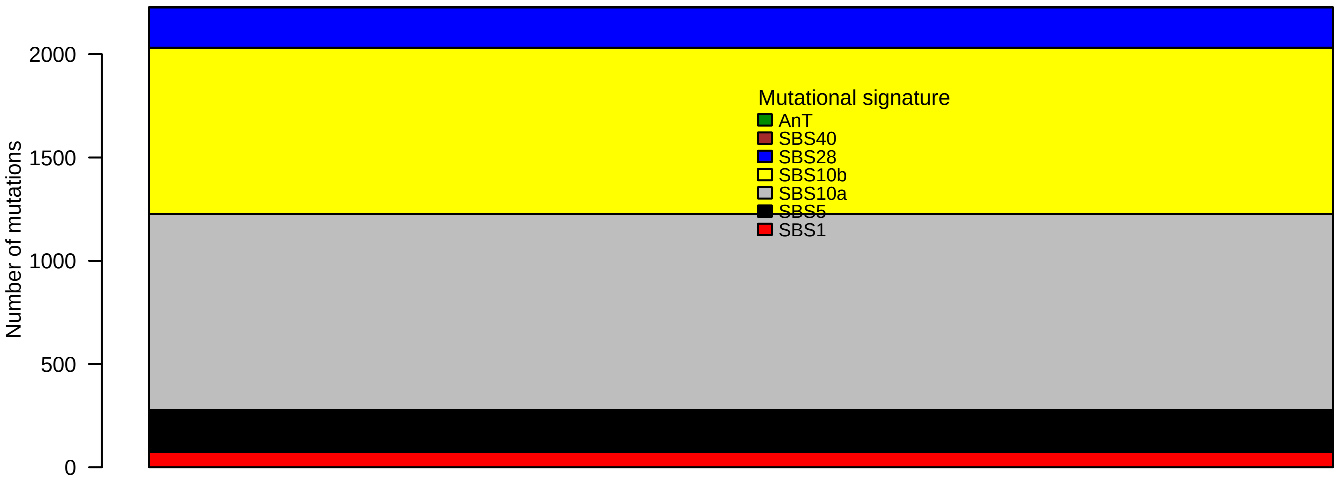

### TCGA-A5-A0GP-01A-11W-A062-09.pval.histogram.pdf

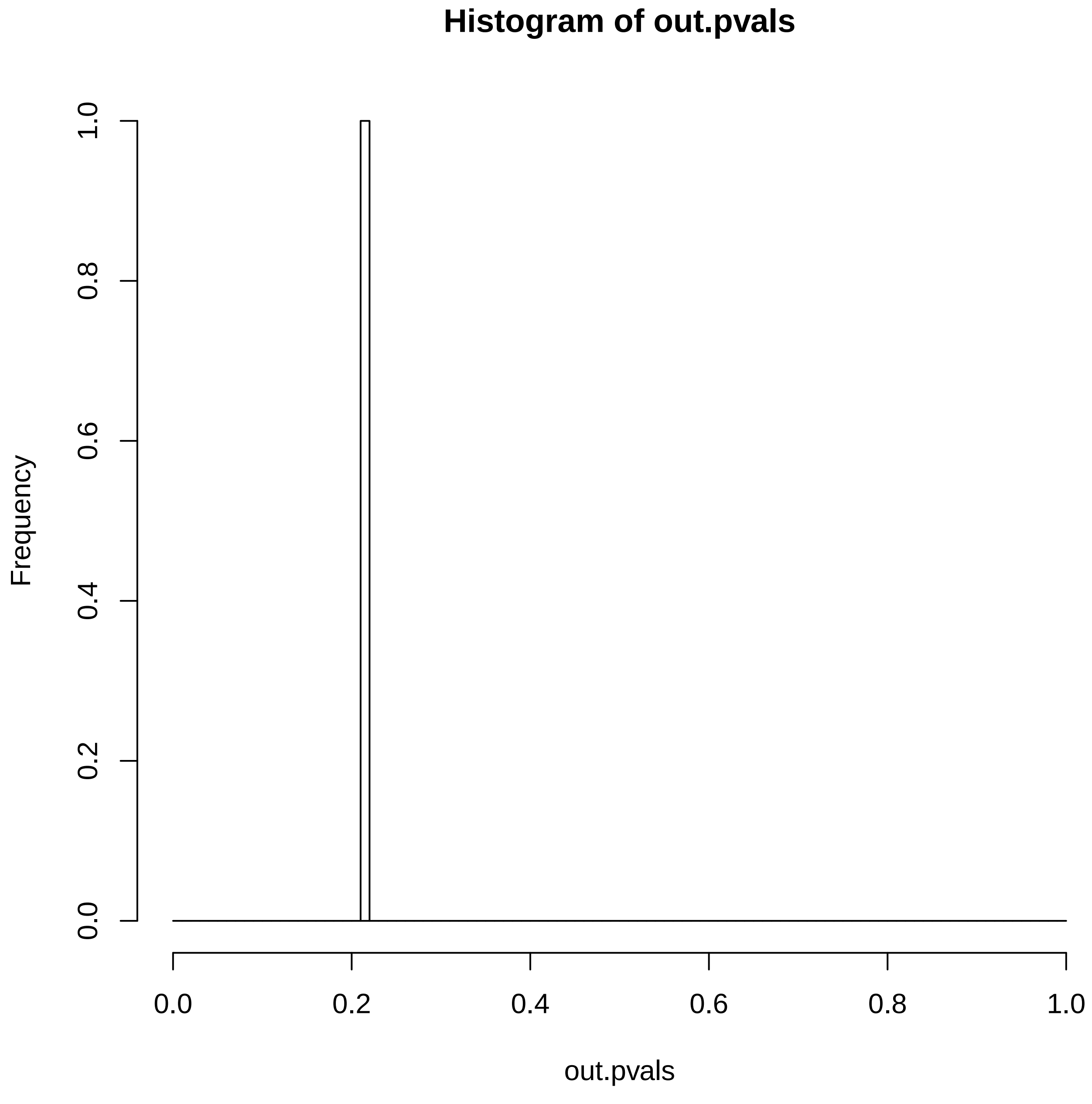

### TCGA-A5-A0GP-01A-11W-A062-09.reconstruction.err.pdf

##### Reconstruction error

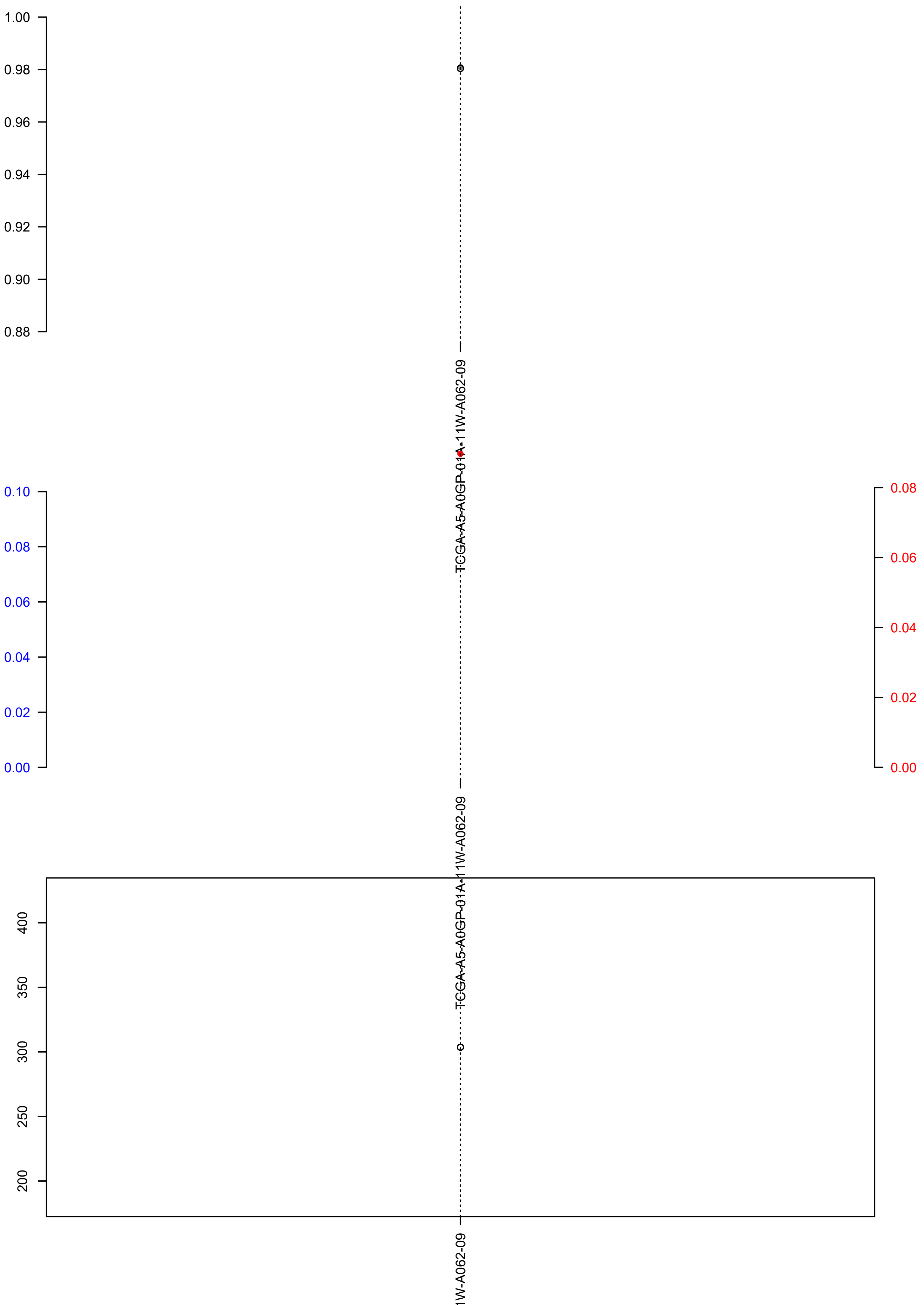
